# Critical contribution of 3’ non-seed base pairing to the *in vivo* function of the evolutionarily conserved *let-7a* microRNA

**DOI:** 10.1101/2021.03.29.437276

**Authors:** Ye Duan, Isana Veksler-Lublinsky, Victor Ambros

## Abstract

Base-pairing of the seed region (g2-g8) is essential for microRNA targeting, however, the *in vivo* function of the 3’ non-seed region (g9-g22) are less well understood. Here we report the first systematic investigation of the *in vivo* roles of 3’ non-seed nucleotides in microRNA *let-7a,* whose entire g9-g22 region is conserved among bilaterians. We found that the 3’ non-seed sequence functionally distinguishes *let-7a* from its family paralogs. The complete pairing of g11-g16 is essential for *let-7a* to fully repress multiple key targets, including evolutionarily conserved *lin-41*, *daf-12* and *hbl-1*. Nucleotides at g17-g22 are less critical but may compensate for mismatches in the g11-g16 region. Interestingly, the 3’ non-seed pairing of *let-7a* can be critically required with certain minimal complementarity for sites with perfect seed pairing. These results provide evidence that the specific configurations of both seed and 3’ non-seed base-pairing can critically influence microRNA-mediated gene regulation *in vivo*.

## Introduction

MicroRNAs (miRNA) are short non-coding RNAs that exist in all metazoans (Friedman et al., 2009; Lee et al., 1993; Nelson and Ambros, 2021). miRNAs were primarily transcribed from their specific genomic loci and processed into the precursor (pre-miRNA) with a hairpin structure consisting of the miRNA and the passenger strand (Bartel, 2018). The pre-miRNAs were further processed, and the mature miRNAs are bound by Argonaute proteins (AGO) to form the miRNA-induced silencing complex (miRISC) and base pair with complementary sites in the 3’ untranslated region (UTR) of target RNAs. miRISC binding leads to post-transcriptional repression of target gene expression through translational inhibition and/or target RNA destabilization (Bartel, 2018). miRNAs are critical for the regulation of diverse physiological processes across species (Ambros, 2004; Bartel, 2018). Up to 60% of human genes are estimated to be regulated by miRNAs, and disfunction of miRNAs is implicated in multiple human diseases (Friedman et al., 2009; Paul et al., 2018).

The miRNA seed corresponds to the 6-7 contiguous nucleotides beginning at the second nucleotide (g2) from 5’ end and is understood to be the dominant determinant of miRNA targeting efficacy and specificity. Structural and biochemical studies indicate that the miRNA seed sequence is critical for target recognition and binding (Salomon et al., 2015; Schirle et al., 2014). Complementarity to the miRNA seed is the most evolutionarily conserved feature of miRNA target sites, and genetically disrupting seed complementarity can result in de-repression of miRNA targets (Lai, 2002; Lewis et al., 2005; Lim et al., 2005).

Organisms commonly contain multiple genes encoding miRNAs with identical seed sequences, which are grouped into seed families. miRNAs of the same family can in principle recognize shared targets through seed pairing and thereby function redundantly (Abbott et al., 2005; Alvarez-Saavedra and Horvitz, 2010; Brenner et al., 2012). Meanwhile, miRNAs with identical seed but divergent 3’ non-seed nucleotides (g9-g22) can exhibit target site selectivity driven by the extent of non-seed base pairing (Broughton et al., 2016; Wahlquist et al., 2014).

*let-7 (lethal-7) family* miRNAs are distributed widely across the bilaterians, consistent with evolutionary origins in bilaterian ancestors (Hertel et al., 2012; Wolter et al., 2017). Similarities in the developmental profile of *let-7 family* miRNAs across diverse species and genetic analyses in model organisms suggest evolutionary conservation of *let-7a family* functionality, which highlights the *let-7 family* as a general model for miRNA studies (Roush and Slack, 2008; Tennessen and Thummel, 2008). In nematodes, *let-7 family* miRNAs function in the heterochronic pathway to promote the stage-specific cell fate transitions during larval development (Abbott et al., 2005; Reinhart et al., 2000). Similarly, *let-7 family* miRNAs control cellular transitions from pluripotency to differentiation and function in tumor suppression in mammals, and the timing of adult fates during metamorphosis in insects (Balzeau et al., 2017; Lee et al., 2016; Sokol et al., 2008).

Notably, almost all bilaterian genomes encode at least one *let-7 family* isoform (*let-7a*), in which all nucleotides in the 3’ non-seed region (g9-g22) are highly conserved (Fig. S1A) (Hertel et al., 2012; Pasquinelli et al., 2003). This deep conservation suggests that the 3’ non-seed region is associated with essential functions which evolutionarily constrained the sequence. Indeed, high-throughput analyses suggested that miRNA 3’ non-seed regions can contribute substantially to *in vivo* miRNA-target interactions (Grosswendt et al., 2014; Helwak et al., 2013). Pairing to the 3’ non-seed region, especially g13-g16, is thought to enhance target repression in certain contexts (Brennecke et al., 2005; Grimson et al., 2007). Recent structural studies of human AGO complexed with miRNA and targets revealed that g13-g16 can form an A-form helix with the target within miRISC, and can thereby increase miRISC-target affinity and specificity (Sheu-Gruttadauria et al., 2019b; Xiao and MacRae, 2020). Notably, some miRNA target sites have imperfect seed complementarity accompanied by extensive non-seed pairing (Chi et al., 2012; Grimson et al., 2007; Vella et al., 2004). In such cases, the 3’ pairing is thought to compensate for the imperfect seed pairing, and this compensatory configuration can determine the target specificity among family isoforms (Brancati and Grosshans, 2018; Brennecke et al., 2005; Doench and Sharp, 2004). However, the relative contributions of specific 3’ non-seed nucleotides to the *in vivo* function of miRNAs remain unclear.

Here, we report a systematic investigation of how the 3’ non-seed nucleotides contribute to the *in vivo* function of the evolutionarily conserved *let-7a*. We employ CRISPR/Cas9 genome editing of *C. elegans* to systematically introduce defined mutations into the *let-7a* and its conserved target *lin-41*. We show that the sequence of g11-g16 is essential for *let-7a in vivo* function, while g17-g22 pairing is less critical than g11-g16 pairing but can partially compensate for mismatches in g11-g16. We confirm that the 3’ non-seed sequence of *let-7a* confers specificity relative to its family paralogs. Furthermore, we show that *lin-41*, as well as heterochronic genes *daf-12* and *hbl-1,* are *let-7a* targets whose proper developmental *in vivo* repression requires 3’ non-seed pairing. Interestingly, we find the 3’ non-seed pairing to *let-7a* is functionally required in certain cases of perfect seed pairing, including the natural *daf-12* 3’ UTR and the reconfigured endogenous *lin-41* 3’ UTR, suggesting that the g11-g16 nucleotides of *let-7a* engage in essential interactions that, in parallel with seed pairing, critically influence miRISC repressing activity.

## Results

### The 3’ non-seed sequence is essential for the *in vivo* functional specificity of *let-7a* compared to family paralogs

An examination of the phylogenetic distribution of *let-7 family* miRNA genes (Fig. 1A, S1B) emphasizes the presence of *let-7 family* in most bilaterians with available miRNA annotations in miRbase 22.1 (Kozomara et al., 2019). 91.5% (n=117) of bilaterian species have at least one *let- 7 family* isoform with no more than 3 nucleotide differences from *let-7a* (Fig. S1C-E). Alignment of the isoforms closest to *let-7a* from each of the 117 bilaterian species indicates that the g9-g22 nucleotides exhibit conservation ranging from 86% to 99%, and 75.2% of the 117 species contain a *let-7* miRNA completely identical to *let-7a* (Fig. 1B, S1F). This degree of conservation is not observed for other conserved miRNAs (Fig. S1D-H). The deep conservation of *let-7a* suggests that *let-7a* is associated with essential functions that depend on the identity of the 3’ non-seed nucleotides.

**Figure 1.**
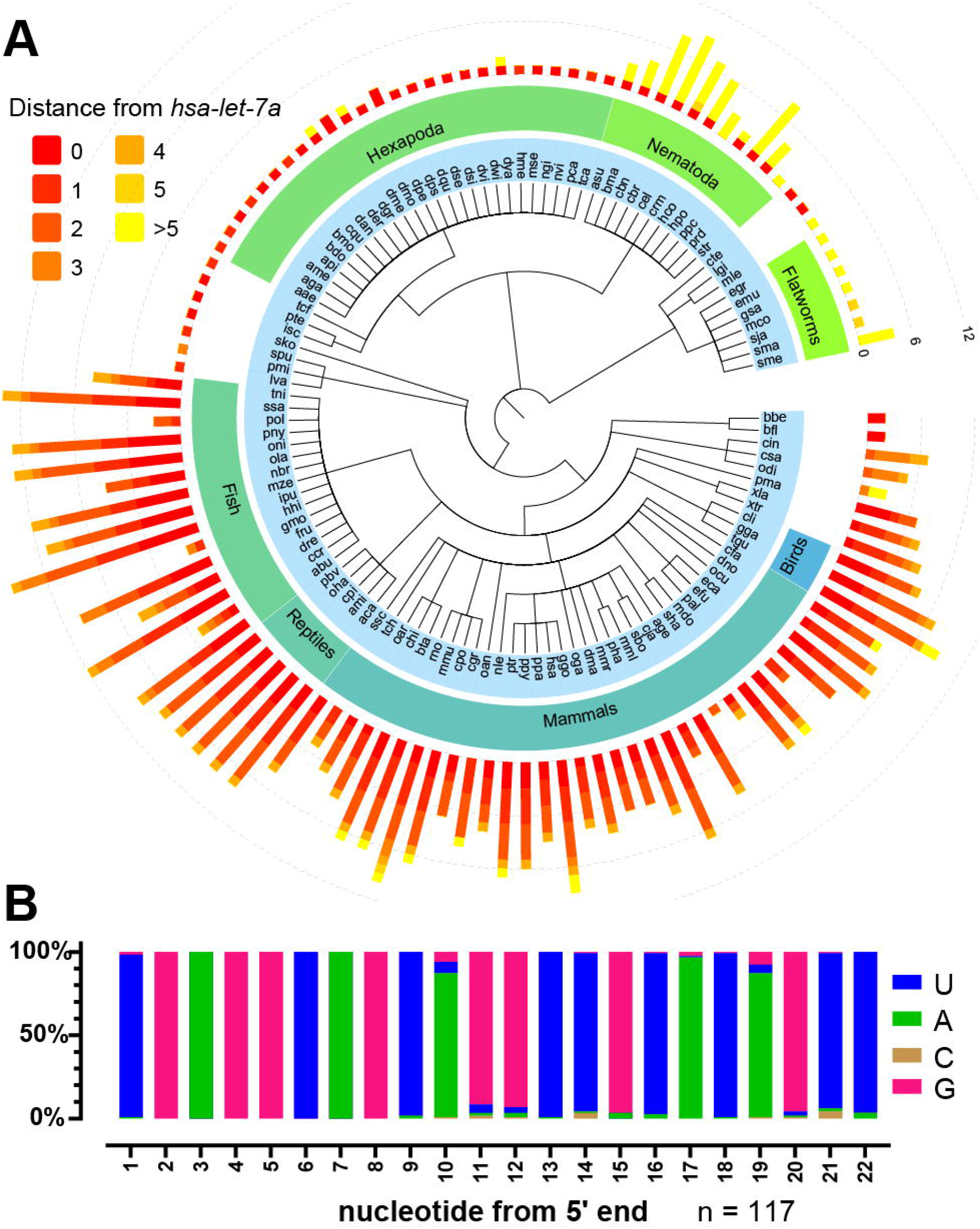
*let-7a* is deeply conserved at all nucleotides across bilaterian species. **A.** Summary of *let-7 family* miRNAs across bilaterian phylogeny. Bar length, numbers of *let-7 family* isoforms; bar color, sequence distances relative to *hsa-let-7a-5p*. **B.** Nucleotide frequency of the *let-7 family* isoforms most similar to *hsa-let-7a-5p* across bilaterians. See also Figure S1.

To test the hypothesis that the 3’ non-seed nucleotides are critical for *let-7a in vivo* function, we used CRISPR/Cas9 to mutate the endogenous *let-7a* in *C. elegans* by swapping its sequence with that of its closest paralog *miR-84* (Fig. 2A, S1A). In the resulting *let-7(ma341*) mutant, both the miRNA and passenger strands were mutated to preserve the pre-miRNA structure. The temporal expression profile of *miR-84* miRNA in *let-7(ma341)* is consistent with expression from the endogenous *mir-84* locus (starting in L2) combined with expression from the edited *let-7* locus (peaking in L4) (Fig. 2B-C). Moreover, in the *mir-84(n4037, null)* background, *miR-84* expression from the *let-7(ma341)* locus alone exhibits a temporal profile like that of normal *let-7a* (Fig. 2D-E). Note that the detection method stringently distinguishes *let-7a* from *miR-84*, as *let-7a* was not detected in *let-7(ma341)* and *miR-84* was not detected in *mir-84(n4037)* (Fig. 2C-E). These results confirm that *let-7(ma341)* expresses *miR-84* miRNA instead of *let-7a*.

**Figure 2.**
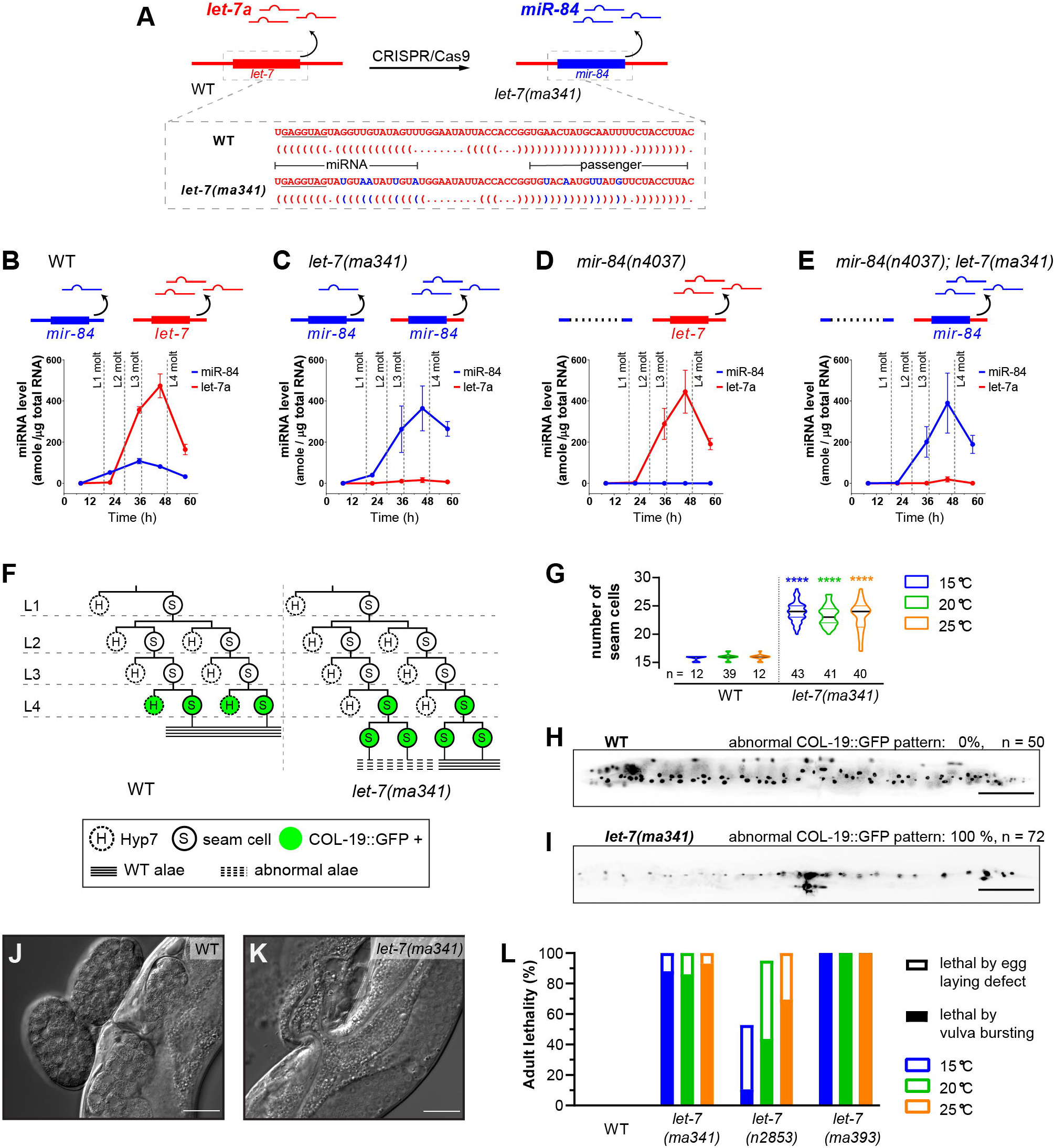
The 3’ non-seed sequence determines the functional specificity of *let-7a* among its paralogs. **A.** Strategy for generation of *let-7*(*ma341)* by CRISPR/Cas9 mutagenesis to swap the *pre-let-7* sequence for the *pre-mir-84* sequence. Dot-bracket notations show the pre-miRNA structures predicted by RNAfold (Denman, 1993). **B-E.** Developmental profiles of *miR-84* and *let- 7a* miRNAs determined by Fireplex assays. Expression is calibrated with synthetic *miR-84* and *let-7a* oligos. Time is hours after feeding starvation-arrested L1 larvae. **F.** Representative lineages and COL-19::GFP expression patterns of V1-V4 seam cells. **G.** Seam cell numbers of young adults. n, numbers of animals tested. **H-I.** Representative expression patterns of COL-19::GFP in adults. Scale bars, 100 µm. **J-K.** DIC images of the vulva region of adults. Scale bars, 25 µm. **L.** Adult lethality for WT, *let-7(ma341)*, *let-7(n2853,* G5C), and *let-7(ma393, null)*. Lethal phenotypes are categorized as severe (via bursting of young adults through the vulva) or mild (via matricide of egg-laying defective gravid adults). Data in **B-E** represents 3 biological replicas. See also Figure S5. Details of the phenotypes are available in Table S1.

In wild-type (WT) animals, the seam cells (hypodermal stem cells) go through asymmetric divisions at each larval stage as one daughter cell differentiates to Hyp7 cells (hypodermis) while the other remains a stem cell (Sulston et al., 1983). At the L4 molt, the seam cells exit the cell cycle, fuse, and produce the adult-specific cuticle structure referred to as adult lateral alae. In *let- 7(ma341)*, we found that seam cells underwent an extra round of cell division after the L4 molt, resulting in extra seam cells in young adults, accompanied by incomplete lateral alae (Fig. 2F-G, Table S1). *let-7(ma341)* adults also exhibit impaired expression of the adult-specific hypodermal reporter COL-19::GFP (Fig. 2H-I). These results indicate that *let-7(ma341)* causes retarded heterochronic phenotypes. *let-7(ma341)* also exhibits adult lethality caused by bursting at the vulva (Fig. 2J-L). The adult lethality and retarded phenotypes are consistent with the *let-7a loss-of-function (lf)* phenotypes observed in the *null (ma393)* and seed (*n2853)* mutants (Liu et al., 1995; Nelson and Ambros, 2019; Reinhart et al., 2000), indicating that the *in vivo* function of *let- 7a* cannot be substituted by its closest paralog *miR-84,* even when *miR-84* is expressed in a developmental profile essentially identical to *let-7a*. Since *let-7a* and *miR-84* share the same seed sequence, these findings indicate that the 3’ non-seed sequence is a critical determinant of the *in vivo* function and specificity of *let-7a*.

### Nucleotides 11-16 are each functionally essential for *let-7a in vivo*

We next sought to characterize how each 3’ non-seed nucleotide contributes to *in vivo* function of *let-7a*. We performed a single nucleotide mutational screen of the *let-7a* non-seed region using the “jump-board” strategy (Duan et al., 2020a). For each g9-g22 nucleotide, we mutated both the miRNA and passenger strands to preserve the pre-miRNA structure (Fig. 3A, S2A). To confirm the expression of the mutant miRNAs, we performed small RNA sequencing of all the mutants and found that the abundance of the mutated *let-7a* miRNA in all cases was greater than 50% of that of the native *let-7a* in WT (Fig. 3B). Since *let-7* is recessive in *C. elegans,* which indicates that 2-fold reduction of *let-7a* dosage is not phenocritical, we reason that any phenotypes exhibited by these mutants can be attributed to sequence but not to reduced expression of miRNA. Note that the strains with mutations at g11-g13 also contained a WT *let-7* allele on a genetic balancer *umnIs25(mnDp1)*, hence both WT and mutant *let-7a* were expressed.

**Figure 3.**
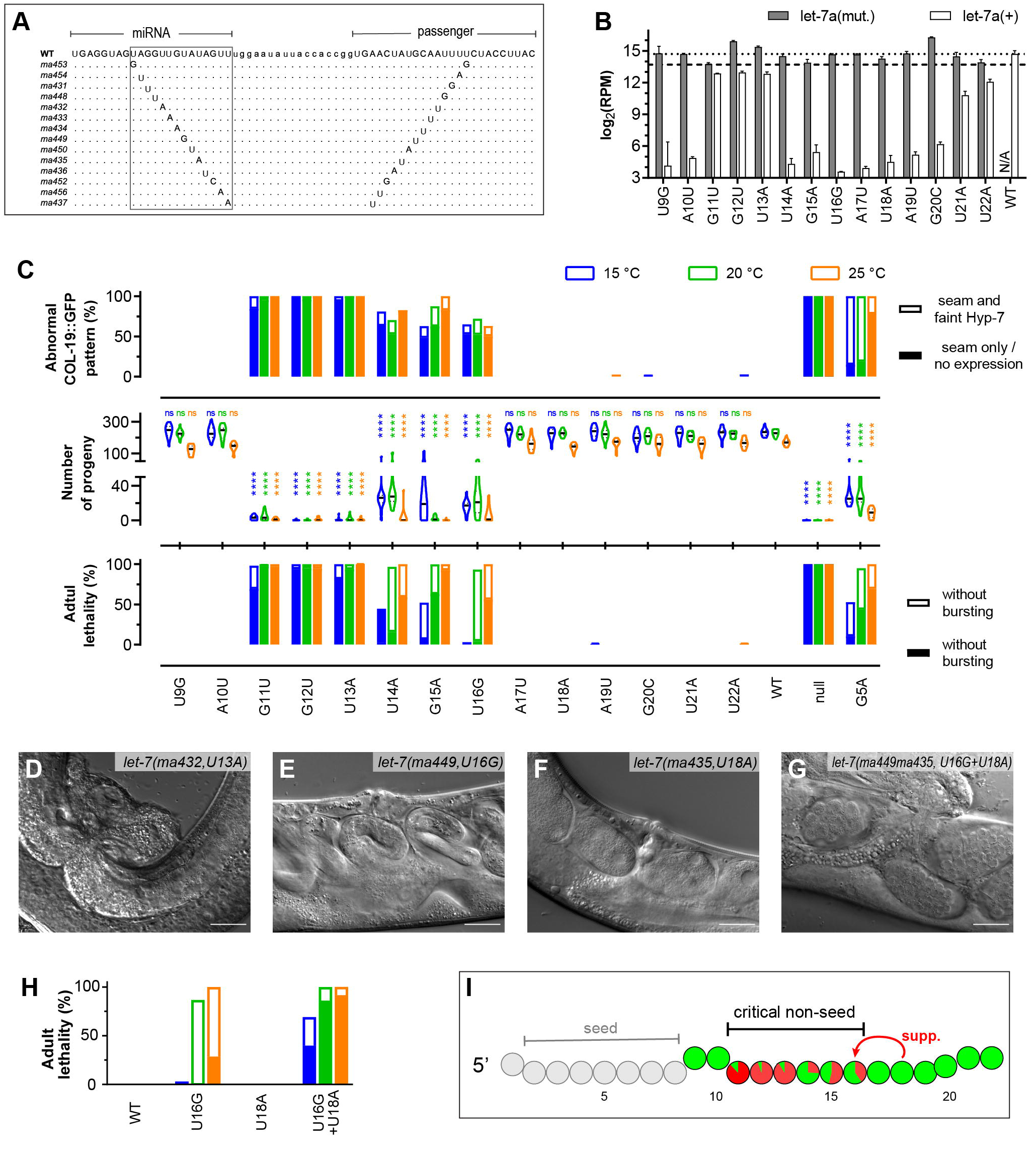
Contribution of single 3’ non-seed nucleotides to the *in vivo* function of *let-7a.* **A.** Alignment of pre-miRNA sequences. Boxed, 3’ non-seed regions. **B.** Small RNA sequencing reads from L4 larvae that mapped to WT or mutant *let-7a* sequences for each single mutant. The reads mapping to WT *let-7a* for strains carrying mutations at g11-g13 include WT *let-7a* miRNA from balancer *umnIs25(mnDp1).* The WT reads from U21A/U22A mutants likely reflect *let-7a(mut.)* miRNAs whose 3’ ends had been trimmed and subsequently uridylated *in vivo*, resulting in artificial *let-7a(+)* reads. Dashed lines, 100% (top) and 50% (bottom) of RPM of *let-7a(+)* in WT. **C.** Quantitation of *let-7a lf* phenotypes: percent animals with abnormal COL-19::GFP expression (top), numbers of progeny per animal (middle), and percent adult lethality (bottom). Lethal phenotypes are categorized as in Fig. 2L. Abnormal COL-19::GFP patterns are categorized as no Hyp7 expression (severe) or faint Hyp7 expression (mild). **D-G.** DIC images of the vulva region in adults at 25 °C. Scale bars, 25 µm. **H.** Functional synergy between g18 and the critical non-seed region based on vulva integrity defect. Labels are identical to **C** (bottom). **I.** Summary of the functional merits of *let-7a* 3’ non-seed nucleotides. Colored circle, 3’ non-seed nucleotide; red proportion, average adult lethality caused by vulva bursting at all temperatures. See also Figures S2, S3. Details of the phenotypes are available in Table S1.

Single nucleotide mutations at g11, g12, or g13 resulted in severe vulva integrity defects characteristic of *let-7 lf* phenotypes, leading to lethality and dramatically reduced numbers of progeny quantitatively similar to *let-7(null*) (Fig. 3C-D). Single nucleotide mutations at g11-g13 also resulted in impaired expression of COL-19::GFP in Hyp7 and adult alae morphogenesis defects, indicative of retarded heterochronic phenotypes (Fig. 3C, S2B, Table S1). Single nucleotide mutations at g14-g16 also resulted in distinct *let-7a lf* phenotypes, although milder than g11-g13 mutations (Fig. 3C-E). By contrast, single nucleotide mutations at g9-g10 or g17-g22 did not cause visible phenotypes (Fig. 3C, 3F, Table S1). These results reveal g11-g16 to be a critical non-seed region of *let-7a*, with g11-g13 are relatively more essential than g14-g16 (Fig. 3G).

### Nucleotides beyond g16 can compensate for mismatches in the critical non-seed region

We found that mutating the g18 of *let-7a* into any of the other 3 nucleotides did not result in *lf* phenotypes and simultaneous mutations of g17-g19, g20-g22, or g17-g22 did not cause apparent *lf* phenotypes, even with sensitized conditions like non-physiological temperatures or pathogen-containing cultures (Fig. S2C and data not shown). These results stand out in contrast to the deep conservation of the *let-7a* sequence, especially with g18 as the second most conserved nucleotide (98.3%). We thus hypothesized that g17-g22 may contribute to target repression in cases where mismatches occur within the g11-g16 region.

To test this possibility, we used the *let-7(ma449,* U16G) mutation as a genetically sensitized background, and generated a compound mutant *let-7(ma449ma435)* with both U16G and U18A mutations. We observed that *let-7(ma449ma435)* exhibited a strong vulva bursting phenotype and retarded adult alae (Fig. 3E-H, S2C-D), which are significantly more penetrant than that of each single mutant. We suggest that g18, and by implication other g17-g22 nucleotides, may contribute to interactions that involve mismatches in the g14-g16 sub-critical region.

It is also possible that g17-g22 pairing could be critical for target repression associated with phenotypes that we have not measured. We accordingly assessed molecular phenotypes of *let- 7(ma435)* by ribosome profiling and RNA-seq (Fig. S3, Table S2). We found that the expression levels of multiple genes are significantly changed by the U18A mutation, among which we identified 3 genes as putative *let-7a* targets associated with g18 pairing, supporting the hypothesis that there are *in vivo* circumstances where g17-g22 nucleotides are critical for proper target regulation.

### De-repression of *lin-41* is a major contributor to *let-7a* critical non-seed mutant phenotypes

The evolutionarily conserved gene *lin-41/Trim71* is a direct target of *let-7a* across divergent animal species, and repression of *lin-41/Trim71* by *let-7a* is essential for normal development in both invertebrates and vertebrates (Ecsedi and Grosshans, 2013; Worringer et al., 2014). In *C. elegans,* robust repression of *lin-41* by *let-7a* is critical for the L4-to-adult cell fate progression, and mutations disrupting the seed pairing between *let-7a* and its complementary sites (LCSs) in the *lin-41* 3’ UTR result in reiteration of L4 cell fates and severe vulva defects (Aeschimann et al., 2019). Notably, both *lin-41* LCSs have imperfect seed complementarity to *let-7a* with a GU pair and a target A-bulge, respectively, while both LCSs contain complementarity to *let-7a* g11-g19, suggesting that 3’ pairing is essential to compensate for weak seed pairing in this context (Fig. 4A).

**Figure 4.**
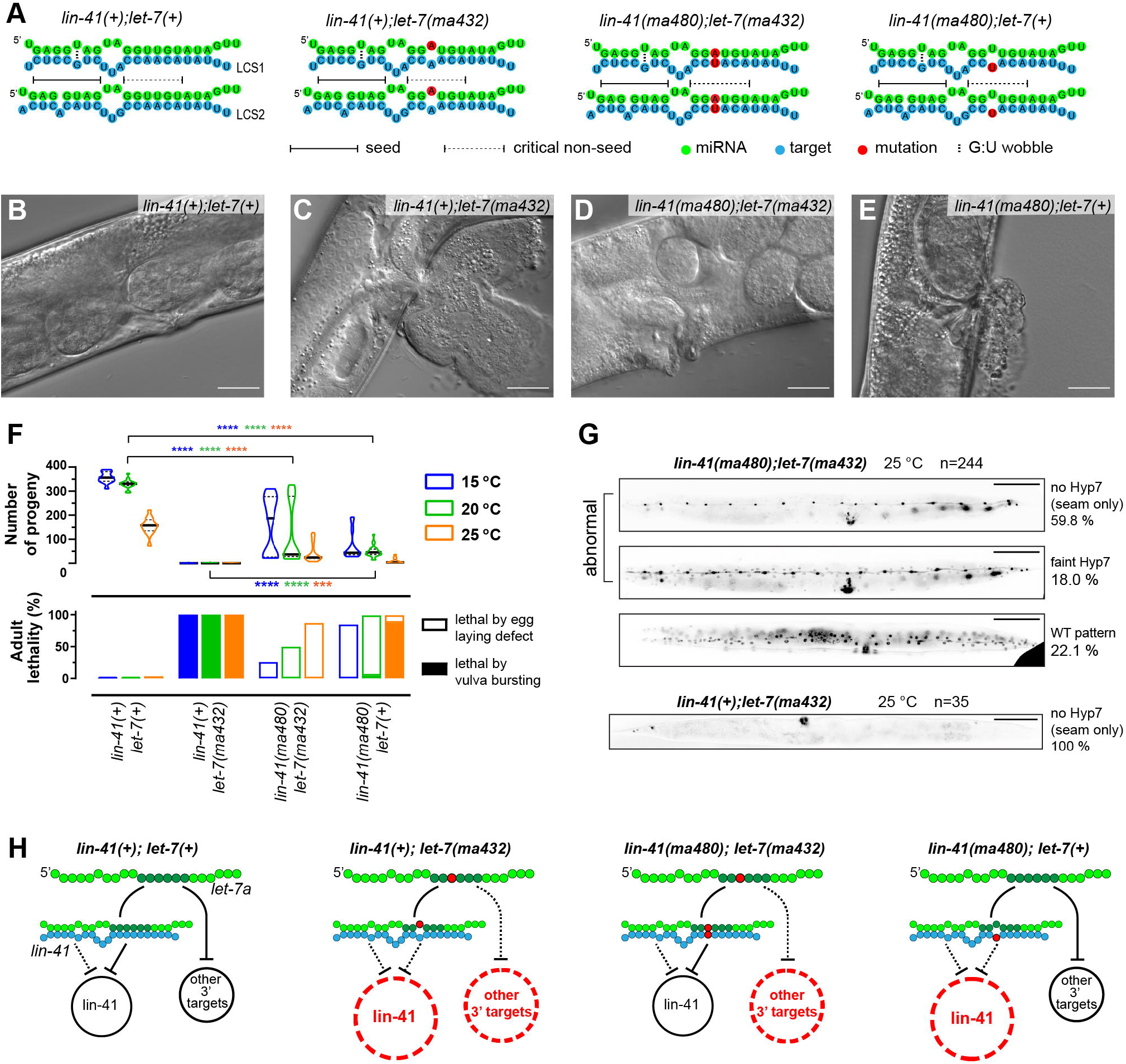
The *let-7a* critical non-seed nucleotides confer *in vivo* function by repressing both *lin-41* and additional 3’ targets. **A.** Pairing configurations between *let-7a* and *lin-41* LCS1/2. **B-E.** DIC images of adult vulva regions representative of *lin-41(tn1541*)(**B**), *lin-41(tn1541);let- 7(ma432)*(**C**), *lin-41(tn1541ma480);let-7(ma432)*(**D**) and *lin-41(tn1541ma480)*(**E**) at 25 °C. Scale bars, 25 µm. **F.** Vulva integrity defects reflected by adult lethality (bottom) and the number of progeny (top). Lethal phenotypes are categorized identically to Fig. 2L. **G.** Representative COL- 19::GFP pattern in young adults at 25 °C. Scale bars, 100 µm. **H.** Illustrative models proposing that both *lin-41* and additional 3’ targets are de-repressed in *let-7a* non-seed mutants. See also Figure S4. All the genotypes tested in this figure include the *lin-41(tn1541,gpf)* allele. Details of the phenotypes are available in Table S1.

We sought to verify whether *lin-41* is de-repressed in the *let-*7a critical non-seed mutants and whether the de-repression causes the *lf* phenotypes. We used the U13A mutant to represent the critical non-seed nucleotides due to its severe phenotypes and the highest conservation of g13. At late L4 stage of WT, endogenously tagged GFP::LIN-41 is undetectable in hypodermal cells due to *let-7a* repression (Fig. S4A) (Spike et al., 2014). In contrast, in *let-7(ma432,g13),* we observed elevated peri-nuclear expression of GFP::LIN-41 in seam and Hyp7 cells, suggesting the de-repression of *lin-41* (Fig. S4B). To confirm that the *lin-41* de-repression in *let-7(ma432,g13)* results specifically from disruption of g13 pairing, we restored the native configurations between *let-7(ma432,g13)* and *lin-41* by introducing compensatory mutations at both LCSs in *lin-41* 3’ UTR (*ma480,t13*) (Fig. 4A), and did not detect the abnormal GFP::LIN-41 expression in *lin- 41(ma480,t13);let-7(ma432,g13)*, indicating the restoration of *lin-41* repression by re-establishing WT 3’ non-seed configuration (Fig. S4C). Moreover, using qRT-PCR, we confirmed that the level of *lin-41* was de-repressed in *let-7(ma432,g13)* compared to the WT, and restored to normal levels in *lin-41(ma480,t13);let-7(ma432,g13*) (Fig. S4E). We found that in *lin-41(ma480,t13);let- 7(ma432,g13)* animals, the *let-7(lf)* phenotypes caused by the *ma432* mutation were substantially rescued, based on the reduced lethality, increased number of progeny, and normal adult alae (Fig. 4B-F, Table S1). The rescue of *let-7(ma432,g13)* phenotypes by restoring non-seed pairing indicates that de-repression of *lin-41* is the major contributor to the *lf* phenotypes of *let-7a* critical non-seed mutants.

In parallel, we found that when present with WT *let-7a*, *lin-41(ma480,t13)* exhibited elevated peri-nuclear expression of GFP::LIN-41, suggesting that disrupting the critical non-seed pairing by mutations in the target also causes *lin-41* de-repression (Fig. S4D). As expected, we also observed vulva integrity defects and retarded heterochronic phenotypes in *lin-41(ma480)* (Fig. 4E-F, Table S1), supporting the conclusion that 3’ non-seed pairing between *let-7a* and *lin-41* is essential for maintaining the robust repression of *lin-41*.

### Disruption of 3’ non-seed pairing of *let-7a* results in de-repression of additional targets including *daf-12* and *hbl-1*

Although *lin-41(ma480,t13);let-7(ma432,g13)* showed alleviated *lf* phenotypes compared to *let-7(ma432,g13),* the double mutant is nevertheless not completely WT and exhibits residual defects in vulva morphogenesis and egg-laying capacity (Fig. 4D, F). Also, COL-19::GFP expression in Hyp7 cells of *lin-41(ma480,t13);let-7(ma432,g13)* adults is reduced at 25 °C, indicating residual retarded heterochronic phenotypes (Fig. 4G). These residual phenotypes suggest that although *lin-41(ma480,t13);let-7(ma432,g13)* restores *lin-41* repression, additional *let-7a* targets may still be de-repressed by the 3’ non-seed mutation (Fig. 4H). We refer to this proposed class of *let-7a* targets that require critical non-seed pairing as “3’ targets”. Note that 3’ targets can contain either perfect or imperfect seed pairing to the miRNA. Consistent with the reasoning that the phenotypes of *let-7(ma432,g13)* result from the over-expression of both *lin-41* and other 3’ targets, the *lin-41(ma480,t13)* mutant, where only *lin-41* is over-expressed since the mutation is on the target, exhibits weaker phenotypes than *let-7(ma432,g13)* (Fig. 4F, Table S1).

To identify *let-7a* 3’ targets in addition to *lin-41*, we screened for genes that are de-repressed by the *let-7a* critical non-seed mutation using ribosomal profiling (Ribo-seq), which assesses gene expression on the translational level (Ingolia, 2016). Accordingly, we compared gene expression between WT and *lin-41(ma480,t13);let-7(ma432,g13),* where *lin-41* repression is restored, yet other *let-7a* 3’ targets are expected to be de-repressed. We found that at L4 stage, 351 genes were significantly over-expressed >1.5 fold in *lin-41(ma480,t13);let-7(ma432,g13)* compared to WT. To apply a stringent test for which of these differentially expressed genes are likely to be perturbed as a result of the *let-7a* mutation specifically, and for which genes observed perturbation could be caused by asynchrony in staging between samples, we applied a filter to remove genes that are known to be highly dynamic over the developmental interval encompassing our sample collections (Aeschimann et al., 2017) (see STAR Methods; Table S3). 203 genes passed this filter and were therefore judged to be likely over-expressed due to the *let-7a* mutation specifically (Fig. 5A, Table S3). The translational levels of *lin-41* and its direct downstream genes were not significantly changed compared to WT, indicating that the de-repressed genes in *lin- 41(ma480);let-7(ma432)* are independent of *lin-41* pathway (Fig. S6A) (Aeschimann et al., 2019).

**Figure 5.**
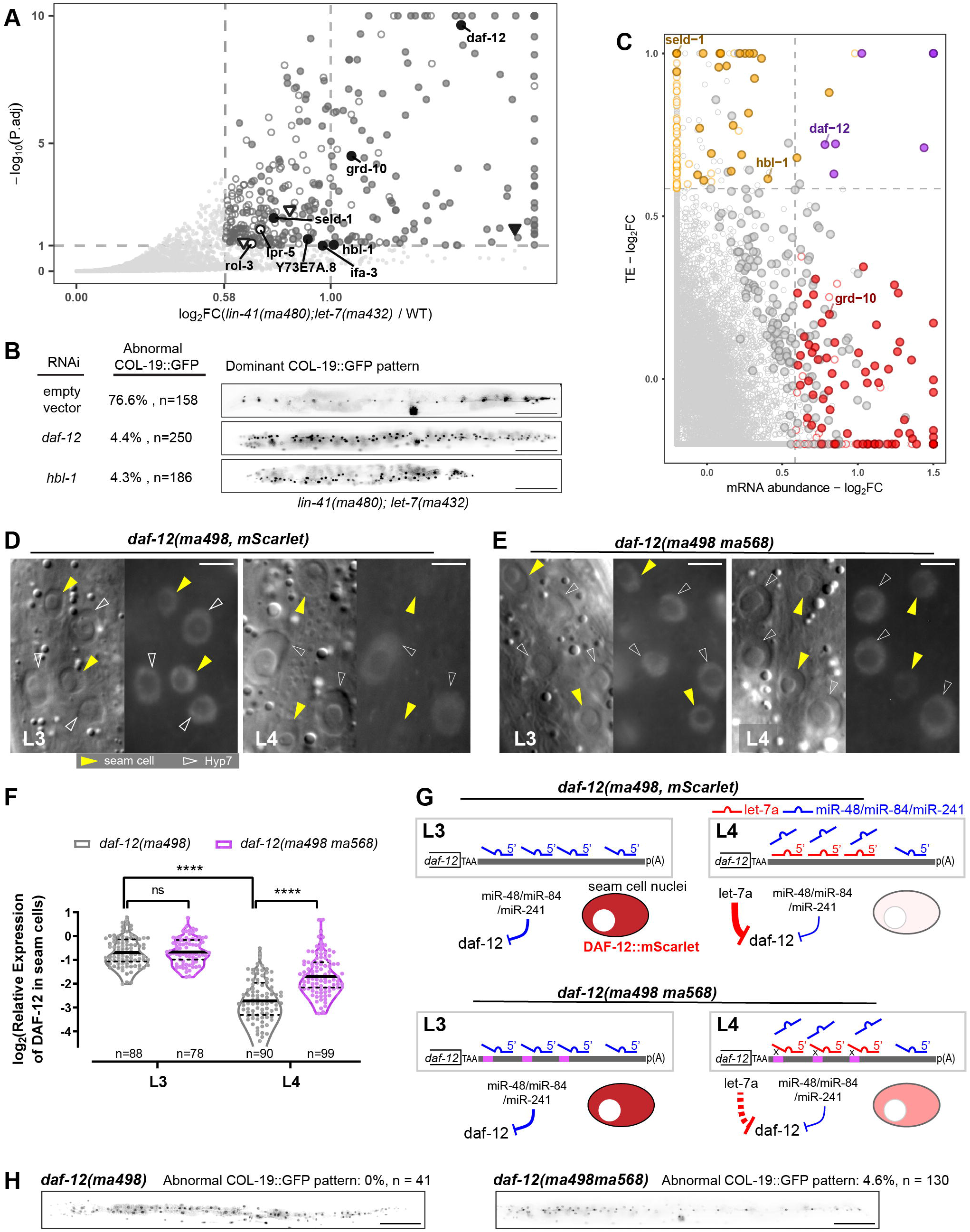
*let-7a* represses a multiplicity of 3’ targets, including *daf-12* and *hbl-1*, through 3’ non-seed pairing. **A.** Differential expression analysis of translatomes of *lin-41(tn1541ma480);let- 7(ma432)* and *lin-41(tn1541)*. Statistical significances were tested by *DESeq2.* Hollow points, developmentally dynamic genes for which the observed perturbation could be caused by asynchrony in staging between samples, independently of *let-7a* genotype; solid points, genes for which the observed perturbation is judged to likely reflect an effect of the *let-7a* mutation specifically (see STAR Methods; Table S3). Circular points with text labels, genes with predicted *let-7a* 3’ sites; triangle points, genes containing only *let-7a* seed-only sites. **B.** Retarded COL- 19::GFP patterns characteristic of *lin-41(ma480);let-7(ma432)* young adults at 25 °C under empty vector condition, *daf-12(RNAi)* or *hbl-1(RNAi)*. Scale bars, 100 µm. **C.** Fold changes of mRNA abundance and translational efficiency (TE). Red points, genes with significantly increased mRNA abundance (FC > 1.5, P.adj < 0.1 by *DESeq2*). Orange points, genes with significantly increased TE (FC > 1.5, P < 0.1 by t-test). Purple points, genes with both significantly increased TE and mRNA abundance. Circled points, genes with significantly increased RPFs in (**A**). Predicted *let- 7a* 3’ targets are text-labeled. **D-E.** DIC (left) and fluorescent (right) microscopy images showing expression of DAF-12::mSCARLET at L3/L4 stages at 25 °C representative of *lin-41(tn1541);daf- 12(ma498)* (**D**) and *lin-41(tn1541);daf-12(ma498ma568)* (**E**). Scale bars, 5 µm. All fluorescent images were generated with identical exposure and processing conditions. **F.** Quantification of relative fluorescent intensity of DAF-12::mSCARLET in seam cell nuclei relative to the adjacent Hyp7 nuclei. The numbers of seam cells quantified per condition are shown under corresponding violin plots. 22 and 28 animals were scored for *daf-12(ma498)* at L3 and L4, and 23 and 31 animals were scored for *daf-12(ma498ma568)* at L3 and L4, respectively. **G.** Illustrative models for the *daf-12* repression patterns and effects at L3/L4. **H.** Representative expression patterns of COL-19::GFP in adult animals. Scale bars, 100 µm. All the genotypes tested in this figure include the *lin-41(tn1541,gpf)* allele. Data in **A** and **C** represent 3 biological replicas. See also Figure S6 and S7.

We next sought to identify potential direct *let-7a* 3’ targets among the over-expressed genes by identifying LCSs in their 3’ UTRs. We developed a computational approach to predict *let-7a* sites involving both seed and critical non-seed complementarity as previously described (Veksler-Lublinsky et al., 2010). We firstly identified candidate sites based on seed complementarity to *let- 7a* allowing for both perfect and imperfect pairing at g2-g7, and then extended these sites to check whether they also included at least 3 consecutive base pairs in g11-g16, among which g13 must be paired. We identified 624 genes whose 3’ UTR contain *let-7a* 3’ sites in *C. elegans,* among which 8 genes are over-expressed in the g13 mutant, including the heterochronic genes *daf-12* and *hbl-1* (Fig. 5A, Table S4).

The 3’ UTRs of *daf-12* and *hbl-1* contain multiple *let-7a* sites with complementarity to the critical non-seed region of *let-7a* (Fig. 5A, S6B-D). We reasoned that the de-repression of *daf-12* and/or *hbl-1* could contribute to the residual heterochronic phenotypes of *lin-41(ma480,t13); let- 7(ma432,g13)*. To test this supposition, we used RNAi to knock down *daf-12* or *hbl-1* during larval development of *lin-41(ma480,t13);let-7(ma432,g13)*, and assayed for suppression of the residual *lf* phenotypes. We found that knocking down either *daf-12* or *hbl-1* by RNAi could rescue the abnormal COL-19::GFP pattern in *lin-41(ma480,t13);let-7(ma432,g13)* (Fig. 5B), suggesting that de-repression of *daf-12* and *hbl-1* contributes to the residual retarded phenotypes of *lin- 41(ma480);let-7(ma432)*, and that the critical non-seed pairing of *let-7a* is required for the repression of not only *lin-41* but also additional targets, including *daf-12* and *hbl-1* for the L4-to-adult transition.

Previous transgenic reporter analyses suggested that *daf-12* can be repressed by *let-7a* as well as the family paralogs *miR-48/miR-84/miR-241* during larval development (Grosshans et al., 2005; Hammell et al., 2009). To gather additional evidence in support of the role of *let-7a* in the down-regulation of DAF-12 expression from the endogenous *daf-12* locus during later developmental stages when *let-7a* is abundantly expressed, we quantified the level of expression of an endogenously-tagged fluorescent reporter *daf-12(ma498,*mScarlet) (Ilbay and Ambros, 2019), and confirmed that endogenous DAF-12::mSCARLET expression is significantly down- regulated in seam cells between the L3 and L4 stages, which confirms the *in vivo* repression of *daf-12* by *let-7a* in later WT development (Fig. 5D, F, S7B-C).

As a further test of whether 3’ pairing to *let-7a* is required for the developmental down- regulation of DAF-12, we employed CRISPR/Cas9 editing of the endogenous *daf- 12(ma498,*mScarlet) 3’ UTR to disrupt the LCS sequences complementary to *let-7a* g11-g13 (Fig. S7A). We found that compared to WT, the L4-specific reduction of DAF-12::mSCARLET in seam cells was alleviated in the mutant *daf-12(ma498ma568)* (Fig. 5E-G). As a comparison, no significant difference of DAF-12::mSCARLET was observed between WT and *daf- 12(ma498ma568)* at the L3 stage, when *let-7a* is not abundantly expressed. Thus, we conclude that 3’ non-seed pairing of *let-7a* contributes to robust repression of *daf-12* at the L4 stage.

For the L4-to-adult cell fate transition, *daf-12(ma498ma568)* exhibits abnormal COL-19::GFP expression with a penetrance of 4.6% (Fig. 5H). This retarded phenotype of *daf-12(ma498ma568)* indicates that the de-repression of *daf-12* observed in *lin-41(ma480,t13);let-7(ma432,g13)* animals could contribute to the residual phenotypes of *lin-41(ma480);let-7(ma432)*, while the low penetrance supports the supposition that unlike *lin-41* as a major contributor, the L4-to-adult cell fate transition in WT requires the proper down-regulation of multiple 3’ targets, including *daf-12 and hbl-1* (Fig. S7E).

### Repression of *let-7a* 3’ targets can involve translational suppression and/or mRNA decay

miRNA-mediated post-transcriptional gene regulation can occur via translational repression or mRNA destabilization, indicated by decreased translational efficiency (TE) or reduced mRNA abundance, respectively (Bazzini et al., 2012; Giraldez et al., 2006). It has been shown that *let- 7a* can regulate *lin-41* through a combination of both modes (Bagga et al., 2005; Nottrott et al., 2006). To assess the target repression mechanisms associated with the *let-7a* 3’ non-seed region, we analyzed our Ribo-seq data in combination with RNA-seq from *lin-41(m480,t13);let- 7(ma432,g13)* larvae (Fig. 5C, S7, Table S3). Previous reports showed that miRNAs may cause deadenylation before mRNA decay, thus RNAseq libraries from poly(A)-enriched mRNA may potentially have a bias against deadenylated but otherwise stable mRNA (Djuranovic et al., 2012; Wu et al., 2006). We thus used ribosomal RNA depletion with anti-sense oligos specifically designed for *C.elegans* to enrich the mRNA (Duan et al., 2020b). We found that the set of de- repressed *let-7a* 3’ targets in *lin-41(ma480,t13);let-7(ma432,g13)* include examples of significantly increased mRNA abundance (e.g., *grd-10*) or TE (e.g., *hbl-1*), or a combination of both modes (e.g., *daf-12*) (Fig. 5C). Thus, target recognition involving the *let-7a* non-seed region can elicit inhibition of TE or mRNA stability, or both, likely depending on context.

### Continuous pairing at g11-g15 is required to compensate for the imperfect seed pairing between *let-7a* and *lin-41*

The involvement of 3’ pairing to compensate for imperfect seed pairing can be essential for *in vivo* targeting efficacy for certain miRNA-target interactions, but how the base-pairing at different positions contributes to this compensatory effect remains unclear (Bartel, 2018). Meanwhile, detailed analyses of how 3’ pairing contributes to endogenous target repression in the context of perfect seed pairing are lacking. To investigate how the 3’ pairing coordinates with seed pairing in endogenous target repression, we systematically introduced mutations in the *lin- 41* 3’ UTR designed to produce various configurations of seed and non-seed pairing with *let-7a* (Fig. 6). Importantly, all the configurations were constructed by mutating both LCSs in *lin-41* and were tested in WT *let-7a* background, so the phenotypes observed should result from de-repression of *lin-41* without confounding effects from other *let-7a* targets.

**Figure 6.**
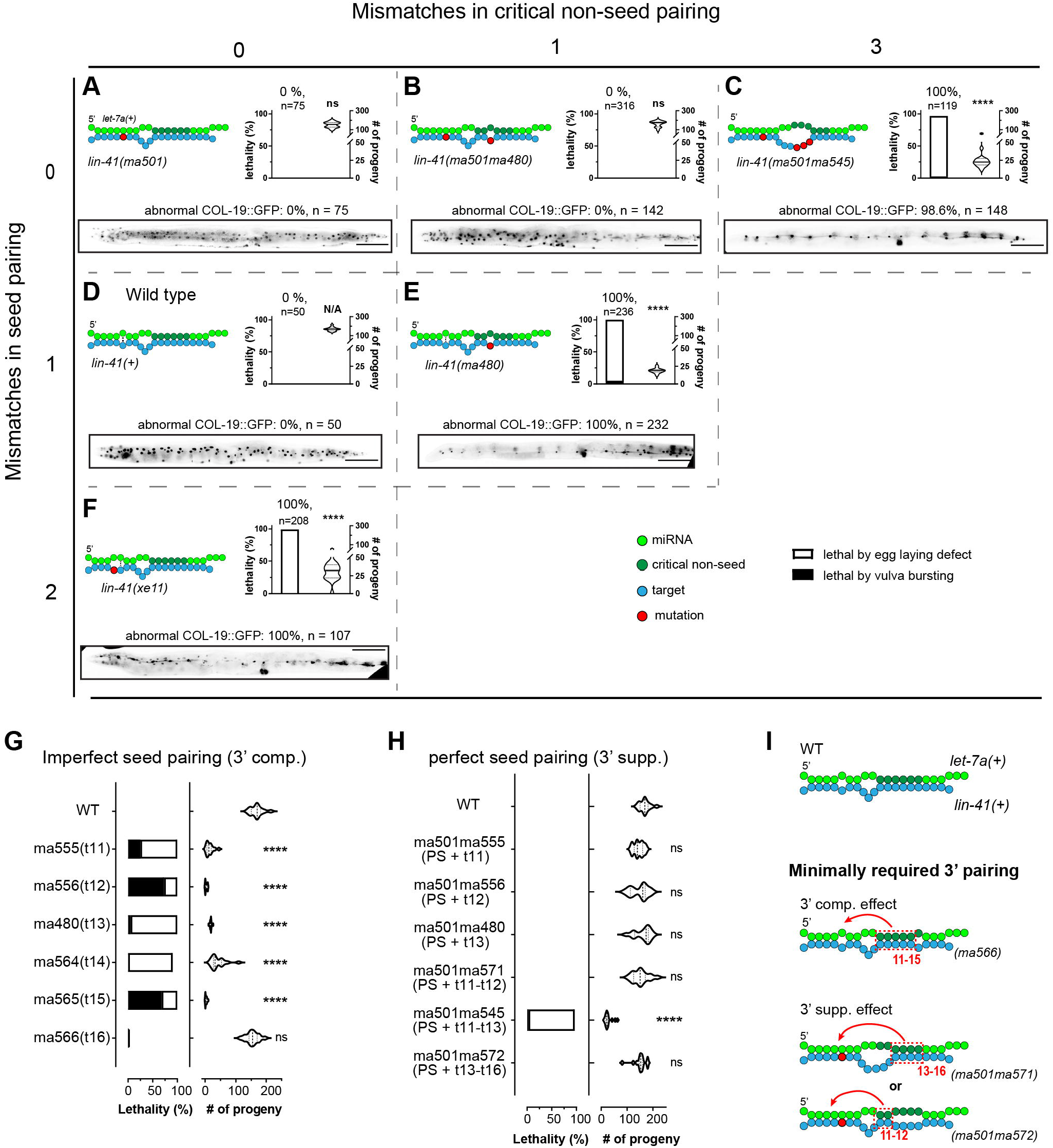
Critical 3’ non-seed pairing and seed pairing together contribute to the function of *let-7a* in repressing *lin-41.* **A-F.** Pairing configurations of *let-7a* to the *lin-41* LCSs, and associated phenotypes for *lin-41(ma501)* (**A**)*; lin-41(ma501ma480)* (**B**); *lin-41(ma501ma545)* (**C**); *lin-41(+)* (**D**); *lin-41(ma480)* (**E**); and *lin-41(ex11)* (**F**), arranged in a matrixed panel based on mismatch numbers in seed (vertical axis) and critical non-seed region (horizontal axis). Each panel includes interacting pattern (top left), vulva defect in terms of young adult lethality and number of progeny (top right), and the heterochronic phenotype (bottom). Scale bars, 100 µm. **G-H.** Vulva integrity of *lin-41* LCSs mutants with 3’ pairing mismatches in the context of imperfect seed pairing (**G**) and perfect seed pairing (**H**). **I.** Illustrative models depicting the minimal pairing requirement for 3’ compensatory and 3’ supplemental effects. All statistical significance indicates comparison with WT. All phenotypes were tested at 25 °C. All the *lin-41* alleles tested in this figure were generated from the *lin-41(+)* without endogenous tag. Details of the phenotypes are available in Table S1.

A previous study employing mutant *lin-41* LCSs constructs with normal imperfect *let-7a* seed matches combined with 3’ pairing restricted to g13-g16, found that the g13-g16 pairing alone is not sufficient to compensate for the imperfect seed pairing between *let-7a* and *lin-41* (Brancati and Grosshans, 2018). Our finding noted above shows that *lin-41(ma480,t13)* exhibited strong *lin-41 gain-of-function, gf* phenotypes (Fig. 6E), suggesting that even quite extensive 3’ pairing (in this case g11-g12 pairing combined with g14-g19 pairing) is insufficient for *lin-41* repression with imperfect seed. To precisely delineate the extent of 3’ pairing required for full *let-7a* repression of *lin-41* in the context of the normal imperfect seed matches, we generated a series of *lin-41* 3’ UTR mutants, each carrying single nucleotide mutations in both *lin-41* LCSs designed to disrupt pairing at a defined position (t11 to t16) (Fig. 6G). We found that disrupting base pairing at any single position in the g11-g15 complementary region results in *lin-41 gf* phenotypes, indicating that continuous pairing from g11-g15 is required to compensate for the imperfect seed pairing between *let-7a* and the *lin-41* (Fig. 6I).

To determine whether the requirement of the g11-g15 pairing for *let-7a* to repress *lin-41* is to compensate for the imperfect seed match characteristic of the WT *lin-41*-*let-7a* interaction, we mutated both LCSs of the t11, t12, and t13 *lin-41* mutants to permit consecutive g2-g8 Watson-Crick pairing (PS) to *let-7a*, whilst maintaining the single 3’ mismatch in each case. We found that the resulting compound mutants *lin-41(ma501ma555,PS+t11), lin-41(ma501ma556,PS+t12)* and *lin-41(ma501ma480*,*PS+t13)* exhibited substantial rescue of the *gf* phenotypes of *lin- 41(ma555,t11), lin-41(ma556,t12)* and *lin-41(ma480,t13),* respectively (Fig. 6B, H). This finding suggests that the requirement for pairing across g11-g15 in large part reflects the compensation for imperfect seed pairing of *let-7a* to the *lin-41* WT LCSs.

### Both seed and 3’ non-seed pairing of *let-7a* are required for full repression *lin-41*

The suppression of the strong *lin-41 gf* phenotypes of single nucleotide mismatches at t11, t12, or t13 by modifying the LCSs to permit perfect seed pairing (Fig. 6H) suggests that the requirements for 3’ pairing may be less stringent with a perfect seed match to *let-7a* compared to the imperfect seed. To test whether 3’ pairing in the g11-g13 region is dispensable in the context of perfect seed pairing, we combined *ma501* with a compound mutation (*ma545*) which simultaneously mismatches g11-g13 of *let-7a* (Fig. 6C). Strikingly, we found that *lin- 41(ma501ma545,PS+t11-13)* exhibited strong vulva integrity defects and retarded COL-19::GFP expression, indicating that *lin-41* was de-repressed despite the perfect seed pairing to *let-7a* (Fig. 6C). This result indicates that even with perfect seed complementarity, 3’ non-seed pairing in the region of g11-g13 still contributes critically to the functional targeting of *lin-41.* We also combined *ma501* with compound mismatch mutations at *t11-t12* and *t13-t16*. Interestingly, the resulting mutants *lin-41(ma501ma571,PS+t11-t12)* and *lin-41(ma501ma572,PS+t13-t16)* did not exhibit observable *lin-41(gf)* phenotypes (Fig. 6H). Taken together, we suggest that in the context of perfect seed pairing, full *lin-41* repression by *let-7a* can occur provided that there is sufficient 3’ pairing at g11-g12 as is the case for *lin-41(ma501ma572*), or g13-g16 as is the case for *lin- 41(ma501ma571*) (Fig. 6I).

The above results, which delineate the pairing requirements at specific *let-7a* 3’ nucleotides for full repression in the context of perfect seed match suggests that *let-7a* family paralogs *miR- 48, miR-84* and *miR-241*, which differ from *let-7a* substantially in their -3’ sequences, should not be able to repress *lin-41* even with engineered perfect seed pairing, even though the expression levels of the combined paralogs have been shown to exceed the level of *let-7a* at L4 stage in *C. elegans* (Nelson and Ambros, 2021) (Fig. 7A, C). Accordingly, *lin-41(ma501,PS);let- 7a(ma393,null)* exhibited vulva integrity defects nearly as severe as *let-7a(ma393,null)*, indicating that *let-7*a family paralogs only render a minor degree of *lin-41* repression with the engineered “perfect-seed-only” configuration (Fig. 7B). This result further indicates that without 3’ critical non-seed pairing, the “perfect-seed-only” pairing configuration is not sufficient to support fully repression of *lin-41*.

**Figure 7.**
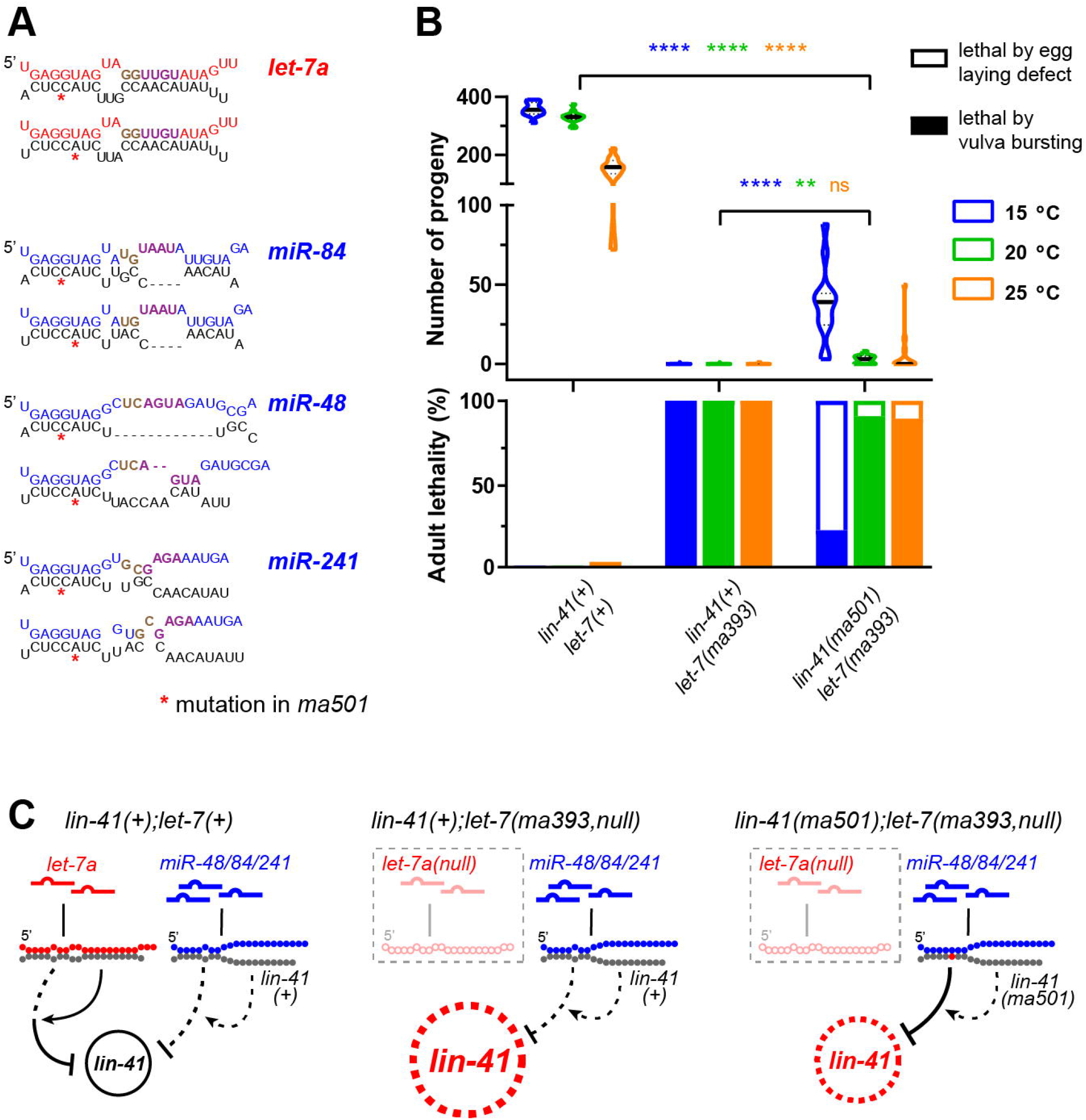
Perfect seed pairing alone is insufficient for *let-7a* family paralogs to confer full repression of *lin-41*. **A.** Predicted pairing configurations of *let-7a* and family paralogs *miR- 48/84/241* to LCS1 (top) and LCS2 (bottom) of *lin-41(ma501)* transcribed from RNAhybrid (Rehmsmeier et al., 2004). Requirements for minimal supplemental pairing (t11-12 pairing or t13- t16 pairing; Fig. 6I) are indicated by brown and purple. Note that *miR-84* is predicted to pair with t11-t12 at LCS2, which should not interfere with the related conclusion. **B.** Vulva integrity defects based on adult lethality (top) and number of progeny (bottom) in *lin-41(tn1541), lin-41(tn1541);let- 7(ma393,null)* and *lin-41(tn1541ma501);let-7(ma393)*. **C.** Schematic diagrams of predicted targeting configurations. Details of the phenotypes are available in Table S1.

The findings demonstrating the importance of non-seed pairing for *lin-41* repression by *let- 7a* prompted us to test whether 3’ non-seed pairing could be sufficient even in the presence of severely-compromised seed pairing. We used a previously reported allele *lin-41(xe11)*, which creates an additional seed mismatch to *let-7a* (Ecsedi et al., 2015). We confirmed that *lin- 41(xe11)*, which has two consecutive seed mismatches, exhibits strong vulva integrity defects and retarded heterochronic phenotypes (Fig. 6F), indicating that seed pairing is also not functionally dispensable, even in the context of extensive 3’ pairing.

## Discussion

### A six-nucleotide critical supplementary pairing region of *let-7a*, involving g11-g16

It has been shown that the 3’ non-seed region of miRNAs can interact with targets and that 3’ non-seed pairing has been implicated in compensating for weak seed pairing, for seed family isoform specificity, and target-dependent miRNA degradation (TDMD) (Bartel, 2018; Pawlica et al., 2020). However, no systematic study has been performed to test the contribution of individual 3’ non-seed nucleotide to the functional efficacy of miRNA-target interactions in intact animals. The functional architecture of base pairing configurations is particularly interesting for miRNAs with evolutionarily conserved 3’ non-seed sequences (e.g., *let-7a*), and evolutionarily conserved miRNA-target interactions (e.g., *lin-41::let-7a*). In this study, we utilized CRISPR/Cas9 genome editing for a systematic genetic investigation of the *in vivo* function of individual 3’ non-seed nucleotide of *let-7a* in the regulation of *lin-41* and other targets in *C. elegans*.

One noteworthy finding from our studies is that each nucleotide at g11-g16 critically contributes to the *in vivo* function of *let-7a*. A recent biochemical study using RNA bind-n-seq (RBNS) with human miRISC also shows that pairing to g11-g16 of *let-7a* is particularly crucial for enhancing the affinity between miRISC and oligonucleotide targets (McGeary et al., 2021). Meanwhile, we found that within g11-g16, the g11-g13 subregion is particularly important to *let- 7a in vivo* function. This finding also agrees with the RBNS results, which show that for 4- nucleotides blocks of target complementarity scanned along the *let-7a* 3’ region, pairing at g11- g14 was the most potent contributor to enhanced target affinity (McGeary et al., 2021). Therefore, both our genetic analysis in *C. elegans* and the RBNS analysis of human AGO2 agree on a 6 nt critical 3’ pairing region (g11-g16) of *let-7a*, expanded by 2 nt compared to the g13-g16 region previously identified by biochemical analysis and co-crystallized structure of miRISC-target complex (Grimson et al., 2007; Sheu-Gruttadauria et al., 2019b).

Our results stand in contrast with a previous study which indicates functional insignificance of the 3’ non-seed region of *let-7a* in *C. elegans*, where a compound mutation at g10, g12 and g13 of *let-7a* exhibited relatively mild developmental defects (Zhang et al., 2015). We note that the results from Zhang et al. (2015) were obtained using exogeneous high-copy transgenes to express mutant *let-7a* miRNAs in the genetic *null* background, and we suggest that over- expression of the transgenic miRNA rescued the poor intrinsic efficacy of the mutant miRNA in Zhang et al. (2015). This contradiction also highlights the importance of using genome engineering of endogenous miRNA loci so that repression efficacy can be assessed in the context of native gene dosage.

Our results suggest that *let-7a-*miRISC may adopt a previously unanticipated conformation that enables g11-g16 to duplex with target. Structural analyses of human miRISC reveal that duplexing of the miRNA seed and target triggers conformational changes of AGO that enable 3’ pairing at g13-g16, whilst g11 and g12 were structurally hindered from duplexing with the target (Sheu-Gruttadauria et al., 2019b). We note that the structural studies thus far have been for instances of perfect seed pairing. By contrast, for the evolutionarily conserved *let-7a* target *Trim71/lin-41,* the LCSs are characterized by imperfect seed pairing accompanied by 3’ pairing involving g11 and g12 in many species, and in some instances even the specific configuration of seed mismatches is conserved (i.e., the g6:t6-GU pair and t5-bulge are conserved across *24 Caenorhabditis* species (Nelson and Ambros, 2021)). We suggest that these mismatched seed configurations may promote the AGO conformation that favors the pairing of g11-g12, in addition to the previously identified g13-g16 pairing. Meanwhile, it has been shown that the miRISC conformation can be affected by certain 3’ pairing configurations. For example, an alternative AGO conformation can be induced when 3’ pairing extends to the 3’ end of the miRNA in the context of TDMD (Sheu-Gruttadauria et al., 2019a). We therefore suggest that *let-7a-*miRISC may adopt a characteristic novel conformation that reflects simultaneous accommodation of both the imperfect seed pairing and the extended 3’ helix.

Although our results confirm that imperfect seed match can be compensated by extensive 3’ pairing, we confirm that the seed pairing between the miRNA and target is also essential for miRISC activity, as *lin-41* LCSs with two nucleotide seed mismatches displayed strong *lin-41 gf* phenotypes despite extensive 3’ pairing at g11-g19 (Fig. 6F). Similarly, disrupting either the seed or 3’ complementarity results in de-repression of *daf-12* in *C. elegans* (Fig. S7D) (Aeschimann et al., 2017). These results suggest that maintaining a certain length of duplex in both the seed and 3’ pairing are critical for *let-7a* repression, at least for *lin-41* and *daf-12*.

### Contribution of the 3’ non-seed pairing to the target specificity of *let-7* family paralogs

Broughton et al (2016) reported that the *C. elegans let-7a* family paralog *miR-48,* which does not regulate *lin-41* in WT, can substitute for *let-7a* in repressing *lin-41* if the *lin-41* LCSs were replaced with a site with a perfect seed match plus extensive 3’ pairing to *miR-48*. It was proposed that the weak seed pairing of *let-7a* to *lin-41* necessitates the 3’ pairing, which differs remarkably between *let-7a* and *miR-48*, and consequently determines the targeting specificity whereby *lin-41* is regulated by *let-7a,* not *miR-48* (Brancati and Grosshans, 2018). Thus, as a general principle, 3’ non-seed pairing combined with imperfect seed pairing is a powerful determinant of specificity for miRNAs of the same seed family. This model is supported by our finding that *miR-84,* the closest family paralog of *let-7a* in *C. elegans*, does not functionally substitute for the absence of *let-7a*. We suggest that our result provides a particularly stringent test of the model because *miR- 84* is expressed at levels and with temporal profile identical to WT *let-7a*; and unlike *miR-48* in the previous reports, which differs from *let-7a* at every non-seed position, *miR-84* has only 5 nt different from *let-7a* (Fig. S1A). We also confirm that the *in vivo* functional distinction between *let- 7a* and *miR-84* primarily reflects the specificity of *let-7a* for targeting *lin-41*, as well as with other targets that require 3’ non-seed pairing (Fig. S5).

Meanwhile, 3’ non-seed pairing may also contribute to the spatiotemporal specificity between *let-7a family* (and possibly other miRNA families) isoforms. In *C. elegans*, *hbl-1* is synergistically repressed by *let-7a* paralogs at L2-to-L3 transition, and by *let-7a* at L4-to-adult transition (Abrahante et al., 2003; Ilbay and Ambros, 2019). Notably, *hbl-1* 3’ UTR contains sites expected to be specifically recognized by *let-7a* (imperfect seed + 3’), which enables the *let-7a*-specific repression of *hbl-1* in particular tissues and at late larval stages when *let-7a* is expressed (Fig. S6D). Meanwhile, *hbl-1* 3’ UTR also has sites that can be recognized by any paralog in *let-7 family* (perfect seed), these sites may mediate repression of *hbl-1* by *let-7 family* paralogs at earlier larval stages and/or in tissues devoid of *let-7a* expression.

### The necessity of non-seed pairing for target repression in the context of perfect seed pairing

It has long been recognized that perfect seed complementarity can be sufficient, in certain contexts, for miRISC to bind and repress a target (Brennecke et al., 2005; Doench and Sharp, 2004; Wee et al., 2012), implying that 3’ pairing may be required only to compensate for weak seed pairing. However, there is also evidence that 3’ pairing can contribute critically to target regulation for instances of perfect seed matches. For example, evolutionarily conserved target sites with perfect seed plus 3’ pairing (3’ supplemental sites) are prevalent in metazoan clades and appear to be more frequent than the 3’ compensatory sites which have imperfect seed pairing (Friedman et al., 2009). Also, competitive binding experiments reveal that 3’ pairing can enhance miRISC affinity to perfect seed sites and enable miRNA isoforms with identical seeds to compete for target recognition (Xiao and MacRae, 2020).

Broughton et al. (2016) reported findings indicating that in the context of perfect g2-g7 seed pairing, the 3’ pairing is required for the specificity of *C. elegans let-7 family* isoforms in the developmental repression of *lin-41*. Brancati & Grosshans (2018) showed that for *lin-41* LCSs modified to contain a modified *miR-48* site from the *dot-1.1* gene with perfect g2-g8 *let-7a* seed pairing, *let-7a* was unable to fully rescue the *mir-48(null)* phenotypes. Our results further enforce this claim by showing that *let-7 family* isoforms can only confer a limited degree of repression of *lin-41* via perfect seed pairing in the context of native 3’ pairing sequence in *let-7(null)* (Fig. 7). Moreover, we investigated the minimal requirements for 3’ pairing for the full repression of *lin-41* containing LCSs with perfect g2-g8 seed pairing, and confirmed that *let-7a* 3’ pairing is necessary to supplement perfect g2-g8 pairing, but that the functional constraints on 3’ pairing are less stringent for the perfect seed pairing compared to the native imperfect seed pairing (Fig. 6). Importantly, we show that the WT 3’ UTRs of additional *let-7a* targets, including the heterochronic gene *daf-12*, contain sites with perfect seed matches combined with extensive supplementary 3’ complementarity to *let-7a*, and supplemental pairing is essential for the down-regulation of DAF- 12 at later developmental stages (Fig. 5). These findings run counter to the assumption that perfect seed pairing should be sufficient for miRISC to confer repression, and emphasize that in the native context of an intact developing animal, the miRNA 3’ non-seed pairing can be critical for the efficacy of evolutionarily conserved miRNA-target interactions.

It is noteworthy that the LCSs with perfect seed plus 3’ complementarity configuration in *daf- 12* 3’ UTR should allow binding of the *let-7* family paralogs by seed pairing, as well as and *let-7a* by seed plus 3’ supplemental pairing. Indeed, it has been reported that *daf-12* is repressed by *let- 7* family paralogs (*mir-48*, *mir-84*, *mir-241*) at the L2/L3 stage transition, and by *let-7a* at the L4 stage (Grosshans et al., 2005; Hammell et al., 2009) (Fig. 5D-F). Since 3’ pairing can render higher target binding affinity to the miRISC, we suggest that the involvement of 3’ non-seed pairing to *let-7a* may enable more efficacious repression of *daf-12* at the L4 stage than that at earlier stages, which is specific to the expression of *let-7a* miRNA at late L3 (Xiao and MacRae, 2020) (Fig. 5G).

Currently, we can only speculate about the mechanistic basis for this requirement for 3’ pairing in the context of perfect seed match: perhaps certain structural peculiarities of a 3’ UTR (e.g., *lin-41* or *daf-12*) may render *let-7a* “seed-only” pairing kinetically unfavorable compared to “seed plus 3’ supplemental” pairing, or even “imperfect seed plus 3’ compensatory” pairing. It is also possible that repression by *let-7a* requires the recruitment of cofactors that recognize certain miRISC conformational features induced by specific 3’ pairing.

### The multiplicity of *C. elegans let-7a* targets with 3’ non-seed pairing

Previous reports have shown that *lin-41* is the major *let-7a* target whose de-repression underlies the *lf* phenotypes of *let-7a* seed mutants (Ecsedi et al., 2015). In our study, we further confirmed that *lin-41* is the major *let-7a* 3’ target by showing the de-repression of LIN-41 by a g13 mutation *let-7(ma432*), and the rescue of LIN-41 de-repression and the associated *lf* phenotypes by a compensatory mutation *lin-41(ma480)*. These results were assessed using three approaches: (1) an endogenously tagged fluorescent LIN-41 reporter for visualization of cell-specific expression (Fig. S4A-D); (2) qRT-PCR to measure the global level of *lin-41* transcript (Fig. S4E); and (3) ribosome profiling to measure the global level of translation of *lin-41* mRNA(Fig. S6A). Note that in ribosome profiling, we were not able to assay *let-7(ma432)* single mutant for technical reasons that *let-7(ma432)* animals exhibit a strong adult lethality which precludes propagating them in numbers adequate for ribosome profiling. However, a previous report showed that the translational level of *lin-41* was up-regulated around 8 times in the *let-7a(n2853)* seed mutant (Aeschimann et al., 2017). Since *let-7(ma432)* exhibits stronger phenotypes than does *let- 7(n2853)*, we anticipate that *lin-41* should be over-expressed to a greater degree in *let-7(ma432)* than in *let-7(n2853).* Thus, the non-significant RPF changes in Fig.S6A can be interpreted to indicate the restoration of normal *lin-41* repression in *lin-41(ma480);let-7(ma432)* animals.

Our computational analysis identified 624 genes in *C. elegans* that contain 3’ UTR sequences predicted to bind *let-7a* with 3’ non-seed pairing. This finding is consistent with previous high-throughput analyses of miRNA-target chimeric ligation products, which also identified multiple *let- 7a* targets with 3’ non-seed complementarity (Broughton et al., 2016; Grosswendt et al., 2014). By ribosome profiling, we confirmed that 3’ non-seed pairing is critical for the *let-7a* repression of at least 8 genes, and that over-expression of *daf-12* and *hbl-1* contribute to the developmental phenotypes of the *let-7a* g13 mutant (Fig. 5). This finding indicates that in addition to *lin-41*, at least *daf-12* and *hbl-1* are also phenocritical *let-7a* targets in *C. elegans* (Ecsedi et al., 2015). We did not test the phenotypic consequences of the other over-expressed *let-7a* 3’ targets, which are not known to be involved in the heterochronic pathway. These genes could contribute to *let-7a* phenotypes not assayed in this study.

It is curious that mutation of g18, which is one of two 3’ nucleotides of *let-7a* that shows the greatest evolutionary conservation, did not cause visible phenotypes. By ribosome profiling, we found 3 genes that were up-regulated in the *let-7a* g18 mutant and that contain 3’ UTR sites with 3’ complementarity that includes g18, suggesting that they could be *let-7a* targets whose proper repression depends on g18 pairing. However, caution is called for in interpreting these genes as g18 targets because their Wormbase annotations indicate that they undergo developmental up- regulation in the L4 of the WT (Harris et al., 2020) and that the up-regulation that we observed for these genes in the *let-7a* g18 mutant compared to WT could reflect differences in staging between samples. However, published data from translational profiling of *C. elegans* larvae (Aeschimann et al., 2017) show that the translational levels of our candidate g18 targets were not significantly increased from L3 to L4 (Fig. S3E). Moreover, in the g18 mutant, 2 of the putative target genes (*chs-1* and *try-1*) were translationally de-repressed without significant perturbance in mRNA levels (Fig. S3B-C). We thus propose that 3’ pairing to *let-7a* at g18 may contribute a dampening of the rise in translational output of these developmentally up-regulated mRNAs. Interestingly, in the Aeschimann et al., 2017 dataset, these putative g18 targets were not translationally de-repressed in *let-7a(n2853, G5C)* (Fig. S3F), suggesting that the mutation at g18 may be more detrimental to the proper repression of these putative targets than the G5C seed mismatch mutation.

There are limitations to the use of ribosome profiling of whole animals for the confirmation of *in vivo* miRNA targeting. In particular, the data could contain numerous false negatives, especially for genes that are broadly expressed but are only repressed by miRNAs in specific tissues or cells. For example, in our analysis, although we identified 8 target genes with 3’ non-seed complementarity that were up-regulated in the *let-7a* g13 mutant, genes with such complementarity as a class were not statistically enriched (Fig. S6E). Similarly, potential *let-7a* target genes with g18 pairing were also not statistically enriched among the de-repressed genes in the *let-7a* g18 mutant (Fig. S3 and data not shown).

### Constraints of the evolutionary conservation of *let-7a* sequence

The entire *let-7a* sequence is almost perfectly conserved across bilaterian phyla (Fig.1) (Wolter et al., 2017), suggesting phylogenetically ubiquitous involvement of *let-7a* in interactions that constrain the g9-g22 sequence. Our results indicate that the conservation of g11-g16 of *let- 7a* could be driven by phenocritical roles for these nucleotides in the repression of conserved targets. Importantly, a multiplicity of targets with a given pairing configuration can also constraint the miRNA sequence (John et al., 2004). Our finding of the multiplicity of *let-7a* targets with critical non-seed pairing supports this hypothesis that the 3’ pairing could constrain the sequence of g11- g16.

In contrast to the g11-g16 sequence, our data do not appreciably illuminate potential mechanisms for the conservation of *let-7a* 3’ distal nucleotides (g17-g22) or the bridge nucleotides (g9-g10). However, although single nucleotide or compound mutations were phenotypically tolerated in g17-g22, we observed phenotypic effects of a g18 mutation in the context of g16 mutation, suggesting the involvement of 3’ distal nucleotides in repressing targets with obligate mismatches to g14-g16 to *let-7a*. It is also possible that *let-7a* 3’ distal nucleotides could interact with miRISC-associated RNA binding proteins which recognize specific sequences. Our finding that mutations at g9-g10 were also phenotypically tolerated is consistent with the observation that *let-7a* is not predicted to engage in target recognition involving g9-g10 paring in *C. elegans*, and that fully complementary target sites that involve the bridge pairing are rare in vertebrates (Bartel, 2018). Interestingly, miRNA bridge nucleotides can be exposed on the miRISC surface even when they are complementary to the target (Sheu-Gruttadauria et al., 2019b), suggesting that these nucleotides could engage in interactions external to AGO. We hypothesize that g9-g10 of *let-7a* may be constrained by association with conserved RNA binding proteins that recognize these nucleotides, possibly in the context of *let-7a* biogenesis or target binding.

It is also possible that the identity of g9-g10 or g17-g22 nucleotides of *let-7a* could be critical for target repression under conditions not assayed in this study, such as various conditions of stress, and/or required for target interactions unrelated to the phenotypes that we monitored. The latter possibility is supported by our analysis that the g18 mutant displays molecular phenotypes by Ribo-seq and RNA-seq (Fig. S3). Such molecular phenotyping of other distal 3’ nucleotides of *let-7a*, and other ‘silent’ nucleotides of other miRNAs, could reveal otherwise experimentally inaccessible functionalities.

## Supporting information

Key Resources Table

## Acknowledgments

We thank Xantha Karp from Central Michigan University for commenting on the manuscript. This research was supported by funding from NIH grants R01GM088365, R01GM034028 and R35GM131741 (VA). Some *C. elegans* strains were provided by the CGC, which is funded by the NIH Office of Research Infrastructure Programs (P40 OD010440).

## Author Contributions

Conceptualization: YD, IVL, VA; Methodology: YD, IVL, VA; Formal analysis: YD, IVL, VA; Investigation: YD, IVL; Resources: IVL, VA; Data curation: YD; Writing -original draft: YD; Writing-review & editing: YD, IVL, VA; Supervision: VA; Project administration: VA; Funding acquisition: VA

## STAR Methods

### Phylogeny and conservation analysis

Precursor and mature miRNA sequences, and a phylogenetic tree of metazoan species were downloaded from miRbase (v22.1) (Kozomara et al., 2019). An in-house code was developed to identify miRNAs that belong to the *let-7 family* based on the presence of the seed sequence “GAGGUA” at g2-g7 on the mature miRNA. For each species in the collection, the number of total miRNA loci and the number *let-7* miRNA loci were counted. To calculate the distance of each member to the conserved *hsa-let-7a*, its sequence was aligned to the *hsa-let-7a* sequence and the number of mismatches in the alignment was counted. The phylogenetic trees were visualized with the program evolView v2 (He et al., 2016).

117 species encode *let-7a* family isoforms. From each species, the sequence of one, the most similar to *let-7a,* was extracted and used to construct a conservation profile. A similar analysis was done with the *mir-1* and *mir-34* families to obtain their conservation profiles.

### Targeted mutagenesis at the *let-7a* genomic locus

CRISPR/Cas9 genome editing of the *let-*7a locus was performed using the “jump board” strategy previously described (Duan et al., 2020a) on strain *VT3742*, which carries the *let- 7(ma393)* insertion of the “jump board” sequence in place of the *pre-let-7* sequence, as well as the genetic balancer *umnIs25(mnDp1)* and a transgene *oxSi1091* expressing Cas9. Templates for dsDNA HR donors were prepared by cloning the WT pre-*let-7a* and 500 bp of flanking sequence into pCR2.1-TOPO vector. The Q5 mutagenesis kit (NEB, Cat: E0554) was used to generate the mutant plasmids. Double-strand DNA donors were generated from the mutant plasmids by PCR with 73/106 base-pairs flanking the pre-*let-7a,* and the PCR product was purified by ethanol precipitation. Injection mixtures containing final concentrations of 30 ng/µl AltR_Cas- 9_crRNA_INPP4A_1/2 each, 10 ng/µl AltR_Cas-9 _crRNA_dpy-10_cn64 as co-CRISPR marker (Arribere et al., 2014), 75 ng/µl Alt-R tracrRNA (IDT, Cat:1072532), 10 ng/µl each dsDNA donor in 1X duplex buffer (IDT, Cat: 11010301) were injected into the gonad of *VT3742* at young adult stage. F1 dumpy animals were isolated and genotyped by PCR with let-7_SEQ_F5/R5 primers followed by an analysis of the PCR product using restriction digestion using EcoRV (NEB, Cat: N3195) and Sanger sequencing as described in (Duan et al., 2020a). Mutants were backcrossed with *N2* (for strains without *maIs105*) or *VT1367*(for strains with *maIs105*) for at least two generations.

### Targeted mutagenesis at the *lin-41* genomic locus

Ultramer single-strand DNA donors (lengths ranging from 115 nt to 117 nt) with 35 nt flanking homology were obtained from IDT. The injection mixture containing final concentrations of 25 ng/µl AltR_Cas-9_crRNA_lin-41_1/2 each, 15 ng/µl AltR_Cas-9_crRNA_dpy-10_cn64, 95 ng/µl Alt-R tracrRNA (IDT, Cat:1072532), 325 ng/µl each ssDNA donor in 1X duplex buffer (IDT, Cat: 11010301) were incubated at room temperature for 10 min for pre-annealing and injected into the gonad of *EG9615, VT3867* or *VT3873* which contain transgene *oxSi1091* expressing Cas9. F1 dumpy animals were isolated and genotyped by PCR with lin-41_SEQ_F2/R2 primers, followed by an analysis of the PCR product using Sanger sequencing. Mutants were backcrossed with N2 for at least two generations.

### Targeted mutagenesis at the *daf-12* genomic locus

Double-stranded DNA donor (length 641 bp) containing the desired mutations plus 65/78 bp flanking homology was obtained from GENEWIZ. The injection mixture, containing final concentrations of 80 ng/µl AltR_Cas-9_crRNA_daf-12_4, 40 ng/µl AltR_Cas-9_crRNA_dpy- 10_cn64, 223.3 ng/µl Alt-R tracrRNA (IDT, Cat:1072532), and 73.3 ng/µl dsDNA donor in 1X duplex buffer (IDT, Cat: 11010301), was incubated at room temperature for 10 min for pre- annealing and injected into the gonad of *VT4126* which contain *daf-12(ma498ma567, mScarlet* with 3’ UTR InDel) as a *daf-12* “jump board” and transgene *oxSi1091* expressing Cas9 (Ilbay and Ambros, 2019). Genotyping and backcrossing were performed same as the *lin-41* locus.

### Worm culturing and synchronization

*C.elegans* were cultured on nematode growth medium (NGM) and fed with *E. coli* HB101 unless specified. To obtain populations of synchronized developing worms, gravid adults were collected and washed twice with water. Pellets of centrifuged worms were treated with 5 ml 1 M NaOH and 1% (v/v) sodium hypochlorite for 5 min with shaking to obtain embryos, and the embryos were rinsed with M9 buffer three times. The embryos were hatched in 10 ml M9 buffer at 20°C for 16-18 hrs with mild shaking. Hatched L1 larvae were transferred to plates at 30-50 worms per plate and replicate plates were cultured at 15°C, 20°C, or 25 °C for defined periods; samples of the population were examined by microscopy to confirm the developmental stage at the time of harvest.

### Phenotypic assays for vulva defects

The adult lethality characteristic of *let-7(lf)*, which results from rupture of the young adult animal at the vulva, was scored approximately 36 hrs (15°C), 24 hrs (20°C) or 16 hrs (25°C) after the animals reached developmental maturation (when at least 90% of the population had reached the adult stage). The young adult (pre-gravid) with vulva bursting phenotypes was distinguished from those without vulva bursting (matricide due to egg-laying defects) based on the visibility of breached intestinal tissues from the ruptured vulva region. To score viable progeny per adult, young adults were transferred to a fresh plate every 12 hrs until those capable of laying eggs had completed egg-laying. Only hatched eggs were counted.

### Microscopy and heterochronic phenotypes

Differential interference contrast and fluorescent images were obtained by Zeiss.Z1 equipped with ZEISS Axiocam 503 camera. COL-19::GFP patterns were scored using a 10X objective. Adult alae and GFP::LIN-41 images were obtained using a 100X objective. DAF-12::mScarlet images were obtained using a 63X objective. Lateral hypodermal heterochronic cell lineage defects were scored by counting the number of seam cells per side of each animal with the aid of the *wIs51* transgenic seam cell reporter. Patterns of expression of the adult-specific COL-19::GFP reporter were scored using the *maIs105* transgenic reporter. Fluorescent images were processed by ImageJ FIJI (Schindelin et al., 2012). To quantify the DAF-12::mScarlet expression, total fluorescence signal and area of individual seam cell and Hyp7 nuclei were quantified by ImageJ FIJI, and the fluorescence signal density of one posterior and one anterior regions adjacent to each nucleus was measured and used to calculate the background signal for that nucleus.

### RNAi

Overnight cultures of HT115 bacteria expressing dsRNA (Timmons et al., 2001) were transferred to LB broth and shaken at 37 °C until OD_600_ was between 0.4 – 0.8. The bacteria cultures were spread on NGM medium containing 100 µg/ml ampicillin and 1 mM IPTG and induced at room temperature for 24-48 hrs. HT115 bacteria strains for RNAi were obtained from the Ahringer library (Kamath et al., 2003).

### Total RNA preparation

Harvested worms were washed with M9 medium, centrifuged, and the worm pellets were flash-frozen in liquid nitrogen. The worm pellets were thawed and lysed by adding 4X volumes of QIAzol (Qiagen, Cat: 79306) and shaking vigorously at room temperature for 15 min. The total RNA was extracted by the addition of 0.85X volume chloroform, centrifugation, and recovery of the aqueous phase, which was then re-extracted with 1 volume phenol:chloroform:isoamyl alcohol (25:24:1, pH = 5.5). Total RNA was then precipitated by add 1 volume of isopropanol and 0.5 µl GlycoBlue (Invitrogen, Cat: AM9516), followed by incubation at -80°C for at least 30 min, and recovery by centrifugation at 25,000 rcf for 10 min at 4 °C. The supernatants were then removed, and the RNA pellets were subsequently washed twice by 70% (v/v) ethanol, dried in air for 5 min, dissolved in water, and stored at -80°C.

### Fireplex miRNA assay

Synchronized populations of developing worms were cultured at 20 °C and harvested at 8 hrs (L1), 22 hrs (L2), 36 hrs (L3), 46 hrs (L4) and 58 hrs (adult) after feeding. Total RNA was extracted and miRNA levels of *miR-84* and *let-7a* were quantified by FirePlex miRNA assay (Abcam) with customized *C.elegans* miRNA panel following the manufacturer’s instructions. Guava easyCyte 8HT (Millipore) was used for readout. To normalize the quantification, synthetic RNA oligonucleotides with sequences of *miR-84* and *let-7a* (IDT) were serially diluted and subjected to FirePlex miRNA assay. Equal amounts of total RNA were used for all the samples and replicates. The amounts of miRNA in experimental samples were calculated from the standard curve generated from the serial dilution of respective synthetic RNA oligonucleotides.

### qRT-PCR analysis

20 – 25 synchronized worms were cultured at 25 °C and harvested at the mid-L4 stage (30 hrs after feeding). Total RNA was extracted as described above. Reverse transcription was performed using HiFiScript gDNA Removal RT MasterMix (CWBio, Cat: CW2020M) with 100 ng total RNA input. qPCR was performed using UltraSYBR Mixture (CWBio, Cat: CW2601) on ViiA 7 platform.

### Small RNA sequencing

Synchronized populations of developing worms were cultured at 20 °C and harvested at the mid-late L4 stage (45 hrs after feeding). Total RNA was extracted as described above. The small RNA sequencing libraries were constructed using NEBNext multiplex small RNA library prep set (NEB, Cat: E7300), and sequenced by Illumina NextSeq 500 system. The adaptor sequences were trimmed from the 3’ end of the raw reads by *Cutadapt/1.9* using default parameters (Martin, 2011). To quantify the wild type or mutant *let-7a* miRNAs, the trimmed reads were mapped with *Bowtie2/2.3.4.3* to either wild type or mutant *let-7a* sequences indexed with *-c* using parameters *--end-to-end -N 0 --no-1mm-upfront -L 22* (Langmead and Salzberg, 2012). To quantify the total small RNA reads, trimmed reads were size filtered by *Cutadapt/1.9,* and reads with a length between 18-25 bp were kept. The filtered reads were mapped to *C. elegans* genome *WBCel235* by *star/2.7.6a* with default parameters (Dobin et al., 2013). The numbers of reads that mapped uniquely to the genome were used to calculate the RPM of wild type and mutant *let-7a.* Gene counting was done by *subread1.6.2/featureCounts* (Liao et al., 2014). The total number of miRNA reads that mapped uniquely to the genome were used to calculate the RPMs of wild type and mutant *let-7a*.

### Ribosome Profiling

#### - Worm harvesting

Synchronized populations of developing worms were cultured at 20 °C for 45 hrs after feeding. Harvested worms were harvested by M9, washed with water three times, and incubated at room temperature for 10 min to allow digestion of intestinal bacteria. Worms were then pelleted by centrifuge at 4,500 rcf for 2 min at room temperature and residual water was removed until the total volumes were twice as the worm pellets. The samples were then flashed frozen by liquid nitrogen and stored at -80 °C.

#### **-** Monosome preparation

Concentrated lysis buffer was add to each frozen sample to final concentration of 20 mM Tris-HCl (pH = 7.4), 150 mM NaCl, 5 mM MgCl_2_, 0.5X Protease Inhibitor (Sigma, Cat:P2714), 1 mM DTT, 0.1 mg/ml cycloheximide (Millipore, Cat:C4859), 1% (v/v) Triton X100 and 5 U/ml Turbo DNase (Invitrogen, Cat:AM2238) (Ingolia et al., 2012), and worm pellets were kept on ice until fully thawed. Suspended worms were transferred to 400 µm silica beads tube (OPS Diagnostic, Cat: PFAW-400-100-04) and lysed in bead beater homogenizer for 4 min at 4 °C. Lysates were then centrifuged at 25,000 rcf for 10 min at 4 °C, and supernatants were collected. To generate monosomes, RNase I (Invitrogen, Cat: AM2294) was added to a final concentration of 0.2 U per µl of harvested worm pellet. The digestion was incubated at room temperature for 40 min with gentle rotation and then quenched by adding SUPERase RNase inhibitor (Invitrogen, Cat: AM2694) at 4 U per RNase I unit. The lysates were then loaded onto 5-40 % (m/v) sucrose gradients prepared with lysis buffer without Triton X100 and centrifuged at 32,000 rpm for 3 hrs at 4 °C in an SW41Ti rotor (Beckman Coulter, Cat:331362). The sucrose gradients were fractionated using BR-188 Density Gradient Fractionation System with 60% (m/v) sucrose as chase solution, and monosome fractions were collected according to OD_254_ profiles.

#### **-** RPF cloning

3.5 X volumes of QIAzol reagent was added to the gradient fractions containing monosomes, and RNA was extracted and separated by 17.5 % denaturing PAGE. A synthetic RNA oligonucleotide with a length of 30 nt was used as a size marker. The gel was stained by Sybr Gold (Invitrogen, Cat: S11494) at room temperature for 10 min, and a gel slice containing RNA of approximately 30 nt was excised and ground by an RNase-free pellet pestle (Fisher Scientific, Cat: 12-141-364). RNA was extracted from the gel slice by adding 500 µl of 300 mM NaAc (pH = 5.5), 1 mM EDTA, and 0.25% (m/v) SDS and mildly shaking overnight at room temperature (Ingolia et al., 2012). The gel granules were excluded using Spin-X tube filter (Millipore, Cat: CLS8160) and the RNA was then concentrated by ethanol precipitation, dissolved in water, and stored at -80 ^°^C. 5’ phosphorylation and 3’ dephosphorylation were performed with T4 PNK (NEB, Cat: M0201S) following the manufacturer’s instructions, and the products were subjected to phenol/chloroform extraction and ethanol precipitation. cDNA libraries were constructed using QIAseq miRNA Library Kit (Qiagen, Cat:331505 & 331595) following the manufacturer’s instructions, except that the amplifying PCR was conducted with 9-12 cycles, and sequencing was performed using Illumina NextSeq 500 system.

#### **-** RPF data analysis

The adaptor sequences were trimmed from the 3’ end of the raw reads by the *Cutadapt/1.9* using default parameters (Martin, 2011). Reads were size filtered to keep only reads with a length between 26-34 bp (a range that fits ribosome-protected fragments) (Aeschimann et al., 2015). The rRNA and tRNA reads were removed by initially mapping with *Bowtie2/2.3.4.3* to *C. elegans* rRNA and tRNA sequences from *WBCel235* with default parameters (Chan and Lowe, 2009, 2016; Langmead and Salzberg, 2012), and the remaining reads were mapped to the *C. elegans* genome *WBcel235* by *Star/2.5.3* with default parameters. The p-offset of the 5’ end of the mapped reads and the monosome periodicity were determined by *plastid/0.4.8* as quality control, and gene counting on exons was generated by *plastid_cs/0.4.8* with p-offset adjusted (Dunn and Weissman, 2016; Santos et al., 2019). Differential expression analysis was performed by *DESeq2* with default parameters (Kucukural et al., 2019; Love et al., 2014). Genes with max raw count smaller than 10 were excluded from the analysis. Volcano plots were generated using *ggplot2* (Hadley, 2016).

To identify genes whose observed expression perturbation between mutant and wild-type samples could be due to slight differences in staging between sample populations, the gene expression changes between the mutants and wild-type were compared with the expression changes between different developmental time points published previously (Aeschimann et al., 2019). Of particular interest were genes reported to change in N2 between 28 and 30 hours at 25°C, and/or between 30 and 32 hours at 25°C (Aeschimann et al., 2017), which corresponds to the L4 harvest time point at 20 °C used in this study. A gene was flagged as potentially non-specifically perturbed owing to imperfect synchrony if the normalized RPF fold change that we observed between mutant and wild-type was smaller than 1.5 times of the reported RPF fold change between two developmental time points in the wild-type (Aeschimann et al., 2017).

### RNAseq of mRNA after ribosomal RNA depletion

Worm samples for RNA-seq were aliquoted from the ribosome profiling harvests before the lysis step and frozen separately. The total RNA was extracted as described above. To enrich for mRNA, rRNA was depleted as described in (Duan et al., 2020b). rRNA-depleted mRNA samples were then purified by RNA Clean & Concentrator-5 Kit (ZYMO, Cat: R1015) (Zhang et al., 2012). The RNA-seq libraries were constructed by NEBNext Ultra II RNA Library Prep kit (NEB, Cat: E7775, E7335, E7500) and sequenced by Illumina NextSeq 500 system (Duan et al., 2020b).

### RNA-seq and translational efficiency data analysis

The adaptor sequences were trimmed from RNA-seq data and reads shorter than 15 nt were filtered out from the analysis by *Cutadapt/1.9*. tRNA, signal recognition particle RNA (srpR), and residual cytoplasmic rRNA reads were removed by initial mapping with *Bowtie2*/*2.3.4.3*, and the remaining reads were mapped to *C. elegans* genome *WBcel235* by *Star/2.5.3* with default parameters. Gene counting was done by *featureCounts(Subread/1.6.2)* (Liao et al., 2014). Differential expression analysis was performed by *DESeq2* with default settings (Kucukural et al., 2019; Love et al., 2014). To calculate the translational efficiency (TE), the genes with max raw count smaller than 10 for either RNA-seq or ribosome profiling were excluded from the analysis. The TE was calculated by dividing normalized ribosome profiling counts by normalized RNA-seq counts for each replica. Significance was calculated by the Student t-test. Volcano plots were generated using *ggplot2*.

### Target prediction

3’ UTR sequences of C. elegans genes were downloaded from the WormBase Parasite website (Howe et al., 2016; Howe et al., 2017) for the genomic version *WBcel235*. The UTRs were sorted and filtered to remove redundant sequences. The final dataset contained 15058 3’UTR sequences for 13975 genes (some genes have more than one 3’UTR isoform). The target sites of *let-7a* sequence that obey one of the following criteria were predicted using the algorithm developed in (Veksler-Lublinsky et al., 2010): (1) perfect Watson-Crick (W/C) complementarity (perfect match) to positions 2-7; (2) W/C match to positions 2-7, but allowing 1 target bulge in positions 5-7; (3) W/C match to positions 2-8, but allowing 1 GU/Mismatch (MM) in positions 5-8; (4) W/C match to positions 2-7, but allowing 1 mRNA bulge in positions 2-4; (5) W/C match to positions 2-8, but allowing 1 GU/MM in positions 2-4.

For sites that obey one of the above criteria, a flanking region of an additional 20 nt after the seed was extracted. The interaction duplex between the full site and the miRNA was then calculated using RNAduplex (Lorenz et al., 2011). The duplex was parsed to identify both the seed type and the non-seed type for each reported interaction as follows: (A) **Seed type**: (1) 2-7 full match ; (2) 2-7 + 1 target bulge on 5-7 or 2-8 with 1GU/MM 5-8; (3) 2-7 + 1 target bulge on 2-4 or 2-8 with 1 GU/MM 2-4. (B) **NonSeed type**: (1) perfect match to positions 11-13 or 12-14 or 13-16; (2) match allowing GUs to positions 11-13 or 12-14 or 13-16; (−1) none of the above.

For genes that have multiple UTR sequences in the analysis, UTRs with redundant duplexes were filtered out from the final report.

### Quantification and statistical analysis

The p-values representation is as follow: 0.05-0.01(*); 0.01-0.001(**); 0.001-0.0001(***); <0.0001(****). Error bars indicate mean ± SD. Significance tests were conducted with Prism 9 if not mentioned.

### Data and software availability

The raw and processed data of ribosome profiling, RNA-seq, and small RNA seq in this study are available in the NCBI Gene Expression Omnibus (GEO) under GEO: GSE171748 (Edgar et al., 2002). Codes for the *let-7a* 3’ target sites prediction algorithm are available at GitHub: https://github.com/IsanaVekslerLublinsky/Let7_Proj_code.git.

