## Supplementary Tables and Figures Legends for "Critical contribution of 3’ non-seed base pairing to the *in vivo* function of the evolutionarily conserved *let-7a* microRNA"

#### SUPPLEMENTAL TABLES AND DATASET:

**Table S1. Results of phenotypic assays.** (Related to **Figure 2, 3, 4, 6, 7, S2, S5**). **A.** Adult alae formation. **B.** Numbers of progeny. **C.** Adult lethality due to vulva defect. **D.** COL-19::GFP expression pattern.

**Table S2. Summary of the ribosome profiling, RNA-seq and translational efficiency (TE) of VT1367(*maIs105*) and VT3797[*maIs105; let-7(ma435,* U18A*)*].** Related to **Figure S3.** For RNA-seq and TEW, significantly changed genes (up or down) indicate |FC| > 1.5 and P.adj < 0.1 by DESeq2 (RNA-seq) or P < 0.1 by t-test (TE).

**Table S3. Summary of the ribosome profiling, RNA-seq and translational efficiency of DG3913[*lin-41(tn1541)*] and VT3878[*lin-41(tn1541ma480); let-7(ma432,* U13A*)*].** Related to **Figures 5, S7.** Significantly changed genes (up or down) indicate |FC| > 1.5 and P.adj < 0.1 by DESeq2 (ribosome profiling and RNA-seq) or P < 0.1 by t-test (TE).

**Table S4. Summary of the 3'-sup target sites identified by the target prediction algorithm.** Related to **Figures 5, S6, S7.** log2FC and P.adj indicate the corresponded values in *lin-41(tn1541ma480);let-7(ma432)* compared to *lin-41(tn1541)***.** Site configurations are categorized in (SeedType, NonSeedType) format. For SeedType, '1' indicates sites with g2-g8 perfect seed pairing; '2' indicates sites with mildly imperfect seed pairing (1 GU/mismatch at g5-g8 or 1 target nucleotide bulge at t5-t7); '3' indicates sites with severely imperfect seed pairing (1 GU/mismatch at g2-g4 or 1 target nucleotide bulge at t2-t4). For NonSeedType, '1' indicates sites with at least 3 consecutive Watson-Crick pairing to g11-g16 of *let-7a* (g13 must be pairing); '2' indicates sites with at least 3 consecutive pairing to g11-g16 of *let-7a* with GU wobble (g13 must be pairing); '-1' indicates other sites which are considered as no critical non-seed pairing. Note that our unpublished data suggest that a single GU in 3' non-seed region may not significantly affect miRNA repressing efficacy.

**Table S5. New alleles generated in this study.**

**Dataset S1. (.html) Results of the *let-7a* 3' target prediction considering both seed and critical non-seed pairing.** The description of site type and over-expression values are identical to Table S4. The analysis is based on *C. elegans* genome *WBcel235*.

### Table S5 New *C. elegans* genetic alleles

#### *let-7a*

| ALLELE | LOCATION/MUTATION | DESCRIPTION | RELATED |
|---|---|---|---|
| ma341 | chr X: 14744250 <– 14744180:<br>WT: acactgtggatccggTGAGGTAGTAGGTTGTATAGTTtggaatattaccaccggTGAACTATGCAATTTTCTACCTTACcggagacagaac<br>ma341: acactgtggatccggTGAGGTAGTATGTAATATTGTAtggaatattaccaccggTGTACAATGTTATGTTCTACCTTACcggagacagaac | *mir-84* swap | Fig.2, S2,S5 |
| ma393 | chr X: 14744250 <– 14744180:<br>WT: acactgtggatccggTGAGGTAGTAGGTTGTATAGTTtggaatattaccaccggTGAACTATGCAATTTTCTACCTTACcggagacagaac<br>ma393: acactgtggatccgg-------------TGTATCAGTTCGATATCTGAcgg------------------------------------------------------cggagacagaac | pre-*let-7* swapped by sgINPP4A jump board. Null. | Fig.2, 3, 7, S2 |
| ma453 | chr X: 14744250 <– 14744180:<br>WT: acactgtggatccggTGAGGTAGTAGGTTGTATAGTTtggaatattaccaccggTGAACTATGCAATTTTCTACCTTACcggagacagaac<br>ma453: acactgtggatccggTGAGGTAGGAGTTTGTATAGTTtggaatattaccaccggTGAACTATGCAATTTGCTACCTTACcggagacagaac | U9G | Fig.3, S2 |
| ma454 | chr X: 14744250 <– 14744180:<br>WT: acactgtggatccggTGAGGTAGTAGGTTGTATAGTTtggaatattaccaccggTGAACTATGCAATTTTCTACCTTACcggagacagaac<br>ma454: acactgtggatccggTGAGGTAGTTGGTTGTATAGTTtggaatattaccaccggTGAACTATGCAATTATCTACCTTACcggagacagaac | A10U | Fig.3, S2 |
| ma431 | chr X: 14744250 <– 14744180:<br>WT: acactgtggatccggTGAGGTAGTAGGTTGTATAGTTtggaatattaccaccggTGAACTATGCAATTTTCTACCTTACcggagacagaac<br>ma431 acactgtggatccggTGAGGTAGTATGTTGTATAGTTtggaatattaccaccggTGAACTATGCAATGTTCTACCTTACcggagacagaac | G11U | Fig.3, S2 |
| ma448 | chr X: 14744250 <– 14744180:<br>WT: acactgtggatccggTGAGGTAGTAGGTTGTATAGTTtggaatattaccaccggTGAACTATGCAATTTTCTACCTTACcggagacagaac<br>ma448: acactgtggatccggTGAGGTAGTAGTTTGTATAGTTtggaatattaccaccggTGAACTATGCAAGTTTCTACCTTACcggagacagaac | G12U | Fig.3, S2 |
| ma432 | chr X: 14744250 <– 14744180:<br>WT: acactgtggatccggTGAGGTAGTAGGTTGTATAGTTtggaatattaccaccggTGAACTATGCAATTTTCTACCTTACcggagacagaac<br>ma432 acactgtggatccggTGAGGTAGTAGGATGTATAGTTtggaatattaccaccggTGAACTATGCATTTTTCTACCTTACcggagacagaac | U13A | Fig.3,4,5 S2,S4,S6 |
| ma433 | chr X: 14744250 <– 14744180:<br>WT: acactgtggatccggTGAGGTAGTAGGTTGTATAGTTtggaatattaccaccggTGAACTATGCAATTTTCTACCTTACcggagacagaac<br>ma433 acactgtggatccggTGAGGTAGTAGGTAGTATAGTTtggaatattaccaccggTGAACTATGCTATTTTCTACCTTACcggagacagaac | U14A | Fig.3, S2 |
| ma434 | chr X: 14744250 <– 14744180:<br>WT: acactgtggatccggTGAGGTAGTAGGTTGTATAGTTtggaatattaccaccggTGAACTATGCAATTTTCTACCTTACcggagacagaac<br>ma434 acactgtggatccggTGAGGTAGTAGGTTATATAGTTtggaatattaccaccggTGAACTATGTAATTTTCTACCTTACcggagacagaac | G15A | Fig.3, S2 |
| ma449 | chr X: 14744250 <– 14744180:<br>WT: acactgtggatccggTGAGGTAGTAGGTTGTATAGTTtggaatattaccaccggTGAACTATGCAATTTTCTACCTTACcggagacagaac<br>ma449: acactgtggatccggTGAGGTAGTAGGTTGGATAGTTtggaatattaccaccggTGAACTATTCAATTTTCTACCTTACcggagacagaac | U16G | Fig.3, S2 |
| ma450 | chr X: 14744250 <– 14744180:<br>WT: acactgtggatccggTGAGGTAGTAGGTTGTATAGTTtggaatattaccaccggTGAACTATGCAATTTTCTACCTTACcggagacagaac<br>ma450: acactgtggatccggTGAGGTAGTAGGTTGTTTAGTTtggaatattaccaccggTGAACTAAGCAATTTTCTACCTTACcggagacagaac | A17U | Fig.3, S2 |
| ma435 | chr X: 14744250 <– 14744180:<br>WT: acactgtggatccggTGAGGTAGTAGGTTGTATAGTTtggaatattaccaccggTGAACTATGCAATTTTCTACCTTACcggagacagaac<br>ma435: acactgtggatccggTGAGGTAGTAGGTTGTAAAGTTtggaatattaccaccggTGAACTTTGCAATTTTCTACCTTACcggagacagaac | U18A | Fig.3, S2, S3 |
| ma436 | chr X: 14744250 <– 14744180:<br>WT: acactgtggatccggTGAGGTAGTAGGTTGTATAGTTtggaatattaccaccggTGAACTATGCAATTTTCTACCTTACcggagacagaac<br>ma436: acactgtggatccggTGAGGTAGTAGGTTGTATTGTTtggaatattaccaccggTGAACAATGCAATTTTCTACCTTACcggagacagaac | A19U | Fig.3, S2 |
| ma452 | chr X: 14744250 <– 14744180:<br>WT: acactgtggatccggTGAGGTAGTAGGTTGTATAGTTtggaatattaccaccggTGAACTATGCAATTTTCTACCTTACcggagacagaac<br>ma452: acactgtggatccggTGAGGTAGTAGGTTGTATACTTtggaatattaccaccggTGAAGTATGCAATTTTCTACCTTACcggagacagaac | G20C | Fig.3, S2 |
| ma456 | chr X: 14744250 <– 14744180:<br>WT: acactgtggatccggTGAGGTAGTAGGTTGTATAGTTtggaatattaccaccggTGAACTATGCAATTTTCTACCTTACcggagacagaac<br>ma456: acactgtggatccggTGAGGTAGTAGGTTGTATAGATtggaatattaccaccggTGATCTATGCAATTTTCTACCTTACcggagacagaac | U21A | Fig.3, S2 |
| ma437 | chr X: 14744250 <– 14744180:<br>WT: acactgtggatccggTGAGGTAGTAGGTTGTATAGTTtggaatattaccaccggTGAACTATGCAATTTTCTACCTTACcggagacagaac<br>ma437 acactgtggatccggTGAGGTAGTAGGTTGTATAGTAtggaatattaccaccggTGTACTATGCAATTTTCTACCTTACcggagacagaac | U22A | Fig.3, S2 |
| ma479 | chr X: 14744250 <– 14744180:<br>WT: acactgtggatccggTGAGGTAGTAGGTTGTATAGTTtggaatattaccaccggTGAACTATGCAATTTTCTACCTTACcggagacagaac<br>ma479: acactgtggatccggTGAGGTAGTAGGTTGGAAAGTTtggaatattaccaccggTGAACTTTTCAATTTTCTACCTTACcggagacagaac | U16G + U18A, equal to *ma449ma435* | Fig.3, S2 |
| ma428 | chr X: 14744250 <– 14744180:<br>WT: acactgtggatccggTGAGGTAGTAGGTTGTATAGTTtggaatattaccaccggTGAACTATGCAATTTTCTACCTTACcggagacagaac<br>ma428: : acactgtggatccggTGAGGTAGTAGGATGTAAAGTTtggaatattaccaccggTGAACTTTGCATTTTTCTACCTTACcggagacagaac | U13A + U18A equal to *ma432ma435* | Fig.S2 |
| ma455 | chr X: 14744250 <– 14744180:<br>WT: acactgtggatccggTGAGGTAGTAGGTTGTATAGTTtggaatattaccaccggTGAACTATGCAATTTTCTACCTTACcggagacagaac<br>ma455: acactgtggatccggTGAGGTAGTAGTTTGTAGAGTTtggaatattaccaccggTGAACTCTGCAATTTTCTACCTTACcggagacagaac | U18G | Fig.S2 |
| ma451 | chr X: 14744250 <– 14744180:<br>WT: acactgtggatccggTGAGGTAGTAGGTTGTATAGTTtggaatattaccaccggTGAACTATGCAATTTTCTACCTTACcggagacagaac<br>ma451: acactgtggatccggTGAGGTAGTAGGTTGTTTAGTTtggaatattaccaccggTGAACTAAGCAATTTTCTACCTTACcggagacagaac | U18C | Fig.S2 |
| ma476 | chr X: 14744250 <– 14744180:<br>WT: acactgtggatccggTGAGGTAGTAGGTTGTATAGTTtggaatattaccaccggTGAACTATGCAATTTTCTACCTTACcggagacagaac<br>ma476: acactgtggatccggTGAGGTAGTAGGTTGTTATCAAtggaatattaccaccggTGTTGATAGCAATTTTCTACCTTACcggagacagaac | A17U + U18A + A19U + G20C + U21A + U22A | Fig.S2 |
| ma477 | chr X: 14744250 <– 14744180:<br>WT: acactgtggatccggTGAGGTAGTAGGTTGTATAGTTtggaatattaccaccggTGAACTATGCAATTTTCTACCTTACcggagacagaac<br>ma477: acactgtggatccggTGAGGTAGTAGGTTGTTATGTTtggaatattaccaccggTGAACATAGCAATTTTCTACCTTACcggagacagaac | A17U + U18A + A19U | Fig.S2 |
| ma478 | chr X: 14744250 <– 14744180:<br>WT: acactgtggatccggTGAGGTAGTAGGTTGTATAGTTtggaatattaccaccggTGAACTATGCAATTTTCTACCTTACcggagacagaac<br>ma478: acactgtggatccggTGAGGTAGTAGGTTGTATACAAtggaatattaccaccggTGTTGTATGCAATTTTCTACCTTACcggagacagaac | G20C + U21A + U22A | Fig.S2 |

*lin-41*

| ALLELE | LOCATION/MUTATION | DESCRIPTION | RELATED |
|---|---|---|---|
| ma480 | chr I. 9335330 <- 9335425<br>WT: caagtatacc**ttttatacaaccgttctacactca**acgcgatgtaaatatcgcaatccct**ttttatacaaccattctgcctc**tgaaccattgaaacc<br>ma480: caagtatacc**ttttatacatccgttctacactca**acgcgatgtaaatatcgcaatccct**ttttatacatccattctgcctc**tgaaccattgaaacc | LCS1 + LCS2 t13 mismatch to *let-7a*, complementary to *ma432* | Fig. 4,5,6, S4,S6 |
| ma378 | chr I. 9335330 <- 9335425<br>WT: caagtatacc**ttttatacaaccgttctacactca**acgcgatgtaaatatcgcaatccct**ttttatacaaccattctgcctc**tgaaccattgaaacc<br>ma378: caagtatacc**attaatattacagttctacactca**acgcgatgtaaatatcgcaatccct**attaatattaacattctgcctc**tgaaccattgaaacc | LCS1 + LCS2 match *ma341* | Fig. S5 |
| ma501 | chr I. 9335330 <- 9335425<br>WT: caagtatacc**ttttatacaaccgttctacactca**acgcgatgtaaatatcgcaatccct**ttttatacaaccattctgcctc**tgaaccattgaaacc<br>ma501: caagtatacc**ttttatacaaccgttctac-ctca**acgcgatgtaaatatcgcaatccct**ttttatacaaccattctacctc**tgaaccattgaaacc | LCS1 + LCS2 perfect seed pairing to *let-7a* * | Fig. 6,7 |
| ma545 | chr I. 9335330 <- 9335425<br>WT: caagtatacc**ttttatacaaccgttctacactca**acgcgatgtaaatatcgcaatccct**ttttatacaaccattctgcctc**tgaaccattgaaacc<br>ma545: caagtatacc**ttttatacatgggttctacactca**acgcgatgtaaatatcgcaatccct**ttttatacatggattctgcctc**tgaaccattgaaacc | LCS1 + LCS2 t11-t13 mismatch to *let-7a* | Fig. 6 |
| ma555 | chr I. 9335330 <- 9335425<br>wt: caagtatacc**ttttatacaaccgttctacactca**acgcgatgtaaatatcgcaatccct**ttttatacaaccattctgcctc**tgaaccattgaaacc<br>ma555: caagtatacc**ttttatacaacggttctacactca**acgcgatgtaaatatcgcaatccct**ttttatacaacgattctgcctc**tgaaccattgaaacc | LCS1 + LCS2 t11 mismatch to *let-7a* | Fig. 6 |
| ma556 | chr I. 9335330 <- 9335425<br>wt: caagtatacc**ttttatacaaccgttctacactca**acgcgatgtaaatatcgcaatccct**ttttatacaaccattctgcctc**tgaaccattgaaacc<br>ma556: caagtatacc**ttttatacaagcgttctacactca**acgcgatgtaaatatcgcaatccct**ttttatacaagcattctgcctc**tgaaccattgaaacc | LCS1 + LCS2 t12 mismatch to *let-7a* | Fig. 6 |
| ma564 | chr I. 9335330 <- 9335425<br>wt: caagtatacc**ttttatacaaccgttctacactca**acgcgatgtaaatatcgcaatccct**ttttatacaaccattctgcctc**tgaaccattgaaacc<br>ma564: caagtatacc**ttttatactaccgttctacactca**acgcgatgtaaatatcgcaatccct**ttttatac taccattctgcctc**tgaaccattgaaacc | LCS1 + LCS2 t14 mismatch to *let-7a* | Fig. 6 |
| ma565 | chr I. 9335330 <- 9335425<br>wt: caagtatacc**ttttatacaaccgttctacactca**acgcgatgtaaatatcgcaatccct**ttttatacaaccattctgcctc**tgaaccattgaaacc<br>ma565: caagtatacc**ttttataaaaccgttctacactca**acgcgatgtaaatatcgcaatccct**ttttataaaaccattctgcctc**tgaaccattgaaacc | LCS1 + LCS2 t15 mismatch to *let-7a* | Fig. 6 |
| ma566 | chr I. 9335330 <- 9335425<br>wt: caagtatacc**ttttatacaaccgttctacactca**acgcgatgtaaatatcgcaatccct**ttttatacaaccattctgcctc**tgaaccattgaaacc<br>ma566: caagtatacc**ttttattcaaccgttctacactca**acgcgatgtaaatatcgcaatccct**ttttatacaaccattctgcctc**tgaaccattgaaacc | LCS1 + LCS2 t16 mismatch to *let-7a* | Fig. 6 |
| ma571 | chr I. 9335330 <- 9335425<br>wt: caagtatacc**ttttatacaaccgttctacactca**acgcgatgtaaatatcgcaatccct**ttttatacaaccattctgcctc**tgaaccattgaaacc<br>ma571: caagtatacc**ttttatacaagggttctacactcaacg**cgatgtaaatatcgcaatccct**ttttatacaaggattctgcctc**tgaaccattgaaacc | LCS1 + LCS2 t11-t12 mismatch to *let-7a* | Fig. 6 |
| ma572 | chr I. 9335330 <- 9335425<br>wt: caagtatacc**ttttatacaaccgttctacactca**acgcgatgtaaatatcgcaatccct**ttttatacaaccattctgcctc**tgaaccattgaaacc<br>ma572: caagtatacc**ttttattgttccgttctacactca**acgcgatgtaaatatcgcaatccct**ttttattgttccattctgcctc**tgaaccattgaaacc | LCS1 + LCS2 t13-t16 mismatch to *let-7a* | Fig. 6 |
| ma504 | chr I. 9335330 <- 9335425<br>WT: caagtatacc**ttttatacaaccgttctacactca**acgcgatgtaaatatscgcaatccct**ttttatacaaccattctgcctc**tgaaccattgaaacc<br>ma504: caagtatacc**ttttatacatccgttctac-ctca**acgcgatgtaaatatcgcaatccct**ttttatacatccattctacctc**tgaaccattgaaacc | Identical to *ma501ma480* | Fig. 6 |
| ma552 | chr I. 9335330 <- 9335425<br>wt: caagtatacc**ttttatacaaccgttctacactca**acgcgatgtaaatatcgcaatccct**ttttatacaaccattctgcctc**tgaaccattgaaacc<br>ma552: caagtatacc**ttttatacaacggttctac-ctca**acgcgatgtaaatatcgcaatccct**ttttatacaacgattctacctc**tgaaccattgaaacc | Identical to *ma501ma555* | Fig. 6 |
| ma553 | chr I. 9335330 <- 9335425<br>wt: caagtatacc**ttttatacaaccgttctacactca**acgcgatgtaaatatcgcaatccct**ttttatacaaccattctgcctc**tgaaccattgaaacc<br>ma553: caagtatacc**ttttatacaagcgttctac-ctca**acgcgatgtaaatatcgcaatccct**ttttatacaagcattctacctc**tgaaccattgaaacc | Identical to *ma501ma456* | Fig. 6 |
| ma511 | chr I. 9335330 <- 9335425<br>WT: caagtatacc**ttttatacaaccgttctacactca**acgcgatgtaaatatcgcaatccct**ttttatacaaccattctgcctc**tgaaccattgaaacc<br>ma511: caagtatacc**ttttatacatgggttctac-ctca**acgcgatgtaaatatcgcaatccct**ttttatacatggattctacctc**tgaaccattgaaacc | Identical to *ma501ma545* | Fig. 6 |
| ma554 | chr I. 9335330 <- 9335425<br>wt: caagtatacc**ttttatacaaccgttctacactca**acgcgatgtaaatatcgcaatccct**ttttatacaaccattctgcctc**tgaaccattgaaacc<br>ma554: caagtatacc**ttttatacaagggttctac - ctca**acgcgatgtaaatatcgcaatccct**ttttatacaaggattctacctc**tgaaccattgaaacc | Identical to *ma501ma571* | Fig. 6 |
| ma525 | chr I. 9335330 <- 9335425<br>wt: caagtatacc**ttttatacaaccgttctacactca**acgcgatgtaaatatcgcaatccct**ttttatacaaccattctgcctc**tgaaccattgaaacc<br>ma525: caagtatacc**ttttattgttccgttctac-ctca**acgcgatgtaaatatcgcaatccct**ttttattgttccattctacctc**tgaaccattgaaacc | Identical to *ma501ma572* | Fig. 6 |

*This allele(*ma501)* is identical to *xe11* (Brancati and Grosshans, 2018), but was generated independently in this study.

*daf-12*

| ALLELE | LOCATION/MUTATION | DESCRIPTION | RELATED |
|---|---|---|---|
| ma567 | chr X.<br>ma567:<br>aaatcatctaccaaacgatgccatgccttc<CTCTTCCTCTTAATTCCTCTCTTAATT >cctacctcttaattccgtcaatactatctcagattttcattgaagaact | daf-12 Jump board | Fig. 5, S7 |
| ma568 | chr X.<br>ma568:<br>ACCTACTAGAAATCATCTATTTATGGTGGTGAATACCTCACATCTTGATTCTATATTGCCTCCATCCAACAAACTCAATCTAGCCAC<br>ATTTCTTCTTTTTCACGTACCTCAACCACCCTTTCCATATTTATGGTGGTGATCTACCTCTTTAACCAATTCATCATCTTTTTATATTG<br>TTTCTTATTTGCATTCAACTTGGAAATAGCCACTATCATATCACTATTGCGTATTTCCTTTCATTTTCTTGTCTTATTTTCTTGAGACC<br>AGCACCAGAAGATTTTTTCGATGGAGAACTAAGCATGATAATTTGAAGTTTTCCATTTAAAAAATGCAGGTAATACGGTTAATTC<br>AATCTGCGAGTTGATGTTCGGTCTCCGGTTTTCATGTTCTACTTCTAATGACTAGAAACCCTTTTATCTAACATCCGGTCCTCCTATC<br>CCTAATGTACCCAGTAGATATTTTTTCCCCGAATGATTAAACCTCCCAGTCAAATATTGTATTTTGATTTATGGTGGTGACCTACCT<br>CTTAATTCCGTCAATACTATCTC | daf-12 LCS1-3 with t11-t13 mismatch to *let-7a* | Fig. 5, S7 |

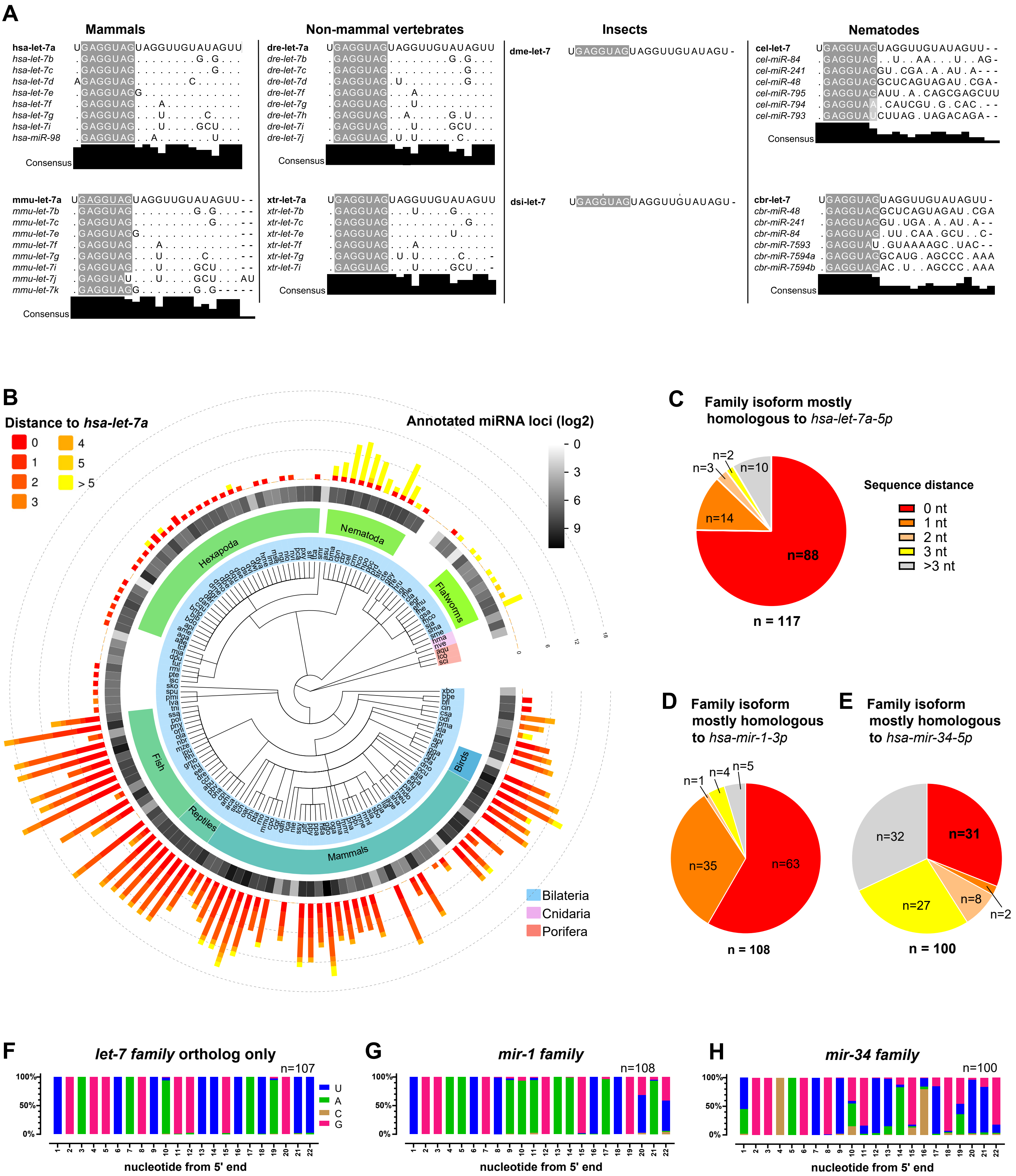
A
Mammals
hsa-let-7a UGAGGUAGUAGGUUGUAUAGUU
hsa-let-7b .GAGGUAG........G.G...
hsa-let-7c .GAGGUAG..........G...
hsa-let-7d AGAGGUAG.......C......
hsa-let-7e .GAGGUAGG.............
hsa-let-7f .GAGGUAG...A..........
hsa-let-7g .GAGGUAG...U.....C....
hsa-let-7i .GAGGUAG...U....GCU...
hsa-miR-98 .GAGGUAG..A.......U...
Consensus
mmu-let-7a UGAGGUAGUAGGUUGUAUAGUU--
mmu-let-7b .GAGGUAG........G.G....--
mmu-let-7c .GAGGUAG..........G....--
mmu-let-7e .GAGGUAGG.............--
mmu-let-7f .GAGGUAG...A...........--
mmu-let-7g .GAGGUAG...U.....C.....--
mmu-let-7i .GAGGUAG...U....GCU....--
mmu-let-7j .GAGGUAU...U....GCU...AU
mmu-let-7k .GAGGUAGG.......G.G-----
Consensus
Non-mammal vertebrates
dre-let-7a UGAGGUAGUAGGUUGUAUAGUU
dre-let-7b .GAGGUAG........G.G...
dre-let-7c .GAGGUAG..........G...
dre-let-7d .GAGGUAG.U........G...
dre-let-7f .GAGGUAG....A.........
dre-let-7g .GAGGUAG...U..........
dre-let-7h .GAGGUAG..A.....G.U...
dre-let-7i .GAGGUAG...U....GCU...
dre-let-7j .GAGGUAG.U.U.....C....
Consensus
xtr-let-7a UGAGGUAGUAGGUUGUAUAGUU
xtr-let-7b .GAGGUAG...U....G.....
xtr-let-7c .GAGGUAG..........G...
xtr-let-7e .GAGGUAG.........U....
xtr-let-7f .GAGGUAG....A.........
xtr-let-7g .GAGGUAG.U.U.....C....-
xtr-let-7i .GAGGUAG...U....GCU..-
Consensus
Insects
dme-let-7 UGAGGUAGUAGGUUGUAUAGU-
dsi-let-7 UGAGGUAGUAGGUUGUAUAGU-
Nematodes
cel-let-7 UGAGGUAGUAGGUUGUAUAGUU--
cel-miR-84 .GAGGUAG..U..AA...U..AG-
cel-miR-241 .GAGGUAGGU.CGA.A.AU.A---
cel-miR-48 .GAGGUAGGCUCAGUAGAU.CGA-
cel-miR-795 .GAGGUAGAUU.A.CAGCGAGCUU
cel-miR-794 .GAGGUAA.CAUCGU.G.CAC.--
cel-miR-793 .GAGGUAUCUUAG.UAGACAGA--
Consensus
cbr-let-7 UGAGGUAGUAGGUUGUAUAGUU-
cbr-miR-48 .GAGGUAGGCUCAGUAGAU.CGA
cbr-miR-241 .GAGGUAGGU.UGA.A.AU.A--
cbr-miR-84 .GAGGUAG.UU.CAA.GCU..C-
cbr-miR-7593 .GAGGUAU.GUAAAAGC.UAC--
cbr-miR-7594a .GAGGUAGGCAUG.AGCCC.AAA
cbr-miR-7594b .GAGGUAGAC.U..AGCCC.AAA
Consensus
B
Distance to hsa-let-7a
0 1 2 3 4 5 > 5
Annotated miRNA loci (log2)
0 3 6 9
Hexapoda
Nematoda
Flatworms
Fish
Reptiles
Mammals
Birds
Bilateria
Cnidaria
Porifera
C
Family isoform mostly homologous to hsa-let-7a-5p
n=2 n=3 n=10 n=14 n=88
n = 117
Sequence distance
0 nt
1 nt
2 nt
3 nt
>3 nt
D
Family isoform mostly homologous to hsa-mir-1-3p
n=1 n=4 n=5 n=35 n=63
n = 108
E
Family isoform mostly homologous to hsa-mir-34-5p
n=32 n=31 n=27 n=8 n=2
n = 100
F
let-7 family ortholog only
n=107
100% 50% 0%
U A C G
nucleotide from 5' end
G
mir-1 family
n=108
nucleotide from 5' end
H
mir-34 family
n=100
nucleotide from 5' end

**Figure S1.** ***let-7a*** **is distinguishably conserved among bilaterian species.** (Related to **Figure 1**).

**A.** Sequence alignments and consensus of *let-7 family* miRNAs from model organisms including *H. sapiens (hsa)*, *M. musculus (mmu), D. rerio (dre), X. tropicalis (xtr), D. melanogaster (dme), D. simulans (dsi), C. elegans (cel) and C. briggsae (cbr)*. Nucleotides identical to *hsa-let-7a-5p* are indicated by dots (.) and gaps in sequence alignment are indicated by dashes (-). *dme* and *dsi* do not have consensus scores because *let-7a* appears singly in these species. Alignments and consensus are scored by JalView (Waterhouse et al., 2009).

**B.** Summary of *let-7 family* miRNAs across animal phylogeny tree which includes all metazoans with *let-7 family* annotation in miRBase v22.1**.** In the outer circle, bar lengths indicate the number of annotated *let-7 family* isoforms, and bar colors indicate the sequence distances to *hsa-let-7a-5p*. The middle circle indicates the quality of the total miRNA annotation of each species. Note that cases of an apparent absence of *let-7 family* miRNA usually correlate with poor miRNA annotation.

**C-E.** Distributions of sequence distances between *hsa-let-7a-5p* (**C**), *hsa-mir-1-3p* (**D**), *hsa-mir-34-5p* (**E**) and their closest isoforms of each species across bilaterians.

**F.** Nucleotide frequency of the *let-7a family* isoforms closest to *hsa-let-7a-5p*, excluding species without the *let-7a* orthologs (closest *let-7* isoform has > 3 nucleotides different from *hsa-let-7a-5p)*.

**G-H.** Nucleotide frequency of *mir-1 family* **(G)** and *mir-34 family* **(H)** isoforms mostly homologous to *hsa-mir-1-3p* **(G)** and *has-mir-34-5p* **(H)**.

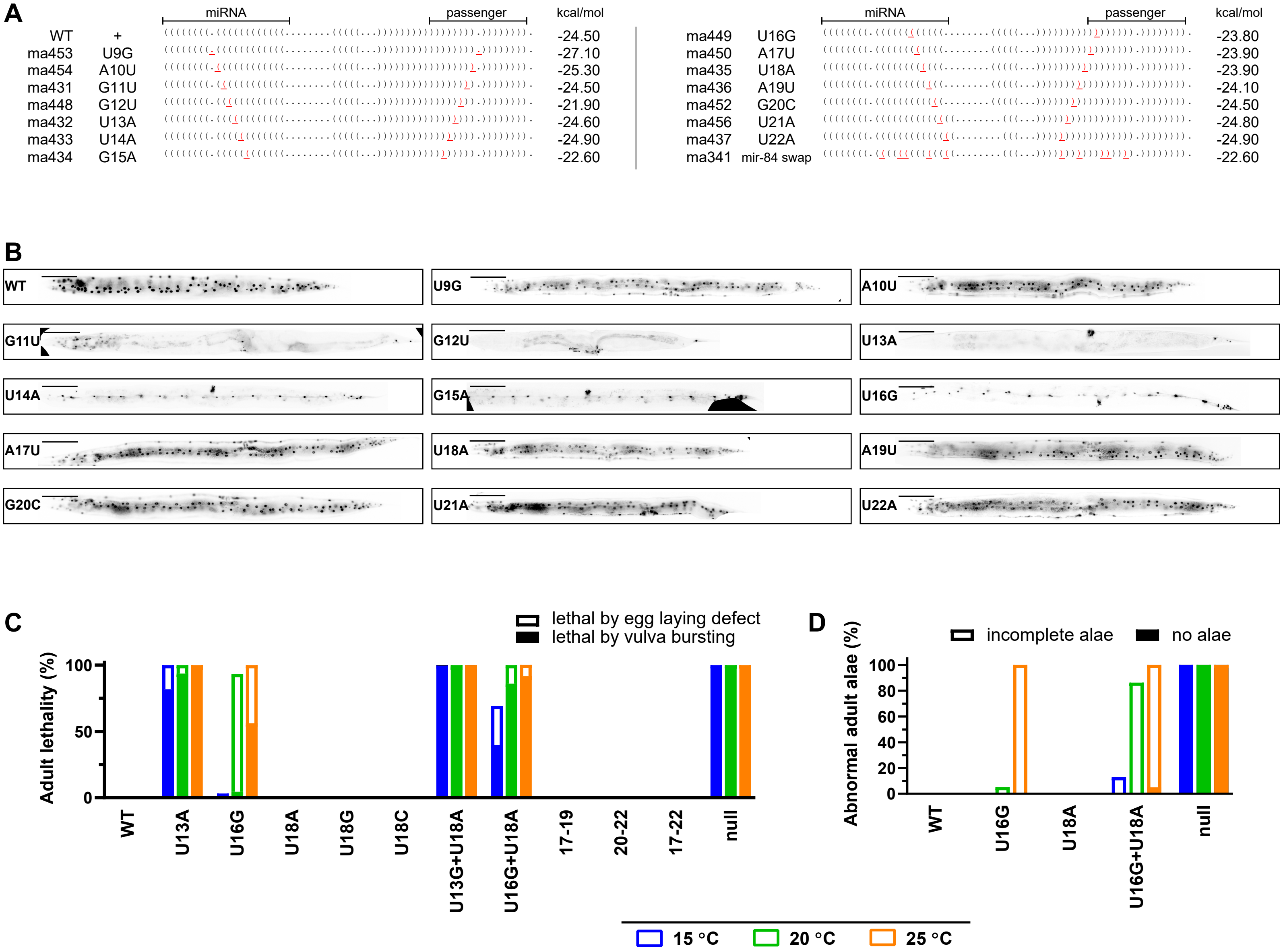
A
miRNA
passenger
kcal/mol
WT + -24.50
ma453 U9G -27.10
ma454 A10U -25.30
ma431 G11U -24.50
ma448 G12U -21.90
ma432 U13A -24.60
ma433 U14A -24.90
ma434 G15A -22.60
ma449 U16G -23.80
ma450 A17U -23.90
ma435 U18A -23.90
ma436 A19U -24.10
ma452 G20C -24.50
ma456 U21A -24.80
ma437 U22A -24.90
ma341 mir-84 swap -22.60
B
WT
U9G
A10U
G11U
G12U
U13A
U14A
G15A
U16G
A17U
U18A
A19U
G20C
U21A
U22A
C
lethal by egg laying defect
lethal by vulva bursting
Adult lethality (%)
100
50
0
WT
U13A
U16G
U18A
U18G
U18C
U13G+U18A
U16G+U18A
17-19
20-22
17-22
null
D
incomplete alae
no alae
Abnormal adult alae (%)
100
80
60
40
20
0
WT
U16G
U18A
U16G+U18A
null
15 °C
20 °C
25 °C

**Figure S2. A phenotypic mutational screen of *let-7a* 3' non-seed region.** (Related to **Figure 3**).

**A.** Dot-bracket notations of optimal secondary structures and the minimum folding free energy of pre-miRNA for *let-7a* mutants (**Fig. 3A**). Secondary structure predictions and free energy were analyzed by RNAfold (Denman, 1993; Kerpedjiev et al., 2015). For each position, the miRNA nucleotide and the corresponding nucleotide in the passenger strand were mutated according to the following criteria (ranked by the order): (1) The resulting nucleotides are identical to the corresponding nucleotides of *let-7a(ma341)*; (2) the resulting sequences cause the minimum change to pre-miRNA structure; (3) the resulting sequences cause the minimum change to pre-miRNA folding energy.

**B**. Representative COL-19::GFP patterns for the *let-7a* 3' non-seed mutants, scored by expression of *maIs105(col-19::gfp)*. Scale bars, 100 µm. Images are processed by ImageJ Fiji (Schindelin et al., 2012).

**C-D.** Synergistic effect between mutations at g17-g22 and the critical non-seed nucleotides at g11-g16. **C.** Vulva integrity phenotypes, reflected by two categories of lethality: bursting of pre-gravid adults (severe), or accumulation of hatched progeny inside the uterus due to egg-laying defects (mild). **D.** Phenotypes in heterochronic pathway based on adult alae phenotypes. The alae phenotypes are categorized as no alae (severe) or incomplete alae (mild). Simultaneously mutating g13 and g18 did not result in enhanced phenotypes compared to mutating g13 alone, likely because the phenotypes of the g13 mutation are already essentially as strong as *let-7(null)*.

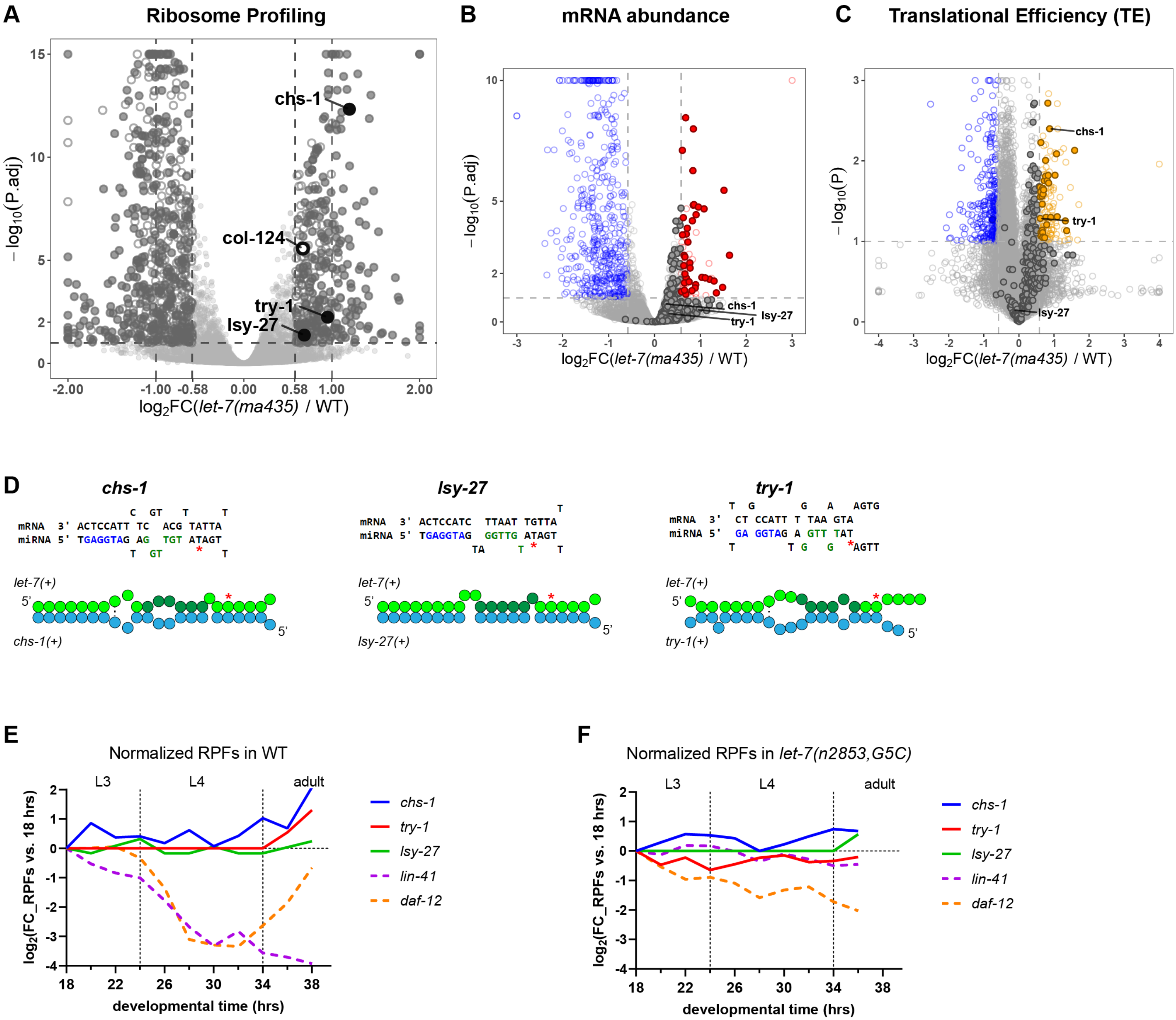
A
Ribosome Profiling
chs-1
col-124
try-1
lsy-27
$-\log_{10}$(P.adj)
$\log_2$FC(let-7(ma435) / WT)
B
mRNA abundance
C
Translational Efficiency (TE)
$-\log_{10}$(P)
D
chs-1
lsy-27
try-1
let-7(+)
chs-1(+)
lsy-27(+)
try-1(+)
E
Normalized RPFs in WT
L3
L4
adult
$\log_2$(FC_RPFs vs. 18 hrs)
developmental time (hrs)
chs-1
try-1
lsy-27
lin-41
daf-12
F
Normalized RPFs in let-7(n2853,G5C)

**Figure S3. Molecular phenotypes of *let-7(ma435, U18A).*** (Related to **Figure 3**).

**A-C**. Differential expression analyses of RPFs **(A)**, mRNA abundance **(B)** and TE **(C)** between *let-7(ma435);maIs105* and *maIs105*. Hollow points indicate developmentally dynamic genes whose perturbation in these experiments could potentially reflect imperfect synchrony between samples (See Figure 3 legend and Star Methods) and not the genotypes; solid points indicate genes likely to be perturbed specifically due to the mutation in **A**. Genes with significantly increased/decreased mRNA (FC > |1.5| and P.adj < 0.1 by DESeq2) are represented by red/blue points in **B**. Genes with significantly increased/decreased TE (FC > |1.5| and P < 0.1 by t-test) are colored orange/blue in **C**. Genes that contain predicted LCSs with g18 pairing are labeled by gene names in **A**. Solid points in **B-C** represent genes with significantly increased RPFs in ribosome profiling.

**D.** Predicted base-pairing configuration between *let-7a* and the predicted LCSs with g18 pairing. g18 are labeled by red asterisks (*).

**E-F.** Developmental profiles of the translation of the putative g18 targets in WT (**E**) and *let-7(n2853, G5C)* (**F**) at L3, L4 and young adult stages. Results were analyzed from (Aeschimann et al., 2017) and normalized by the RPFs at 18 hrs. *lin-41* and *daf-12* are used as validated *let-7a* targets which are down-regulated at L4 stages.

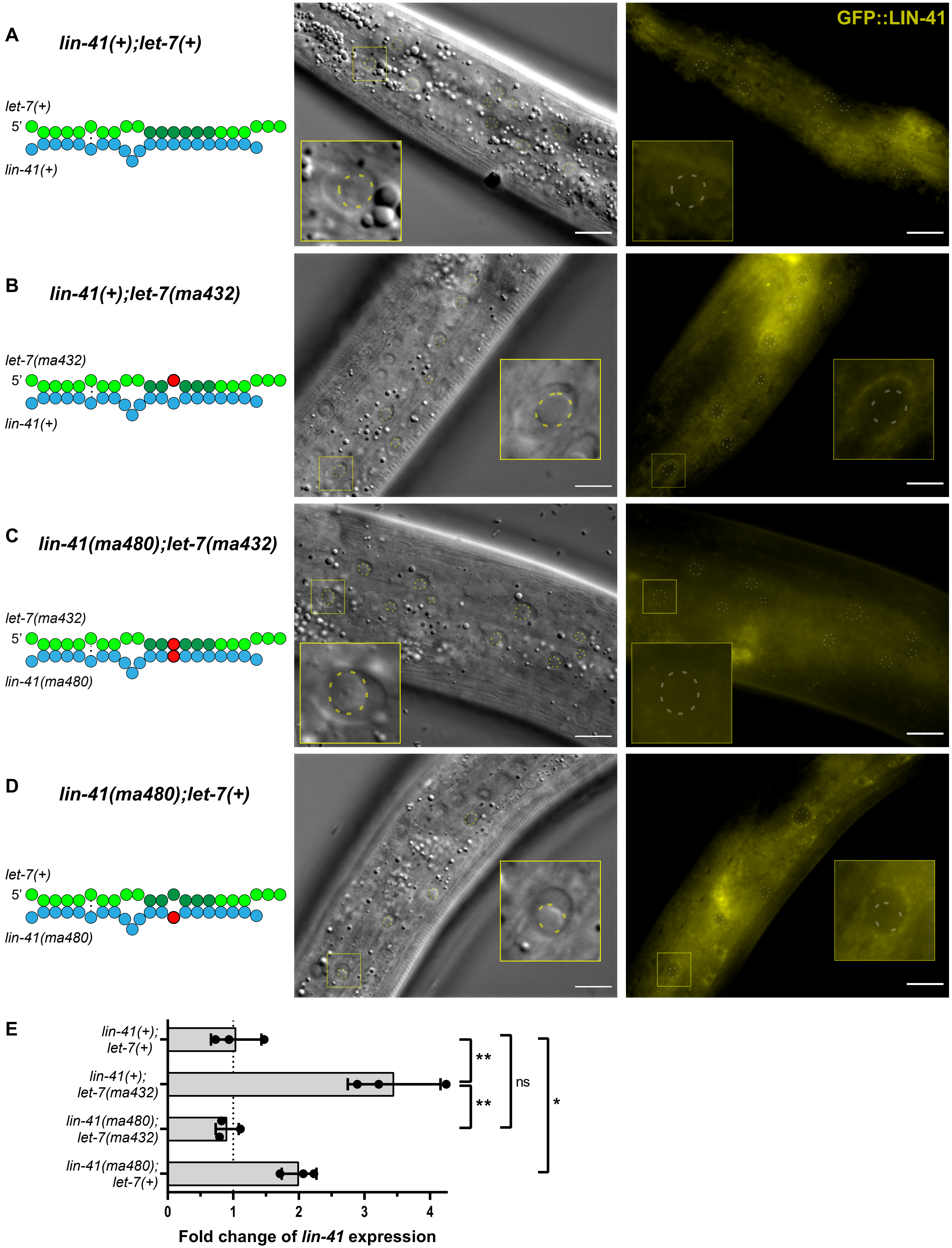

A
lin-41(+);let-7(+)
let-7(+)
5'
lin-41(+)
GFP::LIN-41
B
lin-41(+);let-7(ma432)
let-7(ma432)
5'
lin-41(+)
C
lin-41(ma480);let-7(ma432)
let-7(ma432)
5'
lin-41(ma480)
D
lin-41(ma480);let-7(+)
let-7(+)
5'
lin-41(ma480)
E
lin-41(+); let-7(+)
lin-41(+); let-7(ma432)
lin-41(ma480); let-7(ma432)
lin-41(ma480); let-7(+)
**
**
ns
*
0
1
2
3
4
Fold change of lin-41 expression

**Figure S4. Expression of LIN-41::GFP indicates that *lin-41* is a major target of *let-7a* that requires non-seed pairing.** (Related to **Figure 4**).

LIN-41 expression is visualized based on endogenous GFP-tag (*tn1541)* (Spike et al., 2014). Yellow dashed circles indicate nucleoli of Hyp7 and seam cells, which are identified using the DIC channel (middle panels) and positioned on the GFP channel image (right panels) in register with the DIC channel. The cartoons at left illustrate expected interacting configurations between *let-7a* and *lin-41* LCS1. Note that both LCSs in *lin-41* were modified using the strategy.

**A.** In WT, endogenously GFP tagged LIN-41 is expressed in neither Hyp7 nor in seam cells at the late L4 stage due to repression by *let-7a.*

**B.** In *let-7(ma432,g13*), LIN-41 is expressed abnormally in the peri-nuclear region of both Hyp7 and seam cells due to the loss of *let-7a* repression.

**C.** In *lin-41(ma480,t13);let-7(ma432,g13)*, restoring the interacting configurations between *let-7a* and *lin-41* rescues the *let-7a* repression, thus no abnormal expression of LIN-41 was detected.

**D.** In *lin-41(ma480,t13)*, disruption of non-seed paring to *let-7a* by mutations on the target causes de-repression of *lin-41*, resulting in abnormal expression of LIN-41::GFP similar to (**B**).

All strains are cultured at 25 °C. Scale bars, 10 µm. All GFP images are taken with identical exposure times and microscopy settings.

**E.** Normalized expression of *lin-41* at L4 stage measured by qRT-PCR. The transcription levels were calculated by the ΔΔCt relatively to *gpd-1* and were normalized by to *lin-41(+);let-7(+).* All strains are cultured at 25 °C. Statistical significance indicates t-test from 3 biological replicas.

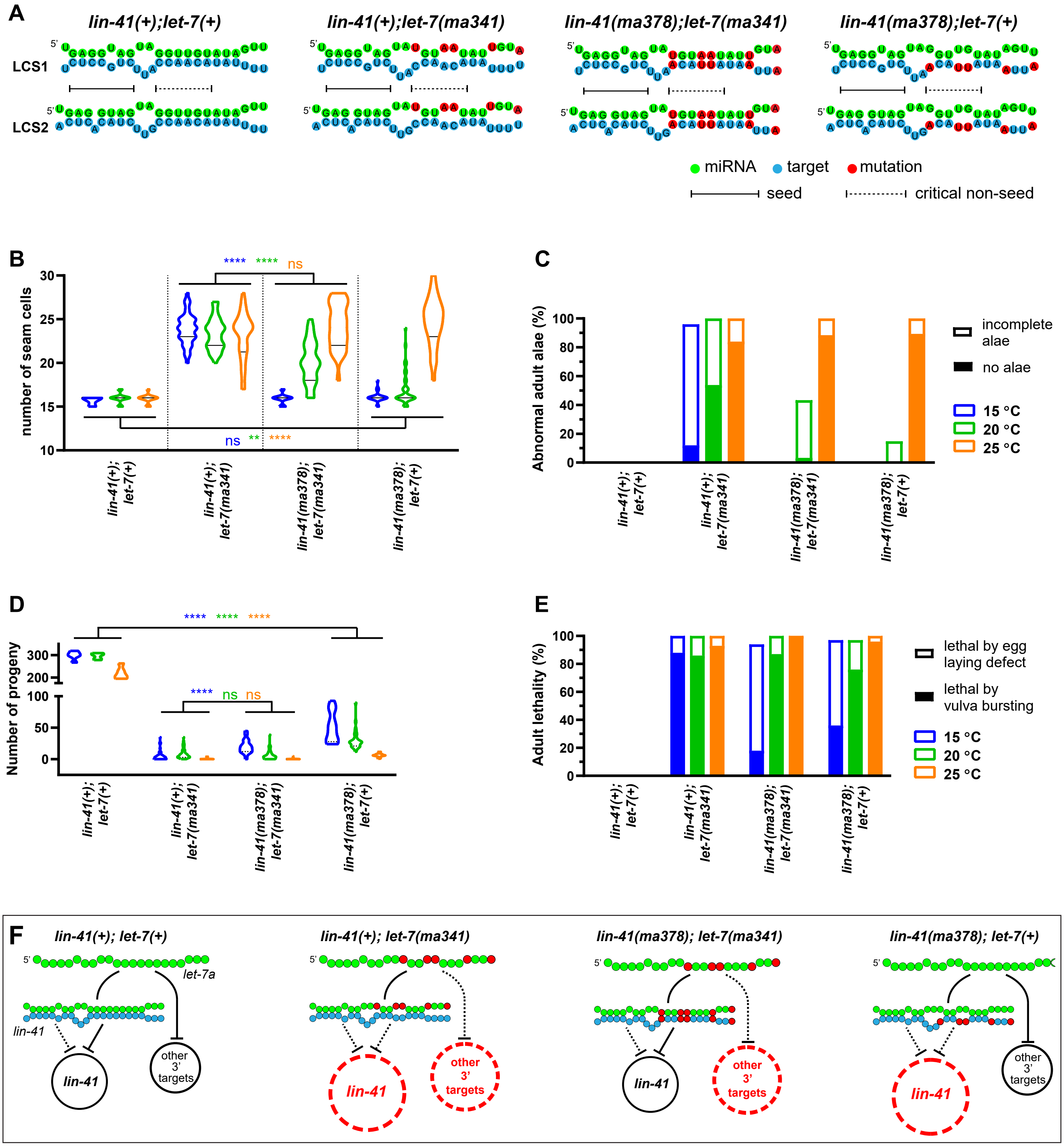
A
lin-41(+);let-7(+)
lin-41(+);let-7(ma341)
lin-41(ma378);let-7(ma341)
lin-41(ma378);let-7(+)
LCS1
LCS2
miRNA
target
mutation
seed
critical non-seed
B
number of seam cells
**** **** ns
ns ** ****
lin-41(+); let-7(+)
lin-41(+); let-7(ma341)
lin-41(ma378); let-7(ma341)
lin-41(ma378); let-7(+)
C
Abnormal adult alae (%)
incomplete alae
no alae
15 °C
20 °C
25 °C
D
Number of progeny
**** **** ****
**** ns ns
E
Adult lethality (%)
lethal by egg laying defect
lethal by vulva bursting
15 °C
20 °C
25 °C
F
lin-41(+); let-7(+)
let-7a
lin-41
other 3' targets
lin-41(+); let-7(ma341)
lin-41(ma378); let-7(ma341)
lin-41(ma378); let-7(+)

**Figure S5. *let-7a* specifically targets *lin-41* and additional 3'-sup. targets compared to the family paralogs.** (Related to **Figures 2, 4**).

**A.** Predicted pairing configurations between *let-7a* and *lin-41* LCS1/2 of WT, *let-7(ma341, mir-84 swap), lin-41(ma378);let-7(ma341)* and *lin-41(ma378)*.

**B-C.** The increase in seam cell numbers of young adult animals(**B**) and the lack of normal adult alae (**C**) are characteristic of *let-7a lf* phenotypes in the heterochronic pathway. Seam cell numbers of young adults were scored using *maIs105 [col-19::gfp].*

**C.** Proportions of animals with abnormal adult alae, categorized as no alae (severe) or incomplete alae (mild).

**D-E.** Penetrance of vulva integrity defects by numbers of progeny (**D**) and adult lethality (**E**)**.** The adult lethality is categorized as vulva bursting (severe) or egg-laying defective (mild).

**F.** Illustrative models propose that the *let-7a* 3'-sup targets additional to *lin-41* are de-repressed in *let-7a(ma341)*. The results indicate that the *lf* phenotypes of *let-7(ma341, miR-84 swap)* result from the de-repression of *lin-41* as a major effector, as well as other putative 3'-sup targets. Thus, *let-7a* confers its functional specificity among family paralogs by regulating a multiplicity of target genes.

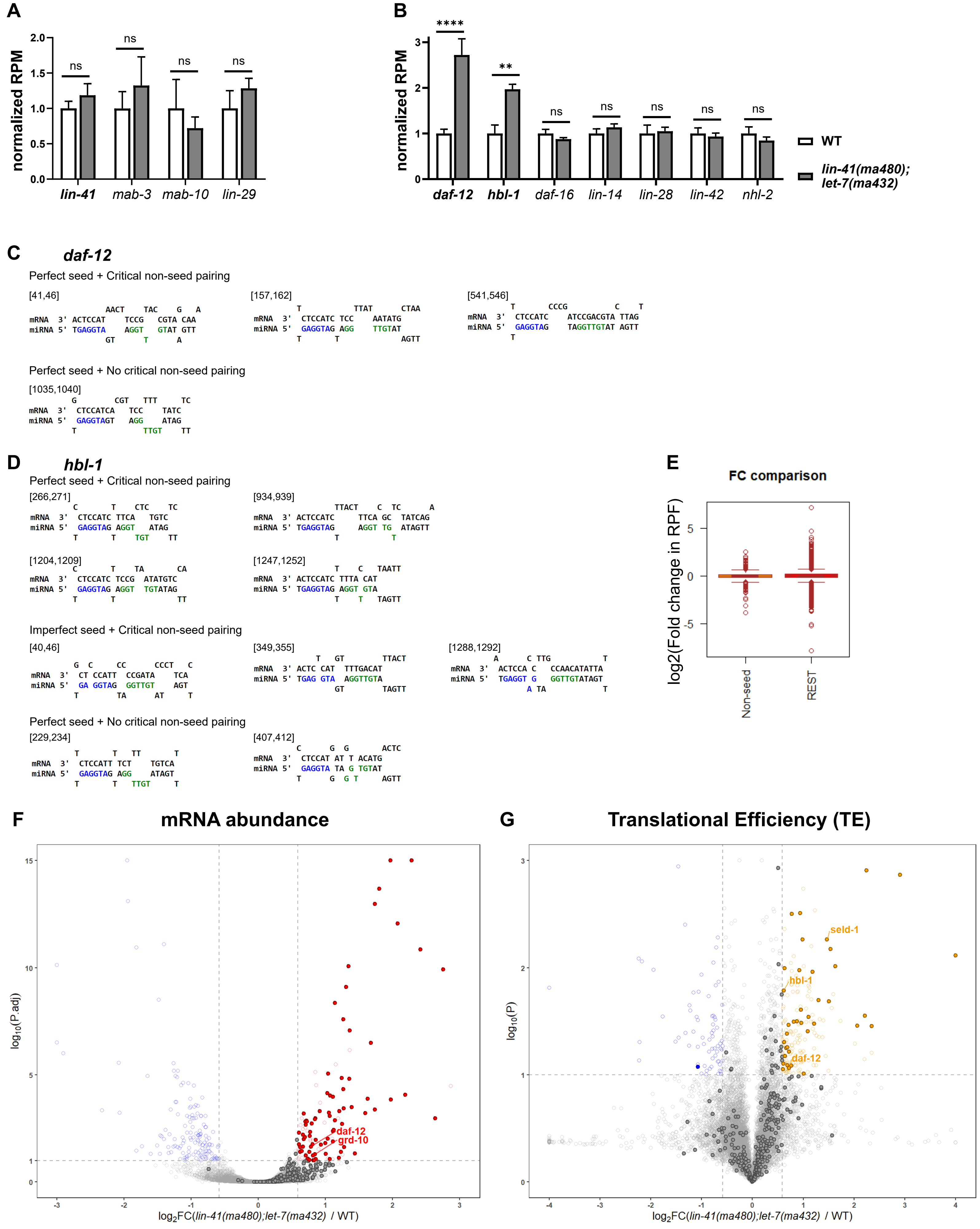
A
normalized RPM
2.0
1.5
1.0
0.5
0.0
ns
ns
ns
ns
lin-41
mab-3
mab-10
lin-29
B
normalized RPM
3
2
1
0
****
**
ns
ns
ns
ns
ns
daf-12
hbl-1
daf-16
lin-14
lin-28
lin-42
nhl-2
WT
lin-41(ma480); let-7(ma432)
C
daf-12
Perfect seed + Critical non-seed pairing
[41,46]
[157,162]
[541,546]
Perfect seed + No critical non-seed pairing
[1035,1040]
D
hbl-1
Perfect seed + Critical non-seed pairing
[266,271]
[934,939]
[1204,1209]
[1247,1252]
Imperfect seed + Critical non-seed pairing
[40,46]
[349,355]
[1288,1292]
Perfect seed + No critical non-seed pairing
[229,234]
[407,412]
E
FC comparison
log2(Fold change in RPF)
5
0
-5
Non-seed
REST
F
mRNA abundance
log10(P.adj)
daf-12
grd-10
log2FC(lin-41(ma480);let-7(ma432) / WT)
G
Translational Efficiency (TE)
log10(P)
seld-1
hbl-1
daf-12
log2FC(lin-41(ma480);let-7(ma432) / WT)

**Figure S6. Heterochronic genes *daf-12* and *hbl-1* are *let-7a* 3'-sup targets that de-repressed in *let-7a* non-seed mutant.** (Related to **Figures 4, 5**).

**A.** Normalized RPFs of *lin-41* and its reported direct downstream targets in WT and *lin-41(ma480);let-7(ma432)*. (Aeschimann et al., 2017). The results further support that repression of *lin-41* has been rescued by restoring the native *let-7a:lin-41* LCSs interacting configuration in Fig. 4. Note that another reported *lin-41* target *dmd-3* is excluded for this analysis due to its low read counts.

**B.** Normalized RPFs of *C.elegans* critical heterochronic genes, including *daf-12* and *hbl-1*, in WT and *lin-41(ma480);let-7(ma432)*. Significance in A-B is evaluated by DESeq2 (Love et al., 2014).

**C-D.** Predicted base pairing configuration between *let-7a* and the predicted LCSs of *daf-12* **(C)** and *hbl-1* **(D)** including sites with pairing in the critical non-seed region (green). The categorization of seed and non-seed types is shown in the Methods section. Sites with "weak seed + no critical non-seed" configuration (2/3, -1) were excluded. The numbers indicate locations of the *let-7a* seed in 3' UTR. Both the interacting configurations and seed locations are further shown in Dataset S1 and Table S4.

**E.** A comparison of fold change (FC) in RPF levels between the genes that are classified as 3'-sup targets (Non-seed) and other genes (REST). Wilcoxon rank test (wilcox.test in R) confirmed that the difference between the two groups is insignificant. We reason that the lack of enrichment of 3' targets among the over-expressed genes was due to the possible scenario that these 3' targets are de-repressed in specific tissues/cells, likely where *let-7a* is highly expressed, while the RPFs in the Ribo-seq were collected from whole animals, thus signals of the significant over-expression of *let-7a* targets in *let-7a*-expressing tissues are diluted. We suggest translational profiles at single-cell resolution to be taken to further investigate the above issue.

**F-G**. Categorization of the repressing mechanisms of *let-7a* non-seed region, shown by volcano plots of the differential expression analysis of RNA-seq **(F)** and TE **(G)** between *lin-41(tn1541ma480);let-7(ma432)* and *lin-41(tn1541).* Significantly increased/decreased genes (FC > |1.5| and P.adj < 0.1 by DESeq2 **(F)** or P < 0.1 by t-test **(G)**) are colored red/blue. Solid points of both plots represent genes with significantly increased RPFs in ribosome profiling in Figure 5, among which genes with predicted *let-7a* non-seed sites are labeled with gene names.

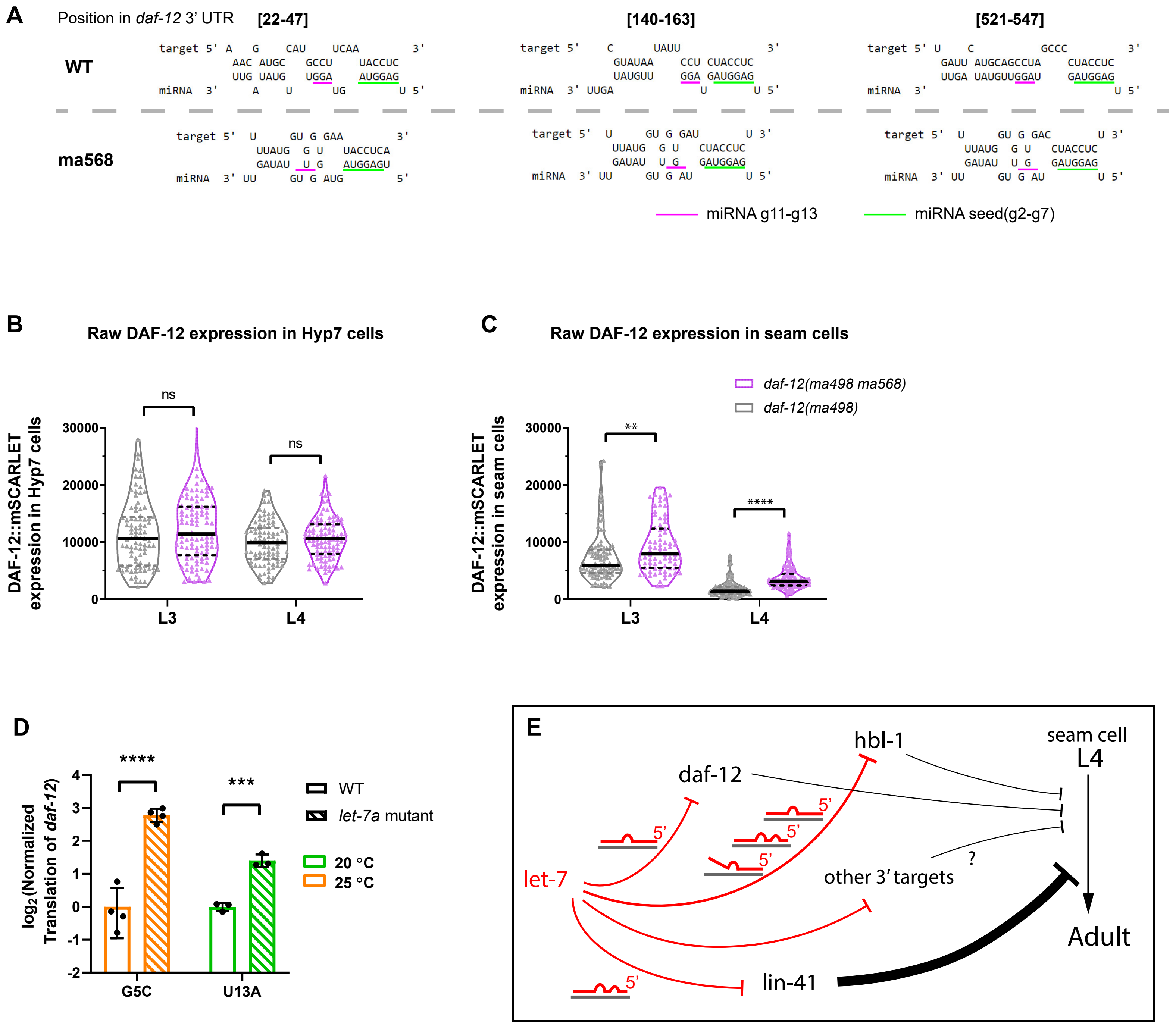

A
Position in daf-12 3' UTR
[22-47]
[140-163]
[521-547]
WT
ma568
miRNA g11-g13
miRNA seed(g2-g7)
B
Raw DAF-12 expression in Hyp7 cells
DAF-12::mSCARLET expression in Hyp7 cells
ns
L3
L4
C
Raw DAF-12 expression in seam cells
daf-12(ma498 ma568)
daf-12(ma498)
DAF-12::mSCARLET expression in seam cells
**
****
D
log2(Normalized Translation of daf-12)
****
***
WT
let-7a mutant
20 °C
25 °C
G5C
U13A
E
hbl-1
daf-12
seam cell L4
let-7
other 3′ targets
?
lin-41
Adult

**Figure S7. *daf-12* is repressed by *let-7 family* miRNAs at different developmental stages with distinct pairing configuration and repressing effect.** (Related to **Figure 5**).

**A.** Base pairing configurations between *let-7a* (miRNA) and the LCSs in *daf-12* 3' UTR which contain 3' pairing at g11-g13 in WT (target). The seed positions in 3' UTR corresponds to those in Dataset S1.

**B-C.** The DAF-12::mSCARLET fluorescent intensity at the L3/L4 stage, in Hyp7 nuclei (**B**) and seam cell nuclei (**C**). These values were used to calculate the normalized expression in Fig. 5F. The fluorescent intensity per cell was quantified as the product of the area of the nucleus (determined in the register of the DIC channel) and the relative mean density; the latter was calculated by removing the average mean density of surrounding regions from the raw mean density of the nucleus region.

**D.** Translational expression levels of *daf-12* in *let-7(n2853, G5C)* and *lin-41(tn1541ma480);let-7(ma432, U13A)* at L4 stages. Data of *lin-41(tn1541ma480);let-7(ma432)* were obtained from Fig. 5A and normalized to *lin-41(tn1541)*. Data of *let-7(n2853, G5C)* were obtained from (Aeschimann et al., 2017) at time points 26-32 hrs and normalized to N2. Note that data for the G5C mutant was obtained at 25 °C, and data of the U13A mutant was obtained at 20 °C.

**E.** Illustrative model for the genetic interactions of *let-7 family* miRNAs with heterochronic pathway genes, and the corresponding base pairing patterns. Red lines indicate the repressive effect of *let-7a*. Black lines indicate the genetic regulation to the L4-to-adult, whereas the line thickness indicates estimated regulation efficacy.

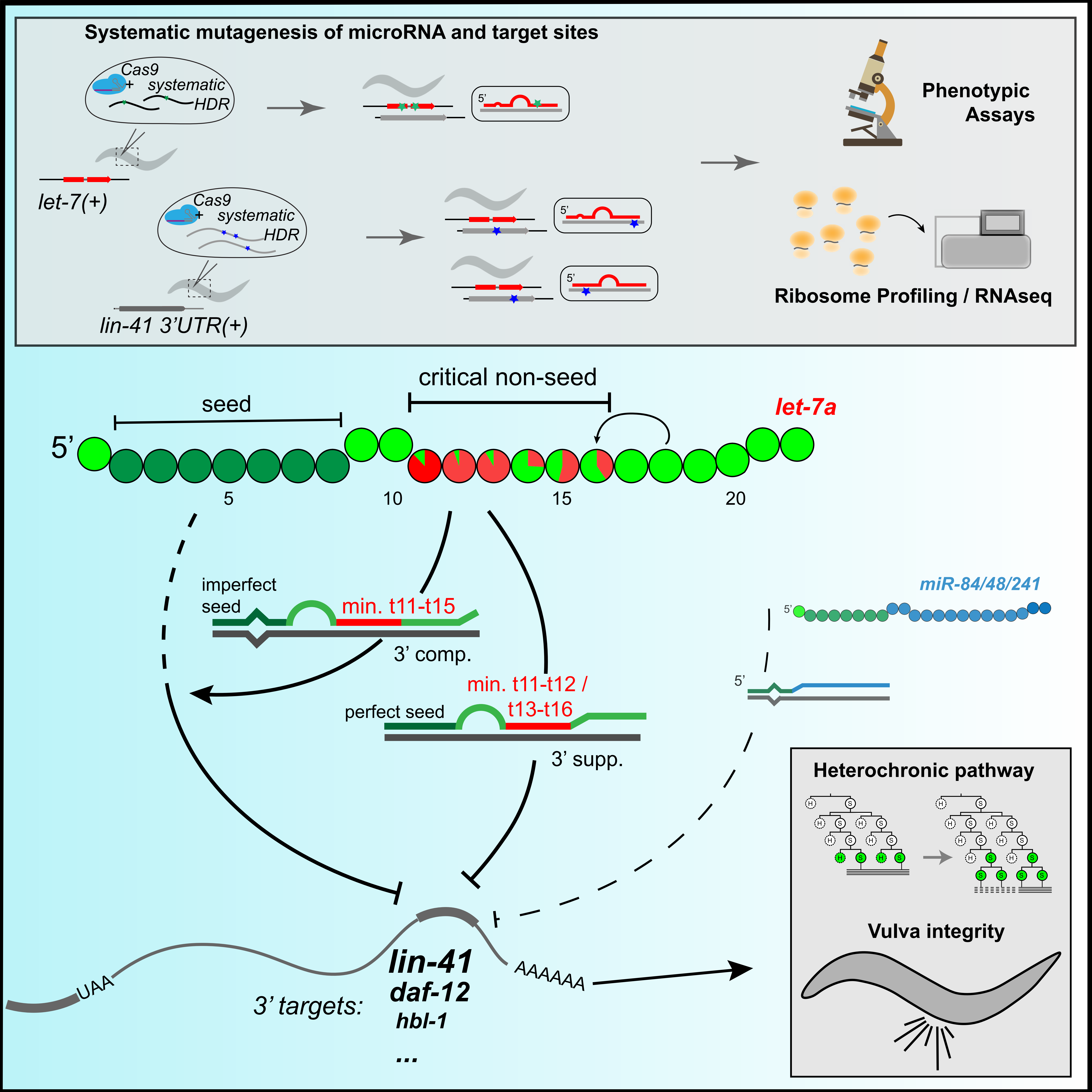

Systematic mutagenesis of microRNA and target sites
Cas9
+ systematic
HDR
let-7(+)
Cas9
+ systematic
HDR
lin-41 3'UTR(+)
5'
5'
5'
Phenotypic Assays
Ribosome Profiling / RNAseq
critical non-seed
seed
let-7a
5'
5
10
15
20
imperfect seed
min. t11-t15
3' comp.
min. t11-t12 / t13-t16
perfect seed
3' supp.
miR-84/48/241
5'
5'
Heterochronic pathway
Vulva integrity
UAA
AAAAAA
3' targets:
lin-41
daf-12
hbl-1
...
