## Supplementary material for "Critical contribution of 3’ non-seed base pairing to the *in vivo* function of the evolutionarily conserved *let-7a* microRNA": Key Resources Table

| REAGENT or RESOURCE | SOURCE | IDENTIFIER |
|---|---|---|
| Antibodies | | |
| N/A | | |
| Bacterial and Virus Strains | | |
| *E.coli* HB101 | | N/A |
| *E.coli* HT115 RNAi clones | (Kamath et al., 2003) | N/A |
| Biological Samples | | |
| N/A | | |
| Chemicals, Peptides, and Recombinant Proteins | | |
| 1X duplex buffer | IDT | Cat# 11010301 |
| QIAzol Lysis Reagent | Qiagen | Cat# 79306 |
| GlycoBlue | Invitrogen | Cat# AM9516 |
| Protease Inhibitor Cocktail | Sigma-Aldrich | Cat# P2714 |
| Turbo DNase | Invitrogen | Cat# AM2238 |
| 400 $\mu$m silica beads | OPS Diagnostic | Cat# PFAW-400-100-04 |
| RNase I | Invitrogen | Cat# AM2294 |
| SUPERase.In RNase Inhibitor | Invitrogen | Cat# AM2694 |
| SYBR Gold Nucleic Acid Gel Stain | Invitrogen | Cat# S11494 |
| T4 PNK | NEB | Cat# M0201S |
| RNase Inhibitor, Murine | NEB | Cat# M0314S |
| Thermostable RNase H | MCLAB | Cat# HTRH-100 |
| Critical Commercial Assays | | |
| Q5 site-directed mutagenesis kit | NEB | Cat# E0552S |
| QIAseq miRNA Library kit | Qiagen | Cat# 331505 |
| QIAseq miRNA NGS 96 Index IL | Qiagen | Cat# 331565 |
| KAPA Library Quantification Kit | KAPA Biosystems | Cat# KK4824 |
| RNA Clean & Concentrator-5 Kit | Zymo | Cat# R1015 |

| | | |
|---|---|---|
| NEBNext Multiplex Small RNA Library Prep Set | NEB | Cat# E7300 |
| NEBNext Ultra II RNA Library Prep kit | NEB | Cat# E7775 |
| NEBNext Multiplex oligo for Illumina | NEB | Cat# E7500 E7335 E7710 E7730 |
| HiFiScript gDNA Removal cDNA Synthesis Kit | CWBIO | CW2020M |
| UltraSYBR Mixture (Low ROX) | CWBIO | CW2601L |
| **Deposited Data** | | |
| Raw and processed data of ribosome profiling | This study | GEO: GSE171726 |
| Raw and processed data of RNA-seq | This study | GEO: GSE171733 |
| Raw and processed data of small RNA seq | This study | GEO: GSE171747 |
| *C.elegans* Reference Genome WBCel235 | Ensemble / iGenome (Illumina) | https://support.illumina.com/sequencing/sequencing_software/igenome.html |
| miRBase release 22.1 | miRBase | http://www.mirbase.org/ftp.shtml |
| **Experimental Models: Cell Lines** | | |
| N/A | | |
| **Experimental Models: Organisms/Strains** | | |
| *C.elegans*: Strain N2 | Caenorhabditis Genetics Center | WormBase Strain: N2 |
| *C.elegans:* Strain VT1313 *(maIs105 V; mir-84(n4037) X)* | this study | VT1313 |
| *C.elegans:* Strain VT1367 (*maIs105 V*) | this study * | VT1367 |
| *C.elegans:* Strain VT2692 *(maIs105 V; let-7(n2853) X)* | this study * | VT2692 |
| *C.elegans:* Strain VT3439 *(maIs105 V; let-7(ma341) X)* | this study * | VT3439 |
| *C.elegans:* Strain VT3455 *(wIs51 V)* | this study * | VT3455 |
| *C.elegans:* Strain VT3460 *(wIs51 V; let-7a(ma341) X)* | this study * | VT3460 |
| *C.elegans:* Strain VT3479 *(maIs105 V; let-7(ma341);mir-84(n4037) X)* | this study * | VT3479 |
| *C.elegans:* Strain VT3585 *(lin-41(ma378) I; maIs105 V; let-7(ma341) X)* | this study * | VT3585 |
| *C.elegans:* Strain VT3639 *(lin-41(ma378) I; maIs105 V)* | this study * | VT3639 |
| *C.elegans:* Strain VT3645 *(mnDp1(umnIs25) (X;V),+ / +,maIs105 V; let-7(ma393) X)* | (Duan et al., 2020) | VT3645 |

| | | |
|---|---|---|
| *C.elegans:* Strain VT3648 *(mnDp1(umnIs25) (X;V),+/+,maIs105 V; let-7(ma341) X)* | this study * | VT3648 |
| *C.elegans:* Strain VT3729 *(mnDp1(umnIs25) (X;V),+/+,maIs105 V; let-7(ma432ma435) X)* | this study * | VT3729<br><br>*ma432ma435* is also labeled as *ma428* |
| *C.elegans:* Strain VT3742 *(oxSi1091(Pmex-5::Cas-9(smu-2 introns) unc-119+) II; mnDp1(umnIs25) (X;V)/+ V; let-7(ma393) X)* | this study * | VT3742 |
| *C.elegans:* Strain VT3793 *(mnDp1(umnIs25) (X;V),+/+,maIs105 V; let-7(ma431) X)* | this study * | VT3793 |
| *C.elegans:* Strain VT3794 *(mnDp1(umnIs25) (X;V),+/+,maIs105 V; let-7(ma432) X)* | this study * | VT3794 |
| *C.elegans:* Strain VT3795 *(mnDp1(umnIs25) (X;V),+/+,maIs105 V; let-7(ma433) X)* | this study * | VT3795 |
| *C.elegans:* Strain VT3796 *(mnDp1(umnIs25) (X;V),+/+,maIs105 V; let-7(ma434) X)* | this study * | VT3796 |
| *C.elegans:* Strain VT3797 *(maIs105 V; let-7(ma435) X)* | this study * | VT3797 |
| *C.elegans:* Strain VT3798 *(maIs105 V; let-7(ma436) X)* | this study * | VT3798 |
| *C.elegans:* Strain VT3799 *(maIs105 V; let-7(ma437) X)* | this study * | VT3799 |
| *C.elegans:* Strain VT3825 *(mnDp1(umnIs25) (X;V),+/+,maIs105 V; let-7(ma448) X)* | this study * | VT3825 |
| *C.elegans:* Strain VT3826 *(mnDp1(umnIs25) (X;V),+/+,maIs105 V; let-7(ma449) X)* | this study * | VT3826 |
| *C.elegans:* Strain VT3827 *(maIs105 V; let-7(ma450) X)* | this study * | VT3827 |
| *C.elegans:* Strain VT3828 *(maIs105 V; let-7(ma451) X)* | this study * | VT3828 |
| *C.elegans:* Strain VT3829 *(maIs105 V; let-7(ma452) X)* | this study * | VT3829 |
| *C.elegans:* Strain VT3835 *(maIs105 V; let-7(ma453) X)* | this study * | VT3835 |
| *C.elegans:* Strain VT3836 *(maIs105 V; let-7(ma454) X)* | this study * | VT3836 |
| *C.elegans:* Strain VT3843 *(maIs105 V; let-7(ma455) X)* | this study * | VT3843 |
| *C.elegans:* Strain VT3844 *(maIs105 V; let-7(ma456) X)* | this study * | VT3844 |
| *C.elegans:* Strain VT3867 *(lin-41(tn1541) I; oxSi1091 II; mnDp1(umnIs25) (X;V)/+ V; let-7(ma432) X)* | this study * | VT3867 |
| *C.elegans:* Strain VT3868 *(lin-41(tn1541) I; mnDp1(umnIs25) (X;V)/+ V; let-7(ma432) X)* | this study * | VT3868 |
| *C.elegans:* Strain VT3873 *(oxSi1091 II; mnDp1(umnIs25) (X;V)/+ V; let-7(ma432) X)* | this study * | VT3873 |
| *C.elegans:* Strain VT3874 *(maIs105 V; let-7(ma476) X)* | this study * | VT3874 |
| *C.elegans:* Strain VT3875 *(maIs105 V; let-7(ma477) X)* | this study * | VT3875 |

| | | |
|---|---|---|
| *C.elegans:* Strain VT3876 *(maIs105 V; let-7a(ma478) X)* | this study * | VT3876 |
| *C.elegans:* Strain VT3877 *(mnDp1(umnIs25) (X;V),+/+,maIs105 V; let-7(ma449ma435) X)* | this study * | VT3877<br>*ma449ma435* is also labeled as *ma479* |
| *C.elegans:* Strain VT3878 *(lin-41(tn1541ma480) I; let-7(ma432) X)* | this study * | VT3878 |
| *C.elegans:* Strain VT3879 *(lin-41(tn1541ma480) I; maIs105 V; let-7(ma432) X)* | this study * | VT3879 |
| *C.elegans:* Strain VT3945 *(lin-41(tn1541ma501) I; maIs105 V)* | this study * | VT3945 |
| *C.elegans:* Strain VT3949 *(lin-41(tn1541ma480) I)* | this study * | VT3949 |
| *C.elegans:* Strain VT3950 *(lin-41(tn1541ma480) I; maIs105 V)* | this study * | VT3950 |
| *C.elegans:* Strain VT3954 *(lin-41(tn1541ma501) I; mnDp1(umnIs25) (X;V),+/+,maIs105 V; let-7(ma393) X)* | this study * | VT3954 |
| *C.elegans:* Strain VT3959 *(lin-41(tn1541) I; mnDp1(umnIs25) (X;V),+/+,maIs105 V; let-7(ma393) X)* | this study * | VT3959 |
| *C.elegans:* Strain VT3974 *(lin-41(ma501ma545) I; maIs105 V)* | this study * | VT3974<br>*ma501ma545* is also labeled as *ma511* |
| *C.elegans:* Strain VT3975 *(lin-41(xe11) I; maIs105 V)* | this study * | VT3975 |
| *C.elegans:* Strain VT4009 *(lin-41(ma501ma572) I; maIs105 V)* | this study * | VT4009<br>*ma501ma572* is also labeled as *ma554* |
| *C.elegans:* Strain VT4058 *(lin-41(ma501ma480) I; maIs105 V)* | this study * | VT4058<br>*ma501ma480* is also labeled as *ma504* |
| *C.elegans:* Strain VT4060 *(lin-41(ma480) I; maIs105 V)* | this study * | VT4060 |
| *C.elegans:* Strain VT4091 *(lin-41(ma501ma555) I; maIs105 V)* | this study * | VT4091<br>*ma501ma555* is also labeled as *ma552* |
| *C.elegans:* Strain VT4092 *(lin-41(ma501ma556) I; maIs105 V)* | this study * | VT4092<br>*ma501ma556* is also labeled as *ma553* |
| *C.elegans:* Strain VT4093 *(lin-41(ma501ma571) I; maIs105 V)* | this study * | VT4093<br>*ma501ma571* is also labeled as *ma554* |
| *C.elegans:* Strain VT4094 *(lin-41(ma555) I; maIs105 V)* | this study * | VT4094 |
| *C.elegans:* Strain VT4095 *(lin-41(ma556) I; maIs105 V)* | this study * | VT4095 |
| *C.elegans:* Strain VT4123 *(lin-41(ma564) I; maIs105 V)* | this study * | VT4123 |
| *C.elegans:* Strain VT4124 *(lin-41(ma565) I; maIs105 V)* | this study * | VT4124 |
| *C.elegans:* Strain VT4125 *(lin-41(ma566) I; maIs105V)* | this study * | VT4125 |
| *C.elegans:* Strain VT4126 *(oxSi1091 II; maIs105 V; daf-12(ma498ma567) X)* | this study * | VT4126 |

| | | |
|---|---|---|
| *C.elegans:* Strain VT4128 *(lin-41(tn1541) I; maIs105 V; daf-12(ma498ma568) X)* | this study * | VT4128 |
| *C.elegans:* Strain VT4129 *(lin-41(tn1541) I; maIs105 V; daf-12(ma498) X)* | this study * | VT4129 |
| *C.elegans:* Strain DG3913 *(lin-41(tn1541) I)* | Caenorhabditis Genetics Center | WormBase: DG3913 |
| **Oligonucleotides** | | |
| 30mer_RPF_marker_RNA<br><br>rArCrUrArGrCrCrUrUrArUrUrUrUrArArCrUrUrGrCrUrArUrGrCrUrCrUrA | this study<br><br>(ordered from IDT) | N/A |
| Alt-R *let-7a* WT crRNA:<br><br>TGTGGATCCGGTGAGGTAGT + Alt-R | this study<br><br>(ordered from IDT) | AltR_Cas-9 _crRNA_let-7_g3 |
| Alt-R *let-7a* jump board crRNA_2:<br><br>ATCAGTTCGATATCTGACGG + Alt-R | this study<br><br>(ordered from IDT) | AltR_Cas-9 _crRNA_INPP4A_1 |
| Alt-R *let-7a* jump board crRNA_2:<br><br>TGTATCAGTTCGATATCTGA + Alt-R | this study<br><br>(ordered from IDT) | AltR_Cas-9 _crRNA_INPP4A_2 |
| Alt-R dpy-10 crRNA as co-CRISPR marker:<br><br>CTACCATAGGCACCACGAG + Alt-R | (Arribere et al., 2014)<br><br>(ordered from IDT) | AltR_Cas-9 _crRNA_dpy-10_cn64 |
| Alt-R *lin-41* crRNA_1:<br><br>CATCGCGTTGAGTGTAGAA + Alt-R | this study<br><br>(ordered from IDT) | AltR_Cas-9 _crRNA_lin-41_1 |
| Alt-R *lin-41* crRNA_2:<br><br>CAATGGTTCAGAGGCAGAA + Alt-R | this study<br><br>(ordered from IDT) | AltR_Cas-9 _crRNA_lin-41_2 |
| Alt-R *daf-12* crRNA_4:<br><br>AGAGAGGAATTAAGAGGAAGGTTTTAGAGCTATGCT + Alt-R | this study<br><br>(ordered from IDT) | AltR_Cas-9 _crRNA_daf-12_4 |
| Alt-R CRISPR-Cas-9 tracrRNA | IDT | Cat# 1072533 |
| *let-7a* cloning primer reverse:<br><br>ATTGAAAATCTGTCATTGAGCAA | this study | let-7_Donor_R4 |
| *let-7a* cloning primer forward:<br><br>GCTCAAGGTTTTCGATCTCTGT | this study | let-7_Donor_F4 |
| *let-7a* HR donor amplifying primer forward:<br><br>ATAGCACAAAATAAAGAAAAACAAAGAGGTGAAAG | this study | let-7_SEQ_F4 |
| *let-7a* HR donor amplifying primer reverse:<br><br>AATTTAACAACAAGTACTAATCCATTTTTCAGGCAAGC | this study | let-7_SEQ_R4 |

| | | |
|---|---|---|
| *let-7a* genotyping primer forward:<br><br>AACGGCTCCATGGATACATTACTCAACAG | this study | let-7_SEQ_F5 |
| *let-7a* genotyping primer reverse:<br><br>GGTTTCTGTTCATATATGAGAAGCGCATCAG | this study | let-7_SEQ_R5 |
| *lin-41* genotyping primer forward:<br><br>CATCCATTCATATGGCTCCGCCCC | this study | lin-41_SEQ_F3 |
| *lin-41* genotyping primer reverse:<br><br>CACTGGGGACATTAGGCAATTGGGAC | this study | lin-41_SEQ_R3 |
| *daf-12* genotyping primer forward:<br><br>CGAGGGACGTCACTCCACCGG | this study | daf-12_SEQ_F1 |
| *daf-12* genotyping primer forward:<br><br>GGCGTTGGGAGTTGAAAGTCTTAAATAG | this study | daf-12_SEQ_R1 |
| *daf-12* genotyping primer forward:<br><br>GATTCCAAAAGCACTGGGATTACTTAATGTAAG | this study | daf-12_SEQ_F2 |
| *daf-12* genotyping primer forward:<br><br>AAACTTCAAATTATCATGCTTAGTTCTCCATCG | this study | daf-12_SEQ_R2 |
| *lin-41* qRT-PCR primer forward*:*<br><br>CGGATACTCGGAATCATCGTGTTCAG | this study<br><br>(ordered from IDT) | lin-41_RTPCR_F |
| *lin-41* qRT-PCR primer reverse*:*<br><br>GACTGCGAGTCGGTGATTGTTGAAG | this study<br><br>(ordered from IDT) | lin-41_RTPCR_R |
| *gpd-1* qRT-PCR primer forward*:*<br><br>TGCTCCAATGTACGTGGTTGGAG | this study<br><br>(ordered from IDT) | gpd-1_RTPCR_F |
| *gpd-1* qRT-PCR primer reverse*:*<br><br>CGTCATGAGTCCTTCGATGATACCG | this study<br><br>(ordered from IDT) | gpd-1_RTPCR_R |
| *lin-41* ssDNA donor for *ma480:*<br><br>TCAAATTGCACCAACTCAAGTATACCTTTTATACATCCGTTCTACACTCAACGCGATGTAAATATC GCAATCCCTTTTTATACATCCATTCTGCCTCTGAACCATTGAAACCTTC | this study<br><br>(IDT, Ultramer) | N/A |
| *lin-41* ssDNA donor for *ma504:*<br><br>CCAACTCAAGTATACCTTTTATACATCCGTTCTACCTCAACGCGATGTAAATATCGCAATCCCTTTT TATACATCCATTCTACCTCTGAACCATTGAAACCTTCTCCCGTACTCCCA | this study<br><br>(IDT, Ultramer) | N/A |
| *lin-41* ssDNA donor for *ma501:*<br><br>CCAACTCAAGTATACCTTTTATACAACCGTTCTACCTCAACGCGATGTAAATATCGCAATCCCTTTT TATACAACCATTCTACCTCTGAACCATTGAAACCTTCTCCCGTACTCCCA | this study<br><br>(IDT, Ultramer) | N/A |

| | | |
|---|---|---|
| *lin-41* ssDNA donor for *ma511:*<br><br>TTGCACCAACTCAAGTATACCTTTTATACATGGGTTCTACCTCAACGCGATGTAAATATCGCAATC<br>CCTTTTTATACATGGATTCTACCTCTGAACCATTGAAACCTTCTCCCGTAC | this study<br><br>(IDT, Ultramer) | N/A |
| *lin-41* ssDNA donor for *ma378:*<br><br>TTCCTCAAATTGCACCAACTCAAGTATACCATTAATATTACAGTTCTACACTCAACGCGATGTAAA<br>TATCGCAATCCCTATTAATATTACAATTCTGCCTCTGAACCATTGAAACCTTCTCCCGTAC | this study<br><br>(IDT, Ultramer) | N/A |
| *lin-41* ssDNA donor for *ma552:*<br><br>TCAAATTGCACCAACTCAAGTATACCTTTTATACAACGGTTCTACCTCAACGCGATGTAAATATCG<br>CAATCCCTTTTTATACAACGATTCTACCTCTGAACCATTGAAACCTTCTCCCGTACTCCCA | this study<br><br>(Genewiz,Oligo-Flex) | N/A |
| *lin-41* ssDNA donor for *ma553:*<br><br>TCAAATTGCACCAACTCAAGTATACCTTTTATACAAGCGTTCTACCTCAACGCGATGTAAATATCG<br>CAATCCCTTTTTATACAAGCATTCTACCTCTGAACCATTGAAACCTTCTCCCGTACTCCCA | this study<br><br>(Genewiz,Oligo-Flex) | N/A |
| *lin-41* ssDNA donor for *ma554:*<br><br>TCAAATTGCACCAACTCAAGTATACCTTTTATACAAGGGTTCTACCTCAACGCGATGTAAATATCG<br>CAATCCCTTTTTATACAAGGATTCTACCTCTGAACCATTGAAACCTTCTCCCGTACTCCCA | this study<br><br>(Genewiz,Oligo-Flex) | N/A |
| *lin-41* ssDNA donor for *ma525:*<br><br>CAAATTGCACCAACTCAAGTATACCTTTTATTGTTCCGTTCTACCTCAACGCGATGTAAATATCGC<br>AATCCCTTTTTATTGTTCCATTCTACCTCTGAACCATTGAAACCTTCTCC | this study<br><br>(Genewiz,Oligo-Flex) | N/A |
| *lin-41* ssDNA donor for *ma555:*<br><br>TCAAATTGCACCAACTCAAGTATACCTTTTATACAACGGTTCTACACTCAACGCGATGTAAATATC<br>GCAATCCCTTTTTATACAACGATTCTGCCTCTGAACCATTGAAACCTTCTCCCGTACTCCCA | this study<br><br>(IDT, Ultramer) | N/A |
| *lin-41* ssDNA donor for *ma556:*<br><br>TCAAATTGCACCAACTCAAGTATACCTTTTATACAAGCGTTCTACACTCAACGCGATGTAAATATC<br>GCAATCCCTTTTTATACAAGCATTCTGCCTCTGAACCATTGAAACCTTCTCCCGTACTCCCA | this study<br><br>(IDT, Ultramer) | N/A |
| *lin-41* ssDNA donor for *ma564:*<br>CCTCTTTTCCTCAAATTGCACCAACTCAAGTATACCTTTTATACTACCGTTCTACACTCAACGCGAT<br>GTAAATATCGCAATCCCTTTTTATACTACCATTCTGCCTCTGAACCATTGAAACCTTCTCCCGTACT<br>CCCA | this study<br><br>(Genewiz,Oligo-Flex) | N/A |
| *lin-41* ssDNA donor for *ma565:*<br><br>CCTCTTTTCCTCAAATTGCACCAACTCAAGTATACCTTTTATAAAACCGTTCTACACTCAACGCGAT<br>GTAAATATCGCAATCCCTTTTTATAAAACCATTCTGCCTCTGAACCATTGAAACCTTCTCCCGTACT<br>CCCA | this study<br><br>(Genewiz,Oligo-Flex) | N/A |
| *lin-41* ssDNA donor for *ma566:*<br><br>CCTCTTTTCCTCAAATTGCACCAACTCAAGTATACCTTTTATTCAACCGTTCTACACTCAACGCGAT<br>GTAAATATCGCAATCCCTTTTTATTCAACCATTCTGCCTCTGAACCATTGAAACCTTCTCCCGTACT<br>CCCA | this study<br><br>(Genewiz,Oligo-Flex) | N/A |

| | | |
|---|---|---|
| *daf-12* dsDNA donor for ma568:<br><br>CGAGGGACGTCACTCCACCGGAGGAATGGACGAGCTCTACAAGTAGACCTACTAGAAATCATCT<br>ATTTATGGTGGTGAATACCTCACATCTTGATTCTATATTGCCTCCATCCAACAAACTCAATCTAGCC<br>ACATTTCTTCTTTTTCACGTACCTCAACCACCCTTTCCATATTTATGGTGGTGATCTACCTCTTTAAC<br>CAATTCATCATCTTTTTATATTGTTTCTTATTTGCATTCAACTTGGAAATAGCCACTATCATATCACT<br>ATTGCGTATTTCCTTTCATTTTCTTGTCTTATTTTCTTGAGACCAGCACCAGAAGATTTTTTCGATG<br>GAGAACTAAGCATGATAATTTGAAGTTTTCCATTTAAAAAATGCAGGTAATACGGTTAATTCAAT<br>CTGCGAGTTGATGTTCGGTCTCCGGTTTTCATGTTCTACTTCTAATGACTAGAAACCCTTTTATCTA<br>ACATCCGGTCCTCCTATCCCTAATGTACCCAGTAGATATTTTTTCCCCGAATGATTAAACCTCCCAG<br>TCAAATATTGTATTTTGATTTATGGTGGTGACCTACCTCTTAATTCCGTCAATACTATCTCAGATTT<br>TCATTGAAGAACTGGTCCGGAATTATTGAATCATCAGCTAG | this study<br><br>(Genewiz, fragmentGENE) | N/A |
| **Recombinant DNA** | | |
| Plasmid: pBlueScript SK(+) *let-7a(mir-84 swap)* | this study | pBS-09 |
| Plasmid: pCR2.1-TOPO *let-7a(U9G)* | this study | pCR-32 |
| Plasmid: pCR2.1-TOPO *let-7a(A10U)* | this study | pCR-33 |
| Plasmid: pCR2.1-TOPO *let-7a(G11U)* | this study | pCR-12 |
| Plasmid: pCR2.1-TOPO *let-7a(G12U)* | this study | pCR-34 |
| Plasmid: pCR2.1-TOPO *let-7a(U13A)* | this study | pCR-25 |
| Plasmid: pCR2.1-TOPO *let-7a(U14A)* | this study | pCR-13 |
| Plasmid: pCR2.1-TOPO *let-7a(G15A)* | this study | pCR-14 |
| Plasmid: pCR2.1-TOPO *let-7a(U16G)* | this study | pCR-35 |
| Plasmid: pCR2.1-TOPO *let-7a(A17U)* | this study | pCR-36 |
| Plasmid: pCR2.1-TOPO *let-7a(U18A)* | this study | pCR-26 |
| Plasmid: pCR2.1-TOPO *let-7a(A19U)* | this study | pCR-15 |
| Plasmid: pCR2.1-TOPO *let-7a(G20C)* | this study | pCR-39 |
| Plasmid: pCR2.1-TOPO *let-7a(U21A)* | this study | pCR-40 |
| Plasmid: pCR2.1-TOPO *let-7a(U22A)* | this study | pCR-16 |
| Plasmid: pCR2.1-TOPO *let-7a(U18C)* | this study | pCR-37 |
| Plasmid: pCR2.1-TOPO *let-7a(U18G)* | this study | pCR-38 |
| Plasmid: pCR2.1-TOPO *let-7a(U16G+U18A)* | this study | pCR-41 |
| Plasmid: pCR2.1-TOPO *let-7a(17-22)* | this study | pCR-42 |
| Plasmid: pCR2.1-TOPO *let-7a(A17U+U18A+A19U)* | this study | pCR-43 |
| Plasmid: pCR2.1-TOPO *let-7a(G20C+U21A+U22A)* | this study | pCR-44 |
| Plasmid: pCR2.1-TOPO *let-7a(INPP4A, Jump Board, null)* | this study | pCR-07 |
| **Software and Algorithms** | | |

| | | |
|---|---|---|
| *let-7a* target prediction algorithm | this study | https://github.com/IsanaVekslerLublinsky/Let7_Proj_code.git |
| ImageJ-FIJI | (Schindelin et al., 2012) | https://imagej.net/Fiji |
| Prism 8 | GraphPad | https://www.graphpad.com/scientific-software/prism/ |
| SnapGene Viewer | SnapGene | https://www.snapgene.com/snapgene-viewer/ |
| Jalview2 | (Waterhouse et al., 2009) | https://www.jalview.org/ |
| Adobe Illustrator CS5 | Adobe Inc. | https://www.adobe.com/products/illustrator.html |
| Cutadapt/1.9 | (Martin, 2011) | https://cutadapt.readthedocs.io/en/stable/ |
| Samtools/1.4.1 | (Li et al., 2009) | http://samtools.sourceforge.net/ |
| Bowtie2/2.3.4.3 | (Langmead and Salzberg, 2012) | http://bowtie-bio.sourceforge.net/bowtie2/index.shtml |
| STAR/2.7.6a | (Dobin et al., 2013) | https://github.com/alexdobin/STAR |
| featureCounts (Subread/1.6.2) | (Liao et al., 2014) | http://subread.sourceforge.net/ |
| plastid/0.4.8 | (Dunn and Weissman, 2016) | https://plastid.readthedocs.io/en/latest/index.html |
| DESeq2 | (Love et al., 2014) | https://bioconductor.org/packages/release/bioc/html/DESeq2.html |
| RStudio(2019) | RStudio | https://rstudio.com/ |
| ggplot2 | (Hadley, 2016) | https://ggplot2.tidyverse.org/ |
| **Other** | | |
| SW 41 Ti swinging-bucket rotor | Beckman-Coulter | Cat# 331362 |
| Spin-X centrifuge tube filter | Millipore | Cat# CLS8160 |
| Fisherbrand™ RNase-Free Disposable Pellet Pestles | Fisher Scientific | Cat# 12-141-364 |

* Information of new alleles generated in this study is available in Table S5.

Some of the *C. elegans* strains will be available at Caenorhabditis Genetic Center (CGC). Materials generated in study are available upon request.
