## Supplementary material for "Critical contribution of 3’ non-seed base pairing to the *in vivo* function of the evolutionarily conserved *let-7a* microRNA": Dataset S1: Dataset S1.html

   
miRNA=cel-let-7-5p (TGAGGTAGTAGGTTGTATAGTT)  
  
1D\_string\_key:   
 1-match   
 2-GU   
 3-mismatch   
4-mRNA\_bulge\_G   
 5-mRNA\_bulge\_other   
 6-miRNA\_bulge   
  
SeedType:   
 1 : (2-7 full match)   
 2 : (2-7 + 1mRNA bulge on 5-7) or (2-8 with 1GU/MM 5-8)   
 3: (2-7 + 1mRNA bulge on 2-4) or (2-8 with 1GU/MM 2-4)   
NonSeedType:   
 1 : match 11-13/12-14/13-16   
 2 : match+GU 11-13/12-14/13-16   
 -1 : other   

**let-7 Targeting Results**

   
| index | gene | RiboSeqInfo | Seed loc | Full alignment | Seed/NonSeed Type | \| 1 \| WBGene00000018:abl-1(11) \| ``` log2FC=-0.135 padj=0.929 ``` \| [seedLoc = 224,230] Seq=CACTGAGTGCAACTAAACGGTGTACCGTCT \| ```          T  G    GTGG  AA        AG   C mRNA  3'  CT CCAT    CA  TCAACGTG  TCA  miRNA 5'  GA GGTA    GT  GGTTGTAT  AGT           T             A              T           311411114544113521111212541113 ``` \| SeedType = 3 NonSeedType = 1 \| \| --- \| --- \| --- \| --- \| --- \| --- \| | \| 2 \| WBGene00000043:acr-3(28) \| ``` log2FC=1.084 padj=1.0 ``` \| [seedLoc = 66,71] Seq=TTCTTCCCAATTCTATCTCG \| ```          G              CCTTCTT mRNA  3'  CTCTATC  TTAAC        miRNA 5'  GAGGTAG  GGTTG                 T       TA     TATAGTT           3111211166221113333333 ``` \| SeedType = 2 NonSeedType = 1 \| \| --- \| --- \| --- \| --- \| --- \| --- \| | \| 3 \| WBGene00000072:add-1(49) \| ``` log2FC=-0.209 padj=0.799 ``` \| [seedLoc = 126,132] Seq=GGATATACTACGCCCAATTTACCGTCA \| ```             G     TAAC   CAT     AGG mRNA  3' ACT CCATT    CCG   CATAT    miRNA 5' TGA GGTAG    GGT   GTATA                      TA     T       GTT           111411112335511235511111333 ``` \| SeedType = 3 NonSeedType = 2 \| \| --- \| --- \| --- \| --- \| --- \| --- \| | \| 4 \| WBGene00000074:adm-2(52) \| ``` log2FC=-0.139 padj=0.911 ``` \| [seedLoc = 444,450] Seq=ATAAATTTTGCCTGGATACACTCA \| ```              A   AG     TTTTAAATA mRNA  3' ACTC CAT  GTCCG          miRNA 5' TGAG GTA  TAGGT                           G      TGTATAGTT           111151113421112333333333 ``` \| SeedType = 2 NonSeedType = 2 \| \| --- \| --- \| --- \| --- \| --- \| --- \| | \| 5 \| WBGene00000079:adr-1(55) \| ``` log2FC=-0.261 padj=0.792 ``` \| [seedLoc = 1751,1756] Seq=TGACTGAATACTATCAAATACCTCC \| ```          C      AAA   T    AG    T mRNA  3'  CTCCAT   CTA CATA  TCAG  miRNA 5'  GAGGTA   GGT GTAT  AGTT           T      GTA   T                      3111111333121311115411125 ``` \| SeedType = 1 NonSeedType = 2 \| \| --- \| --- \| --- \| --- \| --- \| --- \| | \|  \|  \|  \| [seedLoc = 1854,1860] Seq=TATAGTCGAACCAGATACTCTCA \| ```              T   AGA    GC  ATAT mRNA  3' ACTC CAT   CCAA  TG     miRNA 5' TGAG GTA   GGTT  AT                      GTA    GT  AGTT           11115111333111133123333 ``` \| SeedType = 2 NonSeedType = 1 \| \| --- \| --- \| --- \| --- \| --- \| --- \| | \|  \|  \|  \| [seedLoc = 1883,1888] Seq=GCCAATGGAACATCTCTACTTCA \| ```                  TCTA   G   ACCG mRNA  3' ACTTCATC    CAA GTA     miRNA 5' TGAGGTAG    GTT TAT                      TAG    G   AGTT           11121111333511132113333 ``` \| SeedType = 3 NonSeedType = 1 \| \| --- \| --- \| --- \| --- \| --- \| --- \| | \| 6 \| WBGene00000084:aex-1(60) \| ``` log2FC=-0.104 padj=0.953 ``` \| [seedLoc = 437,443] Seq=TGGTTTCGAATTGAACTACTCTCG \| ```          G   T     AG    GCTTTGGT mRNA  3'  CTC CATCA  TTAA         miRNA 5'  GAG GTAGT  GGTT                  T         A     GTATAGTT           311151111134221133333333 ``` \| SeedType = 2 NonSeedType = 2 \| \| --- \| --- \| --- \| --- \| --- \| --- \| | \| 7 \| WBGene00000095:aha-1(70) \| ``` log2FC=-0.091 padj=0.948 ``` \| [seedLoc = 64,69] Seq=GATTCTCATTTCTATCTACTTCT \| ```          T       T    TTACTCTTAG mRNA  3'  CTTCATC ATCT           miRNA 5'  GAGGTAG TAGG                    T            TTGTATAGTT           31121111511123333333333 ``` \| SeedType = 3 NonSeedType = -1 \| \| --- \| --- \| --- \| --- \| --- \| --- \| | \| 8 \| WBGene00000100:ajm-1(75) \| ``` log2FC=-0.331 padj=0.285 ``` \| [seedLoc = 155,160] Seq=CTTTTTTCAATATATTTACCTCT \| ```          T       TATA    TTTTTTC mRNA  3'  CTCCATT    TAAC        miRNA 5'  GAGGTAG    GTTG                 T       TAG     TATAGTT           31111112333521113333333 ``` \| SeedType = 1 NonSeedType = 1 \| \| --- \| --- \| --- \| --- \| --- \| --- \| | \| 9 \| WBGene00000102:akt-1(80) \| ``` log2FC=-0.087 padj=0.957 ``` \| [seedLoc = 1061,1066] Seq=TTCATCTCTTCCTGTTCTGCCTCC \| ```          C       TT    TT TCTACTT mRNA  3'  CTCCGTC  GTCC  C        miRNA 5'  GAGGTAG  TAGG  G                 T             TT TATAGTT           311112115521113313333333 ``` \| SeedType = 2 NonSeedType = -1 \| \| --- \| --- \| --- \| --- \| --- \| --- \| | \| 10 \| WBGene00000105:alg-1(83) \| ``` log2FC=-0.039 padj=0.981 ``` \| [seedLoc = 243,248] Seq=TTCTAATTGCCGCTACCTCT \| ```          T            T      TT mRNA  3'  CTCCATCG CCG TA ATC   miRNA 5'  GAGGTAGT GGT GT TAG            T        A   T  A   TT           3111111126112321611133 ``` \| SeedType = 1 NonSeedType = 2 \| \| --- \| --- \| --- \| --- \| --- \| --- \| | \| 11 \| WBGene00000112:alh-6(90) \| ``` log2FC=0.236 padj=0.541 ``` \| [seedLoc = 334,339] Seq=CCGTCGGCAACATGTACCTCA \| ```                 GTA      GCTGCC mRNA  3' ACTCCAT    CAACG       miRNA 5' TGAGGTA    GTTGT                       GTAG     ATAGTT           1111111333611112333333 ``` \| SeedType = 1 NonSeedType = 1 \| \| --- \| --- \| --- \| --- \| --- \| --- \| | \| 12 \| WBGene00000114:alh-8(95) \| ``` log2FC=0.14 padj=0.816 ``` \| [seedLoc = 405,410] Seq=CTGCGATAACCCAGATATCAAGGCTATCTCA \| ```                   GAACTATAGAC      GC      mRNA  3' ACTCTATCG           CCAATA  GTC   miRNA 5' TGAGGTAGT           GGTTGT  TAG                     A                A    TT           111121112355555554551111213521166 ``` \| SeedType = 2 NonSeedType = 1 \| \| --- \| --- \| --- \| --- \| --- \| --- \| | \|  \|  \|  \| [seedLoc = 1203,1209] Seq=ACGGAAAGGCCGGAATCCAGTTCTACACTCA \| ```              A    TTGACCTAAGG    AAAGG    mRNA  3' ACTC CATC           CCGG     CA  miRNA 5' TGAG GTAG           GGTT     GT                    TA             GTATA  T           11115111133455555544112233333116 ``` \| SeedType = 2 NonSeedType = 2 \| \| --- \| --- \| --- \| --- \| --- \| --- \| | \| 13 \| WBGene00000146:ape-1(116) \| ``` log2FC=-0.077 padj=0.949 ``` \| [seedLoc = 65,70] Seq=AAGTATAATGACCCCATTCACTCTTACCTCT \| ```          T       C  ACTTACCCCAG       AA   mRNA  3'  CTCCATT TC           TAATATG     miRNA 5'  GAGGTAG AG           GTTGTAT              T       T                    AGTT           311111123115555555555421121123366 ``` \| SeedType = 1 NonSeedType = -1 \| \| --- \| --- \| --- \| --- \| --- \| --- \| | \|  \|  \|  \| [seedLoc = 186,191] Seq=ATCTGTGTGAAGAACCCTACTTCA \| ```                        G AGTG    TA mRNA  3' ACTTCATC  CCAA A    TGTC   miRNA 5' TGAGGTAG  GGTT T    ATAG                    TA    G         TT           11121111661111315454121133 ``` \| SeedType = 3 NonSeedType = 1 \| \| --- \| --- \| --- \| --- \| --- \| --- \| | \|  \|  \|  \| [seedLoc = 197,201] Seq=GTGTGAAGAACCCTACTTCAT \| ```          T              G AGTGTG mRNA  3'  ACTTCATC  CCAA A       miRNA 5'  TGAGGTAG  GGTT T                         TA    G ATAGTT           51112111166111131333333 ``` \| SeedType = 3 NonSeedType = 1 \| \| --- \| --- \| --- \| --- \| --- \| --- \| | \| 14 \| WBGene00000149:apl-1(119) \| ``` log2FC=-0.14 padj=0.9 ``` \| [seedLoc = 67,72] Seq=AATGATCCATTCTCCTATCTCA \| ```                  C   T  CTAGTAA mRNA  3' ACTCTATC TCT AC        miRNA 5' TGAGGTAG AGG TG                         T   T  TATAGTT           1111211131123113333333 ``` \| SeedType = 2 NonSeedType = -1 \| \| --- \| --- \| --- \| --- \| --- \| --- \| | \|  \|  \|  \| [seedLoc = 79,83] Seq=TGATCCATTCTCCTATCTCAG \| ```          G        C   T  CTAGT   mRNA  3'  ACTCTATC TCT AC        miRNA 5'  TGAGGTAG AGG TG                          T   T  TATAGTT           41111211131123113333366 ``` \| SeedType = 2 NonSeedType = -1 \| \| --- \| --- \| --- \| --- \| --- \| --- \| | \| 15 \| WBGene00000156:apr-1(123) \|  \| [seedLoc = 91,96] Seq=TTCTCATCTTCTTATTCTACCTCG \| ```          G       TT    TT TACTCTT mRNA  3'  CTCCATC  ATTC  C        miRNA 5'  GAGGTAG  TAGG  G                 T             TT TATAGTT           311111115511213313333333 ``` \| SeedType = 1 NonSeedType = -1 \| \| --- \| --- \| --- \| --- \| --- \| --- \| | \| 16 \| WBGene00000158:apg-1(125) \| ``` log2FC=0.003 padj=0.998 ``` \| [seedLoc = 397,403] Seq=AGGTAGGCAAGCCTACTATCCTCA \| ```               T       G    GATGGA mRNA  3' ACTCC ATCATCC AACG       miRNA 5' TGAGG TAGTAGG TTGT                                  ATAGTT           111115111111141112333333 ``` \| SeedType = 2 NonSeedType = 1 \| \| --- \| --- \| --- \| --- \| --- \| --- \| | \| 17 \| WBGene00000168:apx-1(134) \| ``` log2FC=-0.107 padj=0.974 ``` \| [seedLoc = 191,197] Seq=TATTATTGCTCCCTGTTAACCTCG \| ```          G    A       CT  T     T mRNA  3'  CTCC ATTGTCC  CG TATTA  miRNA 5'  GAGG TAGTAGG  GT ATAGT           T            TT        T           311115112211133125111213 ``` \| SeedType = 2 NonSeedType = -1 \| \| --- \| --- \| --- \| --- \| --- \| --- \| | \| 18 \| WBGene00000170:aqp-2(136) \| ``` log2FC=-0.014 padj=0.994 ``` \| [seedLoc = 23,29] Seq=ACGATTCGACCAATATTCAATGCTACCATCC \| ```          C  A       AACTTATAA     TTAG    mRNA  3'  CT CCATCGT         CCAGC    CA  miRNA 5'  GA GGTAGTA         GGTTG    GT           T                        TATA  T           31151111121555555555111213333116 ``` \| SeedType = 3 NonSeedType = 1 \| \| --- \| --- \| --- \| --- \| --- \| --- \| | \|  \|  \|  \| [seedLoc = 31,36] Seq=TTCAATGCTACCATCCTGCCTCC \| ```          C       CTA   T    ACTT mRNA  3'  CTCCGTC   CCA CGTA     miRNA 5'  GAGGTAG   GGT GTAT              T       TA    T    AGTT           31111211335111312113333 ``` \| SeedType = 2 NonSeedType = 1 \| \| --- \| --- \| --- \| --- \| --- \| --- \| | \|  \|  \|  \| [seedLoc = 55,60] Seq=AAACTTTCTTTCTTCGCTATCTCG \| ```          G        C   TTTCTT    A mRNA  3'  CTCTATCG TTC      TCAA  miRNA 5'  GAGGTAGT AGG      AGTT           T            TTGTAT                311121112512133333311115 ``` \| SeedType = 2 NonSeedType = -1 \| \| --- \| --- \| --- \| --- \| --- \| --- \| | \| 19 \| WBGene00000195:arr-1(160) \| ``` log2FC=-0.276 padj=0.648 ``` \| [seedLoc = 63,67] Seq=CATTCAACTTTTCTACGTCT \| ```          T  G    TT      TTAC    mRNA  3'  CT CATC  TTCAAC        miRNA 5'  GA GTAG  AGGTTG                 T  G    T       TATAGTT           31131111351211113333666 ``` \| SeedType = 3 NonSeedType = 1 \| \| --- \| --- \| --- \| --- \| --- \| --- \| | \| 20 \| WBGene00000223:atf-7(191) \| ``` log2FC=-0.131 padj=0.844 ``` \| [seedLoc = 535,540] Seq=TCAAAAAAGATGTTCTTACCTCT \| ```          T       CT G   AAAAAACT mRNA  3'  CTCCATT  T TAG         miRNA 5'  GAGGTAG  A GTT                  T       T  G   GTATAGTT           31111112351321233333333 ``` \| SeedType = 1 NonSeedType = -1 \| \| --- \| --- \| --- \| --- \| --- \| --- \| | \| 21 \| WBGene00000225:atgp-2(193) \| ``` log2FC=0.317 padj=0.443 ``` \| [seedLoc = 97,102] Seq=TTTCAATTTTTCTATGTTACCTCG \| ```          G      TG     TTTTAACTTT mRNA  3'  CTCCAT  TATCT           miRNA 5'  GAGGTA  GTAGG                    T             TTGTATAGTT           311111154211123333333333 ``` \| SeedType = 1 NonSeedType = -1 \| \| --- \| --- \| --- \| --- \| --- \| --- \| | \| 22 \| WBGene00000227:atm-1(195) \| ``` log2FC=-0.104 padj=0.932 ``` \| [seedLoc = 302,307] Seq=TGTTCAACATTTACATGTTTTTCCTACCTCT \| ```          T       CTTTTTGTACATTTA    T       mRNA  3'  CTCCATC               CAAC TGT    miRNA 5'  GAGGTAG               GTTG ATA             T       TAG                T   GTT           3111111133355545555555511113121666 ``` \| SeedType = 1 NonSeedType = 1 \| \| --- \| --- \| --- \| --- \| --- \| --- \| | \| 23 \| WBGene00000230:atp-3(200) \| ``` log2FC=0.126 padj=0.801 ``` \| [seedLoc = 62,67] Seq=ATCTGTGCTTCAAACCAGTTCCCTGCCTCA \| ```                  CCTTGA    ACTT      TA mRNA  3' ACTCCGTC      CCAA    CGTGTC   miRNA 5' TGAGGTAG      GGTT    GTATAG                    TA                  TT           111112113355451111555512121133 ``` \| SeedType = 2 NonSeedType = 1 \| \| --- \| --- \| --- \| --- \| --- \| --- \| | \| 24 \| WBGene00000231:atx-2(201) \| ``` log2FC=0.066 padj=0.946 ``` \| [seedLoc = 371,376] Seq=GTTCACACCGCCAATACCTCC \| ```          C      AA    C  CACTTG mRNA  3'  CTCCAT   CCG CA       miRNA 5'  GAGGTA   GGT GT                T      GTA   T  ATAGTT           3111111336112311333333 ``` \| SeedType = 1 NonSeedType = 2 \| \| --- \| --- \| --- \| --- \| --- \| --- \| | \|  \|  \|  \| [seedLoc = 592,598] Seq=TTTTTTTGAGCAATACCATCC \| ```          C  A    AA     GTTTTTTT mRNA  3'  CT CCAT    CGA         miRNA 5'  GA GGTA    GTT                  T       GTAG   GTATAGTT           31151111336612133333333 ``` \| SeedType = 3 NonSeedType = -1 \| \| --- \| --- \| --- \| --- \| --- \| --- \| | \| 25 \| WBGene00000254:bli-4(224) \| ``` log2FC=-0.151 padj=0.843 ``` \| [seedLoc = 317,322] Seq=CTTTTTTCAGGCTAAATCCTACTTCT \| ```          T       CTAA   G   TTTTTTC mRNA  3'  CTTCATC    ATC GAC        miRNA 5'  GAGGTAG    TAG TTG                 T              G   TATAGTT           31121111555511132113333333 ``` \| SeedType = 3 NonSeedType = -1 \| \| --- \| --- \| --- \| --- \| --- \| --- \| | \| 26 \| WBGene00000254:bli-4(226) \| ``` log2FC=-0.151 padj=0.843 ``` \| [seedLoc = 275,280] Seq=TAATTATTCATCGTCCTTACCTCT \| ```          T     T  C G T  T      T mRNA  3'  CTCCA TC T C AC TATTAA  miRNA 5'  GAGGT AG A G TG ATAGTT           T        T G T  T                  311111511313131131112115 ``` \| SeedType = 2 NonSeedType = -1 \| \| --- \| --- \| --- \| --- \| --- \| --- \| | \| 27 \| WBGene00000276:byn-1(248) \| ``` log2FC=-0.037 padj=0.98 ``` \| [seedLoc = 168,174] Seq=CATTGTTGGTTATGTTACTCTCT \| ```          T   T      ATTGGT     C mRNA  3'  CTC CATTGT      TGTTA  miRNA 5'  GAG GTAGTA      ATAGT           T          GGTTGT     T           31115111221333333121213 ``` \| SeedType = 2 NonSeedType = -1 \| \| --- \| --- \| --- \| --- \| --- \| --- \| | \| 28 \| WBGene00000277:cab-1(249) \| ``` log2FC=0.147 padj=0.859 ``` \| [seedLoc = 81,86] Seq=CGAATAATTATCAACTACCTTG \| ```          G        A   TTAATAAGC mRNA  3'  TTCCATCA CTA          miRNA 5'  GAGGTAGT GGT                   T        A   TGTATAGTT           3211111113121333333333 ``` \| SeedType = 3 NonSeedType = 2 \| \| --- \| --- \| --- \| --- \| --- \| --- \| | \| 29 \| WBGene00000283:cah-5(253) \| ``` log2FC=0.205 padj=0.693 ``` \| [seedLoc = 27,33] Seq=GGTTGTACATTTTCTATCCTCT \| ```          T    T       T       GG mRNA  3'  CTCC ATC TTT ACATGTT   miRNA 5'  GAGG TAG AGG TGTATAG            T        T   T       TT           31111511161223111121233 ``` \| SeedType = 2 NonSeedType = -1 \| \| --- \| --- \| --- \| --- \| --- \| --- \| | \| 30 \| WBGene00000293:cap-2(261) \| ``` log2FC=-0.022 padj=0.988 ``` \| [seedLoc = 324,330] Seq=TATTTTGCAATGTAAATTATCCTCT \| ```          T    T    AA G      T   T mRNA  3'  CTCC ATTA  T TAACGT TTA  miRNA 5'  GAGG TAGT  A GTTGTA AGT           T            G      T   T           3111151121551321112131213 ``` \| SeedType = 2 NonSeedType = 1 \| \| --- \| --- \| --- \| --- \| --- \| --- \| | \| 31 \| WBGene00000366:cbp-1(269) \| ``` log2FC=-0.093 padj=0.912 ``` \| [seedLoc = 146,152] Seq=TATCTCACAAATTACGTGTAATACACTCT \| ```          T   A   AATGTG    A    CTCTAT mRNA  3'  CTC CAT      CATT AACA       miRNA 5'  GAG GTA      GTAG TTGT                T                 G    ATAGTT           31115111555454111231111333333 ``` \| SeedType = 2 NonSeedType = 1 \| \| --- \| --- \| --- \| --- \| --- \| --- \| | \|  \|  \|  \| [seedLoc = 256,261] Seq=CCCTAATGCACCCAGTAAATACCTCA \| ```                 AAATGA  C    A   CC mRNA  3' ACTCCAT      CC ACGT ATC   miRNA 5' TGAGGTA      GG TGTA TAG                   GTA     T        TT           11111113335451131121511133 ``` \| SeedType = 1 NonSeedType = -1 \| \| --- \| --- \| --- \| --- \| --- \| --- \| | \|  \|  \|  \| [seedLoc = 671,677] Seq=TAACTCCGCCCCCAATACCATCT \| ```          T  A    AA   CC  CC    T mRNA  3'  CT CCAT   CC  CG  TCAA  miRNA 5'  GA GGTA   GG  GT  AGTT           T       GTA  TT  AT                311511113361133123311115 ``` \| SeedType = 3 NonSeedType = -1 \| \| --- \| --- \| --- \| --- \| --- \| --- \| | \| 32 \| WBGene00000368:ccb-1(271) \| ``` log2FC=0.169 padj=0.903 ``` \| [seedLoc = 83,88] Seq=TTCAATATCTCTATCTATCTCT \| ```          T       T    C     ACTT mRNA  3'  CTCTATC ATCT  TATA     miRNA 5'  GAGGTAG TAGG  GTAT              T            TT    AGTT           31112111511123621113333 ``` \| SeedType = 2 NonSeedType = -1 \| \| --- \| --- \| --- \| --- \| --- \| --- \| | \|  \|  \|  \| [seedLoc = 243,248] Seq=TTTTGTATTTTATATGTATACCTCT \| ```          T      ATG   ATTT      TT mRNA  3'  CTCCAT   TAT    TATGTT   miRNA 5'  GAGGTA   GTA    GTATAG            T            GGTT      TT           3111111554211333321121233 ``` \| SeedType = 1 NonSeedType = -1 \| \| --- \| --- \| --- \| --- \| --- \| --- \| | \| 33 \| WBGene00000376:ccr-4(278) \| ``` log2FC=0.036 padj=0.98 ``` \| [seedLoc = 75,80] Seq=CACCATTCTGTCTCATACCTCA \| ```                 AC   GTCTTACCAC mRNA  3' ACTCCAT  TCT           miRNA 5' TGAGGTA  AGG                           GT   TTGTATAGTT           1111111331123333333333 ``` \| SeedType = 1 NonSeedType = -1 \| \| --- \| --- \| --- \| --- \| --- \| --- \| | \|  \|  \|  \| [seedLoc = 929,935] Seq=GTCTATATTCCAATAACTACCTAC \| ```          CA       A     CTT     TG mRNA  3'   TCCATCA  TAAC   ATATC   miRNA 5'   AGGTAGT  GTTG   TATAG            TG       AG            TT           3311111113621115551111133 ``` \| SeedType = 3 NonSeedType = 1 \| \| --- \| --- \| --- \| --- \| --- \| --- \| | \| 34 \| WBGene00000383:cdc-14(291) \| ``` log2FC=-0.258 padj=0.393 ``` \| [seedLoc = 52,57] Seq=GAAAAGAAGAAAAACTGCCTCG \| ```          G        AAAAGAAGAAAAG mRNA  3'  CTCCGTCA              miRNA 5'  GAGGTAGT                       T        AGGTTGTATAGTT           3111121113333333333333 ``` \| SeedType = 2 NonSeedType = -1 \| \| --- \| --- \| --- \| --- \| --- \| --- \| | \| 35 \| WBGene00000403:casy-1(307) \| ``` log2FC=-0.087 padj=0.933 ``` \| [seedLoc = 70,76] Seq=TTTTATTCATTTTAGTCACCTCC \| ```          C     C G    T  T    TT mRNA  3'  CTCCA T ATTT AC TATT   miRNA 5'  GAGGT A TAGG TG ATAG            T       G    T  T    TT           31111151311223113111233 ``` \| SeedType = 2 NonSeedType = -1 \| \| --- \| --- \| --- \| --- \| --- \| --- \| | \|  \|  \|  \| [seedLoc = 243,249] Seq=TACTGATAATAGTGAATACTCTCC \| ```          C   T   AAGTGA          T mRNA  3'  CTC CAT      TAATA GTCA  miRNA 5'  GAG GTA      GTTGT TAGT           T       GTAG       A    T           3111511133334521121621113 ``` \| SeedType = 2 NonSeedType = -1 \| \| --- \| --- \| --- \| --- \| --- \| --- \| | \| 36 \| WBGene00000423:ced-9(326) \| ``` log2FC=-0.052 padj=0.98 ``` \| [seedLoc = 189,194] Seq=TTTGGCCAAACCCTACTTCC \| ```          C             ACCGGTTT mRNA  3'  CTTCATC  CCAA         miRNA 5'  GAGGTAG  GGTT                  T       TA    GTATAGTT           3112111166111133333333 ``` \| SeedType = 3 NonSeedType = 1 \| \| --- \| --- \| --- \| --- \| --- \| --- \| | \| 37 \| WBGene00000437:ceh-13(336) \| ``` log2FC=-0.075 padj=0.982 ``` \| [seedLoc = 37,42] Seq=ATTTTTATAATTTTTTTGTTACCTCT \| ```          T          TTTT      TTTTA mRNA  3'  CTCCATTGTT    TAATAT      miRNA 5'  GAGGTAGTAG    GTTGTA               T                    TAGTT           31111112212555521121133333 ``` \| SeedType = 1 NonSeedType = -1 \| \| --- \| --- \| --- \| --- \| --- \| --- \| | \| 38 \| WBGene00000439:ceh-16(337) \| ``` log2FC=-0.808 padj=1.0 ``` \| [seedLoc = 183,188] Seq=TTATTTGAAATCGAAATACCTCA \| ```                 AAAG    AGTTTATT mRNA  3' ACTCCAT    CTAA         miRNA 5' TGAGGTA    GGTT                         GTA     GTATAGTT           11111113334121133333333 ``` \| SeedType = 1 NonSeedType = 2 \| \| --- \| --- \| --- \| --- \| --- \| --- \| | \| 39 \| WBGene00000441:ceh-18(338) \| ``` log2FC=-0.259 padj=0.567 ``` \| [seedLoc = 58,63] Seq=TCTGTCCACATTTTCCTTATTCATTACCTCA \| ```                   CTTAT   TTTT   CC    T  mRNA  3' ACTCCATTA     TCC    ACA  TGTC   miRNA 5' TGAGGTAGT     AGG    TGT  ATAG                             T            TT           11111112155555111355511155121136 ``` \| SeedType = 1 NonSeedType = -1 \| \| --- \| --- \| --- \| --- \| --- \| --- \| | \| 40 \| WBGene00000443:ceh-20(340) \| ``` log2FC=0.062 padj=0.975 ``` \| [seedLoc = 50,56] Seq=AAATCTCAACCAATCATCCCATCTACCATCA \| ```             A     TACCCTACTAA     TCTAAA  mRNA  3' ACT CCATC           CCAAC        miRNA 5' TGA GGTAG           GGTTG                          TA              TATAGTT           11151111133555555555111113333336 ``` \| SeedType = 3 NonSeedType = 1 \| \| --- \| --- \| --- \| --- \| --- \| --- \| | \| 41 \| WBGene00000444:ceh-21(341) \| ``` log2FC=0.523 padj=0.904 ``` \| [seedLoc = 539,545] Seq=ACATTTGCACACCGGTGGTTATCCTCT \| ```          T    T   GGTGG   C    TTACA mRNA  3'  CTCC ATT     CCA ACGT      miRNA 5'  GAGG TAG     GGT TGTA               T        TA           TAGTT           311115112335441115112133333 ``` \| SeedType = 2 NonSeedType = 1 \| \| --- \| --- \| --- \| --- \| --- \| --- \| | \|  \|  \|  \| [seedLoc = 546,552] Seq=CGGTGGTTATCCTCTCACCTCT \| ```          T     C      TATTGGTGGC mRNA  3'  CTCCA TC TCC           miRNA 5'  GAGGT AG AGG                    T        T   TTGTATAGTT           31111151161113333333333 ``` \| SeedType = 2 NonSeedType = -1 \| \| --- \| --- \| --- \| --- \| --- \| --- \| | \|  \|  \|  \| [seedLoc = 686,692] Seq=AAATATGGCTCCTGTACTCTCG \| ```          G   T        TCG    AAA mRNA  3'  CTC CAT GTCC   GTAT    miRNA 5'  GAG GTA TAGG   TATA             T       G    TTG    GTT           31115111621113332111333 ``` \| SeedType = 2 NonSeedType = -1 \| \| --- \| --- \| --- \| --- \| --- \| --- \| | \| 42 \| WBGene00000460:ceh-39(354) \| ``` log2FC=0.488 padj=0.719 ``` \| [seedLoc = 361,366] Seq=CCCCATCCAATTCCCATACCTCT \| ```          T      ACCC     CTACCCC mRNA  3'  CTCCAT    TTAAC        miRNA 5'  GAGGTA    GGTTG                 T      GTA      TATAGTT           31111113335221113333333 ``` \| SeedType = 1 NonSeedType = 1 \| \| --- \| --- \| --- \| --- \| --- \| --- \| | \| 43 \| WBGene00000464:ceh-44(358) \| ``` log2FC=-0.116 padj=0.882 ``` \| [seedLoc = 202,206] Seq=GGCAGGCAGCTACCTAA \| ```          AA       A G    G      mRNA  3'   TCCATCG C GACG       miRNA 5'   AGGTAGT G TTGT                TG       A G    ATAGTT           3311111123132112366666 ``` \| SeedType = 3 NonSeedType = -1 \| \| --- \| --- \| --- \| --- \| --- \| --- \| | \| 44 \| WBGene00000469:ces-2(362) \| ``` log2FC=0.038 padj=1.0 ``` \| [seedLoc = 189,194] Seq=CTATACCACCCATTCATCTTGATCTATCTCT \| ```          T       TAGTTCTACTTAC   C         mRNA  3'  CTCTATC             CCA CATATC   miRNA 5'  GAGGTAG             GGT GTATAG            T       TA              T      TT           311121113345555555555111311111166 ``` \| SeedType = 2 NonSeedType = 1 \| \| --- \| --- \| --- \| --- \| --- \| --- \| | \|  \|  \|  \| [seedLoc = 213,218] Seq=AAGCTTTCATTCTCCCTACCTCA \| ```                  CC   T  TT     A mRNA  3' ACTCCATC  TCT AC   TCGA  miRNA 5' TGAGGTAG  AGG TG   AGTT                   T    T  TAT                111111113511231133611215 ``` \| SeedType = 1 NonSeedType = -1 \| \| --- \| --- \| --- \| --- \| --- \| --- \| | \| 45 \| WBGene00000472:cey-1(363) \| ``` log2FC=-0.078 padj=0.953 ``` \| [seedLoc = 517,522] Seq=TTGAGAGAATCTTTAATGTTCAAGTACCTCT \| ```          T      G ACTTGTAATT     G GAGTT  mRNA  3'  CTCCAT A          TCTAA A       miRNA 5'  GAGGTA T          AGGTT T                T      G                G ATAGTT           31111113155554555551121131333336 ``` \| SeedType = 1 NonSeedType = 2 \| \| --- \| --- \| --- \| --- \| --- \| --- \| | \| 46 \| WBGene00000478:cfz-2(368) \| ``` log2FC=-0.541 padj=1.0 ``` \| [seedLoc = 43,48] Seq=ATTTGTGATCCCCGCACATCTACCTCT \| ```          T       TACACG  CCTA     TA mRNA  3'  CTCCATC      CC    GTGTT   miRNA 5'  GAGGTAG      GG    TATAG            T       TA      TTG      TT           311111113355541133352121233 ``` \| SeedType = 1 NonSeedType = -1 \| \| --- \| --- \| --- \| --- \| --- \| --- \| | \|  \|  \|  \| [seedLoc = 110,115] Seq=TTTCTCATGAGCTTTTACCTCT \| ```          T       T  G    CTCTTT mRNA  3'  CTCCATT TC AGTA       miRNA 5'  GAGGTAG AG TTGT                T       T  G    ATAGTT           3111111231131221333333 ``` \| SeedType = 1 NonSeedType = -1 \| \| --- \| --- \| --- \| --- \| --- \| --- \| | \| 47 \| WBGene00000479:cgh-1(369) \| ``` log2FC=0.004 padj=0.997 ``` \| [seedLoc = 527,532] Seq=AATCTTGTAATTTCCTATCTCA \| ```                  C      GTTCTAA mRNA  3' ACTCTATC TTTAAT        miRNA 5' TGAGGTAG AGGTTG                         T      TATAGTT           1111211131221123333333 ``` \| SeedType = 2 NonSeedType = 2 \| \| --- \| --- \| --- \| --- \| --- \| --- \| | \| 48 \| WBGene00000480:cgp-1(370) \| ``` log2FC=-0.573 padj=3.78817418824714E-4 ``` \| [seedLoc = 184,189] Seq=CCATCAATCACCACTATCTCC \| ```          C            CTAACTACC mRNA  3'  CTCTATCA CCA          miRNA 5'  GAGGTAGT GGT                   T        A   TGTATAGTT           3111211116111333333333 ``` \| SeedType = 2 NonSeedType = 1 \| \| --- \| --- \| --- \| --- \| --- \| --- \| | \| 49 \| WBGene00000496:chs-1(377) \| ``` log2FC=0.68 padj=0.137 ``` \| [seedLoc = 66,71] Seq=TATTATTGCATGCTCTTACCTCA \| ```                  C  GT   T     T mRNA  3' ACTCCATT TC  ACG TATTA  miRNA 5' TGAGGTAG AG  TGT ATAGT                   T  GT         T           11111112311331125111213 ``` \| SeedType = 1 NonSeedType = -1 \| \| --- \| --- \| --- \| --- \| --- \| --- \| | \|  \|  \|  \| [seedLoc = 72,76] Seq=TATTATTGCATGCTCTTACCTCAT \| ```          T        C  GT   T     T mRNA  3'  ACTCCATT TC  ACG TATTA  miRNA 5'  TGAGGTAG AG  TGT ATAGT                    T  GT         T           511111112311331125111213 ``` \| SeedType = 1 NonSeedType = -1 \| \| --- \| --- \| --- \| --- \| --- \| --- \| | \| 50 \| WBGene00000498:chk-1(380) \| ``` log2FC=0.054 padj=0.973 ``` \| [seedLoc = 193,198] Seq=AGAGTTTGTCCCCACTGCCTCA \| ```                   C  CTGTTTGAGA mRNA  3' ACTCCGTCA CC           miRNA 5' TGAGGTAGT GG                             A  TTGTATAGTT           1111121113113333333333 ``` \| SeedType = 2 NonSeedType = -1 \| \| --- \| --- \| --- \| --- \| --- \| --- \| | \|  \|  \|  \| [seedLoc = 208,211] Seq=TTTGTCCCCACTGCCTCAT \| ```          T         C  CTGTTT     mRNA  3'  ACTCCGTCA CC           miRNA 5'  TGAGGTAGT GG                              A  TTGTATAGTT           51111121113113333336666 ``` \| SeedType = 2 NonSeedType = -1 \| \| --- \| --- \| --- \| --- \| --- \| --- \| | \| 51 \| WBGene00000530:clh-3(409) \| ``` log2FC=0.081 padj=0.973 ``` \| [seedLoc = 153,157] Seq=TTCTTCTTCTCCTCTTCCTCAT \| ```          T     T      TCTTCTTCTT mRNA  3'  ACTCC TC TCC           miRNA 5'  TGAGG AG AGG                          T  T   TTGTATAGTT           51111131161113333333333 ``` \| SeedType = 2 NonSeedType = -1 \| \| --- \| --- \| --- \| --- \| --- \| --- \| | \| 52 \| WBGene00000537:clk-2(416) \| ``` log2FC=-0.213 padj=0.758 ``` \| [seedLoc = 76,81] Seq=CACTTTGATCTCTACCTCG \| ```          G              T     C mRNA  3'  CTCCATC TCTAGT   TCA  miRNA 5'  GAGGTAG AGGTTG   AGT           T       T      TAT   T           3111111161121223661113 ``` \| SeedType = 1 NonSeedType = 2 \| \| --- \| --- \| --- \| --- \| --- \| --- \| | \|  \|  \|  \| [seedLoc = 86,90] Seq=CACTTTGATCTCTACCTCGT \| ```          T               T     C mRNA  3'  GCTCCATC TCTAGT   TCA  miRNA 5'  TGAGGTAG AGGTTG   AGT                    T      TAT   T           52111111161121223661113 ``` \| SeedType = 1 NonSeedType = 2 \| \| --- \| --- \| --- \| --- \| --- \| --- \| | \| 53 \| WBGene00000548:clr-1(428) \| ``` log2FC=0.194 padj=0.732 ``` \| [seedLoc = 238,244] Seq=TACTAGTCAATAGTCTATTCTACACTCC \| ```          C   A    TT   TGA    TG    T mRNA  3'  CTC CATC  ATC   TAAC  ATCA  miRNA 5'  GAG GTAG  TAG   GTTG  TAGT           T                    TA    T           3111511115511154521113311113 ``` \| SeedType = 2 NonSeedType = 1 \| \| --- \| --- \| --- \| --- \| --- \| --- \| | \| 54 \| WBGene00000550:clu-1(431) \| ``` log2FC=0.218 padj=0.448 ``` \| [seedLoc = 484,490] Seq=CTCTGATAATCCGAGTACACTCC \| ```          C   A   G GC          TC mRNA  3'  CTC CAT A  CTAATA GTC   miRNA 5'  GAG GTA T  GGTTGT TAG            T       G A       A   TT           311151113135121121621133 ``` \| SeedType = 2 NonSeedType = 2 \| \| --- \| --- \| --- \| --- \| --- \| --- \| | \| 55 \| WBGene00000552:cmd-1(433) \| ``` log2FC=-0.064 padj=0.955 ``` \| [seedLoc = 391,396] Seq=CGGCTTGACCCTTCCTCAA \| ```          A     T  CCA               mRNA  3'  ACTCC TC   GTTCGGC        miRNA 5'  TGAGG AG   TAGGTTG                       T            TATAGTT           51111131155521212216666666 ``` \| SeedType = 2 NonSeedType = 2 \| \| --- \| --- \| --- \| --- \| --- \| --- \| | \| 56 \| WBGene00000566:cnt-2(445) \| ``` log2FC=-0.137 padj=0.851 ``` \| [seedLoc = 108,113] Seq=TCTCATCAGATCTCTGCCTCG \| ```          G             ACTACTCT mRNA  3'  CTCCGTC TCTAG         miRNA 5'  GAGGTAG AGGTT                  T       T     GTATAGTT           3111121161121233333333 ``` \| SeedType = 2 NonSeedType = 2 \| \| --- \| --- \| --- \| --- \| --- \| --- \| | \| 57 \| WBGene00000591:coh-1(450) \| ``` log2FC=0.025 padj=0.993 ``` \| [seedLoc = 144,149] Seq=TATCTTATTTCTTATTTCTACCTCC \| ```          C       TTT    TT   TCTAT mRNA  3'  CTCCATC   ATTC  TAT      miRNA 5'  GAGGTAG   TAGG  GTA               T              TT   TAGTT           3111111155511213321133333 ``` \| SeedType = 1 NonSeedType = -1 \| \| --- \| --- \| --- \| --- \| --- \| --- \| | \| 58 \| WBGene00000666:col-91(497) \| ``` log2FC=-0.638 padj=0.021 ``` \| [seedLoc = 55,60] Seq=CAACTCTCTTTTCTTTACCTCA \| ```                      TTTCTC    C mRNA  3' ACTCCATT TCT      TCAA  miRNA 5' TGAGGTAG AGG      AGTT                   T   TTGTAT                11111112611233333311115 ``` \| SeedType = 1 NonSeedType = -1 \| \| --- \| --- \| --- \| --- \| --- \| --- \| | \| 59 \| WBGene00000668:col-93(499) \| ``` log2FC=-0.165 padj=0.956 ``` \| [seedLoc = 23,29] Seq=AAGATTTAGTTCTTTACCGTCT \| ```          T  G         TGATTTAGAA mRNA  3'  CT CCATT TCT           miRNA 5'  GA GGTAG AGG                    T        T   TTGTATAGTT           31141111261123333333333 ``` \| SeedType = 3 NonSeedType = -1 \| \| --- \| --- \| --- \| --- \| --- \| --- \| | \| 60 \| WBGene00000698:col-124(522) \| ``` log2FC=0.256 padj=0.945 ``` \| [seedLoc = 146,151] Seq=TCCTCTCCAACTTTTTTACCTCC \| ```          C       TT      CTCTCCT mRNA  3'  CTCCATT  TTCAAC        miRNA 5'  GAGGTAG  AGGTTG                 T       T       TATAGTT           31111112351211113333333 ``` \| SeedType = 1 NonSeedType = 1 \| \| --- \| --- \| --- \| --- \| --- \| --- \| | \|  \|  \|  \| [seedLoc = 225,231] Seq=GAATTATAATATAAATACCATCA \| ```             A    AAA   TA       G mRNA  3' ACT CCAT    TA  ATATTAA  miRNA 5' TGA GGTA    GT  TATAGTT                   GTAG  TG                   111511113336213311112114 ``` \| SeedType = 3 NonSeedType = -1 \| \| --- \| --- \| --- \| --- \| --- \| --- \| | \| 61 \| WBGene00000740:col-167(554) \| ``` log2FC=-0.083 padj=0.984 ``` \| [seedLoc = 52,57] Seq=GCCTGTCTTTCTAATTACCTCA \| ```                  A    TT      CG mRNA  3' ACTCCATT ATCT  C TGTC   miRNA 5' TGAGGTAG TAGG  G ATAG                         TT T    TT           11111112511123316121133 ``` \| SeedType = 1 NonSeedType = -1 \| \| --- \| --- \| --- \| --- \| --- \| --- \| | \| 62 \| WBGene00000743:col-170(557) \| ``` log2FC=-0.109 padj=0.98 ``` \| [seedLoc = 54,59] Seq=ATTTTTGCAAGTCTTTCACATCATACCTCT \| ```          T      ACTACACTT   G     TTTTA mRNA  3'  CTCCAT         TCT AACGT      miRNA 5'  GAGGTA         AGG TTGTA               T      GT                TAGTT           311111133555555511241112133333 ``` \| SeedType = 1 NonSeedType = 1 \| \| --- \| --- \| --- \| --- \| --- \| --- \| | \| 63 \| WBGene00000746:col-173(559) \| ``` log2FC=-0.332 padj=0.456 ``` \| [seedLoc = 140,145] Seq=TTATAAATAAGTTGTTATTTCTACCTCC \| ```          C       TTTATT   G    AATATT mRNA  3'  CTCCATC      GTT AATA       miRNA 5'  GAGGTAG      TAG TTGT                T                G    ATAGTT           3111111155555521231121333333 ``` \| SeedType = 1 NonSeedType = -1 \| \| --- \| --- \| --- \| --- \| --- \| --- \| | \| 64 \| WBGene00000772:cpb-3(579) \| ``` log2FC=-0.209 padj=0.519 ``` \| [seedLoc = 370,375] Seq=AAACTCTCCGCATTGCGTTTTACCTCC \| ```          C       T G  TT   CCTC    A mRNA  3'  CTCCATT T CG  ACG    TCAA  miRNA 5'  GAGGTAG A GT  TGT    AGTT           T       T G       AT                  311111123131255112335511115 ``` \| SeedType = 1 NonSeedType = -1 \| \| --- \| --- \| --- \| --- \| --- \| --- \| | \| 65 \| WBGene00000776:cpl-1(583) \| ``` log2FC=0.037 padj=0.982 ``` \| [seedLoc = 86,91] Seq=TTCTGTCAAGCTTTACCTCG \| ```          G          G        TT mRNA  3'  CTCCATT TC AAC TGTC   miRNA 5'  GAGGTAG AG TTG ATAG            T       T  G   T    TT           3111111261131116121133 ``` \| SeedType = 1 NonSeedType = 1 \| \| --- \| --- \| --- \| --- \| --- \| --- \| | \| 66 \| WBGene00000781:cpr-1(588) \| ``` log2FC=-0.287 padj=0.507 ``` \| [seedLoc = 27,33] Seq=ATGTACACAATTTTTGTAATTACCATCT \| ```          T  A      ATGTT       CATGTA mRNA  3'  CT CCATTA     TTTAACA       miRNA 5'  GA GGTAGT     AGGTTGT                T                     ATAGTT           3115111121554551221111333333 ``` \| SeedType = 3 NonSeedType = 1 \| \| --- \| --- \| --- \| --- \| --- \| --- \| | \| 67 \| WBGene00000792:crb-1(596) \| ``` log2FC=-0.73 padj=0.304 ``` \| [seedLoc = 57,63] Seq=GCCCCCCCCCCCCACTCACCTCT \| ```          T     C   C  CCCCCCCCCG mRNA  3'  CTCCA TCA CC           miRNA 5'  GAGGT AGT GG                    T         A  TTGTATAGTT           31111151113113333333333 ``` \| SeedType = 2 NonSeedType = -1 \| \| --- \| --- \| --- \| --- \| --- \| --- \| | \| 68 \| WBGene00000799:crn-6(603) \| ``` log2FC=-0.043 padj=0.984 ``` \| [seedLoc = 91,96] Seq=CATCCCCTAAAATACTACCTCT \| ```          T         AA   CCCCTAC mRNA  3'  CTCCATCAT  AAT        miRNA 5'  GAGGTAGTA  TTG                 T         GG   TATAGTT           3111111111331123333333 ``` \| SeedType = 1 NonSeedType = -1 \| \| --- \| --- \| --- \| --- \| --- \| --- \| | \| 69 \| WBGene00000854:cux-7(638) \| ``` log2FC=-0.103 padj=0.945 ``` \| [seedLoc = 103,108] Seq=TCTCAATAAACGTATTTACCTCT \| ```          T     T    G   ATAACTCT mRNA  3'  CTCCA TTAT CAA         miRNA 5'  GAGGT AGTA GTT                  T          G   GTATAGTT           31111151211311133333333 ``` \| SeedType = 2 NonSeedType = 1 \| \| --- \| --- \| --- \| --- \| --- \| --- \| | \| 70 \| WBGene00000866:cyb-2.1(647) \| ``` log2FC=-0.032 padj=0.993 ``` \| [seedLoc = 158,163] Seq=TTTTTCGCACCTTCTTTTGTATCTATCTCA \| ```                  TATGTTTTCT       CTTTTT mRNA  3' ACTCTATC          TCC ACG       miRNA 5' TGAGGTAG          AGG TGT                        T            T   ATAGTT           1111211135545555551116112333333 ``` \| SeedType = 2 NonSeedType = -1 \| \| --- \| --- \| --- \| --- \| --- \| --- \| | \| 71 \| WBGene00000869:cyc-1(650) \| ``` log2FC=0.048 padj=0.959 ``` \| [seedLoc = 26,32] Seq=CGCTATTTTTTCATTATCCTCT \| ```          T    T       TTTT     C mRNA  3'  CTCC ATTA CT    TATCG  miRNA 5'  GAGG TAGT GG    ATAGT           T         A  TTGT     T           31111511216123333111123 ``` \| SeedType = 2 NonSeedType = -1 \| \| --- \| --- \| --- \| --- \| --- \| --- \| | \| 72 \| WBGene00000871:cye-1(652) \| ``` log2FC=-0.071 padj=0.955 ``` \| [seedLoc = 444,450] Seq=TACTGTAACTGGTACATATCCTCT \| ```          T    T  A   GGTCA      T mRNA  3'  CTCC AT CAT     ATGTCA  miRNA 5'  GAGG TA GTA     TATAGT           T           GGTTG      T           311115115111333331121113 ``` \| SeedType = 2 NonSeedType = -1 \| \| --- \| --- \| --- \| --- \| --- \| --- \| | \| 73 \| WBGene00000884:cyn-8(664) \| ``` log2FC=0.02 padj=0.994 ``` \| [seedLoc = 271,276] Seq=TCCCATATTCCTATCCATCTATCTCA \| ```                  TACCT    T     CCCT mRNA  3' ACTCTATC     ATCC  TATA     miRNA 5' TGAGGTAG     TAGG  GTAT                               TT    AGTT           111121115555511113621113333 ``` \| SeedType = 2 NonSeedType = -1 \| \| --- \| --- \| --- \| --- \| --- \| --- \| | \| 74 \| WBGene00000894:dab-1(675) \| ``` log2FC=-0.364 padj=0.072 ``` \| [seedLoc = 446,452] Seq=CAACTTGTCAATCTCTCACCTCC \| ```          C     C         TGT    C mRNA  3'  CTCCA TC TCTAAC   TCAA  miRNA 5'  GAGGT AG AGGTTG   AGTT           T        T      TAT                311111511611211133311115 ``` \| SeedType = 2 NonSeedType = 1 \| \| --- \| --- \| --- \| --- \| --- \| --- \| | \| 75 \| WBGene00000897:daf-1(678) \| ``` log2FC=-0.015 padj=0.995 ``` \| [seedLoc = 231,236] Seq=AGACTCCTCACCGTGTACCTCA \| ```                 GTG     TCC    A mRNA  3' ACTCCAT   CC AC   TCAG  miRNA 5' TGAGGTA   GG TG   AGTT                  GTA  T  TAT                11111113331161133311125 ``` \| SeedType = 1 NonSeedType = -1 \| \| --- \| --- \| --- \| --- \| --- \| --- \| | \|  \|  \|  \| [seedLoc = 285,291] Seq=TACTGCATAAGCTTTATTACCCTCC \| ```          C  C      TT  G    C    T mRNA  3'  CT CCATTA  TC AATA GTCA  miRNA 5'  GA GGTAGT  AG TTGT TAGT           T             G    A    T           3115111121551131121321113 ``` \| SeedType = 3 NonSeedType = -1 \| \| --- \| --- \| --- \| --- \| --- \| --- \| | \| 76 \| WBGene00000899:daf-3(680) \| ``` log2FC=-0.381 padj=0.345 ``` \| [seedLoc = 220,225] Seq=AGAAGGACAATGTATCCCATACCTCA \| ```                 ACCC   G     GGAAGA mRNA  3' ACTCCAT    TAT TAACA       miRNA 5' TGAGGTA    GTA GTTGT                              G     ATAGTT           11111115555211321111333333 ``` \| SeedType = 1 NonSeedType = 1 \| \| --- \| --- \| --- \| --- \| --- \| --- \| | \| 77 \| WBGene00000900:daf-4(681) \| ``` log2FC=0.282 padj=0.781 ``` \| [seedLoc = 311,316] Seq=CAACTAACTCAAAAACTTACCTCA \| ```                  CAAA   TCA     C mRNA  3' ACTCCATT    AAC   ATCAA  miRNA 5' TGAGGTAG    TTG   TAGTT                   TAGG   TA                  111111123333111335111115 ``` \| SeedType = 1 NonSeedType = 1 \| \| --- \| --- \| --- \| --- \| --- \| --- \| | \| 78 \| WBGene00000901:daf-5(683) \| ``` log2FC=-0.253 padj=0.649 ``` \| [seedLoc = 158,163] Seq=TTTTTAAATTCATTTTACCTCT \| ```          T       TTACTTAAATTTTT mRNA  3'  CTCCATT               miRNA 5'  GAGGTAG                        T       TAGGTTGTATAGTT           3111111233333333333333 ``` \| SeedType = 1 NonSeedType = -1 \| \| --- \| --- \| --- \| --- \| --- \| --- \| | \|  \|  \|  \| [seedLoc = 226,231] Seq=CACTTGCCATACTATTAGGTACCTCT \| ```          T      GGAT    AT  CGT   C mRNA  3'  CTCCAT    TATC  AC   TCA  miRNA 5'  GAGGTA    GTAG  TG   AGT           T              GT  TAT   T           31111114455211133113331113 ``` \| SeedType = 1 NonSeedType = -1 \| \| --- \| --- \| --- \| --- \| --- \| --- \| | \| 79 \| WBGene00000903:daf-7(685) \| ``` log2FC=0.079 padj=0.981 ``` \| [seedLoc = 296,301] Seq=TCGGATCCTTCCACTACCTCC \| ```          C           TT CTAGGCT mRNA  3'  CTCCATCA CC  C        miRNA 5'  GAGGTAGT GG  G                 T        A  TT TATAGTT           3111111116113313333333 ``` \| SeedType = 1 NonSeedType = -1 \| \| --- \| --- \| --- \| --- \| --- \| --- \| | \| 80 \| WBGene00000908:daf-12(690) \| ``` log2FC=1.512 padj=2.32724916739602E-10 over-expressed ``` \| [seedLoc = 41,46] Seq=AAACGATGCCATGCCTTCAATACCTCA \| ```                 AACT    TAC    G   A mRNA  3' ACTCCAT    TCCG   CGTA CAA  miRNA 5' TGAGGTA    AGGT   GTAT GTT                  GT      T      A               111111133551112355121131115 ``` \| SeedType = 1 NonSeedType = 2 \| \| --- \| --- \| --- \| --- \| --- \| --- \| | \|  \|  \|  \| [seedLoc = 157,162] Seq=AATCGTATAATATTCCTCTACCTCT \| ```          T           TTAT      CTAA mRNA  3'  CTCCATC TCC    AATATG     miRNA 5'  GAGGTAG AGG    TTGTAT              T       T             AGTT           31111111611155551121123333 ``` \| SeedType = 1 NonSeedType = -1 \| \| --- \| --- \| --- \| --- \| --- \| --- \| | \|  \|  \|  \| [seedLoc = 405,410] Seq=CCGGTTTTCATGTTCTACTTCT \| ```          T       T G  CTTTTGGCC mRNA  3'  CTTCATC T TA          miRNA 5'  GAGGTAG A GT                   T       T G  TGTATAGTT           3112111131321333333333 ``` \| SeedType = 3 NonSeedType = -1 \| \| --- \| --- \| --- \| --- \| --- \| --- \| | \|  \|  \|  \| [seedLoc = 541,546] Seq=TGATTCATGCAGCCTAGCCCCTACCTCT \| ```          T       CCCG          C    T mRNA  3'  CTCCATC    ATCCGACGTA TTAG  miRNA 5'  GAGGTAG    TAGGTTGTAT AGTT           T                                      3111111155541111211211512125 ``` \| SeedType = 1 NonSeedType = 2 \| \| --- \| --- \| --- \| --- \| --- \| --- \| | \|  \|  \|  \| [seedLoc = 563,568] Seq=TAATTCCGTCAATACTATCTCA \| ```                    AACTGCCTTAAT mRNA  3' ACTCTATCAT             miRNA 5' TGAGGTAGTA                                GGTTGTATAGTT           1111211111333333333333 ``` \| SeedType = 2 NonSeedType = -1 \| \| --- \| --- \| --- \| --- \| --- \| --- \| | \|  \|  \|  \| [seedLoc = 1035,1040] Seq=CTCTATTTTCCTTGCACTACCTCG \| ```          G        CGT   TTT     TC mRNA  3'  CTCCATCA   TCC    TATC   miRNA 5'  GAGGTAGT   AGG    ATAG            T              TTGT    TT           3111111115451113336111133 ``` \| SeedType = 1 NonSeedType = -1 \| \| --- \| --- \| --- \| --- \| --- \| --- \| | \| 81 \| WBGene00000911:daf-15(692) \| ``` log2FC=0.007 padj=0.996 ``` \| [seedLoc = 136,141] Seq=TTCTTTTCGTTTTCCTACTTCA \| ```                  C   T  TTT  TT mRNA  3' ACTTCATC TTT GC   TC   miRNA 5' TGAGGTAG AGG TG   AG                    T   T  TAT  TT           1112111131223213331133 ``` \| SeedType = 3 NonSeedType = -1 \| \| --- \| --- \| --- \| --- \| --- \| --- \| | \|  \|  \|  \| [seedLoc = 280,286] Seq=AAACTTCTTCTAACTACCTGC \| ```          CG       A  TT T      A mRNA  3'   TCCATCA TC  C   TCAA  miRNA 5'   AGGTAGT GG  G   AGTT           TG       A  TT TAT                33111111132133136611115 ``` \| SeedType = 3 NonSeedType = -1 \| \| --- \| --- \| --- \| --- \| --- \| --- \| | \| 82 \| WBGene00000912:daf-16(693) \| ``` log2FC=-0.131 padj=0.802 ``` \| [seedLoc = 95,101] Seq=ATCTTTTCAAATCCCTACCTAC \| ```          CA      CCTA   TTT  TA mRNA  3'   TCCATC    AAC   TC   miRNA 5'   AGGTAG    TTG   AG            TG      TAGG   TAT  TT           3311111133331113331133 ``` \| SeedType = 3 NonSeedType = 1 \| \| --- \| --- \| --- \| --- \| --- \| --- \| | \|  \|  \|  \| [seedLoc = 278,283] Seq=AATAATATAAAATACCTCT \| ```          T      AA         ATAA mRNA  3'  CTCCAT     AATATA     miRNA 5'  GAGGTA     TTGTAT              T      GTAGG      AGTT           3111111336661121113333 ``` \| SeedType = 1 NonSeedType = -1 \| \| --- \| --- \| --- \| --- \| --- \| --- \| | \|  \|  \|  \| [seedLoc = 332,337] Seq=TATCGAAAAACCGTTGATTATTACCTCT \| ```          T         TAGTTG    AAAGCTAT mRNA  3'  CTCCATTAT      CCAA         miRNA 5'  GAGGTAGTA      GGTT                  T                   GTATAGTT           3111111211554554111133333333 ``` \| SeedType = 1 NonSeedType = 1 \| \| --- \| --- \| --- \| --- \| --- \| --- \| | \|  \|  \|  \| [seedLoc = 402,407] Seq=CACTCTTTAAATGCTACCTCT \| ```          T         A    TTC   C mRNA  3'  CTCCATCGT  AAT   TCA  miRNA 5'  GAGGTAGTA  TTG   AGT           T         GG   TAT   T           3111111121361123331113 ``` \| SeedType = 1 NonSeedType = -1 \| \| --- \| --- \| --- \| --- \| --- \| --- \| | \|  \|  \|  \| [seedLoc = 649,654] Seq=CGTATACACACACACATAGTAATCTACCTCC \| ```          C       TAATGATACACA  C      GC  mRNA  3'  CTCCATC            CA ACATAT    miRNA 5'  GAGGTAG            GT TGTATA             T       TAG                  GTT           31111111333545555555115111111336 ``` \| SeedType = 1 NonSeedType = -1 \| \| --- \| --- \| --- \| --- \| --- \| --- \| | \|  \|  \|  \| [seedLoc = 952,957] Seq=TGCTTGTTTTTTTCTCTACCTCT \| ```          T           TTTTTTGT   T mRNA  3'  CTCCATC TCT        TCG  miRNA 5'  GAGGTAG AGG        AGT           T       T   TTGTAT     T           311111116112333333451123 ``` \| SeedType = 1 NonSeedType = -1 \| \| --- \| --- \| --- \| --- \| --- \| --- \| | \| 83 \| WBGene00000936:dbl-1(708) \| ``` log2FC=0.624 padj=0.524 ``` \| [seedLoc = 321,326] Seq=CTTCTTCCAAAAAATCTATCTCT \| ```          T       TAAAA   CTTCTTC mRNA  3'  CTCTATC     AAC        miRNA 5'  GAGGTAG     TTG                 T       TAGG    TATAGTT           31112111333351113333333 ``` \| SeedType = 2 NonSeedType = 1 \| \| --- \| --- \| --- \| --- \| --- \| --- \| | \| 84 \| WBGene00000961:dgn-1(722) \| ``` log2FC=-0.108 padj=0.927 ``` \| [seedLoc = 60,65] Seq=TCCTGTTATTATCCTCTACCTTT \| ```          T           TATTAT    CT mRNA  3'  TTCCATC TCC      TGTC   miRNA 5'  GAGGTAG AGG      ATAG            T       T   TTGT      TT           321111116111333355121133 ``` \| SeedType = 3 NonSeedType = -1 \| \| --- \| --- \| --- \| --- \| --- \| --- \| | \|  \|  \|  \| [seedLoc = 275,280] Seq=TACTTCAAATCTTTCCCTATCTCT \| ```          T       CCTT     ACT    T mRNA  3'  CTCTATC    TCTAA    TCA  miRNA 5'  GAGGTAG    AGGTT    AGT           T       T        GTAT   T           3111211135551121133361113 ``` \| SeedType = 2 NonSeedType = 2 \| \| --- \| --- \| --- \| --- \| --- \| --- \| | \| 85 \| WBGene00000995:die-1(756) \| ``` log2FC=0.077 padj=0.966 ``` \| [seedLoc = 45,50] Seq=TGCTCTCCCACCATCATAGTATCTACCTTT \| ```          T       TATGATACTA   C CTC   T mRNA  3'  TTCCATC          CCA C   TCG  miRNA 5'  GAGGTAG          GGT G   AGT           T       TA           T TAT   T           321111113354555555111313331123 ``` \| SeedType = 3 NonSeedType = 1 \| \| --- \| --- \| --- \| --- \| --- \| --- \| | \|  \|  \|  \| [seedLoc = 536,541] Seq=AAATTAGGTTATTTTATACCTCC \| ```          C      AT    TTGG     A mRNA  3'  CTCCAT  TTTA    ATTAA  miRNA 5'  GAGGTA  AGGT    TAGTT           T      GT    TGTA                 31111113312213333112115 ``` \| SeedType = 1 NonSeedType = 2 \| \| --- \| --- \| --- \| --- \| --- \| --- \| | \|  \|  \|  \| [seedLoc = 570,576] Seq=ATTCCCAAAACATGCTACCTAC \| ```          CA        A   AACCCTTA mRNA  3'   TCCATCGT CAA         miRNA 5'   AGGTAGTA GTT                  TG        G   GTATAGTT           3311111121311133333333 ``` \| SeedType = 3 NonSeedType = 1 \| \| --- \| --- \| --- \| --- \| --- \| --- \| | \| 86 \| WBGene00001008:dlk-1(765) \| ``` log2FC=0.257 padj=0.578 ``` \| [seedLoc = 47,51] Seq=TTTTGTAGTATCCCTTCCTCC \| ```          C    T  C    TG     TT mRNA  3'  CTCC TC  CTA  ATGTT   miRNA 5'  GAGG AG  GGT  TATAG            T    T  TA   TG     TT           3111131136121331121233 ``` \| SeedType = 2 NonSeedType = 2 \| \| --- \| --- \| --- \| --- \| --- \| --- \| | \| 87 \| WBGene00001020:dnj-2(771) \| ``` log2FC=0.231 padj=0.581 ``` \| [seedLoc = 146,151] Seq=GAGTTTGCGAGCATAATCTACTTCT \| ```          T       TAATA G     TTGAG mRNA  3'  CTTCATC     C AGCGT      miRNA 5'  GAGGTAG     G TTGTA               T       TA    G     TAGTT           3112111133555131212133333 ``` \| SeedType = 3 NonSeedType = -1 \| \| --- \| --- \| --- \| --- \| --- \| --- \| | \| 88 \| WBGene00001025:dnj-7(776) \| ``` log2FC=0.176 padj=0.752 ``` \| [seedLoc = 278,283] Seq=GAATGTATCTTCCCGTTTCCTACCTTT \| ```          T       CTTTG  CTTC     AAG mRNA  3'  TTCCATC     CC    TATGT    miRNA 5'  GAGGTAG     GG    GTATA             T       TA     TT       GTT           321111113355411335521121333 ``` \| SeedType = 3 NonSeedType = -1 \| \| --- \| --- \| --- \| --- \| --- \| --- \| | \| 89 \| WBGene00001026:dnj-8(777) \| ``` log2FC=0.333 padj=0.311 ``` \| [seedLoc = 43,48] Seq=CGTTCATCAATATGTCTACCTTC \| ```          C       TGTA    TACTTGC mRNA  3'  TTCCATC    TAAC        miRNA 5'  GAGGTAG    GTTG                 T       TAG     TATAGTT           32111111333521113333333 ``` \| SeedType = 3 NonSeedType = 1 \| \| --- \| --- \| --- \| --- \| --- \| --- \| | \| 90 \| WBGene00001038:dnj-20(789) \| ``` log2FC=0.099 padj=0.902 ``` \| [seedLoc = 222,226] Seq=GCTGATCTCTACGTCT \| ```          T  G          TCG      mRNA  3'  CT CATC TCTAG         miRNA 5'  GA GTAG AGGTT                  T  G    T     GTATAGTT           3113111161121233366666 ``` \| SeedType = 3 NonSeedType = 2 \| \| --- \| --- \| --- \| --- \| --- \| --- \| | \| 91 \| WBGene00001061:dpl-1(817) \| ``` log2FC=0.037 padj=0.98 ``` \| [seedLoc = 217,222] Seq=GAATTGACGACAGATTTCATAATCTACCTTT \| ```          T       TAATACTTTAGA           G mRNA  3'  TTCCATC            CAGCA GTTAA  miRNA 5'  GAGGTAG            GTTGT TAGTT           T       TAG              A                 32111111333555555545112116212114 ``` \| SeedType = 3 NonSeedType = -1 \| \| --- \| --- \| --- \| --- \| --- \| --- \| | \|  \|  \|  \| [seedLoc = 320,326] Seq=TAACTCAATGCTCCCGATTTTACCATCA \| ```             A     TTAG  CT    AC    T mRNA  3' ACT CCATT    CC  CGTA  TCAA  miRNA 5' TGA GGTAG    GG  GTAT  AGTT                    TA    TT                      1115111123354113312115511115 ``` \| SeedType = 3 NonSeedType = -1 \| \| --- \| --- \| --- \| --- \| --- \| --- \| | \| 92 \| WBGene00001064:dpy-2(819) \| ``` log2FC=0.839 padj=0.004 over-expressed ``` \| [seedLoc = 29,34] Seq=CTCTCTGCATCCTTTTTTGACCGTACCTCA \| ```                 GCCAGTTTTT   T    C  TC mRNA  3' ACTCCAT          TCC ACGT TC   miRNA 5' TGAGGTA          AGG TGTA AG                   GT           T    T  TT           111111133554555551113112131133 ``` \| SeedType = 1 NonSeedType = -1 \| \| --- \| --- \| --- \| --- \| --- \| --- \| | \| 93 \| WBGene00001079:dpy-20(834) \| ``` log2FC=-0.14 padj=0.97 ``` \| [seedLoc = 197,202] Seq=CTTCTCTCTTCTAACTGCCTCT \| ```          T        A  TT TCTCTTC mRNA  3'  CTCCGTCA TC  C        miRNA 5'  GAGGTAGT GG  G                 T        A  TT TATAGTT           3111121113213313333333 ``` \| SeedType = 2 NonSeedType = -1 \| \| --- \| --- \| --- \| --- \| --- \| --- \| | \| 94 \| WBGene00001080:dpy-21(835) \| ``` log2FC=-0.165 padj=0.787 ``` \| [seedLoc = 277,282] Seq=AAAATACAGAAAATCTCATTTTTTTACCTCT \| ```          T       TTTTTACTCTAAAA      AAA  mRNA  3'  CTCCATT              GACATA     miRNA 5'  GAGGTAG              TTGTAT              T       TAGG                AGTT           31111112333355555555552111113336 ``` \| SeedType = 1 NonSeedType = -1 \| \| --- \| --- \| --- \| --- \| --- \| --- \| | \| 95 \| WBGene00001145:eat-16(881) \| ``` log2FC=-0.002 padj=0.999 ``` \| [seedLoc = 263,269] Seq=AAAAGGGAAATCAGTATCCTCT \| ```          T    T  G      AGGGAAAA mRNA  3'  CTCC AT A CTAA         miRNA 5'  GAGG TA T GGTT                  T       G A    GTATAGTT           31111511316121133333333 ``` \| SeedType = 2 NonSeedType = 2 \| \| --- \| --- \| --- \| --- \| --- \| --- \| | \| 96 \| WBGene00001169:eef-1A.2(912) \| ``` log2FC=0.142 padj=0.809 ``` \| [seedLoc = 212,218] Seq=AAATCTCCAATGGATACACTCC \| ```          C   A   AGG     CTCTAAA mRNA  3'  CTC CAT    TAAC        miRNA 5'  GAG GTA    GTTG                 T       GTAG    TATAGTT           31115111333621113333333 ``` \| SeedType = 2 NonSeedType = 1 \| \| --- \| --- \| --- \| --- \| --- \| --- \| | \|  \|  \|  \| [seedLoc = 394,400] Seq=GAGTCCTTCACCGACTACGCTCC \| ```          C   G     G   CTTCCTGAG mRNA  3'  CTC CATCA CCA          miRNA 5'  GAG GTAGT GGT                   T         A   TGTATAGTT           31114111113111333333333 ``` \| SeedType = 2 NonSeedType = 1 \| \| --- \| --- \| --- \| --- \| --- \| --- \| | \| 97 \| WBGene00001172:egl-3(916) \| ``` log2FC=-0.025 padj=0.989 ``` \| [seedLoc = 61,64] Seq=CTAAGAATTTCCTCTTCCTCC \| ```          C    T      TT  AGAATC mRNA  3'  CTCC TC TCC  TA       miRNA 5'  GAGG AG AGG  GT                T    T  T   TT  ATAGTT           3111131161113321333333 ``` \| SeedType = 2 NonSeedType = -1 \| \| --- \| --- \| --- \| --- \| --- \| --- \| | \|  \|  \|  \| [seedLoc = 391,397] Seq=CACAAAACATCATTACCATCT \| ```          T  A             AAACAC mRNA  3'  CT CCATTA CT ACA       miRNA 5'  GA GGTAGT GG TGT                T         A  T   ATAGTT           31151111216126111333333 ``` \| SeedType = 3 NonSeedType = -1 \| \| --- \| --- \| --- \| --- \| --- \| --- \| | \| 98 \| WBGene00001177:egl-8(921) \| ``` log2FC=0.072 padj=0.951 ``` \| [seedLoc = 419,424] Seq=CTCTCTATTGCCACTATCTCT \| ```          T            T   C  TC mRNA  3'  CTCTATCA CCG TAT TC   miRNA 5'  GAGGTAGT GGT GTA AG            T        A   T   T  TT           3111211116112321131133 ``` \| SeedType = 2 NonSeedType = 2 \| \| --- \| --- \| --- \| --- \| --- \| --- \| | \| 99 \| WBGene00001178:egl-9(924) \| ``` log2FC=-0.117 padj=0.953 ``` \| [seedLoc = 357,363] Seq=GATATGGCAACTGATGGTGGTACTCTCA \| ```              T   G  GGTAG      GTATAG mRNA  3' ACTC CAT GT     TCAACG       miRNA 5' TGAG GTA TA     GGTTGT                        G             ATAGTT           1111511132144554211112333333 ``` \| SeedType = 2 NonSeedType = 1 \| \| --- \| --- \| --- \| --- \| --- \| --- \| | \| 100 \| WBGene00001179:egl-10(925) \| ``` log2FC=-0.096 padj=0.959 ``` \| [seedLoc = 448,453] Seq=ATTCAGACAACGTTTACCTCA \| ```                    G     GACTTA mRNA  3' ACTCCATT T CAACA       miRNA 5' TGAGGTAG A GTTGT                        T G     ATAGTT           1111111261311111333333 ``` \| SeedType = 1 NonSeedType = 1 \| \| --- \| --- \| --- \| --- \| --- \| --- \| | \| 101 \| WBGene00001182:egl-13(927) \| ``` log2FC=-0.11 padj=0.961 ``` \| [seedLoc = 335,340] Seq=TTCTATGTGTTTCCTTATTGTACCTCT \| ```          T      GTTAT   TTTGT     TT mRNA  3'  CTCCAT     TCC     GTATC   miRNA 5'  GAGGTA     AGG     TATAG            T      GT      TTG       TT           311111133555111333452111133 ``` \| SeedType = 1 NonSeedType = -1 \| \| --- \| --- \| --- \| --- \| --- \| --- \| | \| 102 \| WBGene00001186:egl-18(936) \| ``` log2FC=0.848 padj=0.404 ``` \| [seedLoc = 244,249] Seq=CGACTGTGAGCTTTACCTCC \| ```          C                     C mRNA  3'  CTCCAT  TTCGA GTGTCAG  miRNA 5'  GAGGTA  AGGTT TATAGTT           T      GT     G                   31111116612121621211125 ``` \| SeedType = 1 NonSeedType = 2 \| \| --- \| --- \| --- \| --- \| --- \| --- \| | \|  \|  \|  \| [seedLoc = 339,345] Seq=CTCTGTGCATGGTTGTGTTATATTCACCTCC \| ```          C     C    ATTGTGTTGGT       TC mRNA  3'  CTCCA TTAT           ACGTGTC   miRNA 5'  GAGGT AGTA           TGTATAG            T          GGT               TT           3111115121133345455445112121133 ``` \| SeedType = 2 NonSeedType = -1 \| \| --- \| --- \| --- \| --- \| --- \| --- \| | \| 103 \| WBGene00001193:egl-26(942) \| ``` log2FC=-0.175 padj=0.912 ``` \| [seedLoc = 70,75] Seq=TTCTATAGTCATTCTGTTCTATCTCA \| ```                  TT    T  TG     TT mRNA  3' ACTCTATC  GTCT AC  ATATC   miRNA 5' TGAGGTAG  TAGG TG  TATAG                          T         TT           11112111552112311541111133 ``` \| SeedType = 2 NonSeedType = -1 \| \| --- \| --- \| --- \| --- \| --- \| --- \| | \| 104 \| WBGene00001230:eif-3.G(964) \| ``` log2FC=0.012 padj=0.994 ``` \| [seedLoc = 255,260] Seq=TCTTCAATGATAAATCTATCTCT \| ```          T       TAAA     ACTTCT mRNA  3'  CTCTATC    TAGTA       miRNA 5'  GAGGTAG    GTTGT                T       TAG      ATAGTT           31112111333521221333333 ``` \| SeedType = 2 NonSeedType = -1 \| \| --- \| --- \| --- \| --- \| --- \| --- \| | \| 105 \| WBGene00001249:elt-1(981) \| ``` log2FC=-0.039 padj=0.989 ``` \| [seedLoc = 70,75] Seq=TTCCATTCTTCTTCTGCCTCC \| ```          C           TT TTACCTT mRNA  3'  CTCCGTC TTC  C        miRNA 5'  GAGGTAG AGG  G                 T       T   TT TATAGTT           3111121161213313333333 ``` \| SeedType = 2 NonSeedType = -1 \| \| --- \| --- \| --- \| --- \| --- \| --- \| | \|  \|  \|  \| [seedLoc = 194,199] Seq=TATTAATTTTCCTGCGGTTTTCATTACCTCA \| ```                 TACTTTTGG     TTTTAATTAT mRNA  3' ACTCCAT         CGTCC           miRNA 5' TGAGGTA         GTAGG                                         TTGTATAGTT           1111111555555544121113333333333 ``` \| SeedType = 1 NonSeedType = -1 \| \| --- \| --- \| --- \| --- \| --- \| --- \| | \| 106 \| WBGene00001320:ent-1(996) \| ``` log2FC=-0.199 padj=0.672 ``` \| [seedLoc = 132,137] Seq=CACTAGTGTTCTTTCCTTAAAATACCTCT \| ```          T      AAAAT   TTTCTTGTG    C mRNA  3'  CTCCAT     TCC         ATCA  miRNA 5'  GAGGTA     AGG         TAGT           T      GT      TTGTA        T           31111113355511133333545411113 ``` \| SeedType = 1 NonSeedType = -1 \| \| --- \| --- \| --- \| --- \| --- \| --- \| | \| 107 \| WBGene00001330:eps-8(1001) \| ``` log2FC=-0.027 padj=0.984 ``` \| [seedLoc = 173,178] Seq=ATATGTTCAATTTACTGCCTCC \| ```          C              T   ATA mRNA  3'  CTCCGTCATTTAAC TGT    miRNA 5'  GAGGTAGTAGGTTG ATA             T              T   GTT           3111121111221113121333 ``` \| SeedType = 2 NonSeedType = 1 \| \| --- \| --- \| --- \| --- \| --- \| --- \| | \| 108 \| WBGene00001345:fos-1(1017) \| ``` log2FC=-0.109 padj=0.932 ``` \| [seedLoc = 384,389] Seq=TCATCATGAATCATTTACCTCA \| ```                  TA    GTACTACT mRNA  3' ACTCCATT  CTAA         miRNA 5' TGAGGTAG  GGTT                          TA    GTATAGTT           1111111233121133333333 ``` \| SeedType = 1 NonSeedType = 2 \| \| --- \| --- \| --- \| --- \| --- \| --- \| | \| 109 \| WBGene00001352:evl-14(1018) \| ``` log2FC=0.01 padj=0.994 ``` \| [seedLoc = 103,108] Seq=TCTTATTTCTCTTCGTACCTCA \| ```                 GC   TCTT    CT mRNA  3' ACTCCAT  TTC    TATT   miRNA 5' TGAGGTA  AGG    ATAG                   GT   TTGT    TT           1111111331213333111233 ``` \| SeedType = 1 NonSeedType = -1 \| \| --- \| --- \| --- \| --- \| --- \| --- \| | \| 110 \| WBGene00001365:exc-4(1020) \| ``` log2FC=0.026 padj=0.988 ``` \| [seedLoc = 972,977] Seq=ACACACACACACATCTATCTCA \| ```                  TA AC   CACACA mRNA  3' ACTCTATC  C  ACA       miRNA 5' TGAGGTAG  G  TGT                        TA GT   ATAGTT           1111211133133111333333 ``` \| SeedType = 2 NonSeedType = -1 \| \| --- \| --- \| --- \| --- \| --- \| --- \| | \| 111 \| WBGene00001366:exc-5(1021) \| ``` log2FC=-0.091 padj=0.956 ``` \| [seedLoc = 315,320] Seq=AAATTGTCTCTTTTACTACCTTC \| ```          C           TCTC      A mRNA  3'  TTCCATCATTT    TGTTAA  miRNA 5'  GAGGTAGTAGG    ATAGTT           T           TTGT                  32111111112233331212115 ``` \| SeedType = 3 NonSeedType = -1 \| \| --- \| --- \| --- \| --- \| --- \| --- \| | \| 112 \| WBGene00001413:fem-3(1055) \| ``` log2FC=0.02 padj=1.0 ``` \| [seedLoc = 108,113] Seq=CACCGATTTGCCTTCTATCTCA \| ```                  T    TTTAGCCAC mRNA  3' ACTCTATC TCCG          miRNA 5' TGAGGTAG AGGT                           T    TGTATAGTT           1111211131112333333333 ``` \| SeedType = 2 NonSeedType = 2 \| \| --- \| --- \| --- \| --- \| --- \| --- \| | \| 113 \| WBGene00001434:fkh-2(1068) \| ``` log2FC=0.126 padj=1.0 ``` \| [seedLoc = 238,242] Seq=CTTGTCATCCTTCTCCCTCC \| ```          C    C  T   T       C  mRNA  3'  CTCC TC TCC AC TGTT   miRNA 5'  GAGG AG AGG TG ATAG            T    T  T   T  T    TT           3111131131113116121236 ``` \| SeedType = 2 NonSeedType = -1 \| \| --- \| --- \| --- \| --- \| --- \| --- \| | \| 114 \| WBGene00001436:fkh-4(1070) \| ``` log2FC=0.548 padj=1.0 ``` \| [seedLoc = 233,238] Seq=TGCGTCAAAAACACTGCCTCA \| ```                   CAAA   TG    T mRNA  3' ACTCCGTCA    AAC    CG  miRNA 5' TGAGGTAGT    TTG    GT                    AGG    TATA  T           11111211133351113366123 ``` \| SeedType = 2 NonSeedType = 1 \| \| --- \| --- \| --- \| --- \| --- \| --- \| | \| 115 \| WBGene00001439:fkh-7(1072) \| ``` log2FC=-0.382 padj=0.531 ``` \| [seedLoc = 487,493] Seq=CACTACACATTTGATCTACTACCATCT \| ```          T  A          GTTT   C    C mRNA  3'  CT CCATCATCTA    ACA ATCA  miRNA 5'  GA GGTAGTAGGT    TGT TAGT           T                    A    T           311511111111214555111311113 ``` \| SeedType = 3 NonSeedType = 2 \| \| --- \| --- \| --- \| --- \| --- \| --- \| | \|  \|  \|  \| [seedLoc = 652,657] Seq=CTTAATATTCACTGAAATTACCTCC \| ```          C        AAG   CT    ATTC mRNA  3'  CTCCATTA   TCA  TATA     miRNA 5'  GAGGTAGT   GGT  GTAT              T        A     T     AGTT           3111111213542113521113333 ``` \| SeedType = 1 NonSeedType = 2 \| \| --- \| --- \| --- \| --- \| --- \| --- \| | \|  \|  \|  \| [seedLoc = 782,787] Seq=TTTTATCATTGAATTACCTCA \| ```                   AG T       TT mRNA  3' ACTCCATTA  T AC TATT   miRNA 5' TGAGGTAGT  G TG ATAG                     AG T  T    TT           1111111213323116111233 ``` \| SeedType = 1 NonSeedType = -1 \| \| --- \| --- \| --- \| --- \| --- \| --- \| | \| 116 \| WBGene00001442:fkh-10(1075) \| ``` log2FC=-0.847 padj=1.0 ``` \| [seedLoc = 71,77] Seq=TCTTCCGTTTCCTGTTACCCTCC \| ```          C  C         TTTGCCTTCT mRNA  3'  CT CCATTGTCC           miRNA 5'  GA GGTAGTAGG                    T            TTGTATAGTT           31151111221113333333333 ``` \| SeedType = 3 NonSeedType = -1 \| \| --- \| --- \| --- \| --- \| --- \| --- \| | \| 117 \| WBGene00001457:flp-14(1090) \| ``` log2FC=0.05 padj=0.985 ``` \| [seedLoc = 123,128] Seq=TTCTATATTTTTCTCTCTATCTCT \| ```          T       TC  TTTT      TT mRNA  3'  CTCTATC  TC    TATATC   miRNA 5'  GAGGTAG  GG    GTATAG            T       TA  TT        TT           311121113321335521111133 ``` \| SeedType = 2 NonSeedType = -1 \| \| --- \| --- \| --- \| --- \| --- \| --- \| | \| 118 \| WBGene00001459:flp-16(1092) \| ``` log2FC=-0.186 padj=0.95 ``` \| [seedLoc = 246,252] Seq=CCCTGTGTGTACCAGTGTACCCTCA \| ```             C    GTGA   TGT     CC mRNA  3' ACT CCAT    CCA   GTGTC   miRNA 5' TGA GGTA    GGT   TATAG                    GTA    TG      TT           1115111133351113352121133 ``` \| SeedType = 3 NonSeedType = 1 \| \| --- \| --- \| --- \| --- \| --- \| --- \| | \| 119 \| WBGene00001460:flp-17(1093) \| ``` log2FC=0.002 padj=0.999 ``` \| [seedLoc = 67,73] Seq=TTTTGAGCAACTCCTACTCTCT \| ```          T   T    C       AGTTTT mRNA  3'  CTC CATC  TCAACG       miRNA 5'  GAG GTAG  GGTTGT                T        TA      ATAGTT           31115111136211112333333 ``` \| SeedType = 2 NonSeedType = 1 \| \| --- \| --- \| --- \| --- \| --- \| --- \| | \| 120 \| WBGene00001464:flp-21(1097) \| ``` log2FC=-0.464 padj=0.911 ``` \| [seedLoc = 168,174] Seq=GCAGAAACAATGTTCTTTTTACTCTCT \| ```          T   T    T   TTGT    AAGACG mRNA  3'  CTC CATT TTC    AACA       miRNA 5'  GAG GTAG AGG    TTGT                T        T           ATAGTT           311151112312155451111333333 ``` \| SeedType = 2 NonSeedType = 1 \| \| --- \| --- \| --- \| --- \| --- \| --- \| | \|  \|  \|  \| [seedLoc = 187,192] Seq=CTCTCTCTCTTTTTCTATCTCA \| ```                  TTTTTCTCTCTCTC mRNA  3' ACTCTATC               miRNA 5' TGAGGTAG                                TAGGTTGTATAGTT           1111211133333333333333 ``` \| SeedType = 2 NonSeedType = -1 \| \| --- \| --- \| --- \| --- \| --- \| --- \| | \|  \|  \|  \| [seedLoc = 193,197] Seq=CTCTCTCTCTTTTTCTATCTCAC \| ```          C        TTTTTCTCTCTCTC mRNA  3'  ACTCTATC               miRNA 5'  TGAGGTAG                                 TAGGTTGTATAGTT           51111211133333333333333 ``` \| SeedType = 2 NonSeedType = -1 \| \| --- \| --- \| --- \| --- \| --- \| --- \| | \| 121 \| WBGene00001466:flr-2(1099) \| ``` log2FC=0.668 padj=0.746 ``` \| [seedLoc = 354,359] Seq=CAATTGACAATGTTGATTGAATGATACCTCT \| ```          T      AGTAAGTTAGT G           C mRNA  3'  CTCCAT           T TAACA GTTAA  miRNA 5'  GAGGTA           A GTTGT TAGTT           T      GT          G     A                 31111113355545554513211116212115 ``` \| SeedType = 1 NonSeedType = 1 \| \| --- \| --- \| --- \| --- \| --- \| --- \| | \| 122 \| WBGene00001510:fzr-1(1138) \| ``` log2FC=-0.188 padj=0.609 ``` \| [seedLoc = 130,135] Seq=TTGTGCCTTTTTAAAGAAAATTATTACCTCC \| ```          C          AAAAGAAATTTTTC         mRNA  3'  CTCCATTATT              CGTGTT   miRNA 5'  GAGGTAGTAG              GTATAG            T          GTT                 TT           311111121123335455555555512121266 ``` \| SeedType = 1 NonSeedType = -1 \| \| --- \| --- \| --- \| --- \| --- \| --- \| | \| 123 \| WBGene00001511:fzy-1(1139) \| ``` log2FC=-0.108 padj=0.901 ``` \| [seedLoc = 172,178] Seq=TCTCTTCTTGTCTTCTCACCTCT \| ```          T     C  T   GTTCTTCTCT mRNA  3'  CTCCA TC TCT           miRNA 5'  GAGGT AG AGG                    T        T   TTGTATAGTT           31111151131123333333333 ``` \| SeedType = 2 NonSeedType = -1 \| \| --- \| --- \| --- \| --- \| --- \| --- \| | \| 124 \| WBGene00001558:gdi-1(1165) \| ``` log2FC=-0.025 padj=0.985 ``` \| [seedLoc = 170,176] Seq=TCTTTTCCAGTCTCTATCCTCA \| ```               T          CTTTTCT mRNA  3' ACTCC ATC TCTGAC        miRNA 5' TGAGG TAG AGGTTG                          T      TATAGTT           11111511161122113333333 ``` \| SeedType = 2 NonSeedType = 2 \| \| --- \| --- \| --- \| --- \| --- \| --- \| | \| 125 \| WBGene00001559:gei-1(1166) \| ``` log2FC=-0.177 padj=0.825 ``` \| [seedLoc = 215,220] Seq=ATCTGGAATACCCTTACCTCC \| ```          C       C    TAAG   TA mRNA  3'  CTCCATT  CCA    GTC   miRNA 5'  GAGGTAG  GGT    TAG            T       TA   TGTA   TT           3111111236111333321133 ``` \| SeedType = 1 NonSeedType = 1 \| \| --- \| --- \| --- \| --- \| --- \| --- \| | \| 126 \| WBGene00001562:lin-66(1169) \| ``` log2FC=-0.05 padj=0.977 ``` \| [seedLoc = 1143,1149] Seq=CTCTGGTCTTCCTCCATTCACCTCT \| ```          T     C   CC   TT TG   TC mRNA  3'  CTCCA TTA  TCC  C  GTC   miRNA 5'  GAGGT AGT  AGG  G  TAG            T              TT TA   TT           3111115121551113313321133 ``` \| SeedType = 2 NonSeedType = -1 \| \| --- \| --- \| --- \| --- \| --- \| --- \| | \| 127 \| WBGene00001564:icl-1(1171) \| ``` log2FC=-0.916 padj=8.14703303514623E-6 ``` \| [seedLoc = 178,183] Seq=AGACTTTCCCACGATTTGAGACTTTACCTCA \| ```                  TCAGA        CCCTT    A mRNA  3' ACTCCATT     GTTTAGCA     TCAG  miRNA 5' TGAGGTAG     TAGGTTGT     AGTT                                AT                   1111111255545212212113355511125 ``` \| SeedType = 1 NonSeedType = 2 \| \| --- \| --- \| --- \| --- \| --- \| --- \| | \|  \|  \|  \| [seedLoc = 195,199] Seq=TGAGACTTTACCTCAT \| ```          T              AGT      mRNA  3'  ACTCCAT  TTCAG         miRNA 5'  TGAGGTA  AGGTT                          GT     GTATAGTT           51111111661211233366666 ``` \| SeedType = 1 NonSeedType = 2 \| \| --- \| --- \| --- \| --- \| --- \| --- \| | \| 128 \| WBGene00001564:icl-1(1172) \| ``` log2FC=-0.916 padj=8.14703303514623E-6 ``` \| [seedLoc = 43,48] Seq=AAATTGGAATTAATTTCAGGATACCTCG \| ```          G      AGGAC      TAAG     A mRNA  3'  CTCCAT     TTTAAT    GTTAA  miRNA 5'  GAGGTA     AGGTTG    TAGTT           T      GT         TA                   3111111334551221123354212115 ``` \| SeedType = 1 NonSeedType = 2 \| \| --- \| --- \| --- \| --- \| --- \| --- \| | \| 129 \| WBGene00001570:gei-13(1177) \| ``` log2FC=-0.06 padj=0.979 ``` \| [seedLoc = 222,227] Seq=TTCTGTCTCCATTCTGATACCTCA \| ```                 A    T  CTC    TT mRNA  3' ACTCCAT GTCT AC   TGTC   miRNA 5' TGAGGTA TAGG TG   ATAG                   G    T  T      TT           111111132112311355121133 ``` \| SeedType = 1 NonSeedType = -1 \| \| --- \| --- \| --- \| --- \| --- \| --- \| | \| 130 \| WBGene00001571:gei-14(1178) \| ``` log2FC=-0.183 padj=0.823 ``` \| [seedLoc = 80,85] Seq=GATAATTATCCCATCTACTTCT \| ```          T       TA  CTATTAATAG mRNA  3'  CTTCATC  CC           miRNA 5'  GAGGTAG  GG                    T       TA  TTGTATAGTT           3112111133113333333333 ``` \| SeedType = 3 NonSeedType = -1 \| \| --- \| --- \| --- \| --- \| --- \| --- \| | \| 131 \| WBGene00001595:gld-1(1205) \| ``` log2FC=-0.102 padj=0.884 ``` \| [seedLoc = 144,150] Seq=ATGTCATCAATTGGTATCTACCATCT \| ```          T  A     TATGG     TACTGTA mRNA  3'  CT CCATC     TTAAC        miRNA 5'  GA GGTAG     GGTTG                 T        TA        TATAGTT           31151111133544221113333333 ``` \| SeedType = 3 NonSeedType = 1 \| \| --- \| --- \| --- \| --- \| --- \| --- \| | \|  \|  \|  \| [seedLoc = 435,440] Seq=CCCTACCCATTCATACTACCTCG \| ```          G         A  T  CC   CC mRNA  3'  CTCCATCAT CT AC  ATC   miRNA 5'  GAGGTAGTA GG TG  TAG            T            T  TA   TT           31111111115123113311133 ``` \| SeedType = 1 NonSeedType = -1 \| \| --- \| --- \| --- \| --- \| --- \| --- \| | \| 132 \| WBGene00001596:gld-2(1206) \| ``` log2FC=-0.123 padj=0.821 ``` \| [seedLoc = 756,761] Seq=TGATTGTACCATAAGTACCTCC \| ```          C      G A   C        T mRNA  3'  CTCCAT A  TA CATGTTAG  miRNA 5'  GAGGTA T  GT GTATAGTT           T      G AG  T                    31111113136213111212125 ``` \| SeedType = 1 NonSeedType = -1 \| \| --- \| --- \| --- \| --- \| --- \| --- \| | \| 133 \| WBGene00001597:gld-3(1208) \| ``` log2FC=0.092 padj=0.927 ``` \| [seedLoc = 29,34] Seq=TTTCCAATAGCTTCTTCTACTTCT \| ```          T       TTC       ACCTTT mRNA  3'  CTTCATC   TTCGATA       miRNA 5'  GAGGTAG   AGGTTGT                T       T         ATAGTT           311211113551212121333333 ``` \| SeedType = 3 NonSeedType = 2 \| \| --- \| --- \| --- \| --- \| --- \| --- \| | \| 134 \| WBGene00001602:gln-1(1213) \| ``` log2FC=-0.217 padj=0.707 ``` \| [seedLoc = 114,120] Seq=TTCTCTCCAATTAGTACCATCC \| ```          C  A    G       CTCTCTT mRNA  3'  CT CCAT AT TAAC        miRNA 5'  GA GGTA TA GTTG                 T       G  G    TATAGTT           31151111311621113333333 ``` \| SeedType = 3 NonSeedType = 1 \| \| --- \| --- \| --- \| --- \| --- \| --- \| | \| 135 \| WBGene00001650:gon-1(1254) \| ``` log2FC=0.132 padj=0.915 ``` \| [seedLoc = 42,47] Seq=TCTCGTATGAATCTCTACTTCT \| ```          T             G    CTCT mRNA  3'  CTTCATC TCTAA TATG     miRNA 5'  GAGGTAG AGGTT GTAT              T       T          AGTT           31121111611211421123333 ``` \| SeedType = 3 NonSeedType = 2 \| \| --- \| --- \| --- \| --- \| --- \| --- \| | \| 136 \| WBGene00001666:gpa-4(1262) \| ``` log2FC=0.262 padj=1.0 ``` \| [seedLoc = 64,70] Seq=ATTTACTCAGATGGATACTCTCG \| ```          G   T   AGGTA   TC   TA mRNA  3'  CTC CAT     GAC  ATT   miRNA 5'  GAG GTA     TTG  TAG            T       GTAGG   TA   TT           31115111333332113311233 ``` \| SeedType = 2 NonSeedType = -1 \| \| --- \| --- \| --- \| --- \| --- \| --- \| | \| 137 \| WBGene00001672:gpa-10(1264) \| ``` log2FC=0.554 padj=1.0 ``` \| [seedLoc = 44,49] Seq=TGTTCTTCAATTTAGTATCTGCCTCA \| ```                  TATG       TTCTTGT mRNA  3' ACTCCGTC    ATTTAAC        miRNA 5' TGAGGTAG    TAGGTTG                                    TATAGTT           11111211555411221113333333 ``` \| SeedType = 2 NonSeedType = 1 \| \| --- \| --- \| --- \| --- \| --- \| --- \| | \| 138 \| WBGene00001679:gpb-1(1268) \| ``` log2FC=0.135 padj=0.821 ``` \| [seedLoc = 190,196] Seq=CTCTTTTTCTTCTCTCACCTCT \| ```          T     C      TCTTTTTCTC mRNA  3'  CTCCA TC TCT           miRNA 5'  GAGGT AG AGG                    T        T   TTGTATAGTT           31111151161123333333333 ``` \| SeedType = 2 NonSeedType = -1 \| \| --- \| --- \| --- \| --- \| --- \| --- \| | \| 139 \| WBGene00001680:gpb-2(1269) \| ``` log2FC=0.077 padj=0.961 ``` \| [seedLoc = 115,120] Seq=CACTATTTTCTCCTATCTCA \| ```                  C   TT       C mRNA  3' ACTCTATC TCT    TATCA  miRNA 5' TGAGGTAG AGG    ATAGT                   T   TTGT     T           1111211131123366111113 ``` \| SeedType = 2 NonSeedType = -1 \| \| --- \| --- \| --- \| --- \| --- \| --- \| | \| 140 \| WBGene00001699:grd-10(1287) \| ``` log2FC=1.081 padj=3.07340887295593E-5 over-expressed ``` \| [seedLoc = 57,62] Seq=TTTTTGACGATCTTCTTTTACCCTATCTCA \| ```                  CCATTTTCT       GTTTTT mRNA  3' ACTCTATC         TCTAGCA       miRNA 5' TGAGGTAG         AGGTTGT                        T               ATAGTT           111121113555555551121211333333 ``` \| SeedType = 2 NonSeedType = 2 \| \| --- \| --- \| --- \| --- \| --- \| --- \| | \| 141 \| WBGene00001743:grp-1(1316) \| ``` log2FC=0.125 padj=0.922 ``` \| [seedLoc = 158,164] Seq=TACTTCTATAATCTGTTATCCTCC \| ```          C    T            CT   T mRNA  3'  CTCC ATTGTCTAATAT  TCA  miRNA 5'  GAGG TAGTAGGTTGTA  AGT           T                 T    T           311115112211211211351113 ``` \| SeedType = 2 NonSeedType = 2 \| \| --- \| --- \| --- \| --- \| --- \| --- \| | \| 142 \| WBGene00001745:gsa-1(1318) \| ``` log2FC=0.205 padj=0.8 ``` \| [seedLoc = 456,460] Seq=AGCTTGCCTGACCTACCTAA \| ```          AA      CA     TT  A      mRNA  3'   TCCATC  GTCCG  CG       miRNA 5'   AGGTAG  TAGGT  GT                TG             T   ATAGTT           3311111155211123512366666 ``` \| SeedType = 3 NonSeedType = 2 \| \| --- \| --- \| --- \| --- \| --- \| --- \| | \| 143 \| WBGene00001748:gsp-2(1321) \| ``` log2FC=0.136 padj=0.839 ``` \| [seedLoc = 246,251] Seq=CGTTGTCAACATCCCTTCCTCT \| ```          T    T  CCTA         GC mRNA  3'  CTCC TC    CAAC TGTT   miRNA 5'  GAGG AG    GTTG ATAG            T    T  TAG     T    TT           31111311333511116121233 ``` \| SeedType = 2 NonSeedType = 1 \| \| --- \| --- \| --- \| --- \| --- \| --- \| | \| 144 \| WBGene00001757:gst-9(1330) \| ``` log2FC=0.252 padj=0.894 ``` \| [seedLoc = 36,41] Seq=TAGTGTTCATTTTTATCTACCTCG \| ```          G       T    TT  T   G T mRNA  3'  CTCCATC ATTT  AC TGT A  miRNA 5'  GAGGTAG TAGG  TG ATA T           T            T   T   G T           311111115112235113121313 ``` \| SeedType = 1 NonSeedType = -1 \| \| --- \| --- \| --- \| --- \| --- \| --- \| | \| 145 \| WBGene00001820:ham-1(1376) \| ``` log2FC=0.473 padj=0.903 ``` \| [seedLoc = 124,129] Seq=ATTTTGTTTTTTTTCTACTTCT \| ```          T       TTTTTTTTGTTTTA mRNA  3'  CTTCATC               miRNA 5'  GAGGTAG                        T       TAGGTTGTATAGTT           3112111133333333333333 ``` \| SeedType = 3 NonSeedType = -1 \| \| --- \| --- \| --- \| --- \| --- \| --- \| | \| 146 \| WBGene00001821:ham-2(1377) \| ``` log2FC=-0.467 padj=0.826 ``` \| [seedLoc = 241,246] Seq=GTACATAATGCCTATTCTCGTAGCTACTTCT \| ```          T        ATGCTCTTA    TAAT          mRNA  3'  CTTCATCG         TCCG    ACATG     miRNA 5'  GAGGTAGT         AGGT    TGTAT              T                              AGTT           31121111255455555511125555111126666 ``` \| SeedType = 3 NonSeedType = 2 \| \| --- \| --- \| --- \| --- \| --- \| --- \| | \| 147 \| WBGene00001824:hbl-1(1379) \| ``` log2FC=1.012 padj=0.093 over-expressed ``` \| [seedLoc = 40,46] Seq=CACTTCCCATAGCCCCTTACCCTCG \| ```          G  C     CC      CCCT   C mRNA  3'  CT CCATT  CCGATA    TCA  miRNA 5'  GA GGTAG  GGTTGT    AGT           T        TA      AT     T           3115111123311212133551113 ``` \| SeedType = 3 NonSeedType = 2 \| \| --- \| --- \| --- \| --- \| --- \| --- \| | \|  \|  \|  \| [seedLoc = 229,234] Seq=TACTGTTTTCTTTTACCTCT \| ```          T       T   TT       T mRNA  3'  CTCCATT TCT    TGTCA  miRNA 5'  GAGGTAG AGG    ATAGT           T       T   TTGT     T           3111111231123366121113 ``` \| SeedType = 1 NonSeedType = -1 \| \| --- \| --- \| --- \| --- \| --- \| --- \| | \|  \|  \|  \| [seedLoc = 266,271] Seq=CTCTGTCTCACTTTCTACCTCC \| ```          C       T    CTC    TC mRNA  3'  CTCCATC TTCA   TGTC   miRNA 5'  GAGGTAG AGGT   ATAG            T       T    TGT    TT           3111111131211333121133 ``` \| SeedType = 1 NonSeedType = 2 \| \| --- \| --- \| --- \| --- \| --- \| --- \| | \|  \|  \|  \| [seedLoc = 349,355] Seq=TCATTTACAGTTTTGTACTCTCA \| ```              T   GT        TTACT mRNA  3' ACTC CAT  TTTGACAT      miRNA 5' TGAG GTA  AGGTTGTA                       GT        TAGTT           11115111331222111133333 ``` \| SeedType = 2 NonSeedType = 2 \| \| --- \| --- \| --- \| --- \| --- \| --- \| | \|  \|  \|  \| [seedLoc = 407,412] Seq=CTCAGTACATGTAGTACCTCC \| ```          C      G  G       ACTC mRNA  3'  CTCCAT AT T ACATG     miRNA 5'  GAGGTA TA G TGTAT              T      G  G T     AGTT           3111111311326111123333 ``` \| SeedType = 1 NonSeedType = -1 \| \| --- \| --- \| --- \| --- \| --- \| --- \| | \|  \|  \|  \| [seedLoc = 916,921] Seq=TGATTTCACATTTTCCAGACTATCTCG \| ```          G        GA  TTTT   CT    T mRNA  3'  CTCTATCA  CC    ACA  TTAG  miRNA 5'  GAGGTAGT  GG    TGT  AGTT           T        A   T      AT                311121111351135551113312125 ``` \| SeedType = 2 NonSeedType = -1 \| \| --- \| --- \| --- \| --- \| --- \| --- \| | \|  \|  \|  \| [seedLoc = 934,939] Seq=AGACTATCTCGCACTTTCATTCTACCTCA \| ```                  TTACT    C  TC      A mRNA  3' ACTCCATC     TTCA GC  TATCAG  miRNA 5' TGAGGTAG     AGGT TG  ATAGTT                   T           T                   11111111355551211521351111125 ``` \| SeedType = 1 NonSeedType = 2 \| \| --- \| --- \| --- \| --- \| --- \| --- \| | \|  \|  \|  \| [seedLoc = 1054,1059] Seq=TTATTCGCTCCCTTTTTTCTATCTCA \| ```                  TTTTT   CT  CTTATT mRNA  3' ACTCTATC     TCC  CG       miRNA 5' TGAGGTAG     AGG  GT                        T       TT  ATAGTT           11112111355551113312333333 ``` \| SeedType = 2 NonSeedType = -1 \| \| --- \| --- \| --- \| --- \| --- \| --- \| | \|  \|  \|  \| [seedLoc = 1119,1124] Seq=AAACTACTTGTCCGCTACCTTA \| ```                      TGTTC     A mRNA  3' ATTCCATCG CC     ATCAA  miRNA 5' TGAGGTAGT GG     TAGTT                    A  TTGTA                 12111111261133333111115 ``` \| SeedType = 3 NonSeedType = -1 \| \| --- \| --- \| --- \| --- \| --- \| --- \| | \|  \|  \|  \| [seedLoc = 1128,1133] Seq=AAACTACTTGTCCGCTACCTTAT \| ```          T            TGTTC     A mRNA  3'  ATTCCATCG CC     ATCAA  miRNA 5'  TGAGGTAGT GG     TAGTT                     A  TTGTA                 512111111261133333111115 ``` \| SeedType = 3 NonSeedType = -1 \| \| --- \| --- \| --- \| --- \| --- \| --- \| | \|  \|  \|  \| [seedLoc = 1204,1209] Seq=ACCTGTATAATGCCTTCTACCTCC \| ```          C       T    TA       CA mRNA  3'  CTCCATC TCCG  ATATGTC   miRNA 5'  GAGGTAG AGGT  TGTATAG            T       T             TT           311111113111255121121133 ``` \| SeedType = 1 NonSeedType = 2 \| \| --- \| --- \| --- \| --- \| --- \| --- \| | \|  \|  \|  \| [seedLoc = 1247,1252] Seq=TTAATTACCATTTTCTACCTCA \| ```                  T    C   TAATT mRNA  3' ACTCCATC TTTA CAT      miRNA 5' TGAGGTAG AGGT GTA                       T    T   TAGTT           1111111131221311133333 ``` \| SeedType = 1 NonSeedType = 2 \| \| --- \| --- \| --- \| --- \| --- \| --- \| | \|  \|  \|  \| [seedLoc = 1288,1292] Seq=TATTATACAACCGTTCCACCTCAA \| ```          A      C TTG           T mRNA  3'  ACTCCA C   CCAACATATTA  miRNA 5'  TGAGGT G   GGTTGTATAGT                  A TA            T           511111131334111111111213 ``` \| SeedType = 2 NonSeedType = 1 \| \| --- \| --- \| --- \| --- \| --- \| --- \| | \| 148 \| WBGene00001827:hcf-1(1380) \| ``` log2FC=-0.053 padj=0.962 ``` \| [seedLoc = 210,215] Seq=ATCAAATCATTCACTACCTTT \| ```          T           T  TAAACTA mRNA  3'  TTCCATCA CT AC        miRNA 5'  GAGGTAGT GG TG                 T        A  T  TATAGTT           3211111116123113333333 ``` \| SeedType = 3 NonSeedType = -1 \| \| --- \| --- \| --- \| --- \| --- \| --- \| | \| 149 \| WBGene00001851:hif-1(1397) \| ``` log2FC=-0.039 padj=0.974 ``` \| [seedLoc = 159,164] Seq=TGACTTTACACCTGCGGTACCTCA \| ```                 GG          T    T mRNA  3' ACTCCAT  CGTCC ACAT TCAG  miRNA 5' TGAGGTA  GTAGG TGTA AGTT                         T    T                1111111441211161111311125 ``` \| SeedType = 1 NonSeedType = -1 \| \| --- \| --- \| --- \| --- \| --- \| --- \| | \| 150 \| WBGene00001852:hil-1(1398) \| ``` log2FC=-0.303 padj=0.819 ``` \| [seedLoc = 30,35] Seq=CATTTTCTCGCTCTCTATCTCT \| ```          T       TC   CTCTTTTAC mRNA  3'  CTCTATC  TCG          miRNA 5'  GAGGTAG  GGT                   T       TA   TGTATAGTT           3111211133212333333333 ``` \| SeedType = 2 NonSeedType = 2 \| \| --- \| --- \| --- \| --- \| --- \| --- \| | \| 151 \| WBGene00001853:hil-2(1399) \| ``` log2FC=-0.518 padj=0.011 ``` \| [seedLoc = 49,54] Seq=TACTGTATTATACCTCC \| ```          C      AT            T mRNA  3'  CTCCAT       TATGTCA  miRNA 5'  GAGGTA       GTATAGT           T      GTAGGTT       T           3111111336666621121113 ``` \| SeedType = 1 NonSeedType = -1 \| \| --- \| --- \| --- \| --- \| --- \| --- \| | \| 152 \| WBGene00001854:hil-3(1400) \| ``` log2FC=-0.481 padj=0.076 ``` \| [seedLoc = 34,38] Seq=TTTAATTTTCTTATACCTCAC \| ```          C       AT   TTTAATTT   mRNA  3'  ACTCCAT  TCT           miRNA 5'  TGAGGTA  AGG                            GT   TTGTATAGTT           51111111331123333333366 ``` \| SeedType = 1 NonSeedType = -1 \| \| --- \| --- \| --- \| --- \| --- \| --- \| | \| 153 \| WBGene00001856:hil-5(1402) \| ``` log2FC=-0.283 padj=0.304 ``` \| [seedLoc = 47,52] Seq=TAATTCGATCTCACCATTGTAATATACCTCC \| ```          C      ATAATGTTAC  C      TTAAT   mRNA  3'  CTCCAT          CA TCTAGC        miRNA 5'  GAGGTA          GT AGGTTG                 T                         TATAGTT           311111155555455551151121213333366 ``` \| SeedType = 1 NonSeedType = 2 \| \| --- \| --- \| --- \| --- \| --- \| --- \| | \| 154 \| WBGene00001858:hil-7(1404) \| ``` log2FC=-0.422 padj=1.0 ``` \| [seedLoc = 100,105] Seq=TTAATGATTCTAAACTACCTCA \| ```                   AATCTTAGTAATT mRNA  3' ACTCCATCA              miRNA 5' TGAGGTAGT                                AGGTTGTATAGTT           1111111113333333333333 ``` \| SeedType = 1 NonSeedType = -1 \| \| --- \| --- \| --- \| --- \| --- \| --- \| | \| 155 \| WBGene00001859:hil-8(1406) \| ``` log2FC=-1.708 padj=1.0 ``` \| [seedLoc = 768,773] Seq=AAGTGCAATTGGCCAGGACACTGCTACTTCT \| ```          T          ACAGGACCGGT       AA   mRNA  3'  CTTCATCGTC           TAACGTG     miRNA 5'  GAGGTAGTAG           GTTGTAT              T                            AGTT           311211112115554455544521112123366 ``` \| SeedType = 3 NonSeedType = 1 \| \| --- \| --- \| --- \| --- \| --- \| --- \| | \| 156 \| WBGene00001898:his-24(1428) \| ``` log2FC=-0.487 padj=0.037 ``` \| [seedLoc = 32,37] Seq=TCTAATATGTTCCTACCTCT \| ```          T           T     ATCT mRNA  3'  CTCCATC  CT GTATA     miRNA 5'  GAGGTAG  GG TGTAT              T       TA  T     AGTT           3111111166123221113333 ``` \| SeedType = 1 NonSeedType = -1 \| \| --- \| --- \| --- \| --- \| --- \| --- \| | \| 157 \| WBGene00001919:his-45(1438) \| ``` log2FC=-0.349 padj=0.127 ``` \| [seedLoc = 75,81] Seq=CTCTTTAGGGCCACAAATATCCTCA \| ```               T  AAA       G  T  TC mRNA  3' ACTCC AT   CA CCGG AT TC   miRNA 5' TGAGG TA   GT GGTT TA AG                         A    G  T  TT           11111511555116112231131133 ``` \| SeedType = 2 NonSeedType = 2 \| \| --- \| --- \| --- \| --- \| --- \| --- \| | \| 158 \| WBGene00001921:his-47(1439) \| ``` log2FC=-0.231 padj=0.443 ``` \| [seedLoc = 163,168] Seq=ATGCAATATTGTTATAAAGTTTTCTACCTTC \| ```          C       TTTTGAAATATTGTTA            mRNA  3'  TTCCATC                TAACGTA     miRNA 5'  GAGGTAG                GTTGTAT              T       TAG                    AGTT           32111111333545555555455521112116666 ``` \| SeedType = 3 NonSeedType = 1 \| \| --- \| --- \| --- \| --- \| --- \| --- \| | \| 159 \| WBGene00001975:hmg-5(1476) \| ``` log2FC=-0.071 padj=0.949 ``` \| [seedLoc = 70,75] Seq=CGATTCATGACACTTTTCTCAACTACCTCT \| ```          T        ACTCTTT  A     CTTAGC mRNA  3'  CTCCATCA       TC CAGTA       miRNA 5'  GAGGTAGT       AG GTTGT                T                       ATAGTT           311111111555555511511221333333 ``` \| SeedType = 1 NonSeedType = -1 \| \| --- \| --- \| --- \| --- \| --- \| --- \| | \| 160 \| WBGene00001980:hmr-1(1482) \| ``` log2FC=-0.11 padj=0.913 ``` \| [seedLoc = 359,364] Seq=TCTTACATAATCACAAGTACCTCT \| ```          T      G ACA      CATTCT mRNA  3'  CTCCAT A   CTAATA       miRNA 5'  GAGGTA T   GGTTGT                T      G A        ATAGTT           311111131355121121333333 ``` \| SeedType = 1 NonSeedType = 2 \| \| --- \| --- \| --- \| --- \| --- \| --- \| | \| 161 \| WBGene00001994:hpk-1(1494) \| ``` log2FC=0.048 padj=0.977 ``` \| [seedLoc = 905,910] Seq=CTGTGCAGTTGTTTCAAAACTGTCTGCCTCC \| ```          C       TG   AAACTTTGTTG          mRNA  3'  CTCCGTC  TCA           ACGTGTC   miRNA 5'  GAGGTAG  GGT           TGTATAG            T       TA                     TT           311112113321155555554554112121166 ``` \| SeedType = 2 NonSeedType = 2 \| \| --- \| --- \| --- \| --- \| --- \| --- \| | \| 162 \| WBGene00001999:hrp-1(1500) \| ``` log2FC=-0.148 padj=0.823 ``` \| [seedLoc = 92,97] Seq=TCTTCTCAACCATTTTTGCTCTGCTATCTCA \| ```                    CTCGTTTTTA     TCT  T  mRNA  3' ACTCTATCGT          CCAAC   TC   miRNA 5' TGAGGTAGTA          GGTTG   AG                                     TAT  TT           11112111215554555555111113331136 ``` \| SeedType = 2 NonSeedType = 1 \| \| --- \| --- \| --- \| --- \| --- \| --- \| | \| 163 \| WBGene00002003:hse-5(1505) \| ``` log2FC=0.413 padj=0.481 ``` \| [seedLoc = 29,34] Seq=ATCATGACACCGGGTACTACTTCT \| ```          T         GGG      GTACTA mRNA  3'  CTTCATCAT   CC ACA       miRNA 5'  GAGGTAGTA   GG TGT                T              T   ATAGTT           3112111111444116111333333 ``` \| SeedType = 3 NonSeedType = -1 \| \| --- \| --- \| --- \| --- \| --- \| --- \| | \|  \|  \|  \| [seedLoc = 101,106] Seq=TTCTGTTCAATTTATAATCCTGCCTCC \| ```          C       CTAAT       T    TT mRNA  3'  CTCCGTC     ATTTAAC TGTC   miRNA 5'  GAGGTAG     TAGGTTG ATAG            T                   T    TT           311112115555511221113121133 ``` \| SeedType = 2 NonSeedType = 1 \| \| --- \| --- \| --- \| --- \| --- \| --- \| | \| 164 \| WBGene00002036:hum-2(1530) \| ``` log2FC=-0.02 padj=0.99 ``` \| [seedLoc = 66,71] Seq=TTCAATTCATTCTCTACCTTT \| ```          T           T  TTAACTT mRNA  3'  TTCCATC TCT AC        miRNA 5'  GAGGTAG AGG TG                 T       T   T  TATAGTT           3211111161123113333333 ``` \| SeedType = 3 NonSeedType = -1 \| \| --- \| --- \| --- \| --- \| --- \| --- \| | \| 165 \| WBGene00002044:hyl-2(1538) \| ``` log2FC=0.181 padj=0.794 ``` \| [seedLoc = 201,206] Seq=TTCTAGAGCCTTTTGTCTTTTTACTACCTTA \| ```                   TTTTTCTGTTT     G  CTT   mRNA  3' ATTCCATCA           TCCGA AT      miRNA 5' TGAGGTAGT           AGGTT TA                                        G  TAGTT           121111111555555545551112131133366 ``` \| SeedType = 3 NonSeedType = 2 \| \| --- \| --- \| --- \| --- \| --- \| --- \| | \| 166 \| WBGene00002051:ifa-3(1544) \| ``` log2FC=0.968 padj=0.1 over-expressed ``` \| [seedLoc = 212,217] Seq=TTCATTATGACCTAAACTACTTCT \| ```          T        AA        TACTT mRNA  3'  CTTCATCA  TCCAGTAT      miRNA 5'  GAGGTAGT  AGGTTGTA               T                  TAGTT           311211111551111221133333 ``` \| SeedType = 3 NonSeedType = 1 \| \| --- \| --- \| --- \| --- \| --- \| --- \| | \| 167 \| WBGene00002052:ifa-4(1545) \| ``` log2FC=-0.198 padj=0.871 ``` \| [seedLoc = 36,42] Seq=TCCTATTGAACCCGGAATTACTCTCG \| ```          G   T     AGGC    GT    CT mRNA  3'  CTC CATTA    CCAA  TATC   miRNA 5'  GAG GTAGT    GGTT  ATAG            T         A       GT    TT           31115111213445111133111133 ``` \| SeedType = 2 NonSeedType = 1 \| \| --- \| --- \| --- \| --- \| --- \| --- \| | \| 168 \| WBGene00002053:ifb-1(1546) \| ``` log2FC=0.163 padj=0.837 ``` \| [seedLoc = 348,353] Seq=TTCTATTCAATGTTTTCTACTTCT \| ```          T       TTT G    T    TT mRNA  3'  CTTCATC   T TAAC TATC   miRNA 5'  GAGGTAG   A GTTG ATAG            T       T   G    T    TT           311211113551321113111133 ``` \| SeedType = 3 NonSeedType = 1 \| \| --- \| --- \| --- \| --- \| --- \| --- \| | \| 169 \| WBGene00002056:ifc-2(1550) \| ``` log2FC=-0.215 padj=0.428 ``` \| [seedLoc = 355,360] Seq=TGATTGTAAATCTCTATTCTATCTCT \| ```          T       TT   T TAA       T mRNA  3'  CTCTATC  ATC C   ATGTTAG  miRNA 5'  GAGGTAG  TAG G   TATAGTT           T              TTG                   31112111551115133311212125 ``` \| SeedType = 2 NonSeedType = -1 \| \| --- \| --- \| --- \| --- \| --- \| --- \| | \| 170 \| WBGene00002066:ifg-1(1562) \| ``` log2FC=0.058 padj=0.949 ``` \| [seedLoc = 896,902] Seq=CAATTGGTTAATTTTAATATCCTCT \| ```          T    T  AAT      TG     C mRNA  3'  CTCC AT   TTTAAT  GTTAA  miRNA 5'  GAGG TA   AGGTTG  TAGTT           T       GT       TA                 3111151133512211233212115 ``` \| SeedType = 2 NonSeedType = 2 \| \| --- \| --- \| --- \| --- \| --- \| --- \| | \| 171 \| WBGene00002073:ima-2(1569) \| ``` log2FC=0.121 padj=0.866 ``` \| [seedLoc = 89,95] Seq=TGTTTTTTTTAGAATTACCCTCA \| ```             C      AGATTTTTTTTGT mRNA  3' ACT CCATTA              miRNA 5' TGA GGTAGT                                 AGGTTGTATAGTT           11151111213333333333333 ``` \| SeedType = 3 NonSeedType = -1 \| \| --- \| --- \| --- \| --- \| --- \| --- \| | \| 172 \| WBGene00002077:imb-3(1573) \| ``` log2FC=0.161 padj=0.766 ``` \| [seedLoc = 92,98] Seq=CGCTTGTTTCGCGGCTATCCTCC \| ```          C    T    GCGCTTTGT   C mRNA  3'  CTCC ATCG         TCG  miRNA 5'  GAGG TAGT         AGT           T         AGGTTGTAT   T           31111511123333333331123 ``` \| SeedType = 2 NonSeedType = -1 \| \| --- \| --- \| --- \| --- \| --- \| --- \| | \|  \|  \|  \| [seedLoc = 208,214] Seq=TGTTGTGTTGAATATTATCCTCA \| ```               T     AAGTT     GT mRNA  3' ACTCC ATTAT     GTGTT   miRNA 5' TGAGG TAGTA     TATAG                       GGTTG     TT           11111511211333332121233 ``` \| SeedType = 2 NonSeedType = -1 \| \| --- \| --- \| --- \| --- \| --- \| --- \| | \|  \|  \|  \| [seedLoc = 390,396] Seq=TTCTGTCACTTGAAGACTATCCTCG \| ```          G    T    GAAGTT  C    TT mRNA  3'  CTCC ATCA      CA TGTC   miRNA 5'  GAGG TAGT      GT ATAG            T         AGGTT        TT           3111151111333335115121133 ``` \| SeedType = 2 NonSeedType = -1 \| \| --- \| --- \| --- \| --- \| --- \| --- \| | \| 173 \| WBGene00002088:ins-5(1584) \| ``` log2FC=0.996 padj=0.129 ``` \| [seedLoc = 96,101] Seq=CATTATTTTCCTCTGTTTCAACTATCTCA \| ```                   ACTTTGTC   TTT      C mRNA  3' ACTCTATCA        TCC    TATTA  miRNA 5' TGAGGTAGT        AGG    ATAGT                               TTGT     T           111121111555554551113336111213 ``` \| SeedType = 2 NonSeedType = -1 \| \| --- \| --- \| --- \| --- \| --- \| --- \| | \|  \|  \|  \| [seedLoc = 115,120] Seq=TCTCATATTTCCAATTCTATCTCA \| ```                  TTAA  TT    CTCT mRNA  3' ACTCTATC    CC  TATA     miRNA 5' TGAGGTAG    GG  GTAT                      TA    TT    AGTT           111121113355113321113333 ``` \| SeedType = 2 NonSeedType = -1 \| \| --- \| --- \| --- \| --- \| --- \| --- \| | \| 174 \| WBGene00002103:ins-20(1592) \| ``` log2FC=0.255 padj=1.0 ``` \| [seedLoc = 37,42] Seq=TTTTTCTCGTGCTATCTACCTCA \| ```                  T   GTGCTCTTTTT mRNA  3' ACTCCATC ATC            miRNA 5' TGAGGTAG TAG                                 GTTGTATAGTT           11111111511133333333333 ``` \| SeedType = 1 NonSeedType = -1 \| \| --- \| --- \| --- \| --- \| --- \| --- \| | \| 175 \| WBGene00002116:ins-33(1601) \| ``` log2FC=0.796 padj=0.515 ``` \| [seedLoc = 52,57] Seq=CATAATATACCAATCTACCTCT \| ```          T       TAA        ATAC mRNA  3'  CTCCATC   CC ATATA     miRNA 5'  GAGGTAG   GG TGTAT              T       TA   T     AGTT           31111111335116121113333 ``` \| SeedType = 1 NonSeedType = -1 \| \| --- \| --- \| --- \| --- \| --- \| --- \| | \| 176 \| WBGene00002118:ins-35(1603) \| ``` log2FC=1.723 padj=0.021 over-expressed ``` \| [seedLoc = 35,40] Seq=TTCTGTTGCGTTTATTGCTACCTCT \| ```          T          ATTT   T    TT mRNA  3'  CTCCATCGTT    GCG TGTC   miRNA 5'  GAGGTAGTAG    TGT ATAG            T          GT          TT           3111111121233552125121133 ``` \| SeedType = 1 NonSeedType = -1 \| \| --- \| --- \| --- \| --- \| --- \| --- \| | \| 177 \| WBGene00002123:inx-1(1604) \| ``` log2FC=0.126 padj=0.973 ``` \| [seedLoc = 319,324] Seq=TTGAGCACAGCTCTTGTTCTACCTTT \| ```          T       TTGTTC      CGAGTT mRNA  3'  TTCCATC      TCGACA       miRNA 5'  GAGGTAG      GGTTGT                T       TA          ATAGTT           32111111334555212111333333 ``` \| SeedType = 3 NonSeedType = 2 \| \| --- \| --- \| --- \| --- \| --- \| --- \| | \| 178 \| WBGene00002129:inx-7(1611) \| ``` log2FC=-0.688 padj=0.081 ``` \| [seedLoc = 91,97] Seq=ATTTCTACGGTCACATACACTCT \| ```          T   A   A          CTTTA mRNA  3'  CTC CAT CA CTGGCAT      miRNA 5'  GAG GTA GT GGTTGTA               T          A       TAGTT           311151115116122211133333 ``` \| SeedType = 2 NonSeedType = 2 \| \| --- \| --- \| --- \| --- \| --- \| --- \| | \| 179 \| WBGene00002130:inx-8(1612) \| ``` log2FC=0.03 padj=0.988 ``` \| [seedLoc = 102,108] Seq=AACGTTCGACCCGTGAATAATACTACCATCG \| ```          G  A       AATAAGTGC     TTG     mRNA  3'  CT CCATCAT         CCAGC    CAA miRNA 5'  GA GGTAGTA         GGTTG    GTT          T                        TATA              31151111111555554545111213336111 ``` \| SeedType = 3 NonSeedType = 1 \| \| --- \| --- \| --- \| --- \| --- \| --- \| | \| 180 \| WBGene00002132:inx-10(1614) \| ``` log2FC=-0.218 padj=0.885 ``` \| [seedLoc = 84,89] Seq=GGTTTTGCAGTCTAATCTCTACCTCT \| ```          T       TCTA         TTTGG mRNA  3'  CTCCATC    ATCTGACGT      miRNA 5'  GAGGTAG    TAGGTTGTA               T                    TAGTT           31111111555511122112133333 ``` \| SeedType = 1 NonSeedType = 2 \| \| --- \| --- \| --- \| --- \| --- \| --- \| | \| 181 \| WBGene00002148:gon-14(1630) \| ``` log2FC=-0.115 padj=0.897 ``` \| [seedLoc = 104,109] Seq=CAGCTCATACATCTGTACCTCA \| ```                            C    C mRNA  3' ACTCCAT GTCT ACATA TCGA  miRNA 5' TGAGGTA TAGG TGTAT AGTT                  G    T                      111111162112611111511215 ``` \| SeedType = 1 NonSeedType = -1 \| \| --- \| --- \| --- \| --- \| --- \| --- \| | \| 182 \| WBGene00002151:irk-3(1634) \| ``` log2FC=-0.066 padj=0.992 ``` \| [seedLoc = 397,402] Seq=ATGTTTCTGACTTAGAGACTACTTCA \| ```                   GAGA     TCTTTGTA mRNA  3' ACTTCATCA    TTCAG         miRNA 5' TGAGGTAGT    AGGTT                                    GTATAGTT           11121111145451211233333333 ``` \| SeedType = 3 NonSeedType = 2 \| \| --- \| --- \| --- \| --- \| --- \| --- \| | \| 183 \| WBGene00002163:ist-1(1639) \| ``` log2FC=-0.083 padj=0.977 ``` \| [seedLoc = 168,173] Seq=CCAATCTCAACCTTTCCTTCATTACCTCT \| ```          T        CTTCCTT      TCTAACC mRNA  3'  CTCCATTA       TCCAAC        miRNA 5'  GAGGTAGT       AGGTTG                 T                     TATAGTT           31111112155555551111113333333 ``` \| SeedType = 1 NonSeedType = 1 \| \| --- \| --- \| --- \| --- \| --- \| --- \| | \| 184 \| WBGene00002169:isw-1(1640) \| ``` log2FC=0.062 padj=0.956 ``` \| [seedLoc = 60,66] Seq=ATTAATGTCGATCTCTATCCTCC \| ```          C    T          T   ATTA mRNA  3'  CTCC ATC TCTAGC GTA     miRNA 5'  GAGG TAG AGGTTG TAT              T        T          AGTT           311115111611212152113333 ``` \| SeedType = 2 NonSeedType = 2 \| \| --- \| --- \| --- \| --- \| --- \| --- \| | \| 185 \| WBGene00002183:kat-1(1652) \| ``` log2FC=0.073 padj=0.929 ``` \| [seedLoc = 122,127] Seq=TATTATTTTGATTATCTACTTCT \| ```          T       TA     TT     T mRNA  3'  CTTCATC  TTAGT  TATTA  miRNA 5'  GAGGTAG  GGTTG  ATAGT           T       TA     T      T           31121111332212235111213 ``` \| SeedType = 3 NonSeedType = 2 \| \| --- \| --- \| --- \| --- \| --- \| --- \| | \| 186 \| WBGene00002189:kin-1(1657) \| ``` log2FC=-0.027 padj=0.986 ``` \| [seedLoc = 47,52] Seq=TGCATATATCTCCACTTCTATCTCT \| ```          T       TTCA  TC     A  T mRNA  3'  CTCTATC    CC  TATAT CG  miRNA 5'  GAGGTAG    GG  GTATA GT           T       TA    TT        T           3111211133551133211115123 ``` \| SeedType = 2 NonSeedType = -1 \| \| --- \| --- \| --- \| --- \| --- \| --- \| | \| 187 \| WBGene00002192:kin-4(1661) \| ``` log2FC=-0.019 padj=0.991 ``` \| [seedLoc = 52,56] Seq=TCATATCAACCAACTACCACAA \| ```          A  A      A         ACT mRNA  3'  AC CCATCA CCAAC TAT    miRNA 5'  TG GGTAGT GGTTG ATA                A      A     T   GTT           51131111113111116111333 ``` \| SeedType = 3 NonSeedType = 1 \| \| --- \| --- \| --- \| --- \| --- \| --- \| | \| 188 \| WBGene00002202:kin-19(1669) \| ``` log2FC=-0.015 padj=0.992 ``` \| [seedLoc = 214,220] Seq=TAACTAAAGATGAATACTCTCT \| ```          T   T   AAG    AA      T mRNA  3'  CTC CAT    TAG   ATCAA  miRNA 5'  GAG GTA    GTT   TAGTT           T       GTAG   GTA                 311151113336212336111115 ``` \| SeedType = 2 NonSeedType = -1 \| \| --- \| --- \| --- \| --- \| --- \| --- \| | \|  \|  \|  \| [seedLoc = 593,597] Seq=AAATGCCTCTCAATACCCTCC \| ```          C  C    AAC  TC    AA   mRNA  3'  CT CCAT   TC  CGTA     miRNA 5'  GA GGTA   GG  GTAT              T       GTA  TT    AGTT           31151111333213312113366 ``` \| SeedType = 3 NonSeedType = -1 \| \| --- \| --- \| --- \| --- \| --- \| --- \| | \| 189 \| WBGene00002203:kin-20(1670) \| ``` log2FC=-0.253 padj=0.364 ``` \| [seedLoc = 652,657] Seq=AAACTGCCTTCATCCTTTTCGGACTATCTCT \| ```          T        GGCTTT   T  TTCC     A mRNA  3'  CTCTATCA      TCC AC    GTCAA  miRNA 5'  GAGGTAGT      AGG TG    TAGTT           T                 T  TA                   3111211114455551113113355211115 ``` \| SeedType = 2 NonSeedType = -1 \| \| --- \| --- \| --- \| --- \| --- \| --- \| | \| 190 \| WBGene00002210:kin-29(1675) \| ``` log2FC=0.147 padj=0.887 ``` \| [seedLoc = 147,152] Seq=TGATTTAGCGAAATTCTACTTCG \| ```          G       TTAA    AT    T mRNA  3'  CTTCATC    AGCG  TTAG  miRNA 5'  GAGGTAG    TTGT  AGTT           T       TAGG    AT                31121111333312123312125 ``` \| SeedType = 3 NonSeedType = -1 \| \| --- \| --- \| --- \| --- \| --- \| --- \| | \| 191 \| WBGene00002216:klp-3(1681) \| ``` log2FC=0.356 padj=0.323 ``` \| [seedLoc = 29,34] Seq=TCAAATATTTCATCTCTACTTCT \| ```          T            CTT    AACT mRNA  3'  CTTCATC TCTA   TATA     miRNA 5'  GAGGTAG AGGT   GTAT              T       T    T      AGTT           311211116112135521113333 ``` \| SeedType = 3 NonSeedType = 2 \| \| --- \| --- \| --- \| --- \| --- \| --- \| | \| 192 \| WBGene00002224:klp-13(1690) \| ``` log2FC=-0.658 padj=1.0 ``` \| [seedLoc = 438,443] Seq=TAGAGGATCGCCTTATGTCTGCCTCA \| ```                  TGTAT    CTAGGAGAT mRNA  3' ACTCCGTC     TCCG          miRNA 5' TGAGGTAG     AGGT                           T        TGTATAGTT           11111211345551112333333333 ``` \| SeedType = 2 NonSeedType = 2 \| \| --- \| --- \| --- \| --- \| --- \| --- \| | \| 193 \| WBGene00002238:kars-1(1703) \| ``` log2FC=0.011 padj=0.994 ``` \| [seedLoc = 110,115] Seq=CTCTATTGCTTTTCTCTCCTATCTCG \| ```          G       C  T TTTT  T    TC mRNA  3'  CTCTATC TC C    CG TATC   miRNA 5'  GAGGTAG AG G    GT ATAG            T       T    TT         TT           31112111311513355125111133 ``` \| SeedType = 2 NonSeedType = -1 \| \| --- \| --- \| --- \| --- \| --- \| --- \| | \| 194 \| WBGene00002251:lat-1(1716) \| ``` log2FC=-0.097 padj=0.938 ``` \| [seedLoc = 198,203] Seq=GAGCCATGTCAGAAATTACCTCG \| ```          G        AA    T   C   G mRNA  3'  CTCCATTA   GAC GTA CGA  miRNA 5'  GAGGTAGT   TTG TAT GTT           T        AGG       A               311111121336211521131214 ``` \| SeedType = 1 NonSeedType = -1 \| \| --- \| --- \| --- \| --- \| --- \| --- \| | \|  \|  \|  \| [seedLoc = 445,450] Seq=TATTTTACTTTTCTTGTTATTTTACCTCC \| ```          C       TTATT    TTTT   T   T mRNA  3'  CTCCATT     GTTC    CAT TTA  miRNA 5'  GAGGTAG     TAGG    GTA AGT           T                TT     T   T           31111112555552121335511131213 ``` \| SeedType = 1 NonSeedType = -1 \| \| --- \| --- \| --- \| --- \| --- \| --- \| | \| 195 \| WBGene00002263:lea-1(1730) \| ``` log2FC=-0.51 padj=0.011 ``` \| [seedLoc = 120,126] Seq=AGTAATTTTATCGCTACCCTCT \| ```          T  C          TTTTAATGA mRNA  3'  CT CCATCG CTA          miRNA 5'  GA GGTAGT GGT                   T         A   TGTATAGTT           31151111126121333333333 ``` \| SeedType = 3 NonSeedType = 2 \| \| --- \| --- \| --- \| --- \| --- \| --- \| | \| 196 \| WBGene00002266:lec-3(1734) \| ``` log2FC=-0.266 padj=0.65 ``` \| [seedLoc = 49,54] Seq=CCCTAGTTTCTTTCTATCTCT \| ```          T       T   TTTG    CC mRNA  3'  CTCTATC TTC     ATC   miRNA 5'  GAGGTAG AGG     TAG            T       T   TTGTA   TT           3111211131213333611133 ``` \| SeedType = 2 NonSeedType = -1 \| \| --- \| --- \| --- \| --- \| --- \| --- \| | \| 197 \| WBGene00002276:lem-3(1744) \| ``` log2FC=-0.152 padj=0.843 ``` \| [seedLoc = 42,47] Seq=GTTTATATTTCAAATTTTACCTCA \| ```                  TTAAA TT      TG mRNA  3' ACTCCATT     C  TATATT   miRNA 5' TGAGGTAG     G  GTATAG                    TAG   TT      TT           111111123335513321111233 ``` \| SeedType = 1 NonSeedType = -1 \| \| --- \| --- \| --- \| --- \| --- \| --- \| | \| 198 \| WBGene00002295:let-19(1749) \| ``` log2FC=0.114 padj=0.918 ``` \| [seedLoc = 185,190] Seq=CCAAATATGATCGTGTTTCTATCTCA \| ```                  TTTGTG        AACC mRNA  3' ACTCTATC      CTAGTATA     miRNA 5' TGAGGTAG      GGTTGTAT                      TA            AGTT           11112111335454121221113333 ``` \| SeedType = 2 NonSeedType = 2 \| \| --- \| --- \| --- \| --- \| --- \| --- \| | \| 199 \| WBGene00002297:ect-2(1750) \| ``` log2FC=-0.071 padj=0.946 ``` \| [seedLoc = 83,88] Seq=CTTTTCCGAGCTTCATACCTCC \| ```          C      AC     GCCTTTTC mRNA  3'  CTCCAT  TTCGA         miRNA 5'  GAGGTA  AGGTT                  T      GT     GTATAGTT           3111111331212133333333 ``` \| SeedType = 1 NonSeedType = 2 \| \| --- \| --- \| --- \| --- \| --- \| --- \| | \| 200 \| WBGene00002632:let-413(1762) \| ``` log2FC=-0.164 padj=0.78 ``` \| [seedLoc = 549,555] Seq=ATATTTCTTCCTTACTAACCTCC \| ```          C    A       CTTCTTTATA mRNA  3'  CTCC ATCATTC           miRNA 5'  GAGG TAGTAGG                    T            TTGTATAGTT           31111511111213333333333 ``` \| SeedType = 2 NonSeedType = -1 \| \| --- \| --- \| --- \| --- \| --- \| --- \| | \| 201 \| WBGene00002694:let-502(1764) \| ``` log2FC=-0.046 padj=0.97 ``` \| [seedLoc = 333,338] Seq=TTTCCTTTGGCGTCTACCTTT \| ```          T         G   TTTCCTTT mRNA  3'  TTCCATC T CGG         miRNA 5'  GAGGTAG A GTT                  T       T G   GTATAGTT           3211111161312233333333 ``` \| SeedType = 3 NonSeedType = -1 \| \| --- \| --- \| --- \| --- \| --- \| --- \| | \| 202 \| WBGene00002717:let-526(1766) \| ``` log2FC=-0.104 padj=0.907 ``` \| [seedLoc = 32,37] Seq=TCATTTCTAATCATACCTCT \| ```          T      A       CTTTACT mRNA  3'  CTCCAT   CTAAT        miRNA 5'  GAGGTA   GGTTG                 T      GTA     TATAGTT           3111111366121123333333 ``` \| SeedType = 1 NonSeedType = 2 \| \| --- \| --- \| --- \| --- \| --- \| --- \| | \| 203 \| WBGene00002881:let-756(1774) \| ``` log2FC=-0.01 padj=0.997 ``` \| [seedLoc = 226,231] Seq=TTCTAATTTTCTATTTCTACCTTC \| ```          C       TTT    TTTA    TT mRNA  3'  TTCCATC   ATCT     ATC   miRNA 5'  GAGGTAG   TAGG     TAG            T              TTGTA   TT           3211111155511123333611133 ``` \| SeedType = 3 NonSeedType = -1 \| \| --- \| --- \| --- \| --- \| --- \| --- \| | \| 204 \| WBGene00002915:let-805(1777) \| ``` log2FC=0.301 padj=0.654 ``` \| [seedLoc = 118,124] Seq=TTCTCTAAAACCTAAAAATTCCCTACCTAC \| ```          CA      CCTTAAAA      AATCTCTT mRNA  3'   TCCATC        ATCCAA         miRNA 5'   AGGTAG        TAGGTT                  TG                    GTATAGTT           331111115555555511111133333333 ``` \| SeedType = 3 NonSeedType = 1 \| \| --- \| --- \| --- \| --- \| --- \| --- \| | \| 205 \| WBGene00002977:lev-10(1785) \|  \| [seedLoc = 139,144] Seq=CAACTGACGATTCTACCTTT \| ```          T                     C mRNA  3'  TTCCATC  TTAGCA GTCAA  miRNA 5'  GAGGTAG  GGTTGT TAGTT           T       TA      A                 32111111662212116211115 ``` \| SeedType = 3 NonSeedType = 2 \| \| --- \| --- \| --- \| --- \| --- \| --- \| | \|  \|  \|  \| [seedLoc = 358,363] Seq=ATGAGTGCAAGTTCCGATTTACCTCG \| ```          G       TAG  TTG      AGTA mRNA  3'  CTCCATT   CC   AACGTG     miRNA 5'  GAGGTAG   GG   TTGTAT              T       TA            AGTT           31111112334115541112123333 ``` \| SeedType = 1 NonSeedType = 1 \| \| --- \| --- \| --- \| --- \| --- \| --- \| | \| 206 \| WBGene00002981:lgg-2(1791) \| ``` log2FC=-0.738 padj=0.01 ``` \| [seedLoc = 197,201] Seq=CCCCTCGCAATTCCCCACCTCAA \| ```          A      C CC      CTCCCC mRNA  3'  ACTCCA C  TTAACG       miRNA 5'  TGAGGT G  GGTTGT                       A TA      ATAGTT           51111113133221112333333 ``` \| SeedType = 2 NonSeedType = 1 \| \| --- \| --- \| --- \| --- \| --- \| --- \| | \| 207 \| WBGene00002985:lig-1(1796) \| ``` log2FC=-0.053 padj=0.973 ``` \| [seedLoc = 83,88] Seq=AATTTTCTAAGTAAATACCTCA \| ```                 AAA G   CTTTTAA mRNA  3' ACTCCAT   T AAT        miRNA 5' TGAGGTA   G TTG                        GTA G   TATAGTT           1111111333231123333333 ``` \| SeedType = 1 NonSeedType = -1 \| \| --- \| --- \| --- \| --- \| --- \| --- \| | \| 208 \| WBGene00002986:lig-4(1797) \| ``` log2FC=-0.025 padj=0.989 ``` \| [seedLoc = 54,59] Seq=TCTTTATTTCCTTTTTGATAGATCTACTTCA \| ```                  TAGATAGTTTT   TT   T  T  mRNA  3' ACTTCATC           TCC  TAT TC   miRNA 5' TGAGGTAG           AGG  GTA AG                    T             TT   T  TT           11121111354555455551113321131136 ``` \| SeedType = 3 NonSeedType = -1 \| \| --- \| --- \| --- \| --- \| --- \| --- \| | \| 209 \| WBGene00003014:lin-28(1821) \| ``` log2FC=0.119 padj=0.925 ``` \| [seedLoc = 315,320] Seq=AAACCATACTACCACCTACCTCC \| ```          C       CA   T    C   A mRNA  3'  CTCCATC  CCA CATA CAA  miRNA 5'  GAGGTAG  GGT GTAT GTT           T       TA   T    A               31111111331113111131115 ``` \| SeedType = 1 NonSeedType = 1 \| \| --- \| --- \| --- \| --- \| --- \| --- \| | \| 210 \| WBGene00003021:lin-36(1826) \| ``` log2FC=-0.045 padj=0.977 ``` \| [seedLoc = 93,98] Seq=TTCTTTCATGTGAACTATCTCT \| ```          T        AGTGTACTTTCTT mRNA  3'  CTCTATCA              miRNA 5'  GAGGTAGT                       T        AGGTTGTATAGTT           3111211113333333333333 ``` \| SeedType = 2 NonSeedType = -1 \| \| --- \| --- \| --- \| --- \| --- \| --- \| | \| 211 \| WBGene00003026:lin-41(1830) \| ``` log2FC=0.298 padj=0.293 ``` \| [seedLoc = 702,708] Seq=TTTTATACAACCGTTCTACACTCA \| ```              A    TTG          TT mRNA  3' ACTC CATC   CCAACATATT   miRNA 5' TGAG GTAG   GGTTGTATAG                     TA           TT           111151111334111111111233 ``` \| SeedType = 2 NonSeedType = 1 \| \| --- \| --- \| --- \| --- \| --- \| --- \| | \|  \|  \|  \| [seedLoc = 751,756] Seq=TTTTATACAACCATTCTGCCTCT \| ```          T       TTA          TT mRNA  3'  CTCCGTC   CCAACATATT   miRNA 5'  GAGGTAG   GGTTGTATAG            T       TA           TT           31111211335111111111233 ``` \| SeedType = 2 NonSeedType = 1 \| \| --- \| --- \| --- \| --- \| --- \| --- \| | \| 212 \| WBGene00003031:lin-46(1833) \| ``` log2FC=-0.696 padj=0.71 ``` \| [seedLoc = 53,58] Seq=TACTCGGTGCTTTTATACCTCC \| ```          C      ATT    TGGC    T mRNA  3'  CTCCAT   TTCG     TCA  miRNA 5'  GAGGTA   AGGT     AGT           T      GT     TGTAT   T           31111113351212333361113 ``` \| SeedType = 1 NonSeedType = 2 \| \| --- \| --- \| --- \| --- \| --- \| --- \| | \| 213 \| WBGene00003052:lmn-1(1848) \| ``` log2FC=-0.226 padj=0.444 ``` \| [seedLoc = 47,52] Seq=CCATTTGGACCTCATTTTTACATCTATCTCT \| ```          T       TACATTTTTAC     GTTTACC  mRNA  3'  CTCTATC           TCCAG         miRNA 5'  GAGGTAG           AGGTT                  T       T               GTATAGTT           31112111355555555551111233333336 ``` \| SeedType = 2 NonSeedType = 1 \| \| --- \| --- \| --- \| --- \| --- \| --- \| | \| 214 \| WBGene00003053:lmp-1(1849) \| ``` log2FC=0.428 padj=0.107 ``` \| [seedLoc = 202,207] Seq=TTTCAAACGAGTAGTACCTCA \| ```                 G  G     AACTTT mRNA  3' ACTCCAT AT  AGCA       miRNA 5' TGAGGTA TA  TTGT                       G  GG    ATAGTT           1111111311361211333333 ``` \| SeedType = 1 NonSeedType = -1 \| \| --- \| --- \| --- \| --- \| --- \| --- \| | \| 215 \| WBGene00003056:lon-2(1851) \| ``` log2FC=0.047 padj=0.982 ``` \| [seedLoc = 352,357] Seq=GTCCGTTTGACACATACCTCA \| ```                 ACA    TTTGCCTG mRNA  3' ACTCCAT    CAG         miRNA 5' TGAGGTA    GTT                         GTAG   GTATAGTT           1111111333611233333333 ``` \| SeedType = 1 NonSeedType = -1 \| \| --- \| --- \| --- \| --- \| --- \| --- \| | \| 216 \| WBGene00003082:lsm-8(1873) \| ``` log2FC=0.168 padj=0.811 ``` \| [seedLoc = 148,154] Seq=TCCTTGACGGTTTTGTATTCATACCATCG \| ```          G  A    ACT   G TTT    GT  CT mRNA  3'  CT CCAT   TAT T   GGCA  TC   miRNA 5'  GA GGTA   GTA G   TTGT  AG            T             G        AT  TT           31151111555211325552211331133 ``` \| SeedType = 3 NonSeedType = -1 \| \| --- \| --- \| --- \| --- \| --- \| --- \| | \| 217 \| WBGene00003087:lsy-2(1880) \| ``` log2FC=0.005 padj=0.997 ``` \| [seedLoc = 151,156] Seq=TTTTGGTTCTTTTCTTACCTCG \| ```          G       CTTTTCTTGGTTTT mRNA  3'  CTCCATT               miRNA 5'  GAGGTAG                        T       TAGGTTGTATAGTT           3111111233333333333333 ``` \| SeedType = 1 NonSeedType = -1 \| \| --- \| --- \| --- \| --- \| --- \| --- \| | \| 218 \| WBGene00003102:mab-5(1891) \| ``` log2FC=0.192 padj=0.953 ``` \| [seedLoc = 41,46] Seq=ATTTGAATTTCTTTTTACCTCG \| ```          G       T   TT  AGTTTA mRNA  3'  CTCCATT TTC  TA       miRNA 5'  GAGGTAG AGG  GT                T       T   TT  ATAGTT           3111111231213321333333 ``` \| SeedType = 1 NonSeedType = -1 \| \| --- \| --- \| --- \| --- \| --- \| --- \| | \| 219 \| WBGene00003111:mab-20(1895) \| ``` log2FC=0.268 padj=0.866 ``` \| [seedLoc = 89,95] Seq=GAAAATGCCTCCCTACCTGC \| ```          CG          TC    AAAG mRNA  3'   TCCATC  CC  CGTA     miRNA 5'   AGGTAG  GG  GTAT              TG      TA  TT    AGTT           3311111166113312113333 ``` \| SeedType = 3 NonSeedType = -1 \| \| --- \| --- \| --- \| --- \| --- \| --- \| | \| 220 \| WBGene00003144:max-2(1911) \| ``` log2FC=-0.069 padj=0.955 ``` \| [seedLoc = 900,906] Seq=CATCCCCCAACAACTACCTGC \| ```          CG       A     CCCCTAC mRNA  3'   TCCATCA  CAAC        miRNA 5'   AGGTAGT  GTTG                 TG       AG    TATAGTT           3311111113611113333333 ``` \| SeedType = 3 NonSeedType = 1 \| \| --- \| --- \| --- \| --- \| --- \| --- \| | \| 221 \| WBGene00003151:mca-1(1916) \| ``` log2FC=0.148 padj=0.87 ``` \| [seedLoc = 196,202] Seq=AGAAACATGACCAAACATATCCTCC \| ```          C    T  A  AA      CAAAGA mRNA  3'  CTCC AT CA  CCAGTA       miRNA 5'  GAGG TA GT  GGTTGT                T          A       ATAGTT           3111151151135111221333333 ``` \| SeedType = 2 NonSeedType = 1 \| \| --- \| --- \| --- \| --- \| --- \| --- \| | \| 222 \| WBGene00003153:mca-3(1917) \| ``` log2FC=0.141 padj=0.798 ``` \| [seedLoc = 432,437] Seq=TCTTTTCTTTTGTCCTACTTCG \| ```          G       CTGTTTTCTTTTCT mRNA  3'  CTTCATC               miRNA 5'  GAGGTAG                        T       TAGGTTGTATAGTT           3112111133333333333333 ``` \| SeedType = 3 NonSeedType = -1 \| \| --- \| --- \| --- \| --- \| --- \| --- \| | \| 223 \| WBGene00003161:mdf-2(1929) \| ``` log2FC=-0.054 padj=0.973 ``` \| [seedLoc = 35,40] Seq=GTCTGTTTTCTCTGCTACCTCT \| ```          T           CTTT    TG mRNA  3'  CTCCATCGTCT    TGTC   miRNA 5'  GAGGTAGTAGG    ATAG            T           TTGT    TT           3111111121123333121133 ``` \| SeedType = 1 NonSeedType = -1 \| \| --- \| --- \| --- \| --- \| --- \| --- \| | \| 224 \| WBGene00003172:mec-8(1943) \| ``` log2FC=-0.087 padj=0.967 ``` \| [seedLoc = 83,88] Seq=CACTCAGCAACTACTACTTCT \| ```          T               AC   C mRNA  3'  CTTCATCAT CAACG  TCA  miRNA 5'  GAGGTAGTA GTTGT  AGT           T         G     AT   T           3112111111611112331113 ``` \| SeedType = 3 NonSeedType = 1 \| \| --- \| --- \| --- \| --- \| --- \| --- \| | \| 225 \| WBGene00003185:mek-1(1956) \| ``` log2FC=0.104 padj=0.918 ``` \| [seedLoc = 64,69] Seq=AAACTATCAATCCCATTCACTATCTCT \| ```          T        CTTACC            A mRNA  3'  CTCTATCA      CTAAC TATCAA  miRNA 5'  GAGGTAGT      GGTTG ATAGTT           T        A          T                  3111211113555551211161111115 ``` \| SeedType = 2 NonSeedType = 1 \| \| --- \| --- \| --- \| --- \| --- \| --- \| | \| 226 \| WBGene00003196:mel-11(1958) \| ``` log2FC=-0.018 padj=0.992 ``` \| [seedLoc = 88,93] Seq=TATTGCACATTTTCTGTCTCGTTACCTCT \| ```          T       GCTCT    TTT   C    T mRNA  3'  CTCCATT     GTCT   ACA GTTA  miRNA 5'  GAGGTAG     TAGG   TGT TAGT           T                T     A    T           31111112455552112355111321213 ``` \| SeedType = 1 NonSeedType = -1 \| \| --- \| --- \| --- \| --- \| --- \| --- \| | \|  \|  \|  \| [seedLoc = 118,123] Seq=ACACACACACACTTCGCCTGCCTCT \| ```          T       CGC    C   CACACA mRNA  3'  CTCCGTC   TTCA ACA       miRNA 5'  GAGGTAG   AGGT TGT                T       T          ATAGTT           3111121134512115111333333 ``` \| SeedType = 2 NonSeedType = 2 \| \| --- \| --- \| --- \| --- \| --- \| --- \| | \| 227 \| WBGene00003209:mel-26(1959) \| ``` log2FC=-0.208 padj=0.696 ``` \| [seedLoc = 294,300] Seq=CACATGTAATTAACTACCATCC \| ```          C  A      A           C mRNA  3'  CT CCATCA TTAATGTA CA  miRNA 5'  GA GGTAGT GGTTGTAT GT           T         A        A  T           31151111113221122116113 ``` \| SeedType = 3 NonSeedType = 2 \| \| --- \| --- \| --- \| --- \| --- \| --- \| | \| 228 \| WBGene00003218:mep-1(1963) \| ``` log2FC=-0.069 padj=0.945 ``` \| [seedLoc = 233,238] Seq=ATTCTCGCATCTTCCCCTACCTTC \| ```          C       CCC   T   CTCTTA mRNA  3'  TTCCATC   TTC ACG       miRNA 5'  GAGGTAG   AGG TGT                T       T     T   ATAGTT           321111113551213112333333 ``` \| SeedType = 3 NonSeedType = -1 \| \| --- \| --- \| --- \| --- \| --- \| --- \| | \| 229 \| WBGene00003225:mev-1(1969) \| ``` log2FC=0.013 padj=0.993 ``` \| [seedLoc = 108,113] Seq=TCTCTCTTTTCCGTAGATTTACCTCT \| ```          T       TAGATG  TTTTCTCTCT mRNA  3'  CTCCATT      CC           miRNA 5'  GAGGTAG      GG                    T       TA      TTGTATAGTT           31111112334554113333333333 ``` \| SeedType = 1 NonSeedType = -1 \| \| --- \| --- \| --- \| --- \| --- \| --- \| | \| 230 \| WBGene00003228:mex-1(1970) \| ``` log2FC=0.219 padj=0.674 ``` \| [seedLoc = 23,29] Seq=CCTCCGTCCACCCCCTACCTAC \| ```          CA      CC   CCTGCCTCC mRNA  3'   TCCATC  CCA          miRNA 5'   AGGTAG  GGT                   TG      TA   TGTATAGTT           3311111133111333333333 ``` \| SeedType = 3 NonSeedType = 1 \| \| --- \| --- \| --- \| --- \| --- \| --- \| | \|  \|  \|  \| [seedLoc = 278,283] Seq=ATTCCCCATTTCTCTATCTCT \| ```          T           TTACCCCTTA mRNA  3'  CTCTATC TCT           miRNA 5'  GAGGTAG AGG                    T       T   TTGTATAGTT           3111211161123333333333 ``` \| SeedType = 2 NonSeedType = -1 \| \| --- \| --- \| --- \| --- \| --- \| --- \| | \|  \|  \|  \| [seedLoc = 295,300] Seq=TATCTCTCTCTCTAGTCTATCTCA \| ```                  TG    CTCTCTCTAT mRNA  3' ACTCTATC  ATCT           miRNA 5' TGAGGTAG  TAGG                                  TTGTATAGTT           111121115411123333333333 ``` \| SeedType = 2 NonSeedType = -1 \| \| --- \| --- \| --- \| --- \| --- \| --- \| | \| 231 \| WBGene00003229:mex-3(1973) \| ``` log2FC=-0.159 padj=0.842 ``` \| [seedLoc = 161,167] Seq=CCATTCCCATTTTCTCACCTCT \| ```          T     C      T  CCTTACC mRNA  3'  CTCCA TC TTT AC        miRNA 5'  GAGGT AG AGG TG                 T        T   T  TATAGTT           31111151161223113333333 ``` \| SeedType = 2 NonSeedType = -1 \| \| --- \| --- \| --- \| --- \| --- \| --- \| | \|  \|  \|  \| [seedLoc = 476,481] Seq=TCCTCTTTTCTCTCTATCTCT \| ```          T           CTTTTCTCCT mRNA  3'  CTCTATC TCT           miRNA 5'  GAGGTAG AGG                    T       T   TTGTATAGTT           3111211161123333333333 ``` \| SeedType = 2 NonSeedType = -1 \| \| --- \| --- \| --- \| --- \| --- \| --- \| | \| 232 \| WBGene00003230:mex-5(1974) \| ``` log2FC=0.215 padj=0.693 ``` \| [seedLoc = 492,496] Seq=TTCTTTCCCCTTCCTCAA \| ```          A     T      CTTTCTT    mRNA  3'  ACTCC TC  CC           miRNA 5'  TGAGG AG  GG                          T  TA  TTGTATAGTT           51111131166113333333666 ``` \| SeedType = 2 NonSeedType = -1 \| \| --- \| --- \| --- \| --- \| --- \| --- \| | \| 233 \| WBGene00003231:mex-6(1975) \| ``` log2FC=0.082 padj=0.959 ``` \| [seedLoc = 348,354] Seq=TTGACAGCAGTCCCTTTACCCTCA \| ```             C     TC  T    ACAGTT mRNA  3' ACT CCATT  CC GACG       miRNA 5' TGA GGTAG  GG TTGT                         TA       ATAGTT           111511112331152112333333 ``` \| SeedType = 3 NonSeedType = -1 \| \| --- \| --- \| --- \| --- \| --- \| --- \| | \| 234 \| WBGene00003239:mig-2(1981) \| ``` log2FC=0.157 padj=0.882 ``` \| [seedLoc = 175,180] Seq=CAGTCATTAGCATTTACCTCG \| ```          G       TA     TACTGAC mRNA  3'  CTCCATT   CGAT        miRNA 5'  GAGGTAG   GTTG                 T       TAG    TATAGTT           3111111233612123333333 ``` \| SeedType = 1 NonSeedType = -1 \| \| --- \| --- \| --- \| --- \| --- \| --- \| | \|  \|  \|  \| [seedLoc = 223,229] Seq=AATTTCTAAATTTACTACTCTCT \| ```          T   T          ATCTTTAA mRNA  3'  CTC CATCATTTAA         miRNA 5'  GAG GTAGTAGGTT                  T              GTATAGTT           31115111111221133333333 ``` \| SeedType = 2 NonSeedType = 2 \| \| --- \| --- \| --- \| --- \| --- \| --- \| | \| 235 \| WBGene00003253:mig-22(1992) \| ``` log2FC=0.387 padj=0.219 ``` \| [seedLoc = 282,287] Seq=CACTAAACAAAGTACATACTATCTCG \| ```          G         ACATGA    A    C mRNA  3'  CTCTATCAT      AACA ATCA  miRNA 5'  GAGGTAGTA      TTGT TAGT           T         GG        A    T           31112111113355451111311113 ``` \| SeedType = 2 NonSeedType = 1 \| \| --- \| --- \| --- \| --- \| --- \| --- \| | \| 236 \| WBGene00003368:mkk-4(1996) \| ``` log2FC=0.252 padj=0.874 ``` \| [seedLoc = 76,81] Seq=TCAAAATTTTCAAACTACTTCT \| ```          T        AACTTTTAAAACT mRNA  3'  CTTCATCA              miRNA 5'  GAGGTAGT                       T        AGGTTGTATAGTT           3112111113333333333333 ``` \| SeedType = 3 NonSeedType = -1 \| \| --- \| --- \| --- \| --- \| --- \| --- \| | \| 237 \| WBGene00003377:mls-2(2008) \| ``` log2FC=-0.488 padj=1.0 ``` \| [seedLoc = 79,84] Seq=CTCATCACTGACTCTATCTCT \| ```          T          AGT  CTACTC mRNA  3'  CTCTATC TC   CA       miRNA 5'  GAGGTAG AG   GT                T       T  GTT  ATAGTT           3111211161133311333333 ``` \| SeedType = 2 NonSeedType = -1 \| \| --- \| --- \| --- \| --- \| --- \| --- \| | \| 238 \| WBGene00003378:mml-1(2009) \| ``` log2FC=-0.084 padj=0.946 ``` \| [seedLoc = 253,258] Seq=AAATGTACATTGTCTCTACCTCT \| ```          T           GTT      AAA mRNA  3'  CTCCATC TCT   ACATGT    miRNA 5'  GAGGTAG AGG   TGTATA             T       T   T        GTT           311111116112355111121333 ``` \| SeedType = 1 NonSeedType = -1 \| \| --- \| --- \| --- \| --- \| --- \| --- \| | \| 239 \| WBGene00003386:mod-1(2014) \| ``` log2FC=-0.479 padj=0.912 ``` \| [seedLoc = 47,52] Seq=TAATTCACAGTTGCCGGTATCTACCTCC \| ```          C       TATGG   TTG   CTTAAT mRNA  3'  CTCCATC     CCG   ACA       miRNA 5'  GAGGTAG     GGT   TGT                T       TA            ATAGTT           3111111133544112554111333333 ``` \| SeedType = 1 NonSeedType = 2 \| \| --- \| --- \| --- \| --- \| --- \| --- \| | \| 240 \| WBGene00003404:mpz-1(2030) \| ``` log2FC=-0.192 padj=0.748 ``` \| [seedLoc = 110,116] Seq=CGATTTCCGGGTTTTACACTCT \| ```          T   A       G   CTTTAGC mRNA  3'  CTC CATT TT GGC        miRNA 5'  GAG GTAG AG TTG                 T        T  G   TATAGTT           31115111261232213333333 ``` \| SeedType = 2 NonSeedType = -1 \| \| --- \| --- \| --- \| --- \| --- \| --- \| | \|  \|  \|  \| [seedLoc = 176,181] Seq=TTCTAGCATTCCTACCTTC \| ```          C           T       TT mRNA  3'  TTCCATC  CT ACG ATC   miRNA 5'  GAGGTAG  GG TGT TAG            T       TA  T   A   TT           3211111166123112611133 ``` \| SeedType = 3 NonSeedType = -1 \| \| --- \| --- \| --- \| --- \| --- \| --- \| | \| 241 \| WBGene00003417:mrt-2(2041) \| ``` log2FC=0.048 padj=0.98 ``` \| [seedLoc = 60,66] Seq=AGTTTTGCAACAAGTGTACCCTCA \| ```             C    GTGAA      TTTGA mRNA  3' ACT CCAT     CAACGT      miRNA 5' TGA GGTA     GTTGTA                       GTAG       TAGTT           111511113333511112133333 ``` \| SeedType = 3 NonSeedType = 1 \| \| --- \| --- \| --- \| --- \| --- \| --- \| | \|  \|  \|  \| [seedLoc = 67,71] Seq=AGTTTTGCAACAAGTGTACCCTCAT \| ```          T   C    GTGAA      TTTGA mRNA  3'  ACT CCAT     CAACGT      miRNA 5'  TGA GGTA     GTTGTA                        GTAG       TAGTT           5111511113333511112133333 ``` \| SeedType = 3 NonSeedType = 1 \| \| --- \| --- \| --- \| --- \| --- \| --- \| | \| 242 \| WBGene00003472:mtk-1(2073) \| ``` log2FC=-0.082 padj=0.951 ``` \| [seedLoc = 74,79] Seq=TTCGTGATAAAAATGTACCTCT \| ```          T      GTAAA    GTGCTT mRNA  3'  CTCCAT     AATA       miRNA 5'  GAGGTA     TTGT                T      GTAGG    ATAGTT           3111111333331121333333 ``` \| SeedType = 1 NonSeedType = -1 \| \| --- \| --- \| --- \| --- \| --- \| --- \| | \| 243 \| WBGene00003495:mup-2(2084) \| ``` log2FC=-0.003 padj=0.998 ``` \| [seedLoc = 174,180] Seq=TCAAGTACTCTGTTTCTGCATACTCTCC \| ```          C   T   A     TTGTCT    AACT mRNA  3'  CTC CAT CGTCT      CATG     miRNA 5'  GAG GTA GTAGG      GTAT              T             TT        AGTT           3111511151211233455511123333 ``` \| SeedType = 2 NonSeedType = -1 \| \| --- \| --- \| --- \| --- \| --- \| --- \| | \| 244 \| WBGene00003548:nas-30(2119) \| ``` log2FC=-0.512 padj=0.098 ``` \| [seedLoc = 60,66] Seq=ATTTGTAAATTTTGTACCATCC \| ```          C  A    GT           TA mRNA  3'  CT CCAT  TTTAA ATGTT   miRNA 5'  GA GGTA  AGGTT TATAG            T       GT     G     TT           31151111331221161121233 ``` \| SeedType = 3 NonSeedType = 2 \| \| --- \| --- \| --- \| --- \| --- \| --- \| | \| 245 \| WBGene00003549:nas-31(2120) \| ``` log2FC=0.24 padj=0.923 ``` \| [seedLoc = 195,200] Seq=CGGGTATGCACTACCTTT \| ```          T                  G  C mRNA  3'  TTCCATCA     CGTAT GG  miRNA 5'  GAGGTAGT     GTATA TT           T        AGGTT     G              32111111166666121113225 ``` \| SeedType = 3 NonSeedType = -1 \| \| --- \| --- \| --- \| --- \| --- \| --- \| | \| 246 \| WBGene00003559:ncl-1(2126) \| ``` log2FC=0.273 padj=0.34 ``` \| [seedLoc = 111,116] Seq=GTGAATAGAATTTTCTACTTCT \| ```          T       T     G   AGTG mRNA  3'  CTTCATC TTTAA ATA     miRNA 5'  GAGGTAG AGGTT TAT              T       T     G   AGTT           3112111131221131113333 ``` \| SeedType = 3 NonSeedType = 2 \| \| --- \| --- \| --- \| --- \| --- \| --- \| | \|  \|  \|  \| [seedLoc = 1070,1076] Seq=AAGCTTTTTAACCGGTTTTGTACCCTCT \| ```          T  C    GTTTTGG     TTT    A mRNA  3'  CT CCAT       CCAAT   TCGA  miRNA 5'  GA GGTA       GGTTG   AGTT           T       GTA         TAT                3115111133355441111233311215 ``` \| SeedType = 3 NonSeedType = 1 \| \| --- \| --- \| --- \| --- \| --- \| --- \| | \| 247 \| WBGene00003561:ncr-1(2127) \| ``` log2FC=-0.238 padj=0.553 ``` \| [seedLoc = 173,179] Seq=AAACTTTATCATATTTTCATTACTCTCA \| ```              T     CT    TAC   T    A mRNA  3' ACTC CATTA  TTTA   TAT TCAA  miRNA 5' TGAG GTAGT  AGGT   GTA AGTT                           T     T                1111511121551221355211311115 ``` \| SeedType = 2 NonSeedType = 2 \| \| --- \| --- \| --- \| --- \| --- \| --- \| | \|  \|  \|  \| [seedLoc = 181,185] Seq=TATCATATTTTCATTACTCTCAA \| ```          A    T        TT    CTAT mRNA  3'  ACTC CATTA CT  TATA     miRNA 5'  TGAG GTAGT GG  GTAT                         A  TT    AGTT           511115111216123321113333 ``` \| SeedType = 2 NonSeedType = -1 \| \| --- \| --- \| --- \| --- \| --- \| --- \| | \| 248 \| WBGene00003567:ncx-2(2135) \| ``` log2FC=0.157 padj=0.866 ``` \| [seedLoc = 253,258] Seq=CGCTTATGTCTTCTTCCTCT \| ```          T    T      TG       C mRNA  3'  CTCC TC TTC  TAT TCG  miRNA 5'  GAGG AG AGG  GTA AGT           T    T  T   TT   T   T           3111131161213321161123 ``` \| SeedType = 2 NonSeedType = -1 \| \| --- \| --- \| --- \| --- \| --- \| --- \| | \| 249 \| WBGene00003599:nhl-3(2165) \| ``` log2FC=-0.274 padj=0.742 ``` \| [seedLoc = 50,55] Seq=GAAAATATAATTTTACCTCT \| ```          T                 AAAG mRNA  3'  CTCCAT  TTTAATATA     miRNA 5'  GAGGTA  AGGTTGTAT              T      GT         AGTT           3111111661221121113333 ``` \| SeedType = 1 NonSeedType = 2 \| \| --- \| --- \| --- \| --- \| --- \| --- \| | \| 250 \| WBGene00003601:nhr-2(2168) \| ``` log2FC=0.127 padj=1.0 ``` \| [seedLoc = 222,227] Seq=GTGAATTTAATTCTCTATCTCA \| ```                  TC     TTAAGTG mRNA  3' ACTCTATC  TTAAT        miRNA 5' TGAGGTAG  GGTTG                         TA     TATAGTT           1111211133221123333333 ``` \| SeedType = 2 NonSeedType = 2 \| \| --- \| --- \| --- \| --- \| --- \| --- \| | \| 251 \| WBGene00003602:nhr-3(2169) \| ``` log2FC=-0.116 padj=0.906 ``` \| [seedLoc = 48,54] Seq=TTGTATACACCAGCCTACCTAC \| ```          CA      CGA         GTT mRNA  3'   TCCATC   CC ACATAT    miRNA 5'   AGGTAG   GG TGTATA             TG      TA   T      GTT           33111111335116111111333 ``` \| SeedType = 3 NonSeedType = -1 \| \| --- \| --- \| --- \| --- \| --- \| --- \| | \|  \|  \|  \| [seedLoc = 277,282] Seq=GAGTTGCCAAACTATTGTTACCTCA \| ```                 TGT    AA  CGTTGAG mRNA  3' ACTCCAT   TATC  AC        miRNA 5' TGAGGTA   GTAG  TG                               GT  TATAGTT           1111111545211133113333333 ``` \| SeedType = 1 NonSeedType = -1 \| \| --- \| --- \| --- \| --- \| --- \| --- \| | \| 252 \| WBGene00003603:nhr-4(2170) \| ``` log2FC=-0.348 padj=0.473 ``` \| [seedLoc = 25,30] Seq=TCGGTTTTTGGATTCTACCTCT \| ```          T       TTAGGTTTTTGGCT mRNA  3'  CTCCATC               miRNA 5'  GAGGTAG                        T       TAGGTTGTATAGTT           3111111133333333333333 ``` \| SeedType = 1 NonSeedType = -1 \| \| --- \| --- \| --- \| --- \| --- \| --- \| | \| 253 \| WBGene00003616:nhr-17(2179) \| ``` log2FC=0.043 padj=0.98 ``` \| [seedLoc = 121,126] Seq=TTTCCCCTTTCCACTACCTCA \| ```                      TTTCCCCTTT mRNA  3' ACTCCATCA CC           miRNA 5' TGAGGTAGT GG                             A  TTGTATAGTT           1111111116113333333333 ``` \| SeedType = 1 NonSeedType = -1 \| \| --- \| --- \| --- \| --- \| --- \| --- \| | \| 254 \| WBGene00003618:nhr-19(2180) \| ``` log2FC=-0.215 padj=0.781 ``` \| [seedLoc = 56,61] Seq=AAATTTTCCTCTTTCTATCTCC \| ```          C       T   TCCTTTTAAA mRNA  3'  CTCTATC TTC           miRNA 5'  GAGGTAG AGG                    T       T   TTGTATAGTT           3111211131213333333333 ``` \| SeedType = 2 NonSeedType = -1 \| \| --- \| --- \| --- \| --- \| --- \| --- \| | \|  \|  \|  \| [seedLoc = 131,136] Seq=GATCACACAACTCAAATACCTCG \| ```          G      AAAC      CACTAG mRNA  3'  CTCCAT    TCAACA       miRNA 5'  GAGGTA    GGTTGT                T      GTA       ATAGTT           31111113335211111333333 ``` \| SeedType = 1 NonSeedType = 1 \| \| --- \| --- \| --- \| --- \| --- \| --- \| | \| 255 \| WBGene00003623:nhr-25(2185) \| ``` log2FC=-0.035 padj=0.993 ``` \| [seedLoc = 187,192] Seq=ATCAATATCATTTTATACCTCA \| ```                 AT    C    ACTA mRNA  3' ACTCCAT  TTTA TATA     miRNA 5' TGAGGTA  AGGT GTAT                     GT    T    AGTT           1111111331221321113333 ``` \| SeedType = 1 NonSeedType = 2 \| \| --- \| --- \| --- \| --- \| --- \| --- \| | \| 256 \| WBGene00003633:nhr-43(2194) \| ``` log2FC=-0.281 padj=0.678 ``` \| [seedLoc = 142,147] Seq=ATCCTTCCAAATGACTATCTCT \| ```          T        GTA   CTTCCTA mRNA  3'  CTCTATCA   AAC        miRNA 5'  GAGGTAGT   TTG                 T        AGG   TATAGTT           3111211113331113333333 ``` \| SeedType = 2 NonSeedType = 1 \| \| --- \| --- \| --- \| --- \| --- \| --- \| | \|  \|  \|  \| [seedLoc = 225,230] Seq=CACTATATTTCCTTACCTCA \| ```                      TT       C mRNA  3' ACTCCAT  TCC  TATATCA  miRNA 5' TGAGGTA  AGG  GTATAGT                  GT   TT       T           1111111661113321111113 ``` \| SeedType = 1 NonSeedType = -1 \| \| --- \| --- \| --- \| --- \| --- \| --- \| | \| 257 \| WBGene00003635:nhr-45(2196) \| ``` log2FC=-0.211 padj=0.681 ``` \| [seedLoc = 191,197] Seq=TTTTCGGCACGTCGTCACCTCT \| ```          T     C GC G     GCTTTT mRNA  3'  CTCCA T  T C ACG       miRNA 5'  GAGGT A  A G TGT                T       GT G T   ATAGTT           31111151331316112333333 ``` \| SeedType = 2 NonSeedType = -1 \| \| --- \| --- \| --- \| --- \| --- \| --- \| | \| 258 \| WBGene00003636:nhr-46(2197) \| ``` log2FC=-0.176 padj=0.854 ``` \| [seedLoc = 205,211] Seq=CACTCACTCACTCTACACTCA \| ```              A       ACT  C    C mRNA  3' ACTC CATC TC   CA  TCA  miRNA 5' TGAG GTAG AG   GT  AGT                    T  GTT  AT   T           11115111161133311361113 ``` \| SeedType = 2 NonSeedType = -1 \| \| --- \| --- \| --- \| --- \| --- \| --- \| | \|  \|  \|  \| [seedLoc = 320,326] Seq=GGCTTTATTGGTTTTTTACTCTCA \| ```              T    TT  G T   T   G mRNA  3' ACTC CATT  TT G TAT TCG  miRNA 5' TGAG GTAG  AG T GTA AGT                    T   G T   T   T           111151112351232321131123 ``` \| SeedType = 2 NonSeedType = -1 \| \| --- \| --- \| --- \| --- \| --- \| --- \| | \| 259 \| WBGene00003656:nhr-66(2216) \| ``` log2FC=-0.09 padj=0.945 ``` \| [seedLoc = 79,84] Seq=CGATTGTACTACAATGATTACCTCT \| ```          T        GTAA  T        C mRNA  3'  CTCCATTA    CA CATGTTAG  miRNA 5'  GAGGTAGT    GT GTATAGTT           T        AG    T                    3111111213355113111212125 ``` \| SeedType = 1 NonSeedType = -1 \| \| --- \| --- \| --- \| --- \| --- \| --- \| | \|  \|  \|  \| [seedLoc = 196,201] Seq=TGTTTTTGGTTTCATTACCTCT \| ```          T        CTTTGGTTTTTGT mRNA  3'  CTCCATTA              miRNA 5'  GAGGTAGT                       T        AGGTTGTATAGTT           3111111213333333333333 ``` \| SeedType = 1 NonSeedType = -1 \| \| --- \| --- \| --- \| --- \| --- \| --- \| | \| 260 \| WBGene00003661:nhr-71(2220) \| ``` log2FC=-0.055 padj=0.982 ``` \| [seedLoc = 155,160] Seq=ACAATTGCTCCCCCTCACTACCTTC \| ```          C        C   CCCT   TAACA mRNA  3'  TTCCATCA TCC    CGT      miRNA 5'  GAGGTAGT AGG    GTA               T            TT     TAGTT           3211111115111335512133333 ``` \| SeedType = 3 NonSeedType = -1 \| \| --- \| --- \| --- \| --- \| --- \| --- \| | \| 261 \| WBGene00003674:nhr-84(2227) \| ``` log2FC=-0.175 padj=0.912 ``` \| [seedLoc = 70,75] Seq=TTTTAGAAGACTTTTTTACCTCA \| ```                  TT     AAG   TT mRNA  3' ACTCCATT  TTCAG   ATT   miRNA 5' TGAGGTAG  AGGTT   TAG                    T      GTA   TT           11111112351211233311233 ``` \| SeedType = 1 NonSeedType = 2 \| \| --- \| --- \| --- \| --- \| --- \| --- \| | \| 262 \| WBGene00003678:nhr-88(2230) \| ``` log2FC=-0.129 padj=0.885 ``` \| [seedLoc = 372,377] Seq=AAATGTGATTCGTGCTGCCTCC \| ```          C         G TTA    AAA mRNA  3'  CTCCGTCGT C   GTGT    miRNA 5'  GAGGTAGTA G   TATA             T         G TTG    GTT           3111121121313332121333 ``` \| SeedType = 2 NonSeedType = -1 \| \| --- \| --- \| --- \| --- \| --- \| --- \| | \| 263 \| WBGene00003681:nhr-91(2234) \| ``` log2FC=-0.193 padj=0.923 ``` \| [seedLoc = 103,108] Seq=TTTGAAATTCCCCTGCCTCT \| ```          T           CTTAAAGTTT mRNA  3'  CTCCGTC  CC           miRNA 5'  GAGGTAG  GG                    T       TA  TTGTATAGTT           3111121166113333333333 ``` \| SeedType = 2 NonSeedType = -1 \| \| --- \| --- \| --- \| --- \| --- \| --- \| | \|  \|  \|  \| [seedLoc = 353,358] Seq=CTCCCCATTCTCTGTTCCTATCTCG \| ```          G       CTT    CTTACCCCTC mRNA  3'  CTCTATC   GTCT           miRNA 5'  GAGGTAG   TAGG                    T              TTGTATAGTT           3111211155521123333333333 ``` \| SeedType = 2 NonSeedType = -1 \| \| --- \| --- \| --- \| --- \| --- \| --- \| | \| 264 \| WBGene00003687:nhr-97(2237) \| ``` log2FC=0.111 padj=0.911 ``` \| [seedLoc = 556,561] Seq=TTTTATTCTTCTTTCTACTTCC \| ```          C       T   TT T    TT mRNA  3'  CTTCATC TTC  C TATT   miRNA 5'  GAGGTAG AGG  G ATAG            T       T   TT T    TT           3112111131213313111233 ``` \| SeedType = 3 NonSeedType = -1 \| \| --- \| --- \| --- \| --- \| --- \| --- \| | \| 265 \| WBGene00003690:nhr-100(2240) \| ``` log2FC=-0.061 padj=0.973 ``` \| [seedLoc = 229,235] Seq=CGTCTTTTTTTGTTATACTCTCC \| ```          C   T   ATTGTTTTTTTCTGC mRNA  3'  CTC CAT                miRNA 5'  GAG GTA                         T       GTAGGTTGTATAGTT           31115111333333333333333 ``` \| SeedType = 2 NonSeedType = -1 \| \| --- \| --- \| --- \| --- \| --- \| --- \| | \| 266 \| WBGene00003728:nhr-138(2276) \| ``` log2FC=0.031 padj=0.993 ``` \| [seedLoc = 81,86] Seq=TGTGATATATTCCCTACCTCT \| ```          T           TT     GTGT mRNA  3'  CTCCATC  CC  ATATA     miRNA 5'  GAGGTAG  GG  TGTAT              T       TA  T      AGTT           31111111661135121113333 ``` \| SeedType = 1 NonSeedType = -1 \| \| --- \| --- \| --- \| --- \| --- \| --- \| | \| 267 \| WBGene00003732:nhx-4(2280) \| ``` log2FC=-0.107 padj=0.946 ``` \| [seedLoc = 458,464] Seq=ACGCCAACAATCGAGCATAATATCCTCT \| ```          T    T  AATA  AG      ACCGCA mRNA  3'  CTCC AT    CG  CTAACA       miRNA 5'  GAGG TA    GT  GGTTGT                T             A       ATAGTT           3111151155551234121111333333 ``` \| SeedType = 2 NonSeedType = 1 \| \| --- \| --- \| --- \| --- \| --- \| --- \| | \| 268 \| WBGene00003754:nlp-16(2303) \| ``` log2FC=0.1 padj=0.961 ``` \| [seedLoc = 119,124] Seq=CGTTTTGCGTATTTCTACCTCT \| ```          T            T    TTTGC mRNA  3'  CTCCATC TTTA GCGT      miRNA 5'  GAGGTAG AGGT TGTA               T       T         TAGTT           31111111612215212133333 ``` \| SeedType = 1 NonSeedType = 2 \| \| --- \| --- \| --- \| --- \| --- \| --- \| | \| 269 \| WBGene00003759:nlp-21(2309) \| ``` log2FC=-0.042 padj=0.99 ``` \| [seedLoc = 38,43] Seq=TACTATCACTTCCACTTCTACTACTTCA \| ```                    CTTCA  TT  C     T mRNA  3' ACTTCATCAT     CC  CA TATCA  miRNA 5' TGAGGTAGTA     GG  GT ATAGT                            TT        T           1112111111555551133115111113 ``` \| SeedType = 3 NonSeedType = -1 \| \| --- \| --- \| --- \| --- \| --- \| --- \| | \|  \|  \|  \| [seedLoc = 44,48] Seq=CACTTCCACTTCTACTACTTCAC \| ```          C            TCACCT   C mRNA  3'  ACTTCATCATCT      TCA  miRNA 5'  TGAGGTAGTAGG      AGT                        TTGTAT   T           51112111111123333331113 ``` \| SeedType = 3 NonSeedType = -1 \| \| --- \| --- \| --- \| --- \| --- \| --- \| | \| 270 \| WBGene00003776:nmy-1(2322) \| ``` log2FC=-0.14 padj=0.84 ``` \| [seedLoc = 124,129] Seq=ATATTTACATACTGTTCTACTTCT \| ```          T       TT   AT    TTATA mRNA  3'  CTTCATC  GTC  ACAT      miRNA 5'  GAGGTAG  TAG  TGTA               T            GT    TAGTT           311211115521133111133333 ``` \| SeedType = 3 NonSeedType = -1 \| \| --- \| --- \| --- \| --- \| --- \| --- \| | \| 271 \| WBGene00003785:nos-3(2328) \| ``` log2FC=0.052 padj=0.961 ``` \| [seedLoc = 430,435] Seq=CTTTCTTTCTTATACTATCTCC \| ```          C         ATTCTTTCTTTC mRNA  3'  CTCTATCAT             miRNA 5'  GAGGTAGTA                      T         GGTTGTATAGTT           3111211111333333333333 ``` \| SeedType = 2 NonSeedType = -1 \| \| --- \| --- \| --- \| --- \| --- \| --- \| | \| 272 \| WBGene00003793:npp-7(2338) \| ``` log2FC=-0.014 padj=0.993 ``` \| [seedLoc = 281,287] Seq=CCCTAGTTATCTTTCTATCCTCA \| ```               T   TT    TTG    CC mRNA  3' ACTCC ATC  TCTA    ATC   miRNA 5' TGAGG TAG  AGGT    TAG                     T     TGTA   TT           111115111351121333611133 ``` \| SeedType = 2 NonSeedType = 2 \| \| --- \| --- \| --- \| --- \| --- \| --- \| | \|  \|  \|  \| [seedLoc = 458,464] Seq=AAATTATTCAAAATTACCATCG \| ```          G  A      A     T      A mRNA  3'  CT CCATTA   AAC TATTAA  miRNA 5'  GA GGTAGT   TTG ATAGTT           T         AGG   T                  311511112136611131112115 ``` \| SeedType = 3 NonSeedType = 1 \| \| --- \| --- \| --- \| --- \| --- \| --- \| | \|  \|  \|  \| [seedLoc = 633,638] Seq=TGAGTTCTAGCTAGAGCTACTTCC \| ```          C        AGA     CTTGAGT mRNA  3'  CTTCATCG   TCGAT        miRNA 5'  GAGGTAGT   GGTTG                 T        A       TATAGTT           311211112345212123333333 ``` \| SeedType = 3 NonSeedType = 2 \| \| --- \| --- \| --- \| --- \| --- \| --- \| | \| 273 \| WBGene00003796:npp-10(2342) \| ``` log2FC=0.109 padj=0.888 ``` \| [seedLoc = 117,123] Seq=CACATGACTGCTTGCCAATACCATCT \| ```          T  A    AAC      T  GTACAC mRNA  3'  CT CCAT   CGTTCG CA       miRNA 5'  GA GGTA   GTAGGT GT                T                T  ATAGTT           31151111555121212311333333 ``` \| SeedType = 3 NonSeedType = 2 \| \| --- \| --- \| --- \| --- \| --- \| --- \| | \| 274 \| WBGene00003826:ntl-3(2365) \| ``` log2FC=-0.038 padj=0.975 ``` \| [seedLoc = 39,44] Seq=TTTTTCTCTGCGTCTATCTCT \| ```          T         G  TCTCTTTTT mRNA  3'  CTCTATC T CG          miRNA 5'  GAGGTAG A GT                   T       T G  TGTATAGTT           3111211161312333333333 ``` \| SeedType = 2 NonSeedType = -1 \| \| --- \| --- \| --- \| --- \| --- \| --- \| | \|  \|  \|  \| [seedLoc = 47,52] Seq=TTCTCTGCGTCTATCTCTCTATCTCT \| ```          T       TC    TCT    C  TT mRNA  3'  CTCTATC  TCTA   GCGT TC   miRNA 5'  GAGGTAG  AGGT   TGTA AG            T       T            T  TT           31112111351121555212131133 ``` \| SeedType = 2 NonSeedType = 2 \| \| --- \| --- \| --- \| --- \| --- \| --- \| | \|  \|  \|  \| [seedLoc = 448,453] Seq=CTTATTATTCCCTTAAAAATACCTCT \| ```          T      AAAAAT   CT   TATTC mRNA  3'  CTCCAT      TCC  TAT      miRNA 5'  GAGGTA      AGG  GTA               T      GT       TT   TAGTT           31111113355551113321133333 ``` \| SeedType = 1 NonSeedType = -1 \| \| --- \| --- \| --- \| --- \| --- \| --- \| | \| 275 \| WBGene00003827:ntl-4(2366) \| ``` log2FC=0.095 padj=0.903 ``` \| [seedLoc = 665,671] Seq=CTTTATTTAAGTTGTACCCTCA \| ```             C        G   T    TC mRNA  3' ACT CCAT GTT AAT TATT   miRNA 5' TGA GGTA TAG TTG ATAG                    G   G   T    TT           11151111621231123111233 ``` \| SeedType = 3 NonSeedType = -1 \| \| --- \| --- \| --- \| --- \| --- \| --- \| | \| 276 \| WBGene00003868:ooc-3(2396) \| ``` log2FC=0.023 padj=0.993 ``` \| [seedLoc = 150,155] Seq=TCCTGTAGATCTTGCCTATTACCTCT \| ```          T            TTCT G     CT mRNA  3'  CTCCATTATCCG    A ATGTC   miRNA 5'  GAGGTAGTAGGT    T TATAG            T                 G     TT           31111112111125555131121133 ``` \| SeedType = 1 NonSeedType = 2 \| \| --- \| --- \| --- \| --- \| --- \| --- \| | \| 277 \| WBGene00003870:ooc-5(2397) \| ``` log2FC=0.365 padj=0.41 ``` \| [seedLoc = 103,108] Seq=AAGCTATTGCATTCCTCGATGATTACCTCA \| ```                   GTAGC   TT   T      A mRNA  3' ACTCCATTA     TCC  ACG TATCGA  miRNA 5' TGAGGTAGT     AGG  TGT ATAGTT                            T                       111111121455451113511251111215 ``` \| SeedType = 1 NonSeedType = -1 \| \| --- \| --- \| --- \| --- \| --- \| --- \| | \| 278 \| WBGene00003876:pept-2(2398) \| ``` log2FC=0.063 padj=0.98 ``` \| [seedLoc = 78,81] Seq=CACTTCAACCCCGTCCACCTCT \| ```          T     C TGCC     T     C mRNA  3'  CTCCA C    CCAAC   TCA  miRNA 5'  GAGGT G    GGTTG   AGT           T     A TA       TAT   T           311111313355111113661113 ``` \| SeedType = 2 NonSeedType = 1 \| \| --- \| --- \| --- \| --- \| --- \| --- \| | \| 279 \| WBGene00003882:orc-2(2402) \| ``` log2FC=0.21 padj=0.71 ``` \| [seedLoc = 40,45] Seq=AAATGAACACACTCTGCCTCC \| ```          C          AC   AGTAAA mRNA  3'  CTCCGTC TC  ACA       miRNA 5'  GAGGTAG AG  TGT                T       T  GT   ATAGTT           3111121161133111333333 ``` \| SeedType = 2 NonSeedType = -1 \| \| --- \| --- \| --- \| --- \| --- \| --- \| | \| 280 \| WBGene00003901:paa-1(2415) \| ``` log2FC=-0.022 padj=0.988 ``` \| [seedLoc = 41,45] Seq=CAAATTGGAATCTCTACATCT \| ```          T  A          G  TAAAC mRNA  3'  CT CATC TCTAA GT      miRNA 5'  GA GTAG AGGTT TA               T  G    T     G  TAGTT           3113111161121132133333 ``` \| SeedType = 3 NonSeedType = 2 \| \| --- \| --- \| --- \| --- \| --- \| --- \| | \|  \|  \|  \| [seedLoc = 95,100] Seq=ACGCGAATAATCTCTATCTCT \| ```          T               AGCGCA mRNA  3'  CTCTATC TCTAATA       miRNA 5'  GAGGTAG AGGTTGT                T       T       ATAGTT           3111211161121121333333 ``` \| SeedType = 2 NonSeedType = 2 \| \| --- \| --- \| --- \| --- \| --- \| --- \| | \|  \|  \|  \| [seedLoc = 275,280] Seq=CCGTATATTTCACCATCCTACTTCT \| ```          T       CTA   CTT     GCC mRNA  3'  CTTCATC   CCA   TATAT    miRNA 5'  GAGGTAG   GGT   GTATA             T       TA    T       GTT           3112111133511135521111333 ``` \| SeedType = 3 NonSeedType = 1 \| \| --- \| --- \| --- \| --- \| --- \| --- \| | \| 281 \| WBGene00003911:pak-1(2423) \| ``` log2FC=0.344 padj=0.286 ``` \| [seedLoc = 272,277] Seq=CAATGCACAATTTGGACTATCTCT \| ```          T        GG       CGTAAC mRNA  3'  CTCTATCA  TTTAACA       miRNA 5'  GAGGTAGT  AGGTTGT                T                 ATAGTT           311121111441221111333333 ``` \| SeedType = 2 NonSeedType = 1 \| \| --- \| --- \| --- \| --- \| --- \| --- \| | \| 282 \| WBGene00003915:pan-1(2426) \| ``` log2FC=0.511 padj=0.061 ``` \| [seedLoc = 78,83] Seq=CATATTCTCCCCGGTTGAATACCTCA \| ```                 AAGTTGG  CCTCTTATAC mRNA  3' ACTCCAT       CC           miRNA 5' TGAGGTA       GG                           GTA      TTGTATAGTT           11111113335544113333333333 ``` \| SeedType = 1 NonSeedType = -1 \| \| --- \| --- \| --- \| --- \| --- \| --- \| | \| 283 \| WBGene00003916:par-1(2427) \| ``` log2FC=0.024 padj=0.987 ``` \| [seedLoc = 234,239] Seq=CAGCTGTGAACGTTACCTCC \| ```          C         G           C mRNA  3'  CTCCAT  T CAA GTGTCGA  miRNA 5'  GAGGTA  A GTT TATAGTT           T      GT G   G                   31111116613111621211215 ``` \| SeedType = 1 NonSeedType = 1 \| \| --- \| --- \| --- \| --- \| --- \| --- \| | \| 284 \| WBGene00003919:par-4(2437) \| ``` log2FC=0.225 padj=0.762 ``` \| [seedLoc = 366,372] Seq=AAAAGTTCGAGTTTTTACACTCA \| ```              A    T  G   TTGAAAA mRNA  3' ACTC CATT TT AGC        miRNA 5' TGAG GTAG AG TTG                          T  G   TATAGTT           11115111231231213333333 ``` \| SeedType = 2 NonSeedType = -1 \| \| --- \| --- \| --- \| --- \| --- \| --- \| | \| 285 \| WBGene00003924:pas-3(2443) \| ``` log2FC=0.054 padj=0.961 ``` \| [seedLoc = 98,103] Seq=TCTTATCACCCATATTCCTATCTCA \| ```                  CTT  A  C       CT mRNA  3' ACTCTATC   AT CC AC TATT   miRNA 5' TGAGGTAG   TA GG TG ATAG                            T  T    TT           11112111555115113116111233 ``` \| SeedType = 2 NonSeedType = -1 \| \| --- \| --- \| --- \| --- \| --- \| --- \| | \| 286 \| WBGene00003933:pat-9(2452) \| ``` log2FC=-0.39 padj=0.382 ``` \| [seedLoc = 280,285] Seq=TCTTAAACAAAAAAGTCTTTACCTCT \| ```          T           GAAAA    AATTCT mRNA  3'  CTCCATT TCT     AACA       miRNA 5'  GAGGTAG AGG     TTGT                T       T            ATAGTT           311111126112455551111333333 ``` \| SeedType = 1 NonSeedType = 1 \| \| --- \| --- \| --- \| --- \| --- \| --- \| | \| 287 \| WBGene00003936:pat-12(2465) \| ``` log2FC=0.085 padj=0.956 ``` \| [seedLoc = 431,436] Seq=CCAAGTGCCTCCGCCTAAGATTTTACCTCA \| ```                  TTAGA     CCTC    AACC mRNA  3' ACTCCATT     ATCCG    CGTG     miRNA 5' TGAGGTAG     TAGGT    GTAT                                T       AGTT           111111125554511112355512123333 ``` \| SeedType = 1 NonSeedType = 2 \| \| --- \| --- \| --- \| --- \| --- \| --- \| | \|  \|  \|  \| [seedLoc = 948,953] Seq=TTCAGTTCAAGTTATCTACCTTT \| ```          T       T   G   TTGACTT mRNA  3'  TTCCATC ATT AAC        miRNA 5'  GAGGTAG TAG TTG                 T           G   TATAGTT           32111111511231113333333 ``` \| SeedType = 3 NonSeedType = 1 \| \| --- \| --- \| --- \| --- \| --- \| --- \| | \|  \|  \|  \| [seedLoc = 1859,1865] Seq=GAGGTGGCGGTGATTACCGTCG \| ```          G  G      GT     GTGGAG mRNA  3'  CT CCATTA   GGCG       miRNA 5'  GA GGTAGT   TTGT                T         AGG    ATAGTT           31141111213362212333333 ``` \| SeedType = 3 NonSeedType = -1 \| \| --- \| --- \| --- \| --- \| --- \| --- \| | \| 288 \| WBGene00003954:pcm-1(2480) \| ``` log2FC=-0.255 padj=0.836 ``` \| [seedLoc = 50,56] Seq=CAACTTGTGCATATTGTATACCATCT \| ```          T  A    AT   AT     T    C mRNA  3'  CT CCAT  GTT  ACGTG TCAA  miRNA 5'  GA GGTA  TAG  TGTAT AGTT           T       G    GT                      31151111352123311212511115 ``` \| SeedType = 3 NonSeedType = -1 \| \| --- \| --- \| --- \| --- \| --- \| --- \| | \| 289 \| WBGene00003963:pdi-2(2489) \| ``` log2FC=0.343 padj=0.42 ``` \| [seedLoc = 115,120] Seq=CCCTGTGCTACTTATTTTGCTATCTCT \| ```          T         TTTAT   T      CC mRNA  3'  CTCTATCGT     TCA CGTGTC   miRNA 5'  GAGGTAGTA     GGT GTATAG            T                 T      TT           311121112155555211312121133 ``` \| SeedType = 2 NonSeedType = 2 \| \| --- \| --- \| --- \| --- \| --- \| --- \| | \| 290 \| WBGene00003968:peb-1(2495) \| ``` log2FC=-0.259 padj=0.925 ``` \| [seedLoc = 605,611] Seq=TTGTTTGGGATTCCAATACTCTCA \| ```              T   AACC    G  TTGTT mRNA  3' ACTC CAT    TTAG GT      miRNA 5' TGAG GTA    GGTT TA                       GTA     G  TAGTT           111151113335221232133333 ``` \| SeedType = 2 NonSeedType = 2 \| \| --- \| --- \| --- \| --- \| --- \| --- \| | \| 291 \| WBGene00003978:pes-4(2500) \| ``` log2FC=-0.25 padj=0.615 ``` \| [seedLoc = 530,536] Seq=GAGCTTTATTGATTTTTCTCATCTACACTCG \| ```          G   A    TAC   TTTTAGT   T    G mRNA  3'  CTC CATC   TCT       TAT TCGA  miRNA 5'  GAG GTAG   AGG       GTA AGTT           T        T     TT        T                3111511113551123355545211311214 ``` \| SeedType = 2 NonSeedType = -1 \| \| --- \| --- \| --- \| --- \| --- \| --- \| | \| 292 \| WBGene00004004:pgp-10(2522) \| ``` log2FC=0.393 padj=0.494 ``` \| [seedLoc = 26,32] Seq=GGCATGCAAAAACTACACTCT \| ```          T   A     AA          G mRNA  3'  CTC CATCA   AACGTA CG  miRNA 5'  GAG GTAGT   TTGTAT GT           T         AGG      A  T           31115111113361112116123 ``` \| SeedType = 2 NonSeedType = 1 \| \| --- \| --- \| --- \| --- \| --- \| --- \| | \| 293 \| WBGene00004017:phg-1(2532) \| ``` log2FC=-0.651 padj=0.142 ``` \| [seedLoc = 81,86] Seq=TTTATTTCTCTCTTCTACTTCT \| ```          T       T   CTCTTTATTT mRNA  3'  CTTCATC TCT           miRNA 5'  GAGGTAG AGG                    T       T   TTGTATAGTT           3112111131123333333333 ``` \| SeedType = 3 NonSeedType = -1 \| \| --- \| --- \| --- \| --- \| --- \| --- \| | \| 294 \| WBGene00004049:parp-1(2563) \|  \| [seedLoc = 39,44] Seq=CGCCCCCCACCCCCAATACCTCG \| ```          G      AACC  C  CCCCCGC mRNA  3'  CTCCAT    CC AC        miRNA 5'  GAGGTA    GG TG                 T      GTA   T  TATAGTT           31111113335113113333333 ``` \| SeedType = 1 NonSeedType = -1 \| \| --- \| --- \| --- \| --- \| --- \| --- \| | \| 295 \| WBGene00004050:parp-2(2564) \|  \| [seedLoc = 41,46] Seq=TCACATTTTATCTGTTTCTACCTCT \| ```          T       TTT     TTTTACACT mRNA  3'  CTCCATC   GTCTA          miRNA 5'  GAGGTAG   TAGGT                   T               TGTATAGTT           3111111155521121333333333 ``` \| SeedType = 1 NonSeedType = 2 \| \| --- \| --- \| --- \| --- \| --- \| --- \| | \| 296 \| WBGene00004060:pmp-3(2574) \| ``` log2FC=-0.002 padj=0.999 ``` \| [seedLoc = 97,102] Seq=TGCTTTTGGACATCCCCTTTTTACTACTTCA \| ```                    TTTTCC  T   GGTTT   T mRNA  3' ACTTCATCAT      CC ACA     TCG  miRNA 5' TGAGGTAGTA      GG TGT     AGT                             T   AT      T           1112111111555555113111335551123 ``` \| SeedType = 3 NonSeedType = -1 \| \| --- \| --- \| --- \| --- \| --- \| --- \| | \| 297 \| WBGene00004086:pph-4.2(2591) \| ``` log2FC=0.217 padj=0.915 ``` \| [seedLoc = 242,248] Seq=CGCTCAAGTTCTCACCTCA \| ```                C     G         C mRNA  3' ACTCCA TC TT AAC   TCG  miRNA 5' TGAGGT AG AG TTG   AGT                    T  G   TAT   T           11111151161231116661123 ``` \| SeedType = 2 NonSeedType = 1 \| \| --- \| --- \| --- \| --- \| --- \| --- \| | \| 298 \| WBGene00004129:pqn-42(2634) \| ``` log2FC=-0.37 padj=0.875 ``` \| [seedLoc = 365,371] Seq=TGACAAGCAACCGCAGTTCGCCTACCTGC \| ```          CG      CGCTTGACG      AACAGT mRNA  3'   TCCATC         CCAACG       miRNA 5'   AGGTAG         GGTTGT                TG      TA             ATAGTT           33111111335554554111112333333 ``` \| SeedType = 3 NonSeedType = 1 \| \| --- \| --- \| --- \| --- \| --- \| --- \| | \| 299 \| WBGene00004132:ifet-1(2638) \| ``` log2FC=-0.015 padj=0.992 ``` \| [seedLoc = 208,214] Seq=TCAAAATTTCGCTTTACCGTCG \| ```          G  G        GCTTTAAAACT mRNA  3'  CT CCATT TC            miRNA 5'  GA GGTAG AG                     T        T  GTTGTATAGTT           31141111261133333333333 ``` \| SeedType = 3 NonSeedType = -1 \| \| --- \| --- \| --- \| --- \| --- \| --- \| | \|  \|  \|  \| [seedLoc = 731,737] Seq=AAATGCGAAAAGTTTTATAAGGGTACACTCA \| ```              A   G GAATATTTTGAAA      AA   mRNA  3' ACTC CAT G             AGCGTA     miRNA 5' TGAG GTA T             TTGTAT                      G AGG                AGTT           111151113233355555545551212113366 ``` \| SeedType = 2 NonSeedType = -1 \| \| --- \| --- \| --- \| --- \| --- \| --- \| | \| 300 \| WBGene00004143:pqn-59(2646) \| ``` log2FC=-0.02 padj=0.988 ``` \| [seedLoc = 49,52] Seq=CGACTTTGCTCCCCCTCCCTCT \| ```          T    C  CC  T    T    C mRNA  3'  CTCC TC  CC  CGT TCAG  miRNA 5'  GAGG AG  GG  GTA AGTT           T    T  TA  TT   T                31111311331136121311125 ``` \| SeedType = 2 NonSeedType = -1 \| \| --- \| --- \| --- \| --- \| --- \| --- \| | \| 301 \| WBGene00004147:larp-5(2650) \| ``` log2FC=-0.111 padj=0.905 ``` \| [seedLoc = 459,464] Seq=TATCGAACACCTCATTGTCTGTCCTATCTCT \| ```          T       CTGTCTGTTAC       AGCTAT mRNA  3'  CTCTATC           TCC ACA       miRNA 5'  GAGGTAG           AGG TGT                T       T             T   ATAGTT           31112111354555455551116111333333 ``` \| SeedType = 2 NonSeedType = -1 \| \| --- \| --- \| --- \| --- \| --- \| --- \| | \| 302 \| WBGene00004164:pqn-83(2667) \| ``` log2FC=0.392 padj=0.297 ``` \| [seedLoc = 124,129] Seq=AAACTCAACATTCCTCCTCACTATCTCG \| ```          G        CTCC   TT   AC    A mRNA  3'  CTCTATCA    TCC  ACA  TCAA  miRNA 5'  GAGGTAGT    AGG  TGT  AGTT           T               T    AT                3111211115555111351113311115 ``` \| SeedType = 2 NonSeedType = -1 \| \| --- \| --- \| --- \| --- \| --- \| --- \| | \| 303 \| WBGene00004166:pqn-85(2669) \| ``` log2FC=0.003 padj=0.998 ``` \| [seedLoc = 313,319] Seq=ATCTGGAATTTCTGGTACACTCT \| ```          T   A   G    TTAAG   TA mRNA  3'  CTC CAT GTCT     GTC   miRNA 5'  GAG GTA TAGG     TAG            T       G    TTGTA   TT           31115111321123333321133 ``` \| SeedType = 2 NonSeedType = -1 \| \| --- \| --- \| --- \| --- \| --- \| --- \| | \| 304 \| WBGene00004184:T19D2.2(2685) \| ``` log2FC=-0.464 padj=0.252 ``` \| [seedLoc = 109,114] Seq=CTCTGTTGCACCATACCTCT \| ```          T      A        T    TC mRNA  3'  CTCCAT   CC ACG TGTC   miRNA 5'  GAGGTA   GG TGT ATAG            T      GTA  T        TT           31111113661161125121133 ``` \| SeedType = 1 NonSeedType = -1 \| \| --- \| --- \| --- \| --- \| --- \| --- \| | \|  \|  \|  \| [seedLoc = 359,365] Seq=CGCTTTTACCATTTCCTTGATACCATCT \| ```          T  A    AGT   TTTAC   TT   C mRNA  3'  CT CCAT   TCC     CAT  TCG  miRNA 5'  GA GGTA   AGG     GTA  AGT           T       GT    TT      T    T           3115111133511133555111351123 ``` \| SeedType = 3 NonSeedType = -1 \| \| --- \| --- \| --- \| --- \| --- \| --- \| | \| 305 \| WBGene00004203:swsn-1(2703) \| ``` log2FC=-0.099 padj=0.907 ``` \| [seedLoc = 622,628] Seq=ACCCCACCAACACATATTACTCTCT \| ```          T   T      A A    CACCCCA mRNA  3'  CTC CATTAT C CAAC        miRNA 5'  GAG GTAGTA G GTTG                 T                 TATAGTT           3111511121151511113333333 ``` \| SeedType = 2 NonSeedType = 1 \| \| --- \| --- \| --- \| --- \| --- \| --- \| | \| 306 \| WBGene00004204:swsn-4(2704) \| ``` log2FC=0.012 padj=0.994 ``` \| [seedLoc = 185,190] Seq=TACTGTTTATCTCTACTTCT \| ```          T            TT      T mRNA  3'  CTTCATC TCTA   TGTCA  miRNA 5'  GAGGTAG AGGT   ATAGT           T       T    TGT     T           3112111161121336121113 ``` \| SeedType = 3 NonSeedType = 2 \| \| --- \| --- \| --- \| --- \| --- \| --- \| | \| 307 \| WBGene00004212:ptl-1(2713) \| ``` log2FC=-0.382 padj=0.676 ``` \| [seedLoc = 88,94] Seq=TTCCAGGCGCCATTTATCCTCT \| ```          T    T   TA      GACCTT mRNA  3'  CTCC ATT  CC GCG       miRNA 5'  GAGG TAG  GG TGT                T        TA  T   ATAGTT           31111511233116212333333 ``` \| SeedType = 2 NonSeedType = -1 \| \| --- \| --- \| --- \| --- \| --- \| --- \| | \| 308 \| WBGene00004213:ptp-1(2714) \| ``` log2FC=-0.228 padj=0.732 ``` \| [seedLoc = 219,224] Seq=ATTTCAACAAAAAATCTATCTCA \| ```                  TAAAA    ACTTTA mRNA  3' ACTCTATC     AACA       miRNA 5' TGAGGTAG     TTGT                        TAGG     ATAGTT           11112111333351111333333 ``` \| SeedType = 2 NonSeedType = 1 \| \| --- \| --- \| --- \| --- \| --- \| --- \| | \| 309 \| WBGene00004220:ptr-5(2722) \| ``` log2FC=-0.052 padj=0.985 ``` \| [seedLoc = 254,259] Seq=GAAATAACAATAACTACTTCC \| ```          C        A      ATAAAG mRNA  3'  CTTCATCA  TAACA       miRNA 5'  GAGGTAGT  GTTGT                T        AG     ATAGTT           3112111113621111333333 ``` \| SeedType = 3 NonSeedType = 1 \| \| --- \| --- \| --- \| --- \| --- \| --- \| | \| 310 \| WBGene00004245:puf-9(2747) \| ``` log2FC=-0.211 padj=0.52 ``` \| [seedLoc = 810,816] Seq=TAATCCAAAACCAATACCATCA \| ```             A    AA     AACCTAAT mRNA  3' ACT CCAT   CCAA         miRNA 5' TGA GGTA   GGTT                          GTA    GTATAGTT           11151111336111133333333 ``` \| SeedType = 3 NonSeedType = 1 \| \| --- \| --- \| --- \| --- \| --- \| --- \| | \|  \|  \|  \| [seedLoc = 1052,1057] Seq=GAACTTTAACGACTACTGTTTTCCTATCTCT \| ```          T       CTTTTGTCA      ATT    G mRNA  3'  CTCTATC         TCAGCA   TCAA  miRNA 5'  GAGGTAG         GGTTGT   AGTT           T       TA             AT                 3111211133555455521121133511114 ``` \| SeedType = 2 NonSeedType = 2 \| \| --- \| --- \| --- \| --- \| --- \| --- \| | \| 311 \| WBGene00004254:pxf-1(2752) \| ``` log2FC=0.098 padj=0.936 ``` \| [seedLoc = 25,31] Seq=CACACAACTTCTTCTTCTTCTACTACCTGC \| ```          CG         TTCTTCTTCTT     CAC    mRNA  3'   TCCATCATC           CAACA       miRNA 5'   AGGTAGTAG           GTTGT                TG                         ATAGTT           331111111115555555555511111333666 ``` \| SeedType = 3 NonSeedType = 1 \| \| --- \| --- \| --- \| --- \| --- \| --- \| | \|  \|  \|  \| [seedLoc = 349,354] Seq=ATTGCCTCAATCGCCGCTACCTCG \| ```          G        CCG     TCCGTTA mRNA  3'  CTCCATCG   CTAAC        miRNA 5'  GAGGTAGT   GGTTG                 T        A       TATAGTT           311111112354121113333333 ``` \| SeedType = 1 NonSeedType = 1 \| \| --- \| --- \| --- \| --- \| --- \| --- \| | \| 312 \| WBGene00004268:rab-5(2764) \| ``` log2FC=0.015 padj=0.993 ``` \| [seedLoc = 43,46] Seq=GTACCATCAACCCTTCTAGCTCC \| ```          C   G   TTC     TACCATG mRNA  3'  CTC ATC   CCAAC        miRNA 5'  GAG TAG   GGTTG                 T   G   TA      TATAGTT           31113111335111113333333 ``` \| SeedType = 2 NonSeedType = 1 \| \| --- \| --- \| --- \| --- \| --- \| --- \| | \| 313 \| WBGene00004271:rab-7(2767) \| ``` log2FC=-0.138 padj=0.794 ``` \| [seedLoc = 27,32] Seq=CGCGCAGAGCCCTCTATCTCT \| ```          T       TC    G CG   C mRNA  3'  CTCTATC  CCGA A   CG  miRNA 5'  GAGGTAG  GGTT T   GT           T       TA    G ATA  T           3111211133112131336123 ``` \| SeedType = 2 NonSeedType = 2 \| \| --- \| --- \| --- \| --- \| --- \| --- \| | \| 314 \| WBGene00004286:rab-39(2780) \| ``` log2FC=-0.225 padj=0.65 ``` \| [seedLoc = 132,138] Seq=GTGTCACCAACCTCACCTACTCTCC \| ```          C   T    CAC      CACTGTG mRNA  3'  CTC CATC   TCCAAC        miRNA 5'  GAG GTAG   AGGTTG                 T        T        TATAGTT           3111511113551111113333333 ``` \| SeedType = 2 NonSeedType = 1 \| \| --- \| --- \| --- \| --- \| --- \| --- \| | \| 315 \| WBGene00004296:rad-50(2781) \| ``` log2FC=-0.101 padj=0.903 ``` \| [seedLoc = 118,123] Seq=GTTTCCAAATGTTGTTACCTCA \| ```                     GTAAACCTTTG mRNA  3' ACTCCATTGTT            miRNA 5' TGAGGTAGTAG                                GTTGTATAGTT           1111111221233333333333 ``` \| SeedType = 1 NonSeedType = -1 \| \| --- \| --- \| --- \| --- \| --- \| --- \| | \| 316 \| WBGene00004313:rbc-1(2797) \| ``` log2FC=0.103 padj=0.895 ``` \| [seedLoc = 114,119] Seq=CGCTTCAAACGTCTACTTACCTCT \| ```          T     T     TG   ACT    C mRNA  3'  CTCCA TCATC  CAA    TCG  miRNA 5'  GAGGT AGTAG  GTT    AGT           T                GTAT   T           3111115111115411133361123 ``` \| SeedType = 2 NonSeedType = 1 \| \| --- \| --- \| --- \| --- \| --- \| --- \| | \| 317 \| WBGene00004328:rdy-2(2810) \| ``` log2FC=-0.32 padj=0.946 ``` \| [seedLoc = 63,69] Seq=TTCGAGGCAATCGTACCATCC \| ```          C  A    G        GAGCTT mRNA  3'  CT CCAT   CTAACG       miRNA 5'  GA GGTA   GGTTGT                T       GTA      ATAGTT           31151111366121112333333 ``` \| SeedType = 3 NonSeedType = 1 \| \| --- \| --- \| --- \| --- \| --- \| --- \| | \| 318 \| WBGene00004335:ref-2(2814) \| ``` log2FC=-0.209 padj=1.0 ``` \| [seedLoc = 110,115] Seq=CAGAGAACATCCCTCATATACCTCC \| ```          C      ATAC   CT   AGAGAC mRNA  3'  CTCCAT    TCC  ACA       miRNA 5'  GAGGTA    AGG  TGT                T      GT     T    ATAGTT           3111111335511135111333333 ``` \| SeedType = 1 NonSeedType = -1 \| \| --- \| --- \| --- \| --- \| --- \| --- \| | \| 319 \| WBGene00004367:ric-8(2842) \| ``` log2FC=0.346 padj=0.345 ``` \| [seedLoc = 262,268] Seq=TTTTGTATCAATTTTACCATCT \| ```          T  A            T     TT mRNA  3'  CT CCAT  TTTAAC ATGTT   miRNA 5'  GA GGTA  AGGTTG TATAG            T       GT            TT           311511116612211151121233 ``` \| SeedType = 3 NonSeedType = 1 \| \| --- \| --- \| --- \| --- \| --- \| --- \| | \| 320 \| WBGene00004370:rig-3(2846) \| ``` log2FC=-0.113 padj=0.952 ``` \| [seedLoc = 124,129] Seq=ATCAATATCATCCCCTATCTCG \| ```          G       CC   C    ACTA mRNA  3'  CTCTATC  CTA TATA     miRNA 5'  GAGGTAG  GGT GTAT              T       TA   T    AGTT           3111211133121321113333 ``` \| SeedType = 2 NonSeedType = 2 \| \| --- \| --- \| --- \| --- \| --- \| --- \| | \|  \|  \|  \| [seedLoc = 132,135] Seq=ATCAATATCATCCCCTATCTCGT \| ```          T        CC   C    ACTA mRNA  3'  GCTCTATC  CTA TATA     miRNA 5'  TGAGGTAG  GGT GTAT                       TA   T    AGTT           52111211133121321113333 ``` \| SeedType = 2 NonSeedType = 2 \| \| --- \| --- \| --- \| --- \| --- \| --- \| | \| 321 \| WBGene00004382:rnh-1.0(2858) \| ``` log2FC=0.041 padj=0.984 ``` \| [seedLoc = 28,33] Seq=ATTTTTACAATTGTTTTTCTGCCTCA \| ```                  TTTTTG       TTTTA mRNA  3' ACTCCGTC      TTAACAT      miRNA 5' TGAGGTAG      GGTTGTA                       TA           TAGTT           11111211335554221111133333 ``` \| SeedType = 2 NonSeedType = 1 \| \| --- \| --- \| --- \| --- \| --- \| --- \| | \| 322 \| WBGene00004383:rnh-2(2859) \| ``` log2FC=-0.004 padj=0.998 ``` \| [seedLoc = 118,123] Seq=CACTTTGGTTGTATACCTCA \| ```                 ATGTTG    T   C mRNA  3' ACTCCAT        GT TCA  miRNA 5' TGAGGTA        TA AGT                  GTAGGTTG  T   T           1111111333333662131113 ``` \| SeedType = 1 NonSeedType = -1 \| \| --- \| --- \| --- \| --- \| --- \| --- \| | \| 323 \| WBGene00004390:rnp-7(2868) \| ``` log2FC=-0.059 padj=0.961 ``` \| [seedLoc = 92,97] Seq=TTTCCTTCTGCTTTCTATCTCG \| ```          G       T    TCTTCCTTT mRNA  3'  CTCTATC TTCG          miRNA 5'  GAGGTAG AGGT                   T       T    TGTATAGTT           3111211131212333333333 ``` \| SeedType = 2 NonSeedType = 2 \| \| --- \| --- \| --- \| --- \| --- \| --- \| | \|  \|  \|  \| [seedLoc = 100,103] Seq=TTTCCTTCTGCTTTCTATCTCGT \| ```          T        T    TCTTCCTTT mRNA  3'  GCTCTATC TTCG          miRNA 5'  TGAGGTAG AGGT                            T    TGTATAGTT           52111211131212333333333 ``` \| SeedType = 2 NonSeedType = 2 \| \| --- \| --- \| --- \| --- \| --- \| --- \| | \| 324 \| WBGene00004393:rnt-1(2872) \| ``` log2FC=-0.497 padj=1.0 ``` \| [seedLoc = 70,75] Seq=CAAATCTCAATTGTTTCCTACCTCT \| ```          T       CTTTG     TCTAAAC mRNA  3'  CTCCATC     TTAAC        miRNA 5'  GAGGTAG     GGTTG                 T       TA        TATAGTT           3111111133554221113333333 ``` \| SeedType = 1 NonSeedType = 1 \| \| --- \| --- \| --- \| --- \| --- \| --- \| | \| 325 \| WBGene00004395:rol-3(2874) \| ``` log2FC=0.69 padj=0.083 over-expressed ``` \| [seedLoc = 49,54] Seq=CATCAAAGAATCTTTGTTTTCTATCTCA \| ```                  TTTTGTT     G AACTAC mRNA  3' ACTCTATC       TCTAA A       miRNA 5' TGAGGTAG       AGGTT T                        T           G ATAGTT           1111211135554551121131333333 ``` \| SeedType = 2 NonSeedType = 2 \| \| --- \| --- \| --- \| --- \| --- \| --- \| | \| 326 \| WBGene00004459:rpn-2(2935) \| ``` log2FC=-0.008 padj=0.994 ``` \| [seedLoc = 27,32] Seq=ATTCGATCAACTCTATCTCG \| ```          G              TAGCTTA mRNA  3'  CTCTATC  TCAAC        miRNA 5'  GAGGTAG  GGTTG                 T       TA     TATAGTT           3111211166211113333333 ``` \| SeedType = 2 NonSeedType = 1 \| \| --- \| --- \| --- \| --- \| --- \| --- \| | \| 327 \| WBGene00004463:rpn-7(2939) \| ``` log2FC=0.009 padj=0.994 ``` \| [seedLoc = 133,139] Seq=AATGAATTAATTTATTACCATCG \| ```          G  A            TAAGTAA mRNA  3'  CT CCATTATTTAAT        miRNA 5'  GA GGTAGTAGGTTG                 T               TATAGTT           31151111211221123333333 ``` \| SeedType = 3 NonSeedType = 2 \| \| --- \| --- \| --- \| --- \| --- \| --- \| | \|  \|  \|  \| [seedLoc = 485,490] Seq=TATTTTAAGCCCTCAACTACTTCA \| ```                   AC   CGAATTTTAT mRNA  3' ACTTCATCA  TCC           miRNA 5' TGAGGTAGT  AGG                                  TTGTATAGTT           111211111551113333333333 ``` \| SeedType = 3 NonSeedType = -1 \| \| --- \| --- \| --- \| --- \| --- \| --- \| | \|  \|  \|  \| [seedLoc = 602,608] Seq=AAATTTTACAAGTAGACTAACCTCA \| ```               A    GA G     T    A mRNA  3' ACTCC ATCA  T AACAT TTAA  miRNA 5' TGAGG TAGT  G TTGTA AGTT                     A  G     T                1111151111352311111312115 ``` \| SeedType = 2 NonSeedType = 1 \| \| --- \| --- \| --- \| --- \| --- \| --- \| | \| 328 \| WBGene00004726:sas-4(3016) \| ``` log2FC=-0.021 padj=0.993 ``` \| [seedLoc = 95,100] Seq=AAATTATAATTAATTCTACCTCC \| ```          C       TTAATTA       A mRNA  3'  CTCCATC       ATATTAA  miRNA 5'  GAGGTAG       TATAGTT           T       TAGGTTG                   31111111333333311112115 ``` \| SeedType = 1 NonSeedType = -1 \| \| --- \| --- \| --- \| --- \| --- \| --- \| | \|  \|  \|  \| [seedLoc = 138,143] Seq=AATAATTCAATCATTTTTTACTATCTCA \| ```                    TTTTTA     TTAATAA mRNA  3' ACTCTATCAT      CTAAC        miRNA 5' TGAGGTAGTA      GGTTG                                      TATAGTT           1111211111555555121113333333 ``` \| SeedType = 2 NonSeedType = 1 \| \| --- \| --- \| --- \| --- \| --- \| --- \| | \| 329 \| WBGene00004727:sax-1(3017) \| ``` log2FC=-0.328 padj=0.414 ``` \| [seedLoc = 239,245] Seq=ACACAAACATTTTCTTACTCTCT \| ```          T   T    C   T   AACACA mRNA  3'  CTC CATT TTT ACA       miRNA 5'  GAG GTAG AGG TGT                T        T   T   ATAGTT           31115111231223111333333 ``` \| SeedType = 2 NonSeedType = -1 \| \| --- \| --- \| --- \| --- \| --- \| --- \| | \| 330 \| WBGene00004739:scd-1(3027) \| ``` log2FC=-0.232 padj=0.644 ``` \| [seedLoc = 355,360] Seq=TTTTGTTTGTTTTTCTACCTTC \| ```          C       TTTTTGTTTGTTTT mRNA  3'  TTCCATC               miRNA 5'  GAGGTAG                        T       TAGGTTGTATAGTT           3211111133333333333333 ``` \| SeedType = 3 NonSeedType = -1 \| \| --- \| --- \| --- \| --- \| --- \| --- \| | \|  \|  \|  \| [seedLoc = 536,541] Seq=ATTGGTGGAACTGTTCTGTACCTCT \| ```          T      GTCTTG    G   GTTA mRNA  3'  CTCCAT      TCAA GTG     miRNA 5'  GAGGTA      GGTT TAT              T      GTA       G   AGTT           3111111333554211132123333 ``` \| SeedType = 1 NonSeedType = 1 \| \| --- \| --- \| --- \| --- \| --- \| --- \| | \|  \|  \|  \| [seedLoc = 597,602] Seq=AAAATTTGTCCCTTCTATCTCA \| ```                  T   CTGTTTAAAA mRNA  3' ACTCTATC TCC           miRNA 5' TGAGGTAG AGG                            T   TTGTATAGTT           1111211131113333333333 ``` \| SeedType = 2 NonSeedType = -1 \| \| --- \| --- \| --- \| --- \| --- \| --- \| | \| 331 \| WBGene00004751:sea-2(3037) \| ``` log2FC=-0.084 padj=0.959 ``` \| [seedLoc = 240,246] Seq=CAGCTGTTGCTGGATGTGCATACGCTCA \| ```              G   A   G   GT  T      C mRNA  3' ACTC CAT CGT TAG  CG TGTCGA  miRNA 5' TGAG GTA GTA GTT  GT ATAGTT                       G                          1111411151213212451251211215 ``` \| SeedType = 2 NonSeedType = -1 \| \| --- \| --- \| --- \| --- \| --- \| --- \| | \| 332 \| WBGene00004756:sec-24.2(3042) \| ``` log2FC=0.171 padj=0.655 ``` \| [seedLoc = 317,322] Seq=CCATTTTTGGCCCTGCCTCA \| ```                        TTTTTACC mRNA  3' ACTCCGTC  CCGG         miRNA 5' TGAGGTAG  GGTT                          TA    GTATAGTT           1111121166112233333333 ``` \| SeedType = 2 NonSeedType = 2 \| \| --- \| --- \| --- \| --- \| --- \| --- \| | \| 333 \| WBGene00004758:sek-1(3043) \| ``` log2FC=-0.044 padj=0.98 ``` \| [seedLoc = 504,509] Seq=TAGTAGTCATTCCTATCTCA \| ```                      T  TG  G T mRNA  3' ACTCTATC  CT AC  AT A  miRNA 5' TGAGGTAG  GG TG  TA T                   TA  T  TA  G T           1111211166123113311313 ``` \| SeedType = 2 NonSeedType = -1 \| \| --- \| --- \| --- \| --- \| --- \| --- \| | \| 334 \| WBGene00004764:sel-7(3048) \| ``` log2FC=-0.224 padj=0.956 ``` \| [seedLoc = 151,156] Seq=CATTTCAAAACCACACCTACCTTA \| ```                  CACA    AACTTTAC mRNA  3' ATTCCATC    CCAA         miRNA 5' TGAGGTAG    GGTT                          TA      GTATAGTT           121111113355111133333333 ``` \| SeedType = 3 NonSeedType = 1 \| \| --- \| --- \| --- \| --- \| --- \| --- \| | \|  \|  \|  \| [seedLoc = 576,580] Seq=GTCCCTACAAAAAAAAACTTCCTCAC \| ```          C     T   AAAAAA     CCCTG mRNA  3'  ACTCC TCA      AACAT      miRNA 5'  TGAGG AGT      TTGTA                     T   AGG        TAGTT           51111131113335551111133333 ``` \| SeedType = 2 NonSeedType = 1 \| \| --- \| --- \| --- \| --- \| --- \| --- \| | \|  \|  \|  \| [seedLoc = 651,656] Seq=CTCATACTTTCCGTTAGTTTTTCCTACCTCG \| ```          G       CTTTTTGATTG  TTT    CTC  mRNA  3'  CTCCATC           CC   CATA     miRNA 5'  GAGGTAG           GG   GTAT              T       TA           TT     AGTT           31111111335555455541133511113336 ``` \| SeedType = 1 NonSeedType = -1 \| \| --- \| --- \| --- \| --- \| --- \| --- \| | \| 335 \| WBGene00004771:sem-2(3054) \| ``` log2FC=-0.639 padj=0.006 ``` \| [seedLoc = 126,132] Seq=ATTTTCTGCTCCTACCTCAA \| ```          A        C  GTCTTTTA    mRNA  3'  ACTCCATC TC            miRNA 5'  TGAGGTAG AG                              T  GTTGTATAGTT           51111111131133333333666 ``` \| SeedType = 1 NonSeedType = -1 \| \| --- \| --- \| --- \| --- \| --- \| --- \| | \| 336 \| WBGene00004774:sem-5(3056) \| ``` log2FC=0.153 padj=0.861 ``` \| [seedLoc = 666,671] Seq=CATGAAGCTTCTCTCTGCCTCC \| ```          C       TC  TT  AAGTAC mRNA  3'  CTCCGTC  TC  CG       miRNA 5'  GAGGTAG  GG  GT                T       TA  TT  ATAGTT           3111121133213312333333 ``` \| SeedType = 2 NonSeedType = -1 \| \| --- \| --- \| --- \| --- \| --- \| --- \| | \| 337 \| WBGene00004788:sft-4(3069) \| ``` log2FC=-0.016 padj=0.993 ``` \| [seedLoc = 52,58] Seq=TTCTATGTCAAATCATTGCATTACACTCA \| ```              A   TA   TA  A   T     TT mRNA  3' ACTC CAT  CGT  CT AAC GTATC   miRNA 5' TGAG GTA  GTA  GG TTG TATAG                                       TT           11115111551215512511152111133 ``` \| SeedType = 2 NonSeedType = 1 \| \| --- \| --- \| --- \| --- \| --- \| --- \| | \| 338 \| WBGene00004795:sid-1(3073) \| ``` log2FC=0.087 padj=0.966 ``` \| [seedLoc = 164,169] Seq=ACTGTATTCATCTCCACGGCTCCCTACCTCC \| ```          C       CC      CCTCTACTTA       mRNA  3'  CTCCATC  TCGGCA          TGTCA  miRNA 5'  GAGGTAG  GGTTGT          ATAGT           T       TA                     T           31111111332122115555555555121116 ``` \| SeedType = 1 NonSeedType = 2 \| \| --- \| --- \| --- \| --- \| --- \| --- \| | \| 339 \| WBGene00004804:skn-1(3080) \| ``` log2FC=0.013 padj=0.993 ``` \| [seedLoc = 165,170] Seq=GATTTCCCAGTGAAATACCTCA \| ```                 AAAGT   CCTTTAG mRNA  3' ACTCCAT     GAC        miRNA 5' TGAGGTA     TTG                        GTAGG   TATAGTT           1111111333332113333333 ``` \| SeedType = 1 NonSeedType = -1 \| \| --- \| --- \| --- \| --- \| --- \| --- \| | \|  \|  \|  \| [seedLoc = 175,179] Seq=GATTTCCCAGTGAAATACCTCAC \| ```          C       AAAGT   CCTTTAG mRNA  3'  ACTCCAT     GAC        miRNA 5'  TGAGGTA     TTG                         GTAGG   TATAGTT           51111111333332113333333 ``` \| SeedType = 1 NonSeedType = -1 \| \| --- \| --- \| --- \| --- \| --- \| --- \| | \| 340 \| WBGene00004809:skr-3(3085) \| ``` log2FC=-0.075 padj=0.945 ``` \| [seedLoc = 49,54] Seq=ATCTGATTCGGTTTACATACCTCT \| ```          T      A        TTA   TA mRNA  3'  CTCCAT CATTTGGC   GTC   miRNA 5'  GAGGTA GTAGGTTG   TAG            T               TA    TT           311111151112222133521133 ``` \| SeedType = 1 NonSeedType = 2 \| \| --- \| --- \| --- \| --- \| --- \| --- \| | \| 341 \| WBGene00004810:skr-4(3086) \| ``` log2FC=-0.012 padj=0.994 ``` \| [seedLoc = 49,54] Seq=ATCTGATTCGGTTTACATACCTCT \| ```          T      A        TTA   TA mRNA  3'  CTCCAT CATTTGGC   GTC   miRNA 5'  GAGGTA GTAGGTTG   TAG            T               TA    TT           311111151112222133521133 ``` \| SeedType = 1 NonSeedType = 2 \| \| --- \| --- \| --- \| --- \| --- \| --- \| | \| 342 \| WBGene00004827:skr-21(3103) \| ``` log2FC=-0.591 padj=0.773 ``` \| [seedLoc = 106,111] Seq=CTTTTCACTGCCTTCCTGCCTCA \| ```                  CT    T  CTTTTC mRNA  3' ACTCCGTC  TCCG CA       miRNA 5' TGAGGTAG  AGGT GT                        T     T  ATAGTT           11111211351112311333333 ``` \| SeedType = 2 NonSeedType = 2 \| \| --- \| --- \| --- \| --- \| --- \| --- \| | \|  \|  \|  \| [seedLoc = 112,116] Seq=CTTTTCACTGCCTTCCTGCCTCAT \| ```          T        CT    T  CTTTTC mRNA  3'  ACTCCGTC  TCCG CA       miRNA 5'  TGAGGTAG  AGGT GT                         T     T  ATAGTT           511111211351112311333333 ``` \| SeedType = 2 NonSeedType = 2 \| \| --- \| --- \| --- \| --- \| --- \| --- \| | \| 343 \| WBGene00004829:sli-1(3104) \| ``` log2FC=0.304 padj=0.68 ``` \| [seedLoc = 266,272] Seq=TCATGTTAATTTCTTACTCTCG \| ```          G   T    C          ACT mRNA  3'  CTC CATT TTTAAT TGT    miRNA 5'  GAG GTAG AGGTTG ATA             T        T      T   GTT           31115111231221126121333 ``` \| SeedType = 2 NonSeedType = 2 \| \| --- \| --- \| --- \| --- \| --- \| --- \| | \| 344 \| WBGene00004856:sma-2(3109) \| ``` log2FC=-0.109 padj=0.955 ``` \| [seedLoc = 28,33] Seq=TTTCAAATAACACTACCTTG \| ```          G               AACTTT mRNA  3'  TTCCATCA  CAATA       miRNA 5'  GAGGTAGT  GTTGT                T        AG     ATAGTT           3211111116611121333333 ``` \| SeedType = 3 NonSeedType = 1 \| \| --- \| --- \| --- \| --- \| --- \| --- \| | \| 345 \| WBGene00004873:smc-3(3117) \| ``` log2FC=-0.033 padj=0.98 ``` \| [seedLoc = 70,75] Seq=CACCACCCACACTATCTACCTTA \| ```                  T   AC  CCACCAC mRNA  3' ATTCCATC ATC  AC        miRNA 5' TGAGGTAG TAG  TG                             GT  TATAGTT           12111111511133113333333 ``` \| SeedType = 3 NonSeedType = -1 \| \| --- \| --- \| --- \| --- \| --- \| --- \| | \| 346 \| WBGene00004879:smg-1(3122) \| ``` log2FC=-0.039 padj=0.98 ``` \| [seedLoc = 30,35] Seq=CACGTGTCATAGTCCGTTCTACTTCT \| ```          T       TTG  T    C   G  C mRNA  3'  CTTCATC   CC GATA TGT CA  miRNA 5'  GAGGTAG   GG TTGT ATA GT           T       TA               T           31121111334115212151214113 ``` \| SeedType = 3 NonSeedType = -1 \| \| --- \| --- \| --- \| --- \| --- \| --- \| | \|  \|  \|  \| [seedLoc = 374,380] Seq=TTGCATTCAACTCATACTCTCC \| ```          C   T   AC      TTACGTT mRNA  3'  CTC CAT   TCAAC        miRNA 5'  GAG GTA   GGTTG                 T       GTA     TATAGTT           31115111336211113333333 ``` \| SeedType = 2 NonSeedType = 1 \| \| --- \| --- \| --- \| --- \| --- \| --- \| | \|  \|  \|  \| [seedLoc = 545,550] Seq=TTCTTATTTGGATATTACCTCG \| ```          G         AGGTTTATTCTT mRNA  3'  CTCCATTAT             miRNA 5'  GAGGTAGTA                      T         GGTTGTATAGTT           3111111211333333333333 ``` \| SeedType = 1 NonSeedType = -1 \| \| --- \| --- \| --- \| --- \| --- \| --- \| | \| 347 \| WBGene00004881:smg-3(3125) \| ``` log2FC=-0.012 padj=0.994 ``` \| [seedLoc = 54,59] Seq=CGACTAGTCTCTTCAACCCTGCCTCT \| ```          T              TTCTCTG     C mRNA  3'  CTCCGTC  CCAAC       ATCAG  miRNA 5'  GAGGTAG  GGTTG       TAGTT           T       TA     TA                      3111121166111113355554111125 ``` \| SeedType = 2 NonSeedType = 1 \| \| --- \| --- \| --- \| --- \| --- \| --- \| | \| 348 \| WBGene00004881:smg-3(3126) \| ``` log2FC=-0.012 padj=0.994 ``` \| [seedLoc = 60,65] Seq=TACTAGTCTCTTCAACCCTGCCTCT \| ```          T              TTCTCTG    T mRNA  3'  CTCCGTC  CCAAC       ATCA  miRNA 5'  GAGGTAG  GGTTG       TAGT           T       TA     TA         T           311112116611111335555411113 ``` \| SeedType = 2 NonSeedType = 1 \| \| --- \| --- \| --- \| --- \| --- \| --- \| | \| 349 \| WBGene00004881:smg-3(3127) \| ``` log2FC=-0.012 padj=0.994 ``` \| [seedLoc = 56,61] Seq=TACTAGTCTCTTCAACCCTGCCTCT \| ```          T              TTCTCTG    T mRNA  3'  CTCCGTC  CCAAC       ATCA  miRNA 5'  GAGGTAG  GGTTG       TAGT           T       TA     TA         T           311112116611111335555411113 ``` \| SeedType = 2 NonSeedType = 1 \| \| --- \| --- \| --- \| --- \| --- \| --- \| | \| 350 \| WBGene00004881:smg-3(3128) \| ``` log2FC=-0.012 padj=0.994 ``` \| [seedLoc = 53,58] Seq=GTCTCTTCAACCCTGCCTCT \| ```          T              TTCTCTG mRNA  3'  CTCCGTC  CCAAC        miRNA 5'  GAGGTAG  GGTTG                 T       TA     TATAGTT           3111121166111113333333 ``` \| SeedType = 2 NonSeedType = 1 \| \| --- \| --- \| --- \| --- \| --- \| --- \| | \| 351 \| WBGene00004882:smg-4(3129) \| ``` log2FC=0.002 padj=0.999 ``` \| [seedLoc = 170,176] Seq=CATCTTATGATTTTCTACCCTCC \| ```          C  C     T        TCTAC mRNA  3'  CT CCATC TTTAGTAT      miRNA 5'  GA GGTAG AGGTTGTA               T        T        TAGTT           31151111131221221133333 ``` \| SeedType = 3 NonSeedType = 2 \| \| --- \| --- \| --- \| --- \| --- \| --- \| | \| 352 \| WBGene00004884:smg-6(3133) \| ``` log2FC=-0.101 padj=0.918 ``` \| [seedLoc = 68,73] Seq=AAAAATATCCCTTTTTTTACCTCG \| ```          G       TTTT   C     AAAA mRNA  3'  CTCCATT    TCC  TATA     miRNA 5'  GAGGTAG    AGG  GTAT              T       T      TT    AGTT           3111111235551113621113333 ``` \| SeedType = 1 NonSeedType = -1 \| \| --- \| --- \| --- \| --- \| --- \| --- \| | \| 353 \| WBGene00004889:smp-1(3138) \| ``` log2FC=0.123 padj=0.955 ``` \| [seedLoc = 74,80] Seq=GAAAGCACGGCTTAATTGTTCTCTACCATCC \| ```          C  A     TCTTGTTA        CGAAAG mRNA  3'  CT CCATC        ATTCGGCA       miRNA 5'  GA GGTAG        TAGGTTGT                T                        ATAGTT           3115111115555455511212211333333 ``` \| SeedType = 3 NonSeedType = 2 \| \| --- \| --- \| --- \| --- \| --- \| --- \| | \|  \|  \|  \| [seedLoc = 476,481] Seq=GTGCAGATCTAACTCAG \| ```          G    A   TA             mRNA  3'  ACTC ATC    GACGTG     miRNA 5'  TGAG TAG    TTGTAT                   G   TAGG      AGTT           41111311133662112126666 ``` \| SeedType = 2 NonSeedType = -1 \| \| --- \| --- \| --- \| --- \| --- \| --- \| | \|  \|  \|  \| [seedLoc = 1296,1301] Seq=GGGACAAAAACGCTATTGTTACCTCA \| ```                 TGT    G   AAACAGGG mRNA  3' ACTCCAT   TATC CAA         miRNA 5' TGAGGTA   GTAG GTT                                    GTATAGTT           11111115452111411133333333 ``` \| SeedType = 1 NonSeedType = 1 \| \| --- \| --- \| --- \| --- \| --- \| --- \| | \| 354 \| WBGene00004893:sms-2(3143) \| ``` log2FC=-0.025 padj=0.994 ``` \| [seedLoc = 185,190] Seq=CCCAAAGCGGTTTTTCCTACTTCT \| ```          T       CTT       AAACCC mRNA  3'  CTTCATC   TTTGGCG       miRNA 5'  GAGGTAG   AGGTTGT                T       T         ATAGTT           311211113551222212333333 ``` \| SeedType = 3 NonSeedType = 2 \| \| --- \| --- \| --- \| --- \| --- \| --- \| | \| 355 \| WBGene00004910:snf-11(3154) \| ``` log2FC=0.287 padj=0.794 ``` \| [seedLoc = 79,84] Seq=CACTTATTCCTAACTATCTCT \| ```          T       A    T        C mRNA  3'  CTCTATC ATCC  TAT TCA  miRNA 5'  GAGGTAG TAGG  GTA AGT           T            TT   T   T           31112111511113621161113 ``` \| SeedType = 2 NonSeedType = -1 \| \| --- \| --- \| --- \| --- \| --- \| --- \| | \| 356 \| WBGene00004915:snr-2(3159) \| ``` log2FC=-0.12 padj=0.861 ``` \| [seedLoc = 117,123] Seq=TAATTATATATTTAATTTATCGATACTCTCT \| ```          T   T   AGC      ATTT         T mRNA  3'  CTC CAT   TATTTA    ATATATTAA  miRNA 5'  GAG GTA   GTAGGT    TGTATAGTT           T                                         3111511154521122155551211112115 ``` \| SeedType = 2 NonSeedType = 2 \| \| --- \| --- \| --- \| --- \| --- \| --- \| | \| 357 \| WBGene00004949:sox-2(3184) \| ``` log2FC=-0.077 padj=0.982 ``` \| [seedLoc = 169,174] Seq=TGCTAAATGACTTTTTACCTCT \| ```          T       T       A    T mRNA  3'  CTCCATT TTCAGTA ATCG  miRNA 5'  GAGGTAG AGGTTGT TAGT           T       T       A    T           3111111231211221311123 ``` \| SeedType = 1 NonSeedType = 2 \| \| --- \| --- \| --- \| --- \| --- \| --- \| | \|  \|  \|  \| [seedLoc = 680,686] Seq=AAAATTTAGATGTTGATACCATCA \| ```             A    AGT G   ATTTAAAA mRNA  3' ACT CCAT   T TAG         miRNA 5' TGA GGTA   A GTT                          GT  G   GTATAGTT           111511113351321233333333 ``` \| SeedType = 3 NonSeedType = -1 \| \| --- \| --- \| --- \| --- \| --- \| --- \| | \| 358 \| WBGene00004980:spk-1(3207) \| ``` log2FC=0.036 padj=0.979 ``` \| [seedLoc = 300,305] Seq=CAACAATCATCCGACTACTTCA \| ```                   G  T     A   C mRNA  3' ACTTCATCA CC AC TA CAA  miRNA 5' TGAGGTAGT GG TG AT GTT                    A  T  T  A               11121111131131161131115 ``` \| SeedType = 3 NonSeedType = -1 \| \| --- \| --- \| --- \| --- \| --- \| --- \| | \|  \|  \|  \| [seedLoc = 306,310] Seq=CAACAATCATCCGACTACTTCAT \| ```          T         G  T     A   C mRNA  3'  ACTTCATCA CC AC TA CAA  miRNA 5'  TGAGGTAGT GG TG AT GTT                     A  T  T  A               511121111131131161131115 ``` \| SeedType = 3 NonSeedType = -1 \| \| --- \| --- \| --- \| --- \| --- \| --- \| | \| 359 \| WBGene00004984:spn-4(3209) \| ``` log2FC=-0.075 padj=0.972 ``` \| [seedLoc = 320,325] Seq=AATATACTACCAGTCCGTTTAACCTACCTCG \| ```          G       CAATTTGCCTGA   T     AA  mRNA  3'  CTCCATC            CCA CATAT    miRNA 5'  GAGGTAG            GGT GTATA             T       TA             T     GTT           31111111335555455545111311111336 ``` \| SeedType = 1 NonSeedType = 1 \| \| --- \| --- \| --- \| --- \| --- \| --- \| | \| 360 \| WBGene00005008:spr-3(3229) \| ``` log2FC=-0.3 padj=0.768 ``` \| [seedLoc = 191,196] Seq=CGCTCAAATAGCTGCCTCA \| ```                   ATA         C mRNA  3' ACTCCGTCG   AAC   TCG  miRNA 5' TGAGGTAGT   TTG   AGT                    AGG   TAT   T           1111121123331116661123 ``` \| SeedType = 2 NonSeedType = 1 \| \| --- \| --- \| --- \| --- \| --- \| --- \| | \|  \|  \|  \| [seedLoc = 288,293] Seq=TTTCATTCATCCTGGTACCTCA \| ```                 G    T  TTACTTT mRNA  3' ACTCCAT GTCC AC        miRNA 5' TGAGGTA TAGG TG                        G    T  TATAGTT           1111111321113113333333 ``` \| SeedType = 1 NonSeedType = -1 \| \| --- \| --- \| --- \| --- \| --- \| --- \| | \| 361 \| WBGene00005040:sra-14(3257) \| ``` log2FC=-0.012 padj=0.998 ``` \| [seedLoc = 46,51] Seq=AAATAAAAATTCTCTACTTCC \| ```          C           TAAAAATAAA mRNA  3'  CTTCATC TCT           miRNA 5'  GAGGTAG AGG                    T       T   TTGTATAGTT           3112111161123333333333 ``` \| SeedType = 3 NonSeedType = -1 \| \| --- \| --- \| --- \| --- \| --- \| --- \| | \| 362 \| WBGene00005057:sra-31(3260) \| ``` log2FC=0.1 padj=1.0 ``` \| [seedLoc = 114,120] Seq=AGTTTTACAGTACACTCA \| ```              A             TTTGA mRNA  3' ACTC CAT     GACAT      miRNA 5' TGAG GTA     TTGTA                       GTAGG     TAGTT           11115111666662111133333 ``` \| SeedType = 2 NonSeedType = -1 \| \| --- \| --- \| --- \| --- \| --- \| --- \| | \| 363 \| WBGene00005085:srd-7(3266) \| ``` log2FC=0.746 padj=1.0 ``` \| [seedLoc = 44,50] Seq=TGACAAGGCGATTTGAAGTTCATACCATCA \| ```             A    A TTGAA        GAA   T mRNA  3' ACT CCAT C     GTTTAGCG   CAG  miRNA 5' TGA GGTA G     TAGGTTGT   GTT                                  ATA               111511115155455212212123331125 ``` \| SeedType = 3 NonSeedType = 2 \| \| --- \| --- \| --- \| --- \| --- \| --- \| | \| 364 \| WBGene00005444:srh-237(3294) \| ``` log2FC=-0.336 padj=0.842 ``` \| [seedLoc = 44,49] Seq=AGAGGTTTTACCGGTTAATTCTACTTCT \| ```          T       TTAATTGG   TTTTGGAGA mRNA  3'  CTTCATC        CCA          miRNA 5'  GAGGTAG        GGT                   T       TA         TGTATAGTT           3112111133555544111333333333 ``` \| SeedType = 3 NonSeedType = 1 \| \| --- \| --- \| --- \| --- \| --- \| --- \| | \| 365 \| WBGene00005712:srv-1(3322) \| ``` log2FC=0.366 padj=0.703 ``` \| [seedLoc = 71,76] Seq=TAATTATCCAACTGCCTCT \| ```          T        A            T mRNA  3'  CTCCGTCA CC    TATTAA  miRNA 5'  GAGGTAGT GG    ATAGTT           T        A  TTGT                  31111211131166661112115 ``` \| SeedType = 2 NonSeedType = -1 \| \| --- \| --- \| --- \| --- \| --- \| --- \| | \| 366 \| WBGene00006062:stn-1(3354) \| ``` log2FC=0.299 padj=0.715 ``` \| [seedLoc = 72,77] Seq=CAGCAATACCACATTCCGTCTCCTGCCTCT \| ```          T       C  TGCCTTA  C    A   C mRNA  3'  CTCCGTC TC       CA CATA CGA  miRNA 5'  GAGGTAG AG       GT GTAT GTT           T       T           T    A               311112113115455555113111131215 ``` \| SeedType = 2 NonSeedType = -1 \| \| --- \| --- \| --- \| --- \| --- \| --- \| | \| 367 \| WBGene00006063:sto-1(3355) \| ``` log2FC=-0.537 padj=0.362 ``` \| [seedLoc = 167,171] Seq=TTTATACTCGATTTCCACCTCAT \| ```          T      C       CT      T  mRNA  3'  ACTCCA C TTTAG  CATATT   miRNA 5'  TGAGGT G AGGTT  GTATAG                   A T             TT           5111111316122125511111236 ``` \| SeedType = 2 NonSeedType = 2 \| \| --- \| --- \| --- \| --- \| --- \| --- \| | \| 368 \| WBGene00006066:sto-4(3357) \| ``` log2FC=-0.096 padj=0.982 ``` \| [seedLoc = 122,127] Seq=TAAATATTAACCAGAAAAATTGACTACCTTG \| ```          G        GTTAAAAAGA    T    AAT  mRNA  3'  TTCCATCA          CCAA TATA     miRNA 5'  GAGGTAGT          GGTT GTAT              T        A                  AGTT           32111111135555555451111521113336 ``` \| SeedType = 3 NonSeedType = 1 \| \| --- \| --- \| --- \| --- \| --- \| --- \| | \| 369 \| WBGene00006067:sto-5(3358) \| ``` log2FC=0.298 padj=0.87 ``` \| [seedLoc = 43,48] Seq=ATCAGTTGGACCGGTAATTATGCTACCTTG \| ```          G         ATTAATGG    GTTGACTA mRNA  3'  TTCCATCGT        CCAG         miRNA 5'  GAGGTAGTA        GGTT                  T                     GTATAGTT           321111112155555544111233333333 ``` \| SeedType = 3 NonSeedType = 1 \| \| --- \| --- \| --- \| --- \| --- \| --- \| | \| 370 \| WBGene00006068:sto-6(3359) \| ``` log2FC=-0.66 padj=1.0 ``` \| [seedLoc = 76,81] Seq=GAATGTGTCAATCTTTCTACTTCC \| ```          C       TT      T    AAG mRNA  3'  CTTCATC  TCTAAC GTGT    miRNA 5'  GAGGTAG  AGGTTG TATA             T       T            GTT           311211113511211152121333 ``` \| SeedType = 3 NonSeedType = 1 \| \| --- \| --- \| --- \| --- \| --- \| --- \| | \| 371 \| WBGene00006127:str-64(3362) \| ``` log2FC=0.1 padj=1.0 ``` \| [seedLoc = 281,286] Seq=CGCTTCCAAATAACTACCTTT \| ```          T        A    ACCT   C mRNA  3'  TTCCATCA  TAA    TCG  miRNA 5'  GAGGTAGT  GTT    AGT           T        AG   GTAT   T           3211111113621133331123 ``` \| SeedType = 3 NonSeedType = -1 \| \| --- \| --- \| --- \| --- \| --- \| --- \| | \|  \|  \|  \| [seedLoc = 342,347] Seq=TTCCACCTTTTCTGTTACCTCC \| ```          C           TTTCCACCTT mRNA  3'  CTCCATTGTCT           miRNA 5'  GAGGTAGTAGG                    T           TTGTATAGTT           3111111221123333333333 ``` \| SeedType = 1 NonSeedType = -1 \| \| --- \| --- \| --- \| --- \| --- \| --- \| | \| 372 \| WBGene00006324:sup-17(3390) \| ``` log2FC=0.32 padj=0.344 ``` \| [seedLoc = 168,173] Seq=TGCCTTTTCCACTCCTACTTCT \| ```          T       C  ACCTTTTCCGT mRNA  3'  CTTCATC TC            miRNA 5'  GAGGTAG AG                     T       T  GTTGTATAGTT           3112111131133333333333 ``` \| SeedType = 3 NonSeedType = -1 \| \| --- \| --- \| --- \| --- \| --- \| --- \| | \| 373 \| WBGene00006331:sup-26(3391) \| ``` log2FC=-0.434 padj=0.071 ``` \| [seedLoc = 271,276] Seq=CCTCCCGAAATCTCTATACCTCT \| ```          T      ATC     AGCCCTCC mRNA  3'  CTCCAT   TCTAA         miRNA 5'  GAGGTA   AGGTT                  T      GT      GTATAGTT           31111113351121133333333 ``` \| SeedType = 1 NonSeedType = 2 \| \| --- \| --- \| --- \| --- \| --- \| --- \| | \|  \|  \|  \| [seedLoc = 472,478] Seq=AAGTTAACAGCACATACACTCT \| ```          T   A   ACA      ATTGAA mRNA  3'  CTC CAT    CGACA       miRNA 5'  GAG GTA    GTTGT                T       GTAG     ATAGTT           31115111333612111333333 ``` \| SeedType = 2 NonSeedType = -1 \| \| --- \| --- \| --- \| --- \| --- \| --- \| | \|  \|  \|  \| [seedLoc = 505,510] Seq=TCTCTCTCTCCCTCTCTATCTCT \| ```          T       TC   CTCTCTCTCT mRNA  3'  CTCTATC  TCC           miRNA 5'  GAGGTAG  AGG                    T       T    TTGTATAGTT           31112111351113333333333 ``` \| SeedType = 2 NonSeedType = -1 \| \| --- \| --- \| --- \| --- \| --- \| --- \| | \| 374 \| WBGene00006342:sup-37(3394) \| ``` log2FC=0.3 padj=0.372 ``` \| [seedLoc = 343,348] Seq=TTGCAATAATCATCTTTTTCGGATTACCTCA \| ```                   GGCTTTTTCTACTAA      T     mRNA  3' ACTCCATTA               TAACGT      miRNA 5' TGAGGTAGT               GTTGTA                        AG                   TAGTT           11111112133555555555555521112136666 ``` \| SeedType = 1 NonSeedType = 1 \| \| --- \| --- \| --- \| --- \| --- \| --- \| | \| 375 \| WBGene00006352:sur-6(3398) \| ``` log2FC=-0.024 padj=0.988 ``` \| [seedLoc = 397,403] Seq=TTCTTTACTCTCTCCCTCT \| ```          T    C           T  TT mRNA  3'  CTCC TC TCT  CAT TC   miRNA 5'  GAGG AG AGG  GTA AG            T    T  T   TT   T  TT           3111131161126611131133 ``` \| SeedType = 2 NonSeedType = -1 \| \| --- \| --- \| --- \| --- \| --- \| --- \| | \| 376 \| WBGene00006359:swp-1(3401) \| ``` log2FC=-0.094 padj=0.954 ``` \| [seedLoc = 543,548] Seq=ACGTCAAGAGCTTCTACTTCA \| ```                        G ACTGCA mRNA  3' ACTTCATC TTCGA A       miRNA 5' TGAGGTAG AGGTT T                        T     G ATAGTT           1112111161212131333333 ``` \| SeedType = 3 NonSeedType = 2 \| \| --- \| --- \| --- \| --- \| --- \| --- \| | \| 377 \| WBGene00006363:syd-1(3404) \| ``` log2FC=0.238 padj=0.966 ``` \| [seedLoc = 198,204] Seq=ATTCAAATAATGTTTACCATCA \| ```             A       G     AACTTA mRNA  3' ACT CCATT T TAATA       miRNA 5' TGA GGTAG A GTTGT                         T G     ATAGTT           11151111261321121333333 ``` \| SeedType = 3 NonSeedType = -1 \| \| --- \| --- \| --- \| --- \| --- \| --- \| | \| 378 \| WBGene00006379:sys-1(3419) \| ``` log2FC=0.052 padj=0.988 ``` \| [seedLoc = 939,944] Seq=TCCAATGCAATTTTCCTACTTCA \| ```                  CT         ACCT mRNA  3' ACTTCATC  TTTAACGTA     miRNA 5' TGAGGTAG  AGGTTGTAT                      T          AGTT           11121111351221112113333 ``` \| SeedType = 3 NonSeedType = 1 \| \| --- \| --- \| --- \| --- \| --- \| --- \| | \|  \|  \|  \| [seedLoc = 953,957] Seq=TGCAATTTTCCTACTTCAC \| ```          C        CT              mRNA  3'  ACTTCATC  TTTAACGT      miRNA 5'  TGAGGTAG  AGGTTGTA                        T         TAGTT           511121111351221112166666 ``` \| SeedType = 3 NonSeedType = 1 \| \| --- \| --- \| --- \| --- \| --- \| --- \| | \| 379 \| WBGene00006384:taf-3(3424) \| ``` log2FC=-0.151 padj=0.914 ``` \| [seedLoc = 346,351] Seq=CCATCCCCATCAAACTTACCTCT \| ```          T     T   AA T  CCCTACC mRNA  3'  CTCCA TCA  C AC        miRNA 5'  GAGGT AGT  G TG                 T         AG T  TATAGTT           31111151113313113333333 ``` \| SeedType = 2 NonSeedType = -1 \| \| --- \| --- \| --- \| --- \| --- \| --- \| | \| 380 \| WBGene00006414:raga-1(3449) \| ``` log2FC=-0.091 padj=0.918 ``` \| [seedLoc = 57,63] Seq=TTATTTTCGTTTAGTTACCATCC \| ```          C  A     GATTTGCTTTTATT mRNA  3'  CT CCATT               miRNA 5'  GA GGTAG                        T        TAGGTTGTATAGTT           31151111233333333333333 ``` \| SeedType = 3 NonSeedType = -1 \| \| --- \| --- \| --- \| --- \| --- \| --- \| | \|  \|  \|  \| [seedLoc = 231,237] Seq=GTTTAATTCTCCCCGTACCATCC \| ```          C  A    GCC  TCTTAATTTG mRNA  3'  CT CCAT   CC           miRNA 5'  GA GGTA   GG                    T       GTA  TTGTATAGTT           31151111333113333333333 ``` \| SeedType = 3 NonSeedType = -1 \| \| --- \| --- \| --- \| --- \| --- \| --- \| | \| 381 \| WBGene00006418:spl-2(3452) \|  \| [seedLoc = 26,31] Seq=AAACTATAATTATATTTGCCTACCTCT \| ```          T      C      TATTA       A mRNA  3'  CTCCAT CGTTTA     ATATCAA  miRNA 5'  GAGGTA GTAGGT     TATAGTT           T             TG                      311111151212213355511111115 ``` \| SeedType = 1 NonSeedType = 2 \| \| --- \| --- \| --- \| --- \| --- \| --- \| | \| 382 \| WBGene00006423:asd-2(3454) \| ``` log2FC=0.123 padj=0.866 ``` \| [seedLoc = 103,108] Seq=TCAGTTATCTCTTCTCTATCTCT \| ```          T       TC   TC   TGACT mRNA  3'  CTCTATC  TTC  TAT      miRNA 5'  GAGGTAG  AGG  GTA               T       T    TT   TAGTT           31112111351213321133333 ``` \| SeedType = 2 NonSeedType = -1 \| \| --- \| --- \| --- \| --- \| --- \| --- \| | \| 383 \| WBGene00006424:ctbp-1(3457) \| ``` log2FC=0.547 padj=0.514 ``` \| [seedLoc = 404,409] Seq=TCTCGCGCATTTCTAAGATACCTCA \| ```                 AGA    TT   CGCTCT mRNA  3' ACTCCAT   ATCT  ACG       miRNA 5' TGAGGTA   TAGG  TGT                       G      T    ATAGTT           1111111345111235112333333 ``` \| SeedType = 1 NonSeedType = -1 \| \| --- \| --- \| --- \| --- \| --- \| --- \| | \|  \|  \|  \| [seedLoc = 418,421] Seq=GCATTTCTAAGATACCTCAT \| ```          T       AGA    TT          mRNA  3'  ACTCCAT   ATCT  ACG       miRNA 5'  TGAGGTA   TAGG  TGT                        G      T    ATAGTT           51111111345111235112666666 ``` \| SeedType = 1 NonSeedType = -1 \| \| --- \| --- \| --- \| --- \| --- \| --- \| | \| 384 \| WBGene00006437:mrck-1(3467) \| ``` log2FC=0.042 padj=0.975 ``` \| [seedLoc = 80,85] Seq=AAACGATGTAAATCTGGTGCTATCTCC \| ```          C        TGG     AT   G   A mRNA  3'  CTCTATCG   TCTAA  GTA CAA  miRNA 5'  GAGGTAGT   AGGTT  TAT GTT           T                G    A               311121112544112113521131115 ``` \| SeedType = 2 NonSeedType = 2 \| \| --- \| --- \| --- \| --- \| --- \| --- \| | \| 385 \| WBGene00006462:svh-5(3489) \|  \| [seedLoc = 411,416] Seq=CTCTGAATTACCATAATTACCTCA \| ```                   ATA   T  A   TC mRNA  3' ACTCCATTA   CCA TA GTC   miRNA 5' TGAGGTAGT   GGT GT TAG                     A     T  A   TT           111111121355111321321133 ``` \| SeedType = 1 NonSeedType = 1 \| \| --- \| --- \| --- \| --- \| --- \| --- \| | \| 386 \| WBGene00006470:tag-120(3495) \| ``` log2FC=-0.139 padj=0.861 ``` \| [seedLoc = 1011,1017] Seq=CCACCGAGAGCATATACCATCA \| ```             A    ATA    G GCCACC mRNA  3' ACT CCAT    CGA A       miRNA 5' TGA GGTA    GTT T                        GTAG   G ATAGTT           11151111333612131333333 ``` \| SeedType = 3 NonSeedType = -1 \| \| --- \| --- \| --- \| --- \| --- \| --- \| | \| 387 \| WBGene00006492:let-391(3513) \|  \| [seedLoc = 67,72] Seq=TAACTAGCATTTTCTACTTCA \| ```                      T         T mRNA  3' ACTTCATC TTT ACG ATCAA  miRNA 5' TGAGGTAG AGG TGT TAGTT                   T   T   A                 11121111612231126111115 ``` \| SeedType = 3 NonSeedType = -1 \| \| --- \| --- \| --- \| --- \| --- \| --- \| | \| 388 \| WBGene00006504:kcc-1(3526) \| ``` log2FC=0.111 padj=0.902 ``` \| [seedLoc = 322,328] Seq=TAAGGTATTCCTTCTTACCATCA \| ```             A     CT   T     GAAT mRNA  3' ACT CCATT  TCC  TATG     miRNA 5' TGA GGTAG  AGG  GTAT                       T    TT    AGTT           111511112351113621123333 ``` \| SeedType = 3 NonSeedType = -1 \| \| --- \| --- \| --- \| --- \| --- \| --- \| | \|  \|  \|  \| [seedLoc = 331,335] Seq=TAAGGTATTCCTTCTTACCATCAC \| ```          C   A     CT   T     GAAT mRNA  3'  ACT CCATT  TCC  TATG     miRNA 5'  TGA GGTAG  AGG  GTAT                        T    TT    AGTT           5111511112351113621123333 ``` \| SeedType = 3 NonSeedType = -1 \| \| --- \| --- \| --- \| --- \| --- \| --- \| | \|  \|  \|  \| [seedLoc = 401,407] Seq=TTTTTGGCGCCCTCCGTACTCTCG \| ```          G   T   G C   C   GTTTTT mRNA  3'  CTC CAT C TCC GCG       miRNA 5'  GAG GTA G AGG TGT                T         T   T   ATAGTT           311151114131113212333333 ``` \| SeedType = 2 NonSeedType = -1 \| \| --- \| --- \| --- \| --- \| --- \| --- \| | \| 389 \| WBGene00006515:txdc-9(3534) \|  \| [seedLoc = 122,127] Seq=TGCTGTTTTATCTACCTTT \| ```          T       T             T mRNA  3'  TTCCATC ATTT    TGTCG  miRNA 5'  GAGGTAG TAGG    ATAGT           T            TTGT     T           32111111511226666121123 ``` \| SeedType = 3 NonSeedType = -1 \| \| --- \| --- \| --- \| --- \| --- \| --- \| | \| 390 \| WBGene00006563:teg-1(3570) \| ``` log2FC=-0.052 padj=0.973 ``` \| [seedLoc = 135,140] Seq=CTCGTTTTCACTCTCTATCTCT \| ```          T       TC   CTTTTGCTC mRNA  3'  CTCTATC  TCA          miRNA 5'  GAGGTAG  GGT                   T       TA   TGTATAGTT           3111211133211333333333 ``` \| SeedType = 2 NonSeedType = 2 \| \| --- \| --- \| --- \| --- \| --- \| --- \| | \| 391 \| WBGene00006577:tlf-1(3581) \| ``` log2FC=-0.009 padj=0.995 ``` \| [seedLoc = 87,91] Seq=ATCTGATTACCTTTCTACATCAT \| ```          T   A    TT    T      TA mRNA  3'  ACT CATC  TCCA TA GTC   miRNA 5'  TGA GTAG  AGGT GT TAG                G    T     T  A   TT           511131111351111321621133 ``` \| SeedType = 3 NonSeedType = 1 \| \| --- \| --- \| --- \| --- \| --- \| --- \| | \|  \|  \|  \| [seedLoc = 217,223] Seq=CTCTGTTTGCCGTTACCATCA \| ```             A     G    TT     TC mRNA  3' ACT CCATT  CCG   TGTC   miRNA 5' TGA GGTAG  GGT   ATAG                     TA   TGT    TT           11151111236112336121133 ``` \| SeedType = 3 NonSeedType = 2 \| \| --- \| --- \| --- \| --- \| --- \| --- \| | \|  \|  \|  \| [seedLoc = 357,362] Seq=CACTATGTGCTCTCTACTTCA \| ```                  TC   T       C mRNA  3' ACTTCATC  TCG  GTATCA  miRNA 5' TGAGGTAG  GGT  TATAGT                   TA   TG      T           1112111133212362111113 ``` \| SeedType = 3 NonSeedType = 2 \| \| --- \| --- \| --- \| --- \| --- \| --- \| | \| 392 \| WBGene00006588:tnt-3(3594) \| ``` log2FC=-0.193 padj=0.676 ``` \| [seedLoc = 390,396] Seq=CGATAAACAAAGATACCATCA \| ```             A    AGA      AATAGC mRNA  3' ACT CCAT     AACA       miRNA 5' TGA GGTA     TTGT                        GTAGG    ATAGTT           11151111333661111333333 ``` \| SeedType = 3 NonSeedType = 1 \| \| --- \| --- \| --- \| --- \| --- \| --- \| | \| 393 \| WBGene00006590:toc-1(3598) \| ``` log2FC=0.445 padj=0.064 ``` \| [seedLoc = 155,160] Seq=CCCAGTACATTTGTTTACCTCT \| ```          T       TG  T     ACCC mRNA  3'  CTCCATT  TT ACATG     miRNA 5'  GAGGTAG  GG TGTAT              T       TA  T     AGTT           3111111233223111123333 ``` \| SeedType = 1 NonSeedType = -1 \| \| --- \| --- \| --- \| --- \| --- \| --- \| | \| 394 \| WBGene00006591:toh-1(3599) \| ``` log2FC=-0.333 padj=0.4 ``` \| [seedLoc = 188,193] Seq=TTCTGTATATTTCTGCCTCT \| ```          T                   TT mRNA  3'  CTCCGTC TTT ATATGTC   miRNA 5'  GAGGTAG AGG TGTATAG            T       T   T       TT           3111121161226121121133 ``` \| SeedType = 2 NonSeedType = -1 \| \| --- \| --- \| --- \| --- \| --- \| --- \| | \| 395 \| WBGene00006602:tps-1(3610) \| ``` log2FC=0.3 padj=0.38 ``` \| [seedLoc = 97,103] Seq=ATCTAAATTCCAGTATTAACCTCT \| ```          T    A     GA  T   A   TA mRNA  3'  CTCC ATTAT  CC  TA ATC   miRNA 5'  GAGG TAGTA  GG  GT TAG            T              TT  A   TT           3111151121145113621311133 ``` \| SeedType = 2 NonSeedType = -1 \| \| --- \| --- \| --- \| --- \| --- \| --- \| | \| 396 \| WBGene00006604:tra-1(3615) \| ``` log2FC=-0.049 padj=0.973 ``` \| [seedLoc = 174,179] Seq=AAATTCACATTTTTCTACCTCT \| ```          T       T   T   CTTAAA mRNA  3'  CTCCATC TTT ACA       miRNA 5'  GAGGTAG AGG TGT                T       T   T   ATAGTT           3111111131223111333333 ``` \| SeedType = 1 NonSeedType = -1 \| \| --- \| --- \| --- \| --- \| --- \| --- \| | \|  \|  \|  \| [seedLoc = 205,210] Seq=CCTTTTTTTTTGTTTTACCTCG \| ```          G       TTGTTTTTTTTTCC mRNA  3'  CTCCATT               miRNA 5'  GAGGTAG                        T       TAGGTTGTATAGTT           3111111233333333333333 ``` \| SeedType = 1 NonSeedType = -1 \| \| --- \| --- \| --- \| --- \| --- \| --- \| | \| 397 \| WBGene00006617:tars-1(3627) \| ``` log2FC=0.095 padj=0.912 ``` \| [seedLoc = 109,115] Seq=TTTTGTGTTCCCTGATTGTTATCCTCT \| ```          T    T   GTTA    CTT     TT mRNA  3'  CTCC ATT    GTCC   GTGTT   miRNA 5'  GAGG TAG    TAGG   TATAG            T                TTG     TT           311115112455521113332121233 ``` \| SeedType = 2 NonSeedType = -1 \| \| --- \| --- \| --- \| --- \| --- \| --- \| | \| 398 \| WBGene00006619:try-1(3629) \| ``` log2FC=0.613 padj=0.543 ``` \| [seedLoc = 97,103] Seq=GTGAATGAAATGTTTACCGTCT \| ```          T  G       G   A   AGTG mRNA  3'  CT CCATT T TAA GTA     miRNA 5'  GA GGTAG A GTT TAT              T        T G   G   AGTT           31141111261321132113333 ``` \| SeedType = 3 NonSeedType = -1 \| \| --- \| --- \| --- \| --- \| --- \| --- \| | \| 399 \| WBGene00006626:tsn-1(3633) \| ``` log2FC=0.217 padj=0.515 ``` \| [seedLoc = 64,70] Seq=CCTCCCCCAACCAGAAAAGTTTTATCCTCA \| ```               T   TTGAAAAGA     CCCCTCC mRNA  3' ACTCC ATT         CCAAC        miRNA 5' TGAGG TAG         GGTTG                          TA            TATAGTT           111115112334555545111113333333 ``` \| SeedType = 2 NonSeedType = 1 \| \| --- \| --- \| --- \| --- \| --- \| --- \| | \| 400 \| WBGene00006635:tsp-9(3640) \| ``` log2FC=0.305 padj=0.526 ``` \| [seedLoc = 198,203] Seq=TCTCATGCTAACTCTGATTTGATCTACCTCG \| ```          G       TAGTTTA   T   T    CTCT mRNA  3'  CTCCATC       GTC CAA CGTA     miRNA 5'  GAGGTAG       TAG GTT GTAT              T                          AGTT           3111111155455552115111512113333 ``` \| SeedType = 1 NonSeedType = 1 \| \| --- \| --- \| --- \| --- \| --- \| --- \| | \| 401 \| WBGene00006638:tsp-12(3645) \| ``` log2FC=-0.181 padj=0.904 ``` \| [seedLoc = 36,41] Seq=CAACGAAATATTCTCTACCTTC \| ```          C           T   AAG   C mRNA  3'  TTCCATC TCT ATA   CAA  miRNA 5'  GAGGTAG AGG TGT   GTT           T       T   T   ATA               32111111611231213331115 ``` \| SeedType = 3 NonSeedType = -1 \| \| --- \| --- \| --- \| --- \| --- \| --- \| | \| 402 \| WBGene00006647:tsr-1(3651) \| ``` log2FC=0.029 padj=0.986 ``` \| [seedLoc = 108,113] Seq=GTTTCATCGATTTTTACCTCG \| ```          G              TACTTTG mRNA  3'  CTCCATT TTTAGC        miRNA 5'  GAGGTAG AGGTTG                 T       T      TATAGTT           3111111261221213333333 ``` \| SeedType = 1 NonSeedType = 2 \| \| --- \| --- \| --- \| --- \| --- \| --- \| | \| 403 \| WBGene00006668:twk-13(3667) \| ``` log2FC=-0.243 padj=0.945 ``` \| [seedLoc = 179,185] Seq=TCAGATTCCTCTTACATATCCTCG \| ```          G    T  A     TCCTTAGACT mRNA  3'  CTCC AT CATTC           miRNA 5'  GAGG TA GTAGG                    T             TTGTATAGTT           311115115111213333333333 ``` \| SeedType = 2 NonSeedType = -1 \| \| --- \| --- \| --- \| --- \| --- \| --- \| | \| 404 \| WBGene00006679:twk-26(3676) \| ``` log2FC=-0.137 padj=0.943 ``` \| [seedLoc = 54,59] Seq=TTTTGAAAAATCAATCTGCCTCA \| ```                  TAA    AAAGTTTT mRNA  3' ACTCCGTC   CTAA         miRNA 5' TGAGGTAG   GGTT                          TA     GTATAGTT           11111211335121133333333 ``` \| SeedType = 2 NonSeedType = 2 \| \| --- \| --- \| --- \| --- \| --- \| --- \| | \| 405 \| WBGene00006683:twk-31(3678) \| ``` log2FC=-0.173 padj=0.945 ``` \| [seedLoc = 142,148] Seq=AATCATCCGGCAATACTCTCT \| ```          T   T   AA      CTACTAA mRNA  3'  CTC CAT    CGGC        miRNA 5'  GAG GTA    GTTG                 T       GTAG    TATAGTT           31115111336612213333333 ``` \| SeedType = 2 NonSeedType = -1 \| \| --- \| --- \| --- \| --- \| --- \| --- \| | \|  \|  \|  \| [seedLoc = 350,356] Seq=ATTCCCCCATCCTTTCTCACCTCA \| ```                C  TT   T  CCCCTTA mRNA  3' ACTCCA TC  TCC AC        miRNA 5' TGAGGT AG  AGG TG                          T    T  TATAGTT           111111511351113113333333 ``` \| SeedType = 2 NonSeedType = -1 \| \| --- \| --- \| --- \| --- \| --- \| --- \| | \| 406 \| WBGene00006698:uaf-2(3687) \| ``` log2FC=-0.062 padj=0.959 ``` \| [seedLoc = 80,85] Seq=TGCTCTTTTTTCTCTATCTCT \| ```          T           TTTTTC   T mRNA  3'  CTCTATC TCT      TCG  miRNA 5'  GAGGTAG AGG      AGT           T       T   TTGTAT   T           3111211161123333331123 ``` \| SeedType = 2 NonSeedType = -1 \| \| --- \| --- \| --- \| --- \| --- \| --- \| | \| 407 \| WBGene00006699:uba-1(3688) \| ``` log2FC=0.17 padj=0.616 ``` \| [seedLoc = 181,186] Seq=TTCACGTTCTCCCCGTACCTCA \| ```                 GCC  TCTTGCACTT mRNA  3' ACTCCAT   CC           miRNA 5' TGAGGTA   GG                           GTA  TTGTATAGTT           1111111333113333333333 ``` \| SeedType = 1 NonSeedType = -1 \| \| --- \| --- \| --- \| --- \| --- \| --- \| | \|  \|  \|  \| [seedLoc = 190,194] Seq=TTCACGTTCTCCCCGTACCTCAT \| ```          T       GCC  TCTTGCACTT mRNA  3'  ACTCCAT   CC           miRNA 5'  TGAGGTA   GG                            GTA  TTGTATAGTT           51111111333113333333333 ``` \| SeedType = 1 NonSeedType = -1 \| \| --- \| --- \| --- \| --- \| --- \| --- \| | \| 408 \| WBGene00006704:ubc-7(3693) \| ``` log2FC=-0.058 padj=0.956 ``` \| [seedLoc = 345,350] Seq=GAATTCACAATCCTCGTGATGCTACTTCT \| ```          T         AGTGCTC      CTTAAG mRNA  3'  CTTCATCGT       CTAACA       miRNA 5'  GAGGTAGTA       GGTTGT                T                      ATAGTT           31121111215454555121111333333 ``` \| SeedType = 3 NonSeedType = 1 \| \| --- \| --- \| --- \| --- \| --- \| --- \| | \| 409 \| WBGene00006720:ubc-25(3708) \| ``` log2FC=-0.313 padj=0.222 ``` \| [seedLoc = 276,281] Seq=CCCCTTCCAATTAATACCTCC \| ```          C      AA      CTTCCCC mRNA  3'  CTCCAT   TTAAC        miRNA 5'  GAGGTA   GGTTG                 T      GTA     TATAGTT           3111111336221113333333 ``` \| SeedType = 1 NonSeedType = 1 \| \| --- \| --- \| --- \| --- \| --- \| --- \| | \| 410 \| WBGene00006725:ubl-1(3714) \| ``` log2FC=-0.087 padj=0.943 ``` \| [seedLoc = 38,42] Seq=CTCTGAGGCCACCATCTACGTCAA \| ```          A   G    TA   C  GA   TC mRNA  3'  ACT CATC  CCA CG  GTC   miRNA 5'  TGA GTAG  GGT GT  TAG                G    TA   T  A    TT           511131111331113123521133 ``` \| SeedType = 3 NonSeedType = 1 \| \| --- \| --- \| --- \| --- \| --- \| --- \| | \| 411 \| WBGene00006730:uev-1(3720) \| ``` log2FC=-0.058 padj=0.959 ``` \| [seedLoc = 31,37] Seq=TTTTGTATACCCAGTTTACTCTCC \| ```          C   T    TGA  C       TT mRNA  3'  CTC CATT   CC ATATGTT   miRNA 5'  GAG GTAG   GG TGTATAG            T        TA   T       TT           311151112335113121121233 ``` \| SeedType = 2 NonSeedType = -1 \| \| --- \| --- \| --- \| --- \| --- \| --- \| | \| 412 \| WBGene00006737:ulp-2(3726) \| ``` log2FC=0.004 padj=0.997 ``` \| [seedLoc = 84,90] Seq=TTTTAAAGATTTTATTATCCTCC \| ```          C    T       TAGAAATTTT mRNA  3'  CTCC ATTATTT           miRNA 5'  GAGG TAGTAGG                    T            TTGTATAGTT           31111511211223333333333 ``` \| SeedType = 2 NonSeedType = -1 \| \| --- \| --- \| --- \| --- \| --- \| --- \| | \| 413 \| WBGene00006739:ulp-4(3727) \| ``` log2FC=-0.15 padj=0.822 ``` \| [seedLoc = 47,53] Seq=TCATATGCCCCGGTTCTTTACCGTCC \| ```          C  G     TCTTGG  C      ACT mRNA  3'  CT CCATT      CC  CGTAT    miRNA 5'  GA GGTAG      GG  GTATA             T        TA      TT     GTT           311411112335544113612111333 ``` \| SeedType = 3 NonSeedType = -1 \| \| --- \| --- \| --- \| --- \| --- \| --- \| | \|  \|  \|  \| [seedLoc = 1204,1210] Seq=TCGTTTTGAGCCAGCTACCATCT \| ```          T  A      A    GTTTTGCT mRNA  3'  CT CCATCG CCGA         miRNA 5'  GA GGTAGT GGTT                  T         A    GTATAGTT           31151111123112133333333 ``` \| SeedType = 3 NonSeedType = 2 \| \| --- \| --- \| --- \| --- \| --- \| --- \| | \| 414 \| WBGene00006742:unc-2(3730) \| ``` log2FC=0.038 padj=0.994 ``` \| [seedLoc = 132,137] Seq=CATCCCCCTTTCTGTCTATCTCA \| ```                  T    TTCCCCCTAC mRNA  3' ACTCTATC GTCT           miRNA 5' TGAGGTAG TAGG                                 TTGTATAGTT           11112111521123333333333 ``` \| SeedType = 2 NonSeedType = -1 \| \| --- \| --- \| --- \| --- \| --- \| --- \| | \|  \|  \|  \| [seedLoc = 606,611] Seq=CATTATAAACAAAATTCTACCTTA \| ```                  TTAA    AA     C mRNA  3' ATTCCATC    AACA  TATTA  miRNA 5' TGAGGTAG    TTGT  ATAGT                   TAGG           T           121111113333111155111213 ``` \| SeedType = 3 NonSeedType = 1 \| \| --- \| --- \| --- \| --- \| --- \| --- \| | \| 415 \| WBGene00006744:unc-4(3733) \| ``` log2FC=1.891 padj=1.0 ``` \| [seedLoc = 47,52] Seq=CATTGGTTATTTGTACCTCC \| ```          C            TTG     C mRNA  3'  CTCCAT GTTTA    GTTA  miRNA 5'  GAGGTA TAGGT    TAGT           T      G     TGTA    T           3111111621221333621213 ``` \| SeedType = 1 NonSeedType = 2 \| \| --- \| --- \| --- \| --- \| --- \| --- \| | \| 416 \| WBGene00006746:unc-6(3737) \| ``` log2FC=0.051 padj=0.988 ``` \| [seedLoc = 217,222] Seq=TACTGTAGCAGTACCTCT \| ```          T      GA            T mRNA  3'  CTCCAT    CG  ATGTCA  miRNA 5'  GAGGTA    GT  TATAGT           T      GTAG  TG      T           3111111336612661121113 ``` \| SeedType = 1 NonSeedType = -1 \| \| --- \| --- \| --- \| --- \| --- \| --- \| | \| 417 \| WBGene00006747:unc-7(3738) \| ``` log2FC=0.102 padj=0.979 ``` \| [seedLoc = 413,418] Seq=TATGTTGTGACTTGCTACCTTA \| ```                           TG  T mRNA  3' ATTCCATCGTTCAGTGT  TA  miRNA 5' TGAGGTAGTAGGTTGTA  GT                            TA  T           1211111121211222133213 ``` \| SeedType = 3 NonSeedType = 2 \| \| --- \| --- \| --- \| --- \| --- \| --- \| | \| 418 \| WBGene00006751:unc-11(3743) \| ``` log2FC=0.017 padj=0.993 ``` \| [seedLoc = 499,504] Seq=GTGATGTTTTCCCCCTATCTCT \| ```          T       CC  TTTTGTAGTG mRNA  3'  CTCTATC  CC           miRNA 5'  GAGGTAG  GG                    T       TA  TTGTATAGTT           3111211133113333333333 ``` \| SeedType = 2 NonSeedType = -1 \| \| --- \| --- \| --- \| --- \| --- \| --- \| | \|  \|  \|  \| [seedLoc = 505,510] Seq=ATGTTTTCCCCCTATCTCTATCTCT \| ```          T       TCT    CCCTTTTGTA mRNA  3'  CTCTATC   ATCC           miRNA 5'  GAGGTAG   TAGG                    T              TTGTATAGTT           3111211155511113333333333 ``` \| SeedType = 2 NonSeedType = -1 \| \| --- \| --- \| --- \| --- \| --- \| --- \| | \| 419 \| WBGene00006754:unc-15(3749) \| ``` log2FC=0.138 padj=0.896 ``` \| [seedLoc = 465,471] Seq=GTCTTGACGAGATCGCTACCTGC \| ```          CG       CTAG    GT  TG mRNA  3'   TCCATCG    AGCA  TC   miRNA 5'   AGGTAGT    TTGT  AG            TG       AGG     AT  TT           33111111233341211331133 ``` \| SeedType = 3 NonSeedType = -1 \| \| --- \| --- \| --- \| --- \| --- \| --- \| | \|  \|  \|  \| [seedLoc = 553,558] Seq=TATCCCATTACTTTCCTACTTCT \| ```          T       CT    TTACCCTAT mRNA  3'  CTTCATC  TTCA          miRNA 5'  GAGGTAG  AGGT                   T       T     TGTATAGTT           31121111351211333333333 ``` \| SeedType = 3 NonSeedType = 2 \| \| --- \| --- \| --- \| --- \| --- \| --- \| | \| 420 \| WBGene00006755:unc-16(3751) \| ``` log2FC=0.019 padj=0.99 ``` \| [seedLoc = 238,241] Seq=CTAATATTCGGTTGCTCCCTCC \| ```          C    C     G CT    ATC  mRNA  3'  CTCC TCGTT G  TATA     miRNA 5'  GAGG AGTAG T  GTAT              T    T     G T     AGTT           31111311212323521113336 ``` \| SeedType = 2 NonSeedType = -1 \| \| --- \| --- \| --- \| --- \| --- \| --- \| | \| 421 \| WBGene00006757:unc-18(3753) \| ``` log2FC=0.136 padj=0.83 ``` \| [seedLoc = 289,293] Seq=TTATTTATTGATATTACCTCAC \| ```          C          AGT    TTATT mRNA  3'  ACTCCATTAT    TAT      miRNA 5'  TGAGGTAGTA    GTA                          GGTT   TAGTT           51111111211333621133333 ``` \| SeedType = 1 NonSeedType = -1 \| \| --- \| --- \| --- \| --- \| --- \| --- \| | \| 422 \| WBGene00006779:unc-43(3781) \| ``` log2FC=0.023 padj=0.988 ``` \| [seedLoc = 59,63] Seq=TCCTAACATCATCTACGTCAT \| ```          T   G    TA          CT mRNA  3'  ACT CATC  CT ACA ATC   miRNA 5'  TGA GTAG  GG TGT TAG                G    TA  T   A   TT           51113111133126111611133 ``` \| SeedType = 3 NonSeedType = -1 \| \| --- \| --- \| --- \| --- \| --- \| --- \| | \| 423 \| WBGene00006784:unc-49(3791) \| ``` log2FC=0.225 padj=0.864 ``` \| [seedLoc = 84,89] Seq=ACTATTTCTCGCCTTTCTCGTTGTTACCTCC \| ```          C       GTTGCTCTT    CTCTT       mRNA  3'  CTCCATT         TCCG     TATCA  miRNA 5'  GAGGTAG         AGGT     ATAGT           T       T            TGT       T           31111112355455555111233355111116 ``` \| SeedType = 1 NonSeedType = 2 \| \| --- \| --- \| --- \| --- \| --- \| --- \| | \| 424 \| WBGene00006791:unc-57(3801) \| ``` log2FC=-0.045 padj=0.975 ``` \| [seedLoc = 383,389] Seq=CCATGTATTCTATGTACTCTCT \| ```          T   T   G            ACC mRNA  3'  CTC CAT TATCT  TATGT    miRNA 5'  GAG GTA GTAGG  GTATA             T             TT     GTT           311151114211126621121333 ``` \| SeedType = 2 NonSeedType = -1 \| \| --- \| --- \| --- \| --- \| --- \| --- \| | \| 425 \| WBGene00006804:unc-71(3814) \| ``` log2FC=0.06 padj=0.985 ``` \| [seedLoc = 99,104] Seq=GTTTTTATTTTTTTTTACCTCA \| ```                  TTTTTTTATTTTTG mRNA  3' ACTCCATT               miRNA 5' TGAGGTAG                                TAGGTTGTATAGTT           1111111233333333333333 ``` \| SeedType = 1 NonSeedType = -1 \| \| --- \| --- \| --- \| --- \| --- \| --- \| | \| 426 \| WBGene00006807:unc-75(3818) \| ``` log2FC=0.197 padj=0.959 ``` \| [seedLoc = 304,309] Seq=CGGCGTTTTTTCCTACCGTACCTCT \| ```          T      GC     TTTTTTG   C mRNA  3'  CTCCAT  CATCC       CGG  miRNA 5'  GAGGTA  GTAGG       GTT           T             TTGTATA               3111111451111133333331225 ``` \| SeedType = 1 NonSeedType = -1 \| \| --- \| --- \| --- \| --- \| --- \| --- \| | \| 427 \| WBGene00006815:unc-83(3826) \| ``` log2FC=0.149 padj=0.854 ``` \| [seedLoc = 458,463] Seq=TTATTTACGAACTACCTTT \| ```          T                TTATT mRNA  3'  TTCCATCA   AGCAT      miRNA 5'  GAGGTAGT   TTGTA               T        AGG     TAGTT           3211111116661211133333 ``` \| SeedType = 3 NonSeedType = -1 \| \| --- \| --- \| --- \| --- \| --- \| --- \| | \| 428 \| WBGene00006833:unc-108(3845) \| ``` log2FC=-0.048 padj=0.962 ``` \| [seedLoc = 176,182] Seq=ATATTGCAATTGTCTCTTATTACTACCATCA \| ```             A        ATTCTCTGT      TATA  mRNA  3' ACT CCATCATT         TAACGT      miRNA 5' TGA GGTAGTAG         GTTGTA                                          TAGTT           11151111111255555554521112133336 ``` \| SeedType = 3 NonSeedType = 1 \| \| --- \| --- \| --- \| --- \| --- \| --- \| | \|  \|  \|  \| [seedLoc = 378,383] Seq=TTGTATGTAAGCTCTACTTCT \| ```          T          G       GTT mRNA  3'  CTTCATC TC AATGTAT    miRNA 5'  GAGGTAG AG TTGTATA             T       T  G       GTT           3112111161131122111333 ``` \| SeedType = 3 NonSeedType = -1 \| \| --- \| --- \| --- \| --- \| --- \| --- \| | \| 429 \| WBGene00006839:unc-115(3847) \| ``` log2FC=0.017 padj=0.993 ``` \| [seedLoc = 915,920] Seq=TAATTGTTTTTTGTTCCTACTTCA \| ```                  C  GTTTTT      T mRNA  3' ACTTCATC TT      TGTTAA  miRNA 5' TGAGGTAG AG      ATAGTT                   T  GTTGT                   111211113123333351212115 ``` \| SeedType = 3 NonSeedType = -1 \| \| --- \| --- \| --- \| --- \| --- \| --- \| | \| 430 \| WBGene00006840:unc-116(3848) \| ``` log2FC=0.053 padj=0.957 ``` \| [seedLoc = 79,84] Seq=AGACTGACATTCAAATGCTTCTCTATCTCT \| ```          T       TC    TAAACTT         A mRNA  3'  CTCTATC  TTCG       ACA GTCAG  miRNA 5'  GAGGTAG  AGGT       TGT TAGTT           T       T               A                 3111211135121255555551116211125 ``` \| SeedType = 2 NonSeedType = 2 \| \| --- \| --- \| --- \| --- \| --- \| --- \| | \| 431 \| WBGene00006852:unc-129(3851) \| ``` log2FC=-0.04 padj=0.989 ``` \| [seedLoc = 81,86] Seq=GTCTGTACATCTTCCCTACCTCT \| ```          T       CC   T       TG mRNA  3'  CTCCATC  TTC ACATGTC   miRNA 5'  GAGGTAG  AGG TGTATAG            T       T    T       TT           31111111351213111121133 ``` \| SeedType = 1 NonSeedType = -1 \| \| --- \| --- \| --- \| --- \| --- \| --- \| | \| 432 \| WBGene00006876:vab-10(3866) \| ``` log2FC=0.072 padj=0.961 ``` \| [seedLoc = 42,47] Seq=ATGTATACCCCGTACCCAGTTTTACCTCT \| ```          T      TTTGACC   G CC     GTA mRNA  3'  CTCCAT       CAT C  CATAT    miRNA 5'  GAGGTA       GTA G  GTATA             T                G TT     GTT           31111115554555111313311111333 ``` \| SeedType = 1 NonSeedType = -1 \| \| --- \| --- \| --- \| --- \| --- \| --- \| | \|  \|  \|  \| [seedLoc = 204,210] Seq=TATTATATATTTTTGTACCCTCA \| ```             C    GT   T        T mRNA  3' ACT CCAT  TTT ATATATTA  miRNA 5' TGA GGTA  AGG TGTATAGT                   GT   T        T           11151111331223121111213 ``` \| SeedType = 3 NonSeedType = -1 \| \| --- \| --- \| --- \| --- \| --- \| --- \| | \| 433 \| WBGene00006887:vav-1(3871) \| ``` log2FC=0.165 padj=0.816 ``` \| [seedLoc = 308,313] Seq=ACCTAATCAGCTCTACTTCA \| ```                         TA   CA mRNA  3' ACTTCATC  TCGAC  ATC   miRNA 5' TGAGGTAG  GGTTG  TAG                    TA     TA   TT           1112111166212113311133 ``` \| SeedType = 3 NonSeedType = 2 \| \| --- \| --- \| --- \| --- \| --- \| --- \| | \| 434 \| WBGene00006916:vha-7(3888) \| ``` log2FC=0.451 padj=0.08 ``` \| [seedLoc = 130,135] Seq=AGTATTTTTTAAATTTACCTCA \| ```                  TAAATTTTTTATGA mRNA  3' ACTCCATT               miRNA 5' TGAGGTAG                                TAGGTTGTATAGTT           1111111233333333333333 ``` \| SeedType = 1 NonSeedType = -1 \| \| --- \| --- \| --- \| --- \| --- \| --- \| | \| 435 \| WBGene00006923:vhp-1(3897) \| ``` log2FC=-0.202 padj=0.673 ``` \| [seedLoc = 124,130] Seq=TCTCTCCCAACATCATTACTCTCA \| ```              T     CTA    CCTCTCT mRNA  3' ACTC CATTA   CAAC        miRNA 5' TGAG GTAGT   GTTG                           AG     TATAGTT           111151112133511113333333 ``` \| SeedType = 2 NonSeedType = 1 \| \| --- \| --- \| --- \| --- \| --- \| --- \| | \| 436 \| WBGene00006931:vps-26(3908) \| ``` log2FC=0.254 padj=0.51 ``` \| [seedLoc = 76,81] Seq=TTAAATGCAAAACCTAACTATCTCC \| ```          C       A     AA     AATT mRNA  3'  CTCTATC ATCCA  ACGTA     miRNA 5'  GAGGTAG TAGGT  TGTAT              T                    AGTT           3111211151111155112113333 ``` \| SeedType = 2 NonSeedType = 1 \| \| --- \| --- \| --- \| --- \| --- \| --- \| | \| 437 \| WBGene00006933:vps-35(3910) \| ``` log2FC=0.126 padj=0.838 ``` \| [seedLoc = 59,64] Seq=TCCTCTTTCTTCTCTATCTCA \| ```                      TCTTTCTCCT mRNA  3' ACTCTATC TCT           miRNA 5' TGAGGTAG AGG                            T   TTGTATAGTT           1111211161123333333333 ``` \| SeedType = 2 NonSeedType = -1 \| \| --- \| --- \| --- \| --- \| --- \| --- \| | \| 438 \| WBGene00006941:wnk-1(3917) \| ``` log2FC=-0.162 padj=0.71 ``` \| [seedLoc = 642,648] Seq=GATGAAATAAAGATACTCTCA \| ```              T   AGA      AAGTAG mRNA  3' ACTC CAT     AATA       miRNA 5' TGAG GTA     TTGT                        GTAGG    ATAGTT           11115111333661121333333 ``` \| SeedType = 2 NonSeedType = -1 \| \| --- \| --- \| --- \| --- \| --- \| --- \| | \| 439 \| WBGene00006942:wrk-1(3919) \| ``` log2FC=0.115 padj=0.915 ``` \| [seedLoc = 38,43] Seq=CAAAACAATTTAGTATGTATTTTCTACCTCT \| ```          T       TTTTATGTATG        AAAC   mRNA  3'  CTCCATC           ATTTAACA       miRNA 5'  GAGGTAG           TAGGTTGT                T                          ATAGTT           311111115555554555411221111333366 ``` \| SeedType = 1 NonSeedType = 1 \| \| --- \| --- \| --- \| --- \| --- \| --- \| | \| 440 \| WBGene00006955:wrt-9(3932) \| ``` log2FC=-0.235 padj=0.839 ``` \| [seedLoc = 328,334] Seq=AATTCATTTCCGTGCTACTCTCC \| ```          C   T      G CTTTACTTAA mRNA  3'  CTC CATCGT C           miRNA 5'  GAG GTAGTA G                    T          G TTGTATAGTT           31115111121313333333333 ``` \| SeedType = 2 NonSeedType = -1 \| \| --- \| --- \| --- \| --- \| --- \| --- \| | \| 441 \| WBGene00006956:wrt-10(3933) \| ``` log2FC=0.658 padj=0.073 over-expressed ``` \| [seedLoc = 26,31] Seq=CACTATTTTTTGTTACCTCT \| ```          T           TT       C mRNA  3'  CTCCATTGTTT    TATCA  miRNA 5'  GAGGTAGTAGG    ATAGT           T           TTGT     T           3111111221223366111113 ``` \| SeedType = 1 NonSeedType = -1 \| \| --- \| --- \| --- \| --- \| --- \| --- \| | \| 442 \| WBGene00006974:zen-4(3945) \| ``` log2FC=-0.076 padj=0.945 ``` \| [seedLoc = 193,198] Seq=CCGTCTTTGATCTTTTTACCTCC \| ```          C       TT     TTTCTGCC mRNA  3'  CTCCATT  TCTAG         miRNA 5'  GAGGTAG  AGGTT                  T       T      GTATAGTT           31111112351121233333333 ``` \| SeedType = 1 NonSeedType = 2 \| \| --- \| --- \| --- \| --- \| --- \| --- \| | \| 443 \| WBGene00006975:zfp-1(3949) \| ``` log2FC=-0.049 padj=0.974 ``` \| [seedLoc = 70,75] Seq=GTGTTTCTAAAATAATACCTCA \| ```                 A  AA   CTTTGTG mRNA  3' ACTCCAT AT  AAT        miRNA 5' TGAGGTA TA  TTG                        G  GG   TATAGTT           1111111311331123333333 ``` \| SeedType = 1 NonSeedType = -1 \| \| --- \| --- \| --- \| --- \| --- \| --- \| | \|  \|  \|  \| [seedLoc = 83,86] Seq=GTTTCTAAAATAATACCTCAA \| ```          A       A  AA   CTTTG   mRNA  3'  ACTCCAT AT  AAT        miRNA 5'  TGAGGTA TA  TTG                         G  GG   TATAGTT           51111111311331123333366 ``` \| SeedType = 1 NonSeedType = -1 \| \| --- \| --- \| --- \| --- \| --- \| --- \| | \|  \|  \|  \| [seedLoc = 228,233] Seq=TTCTATAAAAATCCAATAACTACCTTT \| ```          T        A     CTAAAA     TT mRNA  3'  TTCCATCA  TAAC      ATATC   miRNA 5'  GAGGTAGT  GTTG      TATAG            T        AG               TT           3211111113621115555551111133 ``` \| SeedType = 3 NonSeedType = 1 \| \| --- \| --- \| --- \| --- \| --- \| --- \| | \| 444 \| WBGene00006978:zig-1(3952) \| ``` log2FC=0.352 padj=0.475 ``` \| [seedLoc = 142,147] Seq=AAATTTCATATTTGATACCTCA \| ```                 A     TACTTTAAA mRNA  3' ACTCCAT GTTTA          miRNA 5' TGAGGTA TAGGT                          G     TGTATAGTT           1111111321221333333333 ``` \| SeedType = 1 NonSeedType = 2 \| \| --- \| --- \| --- \| --- \| --- \| --- \| | \| 445 \| WBGene00007008:rfp-1(3973) \| ``` log2FC=-0.101 padj=0.918 ``` \| [seedLoc = 173,178] Seq=AGTTATTGAATCACTATCTCT \| ```          T             GT    GA mRNA  3'  CTCTATCA CTAA  TATT   miRNA 5'  GAGGTAGT GGTT  ATAG            T        A    GT    TT           3111211116121133111233 ``` \| SeedType = 2 NonSeedType = 2 \| \| --- \| --- \| --- \| --- \| --- \| --- \| | \| 446 \| WBGene00007016:mdt-15(3981) \| ``` log2FC=-0.278 padj=0.221 ``` \| [seedLoc = 237,242] Seq=TTTTCCACGAGACTCCTGCCTCC \| ```          C       C  AG    CCTTTT mRNA  3'  CTCCGTC TC  AGCA       miRNA 5'  GAGGTAG AG  TTGT                T       T  G     ATAGTT           31111211311341211333333 ``` \| SeedType = 2 NonSeedType = -1 \| \| --- \| --- \| --- \| --- \| --- \| --- \| | \|  \|  \|  \| [seedLoc = 565,570] Seq=CACTGTTCAAAAATACCTCA \| ```                 AAA     T     C mRNA  3' ACTCCAT     AAC TGTCA  miRNA 5' TGAGGTA     TTG ATAGT                  GTAGG   T     T           1111111333661113121113 ``` \| SeedType = 1 NonSeedType = 1 \| \| --- \| --- \| --- \| --- \| --- \| --- \| | \| 447 \| WBGene00007024:plin-1(3991) \|  \| [seedLoc = 447,452] Seq=TACTAATCGAAGAGCTCTACTTCG \| ```          G          G  AAGCTA    T mRNA  3'  CTTCATC TC AG      ATCA  miRNA 5'  GAGGTAG AG TT      TAGT           T       T  G  GTA       T           3112111161131233355511113 ``` \| SeedType = 3 NonSeedType = -1 \| \| --- \| --- \| --- \| --- \| --- \| --- \| | \| 448 \| WBGene00007030:epc-1(3997) \| ``` log2FC=-0.169 padj=0.728 ``` \| [seedLoc = 272,278] Seq=TGTCTCATCATCTCGGTACTCTCA \| ```              T   G C    CTACTCTGT mRNA  3' ACTC CAT G TCTA          miRNA 5' TGAG GTA T AGGT                           G      TGTATAGTT           111151113251121333333333 ``` \| SeedType = 2 NonSeedType = 2 \| \| --- \| --- \| --- \| --- \| --- \| --- \| | \| 449 \| WBGene00007042:pbrm-1(4000) \| ``` log2FC=-0.028 padj=0.984 ``` \| [seedLoc = 64,69] Seq=TAATTATCCTACCCTATCTCG \| ```          G            T C      T mRNA  3'  CTCTATC  CCA C TATTAA  miRNA 5'  GAGGTAG  GGT G ATAGTT           T       TA   T T                  31112111661113131112115 ``` \| SeedType = 2 NonSeedType = 1 \| \| --- \| --- \| --- \| --- \| --- \| --- \| | \| 450 \| WBGene00007053:chd-7(4009) \| ``` log2FC=0.141 padj=0.848 ``` \| [seedLoc = 269,274] Seq=CAGAAAATTATCTCTCTATCTCT \| ```          T       TC    TTAAAAGAC mRNA  3'  CTCTATC  TCTA          miRNA 5'  GAGGTAG  AGGT                   T       T     TGTATAGTT           31112111351121333333333 ``` \| SeedType = 2 NonSeedType = 2 \| \| --- \| --- \| --- \| --- \| --- \| --- \| | \| 451 \| WBGene00007058:dmd-6(4014) \| ``` log2FC=-0.283 padj=0.553 ``` \| [seedLoc = 151,156] Seq=TTTCATACTTTTTACTATCTCT \| ```          T           TT    CTTT mRNA  3'  CTCTATCATTT  CATA     miRNA 5'  GAGGTAGTAGG  GTAT              T           TT    AGTT           3111211111223311113333 ``` \| SeedType = 2 NonSeedType = -1 \| \| --- \| --- \| --- \| --- \| --- \| --- \| | \| 452 \| WBGene00007109:edg-1(4047) \|  \| [seedLoc = 220,225] Seq=TTTTTCCCGATTTATTCTACCTCA \| ```                  TT       CCTTTTT mRNA  3' ACTCCATC  ATTTAGC        miRNA 5' TGAGGTAG  TAGGTTG                                  TATAGTT           111111115511221213333333 ``` \| SeedType = 1 NonSeedType = 2 \| \| --- \| --- \| --- \| --- \| --- \| --- \| | \| 453 \| WBGene00007131:pals-26(4065) \|  \| [seedLoc = 170,175] Seq=GGCTTTTGTCTGCTACCTTT \| ```          T           GTTT     G mRNA  3'  TTCCATCGTCT      TCG  miRNA 5'  GAGGTAGTAGG      AGT           T           TTGTAT   T           3211111121123333661123 ``` \| SeedType = 3 NonSeedType = -1 \| \| --- \| --- \| --- \| --- \| --- \| --- \| | \|  \|  \|  \| [seedLoc = 328,334] Seq=CCATCCACACTTATCTACCGTCA \| ```             G     T        CCTACC mRNA  3' ACT CCATC ATTC ACA       miRNA 5' TGA GGTAG TAGG TGT                              T   ATAGTT           111411111511216111333333 ``` \| SeedType = 3 NonSeedType = -1 \| \| --- \| --- \| --- \| --- \| --- \| --- \| | \| 454 \| WBGene00007132:pals-27(4066) \|  \| [seedLoc = 51,56] Seq=CATTGTATAAGAACTACCTCC \| ```          C        AG          C mRNA  3'  CTCCATCA   AATATGTTA  miRNA 5'  GAGGTAGT   TTGTATAGT           T        AGG         T           3111111113361121121213 ``` \| SeedType = 1 NonSeedType = -1 \| \| --- \| --- \| --- \| --- \| --- \| --- \| | \| 455 \| WBGene00007135:cdk-12(4068) \| ``` log2FC=0.169 padj=0.745 ``` \| [seedLoc = 196,201] Seq=TCTTCCCCCGCCTCATACCTCC \| ```          C      AC    CCCCCTTCT mRNA  3'  CTCCAT  TCCG          miRNA 5'  GAGGTA  AGGT                   T      GT    TGTATAGTT           3111111331112333333333 ``` \| SeedType = 1 NonSeedType = 2 \| \| --- \| --- \| --- \| --- \| --- \| --- \| | \| 456 \| WBGene00007227:C01G6.5(4137) \| ``` log2FC=-0.071 padj=0.946 ``` \| [seedLoc = 479,485] Seq=AACATCATCATCTTTTACCATCC \| ```          C  A     T    CTACTACAA mRNA  3'  CT CCATT TCTA          miRNA 5'  GA GGTAG AGGT                   T        T    TGTATAGTT           31151111231121333333333 ``` \| SeedType = 3 NonSeedType = 2 \| \| --- \| --- \| --- \| --- \| --- \| --- \| | \| 457 \| WBGene00007255:cnnm-4(4159) \|  \| [seedLoc = 192,197] Seq=TCTCATAATTTATTTTTTTTGATCTACCTTG \| ```          G       TAGTTTTTTTT        CTCT   mRNA  3'  TTCCATC           ATTTAATA       miRNA 5'  GAGGTAG           TAGGTTGT                T                          ATAGTT           321111115545555555511221121333366 ``` \| SeedType = 3 NonSeedType = 2 \| \| --- \| --- \| --- \| --- \| --- \| --- \| | \| 458 \| WBGene00007265:C02F4.5(4167) \| ``` log2FC=0.1 padj=1.0 ``` \| [seedLoc = 241,247] Seq=CGTTTTTTCGTCTTCTACGCTCT \| ```          T   G    T   GCTTTTTTGC mRNA  3'  CTC CATC TCT           miRNA 5'  GAG GTAG AGG                    T        T   TTGTATAGTT           31114111131123333333333 ``` \| SeedType = 2 NonSeedType = -1 \| \| --- \| --- \| --- \| --- \| --- \| --- \| | \| 459 \| WBGene00007287:gck-4(4182) \| ``` log2FC=-0.114 padj=0.893 ``` \| [seedLoc = 109,114] Seq=CCACATTCGACACCTGCCTCG \| ```          G       CA     TTACACC mRNA  3'  CTCCGTC   CAGC        miRNA 5'  GAGGTAG   GTTG                 T       TAG    TATAGTT           3111121133611213333333 ``` \| SeedType = 2 NonSeedType = -1 \| \| --- \| --- \| --- \| --- \| --- \| --- \| | \| 460 \| WBGene00007322:C05B5.5(4207) \| ``` log2FC=0.482 padj=0.899 ``` \| [seedLoc = 264,269] Seq=AGGCCATAAAAATCCGCTACCTCC \| ```          C        C    AA   C   A mRNA  3'  CTCCATCG CTAA  ATA CGG  miRNA 5'  GAGGTAGT GGTT  TAT GTT           T        A    G    A               311111112312113511131225 ``` \| SeedType = 1 NonSeedType = 2 \| \| --- \| --- \| --- \| --- \| --- \| --- \| | \| 461 \| WBGene00007350:sucl-1(4229) \| ``` log2FC=0.099 padj=0.918 ``` \| [seedLoc = 342,347] Seq=CGACTGGACGATGGTGCTACTTCT \| ```          T         GG     G     C mRNA  3'  CTTCATCGT  TAGCA GTCAG  miRNA 5'  GAGGTAGTA  GTTGT TAGTT           T         G      A                 311211112134212113211125 ``` \| SeedType = 3 NonSeedType = -1 \| \| --- \| --- \| --- \| --- \| --- \| --- \| | \| 462 \| WBGene00007358:lact-6(4235) \| ``` log2FC=-0.749 padj=1.0 ``` \| [seedLoc = 34,39] Seq=ATTTTTCTAACAGTTACCTCA \| ```                  GA     CTTTTTA mRNA  3' ACTCCATT   CAAT        miRNA 5' TGAGGTAG   GTTG                         TAG    TATAGTT           1111111233611123333333 ``` \| SeedType = 1 NonSeedType = 1 \| \| --- \| --- \| --- \| --- \| --- \| --- \| | \|  \|  \|  \| [seedLoc = 40,44] Seq=ATTTTTCTAACAGTTACCTCAC \| ```          C        GA     CTTTTTA mRNA  3'  ACTCCATT   CAAT        miRNA 5'  TGAGGTAG   GTTG                          TAG    TATAGTT           51111111233611123333333 ``` \| SeedType = 1 NonSeedType = 1 \| \| --- \| --- \| --- \| --- \| --- \| --- \| | \| 463 \| WBGene00007372:C06B8.7(4244) \| ``` log2FC=-0.189 padj=0.837 ``` \| [seedLoc = 36,41] Seq=GAATTATTTTCATTATTACCTCC \| ```          C          ACTTT      G mRNA  3'  CTCCATTATT     TATTAA  miRNA 5'  GAGGTAGTAG     ATAGTT           T          GTTGT                  31111112112333331112114 ``` \| SeedType = 1 NonSeedType = -1 \| \| --- \| --- \| --- \| --- \| --- \| --- \| | \| 464 \| WBGene00007391:C06H2.7(4255) \| ``` log2FC=0.202 padj=0.966 ``` \| [seedLoc = 66,72] Seq=TATCATATTCATCTGTACCCTCA \| ```             C          CT    CTAT mRNA  3' ACT CCAT GTCTA  TATA     miRNA 5' TGA GGTA TAGGT  GTAT                      G     T     AGTT           111511116211213521113333 ``` \| SeedType = 3 NonSeedType = 2 \| \| --- \| --- \| --- \| --- \| --- \| --- \| | \| 465 \| WBGene00007416:ceh-57(4280) \| ``` log2FC=0.753 padj=1.0 ``` \| [seedLoc = 66,72] Seq=TCTTATATATGTGCTCACCTCG \| ```          G     C    G         CT mRNA  3'  CTCCA TCGT T ATATATT   miRNA 5'  GAGGT AGTA G TGTATAG            T          G T       TT           31111151121326121111233 ``` \| SeedType = 2 NonSeedType = -1 \| \| --- \| --- \| --- \| --- \| --- \| --- \| | \|  \|  \|  \| [seedLoc = 98,103] Seq=TACATATATTAAATACATACCTCA \| ```                 A   AAAT     A  T mRNA  3' ACTCCAT CAT    TATAT CA  miRNA 5' TGAGGTA GTA    GTATA GT                      GGTT        T           111111151113333211115113 ``` \| SeedType = 1 NonSeedType = -1 \| \| --- \| --- \| --- \| --- \| --- \| --- \| | \| 466 \| WBGene00007418:C07E3.8(4282) \| ``` log2FC=0.1 padj=1.0 ``` \| [seedLoc = 74,80] Seq=ATGAGTATAAAAATACTCTCC \| ```          C   T   AAA        AGTA mRNA  3'  CTC CAT     AATATG     miRNA 5'  GAG GTA     TTGTAT              T       GTAGG      AGTT           31115111333661121123333 ``` \| SeedType = 2 NonSeedType = -1 \| \| --- \| --- \| --- \| --- \| --- \| --- \| | \| 467 \| WBGene00007433:swsn-7(4294) \| ``` log2FC=0.005 padj=0.997 ``` \| [seedLoc = 304,309] Seq=CTCTCTCTCTCCATCCTATCTCC \| ```          C       CTA  TCTCTCTCTC mRNA  3'  CTCTATC   CC           miRNA 5'  GAGGTAG   GG                    T       TA   TTGTATAGTT           31112111335113333333333 ``` \| SeedType = 2 NonSeedType = -1 \| \| --- \| --- \| --- \| --- \| --- \| --- \| | \| 468 \| WBGene00007499:C09H10.5(4337) \| ``` log2FC=-0.026 padj=0.993 ``` \| [seedLoc = 414,419] Seq=ATTTGTTTCGCCCAAAAATACCTCT \| ```          T      AAAAAC   CTT    TA mRNA  3'  CTCCAT      CCG   TGTT   miRNA 5'  GAGGTA      GGT   ATAG            T      GTA      TGT    TT           3111111333555112333121233 ``` \| SeedType = 1 NonSeedType = 2 \| \| --- \| --- \| --- \| --- \| --- \| --- \| | \| 469 \| WBGene00007506:C10C5.2(4343) \| ``` log2FC=-0.69 padj=0.039 ``` \| [seedLoc = 395,400] Seq=TGTTCTAAAACACTACTTCC \| ```          C             AATCTTGT mRNA  3'  CTTCATCA  CAA         miRNA 5'  GAGGTAGT  GTT                  T        AG   GTATAGTT           3112111116611133333333 ``` \| SeedType = 3 NonSeedType = 1 \| \| --- \| --- \| --- \| --- \| --- \| --- \| | \| 470 \| WBGene00007600:C15C6.1(4418) \| ``` log2FC=-0.396 padj=0.227 ``` \| [seedLoc = 47,52] Seq=CTAACACTAAGCTTTACCTCT \| ```          T          G   CACAATC mRNA  3'  CTCCATT TC AAT        miRNA 5'  GAGGTAG AG TTG                 T       T  G   TATAGTT           3111111261131123333333 ``` \| SeedType = 1 NonSeedType = -1 \| \| --- \| --- \| --- \| --- \| --- \| --- \| | \| 471 \| WBGene00007606:C15C8.4(4424) \| ``` log2FC=0.35 padj=0.422 ``` \| [seedLoc = 71,77] Seq=ATAATGAGAGCTCATTTCTACCTGC \| ```          CG      TTTAC    G GTAATA mRNA  3'   TCCATC     TCGA A       miRNA 5'   AGGTAG     GGTT T                TG      TA       G ATAGTT           3311111133555212131333333 ``` \| SeedType = 3 NonSeedType = 2 \| \| --- \| --- \| --- \| --- \| --- \| --- \| | \|  \|  \|  \| [seedLoc = 183,188] Seq=CACGTGTATATTTTGTTCGTAAGCTACTTCA \| ```                   AATGCTTG   T      G  C mRNA  3' ACTTCATCG        TTT ATATGT CA  miRNA 5' TGAGGTAGT        AGG TGTATA GT                               T         T           1112111125554555412231211214113 ``` \| SeedType = 3 NonSeedType = -1 \| \| --- \| --- \| --- \| --- \| --- \| --- \| | \| 472 \| WBGene00007620:ocrl-1(4432) \| ``` log2FC=0.119 padj=0.907 ``` \| [seedLoc = 299,304] Seq=TAAGTTATAAATCTACCTCA \| ```                  TA       TGAAT mRNA  3' ACTCCATC    AATAT      miRNA 5' TGAGGTAG    TTGTA                       TAGG     TAGTT           1111111133661121133333 ``` \| SeedType = 1 NonSeedType = -1 \| \| --- \| --- \| --- \| --- \| --- \| --- \| | \| 473 \| WBGene00007635:npr-4(4446) \| ``` log2FC=-0.033 padj=1.0 ``` \| [seedLoc = 219,225] Seq=AAATTATCACGTTGACTCCATACGCTCC \| ```          C   G   ACC    TTG  C      A mRNA  3'  CTC CAT   TCAG   CA TATTAA  miRNA 5'  GAG GTA   GGTT   GT ATAGTT           T       GTA                            3111411133321125541151112115 ``` \| SeedType = 2 NonSeedType = 2 \| \| --- \| --- \| --- \| --- \| --- \| --- \| | \| 474 \| WBGene00007645:C17E4.6(4452) \| ``` log2FC=0.038 padj=0.983 ``` \| [seedLoc = 142,147] Seq=CACTTGGCAGCTCATTTCTCTATCTCT \| ```          T       TCTTTAC      GT   C mRNA  3'  CTCTATC       TCGACG  TCA  miRNA 5'  GAGGTAG       GGTTGT  AGT           T       TA           AT   T           311121113355555212112331113 ``` \| SeedType = 2 NonSeedType = 2 \| \| --- \| --- \| --- \| --- \| --- \| --- \| | \|  \|  \|  \| [seedLoc = 328,334] Seq=CTCGGCGCAGCATATACCATCA \| ```             A    ATA      CGGCTC mRNA  3' ACT CCAT    CGACG       miRNA 5' TGA GGTA    GTTGT                        GTAG     ATAGTT           11151111333612112333333 ``` \| SeedType = 3 NonSeedType = -1 \| \| --- \| --- \| --- \| --- \| --- \| --- \| | \| 475 \| WBGene00007646:nkb-1(4453) \| ``` log2FC=0.298 padj=0.139 ``` \| [seedLoc = 45,50] Seq=ATCGCTAGGATCTAAGTGCATCTATCTCA \| ```                  TACGTGA      G  CGCTA mRNA  3' ACTCTATC       ATCTAG AT      miRNA 5' TGAGGTAG       TAGGTT TA                                    G  TAGTT           11112111555454511121231133333 ``` \| SeedType = 2 NonSeedType = 2 \| \| --- \| --- \| --- \| --- \| --- \| --- \| | \| 476 \| WBGene00007647:C17E4.10(4454) \| ``` log2FC=-0.015 padj=0.994 ``` \| [seedLoc = 158,163] Seq=AATTCCAAAACTTCCTACTTCA \| ```                  C     AACCTTAA mRNA  3' ACTTCATC TTCAA         miRNA 5' TGAGGTAG AGGTT                          T     GTATAGTT           1112111131211133333333 ``` \| SeedType = 3 NonSeedType = 1 \| \| --- \| --- \| --- \| --- \| --- \| --- \| | \| 477 \| WBGene00007651:nra-3(4457) \| ``` log2FC=0.224 padj=0.859 ``` \| [seedLoc = 264,269] Seq=CAACATGCGTGAAAATCCTCAACTACTTCG \| ```          G        AC   TAAAAGT         C mRNA  3'  CTTCATCA  TCC       GCGTA CAA  miRNA 5'  GAGGTAGT  AGG       TGTAT GTT           T             T           A               3112111115511135555452121161115 ``` \| SeedType = 3 NonSeedType = -1 \| \| --- \| --- \| --- \| --- \| --- \| --- \| | \| 478 \| WBGene00007666:C18B12.4(4467) \| ``` log2FC=-0.313 padj=0.464 ``` \| [seedLoc = 96,102] Seq=TTTCCCCCCTCCTGTACACTCC \| ```          C   A        TC CCCCTTT mRNA  3'  CTC CAT GTCC  C        miRNA 5'  GAG GTA TAGG  G                 T       G    TT TATAGTT           31115111621113313333333 ``` \| SeedType = 2 NonSeedType = -1 \| \| --- \| --- \| --- \| --- \| --- \| --- \| | \|  \|  \|  \| [seedLoc = 252,258] Seq=AGTCTTGCAACCTAGTCACCTCT \| ```          T     C G         TCTGA mRNA  3'  CTCCA T ATCCAACGT      miRNA 5'  GAGGT A TAGGTTGTA               T       G         TAGTT           31111151311111112133333 ``` \| SeedType = 2 NonSeedType = 1 \| \| --- \| --- \| --- \| --- \| --- \| --- \| | \| 479 \| WBGene00007675:C18D4.6(4470) \| ``` log2FC=0.369 padj=0.887 ``` \| [seedLoc = 36,41] Seq=CTCAATATTTGATCTACCTCT \| ```          T       TAGTT     ACTC mRNA  3'  CTCCATC      TATA     miRNA 5'  GAGGTAG      GTAT              T       TAGGTT    AGTT           3111111133333621113333 ``` \| SeedType = 1 NonSeedType = -1 \| \| --- \| --- \| --- \| --- \| --- \| --- \| | \| 480 \| WBGene00007679:C18D11.1(4473) \| ``` log2FC=0.054 padj=0.977 ``` \| [seedLoc = 91,96] Seq=TGCTCCTCGCCACTACCTTT \| ```          T            CTCC    T mRNA  3'  TTCCATCA CCG     TCG  miRNA 5'  GAGGTAGT GGT     AGT           T        A   TGTAT   T           3211111116112333361123 ``` \| SeedType = 3 NonSeedType = 2 \| \| --- \| --- \| --- \| --- \| --- \| --- \| | \| 481 \| WBGene00007681:C18D11.3(4475) \| ``` log2FC=0.466 padj=0.088 ``` \| [seedLoc = 225,230] Seq=GGAGGCTCCCCCTATATCTGCCTCC \| ```          C       TAT    CCCTCGGAGG mRNA  3'  CTCCGTC   ATCC           miRNA 5'  GAGGTAG   TAGG                    T              TTGTATAGTT           3111121155511113333333333 ``` \| SeedType = 2 NonSeedType = -1 \| \| --- \| --- \| --- \| --- \| --- \| --- \| | \| 482 \| WBGene00007705:C25A1.1(4500) \| ``` log2FC=0.034 padj=0.989 ``` \| [seedLoc = 76,81] Seq=CTGAATGCATCCAATACCTCT \| ```          T      AA   T     AGTC mRNA  3'  CTCCAT   CC ACGTA     miRNA 5'  GAGGTA   GG TGTAT              T      GTA  T     AGTT           3111111336113112113333 ``` \| SeedType = 1 NonSeedType = -1 \| \| --- \| --- \| --- \| --- \| --- \| --- \| | \| 483 \| WBGene00007707:fath-1(4501) \|  \| [seedLoc = 49,54] Seq=ATTGAGTGAACTCTGCCTCA \| ```                        GTGAGTTA mRNA  3' ACTCCGTC  TCAA         miRNA 5' TGAGGTAG  GGTT                          TA    GTATAGTT           1111121166211133333333 ``` \| SeedType = 2 NonSeedType = 1 \| \| --- \| --- \| --- \| --- \| --- \| --- \| | \|  \|  \|  \| [seedLoc = 63,68] Seq=GAGTGAACTCTGCCTCAC \| ```          C              GTGAG    mRNA  3'  ACTCCGTC  TCAA         miRNA 5'  TGAGGTAG  GGTT                           TA    GTATAGTT           51111121166211133333666 ``` \| SeedType = 2 NonSeedType = 1 \| \| --- \| --- \| --- \| --- \| --- \| --- \| | \|  \|  \|  \| [seedLoc = 223,229] Seq=TACTCAGCACGTGCCTATTATCCTCA \| ```               T        TGC   AC   T mRNA  3' ACTCC ATTATCCG   ACG  TCA  miRNA 5' TGAGG TAGTAGGT   TGT  AGT                               AT   T           11111511211112545112331113 ``` \| SeedType = 2 NonSeedType = 2 \| \| --- \| --- \| --- \| --- \| --- \| --- \| | \|  \|  \|  \| [seedLoc = 230,234] Seq=TACTCAGCACGTGCCTATTATCCTCAC \| ```          C     T        TGC   AC   T mRNA  3'  ACTCC ATTATCCG   ACG  TCA  miRNA 5'  TGAGG TAGTAGGT   TGT  AGT                                AT   T           511111511211112545112331113 ``` \| SeedType = 2 NonSeedType = 2 \| \| --- \| --- \| --- \| --- \| --- \| --- \| | \| 484 \| WBGene00007734:C25G4.7(4522) \| ``` log2FC=0.039 padj=1.0 ``` \| [seedLoc = 93,98] Seq=GTCTCCCCAAGTGCCTACCTCA \| ```                  CG G   CCCTCTG mRNA  3' ACTCCATC  T AAC        miRNA 5' TGAGGTAG  G TTG                         TA G   TATAGTT           1111111133231113333333 ``` \| SeedType = 1 NonSeedType = 1 \| \| --- \| --- \| --- \| --- \| --- \| --- \| | \| 485 \| WBGene00007741:glb-8(4525) \| ``` log2FC=0.269 padj=0.945 ``` \| [seedLoc = 63,68] Seq=TTTTCTGAAATTATCTATCTCA \| ```                  TA    AGTCTTTT mRNA  3' ACTCTATC  TTAA         miRNA 5' TGAGGTAG  GGTT                          TA    GTATAGTT           1111211133221133333333 ``` \| SeedType = 2 NonSeedType = 2 \| \| --- \| --- \| --- \| --- \| --- \| --- \| | \| 486 \| WBGene00007761:rad-26(4542) \| ``` log2FC=-0.029 padj=0.987 ``` \| [seedLoc = 63,68] Seq=TAAAATACAACATCTATCTCA \| ```                  TA        AAAT mRNA  3' ACTCTATC   CAACATA     miRNA 5' TGAGGTAG   GTTGTAT                      TAG       AGTT           1111211133611111113333 ``` \| SeedType = 2 NonSeedType = 1 \| \| --- \| --- \| --- \| --- \| --- \| --- \| | \| 487 \| WBGene00007766:C27C7.1(4547) \| ``` log2FC=-0.158 padj=0.806 ``` \| [seedLoc = 211,217] Seq=GCCCTCAAAACCTGTTGGATTGTACACTCA \| ```              A   GTTAGGTT      AACTCCCG mRNA  3' ACTC CAT        GTCCAA         miRNA 5' TGAG GTA        TAGGTT                          G             GTATAGTT           111151113555445521111133333333 ``` \| SeedType = 2 NonSeedType = 1 \| \| --- \| --- \| --- \| --- \| --- \| --- \| | \| 488 \| WBGene00007773:C27C12.3(4551) \| ``` log2FC=0.964 padj=0.104 ``` \| [seedLoc = 570,575] Seq=TTATTTAGAATAGTATTACCTCT \| ```          T         GA   G  TTATT mRNA  3'  CTCCATTAT  TAA AT      miRNA 5'  GAGGTAGTA  GTT TA               T         G    G  TAGTT           31111112113521131133333 ``` \| SeedType = 1 NonSeedType = -1 \| \| --- \| --- \| --- \| --- \| --- \| --- \| | \| 489 \| WBGene00007787:C27H6.8(4562) \| ``` log2FC=0.084 padj=0.927 ``` \| [seedLoc = 150,156] Seq=TCTTTTTCCATCTCTTACACTCC \| ```          C   A    C    CCTTTTTCT mRNA  3'  CTC CATT TCTA          miRNA 5'  GAG GTAG AGGT                   T        T    TGTATAGTT           31115111231121333333333 ``` \| SeedType = 2 NonSeedType = 2 \| \| --- \| --- \| --- \| --- \| --- \| --- \| | \| 490 \| WBGene00007811:C29F7.2(4579) \| ``` log2FC=-0.554 padj=0.008 ``` \| [seedLoc = 90,95] Seq=TTTTGTTCAAATATACCTCC \| ```          C      ATA     T    TT mRNA  3'  CTCCAT     AAC TGTT   miRNA 5'  GAGGTA     TTG ATAG            T      GTAGG   T    TT           3111111333661113121233 ``` \| SeedType = 1 NonSeedType = 1 \| \| --- \| --- \| --- \| --- \| --- \| --- \| | \| 491 \| WBGene00007813:jmjd-3.3(4581) \| ``` log2FC=-1.541 padj=1.0 ``` \| [seedLoc = 111,116] Seq=GTGTTAGCATCACCTACTTCA \| ```                  CA      ATTGTG mRNA  3' ACTTCATC  CT ACG       miRNA 5' TGAGGTAG  GG TGT                        TA  T   ATAGTT           1112111133126112333333 ``` \| SeedType = 3 NonSeedType = -1 \| \| --- \| --- \| --- \| --- \| --- \| --- \| | \|  \|  \|  \| [seedLoc = 679,685] Seq=GTGAAAGAAGAAAACTACCTGC \| ```          CG       AAAGAAGAAAGTG mRNA  3'   TCCATCA              miRNA 5'   AGGTAGT                       TG       AGGTTGTATAGTT           3311111113333333333333 ``` \| SeedType = 3 NonSeedType = -1 \| \| --- \| --- \| --- \| --- \| --- \| --- \| | \|  \|  \|  \| [seedLoc = 683,688] Seq=AAGAAGAAAACTACCTGCCTCA \| ```                  CA    AAGAAGAA mRNA  3' ACTCCGTC  TCAA         miRNA 5' TGAGGTAG  GGTT                          TA    GTATAGTT           1111121133211133333333 ``` \| SeedType = 2 NonSeedType = 1 \| \| --- \| --- \| --- \| --- \| --- \| --- \| | \| 492 \| WBGene00007898:C33D9.3(4631) \| ``` log2FC=-0.224 padj=0.771 ``` \| [seedLoc = 129,134] Seq=ACCTTCCCGATGTATTTTATCTTACCTCT \| ```          T      TCTATTT   G    CCTTCCA mRNA  3'  CTCCAT       TAT TAGC        miRNA 5'  GAGGTA       GTA GTTG                 T                G    TATAGTT           31111115555555211321213333333 ``` \| SeedType = 1 NonSeedType = -1 \| \| --- \| --- \| --- \| --- \| --- \| --- \| | \| 493 \| WBGene00007900:C33D9.6(4632) \|  \| [seedLoc = 75,80] Seq=CATTTTCCATATTTCTACCTTT \| ```          T            T  CTTTTAC mRNA  3'  TTCCATC TTTA AC        miRNA 5'  GAGGTAG AGGT TG                 T       T       TATAGTT           32111111612215113333333 ``` \| SeedType = 3 NonSeedType = 2 \| \| --- \| --- \| --- \| --- \| --- \| --- \| | \| 494 \| WBGene00007918:sphk-1(4644) \| ``` log2FC=0.228 padj=0.695 ``` \| [seedLoc = 196,201] Seq=TCTCACTTTTCCTCTACTTCG \| ```          G           TTTTCACTCT mRNA  3'  CTTCATC TCC           miRNA 5'  GAGGTAG AGG                    T       T   TTGTATAGTT           3112111161113333333333 ``` \| SeedType = 3 NonSeedType = -1 \| \| --- \| --- \| --- \| --- \| --- \| --- \| | \| 495 \| WBGene00007938:C34F6.1(4661) \| ``` log2FC=-0.101 padj=0.972 ``` \| [seedLoc = 267,273] Seq=TCAAAATTGACCACTACCATCC \| ```          C  A           TTAAAACT mRNA  3'  CT CCATCA CCAG         miRNA 5'  GA GGTAGT GGTT                  T         A    GTATAGTT           31151111116111233333333 ``` \| SeedType = 3 NonSeedType = 1 \| \| --- \| --- \| --- \| --- \| --- \| --- \| | \| 496 \| WBGene00007941:C34F6.7(4663) \| ``` log2FC=0.109 padj=0.952 ``` \| [seedLoc = 173,176] Seq=TTATCCGCGTCTCTCCCTCAT \| ```          T     C     T    CCTATT mRNA  3'  ACTCC TC TC  GCG       miRNA 5'  TGAGG AG AG  TGT                      T  T  GT   ATAGTT           51111131161136212333333 ``` \| SeedType = 2 NonSeedType = -1 \| \| --- \| --- \| --- \| --- \| --- \| --- \| | \| 497 \| WBGene00007975:gls-1(4684) \| ``` log2FC=0.235 padj=0.404 ``` \| [seedLoc = 502,507] Seq=TTTTTTGTAACCCTATCTCT \| ```          T              GTTTTTT mRNA  3'  CTCTATC  CCAAT        miRNA 5'  GAGGTAG  GGTTG                 T       TA     TATAGTT           3111211166111123333333 ``` \| SeedType = 2 NonSeedType = 1 \| \| --- \| --- \| --- \| --- \| --- \| --- \| | \| 498 \| WBGene00007980:C36E8.1(4690) \| ``` log2FC=0.007 padj=0.997 ``` \| [seedLoc = 102,107] Seq=TTTTGTCGTTGATTTACCTCA \| ```                  TAGTT       TT mRNA  3' ACTCCATT     GC TGTT   miRNA 5' TGAGGTAG     TG ATAG                    TAGGT  T    TT           1111111233333216121233 ``` \| SeedType = 1 NonSeedType = -1 \| \| --- \| --- \| --- \| --- \| --- \| --- \| | \| 499 \| WBGene00007993:idhb-1(4702) \| ``` log2FC=-0.024 padj=0.986 ``` \| [seedLoc = 343,349] Seq=AATCGTCTTATCTAATTTTACCATCC \| ```          C  A     TTA     TTCTGCTAA mRNA  3'  CT CCATT   ATCTA          miRNA 5'  GA GGTAG   TAGGT                   T                TGTATAGTT           31151111255511121333333333 ``` \| SeedType = 3 NonSeedType = 2 \| \| --- \| --- \| --- \| --- \| --- \| --- \| | \| 500 \| WBGene00008006:tag-325(4710) \| ``` log2FC=-0.031 padj=0.987 ``` \| [seedLoc = 295,301] Seq=GTATTTTCAATCTTCCTACGCTCA \| ```              G    CT      TTTTATG mRNA  3' ACTC CATC  TCTAAC        miRNA 5' TGAG GTAG  AGGTTG                          T       TATAGTT           111141111351121113333333 ``` \| SeedType = 2 NonSeedType = 1 \| \| --- \| --- \| --- \| --- \| --- \| --- \| | \| 501 \| WBGene00008110:C46C2.2(4768) \| ``` log2FC=0.124 padj=0.916 ``` \| [seedLoc = 45,50] Seq=TTCAAAATTATCTTACCTCG \| ```          G            TTAAAACTT mRNA  3'  CTCCAT  TCTA          miRNA 5'  GAGGTA  AGGT                   T      GT    TGTATAGTT           3111111661121333333333 ``` \| SeedType = 1 NonSeedType = 2 \| \| --- \| --- \| --- \| --- \| --- \| --- \| | \|  \|  \|  \| [seedLoc = 60,66] Seq=AATTATCTTACCTCGT \| ```          T             TTAA      mRNA  3'  GCTCCAT  TCTA          miRNA 5'  TGAGGTA  AGGT                           GT    TGTATAGTT           52111111661121333366666 ``` \| SeedType = 1 NonSeedType = 2 \| \| --- \| --- \| --- \| --- \| --- \| --- \| | \| 502 \| WBGene00008121:C46F11.6(4774) \| ``` log2FC=0.067 padj=0.982 ``` \| [seedLoc = 59,64] Seq=ATTTTTTTTGCCTCTCTACCTCT \| ```          T       TC    TTTTTTTTA mRNA  3'  CTCCATC  TCCG          miRNA 5'  GAGGTAG  AGGT                   T       T     TGTATAGTT           31111111351112333333333 ``` \| SeedType = 1 NonSeedType = 2 \| \| --- \| --- \| --- \| --- \| --- \| --- \| | \| 503 \| WBGene00008175:C48B4.12(4810) \| ``` log2FC=-3.877 padj=1.0 ``` \| [seedLoc = 75,81] Seq=TATTATATAAAAACTTGTTCACCTCC \| ```          C     C       AAA        T mRNA  3'  CTCCA TTGTTCA   ATATATTA  miRNA 5'  GAGGT AGTAGGT   TGTATAGT           T                        T           31111151221211555121111213 ``` \| SeedType = 2 NonSeedType = 2 \| \| --- \| --- \| --- \| --- \| --- \| --- \| | \| 504 \| WBGene00008219:C50B6.3(4840) \| ``` log2FC=0.082 padj=0.98 ``` \| [seedLoc = 163,168] Seq=CGCTGTATTCATCTATCTCG \| ```          G       TACT         C mRNA  3'  CTCTATC      TATGTCG  miRNA 5'  GAGGTAG      GTATAGT           T       TAGGTT       T           3111211133336621121123 ``` \| SeedType = 2 NonSeedType = -1 \| \| --- \| --- \| --- \| --- \| --- \| --- \| | \|  \|  \|  \| [seedLoc = 189,194] Seq=TGTTCCCCGTTTTACCTACCTCG \| ```          G      C     T  CCCTTGT mRNA  3'  CTCCAT CATTT GC        miRNA 5'  GAGGTA GTAGG TG                 T            T  TATAGTT           31111115111223213333333 ``` \| SeedType = 1 NonSeedType = -1 \| \| --- \| --- \| --- \| --- \| --- \| --- \| | \| 505 \| WBGene00008257:C52A11.3(4869) \| ``` log2FC=-0.239 padj=1.0 ``` \| [seedLoc = 361,367] Seq=GCTATAACATTTTTTTTCCTGACTAACCTCA \| ```               A   A    TTTTTTTTACA        mRNA  3' ACTCC ATC GTCC           ATATCG  miRNA 5' TGAGG TAG TAGG           TATAGT                         TTG              T           11111511152111333555555551111126 ``` \| SeedType = 2 NonSeedType = -1 \| \| --- \| --- \| --- \| --- \| --- \| --- \| | \| 506 \| WBGene00008262:ril-1(4873) \| ``` log2FC=0.077 padj=0.933 ``` \| [seedLoc = 30,36] Seq=TCCCATCTAGCCGAATATCCTCT \| ```          T    T  AAG     CTACCCT mRNA  3'  CTCC AT   CCGAT        miRNA 5'  GAGG TA   GGTTG                 T       GTA     TATAGTT           31111511333112123333333 ``` \| SeedType = 2 NonSeedType = 2 \| \| --- \| --- \| --- \| --- \| --- \| --- \| | \| 507 \| WBGene00008311:rskn-2(4895) \| ``` log2FC=-0.051 padj=0.966 ``` \| [seedLoc = 98,104] Seq=TGCTTTCTAGTCTGTACTCTCT \| ```          T   T           CTT   T mRNA  3'  CTC CAT GTCTGAT   TCG  miRNA 5'  GAG GTA TAGGTTG   AGT           T       G       TAT   T           31115111621122123331123 ``` \| SeedType = 2 NonSeedType = 2 \| \| --- \| --- \| --- \| --- \| --- \| --- \| | \| 508 \| WBGene00008363:D1046.2(4932) \| ``` log2FC=0.012 padj=0.994 ``` \| [seedLoc = 157,163] Seq=TTCTGTTCTCTACACTCC \| ```          C   A                TT mRNA  3'  CTC CATC TCT    TGTC   miRNA 5'  GAG GTAG AGG    ATAG            T        T   TTGT    TT           31115111161126666121133 ``` \| SeedType = 2 NonSeedType = -1 \| \| --- \| --- \| --- \| --- \| --- \| --- \| | \| 509 \| WBGene00008410:D2023.6(4971) \| ``` log2FC=0.323 padj=0.253 ``` \| [seedLoc = 89,95] Seq=TTTATTGCATTTCCATTTTTGCTCACCTCT \| ```          T     C    TTTTA  TTT    TATTT mRNA  3'  CTCCA TCGT     CC   ACGT      miRNA 5'  GAGGT AGTA     GG   TGTA               T                 T      TAGTT           311111511215555511355112133333 ``` \| SeedType = 2 NonSeedType = -1 \| \| --- \| --- \| --- \| --- \| --- \| --- \| | \| 510 \| WBGene00008412:D2030.2(4973) \| ``` log2FC=-0.581 padj=0.014 ``` \| [seedLoc = 91,97] Seq=TCCTATTCGTATCTGTACACTCT \| ```          T   A         T  T    CT mRNA  3'  CTC CAT GTCTA GC TATC   miRNA 5'  GAG GTA TAGGT TG ATAG            T       G        T    TT           311151116211215213111133 ``` \| SeedType = 2 NonSeedType = 2 \| \| --- \| --- \| --- \| --- \| --- \| --- \| | \| 511 \| WBGene00008443:pde-3(5004) \| ``` log2FC=-0.096 padj=0.967 ``` \| [seedLoc = 246,252] Seq=TGACTATGTCATTTTTTGTACCCTCT \| ```          T  C    GTT   T  T       T mRNA  3'  CT CCAT   TTT AC GTATCAG  miRNA 5'  GA GGTA   AGG TG TATAGTT           T       GT    T                      31151111335122311521111125 ``` \| SeedType = 3 NonSeedType = -1 \| \| --- \| --- \| --- \| --- \| --- \| --- \| | \| 512 \| WBGene00008454:E02A10.4(5016) \| ``` log2FC=0.415 padj=0.899 ``` \| [seedLoc = 50,55] Seq=GAAATTTTCACCACTACCTCC \| ```          C            CTTTTAAAG mRNA  3'  CTCCATCA CCA          miRNA 5'  GAGGTAGT GGT                   T        A   TGTATAGTT           3111111116111333333333 ``` \| SeedType = 1 NonSeedType = 1 \| \| --- \| --- \| --- \| --- \| --- \| --- \| | \| 513 \| WBGene00008482:cut-4(5031) \| ``` log2FC=-0.221 padj=0.791 ``` \| [seedLoc = 114,119] Seq=ATTCATATCATTGTCTACCTTA \| ```                  TG   C    CTTA mRNA  3' ATTCCATC  TTA TATA     miRNA 5' TGAGGTAG  GGT GTAT                      TA   T    AGTT           1211111133221321113333 ``` \| SeedType = 3 NonSeedType = 2 \| \| --- \| --- \| --- \| --- \| --- \| --- \| | \|  \|  \|  \| [seedLoc = 123,127] Seq=ATTCATATCATTGTCTACCTTAT \| ```          T        TG   C    CTTA mRNA  3'  ATTCCATC  TTA TATA     miRNA 5'  TGAGGTAG  GGT GTAT                       TA   T    AGTT           51211111133221321113333 ``` \| SeedType = 3 NonSeedType = 2 \| \| --- \| --- \| --- \| --- \| --- \| --- \| | \| 514 \| WBGene00008536:F02H6.3(5071) \| ``` log2FC=0.462 padj=1.0 ``` \| [seedLoc = 140,145] Seq=GTCAAATAGTTAAACTACCTCA \| ```                   AATTGATAAACTG mRNA  3' ACTCCATCA              miRNA 5' TGAGGTAGT                                AGGTTGTATAGTT           1111111113333333333333 ``` \| SeedType = 1 NonSeedType = -1 \| \| --- \| --- \| --- \| --- \| --- \| --- \| | \|  \|  \|  \| [seedLoc = 156,160] Seq=TAGTTAAACTACCTCAA \| ```          A         AATTGAT       mRNA  3'  ACTCCATCA              miRNA 5'  TGAGGTAGT                                 AGGTTGTATAGTT           51111111113333333666666 ``` \| SeedType = 1 NonSeedType = -1 \| \| --- \| --- \| --- \| --- \| --- \| --- \| | \| 515 \| WBGene00008555:F07C6.4(5085) \| ``` log2FC=0.106 padj=0.928 ``` \| [seedLoc = 348,354] Seq=TTTTCTTTTTCTTATTCACCTCT \| ```          T     C      TTTTTCTTTT mRNA  3'  CTCCA TTATTC           miRNA 5'  GAGGT AGTAGG                    T            TTGTATAGTT           31111151211213333333333 ``` \| SeedType = 2 NonSeedType = -1 \| \| --- \| --- \| --- \| --- \| --- \| --- \| | \| 516 \| WBGene00008555:F07C6.4(5087) \| ``` log2FC=0.106 padj=0.928 ``` \| [seedLoc = 393,397] Seq=GTTGAAGCATTTTTCACTACACTCC \| ```          C   A     CT   T   AAGTTG mRNA  3'  CTC CATCA  TTT ACG       miRNA 5'  GAG GTAGT  AGG TGT                T              T   ATAGTT           3111511111551223112333333 ``` \| SeedType = 2 NonSeedType = -1 \| \| --- \| --- \| --- \| --- \| --- \| --- \| | \| 517 \| WBGene00008558:F07H5.7(5088) \| ``` log2FC=-0.138 padj=0.966 ``` \| [seedLoc = 30,35] Seq=TAGTATCCAAATTATTACCTCT \| ```          T          A   C   G T mRNA  3'  CTCCATTATT AAC TAT A  miRNA 5'  GAGGTAGTAG TTG ATA T           T          G   T   G T           3111111211231113111313 ``` \| SeedType = 1 NonSeedType = 1 \| \| --- \| --- \| --- \| --- \| --- \| --- \| | \| 518 \| WBGene00008563:F07H5.13(5092) \| ``` log2FC=0.1 padj=1.0 ``` \| [seedLoc = 134,139] Seq=GAGCATGTTGATACTCCTACCTTT \| ```          T       C  AT GT   A   G mRNA  3'  TTCCATC TC  A  TGT CGA  miRNA 5'  GAGGTAG AG  T  ATA GTT           T       T  GT GT                   321111113113313312151214 ``` \| SeedType = 3 NonSeedType = -1 \| \| --- \| --- \| --- \| --- \| --- \| --- \| | \| 519 \| WBGene00008599:F09A5.2(5121) \| ``` log2FC=0.321 padj=0.832 ``` \| [seedLoc = 55,60] Seq=TGATTATTTTTCAAGCTACTTCT \| ```          T        AA TTTT      T mRNA  3'  CTTCATCG  C    TATTAG  miRNA 5'  GAGGTAGT  G    ATAGTT           T        AG TTGT                  31121111233133331112125 ``` \| SeedType = 3 NonSeedType = -1 \| \| --- \| --- \| --- \| --- \| --- \| --- \| | \| 520 \| WBGene00008601:mob-2(5122) \|  \| [seedLoc = 395,401] Seq=TTTGAAATGATAACTACTCTCA \| ```              T     A      AAGTTT mRNA  3' ACTC CATCA  TAGTA       miRNA 5' TGAG GTAGT  GTTGT                          AG     ATAGTT           11115111113621221333333 ``` \| SeedType = 2 NonSeedType = -1 \| \| --- \| --- \| --- \| --- \| --- \| --- \| | \| 521 \| WBGene00008640:F10B5.3(5147) \| ``` log2FC=0.549 padj=1.0 ``` \| [seedLoc = 35,40] Seq=ACCTATATTTTACTACTTCA \| ```                              CA mRNA  3' ACTTCATCATTT  TATATC   miRNA 5' TGAGGTAGTAGG  GTATAG                        TT      TT           1112111111226621111133 ``` \| SeedType = 3 NonSeedType = -1 \| \| --- \| --- \| --- \| --- \| --- \| --- \| | \| 522 \| WBGene00008682:lex-1(5176) \| ``` log2FC=0.09 padj=0.927 ``` \| [seedLoc = 27,32] Seq=TCCTCCTCTTCCAATCTACCTCC \| ```          C       TAA  TT TCC  CT mRNA  3'  CTCCATC   CC  C   TC   miRNA 5'  GAGGTAG   GG  G   AG            T       TA   TT TAT  TT           31111111335113313331133 ``` \| SeedType = 1 NonSeedType = -1 \| \| --- \| --- \| --- \| --- \| --- \| --- \| | \|  \|  \|  \| [seedLoc = 243,248] Seq=CGAGTGAGCATTGCCTGCCTCT \| ```          T       CG      A  G  C mRNA  3'  CTCCGTC  TT ACG GT AG  miRNA 5'  GAGGTAG  GG TGT TA TT           T       TA  T   A  G              31111211332261123213125 ``` \| SeedType = 2 NonSeedType = -1 \| \| --- \| --- \| --- \| --- \| --- \| --- \| | \| 523 \| WBGene00008684:mig-32(5178) \| ``` log2FC=0.199 padj=0.912 ``` \| [seedLoc = 88,93] Seq=TACTGTACCATGTATCTCTATCTCT \| ```          T       TCT  G  C       T mRNA  3'  CTCTATC   AT TA CATGTCA  miRNA 5'  GAGGTAG   TA GT GTATAGT           T            G  T       T           3111211155511321311121113 ``` \| SeedType = 2 NonSeedType = -1 \| \| --- \| --- \| --- \| --- \| --- \| --- \| | \| 524 \| WBGene00008685:mon-2(5179) \| ``` log2FC=0.073 padj=0.943 ``` \| [seedLoc = 179,185] Seq=AGTTTCGAGTACTAGTACCCTCA \| ```             C    G   ATGAGCTTTGA mRNA  3' ACT CCAT ATC            miRNA 5' TGA GGTA TAG                             G   GTTGTATAGTT           11151111311133333333333 ``` \| SeedType = 3 NonSeedType = -1 \| \| --- \| --- \| --- \| --- \| --- \| --- \| | \| 525 \| WBGene00008687:F11A10.6(5181) \| ``` log2FC=0.007 padj=0.997 ``` \| [seedLoc = 68,74] Seq=CTCTAGTTTCGACTACTCTCG \| ```          G   T     G  TTG     TC mRNA  3'  CTC CATCA CT     ATC   miRNA 5'  GAG GTAGT GG     TAG            T         A  TTGTA   TT           31115111113123336611133 ``` \| SeedType = 2 NonSeedType = -1 \| \| --- \| --- \| --- \| --- \| --- \| --- \| | \| 526 \| WBGene00008693:F11C1.4(5186) \| ``` log2FC=-0.434 padj=1.0 ``` \| [seedLoc = 40,45] Seq=TTTTTGCCAGCTTCACTCTACTTCA \| ```                  TCAC      CGTTTTT mRNA  3' ACTTCATC    TTCGAC        miRNA 5' TGAGGTAG    AGGTTG                         T         TATAGTT           1112111135551212113333333 ``` \| SeedType = 3 NonSeedType = 2 \| \| --- \| --- \| --- \| --- \| --- \| --- \| | \| 527 \| WBGene00008694:vwa-8(5188) \|  \| [seedLoc = 25,31] Seq=TTTTGTTTTAGCTGTTACCCTCA \| ```             C        G TTT    TT mRNA  3' ACT CCATTGTC A   TGTT   miRNA 5' TGA GGTAGTAG T   ATAG                        G TGT    TT           11151111221131333121233 ``` \| SeedType = 3 NonSeedType = -1 \| \| --- \| --- \| --- \| --- \| --- \| --- \| | \| 528 \| WBGene00008695:F11C3.1(5190) \| ``` log2FC=-0.709 padj=1.0 ``` \| [seedLoc = 57,62] Seq=CTCTGGGCATATTTTATTTACCTCA \| ```                T      TAT   G   TC mRNA  3' ACTCCA TTATTT   ACG GTC   miRNA 5' TGAGGT AGTAGG   TGT TAG                         T     A   TT           1111115121122355112321133 ``` \| SeedType = 2 NonSeedType = -1 \| \| --- \| --- \| --- \| --- \| --- \| --- \| | \| 529 \| WBGene00008760:F13E9.11(5224) \| ``` log2FC=-0.406 padj=0.874 ``` \| [seedLoc = 82,87] Seq=TGCTTTCATAAAGTTTACCTCA \| ```                  TGA     CTT   T mRNA  3' ACTCCATT    AATA   TCG  miRNA 5' TGAGGTAG    TTGT   AGT                   TAGG    AT    T           11111112333611213351123 ``` \| SeedType = 1 NonSeedType = -1 \| \| --- \| --- \| --- \| --- \| --- \| --- \| | \| 530 \| WBGene00008780:hxk-1(5241) \|  \| [seedLoc = 241,247] Seq=AAACGTATAAACCACTAACCTCC \| ```          C    A              G   A mRNA  3'  CTCC ATCA CCAA ATAT CAA  miRNA 5'  GAGG TAGT GGTT TATA GTT           T         A    G                    3111151111611116111141115 ``` \| SeedType = 2 NonSeedType = 1 \| \| --- \| --- \| --- \| --- \| --- \| --- \| | \| 531 \| WBGene00008807:F14F3.4(5261) \| ``` log2FC=-0.647 padj=1.0 ``` \| [seedLoc = 55,60] Seq=GGATTGTACATATACCTCA \| ```                 AT             G mRNA  3' ACTCCAT      ACATGTTAG  miRNA 5' TGAGGTA      TGTATAGTT                  GTAGGT                     11111113366661111212124 ``` \| SeedType = 1 NonSeedType = -1 \| \| --- \| --- \| --- \| --- \| --- \| --- \| | \| 532 \| WBGene00008834:F14H8.4(5276) \| ``` log2FC=-1.708 padj=1.0 ``` \| [seedLoc = 77,82] Seq=TTAAATAGCACCTTCACTATCTCT \| ```          T        CT    CG   AATT mRNA  3'  CTCTATCA  TCCA  ATA     miRNA 5'  GAGGTAGT  AGGT  TAT              T              TG   AGTT           311121111551111331113333 ``` \| SeedType = 2 NonSeedType = 1 \| \| --- \| --- \| --- \| --- \| --- \| --- \| | \| 533 \| WBGene00008862:F15D4.5(5294) \| ``` log2FC=0.429 padj=1.0 ``` \| [seedLoc = 23,29] Seq=ATACCTCCTACCATCAG \| ```          G   A     C    TA        mRNA  3'  ACT CCATC TCCA          miRNA 5'  TGA GGTAG AGGT                             T    TGTATAGTT           411151111131111336666666 ``` \| SeedType = 3 NonSeedType = 1 \| \| --- \| --- \| --- \| --- \| --- \| --- \| | \|  \|  \|  \| [seedLoc = 36,42] Seq=TCCAATACCATCTCTTCGTACCATCA \| ```             A    GCTTC    C    ACCT mRNA  3' ACT CCAT     TCTA CATA     miRNA 5' TGA GGTA     AGGT GTAT                      GT       T    AGTT           11151111335551121311113333 ``` \| SeedType = 3 NonSeedType = 2 \| \| --- \| --- \| --- \| --- \| --- \| --- \| | \|  \|  \|  \| [seedLoc = 51,57] Seq=ACCATCACTCCTATTACCATCG \| ```          G  A         T   CTACCA mRNA  3'  CT CCATTATCC  CA       miRNA 5'  GA GGTAGTAGG  GT                T            TT  ATAGTT           31151111211113611333333 ``` \| SeedType = 3 NonSeedType = -1 \| \| --- \| --- \| --- \| --- \| --- \| --- \| | \|  \|  \|  \| [seedLoc = 86,92] Seq=CGCTAATTACCTTGATATACCCTCA \| ```             C    ATAGT    T       C mRNA  3' ACT CCAT     TCCA TA ATCG  miRNA 5' TGA GGTA     AGGT GT TAGT                   GT       T  A    T           11151111335451111321611123 ``` \| SeedType = 3 NonSeedType = 1 \| \| --- \| --- \| --- \| --- \| --- \| --- \| | \| 534 \| WBGene00008880:F16B12.4(5304) \| ``` log2FC=-0.331 padj=0.734 ``` \| [seedLoc = 167,172] Seq=CATATTTTCATTTATCTACTTCT \| ```          T       T     CTTTTATAC mRNA  3'  CTTCATC ATTTA          miRNA 5'  GAGGTAG TAGGT                   T             TGTATAGTT           31121111511221333333333 ``` \| SeedType = 3 NonSeedType = 2 \| \| --- \| --- \| --- \| --- \| --- \| --- \| | \| 535 \| WBGene00008900:F16H6.10(5313) \| ``` log2FC=0.055 padj=0.977 ``` \| [seedLoc = 45,51] Seq=AACAACAACTCTTAAAATTTTTATACTCTCC \| ```          C   T   ATTTTTAAAAT  T     ACAA   mRNA  3'  CTC CAT           TC CAACA       miRNA 5'  GAG GTA           AG GTTGT                T       GT                 ATAGTT           311151113355555555511511111333366 ``` \| SeedType = 2 NonSeedType = 1 \| \| --- \| --- \| --- \| --- \| --- \| --- \| | \|  \|  \|  \| [seedLoc = 233,239] Seq=GGGGTGTCAGCTAAATACCGTCT \| ```          T  G    AAA         G  G mRNA  3'  CT CCAT   TCGAC TGT GG  miRNA 5'  GA GGTA   GGTTG ATA TT           T       GTA     T   G              311411113332121161213224 ``` \| SeedType = 3 NonSeedType = 2 \| \| --- \| --- \| --- \| --- \| --- \| --- \| | \| 536 \| WBGene00008904:F17A2.13(5314) \| ``` log2FC=0.566 padj=1.0 ``` \| [seedLoc = 47,52] Seq=TAGCTATTCCACCGCCTACTTCT \| ```          T       CG   C T      T mRNA  3'  CTTCATC  CCA C TATCGA  miRNA 5'  GAGGTAG  GGT G ATAGTT           T       TA   T T                  31121111331113131111215 ``` \| SeedType = 3 NonSeedType = 1 \| \| --- \| --- \| --- \| --- \| --- \| --- \| | \|  \|  \|  \| [seedLoc = 87,91] Seq=GAAGTTTTAATGTCTACCTGAA \| ```          A G        G    TTTGAAG mRNA  3'  A TCCATC T TAAT        miRNA 5'  T AGGTAG A GTTG                   G      T G    TATAGTT           51311111161321123333333 ``` \| SeedType = 3 NonSeedType = -1 \| \| --- \| --- \| --- \| --- \| --- \| --- \| | \| 537 \| WBGene00008905:F17B5.1(5315) \| ``` log2FC=-0.552 padj=0.524 ``` \| [seedLoc = 112,117] Seq=GATCAGTTTTACTAATCTGCCTCT \| ```          T       TA   ATTTTGACTAG mRNA  3'  CTCCGTC  ATC            miRNA 5'  GAGGTAG  TAG                     T            GTTGTATAGTT           311112115511133333333333 ``` \| SeedType = 2 NonSeedType = -1 \| \| --- \| --- \| --- \| --- \| --- \| --- \| | \| 538 \| WBGene00008914:F17C11.2(5319) \| ``` log2FC=0.158 padj=0.945 ``` \| [seedLoc = 270,276] Seq=GCAATACGTCCGGAATAAAGTTATACACTCG \| ```          G   A   ATTGAAATAAGG  T     A    mRNA  3'  CTC CAT            CC GCATA CG  miRNA 5'  GAG GTA            GG TGTAT GT           T       GTA           T     A  T           31115111333455555544113211113126 ``` \| SeedType = 2 NonSeedType = -1 \| \| --- \| --- \| --- \| --- \| --- \| --- \| | \| 539 \| WBGene00008924:F17E5.2(5329) \| ``` log2FC=-0.089 padj=0.973 ``` \| [seedLoc = 286,291] Seq=TCTCAATGTTTCACCTACCTTT \| ```          T       CACTTTGTAACTCT mRNA  3'  TTCCATC               miRNA 5'  GAGGTAG                        T       TAGGTTGTATAGTT           3211111133333333333333 ``` \| SeedType = 3 NonSeedType = -1 \| \| --- \| --- \| --- \| --- \| --- \| --- \| | \| 540 \| WBGene00008951:F19C6.2(5345) \| ``` log2FC=-0.183 padj=0.787 ``` \| [seedLoc = 185,190] Seq=ACTATGCAATTTCAAATTTCAAGATACCTCT \| ```          T      AGAACTTTAAAC              mRNA  3'  CTCCAT            TTTAACGTATCA  miRNA 5'  GAGGTA            AGGTTGTATAGT           T      GT                      T           31111113355555555551221112111116 ``` \| SeedType = 1 NonSeedType = 1 \| \| --- \| --- \| --- \| --- \| --- \| --- \| | \|  \|  \|  \| [seedLoc = 220,225] Seq=TCTTTCAAAAATTACTACTTCT \| ```          T          AAAAACTTTCT mRNA  3'  CTTCATCATT            miRNA 5'  GAGGTAGTAG                     T          GTTGTATAGTT           3112111111233333333333 ``` \| SeedType = 3 NonSeedType = -1 \| \| --- \| --- \| --- \| --- \| --- \| --- \| | \| 541 \| WBGene00008956:nekl-3(5349) \| ``` log2FC=0.157 padj=0.882 ``` \| [seedLoc = 114,120] Seq=TGATGTGCTAAATTTTACTCTCA \| ```              T    TTAAAT     AGT mRNA  3' ACTC CATT      CGTGT    miRNA 5' TGAG GTAG      GTATA                      TAGGTT     GTT           11115111233333312121333 ``` \| SeedType = 2 NonSeedType = -1 \| \| --- \| --- \| --- \| --- \| --- \| --- \| | \| 542 \| WBGene00008971:F20C5.6(5359) \| ``` log2FC=-0.504 padj=0.745 ``` \| [seedLoc = 331,337] Seq=TCAGGGATATTCTGTTTCTCACCTCA \| ```                C  TTT    T   GGGACT mRNA  3' ACTCCA TC   GTCT ATA       miRNA 5' TGAGGT AG   TAGG TGT                                T   ATAGTT           11111151155521123121333333 ``` \| SeedType = 2 NonSeedType = -1 \| \| --- \| --- \| --- \| --- \| --- \| --- \| | \| 543 \| WBGene00008973:F20D1.1(5360) \| ``` log2FC=0.125 padj=0.943 ``` \| [seedLoc = 245,250] Seq=TTTCCTTTTTTCTCACTATCTCT \| ```          T        C   TTTTTCCTTT mRNA  3'  CTCTATCA TCT           miRNA 5'  GAGGTAGT AGG                    T            TTGTATAGTT           31112111151123333333333 ``` \| SeedType = 2 NonSeedType = -1 \| \| --- \| --- \| --- \| --- \| --- \| --- \| | \| 544 \| WBGene00008995:prde-1(5374) \| ``` log2FC=0.029 padj=0.986 ``` \| [seedLoc = 24,30] Seq=AAATTCGTCACTTGTACCATCA \| ```             A          CTGCTTAAA mRNA  3' ACT CCAT GTTCA          miRNA 5' TGA GGTA TAGGT                           G     TGTATAGTT           11151111621211333333333 ``` \| SeedType = 3 NonSeedType = 2 \| \| --- \| --- \| --- \| --- \| --- \| --- \| | \| 545 \| WBGene00009012:F21D5.7(5387) \| ``` log2FC=0.119 padj=0.843 ``` \| [seedLoc = 314,319] Seq=AAACATATGCCTGTTTATTTTCTATCTCA \| ```                  TTTTATTT              A mRNA  3' ACTCTATC        GTCC GTATA CAA  miRNA 5' TGAGGTAG        TAGG TGTAT GTT                               T     A               1111211155555555211162211161115 ``` \| SeedType = 2 NonSeedType = -1 \| \| --- \| --- \| --- \| --- \| --- \| --- \| | \| 546 \| WBGene00009026:F21G4.5(5395) \| ``` log2FC=0.532 padj=0.626 ``` \| [seedLoc = 247,253] Seq=CAATCACCAATTTAAAGGTGGACTACGCTCA \| ```              G     GGTGGAAA      CACTAAC mRNA  3' ACTC CATCA        TTTAAC        miRNA 5' TGAG GTAGT        AGGTTG                                         TATAGTT           1111411111445445551221113333333 ``` \| SeedType = 2 NonSeedType = 1 \| \| --- \| --- \| --- \| --- \| --- \| --- \| | \| 547 \| WBGene00009058:F22E12.1(5416) \| ``` log2FC=0.163 padj=0.888 ``` \| [seedLoc = 137,142] Seq=ATTTTAAAAATCTTACCTCT \| ```          T             AAATTTTA mRNA  3'  CTCCAT  TCTAA         miRNA 5'  GAGGTA  AGGTT                  T      GT     GTATAGTT           3111111661121133333333 ``` \| SeedType = 1 NonSeedType = 2 \| \| --- \| --- \| --- \| --- \| --- \| --- \| | \| 548 \| WBGene00009060:F22E12.3(5418) \| ``` log2FC=-0.91 padj=0.693 ``` \| [seedLoc = 143,148] Seq=TTCTGAATAAAGTCTACCTCA \| ```                  TGA     A   TT mRNA  3' ACTCCATC    AATA GTC   miRNA 5' TGAGGTAG    TTGT TAG                    TAGG    A   TT           1111111133361121321133 ``` \| SeedType = 1 NonSeedType = -1 \| \| --- \| --- \| --- \| --- \| --- \| --- \| | \| 549 \| WBGene00009113:maph-1.2(5454) \|  \| [seedLoc = 178,183] Seq=TGCTGTCCACCATACCTCA \| ```                 A     C       T mRNA  3' ACTCCAT   CCA C TGTCG  miRNA 5' TGAGGTA   GGT G ATAGT                  GTA   T T     T           1111111366111316121123 ``` \| SeedType = 1 NonSeedType = 1 \| \| --- \| --- \| --- \| --- \| --- \| --- \| | \| 550 \| WBGene00009126:pyk-1(5464) \| ``` log2FC=0.325 padj=0.096 ``` \| [seedLoc = 24,30] Seq=CCGTCAGCATACTCGTTTACTCTCA \| ```              T    TGC  AT   ACTGCC mRNA  3' ACTC CATT   TC  ACG       miRNA 5' TGAG GTAG   AG  TGT                         T    GT   ATAGTT           1111511123451133112333333 ``` \| SeedType = 2 NonSeedType = -1 \| \| --- \| --- \| --- \| --- \| --- \| --- \| | \| 551 \| WBGene00009133:bed-3(5470) \| ``` log2FC=-0.384 padj=0.643 ``` \| [seedLoc = 150,156] Seq=CAATTATGTGAAGCTTTACTACCATCA \| ```             A      T     AGT       C mRNA  3' ACT CCATCA TTCGA   GTATTAA  miRNA 5' TGA GGTAGT AGGTT   TATAGTT                           G                     111511111151212134521112115 ``` \| SeedType = 3 NonSeedType = 2 \| \| --- \| --- \| --- \| --- \| --- \| --- \| | \|  \|  \|  \| [seedLoc = 307,313] Seq=TATTGTCATCCTTTTCACCTCT \| ```          T     C  T   T        T mRNA  3'  CTCCA TT TCC AC TGTTA  miRNA 5'  GAGGT AG AGG TG ATAGT           T        T   T  T     T           31111151231113116121213 ``` \| SeedType = 2 NonSeedType = -1 \| \| --- \| --- \| --- \| --- \| --- \| --- \| | \| 552 \| WBGene00009162:F26E4.7(5489) \| ``` log2FC=-0.45 padj=0.426 ``` \| [seedLoc = 162,167] Seq=TTTTTGAAAACATTCTATCTCG \| ```          G       TTA   AAGTTTTT mRNA  3'  CTCTATC   CAA         miRNA 5'  GAGGTAG   GTT                  T       TAG   GTATAGTT           3111211133311133333333 ``` \| SeedType = 2 NonSeedType = 1 \| \| --- \| --- \| --- \| --- \| --- \| --- \| | \| 553 \| WBGene00009190:F27D4.6(5514) \| ``` log2FC=0.007 padj=0.997 ``` \| [seedLoc = 279,285] Seq=TCATTTCTAATTGAGTACCATCT \| ```          T  A    G G     CTTTACT mRNA  3'  CT CCAT A TTAAT        miRNA 5'  GA GGTA T GGTTG                 T       G A     TATAGTT           31151111313221123333333 ``` \| SeedType = 3 NonSeedType = 2 \| \| --- \| --- \| --- \| --- \| --- \| --- \| | \| 554 \| WBGene00009201:F28C1.3(5522) \| ``` log2FC=0.109 padj=0.945 ``` \| [seedLoc = 674,679] Seq=CCCCGTTTTACCTGAAATTACCTCC \| ```          C        AAG    TTTTGCCCC mRNA  3'  CTCCATTA   TCCA          miRNA 5'  GAGGTAGT   AGGT                   T               TGTATAGTT           3111111215541111333333333 ``` \| SeedType = 1 NonSeedType = 1 \| \| --- \| --- \| --- \| --- \| --- \| --- \| | \| 555 \| WBGene00009211:wdr-46(5529) \| ``` log2FC=0.016 padj=0.993 ``` \| [seedLoc = 191,194] Seq=TCGTAATCTTCCTCTTCTCCCTCC \| ```          C    C  TTC   TT TAATGCT mRNA  3'  CTCC TC   TCC  C        miRNA 5'  GAGG AG   AGG  G                 T    T  T     TT TATAGTT           311113113551113313333333 ``` \| SeedType = 2 NonSeedType = -1 \| \| --- \| --- \| --- \| --- \| --- \| --- \| | \| 556 \| WBGene00009234:F28H6.6(5545) \| ``` log2FC=0.031 padj=0.993 ``` \| [seedLoc = 63,69] Seq=TTCTATGTTCTCTTTTACTCTCT \| ```          T   T    T   CTT     TT mRNA  3'  CTC CATT TCT   GTATC   miRNA 5'  GAG GTAG AGG   TATAG            T        T   TTG     TT           31115111231123332111133 ``` \| SeedType = 2 NonSeedType = -1 \| \| --- \| --- \| --- \| --- \| --- \| --- \| | \| 557 \| WBGene00009261:F30A10.2(5560) \| ``` log2FC=-0.029 padj=0.993 ``` \| [seedLoc = 141,146] Seq=GAAAAAGTTTTCTGTACCTCG \| ```          G           TTTGAAAAAG mRNA  3'  CTCCAT GTCT           miRNA 5'  GAGGTA TAGG                    T      G    TTGTATAGTT           3111111621123333333333 ``` \| SeedType = 1 NonSeedType = -1 \| \| --- \| --- \| --- \| --- \| --- \| --- \| | \| 558 \| WBGene00009280:F31B9.3(5573) \| ``` log2FC=0.695 padj=0.252 ``` \| [seedLoc = 313,318] Seq=CCTTTCATTATCTCTACCTTT \| ```          T            TTACTTTCC mRNA  3'  TTCCATC TCTA          miRNA 5'  GAGGTAG AGGT                   T       T    TGTATAGTT           3211111161121333333333 ``` \| SeedType = 3 NonSeedType = 2 \| \| --- \| --- \| --- \| --- \| --- \| --- \| | \| 559 \| WBGene00009289:exos-7(5579) \| ``` log2FC=0.003 padj=0.998 ``` \| [seedLoc = 43,48] Seq=TTTTTTTGAATTGTTACCTCT \| ```          T       G     GTTTTTTT mRNA  3'  CTCCATT  TTAA         miRNA 5'  GAGGTAG  GGTT                  T       TA    GTATAGTT           3111111236221133333333 ``` \| SeedType = 1 NonSeedType = 2 \| \| --- \| --- \| --- \| --- \| --- \| --- \| | \| 560 \| WBGene00009290:F31D4.2(5580) \| ``` log2FC=0.331 padj=0.375 ``` \| [seedLoc = 483,488] Seq=CCCATTGCGCTCCTACCTCA \| ```                  C        TACCC mRNA  3' ACTCCATC  TC GCGT      miRNA 5' TGAGGTAG  GG TGTA                       TA  T    TAGTT           1111111136216212133333 ``` \| SeedType = 1 NonSeedType = -1 \| \| --- \| --- \| --- \| --- \| --- \| --- \| | \| 561 \| WBGene00009299:F31E9.6(5585) \| ``` log2FC=0.495 padj=0.902 ``` \| [seedLoc = 37,42] Seq=CTCAATATTTGATCTACCTCT \| ```          T       TAGTT     ACTC mRNA  3'  CTCCATC      TATA     miRNA 5'  GAGGTAG      GTAT              T       TAGGTT    AGTT           3111111133333621113333 ``` \| SeedType = 1 NonSeedType = -1 \| \| --- \| --- \| --- \| --- \| --- \| --- \| | \| 562 \| WBGene00009306:maph-1.1(5588) \|  \| [seedLoc = 48,54] Seq=TCCTGTGAATTTTTACACTCT \| ```          T   A                CT mRNA  3'  CTC CATT TTTAA GTGTC   miRNA 5'  GAG GTAG AGGTT TATAG            T        T     G     TT           31115111261221162121133 ``` \| SeedType = 2 NonSeedType = 2 \| \| --- \| --- \| --- \| --- \| --- \| --- \| | \| 563 \| WBGene00009385:sas-5(5645) \| ``` log2FC=0.331 padj=0.748 ``` \| [seedLoc = 469,474] Seq=TTTTGAACTATCACTACCTTT \| ```          T            T  AGTTTT mRNA  3'  TTCCATCA CTA CA       miRNA 5'  GAGGTAGT GGT GT                T        A   T  ATAGTT           3211111116121311333333 ``` \| SeedType = 3 NonSeedType = 2 \| \| --- \| --- \| --- \| --- \| --- \| --- \| | \| 564 \| WBGene00009433:F35E12.9(5674) \| ``` log2FC=0.014 padj=0.997 ``` \| [seedLoc = 553,558] Seq=TGATTTTCAATTGTATTACCTCT \| ```          T         GT    TTTTAGT mRNA  3'  CTCCATTAT  TAAC        miRNA 5'  GAGGTAGTA  GTTG                 T         G     TATAGTT           31111112113521113333333 ``` \| SeedType = 1 NonSeedType = 1 \| \| --- \| --- \| --- \| --- \| --- \| --- \| | \| 565 \| WBGene00009435:F35G2.1(5676) \| ``` log2FC=-0.432 padj=0.08 ``` \| [seedLoc = 143,148] Seq=CAAGAAACGACTTTTTTACCTCG \| ```          G       TT       AAGAAC mRNA  3'  CTCCATT  TTCAGCA       miRNA 5'  GAGGTAG  AGGTTGT                T       T        ATAGTT           31111112351211211333333 ``` \| SeedType = 1 NonSeedType = 2 \| \| --- \| --- \| --- \| --- \| --- \| --- \| | \| 566 \| WBGene00009436:marb-1(5677) \| ``` log2FC=0.114 padj=0.861 ``` \| [seedLoc = 49,54] Seq=TAGATTTGTGTTTAGTACCTCA \| ```                 G    GTGTTTAGAT mRNA  3' ACTCCAT ATTT           miRNA 5' TGAGGTA TAGG                           G    TTGTATAGTT           1111111311223333333333 ``` \| SeedType = 1 NonSeedType = -1 \| \| --- \| --- \| --- \| --- \| --- \| --- \| | \| 567 \| WBGene00009440:idhg-1(5680) \| ``` log2FC=0.011 padj=0.994 ``` \| [seedLoc = 195,199] Seq=TTATCTTGTTCCTTTCTTCCTCAT \| ```          T     T  TT   TTGTTCTATT mRNA  3'  ACTCC TC  TCC           miRNA 5'  TGAGG AG  AGG                          T  T    TTGTATAGTT           511111311351113333333333 ``` \| SeedType = 2 NonSeedType = -1 \| \| --- \| --- \| --- \| --- \| --- \| --- \| | \| 568 \| WBGene00009451:cids-2(5691) \| ``` log2FC=-0.043 padj=0.977 ``` \| [seedLoc = 222,227] Seq=CTATGTAATCACCTGGGTTATATCTACCTCT \| ```          T       TATATTGG     CTA          mRNA  3'  CTCCATC        GTCCA   ATGTATC   miRNA 5'  GAGGTAG        TAGGT   TGTATAG            T                              TT           311111115555554421111555122111166 ``` \| SeedType = 1 NonSeedType = 1 \| \| --- \| --- \| --- \| --- \| --- \| --- \| | \| 569 \| WBGene00009465:F36D1.6(5703) \| ``` log2FC=-0.135 padj=0.974 ``` \| [seedLoc = 68,73] Seq=GGACGTGTCGCTTCCAAATAACTACCTTT \| ```          T        ATAAA  TT  C   G   G mRNA  3'  TTCCATCA     CC  CG TGT CAG  miRNA 5'  GAGGTAGT     GG  GT ATA GTT           T        A      TT                      32111111135555113312512141124 ``` \| SeedType = 3 NonSeedType = -1 \| \| --- \| --- \| --- \| --- \| --- \| --- \| | \| 570 \| WBGene00009479:fcp-1(5710) \| ``` log2FC=-0.215 padj=0.616 ``` \| [seedLoc = 136,141] Seq=TTCTGTACTATTTACCTCC \| ```          C       TAT         TT mRNA  3'  CTCCATT      CATGTC   miRNA 5'  GAGGTAG      GTATAG            T       TAGGTT      TT           3111111233366611121133 ``` \| SeedType = 1 NonSeedType = -1 \| \| --- \| --- \| --- \| --- \| --- \| --- \| | \|  \|  \|  \| [seedLoc = 366,371] Seq=TCGATTGCTTACTTAAATACCTCT \| ```          T      AA     TT   TAGCT mRNA  3'  CTCCAT  ATTCA  CGT      miRNA 5'  GAGGTA  TAGGT  GTA               T      G      T    TAGTT           311111135112113512133333 ``` \| SeedType = 1 NonSeedType = 2 \| \| --- \| --- \| --- \| --- \| --- \| --- \| | \| 571 \| WBGene00009503:F37A8.5(5725) \| ``` log2FC=-0.5 padj=0.181 ``` \| [seedLoc = 99,105] Seq=TGAGATGTCGGCCGCAACTACCCTCT \| ```          T  C      ACG     T   GAGT mRNA  3'  CT CCATCA   CCGGC GTA     miRNA 5'  GA GGTAGT   GGTTG TAT              T         A           AGTT           31151111113541122152113333 ``` \| SeedType = 3 NonSeedType = 2 \| \| --- \| --- \| --- \| --- \| --- \| --- \| | \|  \|  \|  \| [seedLoc = 553,559] Seq=TTCCAATTTTTCTATTATCCTCC \| ```          C    T       TTTTAACCTT mRNA  3'  CTCC ATTATCT           miRNA 5'  GAGG TAGTAGG                    T            TTGTATAGTT           31111511211123333333333 ``` \| SeedType = 2 NonSeedType = -1 \| \| --- \| --- \| --- \| --- \| --- \| --- \| | \| 572 \| WBGene00009508:row-1(5729) \|  \| [seedLoc = 70,75] Seq=CGGTTAAAAACTCTACTTCA \| ```                        AAATTGGC mRNA  3' ACTTCATC  TCAA         miRNA 5' TGAGGTAG  GGTT                          TA    GTATAGTT           1112111166211133333333 ``` \| SeedType = 3 NonSeedType = 1 \| \| --- \| --- \| --- \| --- \| --- \| --- \| | \| 573 \| WBGene00009532:ccch-1(5742) \| ``` log2FC=-0.595 padj=0.003 ``` \| [seedLoc = 172,177] Seq=CTTCCCACTTCCCCTACTTCA \| ```                  C   TT  CCCTTC mRNA  3' ACTTCATC  CC  CA       miRNA 5' TGAGGTAG  GG  GT                        TA  TT  ATAGTT           1112111136113311333333 ``` \| SeedType = 3 NonSeedType = -1 \| \| --- \| --- \| --- \| --- \| --- \| --- \| | \| 574 \| WBGene00009559:mtx-1(5761) \| ``` log2FC=0.128 padj=0.874 ``` \| [seedLoc = 49,55] Seq=TTTTTTGCTTGCCCCCTACCTAC \| ```          CA      CC   TT   TTTTT mRNA  3'   TCCATC  CCG  CGT      miRNA 5'   AGGTAG  GGT  GTA               TG      TA   T    TAGTT           33111111331123512133333 ``` \| SeedType = 3 NonSeedType = 2 \| \| --- \| --- \| --- \| --- \| --- \| --- \| | \| 575 \| WBGene00009560:psa-3(5762) \| ``` log2FC=-0.171 padj=0.976 ``` \| [seedLoc = 26,31] Seq=TATTGTAAACAATTTCTACCTCA \| ```                          AA     T mRNA  3' ACTCCATC TTTAACA  TGTTA  miRNA 5' TGAGGTAG AGGTTGT  ATAGT                   T              T           111111116122111155121213 ``` \| SeedType = 1 NonSeedType = 1 \| \| --- \| --- \| --- \| --- \| --- \| --- \| | \| 576 \| WBGene00009561:F39D8.3(5763) \| ``` log2FC=-0.541 padj=0.581 ``` \| [seedLoc = 27,32] Seq=CACTTTTCGAGTTTCCCTAAAATATACCTCT \| ```          T      ATAAA    CTTTG   TTT   C mRNA  3'  CTCCAT     ATCC     AGC   TCA  miRNA 5'  GAGGTA     TAGG     TTG   AGT           T      G                TAT   T           3111111355551111555541213331113 ``` \| SeedType = 1 NonSeedType = -1 \| \| --- \| --- \| --- \| --- \| --- \| --- \| | \|  \|  \|  \| [seedLoc = 131,137] Seq=GTCATTCAGATCTTTACTCTCC \| ```          C   T          ACTTACTG mRNA  3'  CTC CATT TCTAG         miRNA 5'  GAG GTAG AGGTT                  T        T     GTATAGTT           31115111261121233333333 ``` \| SeedType = 2 NonSeedType = 2 \| \| --- \| --- \| --- \| --- \| --- \| --- \| | \| 577 \| WBGene00009635:F42F12.3(5807) \| ``` log2FC=0.042 padj=0.99 ``` \| [seedLoc = 40,46] Seq=TTTTATAGTTTTACTCACCTCA \| ```                C      TG      TT mRNA  3' ACTCCA TCATTT   ATATT   miRNA 5' TGAGGT AGTAGG   TATAG                         TTG     TT           11111151111223361111233 ``` \| SeedType = 2 NonSeedType = -1 \| \| --- \| --- \| --- \| --- \| --- \| --- \| | \| 578 \| WBGene00009677:F44A6.5(5840) \| ``` log2FC=-0.494 padj=0.081 ``` \| [seedLoc = 126,132] Seq=TATTATTATAACCATTGTTTTTACTCTCA \| ```              T    TTTGTTA      T     T mRNA  3' ACTC CATT       CCAATA TATTA  miRNA 5' TGAG GTAG       GGTTGT ATAGT                    TA                 T           11115111233545551111215111213 ``` \| SeedType = 2 NonSeedType = 1 \| \| --- \| --- \| --- \| --- \| --- \| --- \| | \| 579 \| WBGene00009717:dep-1(5867) \| ``` log2FC=0.111 padj=0.949 ``` \| [seedLoc = 45,50] Seq=TTTTTCTCCGGATACTTACCTCT \| ```          T     T    AGGCCTCTTTTT mRNA  3'  CTCCA TCAT             miRNA 5'  GAGGT AGTA                      T          GGTTGTATAGTT           31111151111333333333333 ``` \| SeedType = 2 NonSeedType = -1 \| \| --- \| --- \| --- \| --- \| --- \| --- \| | \| 580 \| WBGene00009727:lips-1(5873) \| ``` log2FC=0.369 padj=1.0 ``` \| [seedLoc = 137,143] Seq=TATTTCTGAATAGACTACCTAC \| ```          CA       GA   GTCTTTAT mRNA  3'   TCCATCA  TAA         miRNA 5'   AGGTAGT  GTT                  TG       AG   GTATAGTT           3311111113321133333333 ``` \| SeedType = 3 NonSeedType = -1 \| \| --- \| --- \| --- \| --- \| --- \| --- \| | \|  \|  \|  \| [seedLoc = 179,185] Seq=TGTTGTATCTTCCGGTTGTATCCTCA \| ```               T  GTTGG  TT T     GT mRNA  3' ACTCC AT     CC  C ATGTT   miRNA 5' TGAGG TA     GG  G TATAG                    GTA    TT       TT           11111511333441133151121233 ``` \| SeedType = 2 NonSeedType = -1 \| \| --- \| --- \| --- \| --- \| --- \| --- \| | \| 581 \| WBGene00009742:F45H10.5(5886) \| ``` log2FC=-0.152 padj=1.0 ``` \| [seedLoc = 30,35] Seq=TGCTTCTTCTTCTTCTTCTACTTCT \| ```          T       TTC   TT TTCT   T mRNA  3'  CTTCATC   TTC  C    TCG  miRNA 5'  GAGGTAG   AGG  G    AGT           T       T     TT TAT    T           3112111135512133133351123 ``` \| SeedType = 3 NonSeedType = -1 \| \| --- \| --- \| --- \| --- \| --- \| --- \| | \| 582 \| WBGene00009745:cox-19(5889) \|  \| [seedLoc = 149,155] Seq=AAGAGTTTTCACTCTACGCTCA \| ```              G       ACTTTTGAGAA mRNA  3' ACTC CATC TC            miRNA 5' TGAG GTAG AG                              T  GTTGTATAGTT           11114111161133333333333 ``` \| SeedType = 2 NonSeedType = -1 \| \| --- \| --- \| --- \| --- \| --- \| --- \| | \| 583 \| WBGene00009774:F46B6.9(5899) \| ``` log2FC=-0.316 padj=1.0 ``` \| [seedLoc = 38,44] Seq=TGTTATATTATTAATACTCTCA \| ```              T   A   AT       GT mRNA  3' ACTC CAT ATT   TATATT   miRNA 5' TGAG GTA TAG   GTATAG                    G   GTT      TT           11115111311233621111233 ``` \| SeedType = 2 NonSeedType = -1 \| \| --- \| --- \| --- \| --- \| --- \| --- \| | \| 584 \| WBGene00009778:F46C5.1(5902) \| ``` log2FC=-0.456 padj=0.726 ``` \| [seedLoc = 126,132] Seq=CATTTGACAGTTTCTGTACACTCA \| ```              A        TT    GT   C mRNA  3' ACTC CAT GTCT  GACA  TTA  miRNA 5' TGAG GTA TAGG  TTGT  AGT                   G          AT   T           1111511162112552111331213 ``` \| SeedType = 2 NonSeedType = -1 \| \| --- \| --- \| --- \| --- \| --- \| --- \| | \| 585 \| WBGene00009800:rrc-1(5918) \| ``` log2FC=0.231 padj=0.842 ``` \| [seedLoc = 131,136] Seq=TCCTCATATTTTTCTTCTACTTCT \| ```          T       T   TTT    C  CT mRNA  3'  CTTCATC TCT   TATA TC   miRNA 5'  GAGGTAG AGG   GTAT AG            T       T   TT        TT           311211113112335211151133 ``` \| SeedType = 3 NonSeedType = -1 \| \| --- \| --- \| --- \| --- \| --- \| --- \| | \| 586 \| WBGene00009801:ipla-2(5919) \|  \| [seedLoc = 323,329] Seq=GTTCTCTCTTCTTTCTACTCTCT \| ```          T   T    T   TT TCTCTTG mRNA  3'  CTC CATC TTC  C        miRNA 5'  GAG GTAG AGG  G                 T        T   TT TATAGTT           31115111131213313333333 ``` \| SeedType = 2 NonSeedType = -1 \| \| --- \| --- \| --- \| --- \| --- \| --- \| | \| 587 \| WBGene00009803:F47B8.2(5920) \| ``` log2FC=-0.117 padj=0.959 ``` \| [seedLoc = 98,104] Seq=ATCACAATAGCATTAAGAATACACTCG \| ```          G   A   AAGAATTA     ACACTA mRNA  3'  CTC CAT        CGATA       miRNA 5'  GAG GTA        GTTGT                T       GTAG         ATAGTT           311151113333555512121333333 ``` \| SeedType = 2 NonSeedType = -1 \| \| --- \| --- \| --- \| --- \| --- \| --- \| | \| 588 \| WBGene00009804:F47B8.3(5921) \| ``` log2FC=-0.011 padj=0.997 ``` \| [seedLoc = 47,52] Seq=TCATTTACTCCGAATACCTCG \| ```          G      AAG  T    TTACT mRNA  3'  CTCCAT   CC  CAT      miRNA 5'  GAGGTA   GG  GTA               T      GTA  TT   TAGTT           3111111333113611133333 ``` \| SeedType = 1 NonSeedType = -1 \| \| --- \| --- \| --- \| --- \| --- \| --- \| | \| 589 \| WBGene00009834:lsy-27(5942) \| ``` log2FC=0.289 padj=0.899 ``` \| [seedLoc = 98,103] Seq=TATTGTTAATTCTACCTCA \| ```                               T mRNA  3' ACTCCATC  TTAAT TGTTA  miRNA 5' TGAGGTAG  GGTTG ATAGT                   TA     T     T           1111111166221126121213 ``` \| SeedType = 1 NonSeedType = 2 \| \| --- \| --- \| --- \| --- \| --- \| --- \| | \| 590 \| WBGene00009843:F48C5.2(5946) \| ``` log2FC=0.004 padj=0.999 ``` \| [seedLoc = 26,32] Seq=AAATTTACTTCTTATTATCCTCT \| ```          T    T       TT   TTAAA mRNA  3'  CTCC ATTATTC  CAT      miRNA 5'  GAGG TAGTAGG  GTA               T            TT   TAGTT           31111511211213311133333 ``` \| SeedType = 2 NonSeedType = -1 \| \| --- \| --- \| --- \| --- \| --- \| --- \| | \| 591 \| WBGene00009888:F49E2.5(5964) \| ``` log2FC=0.209 padj=0.578 ``` \| [seedLoc = 194,199] Seq=TAATAACTAACCTTTTGCACGAAATACCTCT \| ```          T      AAAGCACGTTT      CAATAAT mRNA  3'  CTCCAT           TCCAAT        miRNA 5'  GAGGTA           AGGTTG                 T      GT               TATAGTT           3111111335455545551111123333333 ``` \| SeedType = 1 NonSeedType = 1 \| \| --- \| --- \| --- \| --- \| --- \| --- \| | \| 592 \| WBGene00009899:efl-3(5972) \| ``` log2FC=-0.625 padj=0.045 ``` \| [seedLoc = 224,230] Seq=TTTTCTGCATCTTCTACACTACCCTCT \| ```          T  C      CATC   T    CTTTT mRNA  3'  CT CCATCA    TTC ACGT      miRNA 5'  GA GGTAGT    AGG TGTA               T                T    TAGTT           311511111155551213112133333 ``` \| SeedType = 3 NonSeedType = -1 \| \| --- \| --- \| --- \| --- \| --- \| --- \| | \| 593 \| WBGene00009921:F52B5.2(5984) \| ``` log2FC=0.109 padj=0.914 ``` \| [seedLoc = 234,239] Seq=GTTTATATTTTTATTATATTACCTCA \| ```                    ATTATTTT      TG mRNA  3' ACTCCATTAT        TATATT   miRNA 5' TGAGGTAGTA        GTATAG                      GGTT          TT           11111112113333555521111233 ``` \| SeedType = 1 NonSeedType = -1 \| \| --- \| --- \| --- \| --- \| --- \| --- \| | \| 594 \| WBGene00009935:F52E10.4(5994) \| ``` log2FC=-0.24 padj=0.896 ``` \| [seedLoc = 72,77] Seq=TCATAGACATTCATTGCTGCCTCC \| ```          C          ACTT   GATACT mRNA  3'  CTCCGTCGTT    ACA       miRNA 5'  GAGGTAGTAG    TGT                T          GT     ATAGTT           311112112123355111333333 ``` \| SeedType = 2 NonSeedType = -1 \| \| --- \| --- \| --- \| --- \| --- \| --- \| | \| 595 \| WBGene00009974:F53C11.4(6023) \| ``` log2FC=-0.329 padj=0.21 ``` \| [seedLoc = 321,324] Seq=TTTCTTGGGATCCTTCTCACTTCCTCC \| ```          C    T   CTCT   T  G  TCTTT mRNA  3'  CTCC TCA    TCC AG GT      miRNA 5'  GAGG AGT    AGG TT TA               T    T             G  TAGTT           311113111555511151232133333 ``` \| SeedType = 2 NonSeedType = -1 \| \| --- \| --- \| --- \| --- \| --- \| --- \| | \|  \|  \|  \| [seedLoc = 327,332] Seq=CACTTCCTCCTCTCTATCTCC \| ```          C       TC   TC T     C mRNA  3'  CTCTATC  TCC  C   TCA  miRNA 5'  GAGGTAG  AGG  G   AGT           T       T    TT TAT   T           31112111351113313661113 ``` \| SeedType = 2 NonSeedType = -1 \| \| --- \| --- \| --- \| --- \| --- \| --- \| | \| 596 \| WBGene00010001:F53F8.4(6042) \| ``` log2FC=-0.046 padj=0.993 ``` \| [seedLoc = 45,50] Seq=TCACATTTAGCCGCTGCATATACCTCA \| ```                 ATA   CG     TTACACT mRNA  3' ACTCCAT   CGT  CCGAT        miRNA 5' TGAGGTA   GTA  GGTTG                                     TATAGTT           111111155512154112123333333 ``` \| SeedType = 1 NonSeedType = 2 \| \| --- \| --- \| --- \| --- \| --- \| --- \| | \| 597 \| WBGene00010051:F54D5.5(6077) \| ``` log2FC=0.702 padj=0.486 ``` \| [seedLoc = 98,103] Seq=TTTTTTTTCATTTCTACCTTA \| ```                       CTTTTTTTT mRNA  3' ATTCCATC TTTA          miRNA 5' TGAGGTAG AGGT                           T    TGTATAGTT           1211111161221333333333 ``` \| SeedType = 3 NonSeedType = 2 \| \| --- \| --- \| --- \| --- \| --- \| --- \| | \| 598 \| WBGene00010075:F55A11.1(6097) \| ``` log2FC=0.218 padj=0.596 ``` \| [seedLoc = 141,147] Seq=AATCATTTTATCATACTATCCTCG \| ```          G    T     A   TTTTACTAA mRNA  3'  CTCC ATCAT CTA          miRNA 5'  GAGG TAGTA GGT                   T              TGTATAGTT           311115111115121333333333 ``` \| SeedType = 2 NonSeedType = 2 \| \| --- \| --- \| --- \| --- \| --- \| --- \| | \| 599 \| WBGene00010097:srpa-68(6114) \| ``` log2FC=0.119 padj=0.856 ``` \| [seedLoc = 64,69] Seq=TACTGTTGTTTCGTTGTCCATAGATACCTCT \| ```          T      AG  A  TGTTGCTTTGT     T mRNA  3'  CTCCAT  AT CC           TGTCA  miRNA 5'  GAGGTA  TA GG           ATAGT           T      G      TTGT            T           3111111341151133334555545121113 ``` \| SeedType = 1 NonSeedType = -1 \| \| --- \| --- \| --- \| --- \| --- \| --- \| | \| 600 \| WBGene00010109:F55C10.5(6118) \| ``` log2FC=0.069 padj=0.966 ``` \| [seedLoc = 118,123] Seq=TTCTGTAATAATATTTTTAAAATACCTCT \| ```          T      AAAATTTTTA    A     TT mRNA  3'  CTCCAT          TAAT ATGTC   miRNA 5'  GAGGTA          GTTG TATAG            T      GTAG                TT           31111113333555555211251121133 ``` \| SeedType = 1 NonSeedType = -1 \| \| --- \| --- \| --- \| --- \| --- \| --- \| | \| 601 \| WBGene00010117:nkb-3(6126) \| ``` log2FC=0.033 padj=0.985 ``` \| [seedLoc = 436,442] Seq=ATCTATTTTCCTTCTTCCTCAG \| ```          G     T  T   TTT     TA mRNA  3'  ACTCC TC TCC    TATC   miRNA 5'  TGAGG AG AGG    ATAG                  T  T   TTGT    TT           41111131131113336111133 ``` \| SeedType = 2 NonSeedType = -1 \| \| --- \| --- \| --- \| --- \| --- \| --- \| | \| 602 \| WBGene00010118:F55F3.4(6127) \| ``` log2FC=0.519 padj=1.0 ``` \| [seedLoc = 1026,1031] Seq=TATTCAGCAGATGCCAGAGTACCTCA \| ```                 G GA   TAG   ACTTAT mRNA  3' ACTCCAT A  CCG   ACG       miRNA 5' TGAGGTA T  GGT   TGT                       G A          ATAGTT           11111113135112554112333333 ``` \| SeedType = 1 NonSeedType = 2 \| \| --- \| --- \| --- \| --- \| --- \| --- \| | \| 603 \| WBGene00010122:F55G11.1(6129) \| ``` log2FC=0.108 padj=1.0 ``` \| [seedLoc = 97,103] Seq=TTTTATATGTTGTTACCATCC \| ```          C  A        G        TT mRNA  3'  CT CCATTGTT   TATATT   miRNA 5'  GA GGTAGTAG   GTATAG            T           GTT      TT           31151111221236621111233 ``` \| SeedType = 3 NonSeedType = -1 \| \| --- \| --- \| --- \| --- \| --- \| --- \| | \| 604 \| WBGene00010144:F56C4.1(6149) \| ``` log2FC=-0.318 padj=1.0 ``` \| [seedLoc = 178,183] Seq=TCCATTGCATTAATATTACCTCA \| ```                    AATT    TACCT mRNA  3' ACTCCATTAT    ACGT      miRNA 5' TGAGGTAGTA    TGTA                         GGT     TAGTT           11111112113335112133333 ``` \| SeedType = 1 NonSeedType = -1 \| \| --- \| --- \| --- \| --- \| --- \| --- \| | \| 605 \| WBGene00010177:F57A8.1(6160) \| ``` log2FC=0.143 padj=0.935 ``` \| [seedLoc = 715,720] Seq=AATTTCAATTTATTTTCATCGCAATACCTCC \| ```          C      AACGCTACTTT        TTTAA   mRNA  3'  CTCCAT           TATTTAAC        miRNA 5'  GAGGTA           GTAGGTTG                 T                         TATAGTT           311111155545555555211221113333366 ``` \| SeedType = 1 NonSeedType = 1 \| \| --- \| --- \| --- \| --- \| --- \| --- \| | \| 606 \| WBGene00010199:bet-2(6176) \| ``` log2FC=0.034 padj=0.988 ``` \| [seedLoc = 1211,1216] Seq=TGATTGAATTAGAATTACCTCG \| ```          G        AGATTAAGTTAGT mRNA  3'  CTCCATTA              miRNA 5'  GAGGTAGT                       T        AGGTTGTATAGTT           3111111213333333333333 ``` \| SeedType = 1 NonSeedType = -1 \| \| --- \| --- \| --- \| --- \| --- \| --- \| | \| 607 \| WBGene00010221:F57G12.1(6184) \| ``` log2FC=0.028 padj=0.988 ``` \| [seedLoc = 1205,1210] Seq=TGATTGTATGTTGTTTCTACCTTT \| ```          T       TT G T         T mRNA  3'  TTCCATC  T T GTATGTTAG  miRNA 5'  GAGGTAG  A G TGTATAGTT           T       T  G T                     321111113513232211212125 ``` \| SeedType = 3 NonSeedType = -1 \| \| --- \| --- \| --- \| --- \| --- \| --- \| | \| 608 \| WBGene00010244:F58D5.5(6204) \| ``` log2FC=-0.205 padj=0.817 ``` \| [seedLoc = 1090,1096] Seq=TCAAATACAATGCACTGCGTACACTCG \| ```          G   A   G    ACG       AACT mRNA  3'  CTC CAT CGTC   TAACATA     miRNA 5'  GAG GTA GTAG   GTTGTAT              T                      AGTT           311151114121155421111113333 ``` \| SeedType = 2 NonSeedType = 1 \| \| --- \| --- \| --- \| --- \| --- \| --- \| | \|  \|  \|  \| [seedLoc = 1594,1599] Seq=TCTCTCTCTCACTCTATCTCT \| ```          T          ACTCTCTCTCT mRNA  3'  CTCTATC TC            miRNA 5'  GAGGTAG AG                     T       T  GTTGTATAGTT           3111211161133333333333 ``` \| SeedType = 2 NonSeedType = -1 \| \| --- \| --- \| --- \| --- \| --- \| --- \| | \|  \|  \|  \| [seedLoc = 2069,2075] Seq=CGCATTTCCACCTCTACCATCT \| ```          T  A          CCTTTACGC mRNA  3'  CT CCATC TCCA          miRNA 5'  GA GGTAG AGGT                   T        T    TGTATAGTT           31151111161111333333333 ``` \| SeedType = 3 NonSeedType = 1 \| \| --- \| --- \| --- \| --- \| --- \| --- \| | \| 609 \| WBGene00010272:amph-1(6222) \| ``` log2FC=0.013 padj=0.993 ``` \| [seedLoc = 429,435] Seq=AAACAATTAGCTTAAGCTACCATCA \| ```             A      AA         A   A mRNA  3' ACT CCATCG  TTCGAT TA CAA  miRNA 5' TGA GGTAGT  AGGTTG AT GTT                             T  A               11151111125512121261131115 ``` \| SeedType = 3 NonSeedType = 2 \| \| --- \| --- \| --- \| --- \| --- \| --- \| | \| 610 \| WBGene00010279:letm-1(6225) \| ``` log2FC=0.057 padj=0.956 ``` \| [seedLoc = 110,116] Seq=TCCCCCGCGCGCTCTACACTCT \| ```          T   A       GC   CCCCCT mRNA  3'  CTC CATC TC  GCG       miRNA 5'  GAG GTAG AG  TGT                T        T  GT   ATAGTT           31115111161133212333333 ``` \| SeedType = 2 NonSeedType = -1 \| \| --- \| --- \| --- \| --- \| --- \| --- \| | \|  \|  \|  \| [seedLoc = 164,170] Seq=TCTCGTACCCGCCAAAAATAGTCACCTCC \| ```          C     C G  AAAAA   CC    CTCT mRNA  3'  CTCCA T AT     CCG  CATG     miRNA 5'  GAGGT A TA     GGT  GTAT              T       G          T     AGTT           31111151311555551123511123333 ``` \| SeedType = 2 NonSeedType = 2 \| \| --- \| --- \| --- \| --- \| --- \| --- \| | \| 611 \| WBGene00010288:F58H1.5(6233) \| ``` log2FC=0.792 padj=0.64 ``` \| [seedLoc = 64,69] Seq=TTTAGTTTTTAATTCTACTTCA \| ```                  TTAATTTTTGATTT mRNA  3' ACTTCATC               miRNA 5' TGAGGTAG                                TAGGTTGTATAGTT           1112111133333333333333 ``` \| SeedType = 3 NonSeedType = -1 \| \| --- \| --- \| --- \| --- \| --- \| --- \| | \| 612 \| WBGene00010315:F59B2.13(6250) \| ``` log2FC=0.159 padj=0.971 ``` \| [seedLoc = 36,41] Seq=TTCTGAAAATCAACTACTTCT \| ```          T        A    AA    TT mRNA  3'  CTTCATCA CTAA   GTC   miRNA 5'  GAGGTAGT GGTT   TAG            T        A    GTA   TT           3112111113121133621133 ``` \| SeedType = 3 NonSeedType = 2 \| \| --- \| --- \| --- \| --- \| --- \| --- \| | \| 613 \| WBGene00010369:chd-1(6295) \| ``` log2FC=0.023 padj=0.989 ``` \| [seedLoc = 33,39] Seq=CTACCTGCACACACTACACTCA \| ```              A       AC    CCATC mRNA  3' ACTC CATCA C  ACGT      miRNA 5' TGAG GTAGT G  TGTA                         A GT    TAGTT           11115111116133112133333 ``` \| SeedType = 2 NonSeedType = -1 \| \| --- \| --- \| --- \| --- \| --- \| --- \| | \| 614 \| WBGene00010396:H13N06.2(6303) \| ``` log2FC=0.847 padj=0.155 ``` \| [seedLoc = 62,65] Seq=TTGTCGTGAGCCTTTCTCTCTTCCTCC \| ```          C    T  TCTCTT     GTGCTGTT mRNA  3'  CTCC TC      TCCGA         miRNA 5'  GAGG AG      AGGTT                  T    T  T          GTATAGTT           311113113555551112133333333 ``` \| SeedType = 2 NonSeedType = 2 \| \| --- \| --- \| --- \| --- \| --- \| --- \| | \| 615 \| WBGene00010407:H19N07.3(6309) \| ``` log2FC=-0.473 padj=0.109 ``` \| [seedLoc = 85,90] Seq=CGCATGCAGCTATCTCC \| ```          C                    C mRNA  3'  CTCTATCG    ACGTA CG  miRNA 5'  GAGGTAGT    TGTAT GT           T        AGGT     A  T           3111211126666112116123 ``` \| SeedType = 2 NonSeedType = -1 \| \| --- \| --- \| --- \| --- \| --- \| --- \| | \| 616 \| WBGene00010418:H27A22.1(6314) \| ``` log2FC=0.241 padj=0.71 ``` \| [seedLoc = 59,63] Seq=CATTCCGCTTCTTCCTCCCTCC \| ```          C    C  C   TT  CCTTAC mRNA  3'  CTCC TC TTC  CG       miRNA 5'  GAGG AG AGG  GT                T    T  T   TT  ATAGTT           3111131131213312333333 ``` \| SeedType = 2 NonSeedType = -1 \| \| --- \| --- \| --- \| --- \| --- \| --- \| | \| 617 \| WBGene00010428:dcn-1(6325) \| ``` log2FC=-0.039 padj=0.975 ``` \| [seedLoc = 321,327] Seq=TACTTTTCGACCATATTACTCTCT \| ```          T   T      A     TTT   T mRNA  3'  CTC CATTAT CCAGC   TCA  miRNA 5'  GAG GTAGTA GGTTG   AGT           T                TAT   T           311151112115111213331113 ``` \| SeedType = 2 NonSeedType = 1 \| \| --- \| --- \| --- \| --- \| --- \| --- \| | \|  \|  \|  \| [seedLoc = 358,363] Seq=TCTTTGACGGGTTGCTGCCTCA \| ```                     G    GTTTCT mRNA  3' ACTCCGTCGTT GGCA       miRNA 5' TGAGGTAGTAG TTGT                           G    ATAGTT           1111121121232211333333 ``` \| SeedType = 2 NonSeedType = -1 \| \| --- \| --- \| --- \| --- \| --- \| --- \| | \|  \|  \|  \| [seedLoc = 371,376] Seq=TGACGGGTTGCTGCCTCAT \| ```          T           G    GT     mRNA  3'  ACTCCGTCGTT GGCA       miRNA 5'  TGAGGTAGTAG TTGT                            G    ATAGTT           51111121121232211336666 ``` \| SeedType = 2 NonSeedType = -1 \| \| --- \| --- \| --- \| --- \| --- \| --- \| | \| 618 \| WBGene00010433:H40L08.1(6326) \| ``` log2FC=0.063 padj=0.979 ``` \| [seedLoc = 184,189] Seq=TATTGTAAAGACCTTATTTATTTCTACCTTA \| ```                  TTTATTTAT     AA      T mRNA  3' ATTCCATC         TCCAG  ATGTTA  miRNA 5' TGAGGTAG         AGGTT  TATAGT                   T             G       T           1211111135555555511112351121213 ``` \| SeedType = 3 NonSeedType = 1 \| \| --- \| --- \| --- \| --- \| --- \| --- \| | \| 619 \| WBGene00010453:fozi-1(6338) \| ``` log2FC=-0.032 padj=0.991 ``` \| [seedLoc = 398,403] Seq=CATGAATCTATCTATCTATCTCT \| ```          T       T     TCTAAGTAC mRNA  3'  CTCTATC ATCTA          miRNA 5'  GAGGTAG TAGGT                   T             TGTATAGTT           31112111511121333333333 ``` \| SeedType = 2 NonSeedType = 2 \| \| --- \| --- \| --- \| --- \| --- \| --- \| | \|  \|  \|  \| [seedLoc = 442,448] Seq=TTTTATCAAAAGTATTACTCTCT \| ```          T   T      GAA        TT mRNA  3'  CTC CATTAT   AAC TATT   miRNA 5'  GAG GTAGTA   TTG ATAG            T          GG    T    TT           311151112113351116111233 ``` \| SeedType = 2 NonSeedType = 1 \| \| --- \| --- \| --- \| --- \| --- \| --- \| | \| 620 \| WBGene00010478:K01G5.5(6357) \| ``` log2FC=-0.156 padj=0.736 ``` \| [seedLoc = 30,35] Seq=TTTTCTTTTTTAAATTACCTCT \| ```          T        AATTTTTTCTTTT mRNA  3'  CTCCATTA              miRNA 5'  GAGGTAGT                       T        AGGTTGTATAGTT           3111111213333333333333 ``` \| SeedType = 1 NonSeedType = -1 \| \| --- \| --- \| --- \| --- \| --- \| --- \| | \| 621 \| WBGene00010507:K02E2.6(6380) \| ``` log2FC=2.238 padj=1.0 ``` \| [seedLoc = 86,92] Seq=ATTTTTGCGGCAAGATACTCTCG \| ```          G   T   AGAA      TTTTA mRNA  3'  CTC CAT    CGGCGT      miRNA 5'  GAG GTA    GTTGTA               T       GTAG      TAGTT           31115111333312212133333 ``` \| SeedType = 2 NonSeedType = -1 \| \| --- \| --- \| --- \| --- \| --- \| --- \| | \| 622 \| WBGene00010519:K03A11.1(6389) \| ``` log2FC=2.174 padj=1.0 ``` \| [seedLoc = 164,169] Seq=CAACCGTATCCCCTATCTATCTCC \| ```          C       T    CC    C   C mRNA  3'  CTCTATC ATCC  TATG CAA  miRNA 5'  GAGGTAG TAGG  GTAT GTT           T            TT    A               311121115111133211231115 ``` \| SeedType = 2 NonSeedType = -1 \| \| --- \| --- \| --- \| --- \| --- \| --- \| | \| 623 \| WBGene00010542:K03H1.7(6398) \| ``` log2FC=-0.028 padj=0.994 ``` \| [seedLoc = 46,51] Seq=AATCATGCATCATAACTGCCTCG \| ```          G        ATA T     CTAA mRNA  3'  CTCCGTCA   C ACGTA     miRNA 5'  GAGGTAGT   G TGTAT              T        AG  T     AGTT           31111211133513112113333 ``` \| SeedType = 2 NonSeedType = -1 \| \| --- \| --- \| --- \| --- \| --- \| --- \| | \| 624 \| WBGene00010550:K04B12.2(6407) \| ``` log2FC=0.164 padj=0.884 ``` \| [seedLoc = 47,52] Seq=CATTTTTTCTCCCTACCTCA \| ```                      TCTTTTTTAC mRNA  3' ACTCCATC  CC           miRNA 5' TGAGGTAG  GG                            TA  TTGTATAGTT           1111111166113333333333 ``` \| SeedType = 1 NonSeedType = -1 \| \| --- \| --- \| --- \| --- \| --- \| --- \| | \|  \|  \|  \| [seedLoc = 57,61] Seq=CATTTTTTCTCCCTACCTCAT \| ```          T            TCTTTTTTAC mRNA  3'  ACTCCATC  CC           miRNA 5'  TGAGGTAG  GG                             TA  TTGTATAGTT           51111111166113333333333 ``` \| SeedType = 1 NonSeedType = -1 \| \| --- \| --- \| --- \| --- \| --- \| --- \| | \| 625 \| WBGene00010552:K04C1.2(6409) \| ``` log2FC=0.01 padj=0.997 ``` \| [seedLoc = 336,341] Seq=ATTGAAATCCCCAAATACCTCA \| ```                 AAA  CCTAAAGTTA mRNA  3' ACTCCAT   CC           miRNA 5' TGAGGTA   GG                           GTA  TTGTATAGTT           1111111333113333333333 ``` \| SeedType = 1 NonSeedType = -1 \| \| --- \| --- \| --- \| --- \| --- \| --- \| | \| 626 \| WBGene00010555:K04C1.5(6412) \| ``` log2FC=0.1 padj=1.0 ``` \| [seedLoc = 145,151] Seq=TTGTACTAATTTATCATTATTTTTACTCTCA \| ```              T   TTTTATTAC       T         mRNA  3' ACTC CAT         TATTTAA CATGTT   miRNA 5' TGAG GTA         GTAGGTT GTATAG                                           TT           111151115555555552112211511121266 ``` \| SeedType = 2 NonSeedType = 2 \| \| --- \| --- \| --- \| --- \| --- \| --- \| | \| 627 \| WBGene00010573:K04H4.2(6429) \| ``` log2FC=-0.804 padj=9.51601951934959E-4 ``` \| [seedLoc = 147,152] Seq=TGATTCTAAGTCTTTCCTATCTCC \| ```          C       CT   TGA  C    T mRNA  3'  CTCTATC  TTC   AT TTAG  miRNA 5'  GAGGTAG  AGG   TA AGTT           T       T    TTG  T                311121113512133311312125 ``` \| SeedType = 2 NonSeedType = -1 \| \| --- \| --- \| --- \| --- \| --- \| --- \| | \| 628 \| WBGene00010592:K06A4.2(6440) \| ``` log2FC=0.121 padj=1.0 ``` \| [seedLoc = 126,131] Seq=TAACTTACTTTACTCCTACTTCA \| ```                  C  ATTT        T mRNA  3' ACTTCATC TC    CAT TCAA  miRNA 5' TGAGGTAG AG    GTA AGTT                   T  GTT    T                111211113113335111611115 ``` \| SeedType = 3 NonSeedType = -1 \| \| --- \| --- \| --- \| --- \| --- \| --- \| | \| 629 \| WBGene00010614:cri-1(6451) \| ``` log2FC=2.034 padj=1.0 ``` \| [seedLoc = 67,72] Seq=TTTCTCACTCACTCTCTACCTCT \| ```          T       TC   CT  CTCTTT mRNA  3'  CTCCATC  TCA  CA       miRNA 5'  GAGGTAG  GGT  GT                T       TA   T   ATAGTT           31111111332113511333333 ``` \| SeedType = 1 NonSeedType = 2 \| \| --- \| --- \| --- \| --- \| --- \| --- \| | \|  \|  \|  \| [seedLoc = 141,146] Seq=CTATGTATGACTATAATTCTACCTCG \| ```          G       TTAATA         ATC mRNA  3'  CTCCATC      TCAGTATGT    miRNA 5'  GAGGTAG      GGTTGTATA             T       TA             GTT           31111111335555211221121333 ``` \| SeedType = 1 NonSeedType = 2 \| \| --- \| --- \| --- \| --- \| --- \| --- \| | \|  \|  \|  \| [seedLoc = 348,353] Seq=CAACTATGCAATGTTCTACCTCA \| ```                  T G           C mRNA  3' ACTCCATC T TAACGTATCAA  miRNA 5' TGAGGTAG A GTTGTATAGTT                   T G                       11111111313211121111115 ``` \| SeedType = 1 NonSeedType = 1 \| \| --- \| --- \| --- \| --- \| --- \| --- \| | \| 630 \| WBGene00010631:cash-1(6468) \| ``` log2FC=0.133 padj=0.811 ``` \| [seedLoc = 639,644] Seq=TCATCATTTTCCTCTACTTCA \| ```                      TTTTACTACT mRNA  3' ACTTCATC TCC           miRNA 5' TGAGGTAG AGG                            T   TTGTATAGTT           1112111161113333333333 ``` \| SeedType = 3 NonSeedType = -1 \| \| --- \| --- \| --- \| --- \| --- \| --- \| | \| 631 \| WBGene00010640:K07F5.16(6475) \| ``` log2FC=0.174 padj=0.97 ``` \| [seedLoc = 33,38] Seq=AAACGCACAATAATTGTTGTTACCTCT \| ```          T          GTTAA     CG    A mRNA  3'  CTCCATTGTT     TAACA   CAA  miRNA 5'  GAGGTAGTAG     GTTGT   GTT           T                    ATA               3111111221245555211113361115 ``` \| SeedType = 1 NonSeedType = 1 \| \| --- \| --- \| --- \| --- \| --- \| --- \| | \| 632 \| WBGene00010641:crml-1(6476) \| ``` log2FC=-0.065 padj=0.959 ``` \| [seedLoc = 158,162] Seq=TTCTCGATTCTTCCTCT \| ```          T    T         TCTT    mRNA  3'  CTCC TC  TTAGC        miRNA 5'  GAGG AG  GGTTG                 T    T  TA     TATAGTT           3111131166221213333666 ``` \| SeedType = 2 NonSeedType = 2 \| \| --- \| --- \| --- \| --- \| --- \| --- \| | \| 633 \| WBGene00010657:K08D8.3(6488) \| ``` log2FC=-0.137 padj=0.84 ``` \| [seedLoc = 246,251] Seq=GGAATTTTGAAATGCTATCTCT \| ```          T         AA  TTTTAAGG mRNA  3'  CTCTATCGT  AG         miRNA 5'  GAGGTAGTA  TT                  T         GG  GTATAGTT           3111211121331233333333 ``` \| SeedType = 2 NonSeedType = -1 \| \| --- \| --- \| --- \| --- \| --- \| --- \| | \| 634 \| WBGene00010658:K08D8.4(6489) \| ``` log2FC=0.375 padj=0.658 ``` \| [seedLoc = 231,237] Seq=AAATTATCAAACTTATGGAAACTACACTCA \| ```              A     AAGGTA     AC      A mRNA  3' ACTC CATCA      TTCAA  TATTAA  miRNA 5' TGAG GTAGT      AGGTT  ATAGTT                                GT                  111151111155445512111331112115 ``` \| SeedType = 2 NonSeedType = 1 \| \| --- \| --- \| --- \| --- \| --- \| --- \| | \| 635 \| WBGene00010666:K08E4.2(6497) \| ``` log2FC=0.218 padj=0.934 ``` \| [seedLoc = 27,31] Seq=CTCTTTACTACCCTCT \| ```          T  C         CTC        mRNA  3'  CT CCATCATTT           miRNA 5'  GA GGTAGTAGG                    T            TTGTATAGTT           31151111111223336666666 ``` \| SeedType = 3 NonSeedType = -1 \| \| --- \| --- \| --- \| --- \| --- \| --- \| | \| 636 \| WBGene00010673:K08E7.5(6501) \| ``` log2FC=-0.015 padj=0.994 ``` \| [seedLoc = 38,43] Seq=ACATCGACGGGTTGTACATACCTCA \| ```                 A   GTTG    GCTACA mRNA  3' ACTCCAT CAT    GGCA       miRNA 5' TGAGGTA GTA    TTGT                           GG      ATAGTT           1111111511133542211333333 ``` \| SeedType = 1 NonSeedType = -1 \| \| --- \| --- \| --- \| --- \| --- \| --- \| | \| 637 \| WBGene00010681:mak-1(6508) \| ``` log2FC=-0.296 padj=0.492 ``` \| [seedLoc = 413,418] Seq=CATCAAGAATCCAACTGCCTCA \| ```                   A  TAAGAACTAC mRNA  3' ACTCCGTCA CC           miRNA 5' TGAGGTAGT GG                             A  TTGTATAGTT           1111121113113333333333 ``` \| SeedType = 2 NonSeedType = -1 \| \| --- \| --- \| --- \| --- \| --- \| --- \| | \| 638 \| WBGene00010700:nipi-3(6519) \| ``` log2FC=-0.128 padj=0.842 ``` \| [seedLoc = 162,167] Seq=TCAAGGTCGACAAACTACCTCT \| ```          T        AA    TGGAACT mRNA  3'  CTCCATCA  CAGC        miRNA 5'  GAGGTAGT  GTTG                 T        AG    TATAGTT           3111111113311213333333 ``` \| SeedType = 1 NonSeedType = -1 \| \| --- \| --- \| --- \| --- \| --- \| --- \| | \| 639 \| WBGene00010701:ent-2(6520) \| ``` log2FC=-0.412 padj=0.084 ``` \| [seedLoc = 220,225] Seq=CTTTGTATTTCCCAAATACCTCA \| ```                 AAAC  TT      TC mRNA  3' ACTCCAT    CC  TATGTT   miRNA 5' TGAGGTA    GG  GTATAG                   GTA   TT      TT           11111113335113321121233 ``` \| SeedType = 1 NonSeedType = -1 \| \| --- \| --- \| --- \| --- \| --- \| --- \| | \| 640 \| WBGene00010731:K10C3.4(6544) \| ``` log2FC=0.103 padj=0.977 ``` \| [seedLoc = 397,402] Seq=GTTTTCAGAACCAAATACCTCG \| ```          G      AAA    G CTTTTG mRNA  3'  CTCCAT   CCAA A       miRNA 5'  GAGGTA   GGTT T                T      GTA    G ATAGTT           3111111333111131333333 ``` \| SeedType = 1 NonSeedType = 1 \| \| --- \| --- \| --- \| --- \| --- \| --- \| | \|  \|  \|  \| [seedLoc = 528,533] Seq=CACTATGCCAACTACCTCT \| ```          T        A           C mRNA  3'  CTCCATCA CCG   TATCA  miRNA 5'  GAGGTAGT GGT   ATAGT           T        A   TGT     T           3111111113112666111113 ``` \| SeedType = 1 NonSeedType = 2 \| \| --- \| --- \| --- \| --- \| --- \| --- \| | \|  \|  \|  \| [seedLoc = 697,702] Seq=CTGTTCTCAACCTATTTTTATTGTTACCTCC \| ```          C      TGTTATTTT        TCT       mRNA  3'  CTCCAT         TATCCAAC   TGTC   miRNA 5'  GAGGTA         GTAGGTTG   ATAG            T                       T      TT           311111154555555521111111355121166 ``` \| SeedType = 1 NonSeedType = 1 \| \| --- \| --- \| --- \| --- \| --- \| --- \| | \| 641 \| WBGene00010815:M01F1.9(6605) \| ``` log2FC=0.077 padj=0.974 ``` \| [seedLoc = 416,419] Seq=GTCTGTATAATAGCTCCCTCT \| ```          T    C   A          TG mRNA  3'  CTCC TCG  TAATATGTC   miRNA 5'  GAGG AGT  GTTGTATAG            T    T   AG         TT           3111131123621121121133 ``` \| SeedType = 2 NonSeedType = -1 \| \| --- \| --- \| --- \| --- \| --- \| --- \| | \| 642 \| WBGene00010831:M02G9.2(6611) \| ``` log2FC=-1.027 padj=0.441 ``` \| [seedLoc = 48,54] Seq=TTTTGTATTTATTTATGTACCATCA \| ```             A    G      TT      TT mRNA  3' ACT CCAT TATTTA  TATGTT   miRNA 5' TGA GGTA GTAGGT  GTATAG                           T       TT           1115111142112213521121233 ``` \| SeedType = 3 NonSeedType = 2 \| \| --- \| --- \| --- \| --- \| --- \| --- \| | \| 643 \| WBGene00010870:let-522(6638) \|  \| [seedLoc = 97,102] Seq=AAACTAAATTTTGAACCCTGAAACTGCCTCG \| ```          G        AAGTC    GTTT  A     A mRNA  3'  CTCCGTCA     CCAA    TA ATCAA  miRNA 5'  GAGGTAGT     GGTT    GT TAGTT           T        A              A                 3111121113545511114555213111115 ``` \| SeedType = 2 NonSeedType = 1 \| \| --- \| --- \| --- \| --- \| --- \| --- \| | \| 644 \| WBGene00010873:trpa-2(6641) \| ``` log2FC=0.189 padj=0.936 ``` \| [seedLoc = 35,40] Seq=AAACTGGTTTCTGTACTACCTTT \| ```          T         G CTTTG     A mRNA  3'  TTCCATCAT T     GTCAA  miRNA 5'  GAGGTAGTA G     TAGTT           T         G TTGTA                 32111111113233333211115 ``` \| SeedType = 3 NonSeedType = -1 \| \| --- \| --- \| --- \| --- \| --- \| --- \| | \| 645 \| WBGene00010890:ddb-1(6655) \| ``` log2FC=0.099 padj=0.914 ``` \| [seedLoc = 285,290] Seq=TTGTTGTTTTTGTTCTACCTTT \| ```          T       TTGTTTTTGTTGTT mRNA  3'  TTCCATC               miRNA 5'  GAGGTAG                        T       TAGGTTGTATAGTT           3211111133333333333333 ``` \| SeedType = 3 NonSeedType = -1 \| \| --- \| --- \| --- \| --- \| --- \| --- \| | \|  \|  \|  \| [seedLoc = 352,357] Seq=TTTTTTCTAACCCTCTATCTCG \| ```          G       TC     CTTTTTT mRNA  3'  CTCTATC  CCAAT        miRNA 5'  GAGGTAG  GGTTG                 T       TA     TATAGTT           3111211133111123333333 ``` \| SeedType = 2 NonSeedType = 1 \| \| --- \| --- \| --- \| --- \| --- \| --- \| | \| 646 \| WBGene00010896:M28.5(6660) \| ``` log2FC=-0.071 padj=0.945 ``` \| [seedLoc = 29,34] Seq=TTTTATCAATTTGTTGTTACGTTACCTCA \| ```                  GCATTGTT            TT mRNA  3' ACTCCATT        GTTTAAC TATT   miRNA 5' TGAGGTAG        TAGGTTG ATAG                                   T    TT           111111124555545521221116111233 ``` \| SeedType = 1 NonSeedType = 1 \| \| --- \| --- \| --- \| --- \| --- \| --- \| | \| 647 \| WBGene00010926:M153.3(6682) \| ``` log2FC=-1.04 padj=1.0 ``` \| [seedLoc = 102,107] Seq=GAGAGTACATATCTGGTTTTACCTCC \| ```          C       TTG     T     AGAG mRNA  3'  CTCCATT   GTCTA ACATG     miRNA 5'  GAGGTAG   TAGGT TGTAT              T                     AGTT           31111112554211215111123333 ``` \| SeedType = 1 NonSeedType = 2 \| \| --- \| --- \| --- \| --- \| --- \| --- \| | \| 648 \| WBGene00010935:M163.1(6684) \| ``` log2FC=-0.462 padj=0.359 ``` \| [seedLoc = 205,210] Seq=CAATTGTATTCTGATACCTCC \| ```          C      A              C mRNA  3'  CTCCAT GTCT  TATGTTAA  miRNA 5'  GAGGTA TAGG  GTATAGTT           T      G    TT                    31111113211266211212115 ``` \| SeedType = 1 NonSeedType = -1 \| \| --- \| --- \| --- \| --- \| --- \| --- \| | \| 649 \| WBGene00010936:ztf-14(6685) \| ``` log2FC=-0.044 padj=0.994 ``` \| [seedLoc = 382,387] Seq=TGATTGTGATCTCTTGTAGTTACGTACCTCT \| ```          T      G    G TGTTCTCTA       T mRNA  3'  CTCCAT CATT A         GTGTTAG  miRNA 5'  GAGGTA GTAG T         TATAGTT           T           G TG                          3111111411123133555555521212125 ``` \| SeedType = 1 NonSeedType = -1 \| \| --- \| --- \| --- \| --- \| --- \| --- \| | \|  \|  \|  \| [seedLoc = 460,465] Seq=TACTATTATTATTCTACCTTT \| ```          T       TTATTAT      T mRNA  3'  TTCCATC        TATCA  miRNA 5'  GAGGTAG        ATAGT           T       TAGGTTGT     T           3211111133333336111113 ``` \| SeedType = 3 NonSeedType = -1 \| \| --- \| --- \| --- \| --- \| --- \| --- \| | \| 650 \| WBGene00010993:vha-20(6715) \|  \| [seedLoc = 212,217] Seq=TTTTTCCCATCCTGTTGGTTTATCTACTTCA \| ```                  TATTTGGTT    T  CCTTTTT mRNA  3' ACTTCATC         GTCC AC        miRNA 5' TGAGGTAG         TAGG TG                                      T  TATAGTT           1112111155555445521113113333333 ``` \| SeedType = 3 NonSeedType = -1 \| \| --- \| --- \| --- \| --- \| --- \| --- \| | \| 651 \| WBGene00011015:R04F11.2(6726) \| ``` log2FC=0.004 padj=0.997 ``` \| [seedLoc = 70,75] Seq=TATTAGTTCAATGAATACCTCT \| ```          T      AAG     TTG    T mRNA  3'  CTCCAT    TAAC   ATTA  miRNA 5'  GAGGTA    GTTG   TAGT           T      GTAG    TA     T           31111113336211133411213 ``` \| SeedType = 1 NonSeedType = 1 \| \| --- \| --- \| --- \| --- \| --- \| --- \| | \| 652 \| WBGene00011068:set-14(6773) \| ``` log2FC=0.204 padj=0.816 ``` \| [seedLoc = 76,81] Seq=TGTTATGCAAAATATTATTTATTACCTCA \| ```                       TTATAAA       GT mRNA  3' ACTCCATTATTTA       ACGTATT   miRNA 5' TGAGGTAGTAGGT       TGTATAG                                       TT           11111112112215555555112111233 ``` \| SeedType = 1 NonSeedType = 2 \| \| --- \| --- \| --- \| --- \| --- \| --- \| | \| 653 \| WBGene00011071:R06F6.8(6776) \| ``` log2FC=0.122 padj=0.9 ``` \| [seedLoc = 154,160] Seq=TAACTGTGATGTGTGATACACTCG \| ```          G   A   AGTGTGTA       T mRNA  3'  CTC CAT        GTGTCAA  miRNA 5'  GAG GTA        TATAGTT           T       GTAGGTTG                   311151113333333321211115 ``` \| SeedType = 2 NonSeedType = -1 \| \| --- \| --- \| --- \| --- \| --- \| --- \| | \| 654 \| WBGene00011081:R07B1.9(6783) \| ``` log2FC=-0.639 padj=0.367 ``` \| [seedLoc = 49,54] Seq=CACTAGAAACTGTTAGCTGCCTCA \| ```                   ATTG    AG     C mRNA  3' ACTCCGTCG    TCAA   ATCA  miRNA 5' TGAGGTAGT    GGTT   TAGT                    A       GTA    T           1111121123554211133611113 ``` \| SeedType = 2 NonSeedType = 1 \| \| --- \| --- \| --- \| --- \| --- \| --- \| | \| 655 \| WBGene00011095:gana-1(6794) \| ``` log2FC=-0.034 padj=0.988 ``` \| [seedLoc = 68,73] Seq=TCTTGTATATTGATATATGTACCTCA \| ```                 GTA   AGTT       CT mRNA  3' ACTCCAT   TAT    ATATGTT   miRNA 5' TGAGGTA   GTA    TGTATAG                         GGT        TT           11111114552113335121121233 ``` \| SeedType = 1 NonSeedType = -1 \| \| --- \| --- \| --- \| --- \| --- \| --- \| | \| 656 \| WBGene00011104:cut-5(6801) \| ``` log2FC=-0.381 padj=1.0 ``` \| [seedLoc = 73,78] Seq=ATAAGTGCAATGCTACTTCT \| ```          T                 AATA mRNA  3'  CTTCATCG  TAACGTG     miRNA 5'  GAGGTAGT  GTTGTAT              T        AG       AGTT           3112111126621112123333 ``` \| SeedType = 3 NonSeedType = 1 \| \| --- \| --- \| --- \| --- \| --- \| --- \| | \| 657 \| WBGene00011146:pde-2(6834) \| ``` log2FC=-0.224 padj=0.64 ``` \| [seedLoc = 509,514] Seq=TCGAAAACAAATCGCTTTACCTCG \| ```          G       TCG  A    AAAGCT mRNA  3'  CTCCATT   CT AACA       miRNA 5'  GAGGTAG   GG TTGT                T       TA        ATAGTT           311111123341251111333333 ``` \| SeedType = 1 NonSeedType = 1 \| \| --- \| --- \| --- \| --- \| --- \| --- \| | \| 658 \| WBGene00011198:R10E4.1(6860) \| ``` log2FC=0.213 padj=0.74 ``` \| [seedLoc = 77,83] Seq=TTGAAATCAAGCACATACACTCG \| ```          G   A   A    G   TAAAGTT mRNA  3'  CTC CAT CA C AAC        miRNA 5'  GAG GTA GT G TTG                 T          A G   TATAGTT           311151115116131113333333 ``` \| SeedType = 2 NonSeedType = 1 \| \| --- \| --- \| --- \| --- \| --- \| --- \| | \| 659 \| WBGene00011216:usp-46(6872) \| ``` log2FC=-0.11 padj=0.938 ``` \| [seedLoc = 143,148] Seq=TTTCATACAATTTTCTATCACACTATCTCT \| ```          T        CACTATCT         CTTT mRNA  3'  CTCTATCA        TTTAACATA     miRNA 5'  GAGGTAGT        AGGTTGTAT              T                         AGTT           311121111555555551221111113333 ``` \| SeedType = 2 NonSeedType = 1 \| \| --- \| --- \| --- \| --- \| --- \| --- \| | \| 660 \| WBGene00011240:mask-1(6893) \|  \| [seedLoc = 344,349] Seq=TCTCCCCCGACATACTACTTCT \| ```          T         A    CCCCTCT mRNA  3'  CTTCATCAT CAGC        miRNA 5'  GAGGTAGTA GTTG                 T         G    TATAGTT           3112111111311213333333 ``` \| SeedType = 3 NonSeedType = -1 \| \| --- \| --- \| --- \| --- \| --- \| --- \| | \| 661 \| WBGene00011259:miz-1(6906) \| ``` log2FC=-0.143 padj=0.815 ``` \| [seedLoc = 524,529] Seq=ATCTCTACACCCCTACCTTA \| ```                      C    C  TA mRNA  3' ATTCCATC  CC ACAT TC   miRNA 5' TGAGGTAG  GG TGTA AG                    TA  T    T  TT           1211111166113111131133 ``` \| SeedType = 3 NonSeedType = -1 \| \| --- \| --- \| --- \| --- \| --- \| --- \| | \| 662 \| WBGene00011276:aakg-5(6922) \| ``` log2FC=0.087 padj=0.945 ``` \| [seedLoc = 259,264] Seq=TTACAACACCTAGAGCTTTACTGCTACCTCT \| ```          T          ATTTCGAGATCCA      T     mRNA  3'  CTCCATCGTC             CAACAT      miRNA 5'  GAGGTAGTAG             GTTGTA               T                             TAGTT           31111111211555554545555511111136666 ``` \| SeedType = 1 NonSeedType = 1 \| \| --- \| --- \| --- \| --- \| --- \| --- \| | \| 663 \| WBGene00011293:R102.6(6938) \| ``` log2FC=-0.211 padj=0.966 ``` \| [seedLoc = 48,53] Seq=TCCTCAAAAACCAAATACTACCTCA \| ```                    AAA    AAACTCCT mRNA  3' ACTCCATCAT   CCAA         miRNA 5' TGAGGTAGTA   GGTT                                   GTATAGTT           1111111111555111133333333 ``` \| SeedType = 1 NonSeedType = 1 \| \| --- \| --- \| --- \| --- \| --- \| --- \| | \|  \|  \|  \| [seedLoc = 55,60] Seq=CCTCAAAAACCAAATACTACCTCAT \| ```          T          AAA    AAACTCC  mRNA  3'  ACTCCATCAT   CCAA         miRNA 5'  TGAGGTAGTA   GGTT                                    GTATAGTT           51111111111555111133333336 ``` \| SeedType = 1 NonSeedType = 1 \| \| --- \| --- \| --- \| --- \| --- \| --- \| | \|  \|  \|  \| [seedLoc = 109,114] Seq=AAACTGTAAAGCAACTACTTCC \| ```          C        A    A       A mRNA  3'  CTTCATCA  CGA ATGTCAA  miRNA 5'  GAGGTAGT  GTT TATAGTT           T        AG   G                   31121111136121311211115 ``` \| SeedType = 3 NonSeedType = -1 \| \| --- \| --- \| --- \| --- \| --- \| --- \| | \| 664 \| WBGene00011304:mnk-1(6947) \| ``` log2FC=-0.252 padj=0.389 ``` \| [seedLoc = 1577,1582] Seq=AAATCCTTCCCCGCCTATCTCG \| ```          G       CG  CCTTCCTAAA mRNA  3'  CTCTATC  CC           miRNA 5'  GAGGTAG  GG                    T       TA  TTGTATAGTT           3111211133113333333333 ``` \| SeedType = 2 NonSeedType = -1 \| \| --- \| --- \| --- \| --- \| --- \| --- \| | \|  \|  \|  \| [seedLoc = 1709,1714] Seq=TGGAGTTCACGCTCTATCTCT \| ```          T          GC  TTGAGGT mRNA  3'  CTCTATC TC  AC        miRNA 5'  GAGGTAG AG  TG                 T       T  GT  TATAGTT           3111211161133113333333 ``` \| SeedType = 2 NonSeedType = -1 \| \| --- \| --- \| --- \| --- \| --- \| --- \| | \| 665 \| WBGene00011308:pnn-1(6951) \|  \| [seedLoc = 151,157] Seq=CGTTCTCCAAAATAATTACTCTCT \| ```          T   T     ATAA   CTCTTGC mRNA  3'  CTC CATTA    AAC        miRNA 5'  GAG GTAGT    TTG                 T         AGG    TATAGTT           311151112133351113333333 ``` \| SeedType = 2 NonSeedType = 1 \| \| --- \| --- \| --- \| --- \| --- \| --- \| | \| 666 \| WBGene00011326:T01D3.1(6968) \| ``` log2FC=0.218 padj=0.957 ``` \| [seedLoc = 175,181] Seq=ATCGATATTCATATCTACACTCA \| ```              A    TATACT    GCTA mRNA  3' ACTC CATC      TATA     miRNA 5' TGAG GTAG      GTAT                       TAGGTT    AGTT           11115111133333321113333 ``` \| SeedType = 2 NonSeedType = -1 \| \| --- \| --- \| --- \| --- \| --- \| --- \| | \| 667 \| WBGene00011372:npr-26(6999) \| ``` log2FC=0.691 padj=1.0 ``` \| [seedLoc = 177,183] Seq=CGGTAAATGAATAATTACACTCG \| ```          G   A     ATAAGTAAATGGC mRNA  3'  CTC CATTA              miRNA 5'  GAG GTAGT                       T         AGGTTGTATAGTT           31115111213333333333333 ``` \| SeedType = 2 NonSeedType = -1 \| \| --- \| --- \| --- \| --- \| --- \| --- \| | \| 668 \| WBGene00011383:T02E9.5(7009) \| ``` log2FC=-0.042 padj=0.992 ``` \| [seedLoc = 84,90] Seq=TCAATTTCATATTATTACCCTCA \| ```             C        AT  TTTAACT mRNA  3' ACT CCATTATT  AC        miRNA 5' TGA GGTAGTAG  TG                             GT  TATAGTT           11151111211233113333333 ``` \| SeedType = 3 NonSeedType = -1 \| \| --- \| --- \| --- \| --- \| --- \| --- \| | \| 669 \| WBGene00011409:T04A8.7(7026) \| ``` log2FC=0.126 padj=0.825 ``` \| [seedLoc = 148,153] Seq=TTTTATATATTACCTACCTCT \| ```          T       CATT        TT mRNA  3'  CTCCATC     ATATATT   miRNA 5'  GAGGTAG     TGTATAG            T       TAGGT       TT           3111111133336121111233 ``` \| SeedType = 1 NonSeedType = -1 \| \| --- \| --- \| --- \| --- \| --- \| --- \| | \| 670 \| WBGene00011410:T04A8.8(7027) \| ``` log2FC=-0.18 padj=0.746 ``` \| [seedLoc = 31,36] Seq=TTCTGTAAATTTGACTACCTTA \| ```                  A            TT mRNA  3' ATTCCATC GTTTAA ATGTC   miRNA 5' TGAGGTAG TAGGTT TATAG                           G     TT           12111111521221161121133 ``` \| SeedType = 3 NonSeedType = 2 \| \| --- \| --- \| --- \| --- \| --- \| --- \| | \| 671 \| WBGene00011434:T04D3.5(7041) \| ``` log2FC=-0.4 padj=0.931 ``` \| [seedLoc = 68,73] Seq=AAATGTACAAAACTCTTGATCTACCTCT \| ```          T       TA    TCAA       AAA mRNA  3'  CTCCATC  GTTC    AACATGT    miRNA 5'  GAGGTAG  TAGG    TTGTATA             T                        GTT           3111111155212155551111121333 ``` \| SeedType = 1 NonSeedType = 1 \| \| --- \| --- \| --- \| --- \| --- \| --- \| | \| 672 \| WBGene00011456:T05A1.5(7058) \| ``` log2FC=0.272 padj=1.0 ``` \| [seedLoc = 198,204] Seq=TTGATTCTTGCCTTTTCGCTACTCTCC \| ```          C   T     CTTT    TTCTTAGTT mRNA  3'  CTC CATCG    TCCG          miRNA 5'  GAG GTAGT    AGGT                   T                 TGTATAGTT           311151111255551112333333333 ``` \| SeedType = 2 NonSeedType = 2 \| \| --- \| --- \| --- \| --- \| --- \| --- \| | \| 673 \| WBGene00011464:T05B9.1(7062) \| ``` log2FC=0.067 padj=0.953 ``` \| [seedLoc = 113,116] Seq=GTGAATGCACCCCCCAACCTCC \| ```          C     A CC  C     AGTG mRNA  3'  CTCCA C  CC ACGTA     miRNA 5'  GAGGT G  GG TGTAT              T     A TA  T     AGTT           3111113133113112113333 ``` \| SeedType = 2 NonSeedType = -1 \| \| --- \| --- \| --- \| --- \| --- \| --- \| | \| 674 \| WBGene00011465:T05B9.2(7063) \| ``` log2FC=1.203 padj=1.0 ``` \| [seedLoc = 213,218] Seq=ATTTTTAAAACTATGTACCTCT \| ```          T      GTA    AATTTTTA mRNA  3'  CTCCAT   TCAA         miRNA 5'  GAGGTA   GGTT                  T      GTA    GTATAGTT           3111111333211133333333 ``` \| SeedType = 1 NonSeedType = 1 \| \| --- \| --- \| --- \| --- \| --- \| --- \| | \| 675 \| WBGene00011479:T05E11.2(7070) \| ``` log2FC=-0.236 padj=0.861 ``` \| [seedLoc = 150,155] Seq=TTCTGAATGACATTTGATAGCTACTTCG \| ```          G        ATAGTTTA     A   TT mRNA  3'  CTTCATCG        CAGTA GTC   miRNA 5'  GAGGTAGT        GTTGT TAG            T        AG           A   TT           3112111123354555511221321133 ``` \| SeedType = 3 NonSeedType = -1 \| \| --- \| --- \| --- \| --- \| --- \| --- \| | \| 676 \| WBGene00011531:rsbp-1(7112) \| ``` log2FC=-0.067 padj=0.98 ``` \| [seedLoc = 269,274] Seq=TTCTAGTCAATTACCTCA \| ```                         TG   TT mRNA  3' ACTCCAT    TAAC  ATC   miRNA 5' TGAGGTA    GTTG  TAG                   GTAG    TA   TT           1111111666621113311133 ``` \| SeedType = 1 NonSeedType = 1 \| \| --- \| --- \| --- \| --- \| --- \| --- \| | \| 677 \| WBGene00011547:T06G6.5(7122) \| ``` log2FC=0.1 padj=1.0 ``` \| [seedLoc = 33,38] Seq=TCCTAATCCTCCATTACCTCG \| ```          G           TC TA   CT mRNA  3'  CTCCATTA CC  C  ATC   miRNA 5'  GAGGTAGT GG  G  TAG            T        A  TT TA   TT           3111111216113313311133 ``` \| SeedType = 1 NonSeedType = -1 \| \| --- \| --- \| --- \| --- \| --- \| --- \| | \| 678 \| WBGene00011561:ttr-15(7130) \| ``` log2FC=0.188 padj=0.865 ``` \| [seedLoc = 48,53] Seq=AAGGGAAGGACTTTCTATCTCT \| ```          T       T     G AGGGAA mRNA  3'  CTCTATC TTCAG A       miRNA 5'  GAGGTAG AGGTT T                T       T     G ATAGTT           3111211131211231333333 ``` \| SeedType = 2 NonSeedType = 2 \| \| --- \| --- \| --- \| --- \| --- \| --- \| | \| 679 \| WBGene00011604:T08A11.1(7156) \| ``` log2FC=0.286 padj=0.469 ``` \| [seedLoc = 229,235] Seq=ATTTTTATCTCTTATTACTCTCA \| ```              T        TC   TTTTA mRNA  3' ACTC CATTATTC  TAT      miRNA 5' TGAG GTAGTAGG  GTA                            TT   TAGTT           11115111211213321133333 ``` \| SeedType = 2 NonSeedType = -1 \| \| --- \| --- \| --- \| --- \| --- \| --- \| | \| 680 \| WBGene00011605:sftb-1(7157) \|  \| [seedLoc = 52,57] Seq=ATCTCGATAACTTATTTTTCTATCTCC \| ```          C       TTTTT        GC  TA mRNA  3'  CTCTATC     ATTCAATA  TC   miRNA 5'  GAGGTAG     TAGGTTGT  AG            T                    AT  TT           311121115555511211121331133 ``` \| SeedType = 2 NonSeedType = 1 \| \| --- \| --- \| --- \| --- \| --- \| --- \| | \| 681 \| WBGene00011611:T08D2.6(7158) \| ``` log2FC=-0.298 padj=0.865 ``` \| [seedLoc = 133,138] Seq=TTTTTTAAAATTTTCTACTTCA \| ```                  T     AATTTTTT mRNA  3' ACTTCATC TTTAA         miRNA 5' TGAGGTAG AGGTT                          T     GTATAGTT           1112111131221133333333 ``` \| SeedType = 3 NonSeedType = 2 \| \| --- \| --- \| --- \| --- \| --- \| --- \| | \| 682 \| WBGene00011614:nfya-1(7160) \| ``` log2FC=0.946 padj=0.166 ``` \| [seedLoc = 166,172] Seq=TGTTTTCAAATTTTTTATCCTCA \| ```               T   T     ACTTTTGT mRNA  3' ACTCC ATT TTTAA         miRNA 5' TGAGG TAG AGGTT                           T     GTATAGTT           11111511231221133333333 ``` \| SeedType = 2 NonSeedType = 2 \| \| --- \| --- \| --- \| --- \| --- \| --- \| | \| 683 \| WBGene00011636:cec-3(7170) \| ``` log2FC=-0.224 padj=0.716 ``` \| [seedLoc = 79,84] Seq=CAGTTCAAAGCATATCTATCTCT \| ```          T       TATA   AACTTGAC mRNA  3'  CTCTATC    CGA         miRNA 5'  GAGGTAG    GTT                  T       TAG    GTATAGTT           31112111333512133333333 ``` \| SeedType = 2 NonSeedType = -1 \| \| --- \| --- \| --- \| --- \| --- \| --- \| | \| 684 \| WBGene00011662:T09F3.2(7183) \| ``` log2FC=-0.272 padj=0.373 ``` \| [seedLoc = 283,288] Seq=CAGGAACCAATTTACTACTTCT \| ```          T              CAAGGAC mRNA  3'  CTTCATCATTTAAC        miRNA 5'  GAGGTAGTAGGTTG                 T              TATAGTT           3112111111221113333333 ``` \| SeedType = 3 NonSeedType = 1 \| \| --- \| --- \| --- \| --- \| --- \| --- \| | \| 685 \| WBGene00011713:T11F9.12(7214) \| ``` log2FC=0.444 padj=0.303 ``` \| [seedLoc = 119,124] Seq=TCAAATGGGTTTTGTCTACTTCA \| ```                  T    T G   AACT mRNA  3' ACTTCATC GTTT G GTA     miRNA 5' TGAGGTAG TAGG T TAT                           T G   AGTT           11121111521223232113333 ``` \| SeedType = 3 NonSeedType = -1 \| \| --- \| --- \| --- \| --- \| --- \| --- \| | \| 686 \| WBGene00011719:T11G6.4(7217) \| ``` log2FC=0.281 padj=0.791 ``` \| [seedLoc = 86,92] Seq=TCTTTAAAAACTTTTTAACCTCA \| ```               A   T     AAATTTCT mRNA  3' ACTCC ATT TTCAA         miRNA 5' TGAGG TAG AGGTT                           T     GTATAGTT           11111511231211133333333 ``` \| SeedType = 2 NonSeedType = 1 \| \| --- \| --- \| --- \| --- \| --- \| --- \| | \| 687 \| WBGene00011730:drr-2(7225) \| ``` log2FC=-0.176 padj=0.769 ``` \| [seedLoc = 159,164] Seq=GTCTAATGATCACAAATTACCTCA \| ```                   AACA          TG mRNA  3' ACTCCATTA    CTAGTA ATC   miRNA 5' TGAGGTAGT    GGTTGT TAG                     A         A   TT           1111111213555121221611133 ``` \| SeedType = 1 NonSeedType = 2 \| \| --- \| --- \| --- \| --- \| --- \| --- \| | \| 688 \| WBGene00011734:mlc-5(7229) \| ``` log2FC=0.04 padj=0.977 ``` \| [seedLoc = 169,175] Seq=CAGTGAACAACTTTGTATCCTCC \| ```          C    T  GT       AGTGAC mRNA  3'  CTCC AT  TTCAACA       miRNA 5'  GAGG TA  AGGTTGT                T       GT       ATAGTT           31111511331211111333333 ``` \| SeedType = 2 NonSeedType = 1 \| \| --- \| --- \| --- \| --- \| --- \| --- \| | \| 689 \| WBGene00011752:fbxa-2(7243) \| ``` log2FC=-0.05 padj=0.994 ``` \| [seedLoc = 1119,1124] Seq=TTTTTGTTGTGCTAGTACCTCG \| ```          G      G   GTGTTGTTTTT mRNA  3'  CTCCAT ATC            miRNA 5'  GAGGTA TAG                     T      G   GTTGTATAGTT           3111111311133333333333 ``` \| SeedType = 1 NonSeedType = -1 \| \| --- \| --- \| --- \| --- \| --- \| --- \| | \| 690 \| WBGene00011763:T14B1.1(7253) \| ``` log2FC=-0.026 padj=1.0 ``` \| [seedLoc = 113,118] Seq=TCCATTATAACCCCCTACCTCG \| ```          G       CC       TACCT mRNA  3'  CTCCATC  CCAATAT      miRNA 5'  GAGGTAG  GGTTGTA               T       TA       TAGTT           3111111133111121133333 ``` \| SeedType = 1 NonSeedType = 1 \| \| --- \| --- \| --- \| --- \| --- \| --- \| | \|  \|  \|  \| [seedLoc = 148,153] Seq=TGATTATCACATCTACCTCT \| ```          T       TA            T mRNA  3'  CTCCATC   C AC TATTAG  miRNA 5'  GAGGTAG   G TG ATAGTT           T       TAG T  T                  31111111336161161112125 ``` \| SeedType = 1 NonSeedType = -1 \| \| --- \| --- \| --- \| --- \| --- \| --- \| | \|  \|  \|  \| [seedLoc = 180,185] Seq=CCCCTTTCGACCTCAATCTATCTCT \| ```          T       TAAC      TTTCCCC mRNA  3'  CTCTATC    TCCAGC        miRNA 5'  GAGGTAG    AGGTTG                 T       T         TATAGTT           3111211135551111213333333 ``` \| SeedType = 2 NonSeedType = 1 \| \| --- \| --- \| --- \| --- \| --- \| --- \| | \|  \|  \|  \| [seedLoc = 191,196] Seq=TCGACCTCAATCTATCTCTTTCCTACCTCT \| ```          T       CTTTCTCT       TCCAGCT mRNA  3'  CTCCATC        ATCTAAC        miRNA 5'  GAGGTAG        TAGGTTG                 T                      TATAGTT           311111115555555511121113333333 ``` \| SeedType = 1 NonSeedType = 1 \| \| --- \| --- \| --- \| --- \| --- \| --- \| | \|  \|  \|  \| [seedLoc = 234,239] Seq=TGATTAACAACCCACCAAAGCTACCTCT \| ```          T        AAACCAC            T mRNA  3'  CTCCATCG       CCAACA ATTAG  miRNA 5'  GAGGTAGT       GGTTGT TAGTT           T        A            A                 31111111235555551111116112125 ``` \| SeedType = 1 NonSeedType = 1 \| \| --- \| --- \| --- \| --- \| --- \| --- \| | \|  \|  \|  \| [seedLoc = 285,290] Seq=ACCCTCCTAATCTAAGTCAATTACCTCA \| ```                   ACTGAA      CCTCCCA mRNA  3' ACTCCATTA      TCTAAT        miRNA 5' TGAGGTAGT      AGGTTG                                      TATAGTT           1111111215554551121123333333 ``` \| SeedType = 1 NonSeedType = 2 \| \| --- \| --- \| --- \| --- \| --- \| --- \| | \|  \|  \|  \| [seedLoc = 943,948] Seq=ATTTTTACTCGCTTTGCCTACCTCA \| ```                  CGT    CT   TTTTA mRNA  3' ACTCCATC   TTCG  CAT      miRNA 5' TGAGGTAG   AGGT  GTA                       T      T    TAGTT           1111111134512123511133333 ``` \| SeedType = 1 NonSeedType = 2 \| \| --- \| --- \| --- \| --- \| --- \| --- \| | \|  \|  \|  \| [seedLoc = 1054,1059] Seq=CTCTATTTAATTCTACCTCT \| ```          T              T    TC mRNA  3'  CTCCATC  TTAAT TATC   miRNA 5'  GAGGTAG  GGTTG ATAG            T       TA     T    TT           3111111166221123111133 ``` \| SeedType = 1 NonSeedType = 2 \| \| --- \| --- \| --- \| --- \| --- \| --- \| | \| 691 \| WBGene00011773:ttr-53(7259) \| ``` log2FC=-0.087 padj=0.945 ``` \| [seedLoc = 301,307] Seq=AAGCTATGAACTATCTACTAACCTCT \| ```          T    A        TCAA       A mRNA  3'  CTCC ATCATCTA    GTATCGA  miRNA 5'  GAGG TAGTAGGT    TATAGTT           T             TG                     31111511111121335521111215 ``` \| SeedType = 2 NonSeedType = 2 \| \| --- \| --- \| --- \| --- \| --- \| --- \| | \| 692 \| WBGene00011794:nep-21(7270) \| ``` log2FC=0.271 padj=0.824 ``` \| [seedLoc = 286,291] Seq=CGCTAATTTGATCTCATTAATTCTATCTCC \| ```          C       TTAATTAC      TTA    C mRNA  3'  CTCTATC        TCTAGT   ATCG  miRNA 5'  GAGGTAG        AGGTTG   TAGT           T       T             TA     T           311121113555555511212233511123 ``` \| SeedType = 2 NonSeedType = 2 \| \| --- \| --- \| --- \| --- \| --- \| --- \| | \| 693 \| WBGene00011804:T16G12.3(7278) \| ``` log2FC=0.001 padj=0.999 ``` \| [seedLoc = 390,396] Seq=AAGCAAACGAAAAGTGCTACCTAC \| ```          CA        GAAA    AACGAA mRNA  3'   TCCATCGT    AGCA       miRNA 5'   AGGTAGTA    TTGT                TG        GG      ATAGTT           331111112133551211333333 ``` \| SeedType = 3 NonSeedType = -1 \| \| --- \| --- \| --- \| --- \| --- \| --- \| | \| 694 \| WBGene00011823:T18D3.6(7293) \| ``` log2FC=0.074 padj=0.985 ``` \| [seedLoc = 280,286] Seq=CATTTCTCATCCTGTTTTATACTCTCA \| ```              T   ATTTT    T  TCTTTAC mRNA  3' ACTC CAT     GTCC AC        miRNA 5' TGAG GTA     TAGG TG                         G        T  TATAGTT           111151113555521113113333333 ``` \| SeedType = 2 NonSeedType = -1 \| \| --- \| --- \| --- \| --- \| --- \| --- \| | \| 695 \| WBGene00011824:T18D3.7(7294) \| ``` log2FC=-0.545 padj=0.106 ``` \| [seedLoc = 142,148] Seq=TTTTTCAAAACATACATACTCTCT \| ```          T   T   A   A   AACTTTTT mRNA  3'  CTC CAT CAT CAA         miRNA 5'  GAG GTA GTA GTT                  T           G   GTATAGTT           311151115111311133333333 ``` \| SeedType = 2 NonSeedType = 1 \| \| --- \| --- \| --- \| --- \| --- \| --- \| | \| 696 \| WBGene00011827:T19A6.1(7295) \| ``` log2FC=-0.08 padj=0.956 ``` \| [seedLoc = 578,583] Seq=GGATTATTGATATTTACCTCA \| ```                  TA            G mRNA  3' ACTCCATT   TAGT TATTAG  miRNA 5' TGAGGTAG   GTTG ATAGTT                   TAG    T                  11111112336212261112124 ``` \| SeedType = 1 NonSeedType = -1 \| \| --- \| --- \| --- \| --- \| --- \| --- \| | \| 697 \| WBGene00011857:coa-4(7313) \|  \| [seedLoc = 153,159] Seq=AGAATTGGGAGTGACATACACTCT \| ```          T   A   A  G G  G  TAAGA mRNA  3'  CTC CAT CA T AG GT      miRNA 5'  GAG GTA GT G TT TA               T          A G  G  TAGTT           311151115113231232133333 ``` \| SeedType = 2 NonSeedType = -1 \| \| --- \| --- \| --- \| --- \| --- \| --- \| | \| 698 \| WBGene00011870:T20G5.8(7321) \| ``` log2FC=-0.446 padj=0.619 ``` \| [seedLoc = 187,192] Seq=TTTTATAGTTTCTGAAAGCTACTTCT \| ```          T        AAAG   TTG     TT mRNA  3'  CTTCATCG    TCT   ATATT   miRNA 5'  GAGGTAGT    AGG   TATAG            T               TTG     TT           31121111255541123331111233 ``` \| SeedType = 3 NonSeedType = -1 \| \| --- \| --- \| --- \| --- \| --- \| --- \| | \| 699 \| WBGene00011871:T20G5.9(7322) \| ``` log2FC=0.047 padj=0.982 ``` \| [seedLoc = 29,34] Seq=ATTGCAACGATTAAATATAGATTCTACCTCT \| ```          T       TTAGATATAAATTAG      TA    mRNA  3'  CTCCATC               CAACGT      miRNA 5'  GAGGTAG               GTTGTA               T       TAG                  TAGTT           3111111133345555555555411112133666 ``` \| SeedType = 1 NonSeedType = 1 \| \| --- \| --- \| --- \| --- \| --- \| --- \| | \| 700 \| WBGene00011890:mics-1(7335) \| ``` log2FC=0.055 padj=0.967 ``` \| [seedLoc = 49,54] Seq=TGTAATATTAATTGTATTACCTCT \| ```          T         G TAAT    ATGT mRNA  3'  CTCCATTAT T    TATA     miRNA 5'  GAGGTAGTA G    GTAT              T         G TT      AGTT           311111121132335521113333 ``` \| SeedType = 1 NonSeedType = -1 \| \| --- \| --- \| --- \| --- \| --- \| --- \| | \| 701 \| WBGene00011892:erh-1(7339) \|  \| [seedLoc = 27,32] Seq=GGGTTTCTCACTTCTATCTCT \| ```          T            CTCTTTGGG mRNA  3'  CTCTATC TTCA          miRNA 5'  GAGGTAG AGGT                   T       T    TGTATAGTT           3111211161211333333333 ``` \| SeedType = 2 NonSeedType = 2 \| \| --- \| --- \| --- \| --- \| --- \| --- \| | \| 702 \| WBGene00011906:hsp-12.1(7349) \| ``` log2FC=-0.142 padj=0.933 ``` \| [seedLoc = 139,144] Seq=ATTTTGAAAATAAAGTACCTCG \| ```          G      G AATAAAAGTTTTA mRNA  3'  CTCCAT A              miRNA 5'  GAGGTA T                       T      G AGGTTGTATAGTT           3111111313333333333333 ``` \| SeedType = 1 NonSeedType = -1 \| \| --- \| --- \| --- \| --- \| --- \| --- \| | \| 703 \| WBGene00011916:T22C1.5(7358) \| ``` log2FC=-0.139 padj=0.926 ``` \| [seedLoc = 257,262] Seq=TTTGTTGCTTAAATACTACCTTT \| ```          T         AAATT   TGTTT mRNA  3'  TTCCATCAT     CGT      miRNA 5'  GAGGTAGTA     GTA               T         GGTT    TAGTT           32111111113333512133333 ``` \| SeedType = 3 NonSeedType = -1 \| \| --- \| --- \| --- \| --- \| --- \| --- \| | \| 704 \| WBGene00011932:sptl-3(7369) \| ``` log2FC=0.379 padj=0.168 ``` \| [seedLoc = 24,30] Seq=AGGAGTATTTATTAACTATCCTCG \| ```          G    T    A   TT    AGGA mRNA  3'  CTCC ATCA TTA  TATG     miRNA 5'  GAGG TAGT GGT  GTAT              T         A   T     AGTT           311115111132213521123333 ``` \| SeedType = 2 NonSeedType = 2 \| \| --- \| --- \| --- \| --- \| --- \| --- \| | \| 705 \| WBGene00011956:T23F11.4(7388) \| ``` log2FC=0.694 padj=0.473 ``` \| [seedLoc = 88,94] Seq=TGTAATTATTTCTATAATACTCTCA \| ```              T   AA     TTATTAATGT mRNA  3' ACTC CAT  TATCT           miRNA 5' TGAG GTA  GTAGG                                   TTGTATAGTT           1111511155211123333333333 ``` \| SeedType = 2 NonSeedType = -1 \| \| --- \| --- \| --- \| --- \| --- \| --- \| | \| 706 \| WBGene00011962:T23G5.2(7392) \| ``` log2FC=-0.283 padj=0.331 ``` \| [seedLoc = 74,79] Seq=TCCTTTCTTCTGTACTACTTCC \| ```          C         G CTTCTTTCCT mRNA  3'  CTTCATCAT T           miRNA 5'  GAGGTAGTA G                    T         G TTGTATAGTT           3112111111323333333333 ``` \| SeedType = 3 NonSeedType = -1 \| \| --- \| --- \| --- \| --- \| --- \| --- \| | \| 707 \| WBGene00011970:rlbp-1(7400) \| ``` log2FC=-0.007 padj=0.998 ``` \| [seedLoc = 343,349] Seq=GGGCATAATTTCTCTTTACCATCT \| ```          T  A     TC   TTA       G mRNA  3'  CT CCATT  TCT   ATA CGG  miRNA 5'  GA GGTAG  AGG   TAT GTT           T        T    TTG   A               3115111123511233311161224 ``` \| SeedType = 3 NonSeedType = -1 \| \| --- \| --- \| --- \| --- \| --- \| --- \| | \| 708 \| WBGene00011972:T23G11.7(7404) \| ``` log2FC=-0.218 padj=0.69 ``` \| [seedLoc = 158,163] Seq=TGGTGGGTTAGTCACTACTTCA \| ```                   CTGATTGGGTGGT mRNA  3' ACTTCATCA              miRNA 5' TGAGGTAGT                                AGGTTGTATAGTT           1112111113333333333333 ``` \| SeedType = 3 NonSeedType = -1 \| \| --- \| --- \| --- \| --- \| --- \| --- \| | \| 709 \| WBGene00011977:T24B8.3(7409) \| ``` log2FC=-0.398 padj=0.672 ``` \| [seedLoc = 129,134] Seq=AAGCAATACATCGAATACCTCT \| ```          T      AAG        A   A mRNA  3'  CTCCAT   CT ACATA CGA  miRNA 5'  GAGGTA   GG TGTAT GTT           T      GTA  T     A               31111113331261111131215 ``` \| SeedType = 1 NonSeedType = -1 \| \| --- \| --- \| --- \| --- \| --- \| --- \| | \|  \|  \|  \| [seedLoc = 583,588] Seq=TGCTGATTCGGTTGTACTACCTTT \| ```          T         G TG  TTA    T mRNA  3'  TTCCATCAT T  GC   GTCG  miRNA 5'  GAGGTAGTA G  TG   TAGT           T         G T   TA     T           321111111132342133521123 ``` \| SeedType = 3 NonSeedType = -1 \| \| --- \| --- \| --- \| --- \| --- \| --- \| | \| 710 \| WBGene00011980:T24B8.7(7414) \| ``` log2FC=-0.049 padj=0.968 ``` \| [seedLoc = 1138,1143] Seq=GGTTGTCGAGATATTCTACTTCG \| ```          G       TTATAG        GG mRNA  3'  CTTCATC      AGC TGTT   miRNA 5'  GAGGTAG      TTG ATAG            T       TAGG     T    TT           311211113333541216121233 ``` \| SeedType = 3 NonSeedType = -1 \| \| --- \| --- \| --- \| --- \| --- \| --- \| | \| 711 \| WBGene00011994:samp-1(7421) \| ``` log2FC=0.123 padj=0.918 ``` \| [seedLoc = 227,232] Seq=TTAAATACCTTACTACTTCA \| ```                            AATT mRNA  3' ACTTCATCATTC  CATA     miRNA 5' TGAGGTAGTAGG  GTAT                          TT    AGTT           1112111111216611113333 ``` \| SeedType = 3 NonSeedType = -1 \| \| --- \| --- \| --- \| --- \| --- \| --- \| | \| 712 \| WBGene00012005:jun-1(7429) \|  \| [seedLoc = 24,29] Seq=TATTATCAACTCAAAACTGCCTCA \| ```                   AAAC           T mRNA  3' ACTCCGTCA    TCAAC TATTA  miRNA 5' TGAGGTAGT    GGTTG ATAGT                    A        T     T           1111121113555211116111213 ``` \| SeedType = 2 NonSeedType = 1 \| \| --- \| --- \| --- \| --- \| --- \| --- \| | \| 713 \| WBGene00012033:T26C5.3(7447) \| ``` log2FC=0.063 padj=0.956 ``` \| [seedLoc = 291,294] Seq=TCCTGTGTTCTCTCTTCCTCT \| ```          T    T      CTT     CT mRNA  3'  CTCC TC TCT   GTGTC   miRNA 5'  GAGG AG AGG   TATAG            T    T  T   TTG     TT           3111131161123332121133 ``` \| SeedType = 2 NonSeedType = -1 \| \| --- \| --- \| --- \| --- \| --- \| --- \| | \| 714 \| WBGene00012097:abcf-2(7466) \| ``` log2FC=0.255 padj=0.33 ``` \| [seedLoc = 237,243] Seq=TTTTCATTCATCTTTTACTCTCT \| ```          T   T    T    CTTACTTTT mRNA  3'  CTC CATT TCTA          miRNA 5'  GAG GTAG AGGT                   T        T    TGTATAGTT           31115111231121333333333 ``` \| SeedType = 2 NonSeedType = 2 \| \| --- \| --- \| --- \| --- \| --- \| --- \| | \|  \|  \|  \| [seedLoc = 314,320] Seq=CCCTTTTTTTCCAATTACCCTCC \| ```          C  C      A  TTTTTT  CC mRNA  3'  CT CCATTA CC      TC   miRNA 5'  GA GGTAGT GG      AG            T         A  TTGTAT  TT           31151111213113333331133 ``` \| SeedType = 3 NonSeedType = -1 \| \| --- \| --- \| --- \| --- \| --- \| --- \| | \| 715 \| WBGene00012136:T28F4.1(7504) \| ``` log2FC=-0.117 padj=0.932 ``` \| [seedLoc = 37,42] Seq=TAGGCTACAGTTTTCTTACCTCT \| ```          T       CT        CGGAT mRNA  3'  CTCCATT  TTTGACAT      miRNA 5'  GAGGTAG  AGGTTGTA               T       T         TAGTT           31111112351222111133333 ``` \| SeedType = 1 NonSeedType = 2 \| \| --- \| --- \| --- \| --- \| --- \| --- \| | \|  \|  \|  \| [seedLoc = 65,71] Seq=ATCTATTCATGTTAAATACTCTCT \| ```          T   T   AA   GT  T    TA mRNA  3'  CTC CAT  ATT  AC TATC   miRNA 5'  GAG GTA  TAG  TG ATAG            T       G    GT  T    TT           311151113511233113111133 ``` \| SeedType = 2 NonSeedType = -1 \| \| --- \| --- \| --- \| --- \| --- \| --- \| | \| 716 \| WBGene00012204:mfn-1(7549) \| ``` log2FC=0.127 padj=0.881 ``` \| [seedLoc = 114,120] Seq=GTTTTTTTTTCATATTACTCTCA \| ```              T      ACTTTTTTTTTG mRNA  3' ACTC CATTAT             miRNA 5' TGAG GTAGTA                                 GGTTGTATAGTT           11115111211333333333333 ``` \| SeedType = 2 NonSeedType = -1 \| \| --- \| --- \| --- \| --- \| --- \| --- \| | \| 717 \| WBGene00012205:W02B12.10(7550) \| ``` log2FC=-0.067 padj=0.956 ``` \| [seedLoc = 50,53] Seq=TTTTTCACGAGACTCTCCCTCC \| ```          C    C     AG    CTTTTT mRNA  3'  CTCC TC TC  AGCA       miRNA 5'  GAGG AG AG  TTGT                T    T  T  G     ATAGTT           31111311611341211333333 ``` \| SeedType = 2 NonSeedType = -1 \| \| --- \| --- \| --- \| --- \| --- \| --- \| | \| 718 \| WBGene00012208:W02D9.2(7553) \| ``` log2FC=-0.071 padj=0.955 ``` \| [seedLoc = 143,149] Seq=TAATTCTCCCCCTTTCTACTCTCT \| ```          T   T    TT   CCCTCTTAAT mRNA  3'  CTC CATC  TCC           miRNA 5'  GAG GTAG  AGG                    T        T    TTGTATAGTT           311151111351113333333333 ``` \| SeedType = 2 NonSeedType = -1 \| \| --- \| --- \| --- \| --- \| --- \| --- \| | \| 719 \| WBGene00012216:W02D9.10(7559) \| ``` log2FC=-0.386 padj=0.367 ``` \| [seedLoc = 130,135] Seq=CGATTTTTAAGCCGATACCTCG \| ```          G      AG     ATTT    C mRNA  3'  CTCCAT   CCGA    TTAG  miRNA 5'  GAGGTA   GGTT    AGTT           T      GTA    GTAT                31111113361121333312125 ``` \| SeedType = 1 NonSeedType = 2 \| \| --- \| --- \| --- \| --- \| --- \| --- \| | \| 720 \| WBGene00012256:lpr-5(7590) \| ``` log2FC=0.724 padj=0.023 over-expressed ``` \| [seedLoc = 116,121] Seq=TCTCTTTCAACCCTCGGGTGATTACCTCA \| ```                   GTGGGCTC     TTTCTCT mRNA  3' ACTCCATTA        CCAAC        miRNA 5' TGAGGTAGT        GGTTG                          A            TATAGTT           11111112135444555111113333333 ``` \| SeedType = 1 NonSeedType = 1 \| \| --- \| --- \| --- \| --- \| --- \| --- \| | \| 721 \| WBGene00012275:npr-14(7601) \| ``` log2FC=1.565 padj=1.0 ``` \| [seedLoc = 532,537] Seq=ATGGTGCAGTTGCCAATGATTGGATACCTCC \| ```          C      AGGTTAG  A   TTG     GTA  mRNA  3'  CTCCAT       TA CCG   ACGTG     miRNA 5'  GAGGTA       GT GGT   TGTAT              T               A           AGTT           31111115445554213112554112123336 ``` \| SeedType = 1 NonSeedType = 2 \| \| --- \| --- \| --- \| --- \| --- \| --- \| | \| 722 \| WBGene00012317:ztf-6(7624) \| ``` log2FC=0.195 padj=0.864 ``` \| [seedLoc = 54,60] Seq=TCAAATATTCCTGTACTCTCC \| ```          C   T        T     AACT mRNA  3'  CTC CAT GTCC  TATA     miRNA 5'  GAG GTA TAGG  GTAT              T       G    TT    AGTT           31115111621113621113333 ``` \| SeedType = 2 NonSeedType = -1 \| \| --- \| --- \| --- \| --- \| --- \| --- \| | \|  \|  \|  \| [seedLoc = 508,514] Seq=GGACTTTTTTGTCCAAAATTACCCTCT \| ```          T  C      AAA  TGTTTTT    G mRNA  3'  CT CCATTA   CC       TCAG  miRNA 5'  GA GGTAGT   GG       AGTT           T         A    TTGTAT                 311511112135511333333511124 ``` \| SeedType = 3 NonSeedType = -1 \| \| --- \| --- \| --- \| --- \| --- \| --- \| | \| 723 \| WBGene00012356:W09D6.1(7645) \| ``` log2FC=-0.357 padj=0.773 ``` \| [seedLoc = 233,239] Seq=CATTTTTCGATTTGATCTACCGTCA \| ```             G     TA       TTTTTAC mRNA  3' ACT CCATC  GTTTAGC        miRNA 5' TGA GGTAG  TAGGTTG                                   TATAGTT           1114111115521221213333333 ``` \| SeedType = 3 NonSeedType = 2 \| \| --- \| --- \| --- \| --- \| --- \| --- \| | \|  \|  \|  \| [seedLoc = 247,251] Seq=TCGATTTGATCTACCGTCAC \| ```          C   G     TA       T       mRNA  3'  ACT CCATC  GTTTAGC        miRNA 5'  TGA GGTAG  TAGGTTG                                    TATAGTT           51114111115521221213666666 ``` \| SeedType = 3 NonSeedType = 2 \| \| --- \| --- \| --- \| --- \| --- \| --- \| | \| 724 \| WBGene00012369:W09G3.6(7657) \| ``` log2FC=0.035 padj=0.988 ``` \| [seedLoc = 127,132] Seq=CGTGGAGCGCACTTACTGCCTCA \| ```                       C   AGGTGC mRNA  3' ACTCCGTCATTCA GCG       miRNA 5' TGAGGTAGTAGGT TGT                                 ATAGTT           11111211112115212333333 ``` \| SeedType = 2 NonSeedType = 2 \| \| --- \| --- \| --- \| --- \| --- \| --- \| | \|  \|  \|  \| [seedLoc = 133,137] Seq=CGTGGAGCGCACTTACTGCCTCAT \| ```          T             C   AGGTGC mRNA  3'  ACTCCGTCATTCA GCG       miRNA 5'  TGAGGTAGTAGGT TGT                                  ATAGTT           511111211112115212333333 ``` \| SeedType = 2 NonSeedType = 2 \| \| --- \| --- \| --- \| --- \| --- \| --- \| | \| 725 \| WBGene00012372:W09H1.1(7661) \| ``` log2FC=-1.284 padj=1.0 ``` \| [seedLoc = 119,124] Seq=ATATTTTCAACTCAACTACTTCT \| ```          T        AC     TTTTATA mRNA  3'  CTTCATCA  TCAAC        miRNA 5'  GAGGTAGT  GGTTG                 T        A      TATAGTT           31121111135211113333333 ``` \| SeedType = 3 NonSeedType = 1 \| \| --- \| --- \| --- \| --- \| --- \| --- \| | \| 726 \| WBGene00012374:W09H1.4(7662) \| ``` log2FC=-2.523 padj=1.0 ``` \| [seedLoc = 52,57] Seq=TGACTTTGACTTTTCTTTCAAATTACCTCT \| ```          T        AACTTTCTT      T      T mRNA  3'  CTCCATTA         TTCAGT   TCAG  miRNA 5'  GAGGTAGT         AGGTTG   AGTT           T                       TAT                31111112155555555512112236611125 ``` \| SeedType = 1 NonSeedType = 2 \| \| --- \| --- \| --- \| --- \| --- \| --- \| | \| 727 \| WBGene00012376:nduf-7(7664) \| ``` log2FC=0.091 padj=0.904 ``` \| [seedLoc = 211,216] Seq=TCGAATAGTGTCTATTTTTACCTCA \| ```                 TTT     GTG   AGCT mRNA  3' ACTCCAT   TATCT   ATA     miRNA 5' TGAGGTA   GTAGG   TAT                             TTG   AGTT           1111111555211123331113333 ``` \| SeedType = 1 NonSeedType = -1 \| \| --- \| --- \| --- \| --- \| --- \| --- \| | \| 728 \| WBGene00012405:ztf-25(7677) \| ``` log2FC=0.442 padj=0.902 ``` \| [seedLoc = 130,135] Seq=CTCTATTTTGTCGATACTATCTCA \| ```                    AGCTGTTT    TC mRNA  3' ACTCTATCAT        TATC   miRNA 5' TGAGGTAGTA        ATAG                      GGTTGT      TT           111121111133333355111133 ``` \| SeedType = 2 NonSeedType = -1 \| \| --- \| --- \| --- \| --- \| --- \| --- \| | \| 729 \| WBGene00012435:flh-1(7691) \| ``` log2FC=0.047 padj=0.986 ``` \| [seedLoc = 62,67] Seq=TCCTCCACTCTCCACCTATCTCA \| ```                  CA  TCT  CC  CT mRNA  3' ACTCTATC  CC   CA  TC   miRNA 5' TGAGGTAG  GG   GT  AG                    TA  TT   AT  TT           11112111331133511331133 ``` \| SeedType = 2 NonSeedType = -1 \| \| --- \| --- \| --- \| --- \| --- \| --- \| | \|  \|  \|  \| [seedLoc = 160,165] Seq=TACGATTCAATCTAAATCACATCTACCTCC \| ```          C       TACACTAA       TTAG  T mRNA  3'  CTCCATC        ATCTAAC    CA  miRNA 5'  GAGGTAG        TAGGTTG    GT           T                      TATA  T           311111115555555511121113333113 ``` \| SeedType = 1 NonSeedType = 1 \| \| --- \| --- \| --- \| --- \| --- \| --- \| | \| 730 \| WBGene00012436:flh-3(7692) \| ``` log2FC=0.956 padj=0.668 ``` \| [seedLoc = 228,231] Seq=CATTGTATCGCCAATCTTCCTCC \| ```          C    T  TAA   C       C mRNA  3'  CTCC TC   CCG TATGTTA  miRNA 5'  GAGG AG   GGT GTATAGT           T    T  TA    T       T           31111311335112321121213 ``` \| SeedType = 2 NonSeedType = 2 \| \| --- \| --- \| --- \| --- \| --- \| --- \| | \| 731 \| WBGene00012458:ash-2(7704) \| ``` log2FC=-0.057 padj=0.973 ``` \| [seedLoc = 282,288] Seq=ATTAATTTTTTTTGCTACTCTCC \| ```          C   T        TTTTTAATTA mRNA  3'  CTC CATCGTTT           miRNA 5'  GAG GTAGTAGG                    T            TTGTATAGTT           31115111121223333333333 ``` \| SeedType = 2 NonSeedType = -1 \| \| --- \| --- \| --- \| --- \| --- \| --- \| | \| 732 \| WBGene00012471:Y17G7B.20(7716) \| ``` log2FC=-0.106 padj=0.895 ``` \| [seedLoc = 450,456] Seq=TTCGTTTGGACCCCCCAATACTCTCC \| ```          C   T   AACCCC    GTTTGCTT mRNA  3'  CTC CAT      CCAG         miRNA 5'  GAG GTA      GGTT                  T       GTA       GTATAGTT           31115111333555111233333333 ``` \| SeedType = 2 NonSeedType = 1 \| \| --- \| --- \| --- \| --- \| --- \| --- \| | \| 733 \| WBGene00012514:Y26E6A.3(7736) \| ``` log2FC=0.061 padj=0.97 ``` \| [seedLoc = 120,126] Seq=ATATTTGCAATATTACGCTCT \| ```          T   G             TTATA mRNA  3'  CTC CATTA  TAACGT      miRNA 5'  GAG GTAGT  GTTGTA               T         AG      TAGTT           31114111216621112133333 ``` \| SeedType = 2 NonSeedType = 1 \| \| --- \| --- \| --- \| --- \| --- \| --- \| | \| 734 \| WBGene00012528:pap-1(7742) \| ``` log2FC=-0.062 padj=0.956 ``` \| [seedLoc = 302,307] Seq=TCAAGTATATGCCTTCTACTTCA \| ```                  T    T     AACT mRNA  3' ACTTCATC TCCG ATATG     miRNA 5' TGAGGTAG AGGT TGTAT                      T          AGTT           11121111311125121123333 ``` \| SeedType = 3 NonSeedType = 2 \| \| --- \| --- \| --- \| --- \| --- \| --- \| | \|  \|  \|  \| [seedLoc = 405,411] Seq=TTTTCTTTGACTTCACATACCATCG \| ```          G  A    A  C     TTTCTTTT mRNA  3'  CT CCAT CA TTCAG         miRNA 5'  GA GGTA GT AGGTT                  T                GTATAGTT           3115111151151211233333333 ``` \| SeedType = 3 NonSeedType = 2 \| \| --- \| --- \| --- \| --- \| --- \| --- \| | \|  \|  \|  \| [seedLoc = 471,476] Seq=CTCTGTTACTATCTATCTCT \| ```          T       TA   T      TC mRNA  3'  CTCTATC  TCA   TGTC   miRNA 5'  GAGGTAG  GGT   ATAG            T       TA   TGT    TT           3111211133211366121133 ``` \| SeedType = 2 NonSeedType = 2 \| \| --- \| --- \| --- \| --- \| --- \| --- \| | \| 735 \| WBGene00012559:epg-3(7769) \| ``` log2FC=0.09 padj=0.93 ``` \| [seedLoc = 264,269] Seq=TTCTTCTTCTTTTCCTACCTTT \| ```          T       CTTTTCTTCTTCTT mRNA  3'  TTCCATC               miRNA 5'  GAGGTAG                        T       TAGGTTGTATAGTT           3211111133333333333333 ``` \| SeedType = 3 NonSeedType = -1 \| \| --- \| --- \| --- \| --- \| --- \| --- \| | \| 736 \| WBGene00012584:ceh-100(7779) \| ``` log2FC=-0.013 padj=0.994 ``` \| [seedLoc = 126,132] Seq=TTCCCACCGCCACCACCTCT \| ```          T     C      C  CCCTT  mRNA  3'  CTCCA CA CCG CA       miRNA 5'  GAGGT GT GGT GT                T     A  A   T  ATAGTT           3111113116112311333336 ``` \| SeedType = 2 NonSeedType = 2 \| \| --- \| --- \| --- \| --- \| --- \| --- \| | \| 737 \| WBGene00012621:Y38H6C.8(7801) \| ``` log2FC=-0.077 padj=1.0 ``` \| [seedLoc = 29,34] Seq=TTTATTACAACAAAGTACCTCT \| ```          T      G AA      TATTT mRNA  3'  CTCCAT A  CAACAT      miRNA 5'  GAGGTA T  GTTGTA               T      G AG      TAGTT           3111111313311111133333 ``` \| SeedType = 1 NonSeedType = 1 \| \| --- \| --- \| --- \| --- \| --- \| --- \| | \| 738 \| WBGene00012637:Y38H8A.3(7815) \| ``` log2FC=-0.289 padj=0.362 ``` \| [seedLoc = 27,32] Seq=AGACTTGAAATATAACTGACTACCTTT \| ```          T        G        AAGT    A mRNA  3'  TTCCATCA TCAATATA    TCAG  miRNA 5'  GAGGTAGT GGTTGTAT    AGTT           T        A                            321111111321112111554511125 ``` \| SeedType = 3 NonSeedType = 1 \| \| --- \| --- \| --- \| --- \| --- \| --- \| | \| 739 \| WBGene00012648:Y39A1A.9(7825) \| ``` log2FC=0.887 padj=0.106 ``` \| [seedLoc = 214,219] Seq=ACATATACAAAATTAGTAAAGGTATACCTCC \| ```          C      ATGGAAATGATTAA       ACA mRNA  3'  CTCCAT              AACATAT    miRNA 5'  GAGGTA              TTGTATA             T      GTAGG                GTT           3111111333335554555551111111333 ``` \| SeedType = 1 NonSeedType = 1 \| \| --- \| --- \| --- \| --- \| --- \| --- \| | \| 740 \| WBGene00012717:Y39E4B.6(7864) \| ``` log2FC=0.262 padj=0.818 ``` \| [seedLoc = 254,259] Seq=TCTTGTGCGGTACTCGCTATCTCT \| ```          T        C   TG       CT mRNA  3'  CTCTATCG TCA  GCGTGTT   miRNA 5'  GAGGTAGT GGT  TGTATAG            T        A            TT           311121112321154212121233 ``` \| SeedType = 2 NonSeedType = 2 \| \| --- \| --- \| --- \| --- \| --- \| --- \| | \|  \|  \|  \| [seedLoc = 861,866] Seq=TACTGTTTCTTCCCTGAGGCTGGTACCTCC \| ```          C      G   G  GTCCCTTCTT     T mRNA  3'  CTCCAT GTC GA          TGTCA  miRNA 5'  GAGGTA TAG TT          ATAGT           T      G   G  GT             T           311111132113213355555555121113 ``` \| SeedType = 1 NonSeedType = -1 \| \| --- \| --- \| --- \| --- \| --- \| --- \| | \| 741 \| WBGene00012723:Y39G8B.2(7870) \| ``` log2FC=0.536 padj=0.801 ``` \| [seedLoc = 181,185] Seq=TGCATTTTTCTCTACCACCTCT \| ```          T     C     CTTTTTACGT mRNA  3'  CTCCA CATCT           miRNA 5'  GAGGT GTAGG                    T     A     TTGTATAGTT           3111113111123333333333 ``` \| SeedType = 2 NonSeedType = -1 \| \| --- \| --- \| --- \| --- \| --- \| --- \| | \| 742 \| WBGene00012803:Y43F4B.5(7919) \| ``` log2FC=0.296 padj=0.19 ``` \| [seedLoc = 66,72] Seq=AAACTGGAAGTGTCACTACTCTCC \| ```          C   T     CTGTGAAG     A mRNA  3'  CTC CATCA        GTCAA  miRNA 5'  GAG GTAGT        TAGTT           T         AGGTTGTA                 311151111133333333211115 ``` \| SeedType = 2 NonSeedType = -1 \| \| --- \| --- \| --- \| --- \| --- \| --- \| | \| 743 \| WBGene00012806:Y43F4B.9(7921) \| ``` log2FC=0.114 padj=0.934 ``` \| [seedLoc = 918,924] Seq=CGCGAAGTAGCCGGGTACCATCG \| ```          G  A    G G     GAAG  C mRNA  3'  CT CCAT G CCGAT    CG  miRNA 5'  GA GGTA T GGTTG    GT           T       G A     TATA  T           31151111323112123333123 ``` \| SeedType = 3 NonSeedType = 2 \| \| --- \| --- \| --- \| --- \| --- \| --- \| | \| 744 \| WBGene00012825:Y43F8C.3(7935) \| ``` log2FC=-0.655 padj=0.152 ``` \| [seedLoc = 372,377] Seq=AAAAATGTAACTTTTTCTGCCTCT \| ```          T       TTT         AAAA mRNA  3'  CTCCGTC   TTCAATGTA     miRNA 5'  GAGGTAG   AGGTTGTAT              T       T           AGTT           311112113551211122113333 ``` \| SeedType = 2 NonSeedType = 1 \| \| --- \| --- \| --- \| --- \| --- \| --- \| | \| 745 \| WBGene00012854:Y44A6D.3(7952) \| ``` log2FC=-0.001 padj=1.0 ``` \| [seedLoc = 256,261] Seq=TGACATTATAAAAGATCTACTTCA \| ```                  TAGAA     TA   T mRNA  3' ACTTCATC     AATAT  CAG  miRNA 5' TGAGGTAG     TTGTA  GTT                   TAGG      TA               111211113333511211331125 ``` \| SeedType = 3 NonSeedType = -1 \| \| --- \| --- \| --- \| --- \| --- \| --- \| | \| 746 \| WBGene00012867:seld-1(7963) \| ``` log2FC=0.777 padj=0.008 over-expressed ``` \| [seedLoc = 110,115] Seq=TTTTCTTCAATTCTTTACATACCTCA \| ```                 A    TCT    TTCTTTT mRNA  3' ACTCCAT CATT   TAAC        miRNA 5' TGAGGTA GTAG   GTTG                                    TATAGTT           11111115111255521113333333 ``` \| SeedType = 1 NonSeedType = 1 \| \| --- \| --- \| --- \| --- \| --- \| --- \| | \|  \|  \|  \| [seedLoc = 126,131] Seq=TCTTTACATACCTCAT \| ```          T       A     CT         mRNA  3'  ACTCCAT CATTT           miRNA 5'  TGAGGTA GTAGG                                  TTGTATAGTT           511111115111223366666666 ``` \| SeedType = 1 NonSeedType = -1 \| \| --- \| --- \| --- \| --- \| --- \| --- \| | \| 747 \| WBGene00012894:tbc-17(7978) \| ``` log2FC=-0.023 padj=0.99 ``` \| [seedLoc = 52,57] Seq=CACATCTCGACGGCTACTTCG \| ```          G        G     TCTACAC mRNA  3'  CTTCATCG  CAGC        miRNA 5'  GAGGTAGT  GTTG                 T        AG    TATAGTT           3112111123611213333333 ``` \| SeedType = 3 NonSeedType = -1 \| \| --- \| --- \| --- \| --- \| --- \| --- \| | \| 748 \| WBGene00012908:Y46G5A.18(7991) \| ``` log2FC=-0.041 padj=0.977 ``` \| [seedLoc = 231,237] Seq=TGGAAACTAACTTTTTTTGAAATTACCGTCA \| ```             G      AAGTTTTT      CAAAGGT mRNA  3' ACT CCATTA        TTCAAT        miRNA 5' TGA GGTAGT        AGGTTG                                         TATAGTT           1114111121554555551211123333333 ``` \| SeedType = 3 NonSeedType = 1 \| \| --- \| --- \| --- \| --- \| --- \| --- \| | \| 749 \| WBGene00012929:rsks-1(8008) \| ``` log2FC=-0.032 padj=0.982 ``` \| [seedLoc = 180,186] Seq=TTTTATTAACTTTACACTCT \| ```          T   A                TT mRNA  3'  CTC CAT  TTCAAT TATT   miRNA 5'  GAG GTA  AGGTTG ATAG            T       GT      T    TT           31115111661211126111233 ``` \| SeedType = 2 NonSeedType = 1 \| \| --- \| --- \| --- \| --- \| --- \| --- \| | \| 750 \| WBGene00012932:Y47D3A.21(8011) \| ``` log2FC=-0.031 padj=0.982 ``` \| [seedLoc = 385,390] Seq=TTTGTTACAATTTTTCTCTATCTCC \| ```          C       TCTT        TGTTT mRNA  3'  CTCTATC    TTTAACAT      miRNA 5'  GAGGTAG    AGGTTGTA               T       T           TAGTT           3111211135551221111133333 ``` \| SeedType = 2 NonSeedType = 1 \| \| --- \| --- \| --- \| --- \| --- \| --- \| | \| 751 \| WBGene00012942:Y47D3B.6(8020) \| ``` log2FC=-1.191 padj=8.67143556984441E-7 ``` \| [seedLoc = 103,109] Seq=TCCTGGATACCACTACACTCT \| ```          T   A         TAG    CT mRNA  3'  CTC CATCA CCA    GTC   miRNA 5'  GAG GTAGT GGT    TAG            T         A   TGTA   TT           31115111116111333621133 ``` \| SeedType = 2 NonSeedType = 1 \| \| --- \| --- \| --- \| --- \| --- \| --- \| | \| 752 \| WBGene00012982:jmjd-2(8047) \| ``` log2FC=0.002 padj=0.999 ``` \| [seedLoc = 105,111] Seq=CTCTTTCCCGCCCGTCTACCCTCA \| ```             C     TGC   CCCTTTCTC mRNA  3' ACT CCATC   CCG          miRNA 5' TGA GGTAG   GGT                            TA    TGTATAGTT           111511111335112333333333 ``` \| SeedType = 3 NonSeedType = 2 \| \| --- \| --- \| --- \| --- \| --- \| --- \| | \|  \|  \|  \| [seedLoc = 121,125] Seq=CCGCCCGTCTACCCTCAA \| ```          A   C     TGC   CC        mRNA  3'  ACT CCATC   CCG          miRNA 5'  TGA GGTAG   GGT                             TA    TGTATAGTT           5111511111335112336666666 ``` \| SeedType = 3 NonSeedType = 2 \| \| --- \| --- \| --- \| --- \| --- \| --- \| | \| 753 \| WBGene00013011:snx-14(8071) \| ``` log2FC=0.044 padj=0.986 ``` \| [seedLoc = 110,115] Seq=CTCTATGCGATCTTTCGAATGTACCTCC \| ```          C      GTAAGCTT           TC mRNA  3'  CTCCAT        TCTAGCGTATC   miRNA 5'  GAGGTA        AGGTTGTATAG            T      GT                 TT           3111111335545551121212111133 ``` \| SeedType = 1 NonSeedType = 2 \| \| --- \| --- \| --- \| --- \| --- \| --- \| | \| 754 \| WBGene00013049:Y50E8A.6(8097) \| ``` log2FC=0.078 padj=0.992 ``` \| [seedLoc = 244,248] Seq=CGTCGCCTGATTGACCACCTCGG \| ```          G      C  GTTAG    CTGC      mRNA  3'  GCTCCA CA     TCCG          miRNA 5'  TGAGGT GT     AGGT                          A           TGTATAGTT           4211111311455541112333366666 ``` \| SeedType = 2 NonSeedType = 2 \| \| --- \| --- \| --- \| --- \| --- \| --- \| | \| 755 \| WBGene00013058:clec-247(8102) \| ``` log2FC=0.709 padj=0.444 ``` \| [seedLoc = 88,94] Seq=TCTGAAACAGGAGATCTACACTCT \| ```          T   A    TAGAG    AAGTCT mRNA  3'  CTC CATC     GACA       miRNA 5'  GAG GTAG     TTGT                T        TAGG     ATAGTT           311151111333342111333333 ``` \| SeedType = 2 NonSeedType = -1 \| \| --- \| --- \| --- \| --- \| --- \| --- \| | \| 756 \| WBGene00013082:grdn-1(8114) \| ``` log2FC=-0.376 padj=0.289 ``` \| [seedLoc = 103,109] Seq=TTTTGAAAAATTTTTTCACCTCA \| ```                C  T     AAAGTTTT mRNA  3' ACTCCA TT TTTAA         miRNA 5' TGAGGT AG AGGTT                           T     GTATAGTT           11111151231221133333333 ``` \| SeedType = 2 NonSeedType = 2 \| \| --- \| --- \| --- \| --- \| --- \| --- \| | \| 757 \| WBGene00013106:set-26(8133) \| ``` log2FC=-0.255 padj=0.466 ``` \| [seedLoc = 53,58] Seq=TCCCATATCATCTCTTTCTATCTCG \| ```          G       TTTC    C    CCCT mRNA  3'  CTCTATC    TCTA TATA     miRNA 5'  GAGGTAG    AGGT GTAT              T       T       T    AGTT           3111211135551121321113333 ``` \| SeedType = 2 NonSeedType = 2 \| \| --- \| --- \| --- \| --- \| --- \| --- \| | \| 758 \| WBGene00013111:sta-1(8137) \| ``` log2FC=-0.146 padj=0.903 ``` \| [seedLoc = 34,39] Seq=AACAGTCCCCCCAATTACCTCT \| ```          T        A  CCCCTGACAA mRNA  3'  CTCCATTA CC           miRNA 5'  GAGGTAGT GG                    T        A  TTGTATAGTT           3111111213113333333333 ``` \| SeedType = 1 NonSeedType = -1 \| \| --- \| --- \| --- \| --- \| --- \| --- \| | \| 759 \| WBGene00013136:Y53C10A.5(8154) \| ``` log2FC=0.216 padj=0.715 ``` \| [seedLoc = 74,80] Seq=TACTTGCGCTAGATACTACCATCA \| ```             A       AGA           T mRNA  3' ACT CCATCAT   TC GCGT TCA  miRNA 5' TGA GGTAGTA   GG TGTA AGT                           T    T   T           11151111111545216212161113 ``` \| SeedType = 3 NonSeedType = -1 \| \| --- \| --- \| --- \| --- \| --- \| --- \| | \| 760 \| WBGene00013140:mop-25.2(8157) \| ``` log2FC=-0.022 padj=0.989 ``` \| [seedLoc = 189,194] Seq=TAAGATACAGTAGTACCTCC \| ```          C      G          GAAT mRNA  3'  CTCCAT AT  GACATA     miRNA 5'  GAGGTA TA  TTGTAT              T      G  GG      AGTT           3111111311662111113333 ``` \| SeedType = 1 NonSeedType = -1 \| \| --- \| --- \| --- \| --- \| --- \| --- \| | \|  \|  \|  \| [seedLoc = 255,261] Seq=AGGCATCGATCTTTACCGTCT \| ```          T  G                   A mRNA  3'  CT CCATT TCTAGC TA CGG  miRNA 5'  GA GGTAG AGGTTG AT GTT           T        T      T  A               311411112611212161161225 ``` \| SeedType = 3 NonSeedType = 2 \| \| --- \| --- \| --- \| --- \| --- \| --- \| | \| 761 \| WBGene00013178:Y53H1A.2(8184) \| ``` log2FC=-0.012 padj=0.996 ``` \| [seedLoc = 96,102] Seq=TTCCCAGTGGCCTTATTCATACGCTCA \| ```              G   ACTTAT     TGACCCTT mRNA  3' ACTC CAT      TCCGG         miRNA 5' TGAG GTA      AGGTT                          GT         GTATAGTT           111141113355551112233333333 ``` \| SeedType = 2 NonSeedType = 2 \| \| --- \| --- \| --- \| --- \| --- \| --- \| | \| 762 \| WBGene00013189:Y54E2A.4(8189) \| ``` log2FC=-0.008 padj=0.994 ``` \| [seedLoc = 331,336] Seq=GTTGCAGCCCCCTAAAATTATGTCTGCCTCC \| ```          C       TGTATTAAAATCCC       TG    mRNA  3'  CTCCGTC              CCGACGT      miRNA 5'  GAGGTAG              GGTTGTA               T       TA                   TAGTT           3111121133555555555555112112133666 ``` \| SeedType = 2 NonSeedType = 2 \| \| --- \| --- \| --- \| --- \| --- \| --- \| | \| 763 \| WBGene00013196:tbc-20(8192) \| ``` log2FC=-0.026 padj=0.994 ``` \| [seedLoc = 147,152] Seq=CCAGCAAAAATCGCTACCTCC \| ```          C             AAACGACC mRNA  3'  CTCCATCG CTAA         miRNA 5'  GAGGTAGT GGTT                  T        A    GTATAGTT           3111111126121133333333 ``` \| SeedType = 1 NonSeedType = 2 \| \| --- \| --- \| --- \| --- \| --- \| --- \| | \| 764 \| WBGene00013219:Y54G11A.11(8212) \| ``` log2FC=0.12 padj=0.89 ``` \| [seedLoc = 188,194] Seq=ATTCTCACGAGAATGGTATCCTCG \| ```          G    T  G  AAG    CTCTTA mRNA  3'  CTCC AT GT   AGCA       miRNA 5'  GAGG TA TA   TTGT                T       G  GG     ATAGTT           311115113213341211333333 ``` \| SeedType = 2 NonSeedType = -1 \| \| --- \| --- \| --- \| --- \| --- \| --- \| | \| 765 \| WBGene00013243:Y56A3A.31(8232) \| ``` log2FC=0.065 padj=0.965 ``` \| [seedLoc = 35,40] Seq=TGTTAATTGGCTTGAAAATACCTCA \| ```                 AAAA      TTAATTGT mRNA  3' ACTCCAT    GTTCGG         miRNA 5' TGAGGTA    TAGGTT                         G         GTATAGTT           1111111355521212233333333 ``` \| SeedType = 1 NonSeedType = 2 \| \| --- \| --- \| --- \| --- \| --- \| --- \| | \| 766 \| WBGene00013267:Y57A10A.27(8249) \| ``` log2FC=-0.054 padj=0.959 ``` \| [seedLoc = 155,160] Seq=GGAAATTTTGAATTTTACCTCA \| ```                  TTAAGTTTTAAAGG mRNA  3' ACTCCATT               miRNA 5' TGAGGTAG                                TAGGTTGTATAGTT           1111111233333333333333 ``` \| SeedType = 1 NonSeedType = -1 \| \| --- \| --- \| --- \| --- \| --- \| --- \| | \| 767 \| WBGene00013273:marc-5(8254) \|  \| [seedLoc = 120,125] Seq=AAAACCACATTTTCTTCTACTTCT \| ```          T       T   TTT   CCAAAA mRNA  3'  CTTCATC TCT   ACA       miRNA 5'  GAGGTAG AGG   TGT                T       T   T     ATAGTT           311211113112355111333333 ``` \| SeedType = 3 NonSeedType = -1 \| \| --- \| --- \| --- \| --- \| --- \| --- \| | \| 768 \| WBGene00013307:Y57G11C.9(8274) \| ``` log2FC=-0.208 padj=0.628 ``` \| [seedLoc = 874,878] Seq=TGAAGGAAGAGCTTCCTCAA \| ```          A     T   AGAAGGAAGT    mRNA  3'  ACTCC TCG              miRNA 5'  TGAGG AGT                             T   AGGTTGTATAGTT           51111131123333333333666 ``` \| SeedType = 2 NonSeedType = -1 \| \| --- \| --- \| --- \| --- \| --- \| --- \| | \|  \|  \|  \| [seedLoc = 1558,1564] Seq=AAGCTGGATTCTCTACTCTCC \| ```          C   T        TAG       A mRNA  3'  CTC CATC TCT     GTCGA  miRNA 5'  GAG GTAG AGG     TAGTT           T        T   TTGTA                 311151111611233366211215 ``` \| SeedType = 2 NonSeedType = -1 \| \| --- \| --- \| --- \| --- \| --- \| --- \| | \| 769 \| WBGene00013340:tcc-1(8296) \| ``` log2FC=0.353 padj=0.074 ``` \| [seedLoc = 27,32] Seq=CTCCAGATTTCCTTCAATACCTCC \| ```          C      AACT   TT  GACCTC mRNA  3'  CTCCAT    TCC  TA       miRNA 5'  GAGGTA    AGG  GT                T      GT     TT  ATAGTT           311111133551113321333333 ``` \| SeedType = 1 NonSeedType = -1 \| \| --- \| --- \| --- \| --- \| --- \| --- \| | \| 770 \| WBGene00013405:tdpt-1(8335) \| ``` log2FC=-0.244 padj=0.709 ``` \| [seedLoc = 87,92] Seq=CACTATTTTTACCCAGGAATTACCTCC \| ```          C        AGGAC   TTTT     C mRNA  3'  CTCCATTA     CCA    TATCA  miRNA 5'  GAGGTAGT     GGT    ATAGT           T        A       TGT      T           311111121344551113335111113 ``` \| SeedType = 1 NonSeedType = 1 \| \| --- \| --- \| --- \| --- \| --- \| --- \| | \| 771 \| WBGene00013462:micu-1(8367) \| ``` log2FC=-0.125 padj=0.886 ``` \| [seedLoc = 991,996] Seq=ATAGATGCGAAGCTACTTCA \| ```                   A        GATA mRNA  3' ACTTCATCG   AGCGTA     miRNA 5' TGAGGTAGT   TTGTAT                       AGG      AGTT           1112111123661212113333 ``` \| SeedType = 3 NonSeedType = -1 \| \| --- \| --- \| --- \| --- \| --- \| --- \| | \|  \|  \|  \| [seedLoc = 1024,1029] Seq=GTCAAAGTGACTGGAATACCTCT \| ```          T      AAGG    TGAAACTG mRNA  3'  CTCCAT    TCAG         miRNA 5'  GAGGTA    GGTT                  T      GTA     GTATAGTT           31111113334211233333333 ``` \| SeedType = 1 NonSeedType = 2 \| \| --- \| --- \| --- \| --- \| --- \| --- \| | \| 772 \| WBGene00013545:Y75B8A.8(8415) \| ``` log2FC=-0.018 padj=0.993 ``` \| [seedLoc = 269,275] Seq=AAGGTTGCAAAATTACACTCC \| ```          C   A     A       TGGAA mRNA  3'  CTC CATTA   AACGT      miRNA 5'  GAG GTAGT   TTGTA               T         AGG     TAGTT           31115111213661112133333 ``` \| SeedType = 2 NonSeedType = 1 \| \| --- \| --- \| --- \| --- \| --- \| --- \| | \| 773 \| WBGene00013659:Y105C5B.20(8474) \| ``` log2FC=-0.765 padj=1.0 ``` \| [seedLoc = 158,164] Seq=GGGAAAATAGCGTTTATACTCTCA \| ```              T   ATT G     AAAGGG mRNA  3' ACTC CAT   T CGATA       miRNA 5' TGAG GTA   A GTTGT                        GT  G     ATAGTT           111151113351312121333333 ``` \| SeedType = 2 NonSeedType = -1 \| \| --- \| --- \| --- \| --- \| --- \| --- \| | \| 774 \| WBGene00013674:Y105E8A.14(8485) \| ``` log2FC=-0.187 padj=0.731 ``` \| [seedLoc = 777,782] Seq=TCTCTCATTCCCTCACACTATCTCC \| ```          C        CAC   CTTACTCTCT mRNA  3'  CTCTATCA   TCC           miRNA 5'  GAGGTAGT   AGG                    T              TTGTATAGTT           3111211115551113333333333 ``` \| SeedType = 2 NonSeedType = -1 \| \| --- \| --- \| --- \| --- \| --- \| --- \| | \| 775 \| WBGene00013766:prmt-1(8554) \| ``` log2FC=0.16 padj=0.735 ``` \| [seedLoc = 121,126] Seq=CGATTGTATATACCTCG \| ```          G                     C mRNA  3'  CTCCAT      ATATGTTAG  miRNA 5'  GAGGTA      TGTATAGTT           T      GTAGGT                     31111116666661211212125 ``` \| SeedType = 1 NonSeedType = -1 \| \| --- \| --- \| --- \| --- \| --- \| --- \| | \| 776 \| WBGene00013796:Y116A8C.19(8569) \| ``` log2FC=0.1 padj=1.0 ``` \| [seedLoc = 39,44] Seq=CAACAATTCAACACCAATTACTACTTCC \| ```          C          AACCA    T  A   C mRNA  3'  CTTCATCATT     CAAC TA CAA  miRNA 5'  GAGGTAGTAG     GTTG AT GTT           T                   T  A               3112111111255555111131131115 ``` \| SeedType = 3 NonSeedType = 1 \| \| --- \| --- \| --- \| --- \| --- \| --- \| | \| 777 \| WBGene00013883:npr-13(8622) \| ``` log2FC=-0.703 padj=1.0 ``` \| [seedLoc = 101,106] Seq=TCCTCCGCTCTTCCTATCTCC \| ```          C       C   T   CC  CT mRNA  3'  CTCTATC TTC  CG  TC   miRNA 5'  GAGGTAG AGG  GT  AG            T       T   TT  AT  TT           3111211131213612331133 ``` \| SeedType = 2 NonSeedType = -1 \| \| --- \| --- \| --- \| --- \| --- \| --- \| | \| 778 \| WBGene00013897:ZC443.2(8634) \| ``` log2FC=-0.633 padj=1.0 ``` \| [seedLoc = 54,59] Seq=GTCATTATTCGTCTACTTCT \| ```          T         G T    TACTG mRNA  3'  CTTCATC T C  TAT      miRNA 5'  GAGGTAG A G  GTA               T       T G TT   TAGTT           3112111161313621133333 ``` \| SeedType = 3 NonSeedType = -1 \| \| --- \| --- \| --- \| --- \| --- \| --- \| | \| 779 \| WBGene00013934:ZK131.11(8654) \| ``` log2FC=-0.433 padj=0.18 ``` \| [seedLoc = 39,44] Seq=AATCACAAAACTCTTCTTACCTCC \| ```          C       CT  T   AACACTAA mRNA  3'  CTCCATT  TC CAA         miRNA 5'  GAGGTAG  AG GTT                  T       T       GTATAGTT           311111123511511133333333 ``` \| SeedType = 1 NonSeedType = 1 \| \| --- \| --- \| --- \| --- \| --- \| --- \| | \| 780 \| WBGene00013998:gtf-2E1(8688) \|  \| [seedLoc = 65,71] Seq=GTATAGAGGATTGTTTACTCTCC \| ```          C   T    TG    G GATATG mRNA  3'  CTC CATT  TTAG A       miRNA 5'  GAG GTAG  GGTT T                T        TA    G ATAGTT           31115111233221231333333 ``` \| SeedType = 2 NonSeedType = 2 \| \| --- \| --- \| --- \| --- \| --- \| --- \| | \|  \|  \|  \| [seedLoc = 84,90] Seq=CTCCAATTCTCCTAATACCGTCA \| ```             G    A    TCTTAACCTC mRNA  3' ACT CCAT ATCC           miRNA 5' TGA GGTA TAGG                            G    TTGTATAGTT           11141111311113333333333 ``` \| SeedType = 3 NonSeedType = -1 \| \| --- \| --- \| --- \| --- \| --- \| --- \| | \| 781 \| WBGene00014051:spv-1(8729) \| ``` log2FC=0.045 padj=0.98 ``` \| [seedLoc = 40,45] Seq=CGCATGTCTTCTTCTACTTCT \| ```          T           TT     A  C mRNA  3'  CTTCATC TTC  C TGT CG  miRNA 5'  GAGGTAG AGG  G ATA GT           T       T   TT T      T           31121111612133161215123 ``` \| SeedType = 3 NonSeedType = -1 \| \| --- \| --- \| --- \| --- \| --- \| --- \| | \| 782 \| WBGene00014095:gdh-1(8762) \| ``` log2FC=0.265 padj=0.358 ``` \| [seedLoc = 32,38] Seq=CATCTCATGGCTCTTGTATCCTCT \| ```          T    T  GT  T     CTCTAC mRNA  3'  CTCC AT  TC CGGTA       miRNA 5'  GAGG TA  AG GTTGT                T       GT        ATAGTT           311115113311512221333333 ``` \| SeedType = 2 NonSeedType = -1 \| \| --- \| --- \| --- \| --- \| --- \| --- \| | \| 783 \| WBGene00014099:ZK836.3(8767) \| ``` log2FC=-0.731 padj=0.77 ``` \| [seedLoc = 162,167] Seq=AAATCAATAAATATTTACCTCG \| ```          G       TATA    ACTAAA mRNA  3'  CTCCATT    AATA       miRNA 5'  GAGGTAG    TTGT                T       TAGG    ATAGTT           3111111233331121333333 ``` \| SeedType = 1 NonSeedType = -1 \| \| --- \| --- \| --- \| --- \| --- \| --- \| | \| 784 \| WBGene00014109:chpf-1(8774) \|  \| [seedLoc = 106,111] Seq=TTGAAGACACAAACTATCTCA \| ```                   AA     GAAGTT mRNA  3' ACTCTATCA  C ACA       miRNA 5' TGAGGTAGT  G TGT                         AG T   ATAGTT           1111211113316111333333 ``` \| SeedType = 2 NonSeedType = -1 \| \| --- \| --- \| --- \| --- \| --- \| --- \| | \| 785 \| WBGene00014112:ZK856.11(8776) \| ``` log2FC=-0.214 padj=0.703 ``` \| [seedLoc = 133,138] Seq=GTTTATGCGCCTGCTGCCTCT \| ```          T                   TG mRNA  3'  CTCCGTCGTCC GCGTATT   miRNA 5'  GAGGTAGTAGG TGTATAG            T           T       TT           3111121121116212111233 ``` \| SeedType = 2 NonSeedType = -1 \| \| --- \| --- \| --- \| --- \| --- \| --- \| | \| 786 \| WBGene00014130:ZK892.6(8789) \| ``` log2FC=-0.803 padj=1.0 ``` \| [seedLoc = 219,223] Seq=GTTTTATCCCCCTCCCTCT \| ```          T    C  CC  TATTTTG    mRNA  3'  CTCC TC  CC           miRNA 5'  GAGG AG  GG                    T    T  TA  TTGTATAGTT           3111131133113333333666 ``` \| SeedType = 2 NonSeedType = -1 \| \| --- \| --- \| --- \| --- \| --- \| --- \| | \| 787 \| WBGene00014163:rnh-1.2(8809) \| ``` log2FC=0.0 padj=1.0 ``` \| [seedLoc = 159,165] Seq=CAATTAGTTAATATTACCATCT \| ```          T  A       AATTG       C mRNA  3'  CT CCATTAT       ATTAA  miRNA 5'  GA GGTAGTA       TAGTT           T          GGTTGTA                 311511112113333366112115 ``` \| SeedType = 3 NonSeedType = -1 \| \| --- \| --- \| --- \| --- \| --- \| --- \| | \| 788 \| WBGene00014198:ZK1053.3(8829) \| ``` log2FC=0.415 padj=1.0 ``` \| [seedLoc = 39,44] Seq=TGCTCAATTTTTCTACCTTT \| ```          T       TT            T mRNA  3'  TTCCATC  TTTAAC   TCG  miRNA 5'  GAGGTAG  AGGTTG   AGT           T       T       TAT   T           32111111351221116661123 ``` \| SeedType = 3 NonSeedType = 1 \| \| --- \| --- \| --- \| --- \| --- \| --- \| | \| 789 \| WBGene00014215:obr-3(8843) \| ``` log2FC=-0.187 padj=0.748 ``` \| [seedLoc = 176,181] Seq=TCCATTTCAACGCTACTTCT \| ```          T              TTTACCT mRNA  3'  CTTCATCG  CAAC        miRNA 5'  GAGGTAGT  GTTG                 T        AG    TATAGTT           3112111126611113333333 ``` \| SeedType = 3 NonSeedType = 1 \| \| --- \| --- \| --- \| --- \| --- \| --- \| | \| 790 \| WBGene00014300:D2023.1(8877) \| ``` log2FC=0.369 padj=0.766 ``` \| [seedLoc = 88,92] Seq=AACAAGAAATTCTTCCTCAC \| ```          C     T  TTAAAG    A      mRNA  3'  ACTCC TC      AACA       miRNA 5'  TGAGG AG      TTGT                      T  TAGG      ATAGTT           5111113113333541111366666 ``` \| SeedType = 2 NonSeedType = 1 \| \| --- \| --- \| --- \| --- \| --- \| --- \| | \|  \|  \|  \| [seedLoc = 467,473] Seq=TTTTGTTCAATTCTACAATACCATCT \| ```          T  A    AA    T    T    TT mRNA  3'  CT CCAT  CATC TAAC TGTT   miRNA 5'  GA GGTA  GTAG GTTG ATAG            T                  T    TT           31151111551111521113121233 ``` \| SeedType = 3 NonSeedType = 1 \| \| --- \| --- \| --- \| --- \| --- \| --- \| | \| 791 \| WBGene00015007:ain-2(8900) \| ``` log2FC=-0.026 padj=0.985 ``` \| [seedLoc = 401,407] Seq=TATCCGATTTTTTCTTGTTTAATTACCATCT \| ```          T  A      ATTTGTTCTTT      C       mRNA  3'  CT CCATTA           TTTAGC TAT    miRNA 5'  GA GGTAGT           AGGTTG ATA             T                          T   GTT           3115111121555545555551221213111666 ``` \| SeedType = 3 NonSeedType = 2 \| \| --- \| --- \| --- \| --- \| --- \| --- \| | \| 792 \| WBGene00015117:B0294.1(8978) \| ``` log2FC=-0.686 padj=0.792 ``` \| [seedLoc = 48,53] Seq=CTCCAAACAACCTCCGAACAATCTACTTCA \| ```                  TAACAAGCC       AACCTC mRNA  3' ACTTCATC         TCCAACA       miRNA 5' TGAGGTAG         AGGTTGT                        T               ATAGTT           111211113555554551111111333333 ``` \| SeedType = 3 NonSeedType = 1 \| \| --- \| --- \| --- \| --- \| --- \| --- \| | \| 793 \| WBGene00015235:cdc-26(9071) \| ``` log2FC=-0.035 padj=0.986 ``` \| [seedLoc = 52,57] Seq=CTCCCGGCAATCCTACTTCA \| ```                          GCCCTC mRNA  3' ACTTCATC  CTAACG       miRNA 5' TGAGGTAG  GGTTGT                        TA      ATAGTT           1112111166121112333333 ``` \| SeedType = 3 NonSeedType = 1 \| \| --- \| --- \| --- \| --- \| --- \| --- \| | \| 794 \| WBGene00015251:B0546.4(9081) \| ``` log2FC=-0.668 padj=0.459 ``` \| [seedLoc = 156,161] Seq=TTCTGTACACTTTACCTCT \| ```          T                   TT mRNA  3'  CTCCAT  TTC ACATGTC   miRNA 5'  GAGGTA  AGG TGTATAG            T      GT   T       TT           3111111661216111121133 ``` \| SeedType = 1 NonSeedType = -1 \| \| --- \| --- \| --- \| --- \| --- \| --- \| | \| 795 \| WBGene00015293:C01C4.3(9100) \| ``` log2FC=-0.138 padj=0.946 ``` \| [seedLoc = 72,77] Seq=TTGAAAGCCACCCATTATTATCCTACTTCA \| ```                  CTATTATTAC   C  AAAGTT mRNA  3' ACTTCATC          CCA CG       miRNA 5' TGAGGTAG          GGT GT                        TA           T  ATAGTT           111211113355555555111312333333 ``` \| SeedType = 3 NonSeedType = 1 \| \| --- \| --- \| --- \| --- \| --- \| --- \| | \|  \|  \|  \| [seedLoc = 279,284] Seq=TACTATGTAATGATAGACACTGCCTCA \| ```                   CAGATAG          T mRNA  3' ACTCCGTCA       TAATGTATCA  miRNA 5' TGAGGTAGT       GTTGTATAGT                    AG               T           111112111334555421122111113 ``` \| SeedType = 2 NonSeedType = -1 \| \| --- \| --- \| --- \| --- \| --- \| --- \| | \|  \|  \|  \| [seedLoc = 649,654] Seq=TCTTATTTCATCCACTTCCTCAT \| ```          T     T      T  TT    CT mRNA  3'  ACTCC TCA CC AC  TATT   miRNA 5'  TGAGG AGT GG TG  ATAG                  T   A  T  T     TT           511111311161131135111233 ``` \| SeedType = 2 NonSeedType = -1 \| \| --- \| --- \| --- \| --- \| --- \| --- \| | \| 796 \| WBGene00015301:C01F1.6(9109) \| ``` log2FC=-0.311 padj=0.475 ``` \| [seedLoc = 256,261] Seq=CTCTATGTATTGTATTTACCTCC \| ```          C     T    G T       TC mRNA  3'  CTCCA TTAT T ATGTATC   miRNA 5'  GAGGT AGTA G TGTATAG            T          G T       TT           31111151211323122111133 ``` \| SeedType = 2 NonSeedType = -1 \| \| --- \| --- \| --- \| --- \| --- \| --- \| | \| 797 \| WBGene00015303:rga-6(9111) \| ``` log2FC=0.061 padj=0.994 ``` \| [seedLoc = 196,202] Seq=TTATTTATTTATATTAACCTCA \| ```               A     ATT    TTATT mRNA  3' ACTCC ATTAT    TAT      miRNA 5' TGAGG TAGTA    GTA                          GGTT   TAGTT           11111511211333621133333 ``` \| SeedType = 2 NonSeedType = -1 \| \| --- \| --- \| --- \| --- \| --- \| --- \| | \| 798 \| WBGene00015308:ddi-1(9115) \|  \| [seedLoc = 120,126] Seq=TTCTTTTTTTTCTGCAAATACACTCT \| ```          T   A   AAA     TTTTTTTCTT mRNA  3'  CTC CAT   CGTCT           miRNA 5'  GAG GTA   GTAGG                    T               TTGTATAGTT           31115111555121123333333333 ``` \| SeedType = 2 NonSeedType = -1 \| \| --- \| --- \| --- \| --- \| --- \| --- \| | \| 799 \| WBGene00015328:C02B10.3(9128) \| ``` log2FC=-0.064 padj=0.961 ``` \| [seedLoc = 541,546] Seq=ATATATTTAAGCACTGCCTCT \| ```          T          G   T   ATA mRNA  3'  CTCCGTCA C AAT TAT    miRNA 5'  GAGGTAGT G TTG ATA             T        A G   T   GTT           3111121116131123111333 ``` \| SeedType = 2 NonSeedType = -1 \| \| --- \| --- \| --- \| --- \| --- \| --- \| | \| 800 \| WBGene00015330:C02B10.5(9130) \| ``` log2FC=0.127 padj=0.9 ``` \| [seedLoc = 215,220] Seq=GAACTCAGCTCTACCTTC \| ```          C                     G mRNA  3'  TTCCATC  TCGAC   TCAA  miRNA 5'  GAGGTAG  GGTTG   AGTT           T       TA     TAT                32111111662121166611114 ``` \| SeedType = 3 NonSeedType = 2 \| \| --- \| --- \| --- \| --- \| --- \| --- \| | \| 801 \| WBGene00015418:pac-1(9190) \| ``` log2FC=-0.059 padj=0.966 ``` \| [seedLoc = 51,57] Seq=AGAAGCGCCTCCTCTACCTGC \| ```          CG          TC  CGAAGA mRNA  3'   TCCATC TCC  CG       miRNA 5'   AGGTAG AGG  GT                TG      T   TT  ATAGTT           3311111161113312333333 ``` \| SeedType = 3 NonSeedType = -1 \| \| --- \| --- \| --- \| --- \| --- \| --- \| | \|  \|  \|  \| [seedLoc = 451,456] Seq=CAGAATATCATCCTTTCGTACCTCC \| ```          C      GCTT   T  T   AGAC mRNA  3'  CTCCAT    TCC AC ATA     miRNA 5'  GAGGTA    AGG TG TAT              T      GT     T      AGTT           3111111335511131151113333 ``` \| SeedType = 1 NonSeedType = -1 \| \| --- \| --- \| --- \| --- \| --- \| --- \| | \| 802 \| WBGene00015429:sor-3(9195) \| ``` log2FC=0.015 padj=0.995 ``` \| [seedLoc = 100,105] Seq=GGGTATATCTCTTTTCATACCTCG \| ```          G      ACTT   TC     G G mRNA  3'  CTCCAT    TTC  TATAT G  miRNA 5'  GAGGTA    AGG  GTATA T           T      GT     TT     G T           311111133551213321111323 ``` \| SeedType = 1 NonSeedType = -1 \| \| --- \| --- \| --- \| --- \| --- \| --- \| | \| 803 \| WBGene00015484:atgl-1(9230) \| ``` log2FC=-0.381 padj=0.118 ``` \| [seedLoc = 492,497] Seq=TTACCAATAACCCTTACCTCT \| ```          T       C       ACCATT mRNA  3'  CTCCATT  CCAATA       miRNA 5'  GAGGTAG  GGTTGT                T       TA      ATAGTT           3111111236111121333333 ``` \| SeedType = 1 NonSeedType = 1 \| \| --- \| --- \| --- \| --- \| --- \| --- \| | \| 804 \| WBGene00015494:C05E11.3(9235) \| ``` log2FC=-0.432 padj=0.789 ``` \| [seedLoc = 277,282] Seq=TACTTCACCACTTTACCTCT \| ```          T            C  CT   T mRNA  3'  CTCCAT  TTCA CA  TCA  miRNA 5'  GAGGTA  AGGT GT  AGT           T      GT    T  AT   T           3111111661211311331113 ``` \| SeedType = 1 NonSeedType = 2 \| \| --- \| --- \| --- \| --- \| --- \| --- \| | \| 805 \| WBGene00015501:C06A5.3(9240) \| ``` log2FC=-0.098 padj=0.934 ``` \| [seedLoc = 171,176] Seq=CGACTGTCTACGTTCACTACCTCA \| ```                      T   TC      C mRNA  3' ACTCCATCA CT GCA  TGTCAG  miRNA 5' TGAGGTAGT GG TGT  ATAGTT                    A  T                       1111111116123211551211125 ``` \| SeedType = 1 NonSeedType = -1 \| \| --- \| --- \| --- \| --- \| --- \| --- \| | \| 806 \| WBGene00015504:C06A5.6(9242) \| ``` log2FC=0.134 padj=0.842 ``` \| [seedLoc = 138,143] Seq=CACGGAGATGATCCCTATCTCG \| ```          G       C       GAGG  C mRNA  3'  CTCTATC  CTAGTA    CA  miRNA 5'  GAGGTAG  GGTTGT    GT           T       TA      ATA   T           31112111361212213334113 ``` \| SeedType = 2 NonSeedType = 2 \| \| --- \| --- \| --- \| --- \| --- \| --- \| | \| 807 \| WBGene00015517:C06A8.8(9255) \| ``` log2FC=-0.098 padj=0.966 ``` \| [seedLoc = 234,239] Seq=TATTATGCATTTTATGCTTCAACTATCTCA \| ```                   AC    TATTTT        T mRNA  3' ACTCTATCA  TTCG      ACGTATTA  miRNA 5' TGAGGTAGT  AGGT      TGTATAGT                                        T           111121111551212555555112111213 ``` \| SeedType = 2 NonSeedType = 2 \| \| --- \| --- \| --- \| --- \| --- \| --- \| | \| 808 \| WBGene00015519:doxa-1(9257) \| ``` log2FC=-0.252 padj=0.819 ``` \| [seedLoc = 281,287] Seq=TCCCATAACATTTTGTGTGTAAATACCATCC \| ```          C  A    AAATGTGTGTTTTA     CCCT   mRNA  3'  CT CCAT              CAATA       miRNA 5'  GA GGTA              GTTGT                T       GTAG               ATAGTT           311511113333454545555511121333366 ``` \| SeedType = 3 NonSeedType = 1 \| \| --- \| --- \| --- \| --- \| --- \| --- \| | \| 809 \| WBGene00015538:sams-3(9267) \| ``` log2FC=-0.073 padj=0.933 ``` \| [seedLoc = 127,133] Seq=ACCTGTATTTCCGCAATATCCTCG \| ```          G    T  AACG  TT      CA mRNA  3'  CTCC AT    CC  TATGTC   miRNA 5'  GAGG TA    GG  GTATAG            T       GTA   TT      TT           311115113334113321121133 ``` \| SeedType = 2 NonSeedType = -1 \| \| --- \| --- \| --- \| --- \| --- \| --- \| | \| 810 \| WBGene00015552:C06G3.6(9277) \| ``` log2FC=-0.224 padj=0.715 ``` \| [seedLoc = 47,53] Seq=CAACTTCCATCTTGTACTCTCA \| ```              T        T  CT     C mRNA  3' ACTC CAT GTTC AC   TCAA  miRNA 5' TGAG GTA TAGG TG   AGTT                   G    T  TAT                111151116212131133611115 ``` \| SeedType = 2 NonSeedType = -1 \| \| --- \| --- \| --- \| --- \| --- \| --- \| | \| 811 \| WBGene00015569:C07D10.5(9289) \| ``` log2FC=-0.069 padj=0.982 ``` \| [seedLoc = 79,84] Seq=AAAAATAACTTATTGTTTAAATGATACCTCG \| ```          G      AGTAAATTTGT         AAAA   mRNA  3'  CTCCAT           TATTCAATA       miRNA 5'  GAGGTA           GTAGGTTGT                T                          ATAGTT           311111154555555545211211121333366 ``` \| SeedType = 1 NonSeedType = 1 \| \| --- \| --- \| --- \| --- \| --- \| --- \| | \|  \|  \|  \| [seedLoc = 99,104] Seq=TCGAATGCTTTTCTTCTACTTCA \| ```                  T   TTT    AGCT mRNA  3' ACTTCATC TCT   CGTA     miRNA 5' TGAGGTAG AGG   GTAT                      T   TT     AGTT           11121111311233512113333 ``` \| SeedType = 3 NonSeedType = -1 \| \| --- \| --- \| --- \| --- \| --- \| --- \| | \| 812 \| WBGene00015615:fbxc-48(9310) \| ``` log2FC=-0.681 padj=0.798 ``` \| [seedLoc = 95,100] Seq=CACTGTCGGAGTTTACGACTGCCTCC \| ```          C        GCAT  G  GC     C mRNA  3'  CTCCGTCA    TT AG  TGTCA  miRNA 5'  GAGGTAGT    AG TT  ATAGT           T              G  GT     T           31111211145551231233121113 ``` \| SeedType = 2 NonSeedType = -1 \| \| --- \| --- \| --- \| --- \| --- \| --- \| | \| 813 \| WBGene00015628:best-6(9320) \| ``` log2FC=-0.059 padj=0.989 ``` \| [seedLoc = 101,106] Seq=TACTAGAAATGTTACCTCA \| ```                    G   AG     T mRNA  3' ACTCCAT  T TAA   ATCA  miRNA 5' TGAGGTA  A GTT   TAGT                  GT G   GTA    T           1111111661321133611113 ``` \| SeedType = 1 NonSeedType = -1 \| \| --- \| --- \| --- \| --- \| --- \| --- \| | \| 814 \| WBGene00015676:mct-6(9347) \| ``` log2FC=0.278 padj=0.493 ``` \| [seedLoc = 429,434] Seq=TTTTGTTTTCCTATCTTTAATATACCTCA \| ```                 ATAATTTC     TTT     TT mRNA  3' ACTCCAT        TATCC    TGTT   miRNA 5' TGAGGTA        GTAGG    ATAG                                TTGT    TT           111111155555555211113336121233 ``` \| SeedType = 1 NonSeedType = -1 \| \| --- \| --- \| --- \| --- \| --- \| --- \| | \| 815 \| WBGene00015706:C12D5.3(9371) \| ``` log2FC=-0.317 padj=1.0 ``` \| [seedLoc = 42,47] Seq=TCTCCAAAAACCTCCGAAAAATCTACTTCA \| ```                  TAAAAAGCC     AAACCTCT mRNA  3' ACTTCATC         TCCAA         miRNA 5' TGAGGTAG         AGGTT                          T             GTATAGTT           111211113555554551111133333333 ``` \| SeedType = 3 NonSeedType = 1 \| \| --- \| --- \| --- \| --- \| --- \| --- \| | \| 816 \| WBGene00015707:C12D5.4(9372) \| ``` log2FC=1.359 padj=1.0 ``` \| [seedLoc = 39,45] Seq=AAACTGTTCACTCTTCGCATCTACCTAC \| ```          CA      TACGCT   C  T      A mRNA  3'   TCCATC      TCT AC TGTCAA  miRNA 5'   AGGTAG      AGG TG ATAGTT           TG      T        T  T                  3311111135545511231131211115 ``` \| SeedType = 3 NonSeedType = -1 \| \| --- \| --- \| --- \| --- \| --- \| --- \| | \|  \|  \|  \| [seedLoc = 55,61] Seq=TCGCATCTACCTACATTCAAATCTACACTCA \| ```              A   CTAAACTTA      TCT   CT     mRNA  3' ACTC CAT         CATCCA   ACG       miRNA 5' TGAG GTA         GTAGGT   TGT                                             ATAGTT           11115111555555555111111555112336666 ``` \| SeedType = 2 NonSeedType = 1 \| \| --- \| --- \| --- \| --- \| --- \| --- \| | \|  \|  \|  \| [seedLoc = 77,83] Seq=ATCTGTTCAAGACTACACTCC \| ```          C   A     G     T    TA mRNA  3'  CTC CATCA   AAC TGTC   miRNA 5'  GAG GTAGT   TTG ATAG            T         AGG   T    TT           31115111113661113121133 ``` \| SeedType = 2 NonSeedType = 1 \| \| --- \| --- \| --- \| --- \| --- \| --- \| | \| 817 \| WBGene00015708:C12D5.5(9373) \| ``` log2FC=0.119 padj=1.0 ``` \| [seedLoc = 83,89] Seq=ATCACCTCATCCGACTACACTCC \| ```          C   A     G  T  TCCACTA mRNA  3'  CTC CATCA CC AC        miRNA 5'  GAG GTAGT GG TG                 T         A  T  TATAGTT           31115111113113113333333 ``` \| SeedType = 2 NonSeedType = -1 \| \| --- \| --- \| --- \| --- \| --- \| --- \| | \| 818 \| WBGene00015776:dct-1(9411) \| ``` log2FC=-0.415 padj=0.279 ``` \| [seedLoc = 28,33] Seq=ATTTTAACGTTCCACTTACCTCA \| ```                T      TT   ATTTTA mRNA  3' ACTCCA TCA CC  GCA       miRNA 5' TGAGGT AGT GG  TGT                          A  T    ATAGTT           111111511161135211333333 ``` \| SeedType = 2 NonSeedType = -1 \| \| --- \| --- \| --- \| --- \| --- \| --- \| | \| 819 \| WBGene00015778:got-2.2(9414) \| ``` log2FC=0.18 padj=0.637 ``` \| [seedLoc = 137,142] Seq=CAAGGAACACTCTGTCTACCTCA \| ```                  T    C   AGGAAC mRNA  3' ACTCCATC GTCT ACA       miRNA 5' TGAGGTAG TAGG TGT                             T   ATAGTT           11111111521123111333333 ``` \| SeedType = 1 NonSeedType = -1 \| \| --- \| --- \| --- \| --- \| --- \| --- \| | \| 820 \| WBGene00015780:C14F11.4(9416) \| ``` log2FC=-0.246 padj=0.674 ``` \| [seedLoc = 64,70] Seq=TTTCAATCAATTTTTCTAACCTCC \| ```          C    A   TT      TAACTTT mRNA  3'  CTCC ATC  TTTAAC        miRNA 5'  GAGG TAG  AGGTTG                 T        T       TATAGTT           311115111351221113333333 ``` \| SeedType = 2 NonSeedType = 1 \| \| --- \| --- \| --- \| --- \| --- \| --- \| | \| 821 \| WBGene00015789:syx-6(9424) \| ``` log2FC=0.02 padj=0.994 ``` \| [seedLoc = 162,167] Seq=GTCTGTAGATCAGAACTACTTCT \| ```          T        AGA          TG mRNA  3'  CTTCATCA   CTAG ATGTC   miRNA 5'  GAGGTAGT   GGTT TATAG            T        A      G     TT           311211111345121261121133 ``` \| SeedType = 3 NonSeedType = 2 \| \| --- \| --- \| --- \| --- \| --- \| --- \| | \| 822 \| WBGene00015794:C15F1.1(9429) \| ``` log2FC=-0.142 padj=0.877 ``` \| [seedLoc = 47,52] Seq=TTCTATTCCACTTCTAGTCAACTACTTCC \| ```          C        ACTGATC    C T    TT mRNA  3'  CTTCATCA       TTCA C TATC   miRNA 5'  GAGGTAGT       AGGT G ATAG            T                   T T    TT           31121111155545551211313111133 ``` \| SeedType = 3 NonSeedType = 2 \| \| --- \| --- \| --- \| --- \| --- \| --- \| | \| 823 \| WBGene00015803:C15H9.9(9436) \| ``` log2FC=0.09 padj=0.936 ``` \| [seedLoc = 39,44] Seq=TTTAAAATTGTCTTTACCTCG \| ```          G           GTTAAAATTT mRNA  3'  CTCCATT TCT           miRNA 5'  GAGGTAG AGG                    T       T   TTGTATAGTT           3111111261123333333333 ``` \| SeedType = 1 NonSeedType = -1 \| \| --- \| --- \| --- \| --- \| --- \| --- \| | \| 824 \| WBGene00015818:C16A11.5(9447) \| ``` log2FC=-0.33 padj=0.888 ``` \| [seedLoc = 90,95] Seq=CAACTGACCCTGAATTTCGAATTTTACCTCT \| ```          T       TTAAGCTTTAA              C mRNA  3'  CTCCATT           GTCC  CA GTCAA  miRNA 5'  GAGGTAG           TAGG  GT TAGTT           T                      TT  A                 3111111255554555555211166116211115 ``` \| SeedType = 1 NonSeedType = -1 \| \| --- \| --- \| --- \| --- \| --- \| --- \| | \| 825 \| WBGene00015841:skpo-2(9457) \| ``` log2FC=-0.305 padj=0.64 ``` \| [seedLoc = 40,45] Seq=TTCAATTCAACTCTTCTCATCTGCCTCA \| ```                  TACTCTTC     TTAACTT mRNA  3' ACTCCGTC        TCAAC        miRNA 5' TGAGGTAG        GGTTG                         TA           TATAGTT           1111121133555555211113333333 ``` \| SeedType = 2 NonSeedType = 1 \| \| --- \| --- \| --- \| --- \| --- \| --- \| | \| 826 \| WBGene00015868:C16H3.3(9474) \| ``` log2FC=-0.146 padj=0.936 ``` \| [seedLoc = 121,127] Seq=CTATCAGCCCAAACAAAAAGTTGTACTCTCT \| ```          T   T   GTTGAAAAACAAAC             mRNA  3'  CTC CAT              CCGAC TATC   miRNA 5'  GAG GTA              GGTTG ATAG            T       GTA                T    TT           3111511133345555555555112116111166 ``` \| SeedType = 2 NonSeedType = 2 \| \| --- \| --- \| --- \| --- \| --- \| --- \| | \| 827 \| WBGene00015894:acdh-2(9487) \| ``` log2FC=0.339 padj=0.32 ``` \| [seedLoc = 320,326] Seq=TCAATGGCAAAGTATTACGCTCC \| ```          C   G      GA    GTAACT mRNA  3'  CTC CATTAT  AACG       miRNA 5'  GAG GTAGTA  TTGT                T          GG    ATAGTT           31114111211331112333333 ``` \| SeedType = 2 NonSeedType = 1 \| \| --- \| --- \| --- \| --- \| --- \| --- \| | \| 828 \| WBGene00015920:eif-3.L(9509) \|  \| [seedLoc = 289,295] Seq=TGATTGTTTATATTCTACGCTCA \| ```              G       AT  T      T mRNA  3' ACTC CATC TT  AT TGTTAG  miRNA 5' TGAG GTAG AG  TG ATAGTT                    T  GT  T                  111141111612331231212125 ``` \| SeedType = 2 NonSeedType = -1 \| \| --- \| --- \| --- \| --- \| --- \| --- \| | \|  \|  \|  \| [seedLoc = 304,309] Seq=TTCTACGCTCATCGCAACTACTTCT \| ```          T        ACG   CT  C   TT mRNA  3'  CTTCATCA   CTA  CG ATC   miRNA 5'  GAGGTAGT   GGT  GT TAG            T        A     T   A   TT           3112111113541213512311133 ``` \| SeedType = 3 NonSeedType = 2 \| \| --- \| --- \| --- \| --- \| --- \| --- \| | \| 829 \| WBGene00015971:swsn-2.2(9550) \| ``` log2FC=-0.014 padj=0.994 ``` \| [seedLoc = 118,124] Seq=TCATGTACAAAAAATACGCTCT \| ```          T   G   AAAA        ACT mRNA  3'  CTC CAT     AACATGT    miRNA 5'  GAG GTA     TTGTATA             T       GTAGG       GTT           31114111333361111121333 ``` \| SeedType = 2 NonSeedType = 1 \| \| --- \| --- \| --- \| --- \| --- \| --- \| | \| 830 \| WBGene00015993:C18H7.1(9564) \| ``` log2FC=-2.057 padj=1.0 ``` \| [seedLoc = 167,172] Seq=ATCTGAACATTTTTCTACCTCA \| ```                  T   T   A   TA mRNA  3' ACTCCATC TTT ACA GTC   miRNA 5' TGAGGTAG AGG TGT TAG                    T   T   A   TT           1111111131223111321133 ``` \| SeedType = 1 NonSeedType = -1 \| \| --- \| --- \| --- \| --- \| --- \| --- \| | \| 831 \| WBGene00016002:C18H9.3(9569) \| ``` log2FC=0.044 padj=0.967 ``` \| [seedLoc = 179,185] Seq=TTTCATATCCCTTTACTCTCT \| ```          T   T        C     CTTT mRNA  3'  CTC CATT TCC  TATA     miRNA 5'  GAG GTAG AGG  GTAT              T        T   TT    AGTT           31115111261113621113333 ``` \| SeedType = 2 NonSeedType = -1 \| \| --- \| --- \| --- \| --- \| --- \| --- \| | \| 832 \| WBGene00016067:C24H10.1(9618) \| ``` log2FC=0.025 padj=0.994 ``` \| [seedLoc = 66,71] Seq=TAACTCAAACACTACCTTA \| ```                     AA         T mRNA  3' ATTCCATCA C  AC   TCAA  miRNA 5' TGAGGTAGT G  TG   AGTT                    A GT  TAT                12111111161331166611115 ``` \| SeedType = 3 NonSeedType = -1 \| \| --- \| --- \| --- \| --- \| --- \| --- \| | \| 833 \| WBGene00016111:C25H3.1(9644) \| ``` log2FC=-0.683 padj=1.0 ``` \| [seedLoc = 69,74] Seq=TGTGATATTGTGTTTTACCTCC \| ```          C       TTGTGT    GTGT mRNA  3'  CTCCATT      TATA     miRNA 5'  GAGGTAG      GTAT              T       TAGGTT    AGTT           3111111233333321113333 ``` \| SeedType = 1 NonSeedType = -1 \| \| --- \| --- \| --- \| --- \| --- \| --- \| | \|  \|  \|  \| [seedLoc = 81,86] Seq=TGTTTTACCTCCTCGCCTACTTCA \| ```                  CGC   TC   TTTGT mRNA  3' ACTTCATC   TCC  CAT      miRNA 5' TGAGGTAG   AGG  GTA                       T     TT   TAGTT           111211113451113311133333 ``` \| SeedType = 3 NonSeedType = -1 \| \| --- \| --- \| --- \| --- \| --- \| --- \| | \| 834 \| WBGene00016112:C25H3.3(9645) \| ``` log2FC=-0.186 padj=0.874 ``` \| [seedLoc = 236,241] Seq=AAATTTCGTGTCTGGAATACCTCC \| ```          C      AAG    GTGCTTTAAA mRNA  3'  CTCCAT   GTCT           miRNA 5'  GAGGTA   TAGG                    T      G      TTGTATAGTT           311111135421123333333333 ``` \| SeedType = 1 NonSeedType = -1 \| \| --- \| --- \| --- \| --- \| --- \| --- \| | \| 835 \| WBGene00016119:C25H3.10(9652) \| ``` log2FC=-0.159 padj=0.9 ``` \| [seedLoc = 162,167] Seq=AAGTTCAGATATATCTACCTTC \| ```          C       TATATAGACTTGAA mRNA  3'  TTCCATC               miRNA 5'  GAGGTAG                        T       TAGGTTGTATAGTT           3211111133333333333333 ``` \| SeedType = 3 NonSeedType = -1 \| \| --- \| --- \| --- \| --- \| --- \| --- \| | \| 836 \| WBGene00016120:C25H3.11(9653) \| ``` log2FC=0.138 padj=0.888 ``` \| [seedLoc = 180,186] Seq=TGCTTTACAATTTCTACCCTCA \| ```             C              T   T mRNA  3' ACT CCATC TTTAACAT TCG  miRNA 5' TGA GGTAG AGGTTGTA AGT                    T        T   T           11151111161221111131123 ``` \| SeedType = 3 NonSeedType = 1 \| \| --- \| --- \| --- \| --- \| --- \| --- \| | \| 837 \| WBGene00016131:C26B9.1(9660) \| ``` log2FC=0.465 padj=0.672 ``` \| [seedLoc = 410,416] Seq=TTCTGGTTACTGGCTACCTAC \| ```          CA       G   TTG    TT mRNA  3'   TCCATCG TCA    GTC   miRNA 5'   AGGTAGT GGT    TAG            TG       A   TGTA   TT           3311111123211333621133 ``` \| SeedType = 3 NonSeedType = 2 \| \| --- \| --- \| --- \| --- \| --- \| --- \| | \| 838 \| WBGene00016139:ntl-9(9667) \| ``` log2FC=0.026 padj=0.986 ``` \| [seedLoc = 24,29] Seq=TCTTCTTCGTCCTACTACTTCA \| ```                      TGCTTCTTCT mRNA  3' ACTTCATCATCC           miRNA 5' TGAGGTAGTAGG                                TTGTATAGTT           1112111111113333333333 ``` \| SeedType = 3 NonSeedType = -1 \| \| --- \| --- \| --- \| --- \| --- \| --- \| | \|  \|  \|  \| [seedLoc = 376,382] Seq=TTCAGCAGAATCTCTATCCTCT \| ```          T    T         G CGACTT mRNA  3'  CTCC ATC TCTAA A       miRNA 5'  GAGG TAG AGGTT T                T        T     G ATAGTT           31111511161121131333333 ``` \| SeedType = 2 NonSeedType = 2 \| \| --- \| --- \| --- \| --- \| --- \| --- \| | \| 839 \| WBGene00016140:rpb-2(9668) \| ``` log2FC=-0.012 padj=0.994 ``` \| [seedLoc = 285,291] Seq=GTTTATATGTGAACCTTCTATCCTCC \| ```          C    T   T     GTG      TG mRNA  3'  CTCC ATC TCCAA   TATATT   miRNA 5'  GAGG TAG AGGTT   GTATAG            T        T              TT           31111511131111145421111233 ``` \| SeedType = 2 NonSeedType = 1 \| \| --- \| --- \| --- \| --- \| --- \| --- \| | \| 840 \| WBGene00016146:C26F1.1(9673) \| ``` log2FC=-0.575 padj=0.14 ``` \| [seedLoc = 55,60] Seq=ATTTATGCTCTCTGAAACTTACCTCC \| ```          C       CAAA    CT      TA mRNA  3'  CTCCATT    GTCT  CGTATT   miRNA 5'  GAGGTAG    TAGG  GTATAG            T               TT      TT           31111112555521123312111233 ``` \| SeedType = 1 NonSeedType = -1 \| \| --- \| --- \| --- \| --- \| --- \| --- \| | \| 841 \| WBGene00016158:ari-1.1(9685) \|  \| [seedLoc = 140,146] Seq=CTGTATGTTCATTTCCTGTCATTTACCATCA \| ```             A     TACT    TTTACTTG         mRNA  3' ACT CCATT    GTCC        TATGTC   miRNA 5' TGA GGTAG    TAGG        GTATAG                             TT            TT           111511112555521113355555421121166 ``` \| SeedType = 3 NonSeedType = -1 \| \| --- \| --- \| --- \| --- \| --- \| --- \| | \| 842 \| WBGene00016162:crh-2(9689) \| ``` log2FC=-0.016 padj=0.993 ``` \| [seedLoc = 523,528] Seq=TTTCTTGGGACACTGATTCTACCTCC \| ```          C       TTA   A   G  TCTTT mRNA  3'  CTCCATC   GTC CAG GT      miRNA 5'  GAGGTAG   TAG GTT TA               T                 G  TAGTT           31111111555211511232133333 ``` \| SeedType = 1 NonSeedType = -1 \| \| --- \| --- \| --- \| --- \| --- \| --- \| | \| 843 \| WBGene00016173:fust-1(9700) \| ``` log2FC=-0.233 padj=0.653 ``` \| [seedLoc = 101,106] Seq=TTTTATATCCACCTTTATATTCTACTTCG \| ```          G       TTATATT    CC      TT mRNA  3'  CTTCATC       TCCA  TATATT   miRNA 5'  GAGGTAG       AGGT  GTATAG            T       T          T       TT           31121111355555511113521111233 ``` \| SeedType = 3 NonSeedType = 1 \| \| --- \| --- \| --- \| --- \| --- \| --- \| | \| 844 \| WBGene00016190:C28G1.5(9712) \| ``` log2FC=0.225 padj=0.821 ``` \| [seedLoc = 1105,1110] Seq=AAACTTCTACCGGGTACCTCA \| ```                 G G   TCT      A mRNA  3' ACTCCAT G CCA     TCAA  miRNA 5' TGAGGTA T GGT     AGTT                  G A   TGTAT                11111113231113336611115 ``` \| SeedType = 1 NonSeedType = 1 \| \| --- \| --- \| --- \| --- \| --- \| --- \| | \| 845 \| WBGene00016197:pxl-1(9721) \| ``` log2FC=0.091 padj=0.931 ``` \| [seedLoc = 39,45] Seq=TGTTTTGATATCTAACTACCTGC \| ```          CG      A     TAGTTTTGT mRNA  3'   TCCATC ATCTA          miRNA 5'   AGGTAG TAGGT                   TG            TGTATAGTT           33111111511121333333333 ``` \| SeedType = 3 NonSeedType = 2 \| \| --- \| --- \| --- \| --- \| --- \| --- \| | \| 846 \| WBGene00016235:C29H12.2(9741) \| ``` log2FC=0.318 padj=0.36 ``` \| [seedLoc = 109,114] Seq=AAAACAACGGCATGTGCTACCTTT \| ```          T         GTA     ACAAAA mRNA  3'  TTCCATCGT   CGGCA       miRNA 5'  GAGGTAGTA   GTTGT                T         G       ATAGTT           321111112135512211333333 ``` \| SeedType = 3 NonSeedType = -1 \| \| --- \| --- \| --- \| --- \| --- \| --- \| | \| 847 \| WBGene00016237:C29H12.6(9743) \| ``` log2FC=0.117 padj=0.974 ``` \| [seedLoc = 121,126] Seq=GTACAAAATTGTTCCGGTTCTCCCTATCTCC \| ```          C           TCTTGGCCTTGTTAA           mRNA  3'  CTCTATC  CC               AACATG     miRNA 5'  GAGGTAG  GG               TTGTAT              T       TA                       AGTT           3111211166115555445555455551111126666 ``` \| SeedType = 2 NonSeedType = 1 \| \| --- \| --- \| --- \| --- \| --- \| --- \| | \| 848 \| WBGene00016238:mob-4(9744) \|  \| [seedLoc = 284,289] Seq=CCCTATTCGAATCTATCTCA \| ```                  TA     T    CC mRNA  3' ACTCTATC    AGC TATC   miRNA 5' TGAGGTAG    TTG ATAG                    TAGG   T    TT           1111211133661213111133 ``` \| SeedType = 2 NonSeedType = -1 \| \| --- \| --- \| --- \| --- \| --- \| --- \| | \| 849 \| WBGene00016244:C30B5.2(9747) \| ``` log2FC=-0.135 padj=0.873 ``` \| [seedLoc = 211,216] Seq=TTCGATTTTCCCTATATTCTACTTCG \| ```          G       TTAT    CTTTTAGCTT mRNA  3'  CTTCATC    ATCC           miRNA 5'  GAGGTAG    TAGG                    T               TTGTATAGTT           31121111555511113333333333 ``` \| SeedType = 3 NonSeedType = -1 \| \| --- \| --- \| --- \| --- \| --- \| --- \| | \| 850 \| WBGene00016318:C32D5.11(9789) \| ``` log2FC=-0.176 padj=0.794 ``` \| [seedLoc = 75,81] Seq=ATCTGTATGGATAAATGTACACTCG \| ```          G   A   GTAAATA        TA mRNA  3'  CTC CAT       GGTATGTC   miRNA 5'  GAG GTA       TTGTATAG            T       GTAGG          TT           3111511133333552221121133 ``` \| SeedType = 2 NonSeedType = -1 \| \| --- \| --- \| --- \| --- \| --- \| --- \| | \| 851 \| WBGene00016326:ubr-1(9797) \| ``` log2FC=0.042 padj=0.975 ``` \| [seedLoc = 394,400] Seq=AGTTGTAATTTTCCTTTGTACCCTCT \| ```          T  C    GTT   TTTTA     GA mRNA  3'  CT CCAT   TCC     ATGTT   miRNA 5'  GA GGTA   AGG     TATAG            T       GT    TTG       TT           31151111335111333551121233 ``` \| SeedType = 3 NonSeedType = -1 \| \| --- \| --- \| --- \| --- \| --- \| --- \| | \| 852 \| WBGene00016333:C32F10.8(9803) \| ``` log2FC=0.107 padj=0.85 ``` \| [seedLoc = 93,99] Seq=TCTTAAGGAACGTGCTACACTCG \| ```          G   A      G   G AATTCT mRNA  3'  CTC CATCGT CAA G       miRNA 5'  GAG GTAGTA GTT T                T          G   G ATAGTT           31115111121311132333333 ``` \| SeedType = 2 NonSeedType = 1 \| \| --- \| --- \| --- \| --- \| --- \| --- \| | \| 853 \| WBGene00016343:cnnm-3(9809) \|  \| [seedLoc = 392,398] Seq=CCATTCTTCACTTATATACGCTCT \| ```          T   G   A      CTTCTTACC mRNA  3'  CTC CAT TATTCA          miRNA 5'  GAG GTA GTAGGT                   T              TGTATAGTT           311141115211211333333333 ``` \| SeedType = 2 NonSeedType = 2 \| \| --- \| --- \| --- \| --- \| --- \| --- \| | \| 854 \| WBGene00016374:swd-2.2(9829) \| ``` log2FC=0.135 padj=0.902 ``` \| [seedLoc = 32,37] Seq=CTCAAGAAAATCTCTTTTTAACTATCTCA \| ```                   ATTTTTC     AAGAACTC mRNA  3' ACTCTATCA       TCTAA         miRNA 5' TGAGGTAGT       AGGTT                                       GTATAGTT           11112111155555551121133333333 ``` \| SeedType = 2 NonSeedType = 2 \| \| --- \| --- \| --- \| --- \| --- \| --- \| | \| 855 \| WBGene00016406:C34D10.1(9854) \| ``` log2FC=-0.462 padj=0.369 ``` \| [seedLoc = 258,263] Seq=ATGTATATGACTTCTCTACCTCT \| ```          T       TC          GTA mRNA  3'  CTCCATC  TTCAGTATAT    miRNA 5'  GAGGTAG  AGGTTGTATA             T       T           GTT           31111111351211221111333 ``` \| SeedType = 1 NonSeedType = 2 \| \| --- \| --- \| --- \| --- \| --- \| --- \| | \| 856 \| WBGene00016409:sumv-1(9859) \| ``` log2FC=0.024 padj=0.99 ``` \| [seedLoc = 57,63] Seq=TGTTTCACGATTTCGCTATCCTCA \| ```               T    C       CTTTGT mRNA  3' ACTCC ATCG TTTAGCA       miRNA 5' TGAGG TAGT AGGTTGT                                  ATAGTT           111115111251221211333333 ``` \| SeedType = 2 NonSeedType = 2 \| \| --- \| --- \| --- \| --- \| --- \| --- \| | \|  \|  \|  \| [seedLoc = 123,128] Seq=ATAATTTTCACCTTGTTACCTCC \| ```          C       GT    CTTTTAATA mRNA  3'  CTCCATT  TCCA          miRNA 5'  GAGGTAG  AGGT                   T       T     TGTATAGTT           31111112351111333333333 ``` \| SeedType = 1 NonSeedType = 1 \| \| --- \| --- \| --- \| --- \| --- \| --- \| | \| 857 \| WBGene00016422:noah-1(9871) \| ``` log2FC=1.204 padj=1.11107033653082E-7 over-expressed ``` \| [seedLoc = 231,236] Seq=AAAAAAGATCCCTTTTTCAACTATCTCT \| ```          T        ACTTTT   CTAGAAAAAA mRNA  3'  CTCTATCA      TCC           miRNA 5'  GAGGTAGT      AGG                    T                 TTGTATAGTT           3111211115555551113333333333 ``` \| SeedType = 2 NonSeedType = -1 \| \| --- \| --- \| --- \| --- \| --- \| --- \| | \| 858 \| WBGene00016429:C35A11.2(9877) \| ``` log2FC=-0.597 padj=0.444 ``` \| [seedLoc = 91,96] Seq=TTTGTTGCATTCCTTCTACCTCT \| ```          T       T   TT    TGTTT mRNA  3'  CTCCATC TCC  ACGT      miRNA 5'  GAGGTAG AGG  TGTA               T       T   T     TAGTT           31111111311135112133333 ``` \| SeedType = 1 NonSeedType = -1 \| \| --- \| --- \| --- \| --- \| --- \| --- \| | \| 859 \| WBGene00016436:C35B1.7(9883) \| ``` log2FC=0.86 padj=1.0 ``` \| [seedLoc = 63,68] Seq=TGATAGACCCCGATTTTTCTCATCTACTTCT \| ```          T       TACTCTTTTTAGCC    ATAGT    mRNA  3'  CTTCATC              CCAG         miRNA 5'  GAGGTAG              GGTT                  T       TA                GTATAGTT           3112111133555555555455111233333666 ``` \| SeedType = 3 NonSeedType = 1 \| \| --- \| --- \| --- \| --- \| --- \| --- \| | \| 860 \| WBGene00016454:C35E7.2(9898) \| ``` log2FC=-0.193 padj=0.967 ``` \| [seedLoc = 26,32] Seq=AAACTCAATCAAACTACTCTCA \| ```              T     AA            A mRNA  3' ACTC CATCA  CTAAC   TCAA  miRNA 5' TGAG GTAGT  GGTTG   AGTT                     A      TAT                1111511111351211166611115 ``` \| SeedType = 2 NonSeedType = 1 \| \| --- \| --- \| --- \| --- \| --- \| --- \| | \|  \|  \|  \| [seedLoc = 153,159] Seq=GACAAAACGATTCCATATCCTCA \| ```               T  ACC      AAACAG mRNA  3' ACTCC AT   TTAGCA       miRNA 5' TGAGG TA   GGTTGT                        GTA      ATAGTT           11111511333221211333333 ``` \| SeedType = 2 NonSeedType = 2 \| \| --- \| --- \| --- \| --- \| --- \| --- \| | \|  \|  \|  \| [seedLoc = 201,206] Seq=TACTGTACGTTTCCAGCTACCTTT \| ```          T        A  TTT        T mRNA  3'  TTCCATCG CC   GCATGTCA  miRNA 5'  GAGGTAGT GG   TGTATAGT           T        A  T          T           321111112311355211121113 ``` \| SeedType = 3 NonSeedType = -1 \| \| --- \| --- \| --- \| --- \| --- \| --- \| | \| 861 \| WBGene00016456:C35E7.4(9899) \| ``` log2FC=-0.584 padj=0.598 ``` \| [seedLoc = 62,67] Seq=CATTATTTTTCCCAATACCTCA \| ```                 AAC  TTTT     C mRNA  3' ACTCCAT   CC    TATTA  miRNA 5' TGAGGTA   GG    ATAGT                  GTA  TTGT     T           1111111333113333111213 ``` \| SeedType = 1 NonSeedType = -1 \| \| --- \| --- \| --- \| --- \| --- \| --- \| | \| 862 \| WBGene00016486:fbxa-170(9911) \| ``` log2FC=-0.759 padj=1.0 ``` \| [seedLoc = 211,215] Seq=TCCTTCCACTAACTCT \| ```          T   A       TT CT      mRNA  3'  CTC ATCA CC  C        miRNA 5'  GAG TAGT GG  G                 T   G    A  TT TATAGTT           3111311116113313366666 ``` \| SeedType = 2 NonSeedType = -1 \| \| --- \| --- \| --- \| --- \| --- \| --- \| | \| 863 \| WBGene00016509:adss-1(9931) \| ``` log2FC=0.081 padj=0.925 ``` \| [seedLoc = 287,293] Seq=TTCCCCTTCCCCTAGCGTACCGTCT \| ```          T  G    G G    CCTTCCCCTT mRNA  3'  CT CCAT C ATCC           miRNA 5'  GA GGTA G TAGG                    T              TTGTATAGTT           3114111141411113333333333 ``` \| SeedType = 3 NonSeedType = -1 \| \| --- \| --- \| --- \| --- \| --- \| --- \| | \| 864 \| WBGene00016561:eri-6(9965) \| ``` log2FC=-0.333 padj=0.765 ``` \| [seedLoc = 549,554] Seq=AAATTGTGGATGAAGAACTTACCTCC \| ```          C     T   AGAAGT G       A mRNA  3'  CTCCA TCA      A GTGTTAA  miRNA 5'  GAGGT AGT      T TATAGTT           T         AGGT   G                   31111151113333451321212115 ``` \| SeedType = 2 NonSeedType = -1 \| \| --- \| --- \| --- \| --- \| --- \| --- \| | \| 865 \| WBGene00016575:lido-10(9979) \|  \| [seedLoc = 177,183] Seq=GGAATTGGGAGTTCTACCGTCA \| ```             G        G  G  TAAGG mRNA  3' ACT CCATC TT AG GT      miRNA 5' TGA GGTAG AG TT TA                        T  G  G  TAGTT           11141111161231232133333 ``` \| SeedType = 3 NonSeedType = -1 \| \| --- \| --- \| --- \| --- \| --- \| --- \| | \| 866 \| WBGene00016576:lido-9(9980) \|  \| [seedLoc = 120,125] Seq=TATCACGCGGTCTCACTACTTCA \| ```                   C       CACTAT mRNA  3' ACTTCATCA TCTGGCG       miRNA 5' TGAGGTAGT AGGTTGT                                 ATAGTT           11121111151122212333333 ``` \| SeedType = 3 NonSeedType = 2 \| \| --- \| --- \| --- \| --- \| --- \| --- \| | \| 867 \| WBGene00016609:C43E11.12(10009) \| ``` log2FC=0.009 padj=0.997 ``` \| [seedLoc = 44,49] Seq=CTGAAATAAACTCATATACCTCC \| ```          C      ATAC    ATAAAGTC mRNA  3'  CTCCAT    TCAA         miRNA 5'  GAGGTA    GGTT                  T      GTA     GTATAGTT           31111113335211133333333 ``` \| SeedType = 1 NonSeedType = 1 \| \| --- \| --- \| --- \| --- \| --- \| --- \| | \| 868 \| WBGene00016611:bicd-1(10012) \| ``` log2FC=0.111 padj=0.888 ``` \| [seedLoc = 415,420] Seq=TTTGGCAGCCGACATTTCACGGGTTACCTCA \| ```                 TGGGCACTTTA  G      GTTT   mRNA  3' ACTCCAT           CA CCGACG       miRNA 5' TGAGGTA           GT GGTTGT                                    A      ATAGTT           111111154445555555113112112333366 ``` \| SeedType = 1 NonSeedType = 2 \| \| --- \| --- \| --- \| --- \| --- \| --- \| | \| 869 \| WBGene00016622:mafr-1(10021) \| ``` log2FC=-0.104 padj=0.929 ``` \| [seedLoc = 139,142] Seq=TTCTCATCAATCCTCAACCTCC \| ```          C     A TC     TACTCTT mRNA  3'  CTCCA C  CTAAC        miRNA 5'  GAGGT G  GGTTG                 T     A TA     TATAGTT           3111113133121113333333 ``` \| SeedType = 2 NonSeedType = 1 \| \| --- \| --- \| --- \| --- \| --- \| --- \| | \| 870 \| WBGene00016653:ssb-1(10048) \|  \| [seedLoc = 113,119] Seq=CATTCATTAATCTTGAATACCCTCA \| ```             C    AAGT      TACTTAC mRNA  3' ACT CCAT    TCTAAT        miRNA 5' TGA GGTA    AGGTTG                         GT        TATAGTT           1115111133451121123333333 ``` \| SeedType = 3 NonSeedType = 2 \| \| --- \| --- \| --- \| --- \| --- \| --- \| | \|  \|  \|  \| [seedLoc = 129,133] Seq=AATCTTGAATACCCTCAT \| ```          T   C    AAGT              mRNA  3'  ACT CCAT    TCTAA         miRNA 5'  TGA GGTA    AGGTT                           GT       GTATAGTT           51115111133451121166666666 ``` \| SeedType = 3 NonSeedType = 2 \| \| --- \| --- \| --- \| --- \| --- \| --- \| | \| 871 \| WBGene00016712:C46E10.8(10081) \| ``` log2FC=1.636 padj=1.0 ``` \| [seedLoc = 302,307] Seq=TACTTGGCGAATCTACCTCG \| ```          G       TA      GT   T mRNA  3'  CTCCATC    AGCG  TCA  miRNA 5'  GAGGTAG    TTGT  AGT           T       TAGG    AT   T           3111111133661212331113 ``` \| SeedType = 1 NonSeedType = -1 \| \| --- \| --- \| --- \| --- \| --- \| --- \| | \| 872 \| WBGene00016739:pitr-1(10095) \| ``` log2FC=0.123 padj=0.843 ``` \| [seedLoc = 428,433] Seq=TTTTTTTTGTTTTGTTACCTCA \| ```                      TGTTTTTTTT mRNA  3' ACTCCATTGTTT           miRNA 5' TGAGGTAGTAGG                                TTGTATAGTT           1111111221223333333333 ``` \| SeedType = 1 NonSeedType = -1 \| \| --- \| --- \| --- \| --- \| --- \| --- \| | \|  \|  \|  \| [seedLoc = 448,454] Seq=CATTTCATATTTTATTAACCTCT \| ```          T    A       T   CTTTAC mRNA  3'  CTCC ATTATTT ATA       miRNA 5'  GAGG TAGTAGG TGT                T            T   ATAGTT           31111511211223121333333 ``` \| SeedType = 2 NonSeedType = -1 \| \| --- \| --- \| --- \| --- \| --- \| --- \| | \|  \|  \|  \| [seedLoc = 787,792] Seq=CATTGTTTTATCTTCTACCTTT \| ```          T       T    TTT     C mRNA  3'  TTCCATC TCTA   TGTTA  miRNA 5'  GAGGTAG AGGT   ATAGT           T       T    TGT     T           3211111131121333121213 ``` \| SeedType = 3 NonSeedType = 2 \| \| --- \| --- \| --- \| --- \| --- \| --- \| | \| 873 \| WBGene00016750:let-611(10105) \|  \| [seedLoc = 125,130] Seq=GTTTTTGTTGCTTCTACCTTC \| ```          C            TTGTTTTTG mRNA  3'  TTCCATC TTCG          miRNA 5'  GAGGTAG AGGT                   T       T    TGTATAGTT           3211111161212333333333 ``` \| SeedType = 3 NonSeedType = 2 \| \| --- \| --- \| --- \| --- \| --- \| --- \| | \| 874 \| WBGene00016751:C48E7.6(10106) \| ``` log2FC=0.064 padj=0.982 ``` \| [seedLoc = 91,95] Seq=CAAAATCATTCTCTACGTCT \| ```          T  G        T  TAAAAC  mRNA  3'  CT CATC TCT AC        miRNA 5'  GA GTAG AGG TG                 T  G    T   T  TATAGTT           3113111161123113333336 ``` \| SeedType = 3 NonSeedType = -1 \| \| --- \| --- \| --- \| --- \| --- \| --- \| | \| 875 \| WBGene00016855:C50H11.8(10175) \| ``` log2FC=0.253 padj=1.0 ``` \| [seedLoc = 276,281] Seq=TGATTATCTCCAATTACCTCT \| ```          T        A  T         T mRNA  3'  CTCCATTA CC  C TATTAG  miRNA 5'  GAGGTAGT GG  G ATAGTT           T        A  TT T                  31111112131136161112125 ``` \| SeedType = 1 NonSeedType = -1 \| \| --- \| --- \| --- \| --- \| --- \| --- \| | \| 876 \| WBGene00016865:ets-9(10182) \| ``` log2FC=-0.301 padj=0.71 ``` \| [seedLoc = 124,129] Seq=GTCTGTACAATTGTACCTCC \| ```          C      G            TG mRNA  3'  CTCCAT   TTAACATGTC   miRNA 5'  GAGGTA   GGTTGTATAG            T      GTA          TT           3111111366221111121133 ``` \| SeedType = 1 NonSeedType = 1 \| \| --- \| --- \| --- \| --- \| --- \| --- \| | \| 877 \| WBGene00016902:C53D5.1(10205) \| ``` log2FC=0.209 padj=0.859 ``` \| [seedLoc = 323,328] Seq=ATATGTTTCACCTTGTCTCATTTGTACCTCG \| ```          G      GTTTACTCTGT    CTTTGTATA mRNA  3'  CTCCAT           TCCA          miRNA 5'  GAGGTA           AGGT                   T      GT             TGTATAGTT           3111111335555555451111333333333 ``` \| SeedType = 1 NonSeedType = 1 \| \| --- \| --- \| --- \| --- \| --- \| --- \| | \| 878 \| WBGene00016916:C54D2.2(10213) \| ``` log2FC=0.146 padj=1.0 ``` \| [seedLoc = 43,48] Seq=TACTTCAACACTCCCCATTATCACTACCTTC \| ```          C        CTATTACCCC  A    T     T mRNA  3'  TTCCATCA          TC CAAC   TCA  miRNA 5'  GAGGTAGT          AG GTTG   AGT           T                         TAT   T           321111111555555555511511113661113 ``` \| SeedType = 3 NonSeedType = 1 \| \| --- \| --- \| --- \| --- \| --- \| --- \| | \| 879 \| WBGene00016919:C54E4.4(10215) \| ``` log2FC=0.781 padj=0.661 ``` \| [seedLoc = 63,68] Seq=TTAAATAGAATTTGTACCTCT \| ```          T             G   AATT mRNA  3'  CTCCAT GTTTAA ATA     miRNA 5'  GAGGTA TAGGTT TAT              T      G      G   AGTT           3111111621221131113333 ``` \| SeedType = 1 NonSeedType = 2 \| \| --- \| --- \| --- \| --- \| --- \| --- \| | \| 880 \| WBGene00016937:tag-294(10222) \| ``` log2FC=-0.188 padj=0.939 ``` \| [seedLoc = 217,222] Seq=CAACATATTCTGATTCTACTTCG \| ```          G       TTAG  T         C mRNA  3'  CTTCATC    TC  TATA CAA  miRNA 5'  GAGGTAG    GG  GTAT GTT           T       TA    TT    A               3112111133542136211161115 ``` \| SeedType = 3 NonSeedType = -1 \| \| --- \| --- \| --- \| --- \| --- \| --- \| | \| 881 \| WBGene00016947:C55B7.11(10230) \| ``` log2FC=0.046 padj=0.979 ``` \| [seedLoc = 74,80] Seq=TCCCTTACTTTTATCTACACTCA \| ```              A    T    T    TCCCT mRNA  3' ACTC CATC ATTT  CAT      miRNA 5' TGAG GTAG TAGG  GTA                             TT   TAGTT           111151111511223611133333 ``` \| SeedType = 2 NonSeedType = -1 \| \| --- \| --- \| --- \| --- \| --- \| --- \| | \| 882 \| WBGene00016970:C56E6.2(10241) \| ``` log2FC=0.417 padj=0.363 ``` \| [seedLoc = 203,208] Seq=GTTATCCCATCCAATACCTCA \| ```                 AA   T  CCTATTG mRNA  3' ACTCCAT   CC AC        miRNA 5' TGAGGTA   GG TG                        GTA  T  TATAGTT           1111111336113113333333 ``` \| SeedType = 1 NonSeedType = -1 \| \| --- \| --- \| --- \| --- \| --- \| --- \| | \| 883 \| WBGene00016977:akap-1(10247) \| ``` log2FC=-0.078 padj=0.922 ``` \| [seedLoc = 897,902] Seq=TCGTTTTTATCCACTATCTCC \| ```          C           TATTTTTGCT mRNA  3'  CTCTATCA CC           miRNA 5'  GAGGTAGT GG                    T        A  TTGTATAGTT           3111211116113333333333 ``` \| SeedType = 2 NonSeedType = -1 \| \| --- \| --- \| --- \| --- \| --- \| --- \| | \|  \|  \|  \| [seedLoc = 1042,1047] Seq=TTTACCAGGTTCTTCTACCTTT \| ```          T       T   TGGACCATTT mRNA  3'  TTCCATC TCT           miRNA 5'  GAGGTAG AGG                    T       T   TTGTATAGTT           3211111131123333333333 ``` \| SeedType = 3 NonSeedType = -1 \| \| --- \| --- \| --- \| --- \| --- \| --- \| | \| 884 \| WBGene00016995:acly-1(10262) \| ``` log2FC=0.1 padj=1.0 ``` \| [seedLoc = 172,177] Seq=CACTGTACCCTAAAAGTCCCCTATCTCC \| ```          C       CCCTGAAA             C mRNA  3'  CTCTATC        ATCC  CATGTCA  miRNA 5'  GAGGTAG        TAGG  GTATAGT           T                   TT       T           311121115555455511116611121113 ``` \| SeedType = 2 NonSeedType = -1 \| \| --- \| --- \| --- \| --- \| --- \| --- \| | \|  \|  \|  \| [seedLoc = 247,253] Seq=TATTTCCCGTACTCTACTCTCA \| ```              T       AT  CCTTTAT mRNA  3' ACTC CATC TC  GC        miRNA 5' TGAG GTAG AG  TG                          T  GT  TATAGTT           11115111161133213333333 ``` \| SeedType = 2 NonSeedType = -1 \| \| --- \| --- \| --- \| --- \| --- \| --- \| | \| 885 \| WBGene00017003:D1007.5(10268) \| ``` log2FC=-0.219 padj=0.819 ``` \| [seedLoc = 235,240] Seq=TAATTGCTCAATAATTCTGCCTCT \| ```          T       TTAA    TC     T mRNA  3'  CTCCGTC    TAAC  GTTAA  miRNA 5'  GAGGTAG    GTTG  TAGTT           T       TAG     TA                 311112113335211133212115 ``` \| SeedType = 2 NonSeedType = 1 \| \| --- \| --- \| --- \| --- \| --- \| --- \| | \| 886 \| WBGene00017023:D1022.4(10286) \| ``` log2FC=0.311 padj=0.24 ``` \| [seedLoc = 42,47] Seq=AAATTTTGTGACTTTTCTATCTCT \| ```          T       TT        T    A mRNA  3'  CTCTATC  TTCAGTGT TTAA  miRNA 5'  GAGGTAG  AGGTTGTA AGTT           T       T         T                311121113512112221312115 ``` \| SeedType = 2 NonSeedType = 2 \| \| --- \| --- \| --- \| --- \| --- \| --- \| | \| 887 \| WBGene00017045:atg-13(10300) \| ``` log2FC=-0.429 padj=0.136 ``` \| [seedLoc = 64,70] Seq=CGCATGTACACTTTTCACACGTTTACACTCG \| ```          G   A      G  CACTTTTC      A  C mRNA  3'  CTC CATT T CA        ACATGT CG  miRNA 5'  GAG GTAG A GT        TGTATA GT           T        T G                   T           31115111261311555555551111215123 ``` \| SeedType = 2 NonSeedType = -1 \| \| --- \| --- \| --- \| --- \| --- \| --- \| | \| 888 \| WBGene00017048:D2021.4(10303) \| ``` log2FC=-0.362 padj=0.75 ``` \| [seedLoc = 597,602] Seq=TCTTGTCGCAATACACTATCTCT \| ```          T        CA     C    CT mRNA  3'  CTCTATCA  TAACG TGTT   miRNA 5'  GAGGTAGT  GTTGT ATAG            T        AG          TT           31112111133211125121233 ``` \| SeedType = 2 NonSeedType = 1 \| \| --- \| --- \| --- \| --- \| --- \| --- \| | \| 889 \| WBGene00017066:maco-1(10317) \| ``` log2FC=-0.077 padj=0.956 ``` \| [seedLoc = 263,269] Seq=TATTTCTTTTCCTCTCATTACTCTCA \| ```              T     CTC   TTTTCTTTAT mRNA  3' ACTC CATTA   TCC           miRNA 5' TGAG GTAGT   AGG                                    TTGTATAGTT           11115111215551113333333333 ``` \| SeedType = 2 NonSeedType = -1 \| \| --- \| --- \| --- \| --- \| --- \| --- \| | \| 890 \| WBGene00017074:D2096.7(10323) \| ``` log2FC=-0.107 padj=0.959 ``` \| [seedLoc = 35,40] Seq=TACATGTACACTTGCACTTTTACCTCT \| ```          T      TTTCA            A  T mRNA  3'  CTCCAT     CGTTC ACATGT CA  miRNA 5'  GAGGTA     GTAGG TGTATA GT           T                T         T           3111111555551212161111215113 ``` \| SeedType = 1 NonSeedType = -1 \| \| --- \| --- \| --- \| --- \| --- \| --- \| | \| 891 \| WBGene00017088:akir-1(10336) \| ``` log2FC=0.042 padj=0.973 ``` \| [seedLoc = 296,301] Seq=TCGCCTTTGACCTTTTCTTTTACCTCT \| ```          T       TTCTTT     TTTCCGCT mRNA  3'  CTCCATT      TCCAG         miRNA 5'  GAGGTAG      AGGTT                  T       T          GTATAGTT           311111123555551111233333333 ``` \| SeedType = 1 NonSeedType = 1 \| \| --- \| --- \| --- \| --- \| --- \| --- \| | \| 892 \| WBGene00017093:E02C12.8(10341) \| ``` log2FC=-0.799 padj=0.063 ``` \| [seedLoc = 685,688] Seq=GGCTATGATGCTTCTACCTGT \| ```          TG           TA      G mRNA  3'   TCCATC TTCG  GTATCG  miRNA 5'   AGGTAG AGGT  TATAGT           TG      T    TG      T           3311111161212332111123 ``` \| SeedType = 3 NonSeedType = 2 \| \| --- \| --- \| --- \| --- \| --- \| --- \| | \| 893 \| WBGene00017149:EGAP4.1(10375) \| ``` log2FC=0.436 padj=1.0 ``` \| [seedLoc = 112,117] Seq=TCCTTCACAGCTAATACCTCG \| ```          G      AA       CT  CT mRNA  3'  CTCCAT   TCGACA  TC   miRNA 5'  GAGGTA   GGTTGT  AG            T      GTA      AT  TT           3111111336212111331133 ``` \| SeedType = 1 NonSeedType = 2 \| \| --- \| --- \| --- \| --- \| --- \| --- \| | \|  \|  \|  \| [seedLoc = 132,137] Seq=TGCTCTACATACTACCTCA \| ```                           C   T mRNA  3' ACTCCATCAT   ACAT TCG  miRNA 5' TGAGGTAGTA   TGTA AGT                     GGT    T   T           1111111111666111131123 ``` \| SeedType = 1 NonSeedType = -1 \| \| --- \| --- \| --- \| --- \| --- \| --- \| | \| 894 \| WBGene00017153:E\_BE45912.2(10378) \| ``` log2FC=0.118 padj=0.982 ``` \| [seedLoc = 338,343] Seq=TACTCGCGAACCTATGTCACTACCTCT \| ```          T        CTGTA     GCGC   T mRNA  3'  CTCCATCA     TCCAA    TCA  miRNA 5'  GAGGTAGT     AGGTT    AGT           T                  GTAT   T           311111111554551111133331113 ``` \| SeedType = 1 NonSeedType = 1 \| \| --- \| --- \| --- \| --- \| --- \| --- \| | \| 895 \| WBGene00017164:eng-1(10388) \| ``` log2FC=0.116 padj=0.949 ``` \| [seedLoc = 256,261] Seq=CCCTGACCATAACTTGCACTACTTCC \| ```          C        CG       CCA   CC mRNA  3'  CTTCATCA  TTCAATA   GTC   miRNA 5'  GAGGTAGT  AGGTTGT   TAG            T                 A     TT           31121111154121112135521133 ``` \| SeedType = 3 NonSeedType = 1 \| \| --- \| --- \| --- \| --- \| --- \| --- \| | \| 896 \| WBGene00017257:F08F1.3(10437) \| ``` log2FC=-0.032 padj=0.994 ``` \| [seedLoc = 140,146] Seq=TGGCACCCTCCCTCTATCCTCA \| ```               T       CTCCCACGGT mRNA  3' ACTCC ATC TCC           miRNA 5' TGAGG TAG AGG                             T   TTGTATAGTT           11111511161113333333333 ``` \| SeedType = 2 NonSeedType = -1 \| \| --- \| --- \| --- \| --- \| --- \| --- \| | \| 897 \| WBGene00017285:F09E5.7(10457) \| ``` log2FC=0.09 padj=0.918 ``` \| [seedLoc = 335,341] Seq=TTGTTGACAATGTTTTACCCTCT \| ```          T  C     T G     GTTGTT mRNA  3'  CT CCATT T TAACA       miRNA 5'  GA GGTAG A GTTGT                T        T G     ATAGTT           31151111231321111333333 ``` \| SeedType = 3 NonSeedType = 1 \| \| --- \| --- \| --- \| --- \| --- \| --- \| | \| 898 \| WBGene00017296:F09E10.6(10464) \| ``` log2FC=1.263 padj=1.0 ``` \| [seedLoc = 38,43] Seq=GAACTTTTTCCAATTACCTCC \| ```          C        A  TTTT      G mRNA  3'  CTCCATTA CC      TCAA  miRNA 5'  GAGGTAGT GG      AGTT           T        A  TTGTAT                31111112131133336611114 ``` \| SeedType = 1 NonSeedType = -1 \| \| --- \| --- \| --- \| --- \| --- \| --- \| | \| 899 \| WBGene00017301:hach-1(10468) \|  \| [seedLoc = 315,320] Seq=AAACTGTTATTTTCTGTTTTTGTACCTCT \| ```          T      GTTTT     TTTAT      A mRNA  3'  CTCCAT     TGTCT     TGTCAA  miRNA 5'  GAGGTA     GTAGG     ATAGTT           T                TTGT                   31111114555522112333351211115 ``` \| SeedType = 1 NonSeedType = -1 \| \| --- \| --- \| --- \| --- \| --- \| --- \| | \| 900 \| WBGene00017317:attf-2(10483) \| ``` log2FC=-0.289 padj=0.294 ``` \| [seedLoc = 303,308] Seq=ATTTATTTATCTTATTACCTCC \| ```          C           T  T    TA mRNA  3'  CTCCATTATTC AT TATT   miRNA 5'  GAGGTAGTAGG TG ATAG            T           T  T    TT           3111111211213123111233 ``` \| SeedType = 1 NonSeedType = -1 \| \| --- \| --- \| --- \| --- \| --- \| --- \| | \| 901 \| WBGene00017340:F10D7.3(10500) \| ``` log2FC=-0.314 padj=0.566 ``` \| [seedLoc = 208,213] Seq=TTTGTTGCAATATTTCTGCCTCT \| ```          T       TTTA      TGTTT mRNA  3'  CTCCGTC    TAACGT      miRNA 5'  GAGGTAG    GTTGTA               T       TAG       TAGTT           31111211333521112133333 ``` \| SeedType = 2 NonSeedType = 1 \| \| --- \| --- \| --- \| --- \| --- \| --- \| | \| 902 \| WBGene00017355:F10E9.3(10510) \| ``` log2FC=0.086 padj=0.961 ``` \| [seedLoc = 75,80] Seq=CAAATTGACCCCGTTTTTAAAATCTACTTCA \| ```                  TAAAATTTTTGCC    TTAAAC   mRNA  3' ACTTCATC             CCAG         miRNA 5' TGAGGTAG             GGTT                          TA               GTATAGTT           111211113355555555455111233333366 ``` \| SeedType = 3 NonSeedType = 1 \| \| --- \| --- \| --- \| --- \| --- \| --- \| | \| 903 \| WBGene00017363:F10G2.2(10517) \| ``` log2FC=0.083 padj=1.0 ``` \| [seedLoc = 49,54] Seq=TAACTTCAATAGCCTCTACCTCA \| ```                          ACT    T mRNA  3' ACTCCATC TCCGATA   TCAA  miRNA 5' TGAGGTAG AGGTTGT   AGTT                   T       AT                 111111116111212133511115 ``` \| SeedType = 1 NonSeedType = 2 \| \| --- \| --- \| --- \| --- \| --- \| --- \| | \| 904 \| WBGene00017369:F10G7.5(10520) \| ``` log2FC=0.204 padj=0.816 ``` \| [seedLoc = 79,84] Seq=CATTTGTTTCCCGTTACCTCT \| ```          T       G   CTTTGTTTAC mRNA  3'  CTCCATT  CC           miRNA 5'  GAGGTAG  GG                    T       TA  TTGTATAGTT           3111111236113333333333 ``` \| SeedType = 1 NonSeedType = -1 \| \| --- \| --- \| --- \| --- \| --- \| --- \| | \| 905 \| WBGene00017378:F11C7.6(10525) \| ``` log2FC=-3.169 padj=1.0 ``` \| [seedLoc = 32,37] Seq=TAGCAAAAAATTTACATGTACCTCT \| ```          T      GTA       AAAACGAT mRNA  3'  CTCCAT   CATTTAA         miRNA 5'  GAGGTA   GTAGGTT                  T                GTATAGTT           3111111455111221133333333 ``` \| SeedType = 1 NonSeedType = 2 \| \| --- \| --- \| --- \| --- \| --- \| --- \| | \| 906 \| WBGene00017422:F13C5.1(10550) \| ``` log2FC=0.389 padj=1.0 ``` \| [seedLoc = 188,193] Seq=ATTTGTACATATGTTCTATCTCA \| ```                  T G  T       TA mRNA  3' ACTCTATC T TA ACATGTT   miRNA 5' TGAGGTAG A GT TGTATAG                    T G          TT           11112111313215111121233 ``` \| SeedType = 2 NonSeedType = -1 \| \| --- \| --- \| --- \| --- \| --- \| --- \| | \| 907 \| WBGene00017430:bcl-11(10557) \|  \| [seedLoc = 59,64] Seq=TCTCTCTCAATCCACCTACCTTT \| ```          T       CA  T   TCTCTCT mRNA  3'  TTCCATC  CC AAC        miRNA 5'  GAGGTAG  GG TTG                 T       TA      TATAGTT           32111111331151113333333 ``` \| SeedType = 3 NonSeedType = 1 \| \| --- \| --- \| --- \| --- \| --- \| --- \| | \| 908 \| WBGene00017434:F13H8.1(10560) \| ``` log2FC=-0.694 padj=0.285 ``` \| [seedLoc = 36,41] Seq=CACATAATTCCAACTATCTCA \| ```                   A  TTA      C mRNA  3' ACTCTATCA CC   ATA CA  miRNA 5' TGAGGTAGT GG   TAT GT                    A  TTG   A  T           1111211113113331116113 ``` \| SeedType = 2 NonSeedType = -1 \| \| --- \| --- \| --- \| --- \| --- \| --- \| | \| 909 \| WBGene00017439:F13H8.6(10565) \| ``` log2FC=0.019 padj=1.0 ``` \| [seedLoc = 60,65] Seq=ATCGGTACTTTTCTATCTCG \| ```          G           T     GCTA mRNA  3'  CTCTATC TTT  CATG     miRNA 5'  GAGGTAG AGG  GTAT              T       T   TT    AGTT           3111211161223611123333 ``` \| SeedType = 2 NonSeedType = -1 \| \| --- \| --- \| --- \| --- \| --- \| --- \| | \| 910 \| WBGene00017443:F13H8.11(10569) \| ``` log2FC=0.421 padj=0.654 ``` \| [seedLoc = 159,164] Seq=CGTAATGCATTCTATCTCA \| ```                  TT        ATGC mRNA  3' ACTCTATC     ACGTA     miRNA 5' TGAGGTAG     TGTAT                      TAGGT     AGTT           1111211133666112113333 ``` \| SeedType = 2 NonSeedType = -1 \| \| --- \| --- \| --- \| --- \| --- \| --- \| | \| 911 \| WBGene00017448:F14B8.6(10574) \| ``` log2FC=-0.27 padj=0.67 ``` \| [seedLoc = 72,77] Seq=ATCTGTCAGTTTCATTCCCACATCTACTTCG \| ```          G       TACA  CTTACTTT        TA mRNA  3'  CTTCATC    CC        GAC TGTC   miRNA 5'  GAGGTAG    GG        TTG ATAG            T       TA               T    TT           31121111335511555555552116121133 ``` \| SeedType = 3 NonSeedType = -1 \| \| --- \| --- \| --- \| --- \| --- \| --- \| | \|  \|  \|  \| [seedLoc = 182,188] Seq=GTTGGTATTACCTTACTCTCA \| ```              T         T    GTTG mRNA  3' ACTC CAT  TCCA TATG     miRNA 5' TGAG GTA  AGGT GTAT                      GT    T    AGTT           11115111661111321123333 ``` \| SeedType = 2 NonSeedType = 1 \| \| --- \| --- \| --- \| --- \| --- \| --- \| | \| 912 \| WBGene00017482:set-9(10597) \| ``` log2FC=-0.171 padj=0.742 ``` \| [seedLoc = 50,55] Seq=TCCCATATCTCTTTCTATCTCG \| ```          G       T   TC    CCCT mRNA  3'  CTCTATC TTC  TATA     miRNA 5'  GAGGTAG AGG  GTAT              T       T   TT    AGTT           3111211131213321113333 ``` \| SeedType = 2 NonSeedType = -1 \| \| --- \| --- \| --- \| --- \| --- \| --- \| | \| 913 \| WBGene00017528:F16G10.14(10615) \| ``` log2FC=-1.708 padj=1.0 ``` \| [seedLoc = 259,264] Seq=CGATTTTCAATCGTACCTCC \| ```          C      G       TTTTAGC mRNA  3'  CTCCAT   CTAAC        miRNA 5'  GAGGTA   GGTTG                 T      GTA     TATAGTT           3111111366121113333333 ``` \| SeedType = 1 NonSeedType = 1 \| \| --- \| --- \| --- \| --- \| --- \| --- \| | \| 914 \| WBGene00017534:F17A9.2(10618) \| ``` log2FC=0.005 padj=0.997 ``` \| [seedLoc = 165,171] Seq=ATTTGAAAAAAAAATTACTCTCA \| ```              T     AAAAAAAAGTTTA mRNA  3' ACTC CATTA              miRNA 5' TGAG GTAGT                                 AGGTTGTATAGTT           11115111213333333333333 ``` \| SeedType = 2 NonSeedType = -1 \| \| --- \| --- \| --- \| --- \| --- \| --- \| | \| 915 \| WBGene00017536:F17A9.4(10620) \| ``` log2FC=0.382 padj=0.055 ``` \| [seedLoc = 63,69] Seq=AAATTAAATAAATACTCACCTCC \| ```          C     C    A     A     A mRNA  3'  CTCCA TCAT  AATA ATTAA  miRNA 5'  GAGGT AGTA  TTGT TAGTT           T          GG    A                 311111511113611213112115 ``` \| SeedType = 2 NonSeedType = -1 \| \| --- \| --- \| --- \| --- \| --- \| --- \| | \| 916 \| WBGene00017563:F18C5.10(10638) \| ``` log2FC=-0.346 padj=0.435 ``` \| [seedLoc = 201,206] Seq=AGTAATATAAGTTCCAAACTACCTCC \| ```          C        AACC  G      ATGA mRNA  3'  CTCCATCA    TT AATATA     miRNA 5'  GAGGTAGT    AG TTGTAT              T              G      AGTT           31111111155551231121113333 ``` \| SeedType = 1 NonSeedType = -1 \| \| --- \| --- \| --- \| --- \| --- \| --- \| | \|  \|  \|  \| [seedLoc = 281,286] Seq=TTCAATTTTTCCGATTTACCTCT \| ```          T       TAG  TTTTTAACTT mRNA  3'  CTCCATT   CC           miRNA 5'  GAGGTAG   GG                    T       TA   TTGTATAGTT           31111112334113333333333 ``` \| SeedType = 1 NonSeedType = -1 \| \| --- \| --- \| --- \| --- \| --- \| --- \| | \|  \|  \|  \| [seedLoc = 355,360] Seq=CCCTGTCAATCTACCTTT \| ```          T                   CC mRNA  3'  TTCCATC   TAAC TGTC   miRNA 5'  GAGGTAG   GTTG ATAG            T       TAG    T    TT           3211111166621116121133 ``` \| SeedType = 3 NonSeedType = 1 \| \| --- \| --- \| --- \| --- \| --- \| --- \| | \|  \|  \|  \| [seedLoc = 420,425] Seq=AAAGAAACAACTCTACTTCT \| ```          T               AAGAAA mRNA  3'  CTTCATC  TCAACA       miRNA 5'  GAGGTAG  GGTTGT                T       TA      ATAGTT           3112111166211111333333 ``` \| SeedType = 3 NonSeedType = 1 \| \| --- \| --- \| --- \| --- \| --- \| --- \| | \| 917 \| WBGene00017569:F18E9.3(10641) \| ``` log2FC=0.213 padj=0.859 ``` \| [seedLoc = 72,77] Seq=TACATATACAATTTTAAAATCAATTACCTCT \| ```          T        ACTAAAAT          A  T mRNA  3'  CTCCATTA        TTTAACATAT CA  miRNA 5'  GAGGTAGT        AGGTTGTATA GT           T                             T           3111111215555555512211111115113 ``` \| SeedType = 1 NonSeedType = 1 \| \| --- \| --- \| --- \| --- \| --- \| --- \| | \| 918 \| WBGene00017571:jmjd-3.1(10643) \| ``` log2FC=-0.185 padj=0.862 ``` \| [seedLoc = 190,195] Seq=CAACTCAAGCCTTTCTATCTCA \| ```                  TT     AC      C mRNA  3' ACTCTATC  TCCGA    TCAA  miRNA 5' TGAGGTAG  AGGTT    AGTT                   T      GTAT                111121113511121336611115 ``` \| SeedType = 2 NonSeedType = 2 \| \| --- \| --- \| --- \| --- \| --- \| --- \| | \|  \|  \|  \| [seedLoc = 229,234] Seq=CTTTCGAAGAAAAACGTACCTCG \| ```          G      G  AAAAGAAGCTTTC mRNA  3'  CTCCAT CA              miRNA 5'  GAGGTA GT                       T         AGGTTGTATAGTT           31111114113333333333333 ``` \| SeedType = 1 NonSeedType = -1 \| \| --- \| --- \| --- \| --- \| --- \| --- \| | \|  \|  \|  \| [seedLoc = 310,315] Seq=CACTCCTCGCTCTCGCTATCTCT \| ```          T        C   C  TCC   C mRNA  3'  CTCTATCG TCT GC   TCA  miRNA 5'  GAGGTAGT AGG TG   AGT           T            T  TAT   T           31112111251123213331113 ``` \| SeedType = 2 NonSeedType = -1 \| \| --- \| --- \| --- \| --- \| --- \| --- \| | \| 919 \| WBGene00017583:F19B10.1(10650) \| ``` log2FC=-0.531 padj=0.617 ``` \| [seedLoc = 249,254] Seq=TTTCATATTGCGAAATACCTCC \| ```          C      AAAG  T    CTTT mRNA  3'  CTCCAT    CG TATA     miRNA 5'  GAGGTA    GT GTAT              T      GTAG  T    AGTT           3111111333312321113333 ``` \| SeedType = 1 NonSeedType = -1 \| \| --- \| --- \| --- \| --- \| --- \| --- \| | \| 920 \| WBGene00017601:ets-7(10659) \| ``` log2FC=0.1 padj=1.0 ``` \| [seedLoc = 92,97] Seq=AAACTTTAAGAATTTTATCTGCCTCC \| ```          C       TAT     GA  T    A mRNA  3'  CTCCGTC   TTTAA  AT TCAA  miRNA 5'  GAGGTAG   AGGTT  TA AGTT           T       T       G   T                31111211355122113511311115 ``` \| SeedType = 2 NonSeedType = 2 \| \| --- \| --- \| --- \| --- \| --- \| --- \| | \| 921 \| WBGene00017637:F20D6.8(10677) \| ``` log2FC=0.144 padj=0.974 ``` \| [seedLoc = 486,492] Seq=TCCTGTGCTGCATTTCCCCAATTTACACTCT \| ```          T   A    TAACCCCTTTA  T      CT mRNA  3'  CTC CATT           CG CGTGTC   miRNA 5'  GAG GTAG           GT GTATAG            T        TAG          T      TT           3111511123335555555512312121133 ``` \| SeedType = 2 NonSeedType = -1 \| \| --- \| --- \| --- \| --- \| --- \| --- \| | \| 922 \| WBGene00017641:csr-1(10680) \| ``` log2FC=-0.034 padj=0.977 ``` \| [seedLoc = 359,364] Seq=AAATTGTGGAAAGAATATCCTACCTTT \| ```          T       CT  AAGA  G       A mRNA  3'  TTCCATC  AT    AA GTGTTAA  miRNA 5'  GAGGTAG  TA    TT TATAGTT           T           GG    G                   321111115511334511321212115 ``` \| SeedType = 3 NonSeedType = -1 \| \| --- \| --- \| --- \| --- \| --- \| --- \| | \| 923 \| WBGene00017676:F21F3.7(10704) \| ``` log2FC=0.069 padj=0.982 ``` \| [seedLoc = 292,297] Seq=CTATGTACATAGTTTCTACTACTACCTTA \| ```                     ATCTTTGAT      ATC mRNA  3' ATTCCATCATC         ACATGT    miRNA 5' TGAGGTAGTAG         TGTATA                        GT             GTT           12111111111335555455111121333 ``` \| SeedType = 3 NonSeedType = -1 \| \| --- \| --- \| --- \| --- \| --- \| --- \| | \| 924 \| WBGene00017699:flcn-1(10719) \| ``` log2FC=-0.061 padj=0.975 ``` \| [seedLoc = 556,561] Seq=GTTTCCGTAATTTTCCCTACCTCA \| ```                  CCT      GCCTTTG mRNA  3' ACTCCATC   TTTAAT        miRNA 5' TGAGGTAG   AGGTTG                         T        TATAGTT           111111113551221123333333 ``` \| SeedType = 1 NonSeedType = 2 \| \| --- \| --- \| --- \| --- \| --- \| --- \| | \|  \|  \|  \| [seedLoc = 709,715] Seq=GAGCTTTTTCACCCACAATACCATCT \| ```          T  A    AA  C   CTTTT    G mRNA  3'  CT CCAT  CA CCA     TCGA  miRNA 5'  GA GGTA  GT GGT     AGTT           T           A   TGTAT                31151111551131113333311214 ``` \| SeedType = 3 NonSeedType = 1 \| \| --- \| --- \| --- \| --- \| --- \| --- \| | \| 925 \| WBGene00017717:F22F4.4(10730) \| ``` log2FC=-2.365 padj=1.0 ``` \| [seedLoc = 159,164] Seq=AGACGTGCATTTTGCACTACTTCA \| ```                   CG   T         A mRNA  3' ACTTCATCA  TTT ACGTG CAG  miRNA 5' TGAGGTAGT  AGG TGTAT GTT                         T     A               1112111115412231121261125 ``` \| SeedType = 3 NonSeedType = -1 \| \| --- \| --- \| --- \| --- \| --- \| --- \| | \|  \|  \|  \| [seedLoc = 226,231] Seq=AATTTAGCATGTTTAGTACCTCT \| ```          T      G    GT   ATTTAA mRNA  3'  CTCCAT ATTT  ACG       miRNA 5'  GAGGTA TAGG  TGT                T      G    T    ATAGTT           31111113112235112333333 ``` \| SeedType = 1 NonSeedType = -1 \| \| --- \| --- \| --- \| --- \| --- \| --- \| | \|  \|  \|  \| [seedLoc = 405,410] Seq=TCATATGGTACCAAGTACCTCA \| ```                 G A   TG    ACT mRNA  3' ACTCCAT A CCA  GTAT    miRNA 5' TGAGGTA T GGT  TATA                    G A   TG    GTT           1111111313111332111333 ``` \| SeedType = 1 NonSeedType = 1 \| \| --- \| --- \| --- \| --- \| --- \| --- \| | \| 926 \| WBGene00017760:F23H11.4(10764) \| ``` log2FC=0.082 padj=0.944 ``` \| [seedLoc = 177,182] Seq=TTCTGTATAAAGTCTACTACCTTA \| ```                      GA        TT mRNA  3' ATTCCATCATCT  AATATGTC   miRNA 5' TGAGGTAGTAGG  TTGTATAG                                  TT           121111111112451121121133 ``` \| SeedType = 3 NonSeedType = -1 \| \| --- \| --- \| --- \| --- \| --- \| --- \| | \|  \|  \|  \| [seedLoc = 186,191] Seq=CTGTATAAAGTCTACTACCTTAT \| ```          T            GA           mRNA  3'  ATTCCATCATCT  AATATGTC   miRNA 5'  TGAGGTAGTAGG  TTGTATAG                                   TT           5121111111112451121121166 ``` \| SeedType = 3 NonSeedType = -1 \| \| --- \| --- \| --- \| --- \| --- \| --- \| | \| 927 \| WBGene00017766:peli-1(10768) \|  \| [seedLoc = 123,128] Seq=CCTTTCAAAACCGGTCTCTATCTCT \| ```          T       TCTGG    AACTTTCC mRNA  3'  CTCTATC     CCAA         miRNA 5'  GAGGTAG     GGTT                  T       TA       GTATAGTT           3111211133544111133333333 ``` \| SeedType = 2 NonSeedType = 1 \| \| --- \| --- \| --- \| --- \| --- \| --- \| | \| 928 \| WBGene00017769:hmgs-1(10771) \| ``` log2FC=0.298 padj=0.262 ``` \| [seedLoc = 61,66] Seq=TCGTTTGCCATAGTCTACTTCT \| ```          T       TGA  C   TTGCT mRNA  3'  CTTCATC   TA CGT      miRNA 5'  GAGGTAG   GT GTA               T       TAG  T   TAGTT           3112111133321312133333 ``` \| SeedType = 3 NonSeedType = -1 \| \| --- \| --- \| --- \| --- \| --- \| --- \| | \| 929 \| WBGene00017774:nop-1(10777) \| ``` log2FC=-0.058 padj=0.965 ``` \| [seedLoc = 229,235] Seq=TAGAATGCCCCTGATGTTTCTATCCTCT \| ```          T    T   TTTGTA    C     AGAT mRNA  3'  CTCC ATC      GTCC  CGTA     miRNA 5'  GAGG TAG      TAGG  GTAT              T                  TT    AGTT           31111511155545521113612113333 ``` \| SeedType = 2 NonSeedType = -1 \| \| --- \| --- \| --- \| --- \| --- \| --- \| | \| 930 \| WBGene00017775:F25B5.3(10779) \| ``` log2FC=-0.371 padj=0.146 ``` \| [seedLoc = 142,148] Seq=GCCATTACTGATGAACTACCTAC \| ```          CA       AG   T   TACCG mRNA  3'   TCCATCA  TAG CAT      miRNA 5'   AGGTAGT  GTT GTA               TG       AG       TAGTT           33111111133212511133333 ``` \| SeedType = 3 NonSeedType = -1 \| \| --- \| --- \| --- \| --- \| --- \| --- \| | \| 931 \| WBGene00017814:F26A10.2(10808) \| ``` log2FC=-0.414 padj=0.278 ``` \| [seedLoc = 151,156] Seq=GCACAAATATCCCACTACCTTT \| ```          T        C  T   AACACG mRNA  3'  TTCCATCA CC ATA       miRNA 5'  GAGGTAGT GG TGT                T        A  T   ATAGTT           3211111113113121333333 ``` \| SeedType = 3 NonSeedType = -1 \| \| --- \| --- \| --- \| --- \| --- \| --- \| | \|  \|  \|  \| [seedLoc = 812,817] Seq=CTTTAAACATTTCTTGCTACTTCT \| ```          T           TTT   AATTTC mRNA  3'  CTTCATCGTTC   ACA       miRNA 5'  GAGGTAGTAGG   TGT                T           T     ATAGTT           311211112121355111333333 ``` \| SeedType = 3 NonSeedType = -1 \| \| --- \| --- \| --- \| --- \| --- \| --- \| | \| 932 \| WBGene00017826:tag-340(10815) \| ``` log2FC=0.306 padj=0.559 ``` \| [seedLoc = 134,139] Seq=GACTAATCACTTTTTAAAGTTTCATACCTCC \| ```          C      ACTTTGAAATTT    C         mRNA  3'  CTCCAT            TTCA TA ATCAG miRNA 5'  GAGGTA            AGGT GT TAGTT          T      GT              T  A                31111113355545555551211321611112 ``` \| SeedType = 1 NonSeedType = 2 \| \| --- \| --- \| --- \| --- \| --- \| --- \| | \| 933 \| WBGene00017836:F26F12.5(10826) \| ``` log2FC=-0.32 padj=0.952 ``` \| [seedLoc = 37,43] Seq=ATTTGTATAGTGGTAACCTCC \| ```          C    A  G            TA mRNA  3'  CTCC AT GT  GATATGTT   miRNA 5'  GAGG TA TA  TTGTATAG            T       G  GG        TT           31111511321662121121233 ``` \| SeedType = 2 NonSeedType = -1 \| \| --- \| --- \| --- \| --- \| --- \| --- \| | \| 934 \| WBGene00017864:pcca-1(10842) \| ``` log2FC=0.244 padj=0.414 ``` \| [seedLoc = 160,166] Seq=ATTTTTGATGCCATTACCCTCA \| ```             C          TAGTTTTTA mRNA  3' ACT CCATTA CCG          miRNA 5' TGA GGTAGT GGT                             A   TGTATAGTT           11151111216112333333333 ``` \| SeedType = 3 NonSeedType = 2 \| \| --- \| --- \| --- \| --- \| --- \| --- \| | \| 935 \| WBGene00017885:dip-2(10850) \| ``` log2FC=0.303 padj=0.292 ``` \| [seedLoc = 101,106] Seq=CACTTTTCTTGCCTTTCTACCTTT \| ```          T       TT    TTCTTT   C mRNA  3'  TTCCATC  TCCG      TCA  miRNA 5'  GAGGTAG  AGGT      AGT           T       T     TGTAT    T           321111113511123333351113 ``` \| SeedType = 3 NonSeedType = 2 \| \| --- \| --- \| --- \| --- \| --- \| --- \| | \| 936 \| WBGene00017893:F28B4.4(10859) \| ``` log2FC=-0.564 padj=1.0 ``` \| [seedLoc = 196,201] Seq=ATTTGGAAGGTTCGCTACCTTT \| ```          T        CTTGGAAGGTTTA mRNA  3'  TTCCATCG              miRNA 5'  GAGGTAGT                       T        AGGTTGTATAGTT           3211111123333333333333 ``` \| SeedType = 3 NonSeedType = -1 \| \| --- \| --- \| --- \| --- \| --- \| --- \| | \| 937 \| WBGene00017894:F28B12.1(10860) \| ``` log2FC=0.302 padj=0.942 ``` \| [seedLoc = 29,33] Seq=CGACACGAGCAAATATCCTCC \| ```          C    T  AAA G    CAGC   mRNA  3'  CTCC AT   C AGCA       miRNA 5'  GAGG TA   G TTGT                T       GTA G    ATAGTT           31111511333131211333366 ``` \| SeedType = 2 NonSeedType = -1 \| \| --- \| --- \| --- \| --- \| --- \| --- \| | \| 938 \| WBGene00017950:F31E3.2(10894) \| ``` log2FC=-0.918 padj=1.0 ``` \| [seedLoc = 90,96] Seq=TAAAATACCCTGTTACCATCA \| ```             A               AAAT mRNA  3' ACT CCATTGTCC  CATA     miRNA 5' TGA GGTAGTAGG  GTAT                           TT    AGTT           11151111221116611113333 ``` \| SeedType = 3 NonSeedType = -1 \| \| --- \| --- \| --- \| --- \| --- \| --- \| | \| 939 \| WBGene00017954:F31E8.4(10896) \| ``` log2FC=-0.226 padj=0.885 ``` \| [seedLoc = 100,105] Seq=TTCTGTTGTTACTGGTACCTCA \| ```                 G   ATTGT    TT mRNA  3' ACTCCAT GTC     TGTC   miRNA 5' TGAGGTA TAG     ATAG                   G   GTTGT    TT           1111111321133333121133 ``` \| SeedType = 1 NonSeedType = -1 \| \| --- \| --- \| --- \| --- \| --- \| --- \| | \| 940 \| WBGene00017974:vps-41(10912) \| ``` log2FC=-0.026 padj=0.991 ``` \| [seedLoc = 345,350] Seq=TTTCACTTGGCTTGTTCTACTTCT \| ```          T       TT      TTCACTTT mRNA  3'  CTTCATC  GTTCGG         miRNA 5'  GAGGTAG  TAGGTT                  T               GTATAGTT           311211115521212233333333 ``` \| SeedType = 3 NonSeedType = 2 \| \| --- \| --- \| --- \| --- \| --- \| --- \| | \| 941 \| WBGene00018010:F33E11.2(10945) \| ``` log2FC=-0.1 padj=0.924 ``` \| [seedLoc = 63,69] Seq=ATTCCCACCGGTTATGATACTCTCT \| ```          T   T   AG    G C  CCCTTA mRNA  3'  CTC CAT  TATT G CA       miRNA 5'  GAG GTA  GTAG T GT                T             G T  ATAGTT           3111511154211232311333333 ``` \| SeedType = 2 NonSeedType = -1 \| \| --- \| --- \| --- \| --- \| --- \| --- \| | \| 942 \| WBGene00018015:F33G12.3(10950) \| ``` log2FC=0.104 padj=0.937 ``` \| [seedLoc = 165,171] Seq=ATGAAATTGATGAATTACTCTCC \| ```          C   T     AG   TTAAAGTA mRNA  3'  CTC CATTA  TAG         miRNA 5'  GAG GTAGT  GTT                  T         AG   GTATAGTT           31115111213321233333333 ``` \| SeedType = 2 NonSeedType = -1 \| \| --- \| --- \| --- \| --- \| --- \| --- \| | \| 943 \| WBGene00018041:F35D2.3(10967) \| ``` log2FC=0.955 padj=0.609 ``` \| [seedLoc = 189,195] Seq=TGTTCATCAACAATTAACCTCA \| ```               A    A     TACTTGT mRNA  3' ACTCC ATTA  CAAC        miRNA 5' TGAGG TAGT  GTTG                           AG    TATAGTT           11111511213611113333333 ``` \| SeedType = 2 NonSeedType = 1 \| \| --- \| --- \| --- \| --- \| --- \| --- \| | \| 944 \| WBGene00018055:F35F10.5(10979) \| ``` log2FC=-0.645 padj=0.699 ``` \| [seedLoc = 28,33] Seq=AAACTTTTCGAATTAATTCCTACCTTT \| ```          T       CTTA   A   TTT    A mRNA  3'  TTCCATC    ATT AGC   TCAA  miRNA 5'  GAGGTAG    TAG TTG   AGTT           T              G   TAT                321111115555112312133311115 ``` \| SeedType = 3 NonSeedType = -1 \| \| --- \| --- \| --- \| --- \| --- \| --- \| | \| 945 \| WBGene00018073:F35H10.10(10992) \| ``` log2FC=-0.165 padj=0.8 ``` \| [seedLoc = 170,175] Seq=CAACTACTTAGCCTCTGCCTCC \| ```          C              TC     C mRNA  3'  CTCCGTC TCCGAT  ATCAA  miRNA 5'  GAGGTAG AGGTTG  TAGTT           T       T      TA                 31111211611121233111115 ``` \| SeedType = 2 NonSeedType = 2 \| \| --- \| --- \| --- \| --- \| --- \| --- \| | \| 946 \| WBGene00018077:F35H12.5(10996) \| ``` log2FC=0.593 padj=0.222 ``` \| [seedLoc = 58,64] Seq=GAATTATGAATTTCGATTGTTACTCTCT \| ```          T   T    G      TTTAA       G mRNA  3'  CTC CATT  TTAGC     GTATTAA  miRNA 5'  GAG GTAG  GGTTG     TATAGTT           T        TA                             31115111236221215555521112114 ``` \| SeedType = 2 NonSeedType = 2 \| \| --- \| --- \| --- \| --- \| --- \| --- \| | \| 947 \| WBGene00018090:F36D4.5(11005) \| ``` log2FC=-0.03 padj=0.984 ``` \| [seedLoc = 91,97] Seq=CTATAATTCCTGATAAATCTTCCTACCCTCC \| ```          C  C     CTTCTAAATA    TTA        mRNA  3'  CT CCATC          GTCC   ATATC   miRNA 5'  GA GGTAG          TAGG   TATAG            T                      TTG     TT           311511111555555555521113331111166 ``` \| SeedType = 3 NonSeedType = -1 \| \| --- \| --- \| --- \| --- \| --- \| --- \| | \| 948 \| WBGene00018099:ztf-28(11009) \| ``` log2FC=-0.299 padj=0.518 ``` \| [seedLoc = 38,43] Seq=ATGTTTATAGTTCTACCTCA \| ```                  TT       TTGTA mRNA  3' ACTCCATC    GATAT      miRNA 5' TGAGGTAG    TTGTA                       TAGG     TAGTT           1111111133662121133333 ``` \| SeedType = 1 NonSeedType = -1 \| \| --- \| --- \| --- \| --- \| --- \| --- \| | \| 949 \| WBGene00018101:math-27(11014) \| ``` log2FC=-0.082 padj=0.966 ``` \| [seedLoc = 842,848] Seq=TCGTTCATAATGTGTCATACCGTCA \| ```             G    A T  G     CTTGCT mRNA  3' ACT CCAT C GT TAATA       miRNA 5' TGA GGTA G TA GTTGT                             G     ATAGTT           1114111151521321121333333 ``` \| SeedType = 3 NonSeedType = -1 \| \| --- \| --- \| --- \| --- \| --- \| --- \| | \| 950 \| WBGene00018164:sup-36(11058) \| ``` log2FC=-0.022 padj=0.99 ``` \| [seedLoc = 90,95] Seq=AAGCTCAATACTTTCACTTTTTACCTCC \| ```          C       T    CTTT    AC    A mRNA  3'  CTCCATT TTCA    CATA  TCGA  miRNA 5'  GAGGTAG AGGT    GTAT  AGTT           T       T    T                         3111111231211355511115511215 ``` \| SeedType = 1 NonSeedType = 2 \| \| --- \| --- \| --- \| --- \| --- \| --- \| | \| 951 \| WBGene00018210:F39G3.5(11083) \| ``` log2FC=0.075 padj=0.98 ``` \| [seedLoc = 334,340] Seq=TGTTCATTATTCTCTACACTCT \| ```          T   A        TATTACTTGT mRNA  3'  CTC CATC TCT           miRNA 5'  GAG GTAG AGG                    T        T   TTGTATAGTT           31115111161123333333333 ``` \| SeedType = 2 NonSeedType = -1 \| \| --- \| --- \| --- \| --- \| --- \| --- \| | \| 952 \| WBGene00018213:F39H12.2(11085) \| ``` log2FC=-0.675 padj=1.0 ``` \| [seedLoc = 73,79] Seq=TTAATAACAAAAAAACATTACACTCC \| ```          C   A     CAAAAA    ATAATT mRNA  3'  CTC CATTA      AACA       miRNA 5'  GAG GTAGT      TTGT                T         AGG       ATAGTT           31115111213335551111333333 ``` \| SeedType = 2 NonSeedType = 1 \| \| --- \| --- \| --- \| --- \| --- \| --- \| | \| 953 \| WBGene00018214:F39H12.3(11086) \| ``` log2FC=0.3 padj=0.484 ``` \| [seedLoc = 118,124] Seq=TTCTAACTTCCTATCGTATACACTCG \| ```          G   A   ATGC     TT      TT mRNA  3'  CTC CAT    TATCC  CA ATC   miRNA 5'  GAG GTA    GTAGG  GT TAG            T                TT  A   TT           311151115545211113311611133 ``` \| SeedType = 2 NonSeedType = -1 \| \| --- \| --- \| --- \| --- \| --- \| --- \| | \| 954 \| WBGene00018259:F41B4.3(11115) \| ``` log2FC=0.145 padj=0.945 ``` \| [seedLoc = 39,44] Seq=GTGAATACCACTAACGATACCTCA \| ```                 AG  A   C    AGTG mRNA  3' ACTCCAT  CA TCA CATA     miRNA 5' TGAGGTA  GT GGT GTAT                         A   T    AGTT           111111154113211311113333 ``` \| SeedType = 1 NonSeedType = 2 \| \| --- \| --- \| --- \| --- \| --- \| --- \| | \| 955 \| WBGene00018281:tbc-18(11130) \| ``` log2FC=-0.221 padj=0.934 ``` \| [seedLoc = 28,34] Seq=TAATTGTTAGCATATTACCATCG \| ```          G  A       A           T mRNA  3'  CT CCATTAT CGAT TGTTAA  miRNA 5'  GA GGTAGTA GTTG ATAGTT           T          G    T                  311511112113121261212115 ``` \| SeedType = 3 NonSeedType = -1 \| \| --- \| --- \| --- \| --- \| --- \| --- \| | \|  \|  \|  \| [seedLoc = 45,50] Seq=AGCATATTACCATCGT \| ```          T   A       A            mRNA  3'  GCT CCATTAT CGA         miRNA 5'  TGA GGTAGTA GTT                              G   GTATAGTT           521151111211312166666666 ``` \| SeedType = 3 NonSeedType = -1 \| \| --- \| --- \| --- \| --- \| --- \| --- \| | \| 956 \| WBGene00018285:smk-1(11134) \| ``` log2FC=-0.155 padj=0.773 ``` \| [seedLoc = 28,33] Seq=TTCCCGTCTACCTCTATCTCA \| ```                       TCTGCCCTT mRNA  3' ACTCTATC TCCA          miRNA 5' TGAGGTAG AGGT                           T    TGTATAGTT           1111211161111333333333 ``` \| SeedType = 2 NonSeedType = 1 \| \| --- \| --- \| --- \| --- \| --- \| --- \| | \|  \|  \|  \| [seedLoc = 290,295] Seq=TTTTCCACAAAGCTACCTTG \| ```          G        A      CCTTTT mRNA  3'  TTCCATCG   AACA       miRNA 5'  GAGGTAGT   TTGT                T        AGG    ATAGTT           3211111123661111333333 ``` \| SeedType = 3 NonSeedType = 1 \| \| --- \| --- \| --- \| --- \| --- \| --- \| | \| 957 \| WBGene00018286:F41E6.5(11136) \| ``` log2FC=-0.835 padj=2.61728720131378E-4 ``` \| [seedLoc = 99,104] Seq=CAGCTGTTCTTGGACTACTTCA \| ```                   GGTTCT       C mRNA  3' ACTTCATCA       TGTCGA  miRNA 5' TGAGGTAGT       ATAGTT                    AGGTTGT                  11121111133333361211215 ``` \| SeedType = 3 NonSeedType = -1 \| \| --- \| --- \| --- \| --- \| --- \| --- \| | \| 958 \| WBGene00018304:agr-1(11150) \| ``` log2FC=0.446 padj=0.842 ``` \| [seedLoc = 49,54] Seq=CAATTATGTTTTTTTCTATCTCA \| ```                  T   TTT       C mRNA  3' ACTCTATC TTT   GTATTAA  miRNA 5' TGAGGTAG AGG   TATAGTT                   T   TTG                   11112111312233321112115 ``` \| SeedType = 2 NonSeedType = -1 \| \| --- \| --- \| --- \| --- \| --- \| --- \| | \| 959 \| WBGene00018338:F42A9.9(11170) \| ``` log2FC=0.4 padj=1.0 ``` \| [seedLoc = 188,194] Seq=AAGCAAATAATTAACTAACCTCT \| ```          T    A    A      AACGAA mRNA  3'  CTCC ATCA TTAATA       miRNA 5'  GAGG TAGT GGTTGT                T         A      ATAGTT           31111511113221121333333 ``` \| SeedType = 2 NonSeedType = 2 \| \| --- \| --- \| --- \| --- \| --- \| --- \| | \| 960 \| WBGene00018367:F42H10.3(11192) \| ``` log2FC=0.191 padj=0.73 ``` \| [seedLoc = 25,30] Seq=ACCCACGCACATTTTCTATCTCT \| ```          T       T    C   CACCCA mRNA  3'  CTCTATC TTTA ACG       miRNA 5'  GAGGTAG AGGT TGT                T       T        ATAGTT           31112111312215112333333 ``` \| SeedType = 2 NonSeedType = 2 \| \| --- \| --- \| --- \| --- \| --- \| --- \| | \| 961 \| WBGene00018390:F43E2.1(11208) \| ``` log2FC=-0.118 padj=0.912 ``` \| [seedLoc = 153,159] Seq=GAAGTGGAAAATAAAATACACTCG \| ```          G   A   AAAA   AAG  G  G mRNA  3'  CTC CAT    TAA   GT AA  miRNA 5'  GAG GTA    GTT   TA TT           T       GTAG   GTA  G              311151113333211333213114 ``` \| SeedType = 2 NonSeedType = -1 \| \| --- \| --- \| --- \| --- \| --- \| --- \| | \|  \|  \|  \| [seedLoc = 301,306] Seq=CGATATGCCATGACATTTTTCTCTGCCTCT \| ```          T           TTTTACAGTAC     AGC mRNA  3'  CTCCGTC TCT           CGTAT    miRNA 5'  GAGGTAG AGG           GTATA             T       T   TT              GTT           3111121161123355555455512111333 ``` \| SeedType = 2 NonSeedType = -1 \| \| --- \| --- \| --- \| --- \| --- \| --- \| | \| 962 \| WBGene00018395:mtch-1(11214) \| ``` log2FC=0.008 padj=0.994 ``` \| [seedLoc = 257,263] Seq=TTTTGTAAACAAACTACACTCA \| ```              A     AA         TT mRNA  3' ACTC CATCA  CAA ATGTT   miRNA 5' TGAG GTAGT  GTT TATAG                      AG   G     TT           11115111113311161121233 ``` \| SeedType = 2 NonSeedType = 1 \| \| --- \| --- \| --- \| --- \| --- \| --- \| | \| 963 \| WBGene00018398:sptl-2(11217) \| ``` log2FC=0.557 padj=0.004 ``` \| [seedLoc = 59,64] Seq=ATTCAAACGGGCTTTCCTACTTCT \| ```          T       CTT  G    AACTTA mRNA  3'  CTTCATC   TC GGCA       miRNA 5'  GAGGTAG   AG TTGT                T       T    G    ATAGTT           311211113551132211333333 ``` \| SeedType = 3 NonSeedType = -1 \| \| --- \| --- \| --- \| --- \| --- \| --- \| | \| 964 \| WBGene00018409:F44B9.8(11226) \| ``` log2FC=0.033 padj=0.982 ``` \| [seedLoc = 109,115] Seq=TTTTAAGTGACCTCATTTATTATCCTCG \| ```          G    T    TTTAC     TGAATTTT mRNA  3'  CTCC ATTA     TCCAG         miRNA 5'  GAGG TAGT     AGGTT                  T                   GTATAGTT           3111151121555551111233333333 ``` \| SeedType = 2 NonSeedType = 1 \| \| --- \| --- \| --- \| --- \| --- \| --- \| | \| 965 \| WBGene00018417:F44E2.3(11231) \| ``` log2FC=-0.033 padj=0.985 ``` \| [seedLoc = 35,40] Seq=GCCTGTGCACCTGACTTTACCTCG \| ```          G       TCA            CG mRNA  3'  CTCCATT   GTCC ACGTGTC   miRNA 5'  GAGGTAG   TAGG TGTATAG            T              T       TT           3111111255521116112121133 ``` \| SeedType = 1 NonSeedType = -1 \| \| --- \| --- \| --- \| --- \| --- \| --- \| | \| 966 \| WBGene00018431:F44E7.9(11244) \| ``` log2FC=0.116 padj=0.899 ``` \| [seedLoc = 45,50] Seq=GAATTTTGTTTAGTTTACCTCG \| ```          G       TGATTTGTTTTAAG mRNA  3'  CTCCATT               miRNA 5'  GAGGTAG                        T       TAGGTTGTATAGTT           3111111233333333333333 ``` \| SeedType = 1 NonSeedType = -1 \| \| --- \| --- \| --- \| --- \| --- \| --- \| | \| 967 \| WBGene00018466:F45E1.5(11261) \| ``` log2FC=-0.726 padj=0.563 ``` \| [seedLoc = 54,59] Seq=CGTTGTTAACATTCCTATCTCA \| ```                  CTTA         GC mRNA  3' ACTCTATC    CAAT TGTT   miRNA 5' TGAGGTAG    GTTG ATAG                    TAG     T    TT           11112111333511126121233 ``` \| SeedType = 2 NonSeedType = 1 \| \| --- \| --- \| --- \| --- \| --- \| --- \| | \| 968 \| WBGene00018467:sdpn-1(11262) \| ``` log2FC=-0.187 padj=0.715 ``` \| [seedLoc = 181,187] Seq=TACTTATAATCTGTACTCTCC \| ```          C   T                 T mRNA  3'  CTC CAT GTCTAATAT TCA  miRNA 5'  GAG GTA TAGGTTGTA AGT           T       G         T   T           31115111621121121161113 ``` \| SeedType = 2 NonSeedType = 2 \| \| --- \| --- \| --- \| --- \| --- \| --- \| | \| 969 \| WBGene00018468:cla-1(11264) \|  \| [seedLoc = 57,62] Seq=TTCTGAATACGTGTCTACCTCT \| ```          T       T  G     A   TT mRNA  3'  CTCCATC GT C ATA GTC   miRNA 5'  GAGGTAG TA G TGT TAG            T          G T   A   TT           31111111521316121321133 ``` \| SeedType = 1 NonSeedType = -1 \| \| --- \| --- \| --- \| --- \| --- \| --- \| | \| 970 \| WBGene00018486:glb-16(11277) \| ``` log2FC=-0.16 padj=1.0 ``` \| [seedLoc = 141,147] Seq=AGACTGACCTATTTAACAGATACACTCC \| ```          C   A   AGACAATT               A mRNA  3'  CTC CAT        TATCCAG    TCAG  miRNA 5'  GAG GTA        GTAGGTT    AGTT           T                      GTAT                31115111545555552111112666611125 ``` \| SeedType = 2 NonSeedType = 1 \| \| --- \| --- \| --- \| --- \| --- \| --- \| | \| 971 \| WBGene00018519:F46H5.3(11301) \| ``` log2FC=0.174 padj=0.792 ``` \| [seedLoc = 126,132] Seq=TAGTGTAAAATTTTGTATTACGCTCC \| ```          C   G      GTT    A    G T mRNA  3'  CTC CATTAT   TTAA ATGT A  miRNA 5'  GAG GTAGTA   GGTT TATA T           T                 G    G T           31114111211455221131121313 ``` \| SeedType = 2 NonSeedType = 2 \| \| --- \| --- \| --- \| --- \| --- \| --- \| | \| 972 \| WBGene00018547:clec-78(11317) \| ``` log2FC=-0.321 padj=0.943 ``` \| [seedLoc = 207,212] Seq=TCGTGTAAATACCGTACCTCA \| ```                 G     TAA    GCT mRNA  3' ACTCCAT   CCA   ATGT    miRNA 5' TGAGGTA   GGT   TATA                    GTA   TG     GTT           11111113661113351121333 ``` \| SeedType = 1 NonSeedType = 1 \| \| --- \| --- \| --- \| --- \| --- \| --- \| | \| 973 \| WBGene00018564:F47D12.9(11321) \| ``` log2FC=-0.416 padj=0.069 ``` \| [seedLoc = 140,145] Seq=TTTCTCCCTTTCTCTATCTCT \| ```          T           TTCCCTCTTT mRNA  3'  CTCTATC TCT           miRNA 5'  GAGGTAG AGG                    T       T   TTGTATAGTT           3111211161123333333333 ``` \| SeedType = 2 NonSeedType = -1 \| \| --- \| --- \| --- \| --- \| --- \| --- \| | \| 974 \| WBGene00018565:F47E1.1(11323) \| ``` log2FC=-1.754 padj=1.0 ``` \| [seedLoc = 67,72] Seq=TTTGTTGGAGTCTCTACCTCT \| ```          T             G  TGTTT mRNA  3'  CTCCATC TCTGA GT      miRNA 5'  GAGGTAG AGGTT TA               T       T     G  TAGTT           3111111161122132133333 ``` \| SeedType = 1 NonSeedType = 2 \| \| --- \| --- \| --- \| --- \| --- \| --- \| | \| 975 \| WBGene00018572:lin-42(11328) \| ``` log2FC=-0.052 padj=0.983 ``` \| [seedLoc = 43,48] Seq=AAGGGGACCCTACCCCAA \| ```          A  C           G GAA    mRNA  3'  AC CCATC  CCAG G       miRNA 5'  TG GGTAG  GGTT T                   A     TA    G ATAGTT           51131111166111232333666 ``` \| SeedType = 3 NonSeedType = 1 \| \| --- \| --- \| --- \| --- \| --- \| --- \| | \| 976 \| WBGene00018572:lin-42(11329) \| ``` log2FC=-0.052 padj=0.983 ``` \| [seedLoc = 501,507] Seq=CTTTGTCTAGTTTTTTACCGTCT \| ```          T  G     T      C    TC mRNA  3'  CT CCATT TTTGAT TGTT   miRNA 5'  GA GGTAG AGGTTG ATAG            T        T      T    TT           31141111231222123121233 ``` \| SeedType = 3 NonSeedType = 2 \| \| --- \| --- \| --- \| --- \| --- \| --- \| | \| 977 \| WBGene00018578:F47G3.3(11332) \| ``` log2FC=0.535 padj=0.637 ``` \| [seedLoc = 231,237] Seq=AACAATACCTGTTTTCAAAGTTTTACCCTCC \| ```          C  C     TTGAAACTTTT     TA   A      mRNA  3'  CT CCATT           GTCCA  ACA       miRNA 5'  GA GGTAG           TAGGT  TGT                T                             ATAGTT           311511112554555555552111155111366666 ``` \| SeedType = 3 NonSeedType = 1 \| \| --- \| --- \| --- \| --- \| --- \| --- \| | \| 978 \| WBGene00018602:F48D6.4(11346) \| ``` log2FC=-0.706 padj=0.002 ``` \| [seedLoc = 155,161] Seq=TTGTACTAGAGTAGGAACATTTGTACTCTCT \| ```          T   T   GTTTA   GGATGAGAT         mRNA  3'  CTC CAT     CAA         CATGTT   miRNA 5'  GAG GTA     GTT         GTATAG            T       GTAG                   TT           311151113333511144554545511121266 ``` \| SeedType = 2 NonSeedType = 1 \| \| --- \| --- \| --- \| --- \| --- \| --- \| | \| 979 \| WBGene00018628:F49D11.4(11361) \| ``` log2FC=-1.708 padj=1.0 ``` \| [seedLoc = 36,41] Seq=TGGTGGGCAAATTGCTACCTTA \| ```                     A    GGTGGT mRNA  3' ATTCCATCGTT AACG       miRNA 5' TGAGGTAGTAG TTGT                           G    ATAGTT           1211111121231112333333 ``` \| SeedType = 3 NonSeedType = 1 \| \| --- \| --- \| --- \| --- \| --- \| --- \| | \|  \|  \|  \| [seedLoc = 178,183] Seq=GAATTTTTTTGTTGCTATCTCT \| ```          T          GTTTTTTTAAG mRNA  3'  CTCTATCGTT            miRNA 5'  GAGGTAGTAG                     T          GTTGTATAGTT           3111211121233333333333 ``` \| SeedType = 2 NonSeedType = -1 \| \| --- \| --- \| --- \| --- \| --- \| --- \| | \| 980 \| WBGene00018660:F52C6.3(11379) \| ``` log2FC=0.067 padj=0.982 ``` \| [seedLoc = 140,146] Seq=ACATCTATAATGAGGCTACTCTCC \| ```          C   T     GAG      CTACA mRNA  3'  CTC CATCG   TAATAT      miRNA 5'  GAG GTAGT   GTTGTA               T         AG       TAGTT           311151111233421121133333 ``` \| SeedType = 2 NonSeedType = -1 \| \| --- \| --- \| --- \| --- \| --- \| --- \| | \| 981 \| WBGene00018661:F52C6.4(11380) \| ``` log2FC=0.015 padj=0.998 ``` \| [seedLoc = 110,116] Seq=TCAAAAACGGCAGTACTCTCC \| ```          C   T   GA       AAAACT mRNA  3'  CTC CAT    CGGCA       miRNA 5'  GAG GTA    GTTGT                T       GTAG     ATAGTT           31115111336612211333333 ``` \| SeedType = 2 NonSeedType = -1 \| \| --- \| --- \| --- \| --- \| --- \| --- \| | \| 982 \| WBGene00018667:bath-7(11382) \| ``` log2FC=-0.058 padj=0.989 ``` \| [seedLoc = 29,35] Seq=ATGTATATAATTTTTTAACCTCG \| ```          G    A   T          GTA mRNA  3'  CTCC ATT TTTAATATAT    miRNA 5'  GAGG TAG AGGTTGTATA             T        T          GTT           31111511231221121111333 ``` \| SeedType = 2 NonSeedType = 2 \| \| --- \| --- \| --- \| --- \| --- \| --- \| | \| 983 \| WBGene00018689:F52D2.7(11398) \| ``` log2FC=0.348 padj=0.679 ``` \| [seedLoc = 219,224] Seq=CACCTTTTCCCCTCTACTTCC \| ```          C           CCTTTTCCAC mRNA  3'  CTTCATC TCC           miRNA 5'  GAGGTAG AGG                    T       T   TTGTATAGTT           3112111161113333333333 ``` \| SeedType = 3 NonSeedType = -1 \| \| --- \| --- \| --- \| --- \| --- \| --- \| | \| 984 \| WBGene00018725:kreg-1(11424) \|  \| [seedLoc = 82,88] Seq=ATATGTATTGCGCTACCTGC \| ```          CG           T     ATA mRNA  3'   TCCATCG  CG TATGT    miRNA 5'   AGGTAGT  GT GTATA             TG       AG  T     GTT           3311111126612321121333 ``` \| SeedType = 3 NonSeedType = -1 \| \| --- \| --- \| --- \| --- \| --- \| --- \| | \| 985 \| WBGene00018738:swip-10(11435) \|  \| [seedLoc = 86,91] Seq=TCTTATGTCAAGTTACTACTTCA \| ```                     G   T     CT mRNA  3' ACTTCATCATT AAC GTATT   miRNA 5' TGAGGTAGTAG TTG TATAG                       G         TT           11121111112311152111233 ``` \| SeedType = 3 NonSeedType = 1 \| \| --- \| --- \| --- \| --- \| --- \| --- \| | \| 986 \| WBGene00018747:F53C3.3(11443) \| ``` log2FC=2.552 padj=1.0 ``` \| [seedLoc = 44,49] Seq=TTTCTTTCAACCTTCTTTCTACCTCT \| ```          T       TTTCT      TTTCTTT mRNA  3'  CTCCATC     TCCAAC        miRNA 5'  GAGGTAG     AGGTTG                 T       T          TATAGTT           31111111355551111113333333 ``` \| SeedType = 1 NonSeedType = 1 \| \| --- \| --- \| --- \| --- \| --- \| --- \| | \|  \|  \|  \| [seedLoc = 81,86] Seq=ATTTTTCAAATCACTGCCTCG \| ```          G             ACTTTTTA mRNA  3'  CTCCGTCA CTAA         miRNA 5'  GAGGTAGT GGTT                  T        A    GTATAGTT           3111121116121133333333 ``` \| SeedType = 2 NonSeedType = 2 \| \| --- \| --- \| --- \| --- \| --- \| --- \| | \| 987 \| WBGene00018756:plpp-1.1(11449) \|  \| [seedLoc = 297,302] Seq=AAGCTTTATTCTACGATACCTCT \| ```          T      AG          T    A mRNA  3'  CTCCAT  CATCT  TAT TCGA  miRNA 5'  GAGGTA  GTAGG  GTA AGTT           T             TT   T                3111111541111266211311215 ``` \| SeedType = 1 NonSeedType = -1 \| \| --- \| --- \| --- \| --- \| --- \| --- \| | \| 988 \| WBGene00018764:F53F10.2(11457) \| ``` log2FC=-0.202 padj=0.571 ``` \| [seedLoc = 335,341] Seq=TGAGACACCGCCTTCCTTCGTCTACCTAC \| ```          CA      TGCTTCCT    C  CAGAGT mRNA  3'   TCCATC        TCCG CA       miRNA 5'   AGGTAG        AGGT GT                TG      T           T  ATAGTT           33111111345555551112311333333 ``` \| SeedType = 3 NonSeedType = 2 \| \| --- \| --- \| --- \| --- \| --- \| --- \| | \|  \|  \|  \| [seedLoc = 342,348] Seq=CCTTCGTCTACCTACTACACTCC \| ```          C   A         TCTGCTTCC mRNA  3'  CTC CATCATCCA          miRNA 5'  GAG GTAGTAGGT                   T             TGTATAGTT           31115111111111333333333 ``` \| SeedType = 2 NonSeedType = 1 \| \| --- \| --- \| --- \| --- \| --- \| --- \| | \|  \|  \|  \| [seedLoc = 930,935] Seq=CTTGGTGCAAATGTTACCGGAATCTATCTCG \| ```          G       TAAGG   TTGTAA     GTTC mRNA  3'  CTCTATC     CCA      ACGTG     miRNA 5'  GAGGTAG     GGT      TGTAT              T       TA                 AGTT           3111211133544111554555112123333 ``` \| SeedType = 2 NonSeedType = 1 \| \| --- \| --- \| --- \| --- \| --- \| --- \| | \|  \|  \|  \| [seedLoc = 944,948] Seq=TTACCGGAATCTATCTCGA \| ```          A        TAAGG   TT        mRNA  3'  GCTCTATC     CCA          miRNA 5'  TGAGGTAG     GGT                            TA      TGTATAGTT           52111211133544111336666666 ``` \| SeedType = 2 NonSeedType = 1 \| \| --- \| --- \| --- \| --- \| --- \| --- \| | \| 989 \| WBGene00018782:cct-3(11469) \| ``` log2FC=0.138 padj=0.81 ``` \| [seedLoc = 353,358] Seq=AAAAATGCGATTTTTACCTCG \| ```          G                 AAAA mRNA  3'  CTCCATT TTTAGCGTA     miRNA 5'  GAGGTAG AGGTTGTAT              T       T         AGTT           3111111261221212113333 ``` \| SeedType = 1 NonSeedType = 2 \| \| --- \| --- \| --- \| --- \| --- \| --- \| | \| 990 \| WBGene00018785:F54A3.6(11471) \| ``` log2FC=-0.134 padj=0.916 ``` \| [seedLoc = 690,696] Seq=CAACTGGACAGAGTCGTCTATCCTCC \| ```          C    T   TGCTGA    G     C mRNA  3'  CTCC ATC      GACA GTCAA  miRNA 5'  GAGG TAG      TTGT TAGTT           T        TAGG      A                 31111511133334521113211115 ``` \| SeedType = 2 NonSeedType = -1 \| \| --- \| --- \| --- \| --- \| --- \| --- \| | \| 991 \| WBGene00018788:shc-1(11473) \| ``` log2FC=0.015 padj=0.995 ``` \| [seedLoc = 151,157] Seq=ACCGTAACATGTCATTACTATCCTCT \| ```          T    T      ACTGT   ATGCCA mRNA  3'  CTCC ATCATT     ACA       miRNA 5'  GAGG TAGTAG     TGT                T           GT      ATAGTT           31111511111233545111333333 ``` \| SeedType = 2 NonSeedType = -1 \| \| --- \| --- \| --- \| --- \| --- \| --- \| | \| 992 \| WBGene00018819:rog-1(11493) \| ``` log2FC=0.217 padj=0.766 ``` \| [seedLoc = 240,245] Seq=ACCTATTGCGTACCAGTCATACCTCT \| ```          T      ACTGA   T   T    CA mRNA  3'  CTCCAT     CCA GCG TATC   miRNA 5'  GAGGTA     GGT TGT ATAG            T      GTA              TT           31111113334511152125111133 ``` \| SeedType = 1 NonSeedType = 1 \| \| --- \| --- \| --- \| --- \| --- \| --- \| | \| 993 \| WBGene00018845:F54H12.5(11508) \| ``` log2FC=-0.611 padj=0.86 ``` \| [seedLoc = 51,56] Seq=AATTATAACTGTTCTTGAAATATCTATCTCA \| ```                  TATAAAGTTCTTG       TAA   mRNA  3' ACTCTATC             TCAATAT      miRNA 5' TGAGGTAG             GGTTGTA                       TA                  TAGTT           111121113355554555554211121133366 ``` \| SeedType = 2 NonSeedType = 1 \| \| --- \| --- \| --- \| --- \| --- \| --- \| | \| 994 \| WBGene00018857:stau-1(11516) \| ``` log2FC=0.066 padj=0.973 ``` \| [seedLoc = 108,113] Seq=GCAGAATTAAACTACTGCCTCC \| ```          C          AAATTAAGACG mRNA  3'  CTCCGTCATC            miRNA 5'  GAGGTAGTAG                     T          GTTGTATAGTT           3111121111133333333333 ``` \| SeedType = 2 NonSeedType = -1 \| \| --- \| --- \| --- \| --- \| --- \| --- \| | \| 995 \| WBGene00018864:F55A12.5(11523) \| ``` log2FC=0.12 padj=0.872 ``` \| [seedLoc = 30,35] Seq=ATCTATTATATTTAAATACCTCC \| ```          C      AA     TAT    TA mRNA  3'  CTCCAT  ATTTA   TATC   miRNA 5'  GAGGTA  TAGGT   ATAG            T      G      TGT    TT           31111113511221333111133 ``` \| SeedType = 1 NonSeedType = 2 \| \| --- \| --- \| --- \| --- \| --- \| --- \| | \| 996 \| WBGene00018872:F55C12.5(11530) \| ``` log2FC=0.024 padj=0.988 ``` \| [seedLoc = 141,146] Seq=TTGCGACGATTCTTTCGATTCAAGTACCTCC \| ```          C      GAACTTAGCTTTCTTAG      T     mRNA  3'  CTCCAT                 CAGCGT      miRNA 5'  GAGGTA                 GTTGTA               T      GTAG                   TAGTT           31111113333555455555555411212136666 ``` \| SeedType = 1 NonSeedType = -1 \| \| --- \| --- \| --- \| --- \| --- \| --- \| | \| 997 \| WBGene00018900:rod-1(11551) \| ``` log2FC=0.103 padj=0.912 ``` \| [seedLoc = 95,101] Seq=TTTAATTTAATTTTTTACGCTCT \| ```          T   G    T      TTAATTT mRNA  3'  CTC CATT TTTAAT        miRNA 5'  GAG GTAG AGGTTG                 T        T      TATAGTT           31114111231221123333333 ``` \| SeedType = 2 NonSeedType = 2 \| \| --- \| --- \| --- \| --- \| --- \| --- \| | \| 998 \| WBGene00018926:F56A11.6(11564) \| ``` log2FC=-0.09 padj=0.934 ``` \| [seedLoc = 33,39] Seq=CGCTTATCGGCATTTTATTTTTACTCTCT \| ```          T   T   TTT     T   GCTAT   C mRNA  3'  CTC CAT   TATTT ACG     TCG  miRNA 5'  GAG GTA   GTAGG TGT     AGT           T               T   AT      T           31115111555211223112335551123 ``` \| SeedType = 2 NonSeedType = -1 \| \| --- \| --- \| --- \| --- \| --- \| --- \| | \| 999 \| WBGene00018967:ddx-15(11593) \| ``` log2FC=0.201 padj=0.596 ``` \| [seedLoc = 28,31] Seq=TGCGTTTCAGGCTACTTCAA \| ```          A         GA  TT          mRNA  3'  ACTTCATCG  CT  GCGT      miRNA 5'  TGAGGTAGT  GG  TGTA                         A   T     TAGTT           5111211112351235212166666 ``` \| SeedType = 3 NonSeedType = -1 \| \| --- \| --- \| --- \| --- \| --- \| --- \| | \| 1000 \| WBGene00019018:abts-3(11628) \| ``` log2FC=0.047 padj=0.982 ``` \| [seedLoc = 36,41] Seq=AAAATTTGTTCCTGGATCCTACTTCT \| ```          T       CTAG    TTGTTTAAAA mRNA  3'  CTTCATC    GTCC           miRNA 5'  GAGGTAG    TAGG                    T               TTGTATAGTT           31121111555421113333333333 ``` \| SeedType = 3 NonSeedType = -1 \| \| --- \| --- \| --- \| --- \| --- \| --- \| | \| 1001 \| WBGene00019079:rpa-4(11660) \| ``` log2FC=0.118 padj=0.979 ``` \| [seedLoc = 225,230] Seq=ATGATTCTTATATACTACCTTC \| ```          C         ATATTCTTAGTA mRNA  3'  TTCCATCAT             miRNA 5'  GAGGTAGTA                      T         GGTTGTATAGTT           3211111111333333333333 ``` \| SeedType = 3 NonSeedType = -1 \| \| --- \| --- \| --- \| --- \| --- \| --- \| | \| 1002 \| WBGene00019103:F59D6.1(11675) \| ``` log2FC=0.253 padj=1.0 ``` \| [seedLoc = 58,63] Seq=TCAAATTCAATTACTCTACCTCC \| ```          C       TCA     TTAAACT mRNA  3'  CTCCATC   TTAAC        miRNA 5'  GAGGTAG   GGTTG                 T       TA      TATAGTT           31111111335221113333333 ``` \| SeedType = 1 NonSeedType = 1 \| \| --- \| --- \| --- \| --- \| --- \| --- \| | \|  \|  \|  \| [seedLoc = 75,80] Seq=CTCCTTGTAACCCTTACCTCT \| ```          T       C      GTTCCTC mRNA  3'  CTCCATT  CCAAT        miRNA 5'  GAGGTAG  GGTTG                 T       TA     TATAGTT           3111111236111123333333 ``` \| SeedType = 1 NonSeedType = 1 \| \| --- \| --- \| --- \| --- \| --- \| --- \| | \| 1003 \| WBGene00019127:cgt-3(11689) \| ``` log2FC=0.236 padj=0.52 ``` \| [seedLoc = 205,211] Seq=CGGGTGACATCTGAAATACCATCT \| ```          T  A    AAAG  T   GTGGGC mRNA  3'  CT CCAT    TC ACA       miRNA 5'  GA GGTA    GG TGT                T       GTA   T   ATAGTT           311511113334213111333333 ``` \| SeedType = 3 NonSeedType = -1 \| \| --- \| --- \| --- \| --- \| --- \| --- \| | \| 1004 \| WBGene00019143:H01M10.1(11698) \| ``` log2FC=-0.787 padj=0.678 ``` \| [seedLoc = 152,157] Seq=GTTTAATTGACTTTGAACTACCTTC \| ```          C        AGT     TTAATTTG mRNA  3'  TTCCATCA   TTCAG         miRNA 5'  GAGGTAGT   AGGTT                  T                GTATAGTT           3211111115451211233333333 ``` \| SeedType = 3 NonSeedType = 2 \| \| --- \| --- \| --- \| --- \| --- \| --- \| | \|  \|  \|  \| [seedLoc = 204,210] Seq=CATTGTTTTAATTCATACCATCG \| ```          G  A    AC      TT     C mRNA  3'  CT CCAT   TTAAT  TGTTA  miRNA 5'  GA GGTA   GGTTG  ATAGT           T       GTA     T      T           311511113362211235121213 ``` \| SeedType = 3 NonSeedType = 2 \| \| --- \| --- \| --- \| --- \| --- \| --- \| | \| 1005 \| WBGene00019153:H04M03.3(11703) \| ``` log2FC=0.022 padj=0.994 ``` \| [seedLoc = 60,65] Seq=ACGAGTATTCCTTTTTCTACCTCA \| ```                  TTTT   T     AGCA mRNA  3' ACTCCATC    TCC  TATG     miRNA 5' TGAGGTAG    AGG  GTAT                      T      TT    AGTT           1111111135551113621123333 ``` \| SeedType = 1 NonSeedType = -1 \| \| --- \| --- \| --- \| --- \| --- \| --- \| | \| 1006 \| WBGene00019189:H11L12.1(11727) \| ``` log2FC=-0.081 padj=0.955 ``` \| [seedLoc = 144,150] Seq=GCCCTCAAAACCTGTTGGATTGTACACTCA \| ```              A   GTTAGGTT      AACTCCCG mRNA  3' ACTC CAT        GTCCAA         miRNA 5' TGAG GTA        TAGGTT                          G             GTATAGTT           111151113555445521111133333333 ``` \| SeedType = 2 NonSeedType = 1 \| \| --- \| --- \| --- \| --- \| --- \| --- \| | \| 1007 \| WBGene00019204:H14N18.4(11734) \| ``` log2FC=-0.223 padj=0.633 ``` \| [seedLoc = 64,70] Seq=TAATTATATTCATTACGCTCC \| ```          C   G                  T mRNA  3'  CTC CATTA CT  TATATTAA  miRNA 5'  GAG GTAGT GG  GTATAGTT           T         A  TT                    311141112161266211112115 ``` \| SeedType = 2 NonSeedType = -1 \| \| --- \| --- \| --- \| --- \| --- \| --- \| | \|  \|  \|  \| [seedLoc = 223,229] Seq=AATTCGGTGACCGATCTACACTCC \| ```          C   A    TAG    TGGCTTAA mRNA  3'  CTC CATC   CCAG         miRNA 5'  GAG GTAG   GGTT                  T        TA     GTATAGTT           311151111334111233333333 ``` \| SeedType = 2 NonSeedType = 1 \| \| --- \| --- \| --- \| --- \| --- \| --- \| | \| 1008 \| WBGene00019204:H14N18.4(11735) \| ``` log2FC=-0.223 padj=0.633 ``` \| [seedLoc = 64,70] Seq=TAATTATATTCATTACGCTCC \| ```          C   G                  T mRNA  3'  CTC CATTA CT  TATATTAA  miRNA 5'  GAG GTAGT GG  GTATAGTT           T         A  TT                    311141112161266211112115 ``` \| SeedType = 2 NonSeedType = -1 \| \| --- \| --- \| --- \| --- \| --- \| --- \| | \|  \|  \|  \| [seedLoc = 223,229] Seq=AATTCGGTGACCGATCTACACTCA \| ```              A    TAG    TGGCTTAA mRNA  3' ACTC CATC   CCAG         miRNA 5' TGAG GTAG   GGTT                           TA     GTATAGTT           111151111334111233333333 ``` \| SeedType = 2 NonSeedType = 1 \| \| --- \| --- \| --- \| --- \| --- \| --- \| | \| 1009 \| WBGene00019211:H18N23.2(11739) \| ``` log2FC=0.436 padj=0.073 ``` \| [seedLoc = 157,163] Seq=ATCAGTTCATTTTATTACTCTCT \| ```          T   T        T  TTGACTA mRNA  3'  CTC CATTATTT AC        miRNA 5'  GAG GTAGTAGG TG                 T            T  TATAGTT           31115111211223113333333 ``` \| SeedType = 2 NonSeedType = -1 \| \| --- \| --- \| --- \| --- \| --- \| --- \| | \| 1010 \| WBGene00019259:H34C03.2(11771) \| ``` log2FC=0.213 padj=0.466 ``` \| [seedLoc = 435,440] Seq=ATTTGTGGTATCTTATTTTTTCTATCTCT \| ```          T       TTTTTT    TATG     TA mRNA  3'  CTCTATC      ATTC    GTGTT   miRNA 5'  GAGGTAG      TAGG    TATAG            T                 TTG      TT           31112111555555112133342121233 ``` \| SeedType = 2 NonSeedType = -1 \| \| --- \| --- \| --- \| --- \| --- \| --- \| | \| 1011 \| WBGene00019265:H35N09.1(11777) \| ``` log2FC=2.247 padj=1.0 ``` \| [seedLoc = 69,74] Seq=TGATTTCACCTCCTGTCCCATATACCTCC \| ```          C      ATACC T    TC  CT    T mRNA  3'  CTCCAT     C GTCC  CA  TTAG  miRNA 5'  GAGGTA     G TAGG  GT  AGTT           T                 TT  AT                31111115555515211133113312125 ``` \| SeedType = 1 NonSeedType = -1 \| \| --- \| --- \| --- \| --- \| --- \| --- \| | \| 1012 \| WBGene00019274:H43I07.1(11784) \| ``` log2FC=0.111 padj=0.932 ``` \| [seedLoc = 57,63] Seq=ATATGTACAGTAATCCCTTTTCACCTCC \| ```          C     C  T   CTAAT       ATA mRNA  3'  CTCCA TT TCC     GACATGT    miRNA 5'  GAGGT AG AGG     TTGTATA             T        T               GTT           3111115123111555552111121333 ``` \| SeedType = 2 NonSeedType = -1 \| \| --- \| --- \| --- \| --- \| --- \| --- \| | \| 1013 \| WBGene00019279:K01A2.4(11789) \| ``` log2FC=0.423 padj=1.0 ``` \| [seedLoc = 921,926] Seq=GGGAATACAATTCACTCTTCTACCTCC \| ```          C       T  TCACT       AGGG mRNA  3'  CTCCATC TC     TAACATA     miRNA 5'  GAGGTAG AG     GTTGTAT              T       T              AGTT           311111113115555521111113333 ``` \| SeedType = 1 NonSeedType = 1 \| \| --- \| --- \| --- \| --- \| --- \| --- \| | \|  \|  \|  \| [seedLoc = 1478,1484] Seq=TATTGTGGAGTACTACTCTCA \| ```              T          G      T mRNA  3' ACTC CATCAT  GA GTGTTA  miRNA 5' TGAG GTAGTA  TT TATAGT                      GG  G      T           11115111111662132121213 ``` \| SeedType = 2 NonSeedType = -1 \| \| --- \| --- \| --- \| --- \| --- \| --- \| | \|  \|  \|  \| [seedLoc = 1512,1517] Seq=GAGAGTGCACGAAAAAAAGAGATACCTCC \| ```          C      AGAGAAAAAAAG       AGAG mRNA  3'  CTCCAT            C ACGTG     miRNA 5'  GAGGTA            G TGTAT              T      GTAG         T     AGTT           311111133335555555416112123333 ``` \| SeedType = 1 NonSeedType = -1 \| \| --- \| --- \| --- \| --- \| --- \| --- \| | \| 1014 \| WBGene00019284:K01A2.10(11792) \| ``` log2FC=-0.7 padj=0.075 ``` \| [seedLoc = 74,79] Seq=TATTGTACATATATACCTCA \| ```                 ATAT          T mRNA  3' ACTCCAT      ACATGTTA  miRNA 5' TGAGGTA      TGTATAGT                  GTAGGT        T           1111111333366111121213 ``` \| SeedType = 1 NonSeedType = -1 \| \| --- \| --- \| --- \| --- \| --- \| --- \| | \| 1015 \| WBGene00019301:K02D10.1(11804) \| ``` log2FC=0.09 padj=0.902 ``` \| [seedLoc = 36,41] Seq=TTTTTTAGGAATCTATCTCT \| ```          T       TA    G  TTTTT mRNA  3'  CTCTATC    AG AT      miRNA 5'  GAGGTAG    TT TA               T       TAGG  G  TAGTT           3111211133661231133333 ``` \| SeedType = 2 NonSeedType = -1 \| \| --- \| --- \| --- \| --- \| --- \| --- \| | \| 1016 \| WBGene00019318:K02E10.4(11813) \| ``` log2FC=0.438 padj=0.704 ``` \| [seedLoc = 68,73] Seq=TTTTATTTAAATAAATACCTCA \| ```                 AAATA   T    TT mRNA  3' ACTCCAT     AAT TATT   miRNA 5' TGAGGTA     TTG ATAG                   GTAGG   T    TT           1111111333331123111233 ``` \| SeedType = 1 NonSeedType = -1 \| \| --- \| --- \| --- \| --- \| --- \| --- \| | \| 1017 \| WBGene00019399:K04F10.7(11856) \| ``` log2FC=-0.223 padj=0.701 ``` \| [seedLoc = 295,300] Seq=CTAATTACAATCTTTGGACGTAGCTACTTCT \| ```          T        ATGCAGGTT        TAATC mRNA  3'  CTTCATCG         TCTAACAT      miRNA 5'  GAGGTAGT         AGGTTGTA               T                         TAGTT           3112111125545544551121111133333 ``` \| SeedType = 3 NonSeedType = 1 \| \| --- \| --- \| --- \| --- \| --- \| --- \| | \| 1018 \| WBGene00019464:K07B1.4(11893) \| ``` log2FC=0.554 padj=0.109 ``` \| [seedLoc = 55,60] Seq=ATATTTACATCTCTATTATACCTCA \| ```                 AT    T T    TTATA mRNA  3' ACTCCAT  TATC C ACAT      miRNA 5' TGAGGTA  GTAG G TGTA                              T    TAGTT           1111111552111513111133333 ``` \| SeedType = 1 NonSeedType = -1 \| \| --- \| --- \| --- \| --- \| --- \| --- \| | \| 1019 \| WBGene00019479:K07C11.7(11902) \| ``` log2FC=-0.199 padj=0.695 ``` \| [seedLoc = 233,239] Seq=CAAAGTAGAATTTCAAATACTCTCT \| ```          T   T   AAAC     G   AAAC mRNA  3'  CTC CAT    TTTAA ATG     miRNA 5'  GAG GTA    AGGTT TAT              T       GT       G   AGTT           3111511133551221131123333 ``` \| SeedType = 2 NonSeedType = 2 \| \| --- \| --- \| --- \| --- \| --- \| --- \| | \| 1020 \| WBGene00019487:ephx-1(11907) \| ``` log2FC=-0.527 padj=0.034 ``` \| [seedLoc = 253,258] Seq=TATCATAGAGATTTATGGATACCTCC \| ```          C      AGG       AG   CTAT mRNA  3'  CTCCAT   TATTTAG  ATA     miRNA 5'  GAGGTA   GTAGGTT  TAT              T                G    AGTT           31111115442112212341113333 ``` \| SeedType = 1 NonSeedType = 2 \| \| --- \| --- \| --- \| --- \| --- \| --- \| | \| 1021 \| WBGene00019489:K07E1.1(11909) \| ``` log2FC=-0.056 padj=0.966 ``` \| [seedLoc = 195,200] Seq=GAGCTCTCTTCATCTCTATCTCG \| ```          G            CTTCTC    G mRNA  3'  CTCTATC TCTA      TCGA  miRNA 5'  GAGGTAG AGGT      AGTT           T       T    TGTAT                 311121116112133333511214 ``` \| SeedType = 2 NonSeedType = 2 \| \| --- \| --- \| --- \| --- \| --- \| --- \| | \| 1022 \| WBGene00019495:sdz-24(11914) \| ``` log2FC=0.057 padj=0.98 ``` \| [seedLoc = 79,85] Seq=TGCTTCTCAACCCCGCCGACTACCCTCC \| ```          C  C      GCCGCC     TCT   T mRNA  3'  CT CCATCA      CCAAC   TCG  miRNA 5'  GA GGTAGT      GGTTG   AGT           T         A          TAT   T           3115111111355455111113331123 ``` \| SeedType = 3 NonSeedType = 1 \| \| --- \| --- \| --- \| --- \| --- \| --- \| | \| 1023 \| WBGene00019510:K07H8.10(11923) \| ``` log2FC=-0.2 padj=0.747 ``` \| [seedLoc = 185,190] Seq=CATTGTTTTCCCCTGCCTCT \| ```          T       C   TTT      C mRNA  3'  CTCCGTC  CC    TGTTA  miRNA 5'  GAGGTAG  GG    ATAGT           T       TA  TTGT     T           3111121136113336121213 ``` \| SeedType = 2 NonSeedType = -1 \| \| --- \| --- \| --- \| --- \| --- \| --- \| | \| 1024 \| WBGene00019518:K08B5.1(11929) \| ``` log2FC=-0.255 padj=0.918 ``` \| [seedLoc = 58,63] Seq=GGTTGAACATACAGTACCTCG \| ```          G      G   AT   AGTTGG mRNA  3'  CTCCAT A C  ACA       miRNA 5'  GAGGTA T G  TGT                T      G A GT   ATAGTT           3111111316133111333333 ``` \| SeedType = 1 NonSeedType = -1 \| \| --- \| --- \| --- \| --- \| --- \| --- \| | \| 1025 \| WBGene00019537:K08D12.3(11941) \| ``` log2FC=0.033 padj=0.98 ``` \| [seedLoc = 141,147] Seq=CAACTGCGGAGGACGTGGACACTACTCTCG \| ```          G   T     CAGG G   G GGC     C mRNA  3'  CTC CATCA    T CAG A   GTCAA  miRNA 5'  GAG GTAGT    A GTT T   TAGTT           T              G   G A                   311151111155441311231345211115 ``` \| SeedType = 2 NonSeedType = -1 \| \| --- \| --- \| --- \| --- \| --- \| --- \| | \| 1026 \| WBGene00019543:cif-1(11945) \| ``` log2FC=0.123 padj=0.806 ``` \| [seedLoc = 108,113] Seq=TCCTCTATAATTTCTATCTCT \| ```          T                C  CT mRNA  3'  CTCTATC TTTAATAT TC   miRNA 5'  GAGGTAG AGGTTGTA AG            T       T        T  TT           3111211161221121131133 ``` \| SeedType = 2 NonSeedType = 2 \| \| --- \| --- \| --- \| --- \| --- \| --- \| | \| 1027 \| WBGene00019606:clec-88(11977) \| ``` log2FC=0.006 padj=0.997 ``` \| [seedLoc = 26,32] Seq=TGTAAATAATCTTACATTACAATTACCATCT \| ```          T  A      ACATTACAT       AATGT  mRNA  3'  CT CCATTA         TCTAATA       miRNA 5'  GA GGTAGT         AGGTTGT                T                         ATAGTT           31151111215555555551121121333336 ``` \| SeedType = 3 NonSeedType = 2 \| \| --- \| --- \| --- \| --- \| --- \| --- \| | \| 1028 \| WBGene00019622:K10C2.7(11990) \| ``` log2FC=0.577 padj=0.786 ``` \| [seedLoc = 36,41] Seq=AAAAATTCATGAAATTACCTCT \| ```          T        AAGTACTTAAAAA mRNA  3'  CTCCATTA              miRNA 5'  GAGGTAGT                       T        AGGTTGTATAGTT           3111111213333333333333 ``` \| SeedType = 1 NonSeedType = -1 \| \| --- \| --- \| --- \| --- \| --- \| --- \| | \| 1029 \| WBGene00019628:pup-2(11994) \| ``` log2FC=0.273 padj=0.509 ``` \| [seedLoc = 583,589] Seq=TAAGTCTGAGCTTAATTACCCTCC \| ```          C  C     A      GTCTGAAT mRNA  3'  CT CCATT ATTCGA         miRNA 5'  GA GGTAG TAGGTT                  T               GTATAGTT           311511112511212133333333 ``` \| SeedType = 3 NonSeedType = 2 \| \| --- \| --- \| --- \| --- \| --- \| --- \| | \| 1030 \| WBGene00019697:M01B12.4(12042) \| ``` log2FC=-0.152 padj=0.766 ``` \| [seedLoc = 201,207] Seq=TCCTGTACACTATTCTGTACTCTCT \| ```          T   T        TATC       CT mRNA  3'  CTC CAT GTCT    ACATGTC   miRNA 5'  GAG GTA TAGG    TGTATAG            T       G    T          TT           31115111621123555111121133 ``` \| SeedType = 2 NonSeedType = -1 \| \| --- \| --- \| --- \| --- \| --- \| --- \| | \| 1031 \| WBGene00019698:riok-1(12043) \| ``` log2FC=0.036 padj=0.982 ``` \| [seedLoc = 356,361] Seq=GAATATACCAGTTTTCTGCCTCC \| ```          C       TT  G C     AAG mRNA  3'  CTCCGTC  TT A CATAT    miRNA 5'  GAGGTAG  AG T GTATA             T       T   G T     GTT           31111211351231311111333 ``` \| SeedType = 2 NonSeedType = -1 \| \| --- \| --- \| --- \| --- \| --- \| --- \| | \| 1032 \| WBGene00019713:pqn-52(12052) \| ``` log2FC=-0.195 padj=0.717 ``` \| [seedLoc = 79,84] Seq=TCATGTACAATCGTTCATGTACTGCCTCT \| ```          T         GTACTTG         ACT mRNA  3'  CTCCGTCAT       CTAACATGT    miRNA 5'  GAGGTAGTA       GGTTGTATA             T                         GTT           31111211114555554121111121333 ``` \| SeedType = 2 NonSeedType = 1 \| \| --- \| --- \| --- \| --- \| --- \| --- \| | \| 1033 \| WBGene00019715:M01G5.1(12053) \| ``` log2FC=-0.042 padj=0.977 ``` \| [seedLoc = 62,68] Seq=TTCTAATGGCCTTATTTATCCTCG \| ```          G    T   TAT           TT mRNA  3'  CTCC ATT   TCCGGTA ATC   miRNA 5'  GAGG TAG   AGGTTGT TAG            T        T         A   TT           3111151123551112221611133 ``` \| SeedType = 2 NonSeedType = 2 \| \| --- \| --- \| --- \| --- \| --- \| --- \| | \| 1034 \| WBGene00019732:M02D8.6(12069) \| ``` log2FC=-0.406 padj=0.662 ``` \| [seedLoc = 55,61] Seq=TTATTAACAAACACCTACCGTCA \| ```             G     CA AA   ATTATT mRNA  3' ACT CCATC  C  ACA       miRNA 5' TGA GGTAG  G  TGT                         TA GT   ATAGTT           11141111133133111333333 ``` \| SeedType = 3 NonSeedType = -1 \| \| --- \| --- \| --- \| --- \| --- \| --- \| | \| 1035 \| WBGene00019771:M04F3.5(12098) \| ``` log2FC=-0.026 padj=0.989 ``` \| [seedLoc = 96,101] Seq=AAACGGGCATTTTCTATCTCT \| ```          T           T   GG    A mRNA  3'  CTCTATC TTT ACG   CAA  miRNA 5'  GAGGTAG AGG TGT   GTT           T       T   T   ATA               31112111612231123361115 ``` \| SeedType = 2 NonSeedType = -1 \| \| --- \| --- \| --- \| --- \| --- \| --- \| | \| 1036 \| WBGene00019778:M57.2(12104) \| ``` log2FC=0.067 padj=0.951 ``` \| [seedLoc = 194,199] Seq=CTTCTTACACTTCTTGTAATAAACTACTTCT \| ```          T        AATAATGTTC        TCTTC mRNA  3'  CTTCATCA          TTC ACAT      miRNA 5'  GAGGTAGT          AGG TGTA               T                     T    TAGTT           31121111155555545551216111133333 ``` \| SeedType = 3 NonSeedType = -1 \| \| --- \| --- \| --- \| --- \| --- \| --- \| | \| 1037 \| WBGene00019782:M60.7(12108) \| ``` log2FC=-0.845 padj=0.126 ``` \| [seedLoc = 159,164] Seq=AAGTGTACAAAACCAGCACATTGTTACCTCA \| ```                    TACACGA   AA      G   mRNA  3' ACTCCATTGT       CCA  ACATGT AA miRNA 5' TGAGGTAGTA       GGT  TGTATA TT                                      G             1111111221555554511155111121311 ``` \| SeedType = 1 NonSeedType = 1 \| \| --- \| --- \| --- \| --- \| --- \| --- \| | \| 1038 \| WBGene00019808:R01B10.6(12127) \| ``` log2FC=-0.353 padj=0.603 ``` \| [seedLoc = 67,72] Seq=CGGTATACCTACCTCG \| ```          G                  G C mRNA  3'  CTCCATC      CATAT G  miRNA 5'  GAGGTAG      GTATA T           T       TAGGTT     G T           3111111166666611111323 ``` \| SeedType = 1 NonSeedType = -1 \| \| --- \| --- \| --- \| --- \| --- \| --- \| | \| 1039 \| WBGene00019831:R02F2.1(12145) \| ``` log2FC=0.066 padj=0.966 ``` \| [seedLoc = 217,222] Seq=GAGCTGGTTCTCTTCCTTTCTCTATCTCT \| ```          T       TCTT   TT TCTTG     G mRNA  3'  CTCTATC    TCC  C     GTCGA  miRNA 5'  GAGGTAG    AGG  G     TAGTT           T       T      TT TA                    31112111355511133133554211214 ``` \| SeedType = 2 NonSeedType = -1 \| \| --- \| --- \| --- \| --- \| --- \| --- \| | \| 1040 \| WBGene00019844:abts-4(12159) \| ``` log2FC=-0.148 padj=0.842 ``` \| [seedLoc = 158,163] Seq=TTCTCTACAAATTTCGTGTCTACTTCT \| ```          T       T  G TTTA     C  TT mRNA  3'  CTTCATC GT C    AACAT TC   miRNA 5'  GAGGTAG TA G    TTGTA AG            T          G          T  TT           311211115213155551111131133 ``` \| SeedType = 3 NonSeedType = 1 \| \| --- \| --- \| --- \| --- \| --- \| --- \| | \| 1041 \| WBGene00019858:R03H10.6(12166) \| ``` log2FC=-0.059 padj=0.992 ``` \| [seedLoc = 416,421] Seq=AAATTACCAATTTATTTTACCTCA \| ```                 TT        CATTAAA mRNA  3' ACTCCAT  TATTTAAC        miRNA 5' TGAGGTA  GTAGGTTG                                  TATAGTT           111111155211221113333333 ``` \| SeedType = 1 NonSeedType = 1 \| \| --- \| --- \| --- \| --- \| --- \| --- \| | \| 1042 \| WBGene00019865:R04A9.6(12171) \| ``` log2FC=-0.285 padj=0.932 ``` \| [seedLoc = 143,148] Seq=AAATTATATTGTGATTGTTACCTCA \| ```                     AGTGT        A mRNA  3' ACTCCATTGTT     TATATTAA  miRNA 5' TGAGGTAGTAG     GTATAGTT                      GTT                      1111111221233345211112115 ``` \| SeedType = 1 NonSeedType = -1 \| \| --- \| --- \| --- \| --- \| --- \| --- \| | \| 1043 \| WBGene00019871:R04E5.8(12175) \| ``` log2FC=0.011 padj=0.996 ``` \| [seedLoc = 781,786] Seq=TTTTTTGCTACTGCTACTTCT \| ```          T          AT    TTTTT mRNA  3'  CTTCATCGTC   CGT      miRNA 5'  GAGGTAGTAG   GTA               T          GTT   TAGTT           3112111121133612133333 ``` \| SeedType = 3 NonSeedType = -1 \| \| --- \| --- \| --- \| --- \| --- \| --- \| | \| 1044 \| WBGene00019875:mca-2(12178) \| ``` log2FC=0.072 padj=0.959 ``` \| [seedLoc = 208,211] Seq=CAAATTCCTTCCCACTTCCTCC \| ```          C    T   C  TT CTTAAAC mRNA  3'  CTCC TCA CC  C        miRNA 5'  GAGG AGT GG  G                 T    T   A  TT TATAGTT           3111131113113313333333 ``` \| SeedType = 2 NonSeedType = -1 \| \| --- \| --- \| --- \| --- \| --- \| --- \| | \| 1045 \| WBGene00019883:met-2(12183) \| ``` log2FC=0.075 padj=0.945 ``` \| [seedLoc = 147,152] Seq=ATATGTATGATCTAAACACGTATTTACCTCG \| ```          G       TATGCACAA           ATA mRNA  3'  CTCCATT         ATCTAGTATGT    miRNA 5'  GAGGTAG         TAGGTTGTATA             T                           GTT           3111111255545555511121221121333 ``` \| SeedType = 1 NonSeedType = 2 \| \| --- \| --- \| --- \| --- \| --- \| --- \| | \| 1046 \| WBGene00019915:tbc-10(12208) \| ``` log2FC=0.059 padj=0.982 ``` \| [seedLoc = 126,131] Seq=TGCTTTCGCATCTTGCCATATACCTCA \| ```                 ATAC     T   CTT   T mRNA  3' ACTCCAT    CGTTC ACG   TCG  miRNA 5' TGAGGTA    GTAGG TGT   AGT                           T   AT    T           111111155551212131123351123 ``` \| SeedType = 1 NonSeedType = -1 \| \| --- \| --- \| --- \| --- \| --- \| --- \| | \| 1047 \| WBGene00019947:htz-1(12228) \| ``` log2FC=-0.071 padj=0.953 ``` \| [seedLoc = 109,114] Seq=ATTAATTTGATCACTGCCTCA \| ```                        TTTAATTA mRNA  3' ACTCCGTCA CTAG         miRNA 5' TGAGGTAGT GGTT                           A    GTATAGTT           1111121116121233333333 ``` \| SeedType = 2 NonSeedType = 2 \| \| --- \| --- \| --- \| --- \| --- \| --- \| | \| 1048 \| WBGene00019952:fbxb-74(12232) \| ``` log2FC=1.203 padj=1.0 ``` \| [seedLoc = 170,176] Seq=CACTATAATTCTCTTATCTCTACCATCA \| ```             A          TTCTCTTA      C mRNA  3' ACT CCATC TCTA        ATATCA  miRNA 5' TGA GGTAG AGGT        TATAGT                    T    TG            T           11151111161121335555551111113 ``` \| SeedType = 3 NonSeedType = 2 \| \| --- \| --- \| --- \| --- \| --- \| --- \| | \|  \|  \|  \| [seedLoc = 179,184] Seq=AATTCTCTTATCTCTACCATCAA \| ```          A   A          TTCTCTTAA mRNA  3'  ACT CCATC TCTA          miRNA 5'  TGA GGTAG AGGT                             T    TGTATAGTT           511151111161121333333333 ``` \| SeedType = 3 NonSeedType = 2 \| \| --- \| --- \| --- \| --- \| --- \| --- \| | \| 1049 \| WBGene00019960:ztf-16(12240) \| ``` log2FC=-0.105 padj=0.949 ``` \| [seedLoc = 162,167] Seq=TCACAGTCAATTCTGATTACCTCT \| ```          T        G  T    TGACACT mRNA  3'  CTCCATTA TC TAAC        miRNA 5'  GAGGTAGT AG GTTG                 T                TATAGTT           311111121411521113333333 ``` \| SeedType = 1 NonSeedType = 1 \| \| --- \| --- \| --- \| --- \| --- \| --- \| | \| 1050 \| WBGene00020016:lipl-3(12274) \| ``` log2FC=-1.5 padj=3.24610274292036E-5 ``` \| [seedLoc = 49,54] Seq=TCAAGGAATGCGTCTGCCTCC \| ```          C         G  TAAGGAACT mRNA  3'  CTCCGTC T CG          miRNA 5'  GAGGTAG A GT                   T       T G  TGTATAGTT           3111121161312333333333 ``` \| SeedType = 2 NonSeedType = -1 \| \| --- \| --- \| --- \| --- \| --- \| --- \| | \| 1051 \| WBGene00020073:sdz-28(12313) \| ``` log2FC=-1.441 padj=1.0 ``` \| [seedLoc = 40,45] Seq=GGCTCATCGATCACACAAATAAGTTACCTCA \| ```                 TGAATAAACA        TAC   G mRNA  3' ACTCCAT          CA CTAGC   TCG  miRNA 5' TGAGGTA          GT GGTTG   AGT                              A     TAT   T           11111115455555555116121213331123 ``` \| SeedType = 1 NonSeedType = 2 \| \| --- \| --- \| --- \| --- \| --- \| --- \| | \| 1052 \| WBGene00020097:larp-1(12328) \| ``` log2FC=0.029 padj=0.982 ``` \| [seedLoc = 49,55] Seq=TCCTATCACTTTCACCTAAGTTACCATCA \| ```             A     GA     CTTTC       CT mRNA  3' ACT CCATT  ATCCA     AC TATC   miRNA 5' TGA GGTAG  TAGGT     TG ATAG                                   T    TT           111511112451111155555116111133 ``` \| SeedType = 3 NonSeedType = 1 \| \| --- \| --- \| --- \| --- \| --- \| --- \| | \| 1053 \| WBGene00020111:R151.8(12345) \| ``` log2FC=-0.09 padj=0.956 ``` \| [seedLoc = 74,80] Seq=ATTCATTTTCTGTACTACACTCA \| ```              A      G CTTTTACTTA mRNA  3' ACTC CATCAT T           miRNA 5' TGAG GTAGTA G                               G TTGTATAGTT           11115111111323333333333 ``` \| SeedType = 2 NonSeedType = -1 \| \| --- \| --- \| --- \| --- \| --- \| --- \| | \| 1054 \| WBGene00020142:aak-2(12367) \| ``` log2FC=-0.07 padj=0.945 ``` \| [seedLoc = 406,411] Seq=AAACTCCCCTTCTCCTACTTCT \| ```          T       C   TCCCC     A mRNA  3'  CTTCATC TCT      TCAA  miRNA 5'  GAGGTAG AGG      AGTT           T       T   TTGTAT                31121111311233333611115 ``` \| SeedType = 3 NonSeedType = -1 \| \| --- \| --- \| --- \| --- \| --- \| --- \| | \| 1055 \| WBGene00020149:T01D1.4(12371) \| ``` log2FC=-0.274 padj=0.426 ``` \| [seedLoc = 120,126] Seq=TTTTGTACTCCTTTTTCTAAATACCATCC \| ```          C  A    AAATCTTTT   T       TT mRNA  3'  CT CCAT         TCC  CATGTT   miRNA 5'  GA GGTA         AGG  GTATAG            T       GT          TT      TT           311511113355555551113611121233 ``` \| SeedType = 3 NonSeedType = -1 \| \| --- \| --- \| --- \| --- \| --- \| --- \| | \| 1056 \| WBGene00020160:igcm-3(12378) \| ``` log2FC=-0.032 padj=0.99 ``` \| [seedLoc = 703,709] Seq=ATGTGTACATCCACTATCCTCC \| ```          C    T       T      GTA mRNA  3'  CTCC ATCA CC ACATGT    miRNA 5'  GAGG TAGT GG TGTATA             T         A  T      GTT           31111511116113111121333 ``` \| SeedType = 2 NonSeedType = -1 \| \| --- \| --- \| --- \| --- \| --- \| --- \| | \|  \|  \|  \| [seedLoc = 900,905] Seq=CATGCAACTTCGGGTATTATATGCTATCTCT \| ```          T         ATATTATGGGCT        C    mRNA  3'  CTCTATCGT            TCAACGTA     miRNA 5'  GAGGTAGTA            GGTTGTAT              T                             AGTT           3111211121555555544455211112113666 ``` \| SeedType = 2 NonSeedType = 1 \| \| --- \| --- \| --- \| --- \| --- \| --- \| | \| 1057 \| WBGene00020171:T02H6.1(12384) \| ``` log2FC=-0.065 padj=0.959 ``` \| [seedLoc = 45,49] Seq=TCCGCCTACCTACCTACAC \| ```          C  A     C      CCT       mRNA  3'  AC TCCAT CATCCG          miRNA 5'  TG AGGTA GTAGGT                                   TGTATAGTT           5115111115111112333666666 ``` \| SeedType = 3 NonSeedType = 2 \| \| --- \| --- \| --- \| --- \| --- \| --- \| | \| 1058 \| WBGene00020183:T03D3.5(12388) \| ``` log2FC=-1.565 padj=9.06463570422657E-13 ``` \| [seedLoc = 179,184] Seq=TTCTATAATTTTGATGTACTTCTGCCTCT \| ```          T            TGTAGTTTTA     TT mRNA  3'  CTCCGTC TTCA          ATATC   miRNA 5'  GAGGTAG AGGT          TATAG            T       T    TG             TT           311112116121133554555551111133 ``` \| SeedType = 2 NonSeedType = 2 \| \| --- \| --- \| --- \| --- \| --- \| --- \| | \| 1059 \| WBGene00020195:T03G6.3(12399) \| ``` log2FC=0.091 padj=0.961 ``` \| [seedLoc = 140,146] Seq=TTAGTAATAAGAAATACGCTCA \| ```              G   AAAG     ATGATT mRNA  3' ACTC CAT     AATA       miRNA 5' TGAG GTA     TTGT                        GTAGG    ATAGTT           11114111333361121333333 ``` \| SeedType = 2 NonSeedType = -1 \| \| --- \| --- \| --- \| --- \| --- \| --- \| | \| 1060 \| WBGene00020235:T05A12.4(12426) \| ``` log2FC=-0.021 padj=0.994 ``` \| [seedLoc = 108,113] Seq=AAAGTTTCACCCTGTTACCTCG \| ```          G           C  TTTGAAA mRNA  3'  CTCCATTGTCC AC        miRNA 5'  GAGGTAGTAGG TG                 T           T  TATAGTT           3111111221113113333333 ``` \| SeedType = 1 NonSeedType = -1 \| \| --- \| --- \| --- \| --- \| --- \| --- \| | \| 1061 \| WBGene00020261:vet-1(12441) \| ``` log2FC=0.714 padj=1.0 ``` \| [seedLoc = 195,201] Seq=TTCAATCCATCTTAAAATGTTACTCTCT \| ```          T   T    GTAAA    T  CTAACTT mRNA  3'  CTC CATT     ATTC AC        miRNA 5'  GAG GTAG     TAGG TG                 T                 T  TATAGTT           3111511124555511213113333333 ``` \| SeedType = 2 NonSeedType = -1 \| \| --- \| --- \| --- \| --- \| --- \| --- \| | \| 1062 \| WBGene00020263:let-355(12442) \|  \| [seedLoc = 142,147] Seq=TTCTGTTTTTCCCCCTACCTCT \| ```          T       CC  TTTT    TT mRNA  3'  CTCCATC  CC    TGTC   miRNA 5'  GAGGTAG  GG    ATAG            T       TA  TTGT    TT           3111111133113333121133 ``` \| SeedType = 1 NonSeedType = -1 \| \| --- \| --- \| --- \| --- \| --- \| --- \| | \| 1063 \| WBGene00020299:T07A9.12(12472) \| ``` log2FC=-0.009 padj=0.998 ``` \| [seedLoc = 95,101] Seq=AGAAATACAAATAAATACCATCT \| ```          T  A    AAATA      AAGA mRNA  3'  CT CCAT     AACATA     miRNA 5'  GA GGTA     TTGTAT              T       GTAGG      AGTT           31151111333331111113333 ``` \| SeedType = 3 NonSeedType = 1 \| \| --- \| --- \| --- \| --- \| --- \| --- \| | \| 1064 \| WBGene00020314:T07E3.3(12478) \| ``` log2FC=0.555 padj=0.023 ``` \| [seedLoc = 38,42] Seq=GTTTCCAATTCTTCCTTCCTCAC \| ```          C     T  CT  T    CTTTG   mRNA  3'  ACTCC TC  TC TAAC        miRNA 5'  TGAGG AG  AG GTTG                       T  T        TATAGTT           5111113113511521113333366 ``` \| SeedType = 2 NonSeedType = 1 \| \| --- \| --- \| --- \| --- \| --- \| --- \| | \|  \|  \|  \| [seedLoc = 114,120] Seq=TTTCCCAAATCCTGTACCCTCT \| ```          T  C         TAAACCCTTT mRNA  3'  CT CCAT GTCC           miRNA 5'  GA GGTA TAGG                    T       G    TTGTATAGTT           31151111621113333333333 ``` \| SeedType = 3 NonSeedType = -1 \| \| --- \| --- \| --- \| --- \| --- \| --- \| | \| 1065 \| WBGene00020334:epg-7(12489) \|  \| [seedLoc = 121,126] Seq=CTTTATACGTTCATGTCCCCTTTACCTCT \| ```          T           CCTGTACTT       TC mRNA  3'  CTCCATT TCC         GCATATT   miRNA 5'  GAGGTAG AGG         TGTATAG            T       T   T               TT           311111126111355455555211111233 ``` \| SeedType = 1 NonSeedType = -1 \| \| --- \| --- \| --- \| --- \| --- \| --- \| | \| 1066 \| WBGene00020342:fbxa-201(12495) \| ``` log2FC=0.1 padj=1.0 ``` \| [seedLoc = 142,147] Seq=CACGTTGCTCACTTTACCTCA \| ```                     ACT   TG  C mRNA  3' ACTCCATT TC   CGT  CA  miRNA 5' TGAGGTAG AG   GTA  GT                   T  GTT   TA  T           1111111261133312133113 ``` \| SeedType = 1 NonSeedType = -1 \| \| --- \| --- \| --- \| --- \| --- \| --- \| | \| 1067 \| WBGene00020346:rbm-5(12498) \|  \| [seedLoc = 1711,1716] Seq=TTCTGAAGAACCGTCTACTTCA \| ```                  TG    G A   TT mRNA  3' ACTTCATC  CCAA A GTC   miRNA 5' TGAGGTAG  GGTT T TAG                    TA    G A   TT           1112111133111131321133 ``` \| SeedType = 3 NonSeedType = 1 \| \| --- \| --- \| --- \| --- \| --- \| --- \| | \| 1068 \| WBGene00020368:ast-1(12516) \| ``` log2FC=-0.11 padj=1.0 ``` \| [seedLoc = 93,98] Seq=ATACCAACTTCTTTATTTTTGCTCTACCTTA \| ```                  TCGTTTTTATTTC      CATA    mRNA  3' ATTCCATC             TTCAAC        miRNA 5' TGAGGTAG             AGGTTG                         T                  TATAGTT           1211111135455555555551211113333666 ``` \| SeedType = 3 NonSeedType = 1 \| \| --- \| --- \| --- \| --- \| --- \| --- \| | \|  \|  \|  \| [seedLoc = 180,185] Seq=AGTTGTACATTTTGGTTTTGCACATACCTCT \| ```          T      ACA   TTTTGGTTTT       GA mRNA  3'  CTCCAT    CG          ACATGTT   miRNA 5'  GAGGTA    GT          TGTATAG            T      GTAG                   TT           31111113336125555445555111121233 ``` \| SeedType = 1 NonSeedType = -1 \| \| --- \| --- \| --- \| --- \| --- \| --- \| | \| 1069 \| WBGene00020379:T09B4.5(12525) \| ``` log2FC=-0.097 padj=0.981 ``` \| [seedLoc = 255,260] Seq=CCAATTACATTTCAAATACCTCC \| ```          C      AAA  TT    TAACC mRNA  3'  CTCCAT   CT  ACAT      miRNA 5'  GAGGTA   GG  TGTA               T      GTA  T     TAGTT           31111113331235111133333 ``` \| SeedType = 1 NonSeedType = -1 \| \| --- \| --- \| --- \| --- \| --- \| --- \| | \|  \|  \|  \| [seedLoc = 325,330] Seq=TTTTCCAAAATCCCTACTTCT \| ```          T       C     AACCTTTT mRNA  3'  CTTCATC  CTAA         miRNA 5'  GAGGTAG  GGTT                  T       TA    GTATAGTT           3112111136121133333333 ``` \| SeedType = 3 NonSeedType = 2 \| \| --- \| --- \| --- \| --- \| --- \| --- \| | \| 1070 \| WBGene00020419:acdh-4(12556) \| ``` log2FC=-0.049 padj=0.975 ``` \| [seedLoc = 52,57] Seq=AAACAGAAGGCCTGAAAATGTACCTCT \| ```          T      GTAAAA      AAGA    A mRNA  3'  CTCCAT      GTCCGG     CAA  miRNA 5'  GAGGTA      TAGGTT     GTT           T      G           GTATA               3111111355555211122333361115 ``` \| SeedType = 1 NonSeedType = 2 \| \| --- \| --- \| --- \| --- \| --- \| --- \| | \| 1071 \| WBGene00020426:T10H10.2(12561) \| ``` log2FC=-0.036 padj=0.988 ``` \| [seedLoc = 50,55] Seq=GTTCTCACAACTTTGGCTACCTTT \| ```          T        GT       CTCTTG mRNA  3'  TTCCATCG  TTCAACA       miRNA 5'  GAGGTAGT  AGGTTGT                T                 ATAGTT           321111112451211111333333 ``` \| SeedType = 3 NonSeedType = 1 \| \| --- \| --- \| --- \| --- \| --- \| --- \| | \| 1072 \| WBGene00020437:stt-3(12566) \| ``` log2FC=0.082 padj=0.937 ``` \| [seedLoc = 162,168] Seq=CACTCTTGCGTTCTCTTACCATCT \| ```          T  A     C   T    TC   C mRNA  3'  CT CCATT TCT GCGT  TCA  miRNA 5'  GA GGTAG AGG TGTA  AGT           T        T   T    T    T           311511112311232121351113 ``` \| SeedType = 3 NonSeedType = -1 \| \| --- \| --- \| --- \| --- \| --- \| --- \| | \| 1073 \| WBGene00020443:esyt-2(12570) \| ``` log2FC=-0.081 padj=0.953 ``` \| [seedLoc = 88,93] Seq=AAATGTCTGATTTTTCCTGCCTCT \| ```          T       CTT      C   AAA mRNA  3'  CTCCGTC   TTTAGT TGT    miRNA 5'  GAGGTAG   AGGTTG ATA             T       T        T   GTT           311112113551221223121333 ``` \| SeedType = 2 NonSeedType = 2 \| \| --- \| --- \| --- \| --- \| --- \| --- \| | \| 1074 \| WBGene00020462:ari-2(12582) \|  \| [seedLoc = 66,71] Seq=ATCTAAAAAACAATTTACCTCC \| ```          C       TAA   AAA   TA mRNA  3'  CTCCATT   CAA   ATC   miRNA 5'  GAGGTAG   GTT   TAG            T       TAG   GTA   TT           3111111233311133311133 ``` \| SeedType = 1 NonSeedType = 1 \| \| --- \| --- \| --- \| --- \| --- \| --- \| | \| 1075 \| WBGene00020464:T12E12.3(12584) \| ``` log2FC=-0.126 padj=0.866 ``` \| [seedLoc = 233,238] Seq=AAGTTTTCATTCTCTACCTCT \| ```          T           T  TTTTGAA mRNA  3'  CTCCATC TCT AC        miRNA 5'  GAGGTAG AGG TG                 T       T   T  TATAGTT           3111111161123113333333 ``` \| SeedType = 1 NonSeedType = -1 \| \| --- \| --- \| --- \| --- \| --- \| --- \| | \| 1076 \| WBGene00020467:T12F5.2(12587) \| ``` log2FC=0.154 padj=0.956 ``` \| [seedLoc = 25,31] Seq=TGTTGTACATTTATATTTATCTACCTAC \| ```          CA      T     TATTT       GT mRNA  3'   TCCATC ATTTA     ACATGTT   miRNA 5'   AGGTAG TAGGT     TGTATAG            TG                        TT           3311111151122155555111121233 ``` \| SeedType = 3 NonSeedType = 2 \| \| --- \| --- \| --- \| --- \| --- \| --- \| | \| 1077 \| WBGene00020496:spat-3(12605) \| ``` log2FC=0.38 padj=0.313 ``` \| [seedLoc = 85,91] Seq=TATTGAGCACCACATTACCATCT \| ```          T  A      CA      A    T mRNA  3'  CT CCATTA  CC ACG GTTA  miRNA 5'  GA GGTAGT  GG TGT TAGT           T         A   T   A    T           311511112135116112321213 ``` \| SeedType = 3 NonSeedType = -1 \| \| --- \| --- \| --- \| --- \| --- \| --- \| | \| 1078 \| WBGene00020507:vha-15(12616) \| ``` log2FC=0.241 padj=0.435 ``` \| [seedLoc = 209,215] Seq=ATCTAGTTTTTACCTTCTACCTAC \| ```          CA      T    TTTTTG   TA mRNA  3'   TCCATC TCCA      ATC   miRNA 5'   AGGTAG AGGT      TAG            TG      T    TGTA     TT           331111113111133335411133 ``` \| SeedType = 3 NonSeedType = 1 \| \| --- \| --- \| --- \| --- \| --- \| --- \| | \| 1079 \| WBGene00020511:immt-1(12621) \| ``` log2FC=0.16 padj=0.715 ``` \| [seedLoc = 56,62] Seq=TACTGGTTTTCACTACACTCA \| ```              A        TTTG     T mRNA  3' ACTC CATCA CT     GTCA  miRNA 5' TGAG GTAGT GG     TAGT                     A  TTGTA    T           11115111116123333621113 ``` \| SeedType = 2 NonSeedType = -1 \| \| --- \| --- \| --- \| --- \| --- \| --- \| | \| 1080 \| WBGene00020551:T17H7.7(12636) \| ``` log2FC=-0.922 padj=3.0103780524743E-5 ``` \| [seedLoc = 31,37] Seq=AAATTAATTTATTACTATCCTCT \| ```          T    T      ATT        A mRNA  3'  CTCC ATCATT   TA ATTAA  miRNA 5'  GAGG TAGTAG   GT TAGTT           T           GTT  A                 311115111112333216112115 ``` \| SeedType = 2 NonSeedType = -1 \| \| --- \| --- \| --- \| --- \| --- \| --- \| | \|  \|  \|  \| [seedLoc = 90,95] Seq=GTCTGTATATATCTATCTCG \| ```          G       TAT         TG mRNA  3'  CTCTATC     ATATGTC   miRNA 5'  GAGGTAG     TGTATAG            T       TAGGT       TT           3111211133366121121133 ``` \| SeedType = 2 NonSeedType = -1 \| \| --- \| --- \| --- \| --- \| --- \| --- \| | \|  \|  \|  \| [seedLoc = 121,126] Seq=AAAAAATCTCGCTCTACCTTT \| ```          T          GCTCTAAAAAA mRNA  3'  TTCCATC TC            miRNA 5'  GAGGTAG AG                     T       T  GTTGTATAGTT           3211111161133333333333 ``` \| SeedType = 3 NonSeedType = -1 \| \| --- \| --- \| --- \| --- \| --- \| --- \| | \| 1081 \| WBGene00020605:lgc-50(12676) \| ``` log2FC=0.2 padj=0.939 ``` \| [seedLoc = 37,40] Seq=TTCCCCCATGCCCTCCCTCC \| ```          C    C       TACCCCCTT mRNA  3'  CTCC TC  CCG          miRNA 5'  GAGG AG  GGT                   T    T  TA   TGTATAGTT           3111131166112333333333 ``` \| SeedType = 2 NonSeedType = 2 \| \| --- \| --- \| --- \| --- \| --- \| --- \| | \| 1082 \| WBGene00020612:arrd-22(12678) \| ``` log2FC=-0.147 padj=0.933 ``` \| [seedLoc = 78,83] Seq=GCCTGTATTCAAGCTACATATTTACCTCA \| ```                 TTATA    G   TT     CG mRNA  3' ACTCCAT     CATC AAC  ATGTC   miRNA 5' TGAGGTA     GTAG TTG  TATAG                            G          TT           11111115555511113111551121133 ``` \| SeedType = 1 NonSeedType = 1 \| \| --- \| --- \| --- \| --- \| --- \| --- \| | \| 1083 \| WBGene00020703:T22F7.4(12728) \| ``` log2FC=0.275 padj=0.893 ``` \| [seedLoc = 106,111] Seq=CGCACAACAATCCTTACCTCG \| ```          G       C       ACACGC mRNA  3'  CTCCATT  CTAACA       miRNA 5'  GAGGTAG  GGTTGT                T       TA      ATAGTT           3111111236121111333333 ``` \| SeedType = 1 NonSeedType = 1 \| \| --- \| --- \| --- \| --- \| --- \| --- \| | \|  \|  \|  \| [seedLoc = 121,126] Seq=AACAATCCTTACCTCGA \| ```          A        C       A      mRNA  3'  GCTCCATT  CTAACA       miRNA 5'  TGAGGTAG  GGTTGT                         TA      ATAGTT           52111111236121111366666 ``` \| SeedType = 1 NonSeedType = 1 \| \| --- \| --- \| --- \| --- \| --- \| --- \| | \| 1084 \| WBGene00020740:T23F4.2(12758) \| ``` log2FC=-0.131 padj=0.935 ``` \| [seedLoc = 58,64] Seq=TTTTTCTCAACTTTTACTCTCA \| ```              T           TCTTTTT mRNA  3' ACTC CATT TTCAAC        miRNA 5' TGAG GTAG AGGTTG                          T      TATAGTT           11115111261211113333333 ``` \| SeedType = 2 NonSeedType = 1 \| \| --- \| --- \| --- \| --- \| --- \| --- \| | \| 1085 \| WBGene00020777:T24E12.10(12778) \| ``` log2FC=-1.407 padj=1.0 ``` \| [seedLoc = 108,113] Seq=AAATTCCCCGCTGAACTACCTCT \| ```          T        AG   CCCCTTAAA mRNA  3'  CTCCATCA  TCG          miRNA 5'  GAGGTAGT  GGT                   T        A    TGTATAGTT           31111111134212333333333 ``` \| SeedType = 1 NonSeedType = 2 \| \| --- \| --- \| --- \| --- \| --- \| --- \| | \| 1086 \| WBGene00020779:T24G10.2(12780) \| ``` log2FC=-0.088 padj=0.939 ``` \| [seedLoc = 39,45] Seq=TGTTATAATTATCTCCTACACTCA \| ```              A    C    TTA     GT mRNA  3' ACTC CATC TCTA   ATATT   miRNA 5' TGAG GTAG AGGT   TATAG                     T    TG      TT           111151111311213351111233 ``` \| SeedType = 2 NonSeedType = 2 \| \| --- \| --- \| --- \| --- \| --- \| --- \| | \| 1087 \| WBGene00020784:tat-4(12785) \| ``` log2FC=-0.254 padj=0.534 ``` \| [seedLoc = 77,82] Seq=ACACAAACGGGTACTATCTCT \| ```          T         G     AACACA mRNA  3'  CTCTATCAT  GGCA       miRNA 5'  GAGGTAGTA  TTGT                T         GG    ATAGTT           3111211111362211333333 ``` \| SeedType = 2 NonSeedType = -1 \| \| --- \| --- \| --- \| --- \| --- \| --- \| | \| 1088 \| WBGene00020803:T25E4.1(12797) \| ``` log2FC=-1.203 padj=5.96953608835658E-7 ``` \| [seedLoc = 141,146] Seq=AAATTGTTTTCAAGTATTACCTCT \| ```          T         G    TTT      A mRNA  3'  CTCCATTAT  AAC   TGTTAA  miRNA 5'  GAGGTAGTA  TTG   ATAGTT           T         GG   T                    3111111211361113551212115 ``` \| SeedType = 1 NonSeedType = 1 \| \| --- \| --- \| --- \| --- \| --- \| --- \| | \| 1089 \| WBGene00020806:T25F10.3(12798) \| ``` log2FC=-0.557 padj=0.095 ``` \| [seedLoc = 87,92] Seq=ATCTGTCCTCAGGACCTGTACCTCA \| ```                        G CTCC    TA mRNA  3' ACTCCAT GTCCAG A    TGTC   miRNA 5' TGAGGTA TAGGTT T    ATAG                   G      G         TT           11111116211112315555121133 ``` \| SeedType = 1 NonSeedType = 1 \| \| --- \| --- \| --- \| --- \| --- \| --- \| | \| 1090 \| WBGene00020821:jhdm-1(12809) \| ``` log2FC=-0.003 padj=0.998 ``` \| [seedLoc = 48,53] Seq=CACGTTCAATCTCCTTCACTATCTCA \| ```                   CTTCC      TTG   C mRNA  3' ACTCTATCA     TCTAAC    CA  miRNA 5' TGAGGTAGT     AGGTTG    GT                               TATA  T           111121111555551121113336113 ``` \| SeedType = 2 NonSeedType = 1 \| \| --- \| --- \| --- \| --- \| --- \| --- \| | \|  \|  \|  \| [seedLoc = 63,67] Seq=TCTCCTTCACTATCTCAC \| ```          C         CT   TCT        mRNA  3'  ACTCTATCA  TCC           miRNA 5'  TGAGGTAGT  AGG                                   TTGTATAGTT           5111121111551113336666666 ``` \| SeedType = 2 NonSeedType = -1 \| \| --- \| --- \| --- \| --- \| --- \| --- \| | \| 1091 \| WBGene00020821:jhdm-1(12808) \| ``` log2FC=-0.003 padj=0.998 ``` \| [seedLoc = 265,270] Seq=CATTGTTTTTTCAAGCTACCTCA \| ```                   AA TTTTT     C mRNA  3' ACTCCATCG  C     TGTTA  miRNA 5' TGAGGTAGT  G     ATAGT                    AG TTGT      T           11111111233133335121213 ``` \| SeedType = 1 NonSeedType = -1 \| \| --- \| --- \| --- \| --- \| --- \| --- \| | \| 1092 \| WBGene00020843:T27A3.7(12829) \| ``` log2FC=-0.004 padj=0.998 ``` \| [seedLoc = 208,213] Seq=TCCCTCTTCACTTTCCTACCTTC \| ```          C       CT    CTTCTCCCT mRNA  3'  TTCCATC  TTCA          miRNA 5'  GAGGTAG  AGGT                   T       T     TGTATAGTT           32111111351211333333333 ``` \| SeedType = 3 NonSeedType = 2 \| \| --- \| --- \| --- \| --- \| --- \| --- \| | \|  \|  \|  \| [seedLoc = 237,241] Seq=TTCCCACACACGTGTCAAATACCTCAC \| ```          C       AAA T  G  C   CCCTT  mRNA  3'  ACTCCAT   C GT CA ACA       miRNA 5'  TGAGGTA   G TA GT TGT                               G      ATAGTT           5111111155515213115111333336 ``` \| SeedType = 1 NonSeedType = -1 \| \| --- \| --- \| --- \| --- \| --- \| --- \| | \| 1093 \| WBGene00020867:shc-2(12846) \| ``` log2FC=-0.113 padj=0.949 ``` \| [seedLoc = 460,466] Seq=ACGTAACCGCCACCTCAA \| ```          A      C        GCA     mRNA  3'  ACTCCA CG CCAAT        miRNA 5'  TGAGGT GT GGTTG                        A  A     TATAGTT           51111113126111123336666 ``` \| SeedType = 2 NonSeedType = 1 \| \| --- \| --- \| --- \| --- \| --- \| --- \| | \| 1094 \| WBGene00020902:jmjc-1(12869) \| ``` log2FC=0.469 padj=0.154 ``` \| [seedLoc = 75,81] Seq=TTGTTTTTTTCTTGTTACCATCT \| ```          T  A         TTTTTTTGTT mRNA  3'  CT CCATTGTTC           miRNA 5'  GA GGTAGTAGG                    T            TTGTATAGTT           31151111221213333333333 ``` \| SeedType = 3 NonSeedType = -1 \| \| --- \| --- \| --- \| --- \| --- \| --- \| | \| 1095 \| WBGene00020935:fnci-1(12891) \| ``` log2FC=-0.242 padj=0.649 ``` \| [seedLoc = 177,183] Seq=TGACTGCTACAGTATACCGTCT \| ```          T  G    AT        C     T mRNA  3'  CT CCAT     GACAT GTCAG  miRNA 5'  GA GGTA     TTGTA TAGTT           T       GTAGG                       3114111133666211115211125 ``` \| SeedType = 3 NonSeedType = -1 \| \| --- \| --- \| --- \| --- \| --- \| --- \| | \| 1096 \| WBGene00020936:hrpf-1(12892) \| ``` log2FC=-0.038 padj=0.98 ``` \| [seedLoc = 315,321] Seq=CCGGACAATGGAAGATATTAAATTACCATCT \| ```          T  A      AATTATAGAAGG     GGCC   mRNA  3'  CT CCATTA            TAACA       miRNA 5'  GA GGTAGT            GTTGT                T         AG               ATAGTT           311511112133555554554421111333366 ``` \| SeedType = 3 NonSeedType = 1 \| \| --- \| --- \| --- \| --- \| --- \| --- \| | \| 1097 \| WBGene00020999:W03F9.1(12927) \| ``` log2FC=0.002 padj=0.998 ``` \| [seedLoc = 215,221] Seq=GAAAAACCAAACTACAGTACCCTCT \| ```          T  C    GA    AA  CAAAAAG mRNA  3'  CT CCAT  CATC  AC        miRNA 5'  GA GGTA  GTAG  TG                 T             GT  TATAGTT           3115111145111133113333333 ``` \| SeedType = 3 NonSeedType = -1 \| \| --- \| --- \| --- \| --- \| --- \| --- \| | \| 1098 \| WBGene00021009:afd-1(12935) \| ``` log2FC=-0.196 padj=0.768 ``` \| [seedLoc = 216,221] Seq=ATATTTACAAATTCCACGCCTCCTGCCTCC \| ```          C       C    CACCTTAA    TTATA mRNA  3'  CTCCGTC TCCG        ACAT      miRNA 5'  GAGGTAG AGGT        TGTA               T       T                TAGTT           311112113111255555555111133333 ``` \| SeedType = 2 NonSeedType = 2 \| \| --- \| --- \| --- \| --- \| --- \| --- \| | \| 1099 \| WBGene00021019:W04B5.2(12940) \| ``` log2FC=0.195 padj=1.0 ``` \| [seedLoc = 209,215] Seq=CCGAAATGGCTTAGGTTAAACTTTACGCTCA \| ```              G    TCAAATTGG        AAGCC  mRNA  3' ACTC CATT         ATTCGGTA       miRNA 5' TGAG GTAG         TAGGTTGT                                          ATAGTT           11114111255555554411212221333336 ``` \| SeedType = 2 NonSeedType = 2 \| \| --- \| --- \| --- \| --- \| --- \| --- \| | \| 1100 \| WBGene00021020:W04B5.3(12941) \| ``` log2FC=0.085 padj=0.945 ``` \| [seedLoc = 104,108] Seq=AAATATTTAACTCTAGCTCGT \| ```          T    G          T   AAA mRNA  3'  GCTC ATC  TCAAT TAT    miRNA 5'  TGAG TAG  GGTTG ATA                  G   TA     T   GTT           52111311166211123111333 ``` \| SeedType = 2 NonSeedType = 1 \| \| --- \| --- \| --- \| --- \| --- \| --- \| | \| 1101 \| WBGene00021047:zfp-3(12958) \| ``` log2FC=-0.437 padj=0.343 ``` \| [seedLoc = 677,683] Seq=AAGCTGTGGCCTATTTTTTTTTCTATCCTCT \| ```          T    T   TTTTTTTTT              A mRNA  3'  CTCC ATC         ATCCG  GTGTCGA  miRNA 5'  GAGG TAG         TAGGT  TATAGTT           T                      TG                   311115111555555555111126621211215 ``` \| SeedType = 2 NonSeedType = 2 \| \| --- \| --- \| --- \| --- \| --- \| --- \| | \|  \|  \|  \| [seedLoc = 1741,1746] Seq=TCTCCGTGGTCCAGCTATCTCC \| ```          C        A  TGGTGCCTCT mRNA  3'  CTCTATCG CC           miRNA 5'  GAGGTAGT GG                    T        A  TTGTATAGTT           3111211123113333333333 ``` \| SeedType = 2 NonSeedType = -1 \| \| --- \| --- \| --- \| --- \| --- \| --- \| | \| 1102 \| WBGene00021068:W06H8.6(12974) \| ``` log2FC=-0.151 padj=0.873 ``` \| [seedLoc = 199,204] Seq=AAACTCCAAACTCTTGTCATCTACCTTT \| ```          T       TACTGTTC    ACC     A mRNA  3'  TTCCATC        TCAA    TCAA  miRNA 5'  GAGGTAG        GGTT    AGTT           T       TA          GTAT                32111111335545552111333611115 ``` \| SeedType = 3 NonSeedType = 1 \| \| --- \| --- \| --- \| --- \| --- \| --- \| | \|  \|  \|  \| [seedLoc = 402,407] Seq=GATCTCCCGACCGGGCCTATCTCC \| ```          C       CGGG     CCTCTAG mRNA  3'  CTCTATC    CCAGC        miRNA 5'  GAGGTAG    GGTTG                 T       TA       TATAGTT           311121113344111213333333 ``` \| SeedType = 2 NonSeedType = 1 \| \| --- \| --- \| --- \| --- \| --- \| --- \| | \|  \|  \|  \| [seedLoc = 2721,2726] Seq=CCCAATAGTGTCTTATTTCTATCTCT \| ```          T       TTT    TGTG   ACCC mRNA  3'  CTCTATC   ATTC    ATA     miRNA 5'  GAGGTAG   TAGG    TAT              T              TTG    AGTT           31112111555112133341113333 ``` \| SeedType = 2 NonSeedType = -1 \| \| --- \| --- \| --- \| --- \| --- \| --- \| | \| 1103 \| WBGene00021088:W08E12.7(12987) \| ``` log2FC=-0.084 padj=0.911 ``` \| [seedLoc = 66,71] Seq=TACTATCCCCTATCTGCCTCT \| ```          T       T    C        T mRNA  3'  CTCCGTC ATCC  C TATCA  miRNA 5'  GAGGTAG TAGG  G ATAGT           T            TT T     T           31111211511113616111113 ``` \| SeedType = 2 NonSeedType = -1 \| \| --- \| --- \| --- \| --- \| --- \| --- \| | \|  \|  \|  \| [seedLoc = 168,174] Seq=GGTAAAAGCCGTAGTCTCAGAACTACGCTCA \| ```              G     AGACTCTGATG    AA   G    mRNA  3' ACTC CATCA           CCGA  ATG     miRNA 5' TGAG GTAGT           GGTT  TAT                        A              G    AGTT           1111411111345555545541121351123666 ``` \| SeedType = 2 NonSeedType = 2 \| \| --- \| --- \| --- \| --- \| --- \| --- \| | \| 1104 \| WBGene00021093:W08F4.3(12989) \| ``` log2FC=0.547 padj=0.098 ``` \| [seedLoc = 96,101] Seq=CAATGTCCCCCCTCTACTTCC \| ```          C           CCCCTGTAAC mRNA  3'  CTTCATC TCC           miRNA 5'  GAGGTAG AGG                    T       T   TTGTATAGTT           3112111161113333333333 ``` \| SeedType = 3 NonSeedType = -1 \| \| --- \| --- \| --- \| --- \| --- \| --- \| | \| 1105 \| WBGene00021214:hasp-2(13050) \|  \| [seedLoc = 819,825] Seq=AAATCGGTGGCCGAATATCCTCT \| ```          T    T  AAG    TGGCTAAA mRNA  3'  CTCC AT   CCGG         miRNA 5'  GAGG TA   GGTT                  T       GTA    GTATAGTT           31111511333112233333333 ``` \| SeedType = 2 NonSeedType = 2 \| \| --- \| --- \| --- \| --- \| --- \| --- \| | \| 1106 \| WBGene00021237:crtc-1(13053) \| ``` log2FC=-0.454 padj=0.264 ``` \| [seedLoc = 133,138] Seq=GTGATGGGCCTGGCACCATGTCTCTATCTCT \| ```          T       TCTGTACCACG      GTAGTG   mRNA  3'  CTCTATC           GTCCGG         miRNA 5'  GAGGTAG           TAGGTT                  T                        GTATAGTT           311121115554555555421112233333366 ``` \| SeedType = 2 NonSeedType = 2 \| \| --- \| --- \| --- \| --- \| --- \| --- \| | \|  \|  \|  \| [seedLoc = 357,363] Seq=AAATCGATAATTTTATTACTCTCT \| ```          T   T     T       GCTAAA mRNA  3'  CTC CATTA TTTAATA       miRNA 5'  GAG GTAGT AGGTTGT                T                 ATAGTT           311151112151221121333333 ``` \| SeedType = 2 NonSeedType = 2 \| \| --- \| --- \| --- \| --- \| --- \| --- \| | \| 1107 \| WBGene00021238:gap-3(13054) \| ``` log2FC=-0.204 padj=0.832 ``` \| [seedLoc = 229,235] Seq=ATATGTGCATCGTTTTACCCTCC \| ```          C  C     T G T      ATA mRNA  3'  CT CCATT T C ACGTGT    miRNA 5'  GA GGTAG A G TGTATA             T        T G T      GTT           31151111231313112121333 ``` \| SeedType = 3 NonSeedType = -1 \| \| --- \| --- \| --- \| --- \| --- \| --- \| | \| 1108 \| WBGene00021326:Y34B4A.10(13114) \| ``` log2FC=-0.059 padj=0.98 ``` \| [seedLoc = 99,104] Seq=AAATGTATTCTCTACCTTG \| ```          G                  AAA mRNA  3'  TTCCATC TCT  TATGT    miRNA 5'  GAGGTAG AGG  GTATA             T       T   TT     GTT           3211111161126621121333 ``` \| SeedType = 3 NonSeedType = -1 \| \| --- \| --- \| --- \| --- \| --- \| --- \| | \| 1109 \| WBGene00021328:npr-23(13116) \| ``` log2FC=0.168 padj=0.973 ``` \| [seedLoc = 294,299] Seq=GCAATGAGACTACCTGAC \| ```          C G       GAG            mRNA  3'  A TCCATCA   TAACG       miRNA 5'  T AGGTAGT   GTTGT                  G       AG      ATAGTT           513111111133421112666666 ``` \| SeedType = 3 NonSeedType = 1 \| \| --- \| --- \| --- \| --- \| --- \| --- \| | \| 1110 \| WBGene00021465:ekl-7(13197) \|  \| [seedLoc = 249,254] Seq=TTCTCCTAATTTTACTATCTCC \| ```          C           TAATCCTCTT mRNA  3'  CTCTATCATTT           miRNA 5'  GAGGTAGTAGG                    T           TTGTATAGTT           3111211111223333333333 ``` \| SeedType = 2 NonSeedType = -1 \| \| --- \| --- \| --- \| --- \| --- \| --- \| | \| 1111 \| WBGene00021474:dot-1.1(13208) \| ``` log2FC=0.037 padj=0.981 ``` \| [seedLoc = 464,469] Seq=ATCTGAACTCCCTTACCTCA \| ```                      CT  A   TA mRNA  3' ACTCCAT  TCC  CA GTC   miRNA 5' TGAGGTA  AGG  GT TAG                   GT   TT  A   TT           1111111661113311321133 ``` \| SeedType = 1 NonSeedType = -1 \| \| --- \| --- \| --- \| --- \| --- \| --- \| | \| 1112 \| WBGene00021485:Y40A1A.3(13215) \| ``` log2FC=0.1 padj=1.0 ``` \| [seedLoc = 244,249] Seq=ATCCGTCTCACCTTGCGGCTACTTCC \| ```          C        GCGT    CTCTGCCTA mRNA  3'  CTTCATCG    TCCA          miRNA 5'  GAGGTAGT    AGGT                   T                TGTATAGTT           31121111245451111333333333 ``` \| SeedType = 3 NonSeedType = 1 \| \| --- \| --- \| --- \| --- \| --- \| --- \| | \| 1113 \| WBGene00021495:sqst-2(13221) \| ``` log2FC=0.16 padj=0.851 ``` \| [seedLoc = 168,173] Seq=ATTTTAATTTCCTGAAATCTATCTCG \| ```          G       TAAA    TT  ATTTTA mRNA  3'  CTCTATC    GTCC  TA       miRNA 5'  GAGGTAG    TAGG  GT                T               TT  ATAGTT           31112111555521113321333333 ``` \| SeedType = 2 NonSeedType = -1 \| \| --- \| --- \| --- \| --- \| --- \| --- \| | \| 1114 \| WBGene00021506:Y41D4A.4(13230) \| ``` log2FC=-0.054 padj=0.965 ``` \| [seedLoc = 64,67] Seq=CTGAATCTCCCCTTCTCCCTCC \| ```          C    C  T   CCTCTAAGTC mRNA  3'  CTCC TC TCC           miRNA 5'  GAGG AG AGG                    T    T  T   TTGTATAGTT           3111131131113333333333 ``` \| SeedType = 2 NonSeedType = -1 \| \| --- \| --- \| --- \| --- \| --- \| --- \| | \| 1115 \| WBGene00021522:nhr-274(13242) \| ``` log2FC=-0.056 padj=0.98 ``` \| [seedLoc = 443,448] Seq=CTCTGTGATAAGTCACTCTCTACCTCC \| ```          C           C  TGAATA     TC mRNA  3'  CTCCATC TCT AC      GTGTC   miRNA 5'  GAGGTAG AGG TG      TATAG            T       T   T             TT           3111111161123115455552121133 ``` \| SeedType = 1 NonSeedType = -1 \| \| --- \| --- \| --- \| --- \| --- \| --- \| | \| 1116 \| WBGene00021528:gbb-1(13246) \| ``` log2FC=0.195 padj=0.961 ``` \| [seedLoc = 358,363] Seq=TTCCCCACGGCCGCTACCTCC \| ```          C               CCCCTT mRNA  3'  CTCCATCG CCGGCA       miRNA 5'  GAGGTAGT GGTTGT                T        A      ATAGTT           3111111126112211333333 ``` \| SeedType = 1 NonSeedType = 2 \| \| --- \| --- \| --- \| --- \| --- \| --- \| | \| 1117 \| WBGene00021531:Y42G9A.1(13249) \| ``` log2FC=-0.434 padj=0.541 ``` \| [seedLoc = 48,53] Seq=TATTATATTTAAGCACTTACCTCG \| ```          G     T     G  TT       T mRNA  3'  CTCCA TCA C AA  TATATTA  miRNA 5'  GAGGT AGT G TT  GTATAGT           T         A G           T           3111115111613115521111213 ``` \| SeedType = 2 NonSeedType = -1 \| \| --- \| --- \| --- \| --- \| --- \| --- \| | \| 1118 \| WBGene00021539:Y42H9AR.4(13258) \| ``` log2FC=-0.024 padj=0.992 ``` \| [seedLoc = 219,225] Seq=TTTTCGTGTTCCTCTCTCACCTCC \| ```          C     C  TC   TTGTGCTTTT mRNA  3'  CTCCA TC  TCC           miRNA 5'  GAGGT AG  AGG                    T        T    TTGTATAGTT           311111511351113333333333 ``` \| SeedType = 2 NonSeedType = -1 \| \| --- \| --- \| --- \| --- \| --- \| --- \| | \| 1119 \| WBGene00021546:laat-1(13266) \| ``` log2FC=0.462 padj=0.121 ``` \| [seedLoc = 186,192] Seq=ATCTATATATACATATTTTGTACTAACCTCC \| ```          C    A     GTTTTATA AT       TA mRNA  3'  CTCC ATCAT        C  ATATATC   miRNA 5'  GAGG TAGTA        G  TGTATAG            T                   GT       TT           3111151111145555555133121111133 ``` \| SeedType = 2 NonSeedType = -1 \| \| --- \| --- \| --- \| --- \| --- \| --- \| | \| 1120 \| WBGene00021579:clec-73(13286) \| ``` log2FC=-0.008 padj=0.994 ``` \| [seedLoc = 117,122] Seq=GTTTAGTTGTTCAAGTACCTCT \| ```          T      G ACTTGTTGATTTG mRNA  3'  CTCCAT A              miRNA 5'  GAGGTA T                       T      G AGGTTGTATAGTT           3111111313333333333333 ``` \| SeedType = 1 NonSeedType = -1 \| \| --- \| --- \| --- \| --- \| --- \| --- \| | \| 1121 \| WBGene00021598:Y46E12BR.1(13297) \| ``` log2FC=-1.346 padj=0.17 ``` \| [seedLoc = 431,436] Seq=CTTTTGTTGATAAATACCTCA \| ```                 AAA    TTGTTTTC mRNA  3' ACTCCAT    TAG         miRNA 5' TGAGGTA    GTT                         GTAG   GTATAGTT           1111111333621233333333 ``` \| SeedType = 1 NonSeedType = -1 \| \| --- \| --- \| --- \| --- \| --- \| --- \| | \| 1122 \| WBGene00021629:scpl-3(13314) \| ``` log2FC=0.091 padj=0.982 ``` \| [seedLoc = 93,98] Seq=TTCTTTCCAAGAAAATACCTCA \| ```                 AAAAG   CTTTCTT mRNA  3' ACTCCAT     AAC        miRNA 5' TGAGGTA     TTG                        GTAGG   TATAGTT           1111111333331113333333 ``` \| SeedType = 1 NonSeedType = 1 \| \| --- \| --- \| --- \| --- \| --- \| --- \| | \|  \|  \|  \| [seedLoc = 226,232] Seq=CCCCAAAAAACCGTCTCAGATACTCTCT \| ```          T   T   AGACTCTG    AAAACCCC mRNA  3'  CTC CAT        CCAA         miRNA 5'  GAG GTA        GGTT                  T       GTA         GTATAGTT           3111511133355554111133333333 ``` \| SeedType = 2 NonSeedType = 1 \| \| --- \| --- \| --- \| --- \| --- \| --- \| | \| 1123 \| WBGene00021630:Y47D9A.3(13315) \| ``` log2FC=-0.642 padj=0.666 ``` \| [seedLoc = 365,370] Seq=AAACATGGGATCCGTTTACCTCG \| ```          G       TG  T  G       A mRNA  3'  CTCCATT  CC AG GTA CAA  miRNA 5'  GAGGTAG  GG TT TAT GTT           T       TA     G   A               311111123311512321161115 ``` \| SeedType = 1 NonSeedType = -1 \| \| --- \| --- \| --- \| --- \| --- \| --- \| | \| 1124 \| WBGene00021644:Y47G6A.18(13328) \| ``` log2FC=-0.01 padj=0.994 ``` \| [seedLoc = 57,63] Seq=CAAGTCTAAACCGGTACCATCA \| ```             A    GG     ATCTGAAC mRNA  3' ACT CCAT   CCAA         miRNA 5' TGA GGTA   GGTT                          GTA    GTATAGTT           11151111336111133333333 ``` \| SeedType = 3 NonSeedType = 1 \| \| --- \| --- \| --- \| --- \| --- \| --- \| | \| 1125 \| WBGene00021675:Y48G1BR.1(13347) \| ``` log2FC=-0.178 padj=0.812 ``` \| [seedLoc = 249,254] Seq=CATCTCATTTTCTAGGTTACCTCT \| ```          T       GG    TTTACTCTAC mRNA  3'  CTCCATT  ATCT           miRNA 5'  GAGGTAG  TAGG                    T             TTGTATAGTT           311111124411123333333333 ``` \| SeedType = 1 NonSeedType = -1 \| \| --- \| --- \| --- \| --- \| --- \| --- \| | \| 1126 \| WBGene00021679:Y48G1C.6(13351) \| ``` log2FC=0.232 padj=0.933 ``` \| [seedLoc = 27,32] Seq=TATTTTGAAAATTGCTATCTCT \| ```          T          AAAAGTTTTAT mRNA  3'  CTCTATCGTT            miRNA 5'  GAGGTAGTAG                     T          GTTGTATAGTT           3111211121233333333333 ``` \| SeedType = 2 NonSeedType = -1 \| \| --- \| --- \| --- \| --- \| --- \| --- \| | \| 1127 \| WBGene00021732:Y49G5B.1(13381) \| ``` log2FC=-0.234 padj=0.742 ``` \| [seedLoc = 535,541] Seq=CCTAGTGCTTCCTGCCGTAGTTACCATCT \| ```          T  A    TGATGC     TT    ATCC mRNA  3'  CT CCAT      CGTCC  CGTG     miRNA 5'  GA GGTA      GTAGG  GTAT              T                  TT    AGTT           31151111545545121113312123333 ``` \| SeedType = 3 NonSeedType = -1 \| \| --- \| --- \| --- \| --- \| --- \| --- \| | \| 1128 \| WBGene00021810:phf-15(13431) \| ``` log2FC=-0.014 padj=0.994 ``` \| [seedLoc = 425,430] Seq=ATTTTTCAGAGCTCTGCCTCT \| ```          T          G  ACTTTTTA mRNA  3'  CTCCGTC TC AG         miRNA 5'  GAGGTAG AG TT                  T       T  G  GTATAGTT           3111121161131233333333 ``` \| SeedType = 2 NonSeedType = -1 \| \| --- \| --- \| --- \| --- \| --- \| --- \| | \| 1129 \| WBGene00021834:mab-31(13448) \| ``` log2FC=-0.229 padj=0.459 ``` \| [seedLoc = 92,97] Seq=AGTCGTGGATCCCGTTTACCTCA \| ```                  TG  CT G   CTGA mRNA  3' ACTCCATT  CC  A GTG     miRNA 5' TGAGGTAG  GG  T TAT                      TA  T  G   AGTT           11111112331135132123333 ``` \| SeedType = 1 NonSeedType = -1 \| \| --- \| --- \| --- \| --- \| --- \| --- \| | \| 1130 \| WBGene00021841:Y54E10BR.2(13455) \| ``` log2FC=0.038 padj=0.989 ``` \| [seedLoc = 76,82] Seq=TCCTGTGATGATTTCCTAACCTCA \| ```               A   C       G    CT mRNA  3' ACTCC ATC TTTAGTA TGTC   miRNA 5' TGAGG TAG AGGTTGT ATAG                     T            TT           111115111312212214121133 ``` \| SeedType = 2 NonSeedType = 2 \| \| --- \| --- \| --- \| --- \| --- \| --- \| | \| 1131 \| WBGene00021890:Y54G2A.26(13495) \| ``` log2FC=-0.317 padj=0.402 ``` \| [seedLoc = 471,476] Seq=CCCTGTACTCGTGAAAATGTTTTACCTCC \| ```          C       T G   AAGTGCT      CC mRNA  3'  CTCCATT T TAA       CATGTC   miRNA 5'  GAGGTAG A GTT       GTATAG            T       T G                TT           31111112313211554545511121133 ``` \| SeedType = 1 NonSeedType = -1 \| \| --- \| --- \| --- \| --- \| --- \| --- \| | \| 1132 \| WBGene00021952:vha-19(13542) \| ``` log2FC=0.236 padj=0.597 ``` \| [seedLoc = 224,229] Seq=TCGAAATTTTTCTATTTCTATCTCT \| ```          T       TTT    TTTTAAAGCT mRNA  3'  CTCTATC   ATCT           miRNA 5'  GAGGTAG   TAGG                    T              TTGTATAGTT           3111211155511123333333333 ``` \| SeedType = 2 NonSeedType = -1 \| \| --- \| --- \| --- \| --- \| --- \| --- \| | \| 1133 \| WBGene00022014:Y61A9LA.1(13571) \| ``` log2FC=0.402 padj=0.521 ``` \| [seedLoc = 371,376] Seq=TAAATTTGAATTTACTATCTCT \| ```          T             GTTTAAAT mRNA  3'  CTCTATCATTTAA         miRNA 5'  GAGGTAGTAGGTT                  T             GTATAGTT           3111211111221133333333 ``` \| SeedType = 2 NonSeedType = 2 \| \| --- \| --- \| --- \| --- \| --- \| --- \| | \| 1134 \| WBGene00022078:epg-9(13622) \| ``` log2FC=-0.653 padj=0.113 ``` \| [seedLoc = 395,400] Seq=AATTCCACCACTGAAATTACCTCT \| ```          T        AAG   C  CCTTAA mRNA  3'  CTCCATTA   TCA CA       miRNA 5'  GAGGTAGT   GGT GT                T        A     T  ATAGTT           311111121354211311333333 ``` \| SeedType = 1 NonSeedType = 2 \| \| --- \| --- \| --- \| --- \| --- \| --- \| | \| 1135 \| WBGene00022087:Y69A2AR.16(13624) \| ``` log2FC=0.199 padj=0.657 ``` \| [seedLoc = 129,135] Seq=TTTTTTTTTTCTTTTTACCATCA \| ```             A     T   TTTTTTTTTT mRNA  3' ACT CCATT TTC           miRNA 5' TGA GGTAG AGG                             T   TTGTATAGTT           11151111231213333333333 ``` \| SeedType = 3 NonSeedType = -1 \| \| --- \| --- \| --- \| --- \| --- \| --- \| | \| 1136 \| WBGene00022099:Y69A2AR.28(13633) \| ``` log2FC=-0.259 padj=0.513 ``` \| [seedLoc = 62,67] Seq=TACCATACTCTCTCTCTTCTATCTCT \| ```          T       T  T TCTCT    C  T mRNA  3'  CTCTATC TC C     CATA CA  miRNA 5'  GAGGTAG AG G     GTAT GT           T       T    TT       A  T           31112111311513355511113113 ``` \| SeedType = 2 NonSeedType = -1 \| \| --- \| --- \| --- \| --- \| --- \| --- \| | \| 1137 \| WBGene00022128:Y71F9B.6(13656) \| ``` log2FC=-0.037 padj=0.988 ``` \| [seedLoc = 97,103] Seq=TCCTTAAAGGTGGACTACGCTCA \| ```              G     GGTGGAAATTCCT mRNA  3' ACTC CATCA              miRNA 5' TGAG GTAGT                                 AGGTTGTATAGTT           11114111113333333333333 ``` \| SeedType = 2 NonSeedType = -1 \| \| --- \| --- \| --- \| --- \| --- \| --- \| | \| 1138 \| WBGene00022139:tub-2(13662) \| ``` log2FC=-0.194 padj=0.766 ``` \| [seedLoc = 32,37] Seq=GGCTCATATCACCAAATCCTACCTTT \| ```          T       CTAAA   C    C   G mRNA  3'  TTCCATC     CCA TATA TCG  miRNA 5'  GAGGTAG     GGT GTAT AGT           T       TA      T        T           32111111335551113211151123 ``` \| SeedType = 3 NonSeedType = 1 \| \| --- \| --- \| --- \| --- \| --- \| --- \| | \| 1139 \| WBGene00022177:Y71H2AM.12(13691) \| ``` log2FC=-0.328 padj=0.786 ``` \| [seedLoc = 26,32] Seq=GGTGCTGCATTCCGGAATACTCTCA \| ```              T   AAGG  TT    CGTGG mRNA  3' ACTC CAT    CC  ACGT      miRNA 5' TGAG GTA    GG  TGTA                       GTA   T     TAGTT           1111511133341135112133333 ``` \| SeedType = 2 NonSeedType = -1 \| \| --- \| --- \| --- \| --- \| --- \| --- \| | \| 1140 \| WBGene00022203:Y73B3A.1(13716) \| ``` log2FC=-0.17 padj=0.791 ``` \| [seedLoc = 54,59] Seq=TCCCATATCTCTTTCTATCTCG \| ```          G       T   TC    CCCT mRNA  3'  CTCTATC TTC  TATA     miRNA 5'  GAGGTAG AGG  GTAT              T       T   TT    AGTT           3111211131213321113333 ``` \| SeedType = 2 NonSeedType = -1 \| \| --- \| --- \| --- \| --- \| --- \| --- \| | \| 1141 \| WBGene00022221:Y73B3A.20(13720) \| ``` log2FC=-0.622 padj=0.836 ``` \| [seedLoc = 30,36] Seq=TGGATCGAGACGGTTATACACTCA \| ```              A   ATTGG   AGCTAGGT mRNA  3' ACTC CAT     CAG         miRNA 5' TGAG GTA     GTT                          GTAG    GTATAGTT           111151113333411233333333 ``` \| SeedType = 2 NonSeedType = -1 \| \| --- \| --- \| --- \| --- \| --- \| --- \| | \| 1142 \| WBGene00022235:sqd-1(13725) \| ``` log2FC=0.155 padj=0.76 ``` \| [seedLoc = 858,863] Seq=CCCCCCCCAACCGATTCTAATAACTGCCTCT \| ```          T        ATAATCTTAG     CCCCCCC mRNA  3'  CTCCGTCA          CCAAC        miRNA 5'  GAGGTAGT          GGTTG                 T        A              TATAGTT           3111121113555555554111113333333 ``` \| SeedType = 2 NonSeedType = 1 \| \| --- \| --- \| --- \| --- \| --- \| --- \| | \| 1143 \| WBGene00022253:hrpf-2(13738) \| ``` log2FC=-0.143 padj=0.812 ``` \| [seedLoc = 196,201] Seq=ACGCGAACGTCTTATTTTACCTCT \| ```          T      TT     T   AGCGCA mRNA  3'  CTCCAT  TATTC GCA       miRNA 5'  GAGGTA  GTAGG TGT                T             T   ATAGTT           311111155211213211333333 ``` \| SeedType = 1 NonSeedType = -1 \| \| --- \| --- \| --- \| --- \| --- \| --- \| | \| 1144 \| WBGene00022270:Y73E7A.3(13750) \| ``` log2FC=0.301 padj=0.661 ``` \| [seedLoc = 449,455] Seq=TTTGTTACGTGTTACCAAGATTCACCTCT \| ```          T     C   GAA   TTGT    TGTTT mRNA  3'  CTCCA TTA   CCA    GCAT      miRNA 5'  GAGGT AGT   GGT    TGTA               T         A             TAGTT           31111151213551115545211133333 ``` \| SeedType = 2 NonSeedType = 1 \| \| --- \| --- \| --- \| --- \| --- \| --- \| | \| 1145 \| WBGene00022274:Y73E7A.8(13753) \| ``` log2FC=0.91 padj=0.056 over-expressed ``` \| [seedLoc = 61,66] Seq=AAAAAACCAGCAGATCTTACCTCA \| ```                 TC  GA    CAAAAAA mRNA  3' ACTCCAT  TA  CGAC        miRNA 5' TGAGGTA  GT  GTTG                            AG    TATAGTT           111111155213312113333333 ``` \| SeedType = 1 NonSeedType = -1 \| \| --- \| --- \| --- \| --- \| --- \| --- \| | \|  \|  \|  \| [seedLoc = 76,80] Seq=CAGCAGATCTTACCTCAA \| ```          A             G   AC     mRNA  3'  ACTCCAT  TCTA ACG       miRNA 5'  TGAGGTA  AGGT TGT                        GT        ATAGTT           511111116611214112336666 ``` \| SeedType = 1 NonSeedType = 2 \| \| --- \| --- \| --- \| --- \| --- \| --- \| | \| 1146 \| WBGene00022279:sesn-1(13757) \| ``` log2FC=0.189 padj=0.74 ``` \| [seedLoc = 657,663] Seq=TTTCTTTTTCACTAATTCTACACTCA \| ```              A    TTA   ACTTTTTCTTT mRNA  3' ACTC CATC   ATC            miRNA 5' TGAG GTAG   TAG                                    GTTGTATAGTT           11115111155511133333333333 ``` \| SeedType = 2 NonSeedType = -1 \| \| --- \| --- \| --- \| --- \| --- \| --- \| | \| 1147 \| WBGene00022300:Y76B12C.6(13772) \| ``` log2FC=0.272 padj=0.705 ``` \| [seedLoc = 574,580] Seq=CACTAAGAAATGTGAATACCCTCC \| ```          C  C    AA  G   AGA    C mRNA  3'  CT CCAT  GT TAA   ATCA  miRNA 5'  GA GGTA  TA GTT   TAGT           T       G   G   GTA    T           311511113521321133311113 ``` \| SeedType = 3 NonSeedType = -1 \| \| --- \| --- \| --- \| --- \| --- \| --- \| | \| 1148 \| WBGene00022310:Y77E11A.7(13779) \| ``` log2FC=-0.009 padj=0.994 ``` \| [seedLoc = 176,182] Seq=TAATAATTTCCCCCCTACCATCC \| ```          C  A     CC  CTTTAATAAT mRNA  3'  CT CCATC  CC           miRNA 5'  GA GGTAG  GG                    T        TA  TTGTATAGTT           31151111133113333333333 ``` \| SeedType = 3 NonSeedType = -1 \| \| --- \| --- \| --- \| --- \| --- \| --- \| | \| 1149 \| WBGene00022352:Y82E9BR.20(13798) \| ``` log2FC=0.1 padj=1.0 ``` \| [seedLoc = 74,79] Seq=CTCCGTACCAACCTAGCTGGGCACTACTTCT \| ```          T        CGGGTCGA     C    CCTC mRNA  3'  CTTCATCA        TCCAA CATG     miRNA 5'  GAGGTAGT        AGGTT GTAT              T                          AGTT           3112111115444554511111511123333 ``` \| SeedType = 3 NonSeedType = 1 \| \| --- \| --- \| --- \| --- \| --- \| --- \| | \| 1150 \| WBGene00022383:Y95B8A.2(13824) \| ``` log2FC=-0.662 padj=0.026 ``` \| [seedLoc = 264,270] Seq=GAATTGTGTTGGTTTAATTTTACCGTCG \| ```          G  G            TTGGTT       G mRNA  3'  CT CCAT  TTTAAT      GTGTTAA  miRNA 5'  GA GGTA  AGGTTG      TATAGTT           T       GT                               311411116612211255445521212114 ``` \| SeedType = 3 NonSeedType = 2 \| \| --- \| --- \| --- \| --- \| --- \| --- \| | \| 1151 \| WBGene00022386:Y95B8A.6(13826) \| ``` log2FC=0.406 padj=0.905 ``` \| [seedLoc = 372,378] Seq=CCTCGAACAGTTTGCTAGCCTCG \| ```          G    G           AGCTCC mRNA  3'  CTCC ATCGTTTGACA       miRNA 5'  GAGG TAGTAGGTTGT                T                ATAGTT           31111411121222111333333 ``` \| SeedType = 2 NonSeedType = 2 \| \| --- \| --- \| --- \| --- \| --- \| --- \| | \| 1152 \| WBGene00022388:Y95B8A.8(13827) \| ``` log2FC=-0.068 padj=0.949 ``` \| [seedLoc = 150,155] Seq=TTATTTTTCTACTATTACCTCC \| ```          C          ATCTTTTTATT mRNA  3'  CTCCATTATC            miRNA 5'  GAGGTAGTAG                     T          GTTGTATAGTT           3111111211133333333333 ``` \| SeedType = 1 NonSeedType = -1 \| \| --- \| --- \| --- \| --- \| --- \| --- \| | \| 1153 \| WBGene00022389:pde-6(13829) \| ``` log2FC=0.192 padj=0.822 ``` \| [seedLoc = 154,160] Seq=ATTTTTCAAATCTATCGTTATCCTCT \| ```          T    T  TGC       ACTTTTTA mRNA  3'  CTCC AT   TATCTAA         miRNA 5'  GAGG TA   GTAGGTT                  T                 GTATAGTT           31111511545211121133333333 ``` \| SeedType = 2 NonSeedType = 2 \| \| --- \| --- \| --- \| --- \| --- \| --- \| | \| 1154 \| WBGene00022397:Y97E10AR.2(13835) \| ``` log2FC=0.133 padj=0.968 ``` \| [seedLoc = 532,537] Seq=TTAGTAGAAACTGGAGTACCTCA \| ```                 G GG    AGATGATT mRNA  3' ACTCCAT A  TCAA         miRNA 5' TGAGGTA T  GGTT                         G A     GTATAGTT           11111113134211133333333 ``` \| SeedType = 1 NonSeedType = 1 \| \| --- \| --- \| --- \| --- \| --- \| --- \| | \| 1155 \| WBGene00022401:Y97E10AR.6(13839) \| ``` log2FC=-0.142 padj=0.795 ``` \| [seedLoc = 79,84] Seq=CACTGTCACTCCTATCTCT \| ```          T       C            C mRNA  3'  CTCTATC TC  AC TGTCA  miRNA 5'  GAGGTAG AG  TG ATAGT           T       T  GT  T     T           3111211131166116121113 ``` \| SeedType = 2 NonSeedType = -1 \| \| --- \| --- \| --- \| --- \| --- \| --- \| | \| 1156 \| WBGene00022410:Y97E10C.1(13841) \| ``` log2FC=-1.592 padj=1.0 ``` \| [seedLoc = 561,567] Seq=TCTGATACTCGTTCCCAATACTACTCTCA \| ```              T      AAC  TTGCT    GTCT mRNA  3' ACTC CATCAT   CC     CATA     miRNA 5' TGAG GTAGTA   GG     GTAT                              TT       AGTT           11115111111555113345511113333 ``` \| SeedType = 2 NonSeedType = -1 \| \| --- \| --- \| --- \| --- \| --- \| --- \| | \| 1157 \| WBGene00022423:nhr-41(13851) \| ``` log2FC=-0.247 padj=0.617 ``` \| [seedLoc = 178,183] Seq=CAATTCCCAGTTTGAATACCTCT \| ```          T      AA       CCTTAAC mRNA  3'  CTCCAT  GTTTGAC        miRNA 5'  GAGGTA  TAGGTTG                 T      G        TATAGTT           31111113521222113333333 ``` \| SeedType = 1 NonSeedType = 2 \| \| --- \| --- \| --- \| --- \| --- \| --- \| | \| 1158 \| WBGene00022428:Y104H12D.4(13855) \| ``` log2FC=-0.162 padj=0.926 ``` \| [seedLoc = 96,100] Seq=CTCTGTTTCTCTGCTTCCTCGA \| ```          A     T      CTT     TC mRNA  3'  GCTCC TCGTCT    TGTC   miRNA 5'  TGAGG AGTAGG    ATAG                  T      TTGT    TT           52111131121123336121133 ``` \| SeedType = 2 NonSeedType = -1 \| \| --- \| --- \| --- \| --- \| --- \| --- \| | \| 1159 \| WBGene00022452:Y110A2AR.1(13866) \| ``` log2FC=0.024 padj=0.993 ``` \| [seedLoc = 65,70] Seq=TCCTGTATATCACTAACACCTACCTTT \| ```          T       CACA   A T       CT mRNA  3'  TTCCATC    ATC C ATATGTC   miRNA 5'  GAGGTAG    TAG G TGTATAG            T                T       TT           321111115555111513121121133 ``` \| SeedType = 3 NonSeedType = -1 \| \| --- \| --- \| --- \| --- \| --- \| --- \| | \| 1160 \| WBGene00022462:Y110A7A.15(13875) \| ``` log2FC=-0.06 padj=0.959 ``` \| [seedLoc = 131,136] Seq=TCTTCAACATTCATAAACTACCTCC \| ```          C        AATA  T   ACTTCT mRNA  3'  CTCCATCA    CT ACA       miRNA 5'  GAGGTAGT    GG TGT                T        A     T   ATAGTT           3111111113555123111333333 ``` \| SeedType = 1 NonSeedType = -1 \| \| --- \| --- \| --- \| --- \| --- \| --- \| | \| 1161 \| WBGene00022474:zig-9(13883) \| ``` log2FC=-0.223 padj=0.679 ``` \| [seedLoc = 23,29] Seq=TGTTATCAGATTTATTTTTTCTACTCTCC \| ```          C   T    TTTTTT      AC    GT mRNA  3'  CTC CATC      ATTTAG  TATT   miRNA 5'  GAG GTAG      TAGGTT  ATAG            T                    GT    TT           31115111155555511221233111233 ``` \| SeedType = 2 NonSeedType = 2 \| \| --- \| --- \| --- \| --- \| --- \| --- \| | \| 1162 \| WBGene00022531:morc-1(13918) \| ``` log2FC=0.064 padj=0.953 ``` \| [seedLoc = 109,114] Seq=CATTATCGTTTTTACCTCA \| ```                  TTT          C mRNA  3' ACTCCATT     GC TATTA  miRNA 5' TGAGGTAG     TG ATAGT                   TAGGT  T     T           1111111233366216111213 ``` \| SeedType = 1 NonSeedType = -1 \| \| --- \| --- \| --- \| --- \| --- \| --- \| | \| 1163 \| WBGene00022538:ZC190.4(13923) \| ``` log2FC=-0.058 padj=0.98 ``` \| [seedLoc = 378,383] Seq=AAACCGTATCCATATATACCTCT \| ```          T      A   A        C   A mRNA  3'  CTCCAT TAT CC  TATG CAA  miRNA 5'  GAGGTA GTA GG  GTAT GTT           T             TT    A               3111111521151166211231115 ``` \| SeedType = 1 NonSeedType = -1 \| \| --- \| --- \| --- \| --- \| --- \| --- \| | \|  \|  \|  \| [seedLoc = 400,405] Seq=CTCTAAACATCTCTTCAAATACCTCT \| ```          T      AAACT  T T   A   TC mRNA  3'  CTCCAT     TC C ACA ATC   miRNA 5'  GAGGTA     AG G TGT TAG            T      GT       T   A   TT           31111113355511513111311133 ``` \| SeedType = 1 NonSeedType = -1 \| \| --- \| --- \| --- \| --- \| --- \| --- \| | \| 1164 \| WBGene00022591:cuti-1(13952) \| ``` log2FC=0.498 padj=0.26 ``` \| [seedLoc = 60,66] Seq=CATTATAATTTCTCTACCATCA \| ```             A         TTA      C mRNA  3' ACT CCATC TCT   ATATTA  miRNA 5' TGA GGTAG AGG   TATAGT                    T   TTG      T           11151111161123331111213 ``` \| SeedType = 3 NonSeedType = -1 \| \| --- \| --- \| --- \| --- \| --- \| --- \| | \| 1165 \| WBGene00022592:klu-2(13953) \|  \| [seedLoc = 73,78] Seq=TATTATAAGCTTCAGTACCTCT \| ```          T      G C            T mRNA  3'  CTCCAT A TTCGA ATATTA  miRNA 5'  GAGGTA T AGGTT TATAGT           T      G       G      T           31111113151212161111213 ``` \| SeedType = 1 NonSeedType = 2 \| \| --- \| --- \| --- \| --- \| --- \| --- \| | \| 1166 \| WBGene00022608:madf-9(13960) \| ``` log2FC=-0.307 padj=0.616 ``` \| [seedLoc = 181,186] Seq=TTAAAAACTTTTTGCTACCTTC \| ```          C           TT  AAAATT mRNA  3'  TTCCATCGTTT  CA       miRNA 5'  GAGGTAGTAGG  GT                T           TT  ATAGTT           3211111121223311333333 ``` \| SeedType = 3 NonSeedType = -1 \| \| --- \| --- \| --- \| --- \| --- \| --- \| | \| 1167 \| WBGene00022648:ZK54.3(13987) \| ``` log2FC=-0.1 padj=1.0 ``` \| [seedLoc = 100,105] Seq=TTAGTTACAACTACCTCC \| ```          C                TGATT mRNA  3'  CTCCATCA    ACAT      miRNA 5'  GAGGTAGT    TGTA               T        AGGT    TAGTT           3111111116666111133333 ``` \| SeedType = 1 NonSeedType = -1 \| \| --- \| --- \| --- \| --- \| --- \| --- \| | \| 1168 \| WBGene00022673:ZK177.8(14001) \| ``` log2FC=0.066 padj=0.945 ``` \| [seedLoc = 52,57] Seq=AAATTATACATTTTACTGCCTCC \| ```          C           T         A mRNA  3'  CTCCGTCATTT ACATATTAA  miRNA 5'  GAGGTAGTAGG TGTATAGTT           T           T                     31111211112231111112115 ``` \| SeedType = 2 NonSeedType = -1 \| \| --- \| --- \| --- \| --- \| --- \| --- \| | \| 1169 \| WBGene00022677:rfth-1(14003) \|  \| [seedLoc = 32,37] Seq=TTTTCATTTTTTTTGTACCTCG \| ```          G      GTTTTTTTTACTTTT mRNA  3'  CTCCAT                miRNA 5'  GAGGTA                         T      GTAGGTTGTATAGTT           3111111333333333333333 ``` \| SeedType = 1 NonSeedType = -1 \| \| --- \| --- \| --- \| --- \| --- \| --- \| | \| 1170 \| WBGene00022697:cyy-1(14013) \| ``` log2FC=0.003 padj=0.998 ``` \| [seedLoc = 410,415] Seq=GGCTATTTAATTGACTCAATACCTATCTCA \| ```                  CATAACTCAG     T     G mRNA  3' ACTCTATC          TTAAT TATCG  miRNA 5' TGAGGTAG          GGTTG ATAGT                   TA             T     T           111121113355555554221123111123 ``` \| SeedType = 2 NonSeedType = 2 \| \| --- \| --- \| --- \| --- \| --- \| --- \| | \|  \|  \|  \| [seedLoc = 421,425] Seq=TGACTCAATACCTATCTCAG \| ```          G        CA            T mRNA  3'  ACTCTATC   TAAC   TCAG  miRNA 5'  TGAGGTAG   GTTG   AGTT                    TAG    TAT                411112111336211166611125 ``` \| SeedType = 2 NonSeedType = 1 \| \| --- \| --- \| --- \| --- \| --- \| --- \| | \| 1171 \| WBGene00022698:ZK353.2(14014) \| ``` log2FC=0.353 padj=0.855 ``` \| [seedLoc = 57,62] Seq=GGTTTCGATATGTATTTACCTCA \| ```                T    G  TAGCTTTGG mRNA  3' ACTCCA TTAT TA          miRNA 5' TGAGGT AGTA GT                              G  TGTATAGTT           11111151211321333333333 ``` \| SeedType = 2 NonSeedType = -1 \| \| --- \| --- \| --- \| --- \| --- \| --- \| | \| 1172 \| WBGene00022702:cutc-1(14017) \| ``` log2FC=0.218 padj=0.663 ``` \| [seedLoc = 437,443] Seq=CTCTGGTCACTCTTTCTCTACCTGC \| ```          CG      TCT   TC  TG   TC mRNA  3'   TCCATC   TTC  AC  GTC   miRNA 5'   AGGTAG   AGG  TG  TAG            TG      T     T   TA   TT           3311111135512135113321133 ``` \| SeedType = 3 NonSeedType = -1 \| \| --- \| --- \| --- \| --- \| --- \| --- \| | \| 1173 \| WBGene00022712:ZK355.2(14023) \| ``` log2FC=0.101 padj=0.953 ``` \| [seedLoc = 30,35] Seq=CGCTGGACACCTTCTCAACAGTACCTCT \| ```          T      G CAACTCT       G    C mRNA  3'  CTCCAT A       TCC ACA GTCG  miRNA 5'  GAGGTA T       AGG TGT TAGT           T      G           T   A    T           31111113155555551116111321123 ``` \| SeedType = 1 NonSeedType = -1 \| \| --- \| --- \| --- \| --- \| --- \| --- \| | \| 1174 \| WBGene00022735:ZK418.6(14040) \| ``` log2FC=-0.181 padj=0.959 ``` \| [seedLoc = 31,36] Seq=TTTTACACAATAAAAAATCTACCTCC \| ```          C       TAAAAAA     CATTTT mRNA  3'  CTCCATC       TAACA       miRNA 5'  GAGGTAG       GTTGT                T       TAG         ATAGTT           31111111333555521111333333 ``` \| SeedType = 1 NonSeedType = 1 \| \| --- \| --- \| --- \| --- \| --- \| --- \| | \| 1175 \| WBGene00022747:ZK470.6(14047) \| ``` log2FC=-1.489 padj=0.046 ``` \| [seedLoc = 99,104] Seq=ATTTAATTCCCCTTCCCTACCTTC \| ```          C       CCT   CCTTAATTTA mRNA  3'  TTCCATC   TCC           miRNA 5'  GAGGTAG   AGG                    T       T     TTGTATAGTT           321111113551113333333333 ``` \| SeedType = 3 NonSeedType = -1 \| \| --- \| --- \| --- \| --- \| --- \| --- \| | \| 1176 \| WBGene00022749:ZK484.3(14051) \| ``` log2FC=-0.272 padj=0.524 ``` \| [seedLoc = 281,286] Seq=ATCATGTTTCATTGCTATCTCA \| ```                     ACTTTGTACTA mRNA  3' ACTCTATCGTT            miRNA 5' TGAGGTAGTAG                                GTTGTATAGTT           1111211121233333333333 ``` \| SeedType = 2 NonSeedType = -1 \| \| --- \| --- \| --- \| --- \| --- \| --- \| | \| 1177 \| WBGene00022753:ZK484.7(14055) \| ``` log2FC=-0.073 padj=0.966 ``` \| [seedLoc = 63,68] Seq=GAATAAAAAAGTTCTACCTTT \| ```          T          G  AAAATAAG mRNA  3'  TTCCATC TT AA         miRNA 5'  GAGGTAG AG TT                  T       T  G  GTATAGTT           3211111161231133333333 ``` \| SeedType = 3 NonSeedType = -1 \| \| --- \| --- \| --- \| --- \| --- \| --- \| | \| 1178 \| WBGene00022775:ZK616.5(14068) \| ``` log2FC=-0.259 padj=0.407 ``` \| [seedLoc = 242,248] Seq=CATAATACGATCGAAGGTTATTTACCATCG \| ```          G  A     TATTGGAAG        ATAC mRNA  3'  CT CCATT         CTAGCATA     miRNA 5'  GA GGTAG         GGTTGTAT              T        TA               AGTT           311511112335544554121211113333 ``` \| SeedType = 3 NonSeedType = 2 \| \| --- \| --- \| --- \| --- \| --- \| --- \| | \| 1179 \| WBGene00022791:ZK686.1(14078) \| ``` log2FC=0.263 padj=0.902 ``` \| [seedLoc = 60,65] Seq=CCTTGGATTTCCGCTATCTCT \| ```          T           TT  GGTTCC mRNA  3'  CTCTATCG CC  TA       miRNA 5'  GAGGTAGT GG  GT                T        A  TT  ATAGTT           3111211126113321333333 ``` \| SeedType = 2 NonSeedType = -1 \| \| --- \| --- \| --- \| --- \| --- \| --- \| | \| 1180 \| WBGene00022825:ZK816.1(14104) \| ``` log2FC=0.349 padj=1.0 ``` \| [seedLoc = 33,38] Seq=TTGAAAGTTACCCCTACCTTA \| ```                  C    TTGAAAGTT mRNA  3' ATTCCATC  CCA          miRNA 5' TGAGGTAG  GGT                           TA   TGTATAGTT           1211111136111333333333 ``` \| SeedType = 3 NonSeedType = 1 \| \| --- \| --- \| --- \| --- \| --- \| --- \| | \| 1181 \| WBGene00022855:tcer-1(14123) \| ``` log2FC=0.004 padj=0.997 ``` \| [seedLoc = 412,417] Seq=CCCCGGATGGCCGGAAATACTACTTCA \| ```                    AAAGG      GGCCCC mRNA  3' ACTTCATCAT     CCGGTA       miRNA 5' TGAGGTAGTA     GGTTGT                                     ATAGTT           111211111155544112221333333 ``` \| SeedType = 3 NonSeedType = 2 \| \| --- \| --- \| --- \| --- \| --- \| --- \| | \| 1182 \| WBGene00022861:dve-1(14128) \| ``` log2FC=-0.283 padj=0.52 ``` \| [seedLoc = 164,169] Seq=AAGAGCAAAATCACTACTTCA \| ```                        AACGAGAA mRNA  3' ACTTCATCA CTAA         miRNA 5' TGAGGTAGT GGTT                           A    GTATAGTT           1112111116121133333333 ``` \| SeedType = 3 NonSeedType = 2 \| \| --- \| --- \| --- \| --- \| --- \| --- \| | \|  \|  \|  \| [seedLoc = 412,418] Seq=TCAATGTTTCCCTTCTTACCGTCA \| ```             G     CT   CTTTGTAACT mRNA  3' ACT CCATT  TCC           miRNA 5' TGA GGTAG  AGG                             T    TTGTATAGTT           111411112351113333333333 ``` \| SeedType = 3 NonSeedType = -1 \| \| --- \| --- \| --- \| --- \| --- \| --- \| | \| 1183 \| WBGene00022881:ZK1248.11(14140) \| ``` log2FC=-0.098 padj=0.938 ``` \| [seedLoc = 83,88] Seq=CACTTCACTTCTCAATACCTCT \| ```          T      AAC  TT  CT   C mRNA  3'  CTCCAT   TC  CA  TCA  miRNA 5'  GAGGTA   GG  GT  AGT           T      GTA  TT  AT   T           3111111333213311331113 ``` \| SeedType = 1 NonSeedType = -1 \| \| --- \| --- \| --- \| --- \| --- \| --- \| | \| 1184 \| WBGene00023405:sor-1(14153) \| ``` log2FC=-0.071 padj=0.959 ``` \| [seedLoc = 58,63] Seq=TGTTACCAAACCATATTTTGTGTACCTCT \| ```          T      GTGTTT   A    ACCATTGT mRNA  3'  CTCCAT      TAT CCAA         miRNA 5'  GAGGTA      GTA GGTT                  T                    GTATAGTT           31111114545552115111133333333 ``` \| SeedType = 1 NonSeedType = 1 \| \| --- \| --- \| --- \| --- \| --- \| --- \| | \| 1185 \| WBGene00023491:tsp-21(14180) \| ``` log2FC=0.242 padj=0.934 ``` \| [seedLoc = 144,149] Seq=CTCTGTATAAAATTTGCCTCTACTTCA \| ```                       TTTAAA       TC mRNA  3' ACTTCATC TCCG      ATATGTC   miRNA 5' TGAGGTAG AGGT      TGTATAG                    T                 TT           1112111161112555555121121133 ``` \| SeedType = 3 NonSeedType = 2 \| \| --- \| --- \| --- \| --- \| --- \| --- \| | \| 1186 \| WBGene00023500:C11H1.9(14186) \| ``` log2FC=0.761 padj=8.64117179257199E-4 over-expressed ``` \| [seedLoc = 96,102] Seq=TATTATATTTGACAATACTCTCT \| ```          T   T   AA     TT       T mRNA  3'  CTC CAT    CAG  TATATTA  miRNA 5'  GAG GTA    GTT  GTATAGT           T       GTAG            T           3111511133661125521111213 ``` \| SeedType = 2 NonSeedType = -1 \| \| --- \| --- \| --- \| --- \| --- \| --- \| | \| 1187 \| WBGene00023520:oig-5(14193) \| ``` log2FC=0.187 padj=0.962 ``` \| [seedLoc = 118,124] Seq=TGCGTGGTAGCCTAAAGAGGTTACCGTCT \| ```          T  G     GGAGAA       GGTG  T mRNA  3'  CT CCATT      ATCCGAT    CG  miRNA 5'  GA GGTAG      TAGGTTG    GT           T                     TATA  T           31141111244545511112123333123 ``` \| SeedType = 3 NonSeedType = 2 \| \| --- \| --- \| --- \| --- \| --- \| --- \| | \| 1188 \| WBGene00043062:clec-80(14200) \| ``` log2FC=0.144 padj=0.918 ``` \| [seedLoc = 289,295] Seq=CACCAGGCGGACCCGGGGTACACTCT \| ```          T   A   G GGC   G   GACCAC mRNA  3'  CTC CAT G   CCA GCG       miRNA 5'  GAG GTA T   GGT TGT                T       G A         ATAGTT           31115111323451114212333333 ``` \| SeedType = 2 NonSeedType = 1 \| \| --- \| --- \| --- \| --- \| --- \| --- \| | \| 1189 \| WBGene00043064:nbet-1(14202) \| ``` log2FC=0.165 padj=0.824 ``` \| [seedLoc = 196,202] Seq=GAAATTTTGGAAGATTACACTCA \| ```              A     GAAGGTTTTAAAG mRNA  3' ACTC CATTA              miRNA 5' TGAG GTAGT                                 AGGTTGTATAGTT           11115111213333333333333 ``` \| SeedType = 2 NonSeedType = -1 \| \| --- \| --- \| --- \| --- \| --- \| --- \| | \| 1190 \| WBGene00044068:syd-9(14253) \| ``` log2FC=-0.02 padj=0.996 ``` \| [seedLoc = 286,291] Seq=TTTCTCCCGGAATGCATACCTCA \| ```                 A   AA   CCTCTTT mRNA  3' ACTCCAT CGT  GGC        miRNA 5' TGAGGTA GTA  TTG                            GG   TATAGTT           11111115121332213333333 ``` \| SeedType = 1 NonSeedType = -1 \| \| --- \| --- \| --- \| --- \| --- \| --- \| | \| 1191 \| WBGene00044071:dhhc-14(14257) \| ``` log2FC=-0.331 padj=0.547 ``` \| [seedLoc = 269,274] Seq=TTAAATGCTCTCTATCTCC \| ```          C                 AATT mRNA  3'  CTCTATC TCT  CGTA     miRNA 5'  GAGGTAG AGG  GTAT              T       T   TT    AGTT           3111211161126612113333 ``` \| SeedType = 2 NonSeedType = -1 \| \| --- \| --- \| --- \| --- \| --- \| --- \| | \| 1192 \| WBGene00044158:Y53C12A.8(14283) \| ``` log2FC=-0.086 padj=0.984 ``` \| [seedLoc = 63,69] Seq=GTGCCCACAACTTTTTATCCTCT \| ```          T    T   T       CCCGTG mRNA  3'  CTCC ATT TTCAACA       miRNA 5'  GAGG TAG AGGTTGT                T        T       ATAGTT           31111511231211111333333 ``` \| SeedType = 2 NonSeedType = 1 \| \| --- \| --- \| --- \| --- \| --- \| --- \| | \| 1193 \| WBGene00044319:tag-266(14343) \| ``` log2FC=-0.055 padj=0.977 ``` \| [seedLoc = 530,536] Seq=AAACACCCATTCTTCTCACCTCA \| ```                C  T   T  CCACAAA mRNA  3' ACTCCA TC TCT AC        miRNA 5' TGAGGT AG AGG TG                          T   T  TATAGTT           11111151131123113333333 ``` \| SeedType = 2 NonSeedType = -1 \| \| --- \| --- \| --- \| --- \| --- \| --- \| | \| 1194 \| WBGene00044330:alr-1(14351) \| ``` log2FC=-0.063 padj=0.989 ``` \| [seedLoc = 232,237] Seq=TCCTGACCGCCAATGTTCTCGATTACCTCG \| ```          G        GCTCTTGTAA   C      CT mRNA  3'  CTCCATTA          CCG CA GTC   miRNA 5'  GAGGTAGT          GGT GT TAG            T        A            T  A   TT           3111111213555554555112311621133 ``` \| SeedType = 1 NonSeedType = 2 \| \| --- \| --- \| --- \| --- \| --- \| --- \| | \| 1195 \| WBGene00044386:C27A2.7(14369) \| ``` log2FC=-1.708 padj=1.0 ``` \| [seedLoc = 49,54] Seq=CTCTATCTATCTTATTACGATTACCTCT \| ```          T        GCATTAT    T      TC mRNA  3'  CTCCATTA       TCTA C TATC   miRNA 5'  GAGGTAGT       AGGT G ATAG            T                   T T    TT           31111112145555551121316111133 ``` \| SeedType = 1 NonSeedType = 2 \| \| --- \| --- \| --- \| --- \| --- \| --- \| | \| 1196 \| WBGene00044467:R05C11.4(14407) \| ``` log2FC=-0.242 padj=0.887 ``` \| [seedLoc = 70,76] Seq=GTCTTGACACGAAGTACTCTCA \| ```              T   G AG     GT  TG mRNA  3' ACTC CAT A  C ACA  TC   miRNA 5' TGAG GTA T  G TGT  AG                    G AG T   AT  TT           11115111313316111331133 ``` \| SeedType = 2 NonSeedType = -1 \| \| --- \| --- \| --- \| --- \| --- \| --- \| | \| 1197 \| WBGene00044509:K07H8.12(14429) \| ``` log2FC=-0.036 padj=0.994 ``` \| [seedLoc = 82,87] Seq=TCCTATTTTTTCCATTTCTACTTCC \| ```          C       TTTA  TTTTT    CT mRNA  3'  CTTCATC    CC     TATC   miRNA 5'  GAGGTAG    GG     ATAG            T       TA    TTGT     TT           3112111133551133335111133 ``` \| SeedType = 3 NonSeedType = -1 \| \| --- \| --- \| --- \| --- \| --- \| --- \| | \| 1198 \| WBGene00044533:F41F3.8(14438) \| ``` log2FC=-0.561 padj=0.305 ``` \| [seedLoc = 154,160] Seq=GGTTGTTAGCTTGATTTTTCTCACCTCG \| ```          G     C  TTTTTA            GG mRNA  3'  CTCCA TC      GTTCGAT TGTT   miRNA 5'  GAGGT AG      TAGGTTG ATAG            T                     T    TT           31111151155555521212126121233 ``` \| SeedType = 2 NonSeedType = 2 \| \| --- \| --- \| --- \| --- \| --- \| --- \| | \| 1199 \| WBGene00044623:bus-8(14463) \| ``` log2FC=-0.204 padj=0.805 ``` \| [seedLoc = 287,292] Seq=TTTTAAGCATTCTTCTACTTCT \| ```          T       T   T   AATTTT mRNA  3'  CTTCATC TCT ACG       miRNA 5'  GAGGTAG AGG TGT                T       T   T   ATAGTT           3112111131123112333333 ``` \| SeedType = 3 NonSeedType = -1 \| \| --- \| --- \| --- \| --- \| --- \| --- \| | \| 1200 \| WBGene00044699:nhr-286(14499) \| ``` log2FC=0.306 padj=0.859 ``` \| [seedLoc = 40,46] Seq=TTTTTTTAAATTTCCTACCGTCA \| ```             G     C     ATTTTTTT mRNA  3' ACT CCATC TTTAA         miRNA 5' TGA GGTAG AGGTT                           T     GTATAGTT           11141111131221133333333 ``` \| SeedType = 3 NonSeedType = 2 \| \| --- \| --- \| --- \| --- \| --- \| --- \| | \| 1201 \| WBGene00045276:F18E3.13(14594) \| ``` log2FC=-0.018 padj=0.993 ``` \| [seedLoc = 84,89] Seq=ACAAGTATAATTTACCTCT \| ```          T                 AACA mRNA  3'  CTCCATT   TAATATG     miRNA 5'  GAGGTAG   GTTGTAT              T       TAG       AGTT           3111111266621121123333 ``` \| SeedType = 1 NonSeedType = -1 \| \| --- \| --- \| --- \| --- \| --- \| --- \| | \| 1202 \| WBGene00045305:ZC250.4(14596) \| ``` log2FC=0.724 padj=0.098 over-expressed ``` \| [seedLoc = 90,96] Seq=GTTGGTAGATTCTTCCTACCATCA \| ```             A     CT   T G   GTTG mRNA  3' ACT CCATC  TCT A ATG     miRNA 5' TGA GGTAG  AGG T TAT                       T    T G   AGTT           111511111351123131123333 ``` \| SeedType = 3 NonSeedType = -1 \| \| --- \| --- \| --- \| --- \| --- \| --- \| | \| 1203 \| WBGene00045362:F11D11.12(14604) \| ``` log2FC=0.1 padj=1.0 ``` \| [seedLoc = 93,97] Seq=CAATGGCCCGCCTCTACGTCG \| ```          G  G         CCCGGTAAC mRNA  3'  CT CATC TCCG          miRNA 5'  GA GTAG AGGT                   T  G    T    TGTATAGTT           3113111161112333333333 ``` \| SeedType = 3 NonSeedType = 2 \| \| --- \| --- \| --- \| --- \| --- \| --- \| | \| 1204 \| WBGene00045483:oxy-5(14638) \| ``` log2FC=-0.016 padj=0.993 ``` \| [seedLoc = 54,60] Seq=TAATTGTCGCTTTCTCACCTCT \| ```          T     C  T    C        T mRNA  3'  CTCCA TC TTCG   TGTTAA  miRNA 5'  GAGGT AG AGGT   ATAGTT           T        T    TGT                  311111511312123661212115 ``` \| SeedType = 2 NonSeedType = 2 \| \| --- \| --- \| --- \| --- \| --- \| --- \| | \| 1205 \| WBGene00050941:C33F10.14(14665) \| ``` log2FC=-0.004 padj=0.998 ``` \| [seedLoc = 58,63] Seq=CTCTATTTTGCTACTTCA \| ```                              TC mRNA  3' ACTTCATCGTTT    TATC   miRNA 5' TGAGGTAGTAGG    ATAG                        TTGT    TT           1112111121226666111133 ``` \| SeedType = 3 NonSeedType = -1 \| \| --- \| --- \| --- \| --- \| --- \| --- \| | \| 1206 \| WBGene00050944:C14A11.9(14667) \| ``` log2FC=-0.06 padj=0.98 ``` \| [seedLoc = 77,82] Seq=CCTCGTGCAATACTACTTCA \| ```                            CTCC mRNA  3' ACTTCATCA  TAACGTG     miRNA 5' TGAGGTAGT  GTTGTAT                       AG       AGTT           1112111116621112123333 ``` \| SeedType = 3 NonSeedType = 1 \| \| --- \| --- \| --- \| --- \| --- \| --- \| | \| 1207 \| WBGene00077490:M03A1.8(14671) \| ``` log2FC=0.028 padj=1.0 ``` \| [seedLoc = 116,121] Seq=CACTTTATCACCAACTATCTCT \| ```          T        A   C   T   C mRNA  3'  CTCTATCA CCA TAT TCA  miRNA 5'  GAGGTAGT GGT GTA AGT           T        A   T   T   T           3111211113111321131113 ``` \| SeedType = 2 NonSeedType = 1 \| \| --- \| --- \| --- \| --- \| --- \| --- \| | \| 1208 \| WBGene00077521:maf-1(14676) \| ``` log2FC=1.626 padj=1.0 ``` \| [seedLoc = 203,208] Seq=GCGCCAACAACAATACCTCT \| ```          T      AA       ACCGCG mRNA  3'  CTCCAT    CAACA       miRNA 5'  GAGGTA    GTTGT                T      GTAG     ATAGTT           3111111336611111333333 ``` \| SeedType = 1 NonSeedType = 1 \| \| --- \| --- \| --- \| --- \| --- \| --- \| | \| 1209 \| WBGene00086546:cil-1(14718) \| ``` log2FC=-0.192 padj=0.86 ``` \| [seedLoc = 125,131] Seq=TATGGTACTCCCTATTAGTGTTACACTCC \| ```          C   A   TGTGAT     CT    GTAT mRNA  3'  CTC CAT      TATCC  CATG     miRNA 5'  GAG GTA      GTAGG  GTAT              T                  TT    AGTT           31115111545455211113311123333 ``` \| SeedType = 2 NonSeedType = -1 \| \| --- \| --- \| --- \| --- \| --- \| --- \| | \| 1210 \| WBGene00175030:Y53C12A.10(14739) \| ``` log2FC=-0.106 padj=0.934 ``` \| [seedLoc = 109,115] Seq=GTTGTTTGTTACTTTACACTCA \| ```              A       ATTGTTTGTTG mRNA  3' ACTC CATT TC            miRNA 5' TGAG GTAG AG                              T  GTTGTATAGTT           11115111261133333333333 ``` \| SeedType = 2 NonSeedType = -1 \| \| --- \| --- \| --- \| --- \| --- \| --- \| | \| 1211 \| WBGene00194651:C33D9.13(14774) \| ``` log2FC=0.131 padj=0.973 ``` \| [seedLoc = 50,55] Seq=TGACATGAACTCACATACTACCTCC \| ```          C         ACAC            T mRNA  3'  CTCCATCAT    TCAA GTA CAG  miRNA 5'  GAGGTAGTA    GGTT TAT GTT           T                 G   A               311111111155552111621161125 ``` \| SeedType = 1 NonSeedType = 1 \| \| --- \| --- \| --- \| --- \| --- \| --- \| | \| 1212 \| WBGene00194908:F22F7.8(14815) \| ``` log2FC=1.306 padj=1.0 ``` \| [seedLoc = 128,133] Seq=TCAAATATAACTCAGGATTTTGTCTACCTCA \| ```                  TGTTTTAGGAC        AACT mRNA  3' ACTCCATC           TCAATATA     miRNA 5' TGAGGTAG           GGTTGTAT                      TA                 AGTT           1111111133555554455211121113333 ``` \| SeedType = 1 NonSeedType = 1 \| \| --- \| --- \| --- \| --- \| --- \| --- \| | \| 1213 \| WBGene00206487:best-19(14894) \| ``` log2FC=-0.147 padj=0.939 ``` \| [seedLoc = 47,52] Seq=TAACTGTACGTGTGATCTACCTCT \| ```          T       TAGTGT         T mRNA  3'  CTCCATC      GCATGTCAA  miRNA 5'  GAGGTAG      TGTATAGTT           T       TAGGT                      311111113333352111211115 ``` \| SeedType = 1 NonSeedType = -1 \| \| --- \| --- \| --- \| --- \| --- \| --- \| | \| 1214 \| WBGene00206522:F14D2.19(14902) \| ``` log2FC=1.636 padj=1.0 ``` \| [seedLoc = 80,86] Seq=AAACCGAAAAACCACCTACCATCA \| ```             A     CA    AAAGC   A mRNA  3' ACT CCATC  CCAA     CAA  miRNA 5' TGA GGTAG  GGTT     GTT                    TA    GTATA               111511111331111333331115 ``` \| SeedType = 3 NonSeedType = 1 \| \| --- \| --- \| --- \| --- \| --- \| --- \| | \| 1215 \| WBGene00219313:C17E4.20(14925) \| ``` log2FC=0.317 padj=0.911 ``` \| [seedLoc = 116,121] Seq=TCAAGTACAAAATCTACCTCC \| ```          C       TAA       AACT mRNA  3'  CTCCATC    AACATG     miRNA 5'  GAGGTAG    TTGTAT              T       TAGG      AGTT           3111111133361111123333 ``` \| SeedType = 1 NonSeedType = 1 \| \| --- \| --- \| --- \| --- \| --- \| --- \| | \| 1216 \| WBGene00219319:H25P06.5(14927) \| ``` log2FC=0.1 padj=1.0 ``` \| [seedLoc = 24,29] Seq=CGAATTGGAACGCTATCTCG \| ```          G             G  TAAGC mRNA  3'  CTCTATCG  CAA GT      miRNA 5'  GAGGTAGT  GTT TA               T        AG   G  TAGTT           3111211126611132133333 ``` \| SeedType = 2 NonSeedType = 1 \| \| --- \| --- \| --- \| --- \| --- \| --- \| | \| 1217 \| WBGene00219434:F54D5.23(14937) \| ``` log2FC=-0.314 padj=1.0 ``` \| [seedLoc = 26,31] Seq=TCCTGTGTGAACGACTACCTTC \| ```          C        G    GT     CT mRNA  3'  TTCCATCA  CAA  GTGTC   miRNA 5'  GAGGTAGT  GTT  TATAG            T        AG   G      TT           32111111136111352121133 ``` \| SeedType = 3 NonSeedType = 1 \| \| --- \| --- \| --- \| --- \| --- \| --- \| | \| 1218 \| WBGene00270293:F36H5.17(14989) \|  \| [seedLoc = 143,148] Seq=TCATTCACATTTCTTACCTCA \| ```                  C       CTTACT mRNA  3' ACTCCATT TTT ACA       miRNA 5' TGAGGTAG AGG TGT                        T   T   ATAGTT           1111111231226111333333 ``` \| SeedType = 1 NonSeedType = -1 \| \| --- \| --- \| --- \| --- \| --- \| --- \| | \|  \|  \|  \| [seedLoc = 149,153] Seq=TCATTCACATTTCTTACCTCAT \| ```          T        C       CTTACT mRNA  3'  ACTCCATT TTT ACA       miRNA 5'  TGAGGTAG AGG TGT                         T   T   ATAGTT           51111111231226111333333 ``` \| SeedType = 1 NonSeedType = -1 \| \| --- \| --- \| --- \| --- \| --- \| --- \| | \| 1219 \| WBGene00271605:H05L14.3(14995) \|  \| [seedLoc = 115,120] Seq=TCTTTTTCGTTTTATCTACCTCA \| ```                  T    T  TTTTTCT mRNA  3' ACTCCATC ATTT GC        miRNA 5' TGAGGTAG TAGG TG                              T  TATAGTT           11111111511223213333333 ``` \| SeedType = 1 NonSeedType = -1 \| \| --- \| --- \| --- \| --- \| --- \| --- \| | \|  \|  \|  \| [seedLoc = 249,254] Seq=ACACCCGCGCCTTCTGAATACCTACCTCT \| ```          T       CATAAGTCT       CCCACA mRNA  3'  CTCCATC         TCC GCG       miRNA 5'  GAGGTAG         AGG TGT                T       T           T   ATAGTT           311111113555545551116212333333 ``` \| SeedType = 1 NonSeedType = -1 \| \| --- \| --- \| --- \| --- \| --- \| --- \| | \|  \|  \|  \| [seedLoc = 255,260] Seq=CTGAATACCTACCTCTACCTTT \| ```          T            TC    AGTC mRNA  3'  TTCCATC TCCA  CATA     miRNA 5'  GAGGTAG AGGT  GTAT              T       T    T     AGTT           32111111611113511113333 ``` \| SeedType = 3 NonSeedType = 1 \| \| --- \| --- \| --- \| --- \| --- \| --- \| | \| 1220 \| WBGene00271796:C54F6.20(15005) \|  \| [seedLoc = 31,36] Seq=AAAAATGAAACGTTACCTCC \| ```          C         G   A   AAAA mRNA  3'  CTCCAT  T CAA GTA     miRNA 5'  GAGGTA  A GTT TAT              T      GT G   G   AGTT           3111111661311132113333 ``` \| SeedType = 1 NonSeedType = 1 \| \| --- \| --- \| --- \| --- \| --- \| --- \| | \| 1221 \| WBGene00271803:C09G5.14(15007) \|  \| [seedLoc = 80,85] Seq=CTCGTGTTCCACTGTACCTCC \| ```          C          ACCTTGTGCTC mRNA  3'  CTCCAT GTC            miRNA 5'  GAGGTA TAG                     T      G   GTTGTATAGTT           3111111621133333333333 ``` \| SeedType = 1 NonSeedType = -1 \| \| --- \| --- \| --- \| --- \| --- \| --- \| | \| 1222 \| WBGene00271818:K05F1.14(15010) \|  \| [seedLoc = 84,90] Seq=ATTAGTATAAAAAAGTTTCTCACCTCT \| ```          T     C  TTTGAAAA      ATTA mRNA  3'  CTCCA TC        AATATG     miRNA 5'  GAGGT AG        TTGTAT              T        TAGG          AGTT           311111511333355551121123333 ``` \| SeedType = 2 NonSeedType = -1 \| \| --- \| --- \| --- \| --- \| --- \| --- \| | \| 1223 \| WBGene00302971:F59E12.17(15022) \|  \| [seedLoc = 261,267] Seq=AATCAAGGAGCTTCTTACACTCA \| ```              A    C     G AACTAA mRNA  3' ACTC CATT TTCGA G       miRNA 5' TGAG GTAG AGGTT T                         T     G ATAGTT           11115111231212132333333 ``` \| SeedType = 2 NonSeedType = 2 \| \| --- \| --- \| --- \| --- \| --- \| --- \| | \| 1224 \| WBGene00303026:T20D3.14(15044) \|  \| [seedLoc = 56,61] Seq=GAAAAACCAACTTATCTACCTTT \| ```          T       T       CAAAAAG mRNA  3'  TTCCATC ATTCAAC        miRNA 5'  GAGGTAG TAGGTTG                 T               TATAGTT           32111111511211113333333 ``` \| SeedType = 3 NonSeedType = 1 \| \| --- \| --- \| --- \| --- \| --- \| --- \| |

 
